# Evolutionary dynamics of microbial communities in bioelectrochemical systems

**DOI:** 10.1101/725580

**Authors:** Lukasz Szydlowski, Anatoly Sorokin, Olga Vasieva, Susan Boerner, Veyacheslav Fedorovich, Igor Goryanin

## Abstract

Bio-electrochemical systems can generate electricity by virtue of mature microbial consortia that gradually and spontaneously optimize performance. To evaluate selective enrichment of these electrogenic microbial communities, five, 3-electrode reactors were inoculated with microbes derived from rice wash wastewater and incubated under a range of applied potentials. Reactors were sampled over a 12-week period and DNA extracted from anodal, cathodal, and planktonic bacterial communities was interrogated using a custom-made bioinformatics pipeline that combined 16S and metagenomic samples to monitor temporal changes in community composition. Some genera that constituted a minor proportion of the initial inoculum dominated within weeks following inoculation and correlated with applied potential. For instance, the abundance of *Geobacter* increased from 423-fold to 766-fold between −350 mV and −50 mV, respectively. Full metagenomic profiles of bacterial communities were obtained from reactors operating for 12 weeks. Functional analyses of metagenomes revealed metabolic changes between different species of the dominant genus, *Geobacter*, suggesting that optimal nutrient utilization at the lowest electrode potential is achieved via genome rearrangements and a strong inter-strain selection, as well as adjustment of the characteristic syntrophic relationships. These results reveal a certain degree of metabolic plasticity of electrochemically active bacteria and their communities in adaptation to adverse anodic and cathodic environments.

## 1 INTRODUCTION

Bio-electrochemical Systems (BESs) refer to microbial communities that either generate electricity, as in Microbial Fuel Cells (MFCs), or utilize electricity, as in Microbial Electrolysis Cells (MECs) (Santoro *et al*., 2017; Rittmann and Asce, 2017). BES is a well-known technology allowing simultaneous wastewater treatment and electricity production. BES performance depends on activities of electrode-associated bacteria (EAB) that form biofilms on anodal surfaces (e.g. Allen and Bennetto, 1993). Various factors account for EAB enrichment: organic substrates, pH, temperature, electrode composition, and electrical potential (Logan *et al*., 2006; Aelterman *et al*., 2008; Torres *et al*., 2009; Dennis *et al*., 2016; Rittmann and Asce, 2017).

EAB reach maximum power density when reactors operate at near-neutral pH, at ambient temperatures (25-40°C), and are fed with acetate. Although some studies indicate optimum electrode potentials at ca. 0.3 mV in comparison with a standard hydrogen electrode (SHE) (Aelterman *et al*., 2008), based upon thermodynamics of acetate consumption, others have found that the most efficient EAB prefer lower anode potentials (Torres *et al*., 2009). Microbial communities used to inoculate BESs were derived mainly from sludge from wastewater treatment plants (Torres *et al*., 2009; Ishii *et al*., 2013; Paitier *et al*., 2017), aquatic sediments (Holmes *et al*., 2004), biogas digestate (Daghio *et al*., 2015), and various environmental samples (e.g. Yates *et al*., 2012; Ieropoulos *et al*. 2010). EAB are abundant in many environments, such that virtually all environmental inocula can eventually give rise to stable EAB consortia within 60 days (Yates *et al*., 2012). However, further changes and details of community structure are not well understood.

Previous studies analyzed changes within microbial populations for up to several weeks (reviewed in Daghio *et al*., 2015; Philips *et al*., 2015; Khater *et al*., 2017), and employed mainly 16S sequencing (Ishii *et al*., 2013; Ishii *et al*., 2014; Dennis *et al*., 2016), Ribosomal Intergenic Spacer Analysis (RISA) (Paitier *et al*., 2017), or Denaturing Gradient Gel Electrophoresis (DGGE) (Beecroft *et al*., 2012). In another study, community shifts were tracked during 90-days of operation (Beecroft *et al*., 2012). However, those reactors were fed with sucrose, which cannot be metabolized as efficiently (e.g. Schroder, 2007) as acetate (Bond *et al*., 2002; Bond and Lovley, 2003; Logan *et al*., 2006; Fedorovich *et al*., 2009; Daghio *et al*., 2015). Sucrose-fed systems developed fermentative communities that did not participate in electron transfer. However, these population studies may significantly underestimate microbial diversity, with novel taxonomic groups not detected due to low compatibility with universal primers (e.g. Poretsky *et al*., 2014; Roselli *et al*., 2016). Moreover, the type of inoculum also influences community development. In our previous work (Khylias *et al*., 2015), electrogenic communities derived from different sources exhibit different properties in terms of COD consumption, as well as coulombic efficiencies. Therefore, communities present in particular waste streams should already contain some electrogenic bacterial taxa. The minimal number of EAB is unknown and it remains unclear whether conclusions drawn by Yates et al. (2012) are valid for all inocula. Thus, complex studies examining long-term community changes across a range of EAB-selective electrode potentials have not been attempted.

In this study, we investigated enrichment of EAB from rice wash inoculum in single-chamber, three-electrode BES. Normalization of conditions among reactors utilizing acetate feeding provided selection pressure for metabolic pathways relevant to electrogenic respiration. Application of potentials ranging from −50 mV to −350 mV vs Ag/AgCl (147 mV to −153 mV vs SHE) on anodes and compositional and functional changes in the resulting bacterial communities were tracked using detailed metagenomics.

## 2 RESULTS

### 2.1 Conditions within reactors

The single-chamber, 3-electrode BES reactors (M1-M4) were connected to a potentiostat to apply fixed potentials (147 mV to −153 mV vs SHE) to the working electrodes (anodes). We also measured potentials observed (792 mV to 1182 mV vs SHE) on counter electrodes (cathodes). In addition, we prepared one reactor (M5) with only one set of electrodes to operate under open circuit potential (OCP) (Table 1). Our BESs were designed to develop electroactive biofilms, with high volume-to-electrode surface ratios in order to minimize nutrient limitations. Thus, after setting a constant potential, the potentiostat maintained current flow automatically and we did not measure current. One channel of the potentiostat, attached to reactor M3, exhibited an overload error after 8 weeks of operation, resulting in turbidity. This caused a shift in community structure (Fig. S2); hence, we excluded this reactor from our analysis.

**Table 1.** Potentials (measured with Ag/AgCl reference electrode in saturated KCl; +0.197 mV vs SHE) vs. SHE on anode **(A)** and cathode **(C)** of each reactor.^a)^ Electrode potential measured in the control (OCP) reactor after every 2 weeks. * Around week 8, we experienced a technical fault in M3, resulting in overcharge of the reactor, which caused a shift in the microbial community; results for this reactor are in supplementary material, Fig. S2.

| Reactor | Anode potential mV (vs SHE) | Cathode potential mV (vs SHE) |
| --- | --- | --- |
| M1 | -50 (147) | 595 (792) |
| M2 | -150 (47) | 775 (972) |
| M3* | -250 (-53) | 885 (1082) |
| M4 | -250 (-153) | 985 (1182) |
| M5 | 0 to -530 (0 to -333)<sup>a</sup> | NA |

### 2.2 Community analysis

Samples were collected every two weeks from anodes (M1-5A), cathodes (M1-4C) and planktonic (free swimming) fractions (M1-5P). Microscopic imaging (Fig. 1) revealed that anodes from M1-M4 reactors developed communities that formed thick biofilms (Fig. 1a-c), and they were more abundantly populated than cathodes (Fig. 1e-g). No comparable community formed on electrode strips in M5 (Fig. 1d), due to its OCP mode. Negative charge accumulation on the electrode eventually inhibited microbial growth (Table 1).

**Fig. 1.**
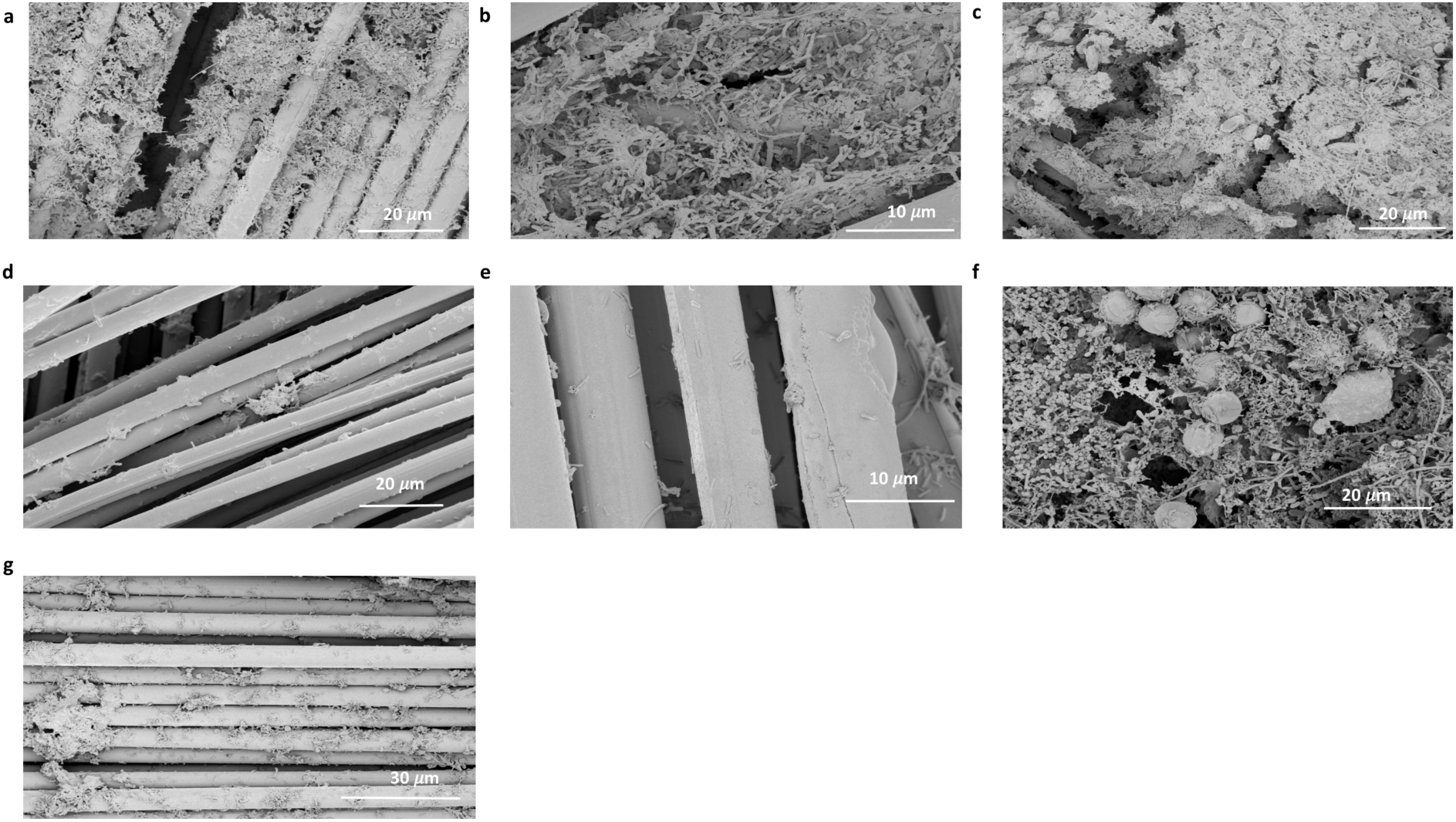
SEM images of electrodes from a) M1A, b) M2A, c) M4A d) M5A e) M1C, f) M2C and g) M4C after 12 weeks of operation. Scale bars are included.

### 2.3 Taxonomic analysis of anodes

We analyzed organismal abundances on each anode (Fig.2) during the 12-week operation and compared them with initial community compositions (Table S2). Although rice inoculum is nutrient-rich, the abundance of electrogenic taxa were very low, with *Geobacter* and *Shewanella* spp., two of the most efficient EABs, comprising less than 0.09% and 0.02% of the total communities, respectively. Enrichment data indicated that the abundance of *Geobacter* spp. rapidly increased over the first 6-8 weeks under poised anode potential (M1-4). Between 8 and 12 weeks, *Geobacter* growth rates plateaued and the change in abundance of *Geobacter* after 12 weeks was 766-fold, 598-fold and 423-fold in M1A, M2A and M4A, respectively, whereas in M5A it only increased 1.2-fold. The relative abundance of *Geobacter* was significantly higher (ANOVA, p < 0.05) in M1A at −50 mV and decreased with decreasing potential. After 12 weeks of operation, the abundance *of Shewanella* remained unchanged in all reactors. Control reactor M5 showed an oscillating pattern of the most abundant genera *Methanosaeta, Methanobacterium*, and *Acinetobacter*, with an opposite oscillation pattern of *Prevotella* (Fig.2d), although the changes are not as significant as in the other reactors. With regard to generic abundance differences between the reactors, for M1A, *Geobacter* was the only genus with an abundance over 1% after 12 weeks’ operation. M2A had 2 such genera (*Geobacter* and *Denitrovibrio*) and in reactor M4 and M5 anodes, the number of significantly abundant (i.e. > 1%) genera reached 10. In the initial inoculum, 5 genera showed abundances >1%.

**Fig. 2.**
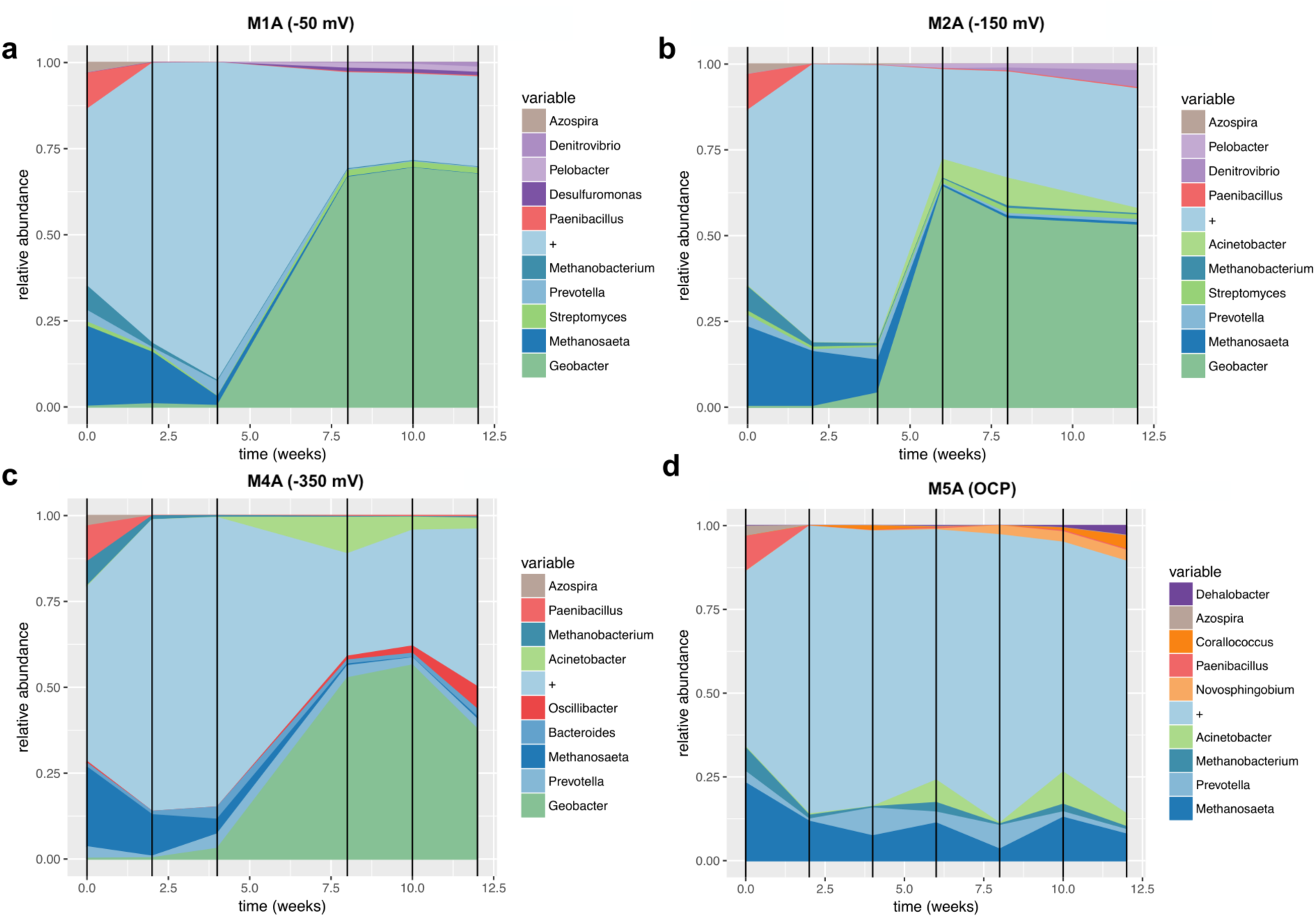
Relative abundances of anodic dominant genera collected from a) M1A, b) M2A, c) M4A and d) M5A during the experiment. Colors represent specific taxonomic groups, “+” refers to all other organisms.

### 2.4 Taxonomic analysis of cathodes

On the M1 and M2 cathodes, the most abundant organisms were methanogenic archaea (Fig.3a-b); however, a proportion of *Methanosaeta*, the most abundant methanogenic genus from the initial inoculum, decreased during the course of the experiment with the subsequent growth of *Methanobacter* spp. The growth of the latter was in turn inversely correlated with that of *Methanoregula*. Unexpectedly, *Geobacter* was the most abundant genus on M4C, reaching a peak abundance of 16%, 12 weeks after inoculation, a level four times higher than that of the next most abundant genus, *Methanobacterium* (Fig.3c). *Geobacter* remained scarce on the M1 and M2 cathodes (< 0.1%), not exceeding its abundance in the inoculum. On the M4 cathode, a rapid increase in *Geobacter* abundance after 10 weeks was accompanied by a concomitant decrease of *Acinetobacter* from 25% to 2.5%(Fig.3c).

**Fig. 3.**
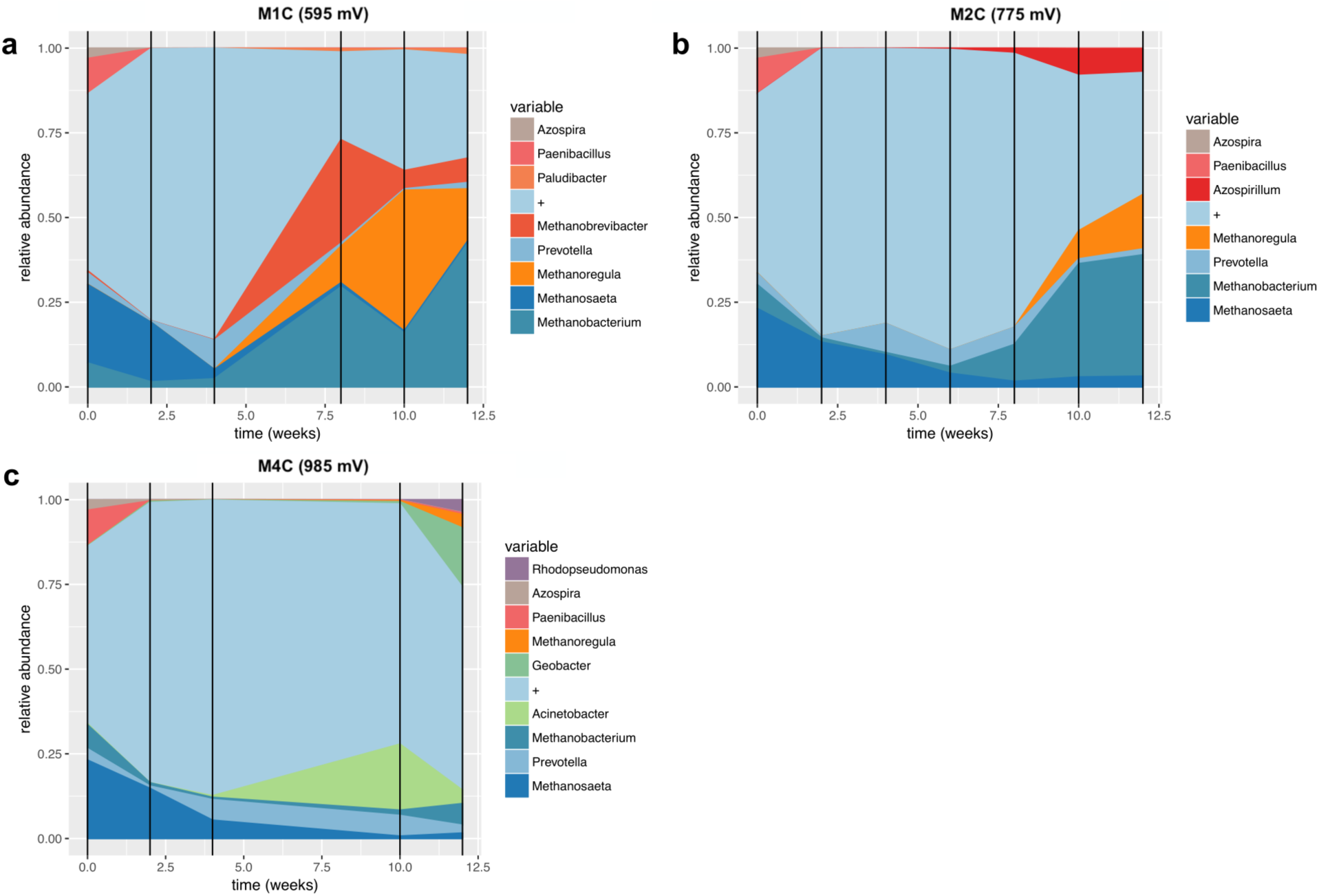
Relative abundances of cathodic dominant genera collected a) M1C, b) M2C and c) M4C during the experiment. Colors represent specific taxonomic groups, “+” refers to all other organisms.

### 2.5 Taxonomic analysis of planktonic communities

In the case of planktonic samples (Fig.4), in M1P, a pattern of sudden growth around week 8, similar to that observed on M1A (although with much lower abundances) was observed with the genera, *Pelomonas, Paludibacter*, and *Bacteroides*. In all planktonic samples, *Methanosaeta*, the most abundant genus in the inoculum (Table S1), decreased within the first weeks of operation, as did *Porphyromonas and Azospirillum*. The proportion *of Pelomonas*, a genus comprising 0.01% of the initial community, rose to about 20% of the total planktonic community in M2 after 12 weeks. In M5P, abundances did not reflect the initial community profile, as *Methanosaeta* decreased within 4 weeks from 24 % to ~1%, whereas *Azospirillum* abundance reached ~25%. *Geobacter* abundance was 6.54%, 1.92%, 4.27% and < 0.05% in M1, M2, M4 and M5 planktonic communities, respectively.

**Fig. 4.**
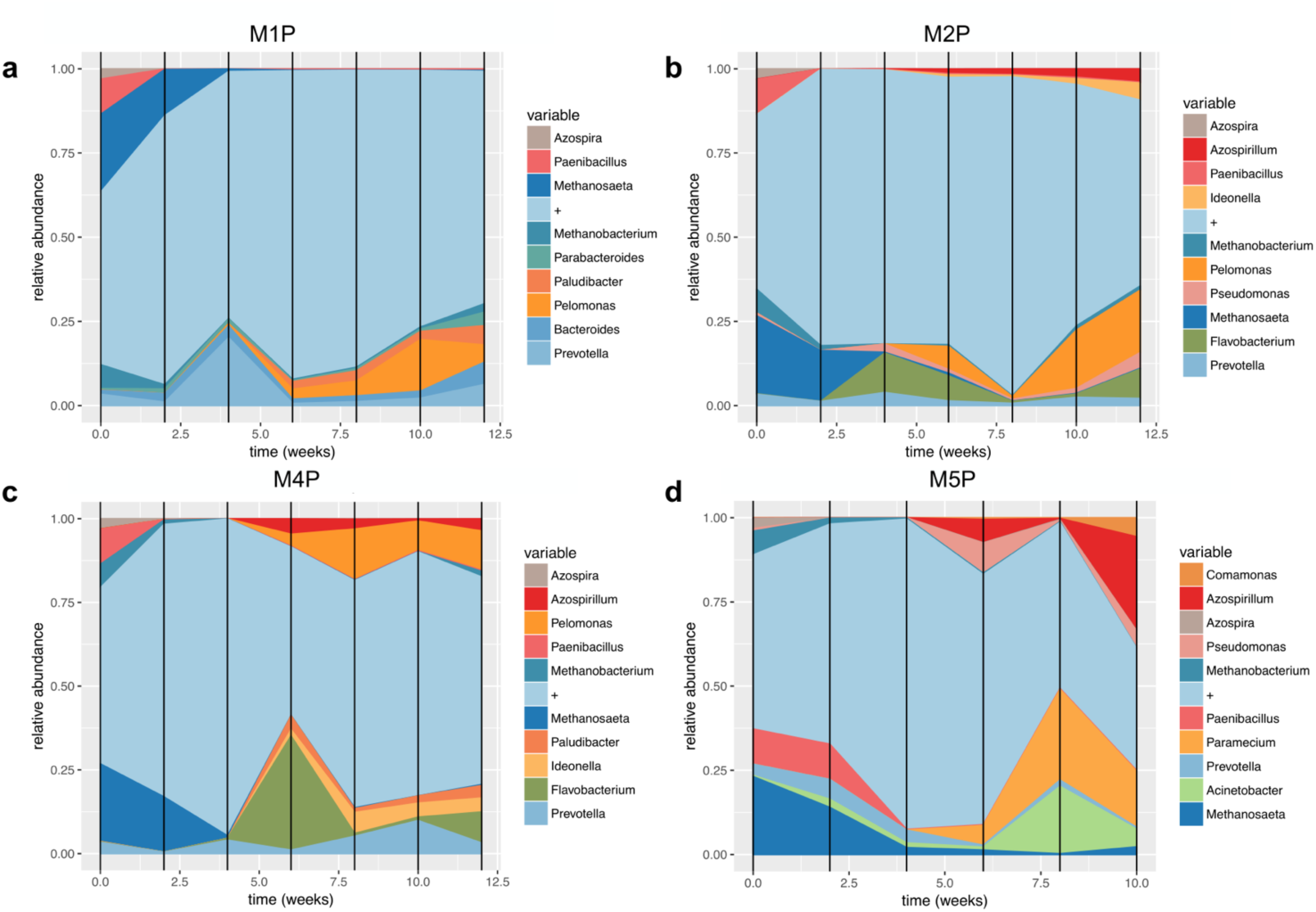
Relative abundances of planktonic dominant genera collected from a) M1P b) M2P c) M4P and d) M5P during the experiment. Colors represent taxonomic groups, “+” refers to all other organisms.

### 2.6 Diversity of metagenomes

We used multidimensional scaling to look at how the communities clustered based on sample type (initial sludge, anode, cathode, plankton and OCP) at both the genus level (Fig. 5a) and functional levels (Fig. 5b) after 12 weeks. Significant differences were found for both levels, with 72% of the distance variability due to sample-type and applied potential at the genus level (Fig. 5a, p-value 0.01), and 65% (Fig. 5b, p-value 0.04) at the functional level. We also compared the percentage of unclassified reads (at the generic level) from each metagenome to that from the initial inoculum (Fig. S3). Results indicate almost a 2-fold increase of unclassified taxa after 12 weeks in all sampled metagenomes, with the highest being reported in M4P (34.9%), followed by M1P (31.9%) and M5P (30.0%), with 17% of unclassified genera in the initial inoculum.

**Figure 5.**
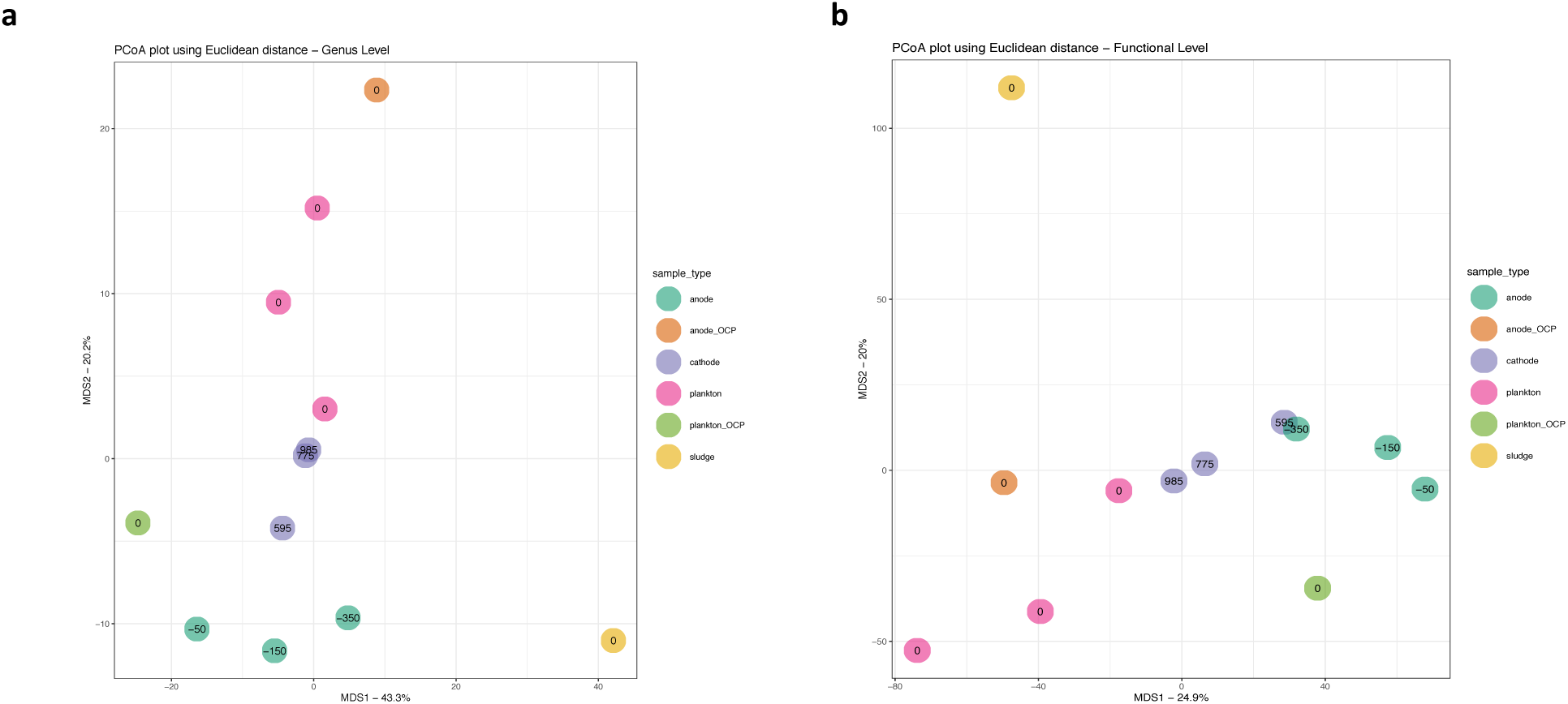
PCoA plots based on **a)** taxonomy **b)** function.

### 2.7 Functional overview of metagenomes

Metagenomic analysis also revealed changes in abundances of functions mapped to genomes that were identified via PALADIN analysis. *Methanosaeta* functions were predominant in the initial communities and the M5 reactor, where they ranked in the top 200 by read count (Tables S3 and S7, respectively). *Geobacter* functions dominated the ranked lists for the M1, M2, and M4 reactors. To gain a deeper understanding of the communities, we compared abundances in greater detail and ranked the top 200 *Geobacter* functions from each reactor (Supplementary tables S3-S7). The rank of each gene was established in relation to the normalized abundance of its mapped reads (See Methods for a gene/function abundance calculation) for each species. The function with the highest number of mapped reads was assigned a rank of 1. Functions with lower numbers of mapped reads had lower ranks with larger assigned values. The main abundance trend defined by the taxonomic analysis, correlates with the occurrence of *Geobacter* spp. in reactors, with counts for almost all mapped genes decreasing in the order M1>M4>M2>M5. However, we also noticed changes in ranks of several *G. metallireducens* and *G. sulfurreducens* genes that may reflect changes in the proportion of these functionally significant genes in *Geobacteraceae* populations in different reactors. A comparison of M4A to M1A revealed that 14 genes increase and 17 genes decrease in rank in M4A with respect to M1A (Fig. 6). An increase in rank order suggests potentially favorable genomic changes and may help to identify species-specific significant functions for specific reactor conditions. The comparative nature of the analysis also helps to avoid biases caused by gene length due to translation of the number of reads into a gene presentation ratio. Interestingly, the results of such a comparison of *Geobacter* fractions at the M1 and M4 metagenomes (Fig. 6) suggests a positive selection of bacteria for functions/genes involved in electrogenic metabolism. Metagenomic changes in *G. sulfurreducens* were related to genes encoding ATP synthase, NADH-quinone oxidoreductase, and the acetate utilization pathway, with 2-fold, 1.8-fold, and 1.5-fold rank increases in M4A compared to M1A. Conversely, *G. sulfurreducens* genes encoding ATPase (*prkA*), citramalate synthase (*cimA*), sodium symporter (*aplC*), aldehyde dehydrogenase (*aldh*), and Fe-S binding protein increased 4-fold, 2.02-fold, 2-fold, 1.52-fold, and 1.51-fold in rank in M1A, respectively. Changes in the *G. metallireducens* metagenome included a wide range of functions involved in conductive pilin assembly (*pilB*), flagella biosynthesis regulation (*fgrM*), pyruvate metabolism (*leuA*), electron transfer (*nuoB/C/G/L* and *por*), utilization of ammonia (*carb-1*) efflux pump (cusA), and aspartokinase (*asd-1*) between 2-fold and 1.5-fold in M4A, whereas periplasmic Ni-Fe dehydrogenase (*hybL*) and NADH-quinone oxidoreductase (*nuoD*) show 1.95 and 1.76-fold increase in M1A.

**Figure 6.**
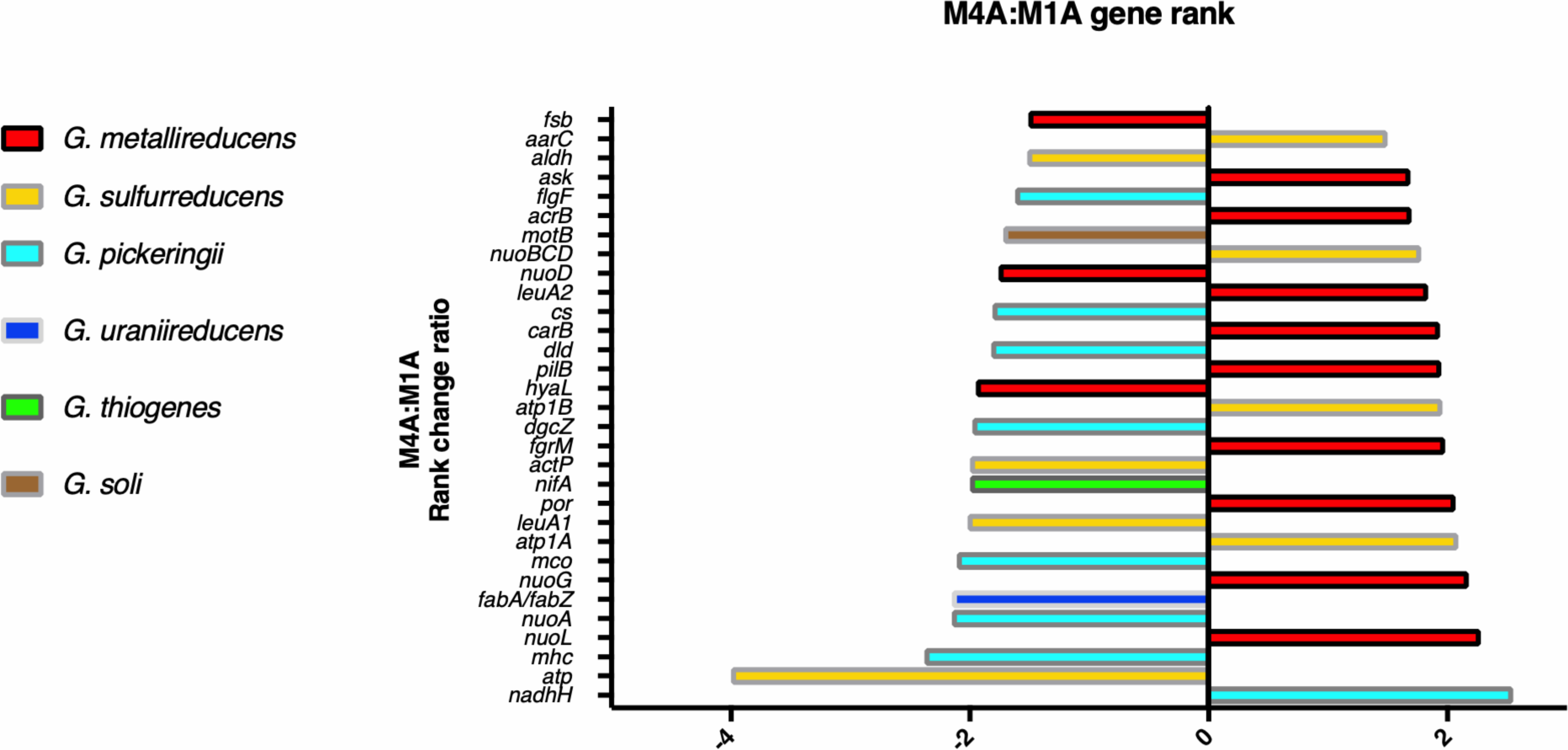
Rank change ratio between M4A and M1A. Colours represent hits with the highest match to one of *Geobacter* spp. Fold change ratio 1.5 was chosen as a threshold. Represented genes are as follows:***fsb*,** Fe-S binding protein; ***aarC*,** Succinyl:acetate coenzyme A transferase; ***aldh*,** aldehyde dehydrogenase; ***ask*,** aspartokinase; ***flgF*,** flagellar basal-body rod protein; ***acrB*,** Efflux pump, RND family, inner membrane protein; ***motB*,** flagellar basal body stator protein; ***nuoBCD*,** NADH-quinone oxidoreductase subunit B/C/D (EC 1.6.5.11); ***leuA2*,** 2-isopropylmalate synthase (EC 2.3.3.13); ***cs*,** type I citrate synthase (EC 2.3.3.1); ***carB***, Carbamoyl-phosphate synthase; ***dld*,** Dihydrolipoyl dehydrogenase (EC 1.8.1.4); ***pilB*,** Type IV pilus biogenesis ATPase; ***hyaL*,** Periplasmically oriented, membrane-bound [NiFe]-hydrogenase; ***atp1B*,** ATP synthase subunit beta (EC 3.6.3.14); ***dgcZ*,** Diguanylate cyclase; ***fgrM*,** Flagellar biogenesis master sigma-54-dependent transcriptional regulator; ***actP*,** Sodium/solute symporter family protein; ***nifA*,** Nif-specific regulatory protein; ***por*,** Pyruvate-flavodoxin oxidoreductase; ***leuA1*,** (R)-citramalate synthase (EC 2.3.1.182); ***atp1A*,** ATP synthase subunit alpha; ***mco*,** multicopper oxidase; ***nuoG*,** NADH dehydrogenase I, G subunit; ***fabA/fabZ*,** Beta-hydroxyacyl dehydratase; ***nuoA*,** NADH-ubiquinone oxidoreductase subunit 3; ***nuoL*,** NADH dehydrogenase, subunit L; ***mhc*,** multiheme cytochrome; ***atp*,** ATPase; ***nadhH*,** NADH-quinone oxidoreductase subunit H (EC 1.6.5.11).

Genes from the *Geobacteraceae* (*G. pickeringii, G. uraniireducens, G. thiogenes*, and *G. soli*) mostly increased in rank in M1A, with outer membrane multiheme cytochrome c (*omc*), NADH-ubiquinone oxidoreductase (*nuoA*), β-hydroxyacyl dehydratase (*fabA/Z*), multicopper oxidase (*ompB*), Nif-regulatory protein (*nifA*) exhibiting more than a 2-fold increase in rank, as well as dihydrolipoyl dehydrogenase (*lpdA*), type I citrate synthase (*gltA*), and flagellar components (*motB,flgEF*), which exhibited 1.82, 1.81 and 1.7-fold shifts in rank, respectively. In M4A, a 2.55-fold rank increase was observed for the NADH quinone oxidoreductase gene (*nuoH*) from *G. pickeringii*.

In all planktonic samples, as well as in the initial community, top ranked genes are those involved in genome rearrangement (transposases, reverse transcriptases and endonucleases, see Tables S3-S7), which indicates selective pressure for adaptation to a more competitive environment.

## 3 DISCUSSION

### 3.1 Abundance of *Geobacter* spp. at anodes is directly proportional to the applied voltage

*Geobacter* is a well-characterized genus of EAB that populates BES anodes abundantly (Bond and Lovley, 2003). It can comprise ≤99% of bacterial communities isolated from BES electrodes operating at the lowest potential (Torres *et al*., 2009). However, our work indicates that the abundance of *Geobacter* increases at anodes with increasing applied potential, meaning that as the electrode potential increases, *Geobacter* competes more effectively with other genera in the community. In contrast, *Shewanella* did not increase in abundance under these conditions, likely due to the fact that *Shewanella* primarily utilizes lactate as a carbon source (e.g. Kim *et al*., 1999; Pinchuk *et al*., 2009).

Our abundance results contrast with those of Ishii *et al*. (2014), in which the highest abundance of *Geobacter spp*. in acetate-fed, set-potential reactors was observed when anodes were held at −50 mV vs SHE (−247 mV vs Ag/AgCl), whereas in our study the highest abundance was observed on anodes poised at 147 mV vs SHE (−50 mV vs Ag/AgCl). However, abundances from our study resemble those reported by Dennis *et al*. (2016), where highest *Geobacter* abundance was observed on anodes poised at 300 mV vs SHE (103 mV vs Ag/AgCl). Also, the low initial population of *Geobacter* in the inoculum (see Table S2) may explain the slower growth of *Geobacter* spp. at M4 compared to M1 and M2. Although periodic metagenomic sequencing reveals changes in the most abundant genera, it also indicates a large number of potentially undetected bacterial taxa. Changes in this community, as well as interactions among the most abundant EAB, will remain enigmatic, however, until genome assembly and isolation methods are improved to identify and characterize new strains.

### 3.2 Presence of *Geobacter* spp. at cathodes

Apart from the *Geobacter* presence at anodes, *Geobacter* also dominated the M4 cathode community after 12 weeks, although it was scarcely present at other cathodes (0.06% and 0.07% in M1 and M2 cathodes, respectively) as well as at the M5 electrode (0.1%), being close to the inoculum abundance (0.08%). Such an increase in *Geobacter* spp. abundance in compartments with opposite conditions reflects its ability to both donate and accept electrons in association with electrodes (Holmes *et al*., 2004; Gregory *et al*., 2004). However, *Geobacter* was not found at cathodes in other studies (e.g. Daghio *et al*., 2015), which may reflect competition with different bacterial taxa, as well as differences in operating conditions, initial community structure, etc. Moreover, rank shifts of flagellar biosynthesis genes demonstrate ongoing colonization of new environmental niches by *Geobacter* spp. Recently, Rittmann and Asce (2017) concluded that the best-performing EAB has the lowest anode potential, but noted that such conditions are in fact stressful to the bacteria. Perhaps the M4 cathode offered less deleterious conditions for *Geobacteriae* growth. The continuous decrease of methanogenic archaea at the M4 cathode may hint at competition for electrons with *Geobacter* spp., a known electrotroph (Strycharz *et al*., 2011). It is tempting to suggest that incidental oxygen formation, due to the potential difference between M4 electrodes, exceeded the potential difference at which electrolysis of water can occur (1.33 V vs 1.23 V). However, we did not observe bubble formation; therefore, the latter hypothesis may be discarded. Moreover, there are no abundance shifts due to oxygen stress between the M1 and M4 cathodic metagenomes (Table S2), which would certainly follow electrolysis by BES. There is also no evidence of rank shifts in genes involved in hydrogenotrophic methanogenesis, e.g., fumarate reductase, hydrogenase within. Microscopic observations (Fig. 1e-g) suggest some other relationship between *Geobacter* and methanogens in the cathodic community, for example via direct interspecies electron transfer (Lovley, 2017).

### 3.3 Functional analysis and evidence of differential selection pressure on *Geobacter* at low electrode potentials

We were intrigued by the dependence of observed genomic shifts in *Geobacter* metabolic functions on reactor conditions. The changes themselves, relevant to the main pathways of electrogenic organisms in MFCs, suggest that bacterial genomes evolve rapidly due to metabolic competition. Acetate metabolism is central to the metabolism of *G. sulfurreducens* under electrogenic conditions (Bond and Lovley, 2003; Ieropoulos *et al*., 2010). This metabolic feature allows these bacteria to dominate anodic communities if acetate is provided or generated by other members of the syntrophic bacterial community (e.g. *G. metallireducens)*. Interestingly, *G. sulfurreducens* functions required for acetate utilization (*ato*) were the only ones from this species that strongly changed rank at lower anodic potential (M4), with aldehyde dehydrogenase rank decreasing (Fig. 6). Acetate utilization by *G. sulfurreducens* may be supported by a syntrophic association with *Pelobacter* spp. (Sreshtha *et al*., 2013), and enrichment of this genus was observed on anodes M1 and M2 (Fig. 2a-b). Though acetate was provided in the medium, local interactions between bacteria and bacterial clusters may be significant. Sequences corresponding to ATP synthase subunits also increased in abundance in M4A, with a 4-fold decrease in ATPase (Fig. 6), suggesting higher pressure for energy generation.

At the same time, adaptation and evolution of highly electrogenic *G. metallireducens* at M4 seems to proceed due to a requirement for conductive pili, respiratory NADH dehydrogenases, and pyruvate metabolism, which increase in rank at M4A (Fig. 6). NuoL, for which metagenomic rank changed most at the M4 anode compared to M1A, is responsible for the reverse electron transfer and H^+^/e^-^ stoichiometry (Steimle *et al*., 2011), which may be important for balancing electron flow between NAD+ and ferredoxin pools. PilB is an ATPase required for polymerization of conductive e-pili (McCallum *et al*., 2017), and *pilB* mutants are reported to generate lower current and form thinner biofilms (Steidl *et al*., 2016). Additionally, it was observed by Ishii *et al*. (2018) that *pilA* expression increased at lower surface potential. Change in *pilB expression* has not been reported in the aforementioned study, although it could be a result of a normalization procedure: with an increase in the abundance of DNA reads, changes in abundance of RNA reads/expression could not be observed.

Flagellar response regulator (*fgrM*), which increased in rank in M4A, regulates flagellar growth, a known feature of *G. metallireducens* when grown with an insoluble Fe(III) source (Ueki *et al*., 2012). This feature corresponds to increased motility of cells when Fe(III) sources are sparse. Cells can store electrons in their numerous cytochromes, acting as capacitors, so that they can discharge them upon the next available Fe(III) cluster. Similarly, lower surface potential of the anode in M4A could lead to formation of dispersed local spots for electron release. Thus, *G. metallireducens* cells with regulated expression of flagella proteins could possess higher and more ordered motility, which, together with the higher capacity to polymerize e-pili by PilB, should give them an advantage over competitors.

This study shows a decrease in rank of genes belonging to other taxa (*G. pickeringii, G. uraniireducens, G. thiogenes* and *G. soli*) at lower surface potential (Fig. 6b). Citrate synthase (*G. pickeringii*), a proposed indicator of Geobacteraceae metabolic activity (Holmes *et al*., 2005) decreased almost 2-fold at M4A. Also, genes encoding components of EET are lower in rank at M4A (Fig. 6b). This correlates with the limited capacity of these Geobacteraceae to adapt to low surface potential, as their electrochemical activity was reported for rather more positive redox potentials (Ishii *et al*., 2018).

We conclude that the comparative rank measurement gives a better estimate of a functional genomic shift for a particular metagenome in relation to a reactor condition, which is not surprising when one takes into account that ranks are known to be a robust characterisation of the population for analysis of covariance (Conover and Iman, 1982; Quade, 1967). The rank transformation has been widely used since Charles Spearman defined the correlation coefficient in 1904 and more recently it has been adapted in metagenomics studies of microbial communities. (e.g. Saeedghalati et al., 2017). Certainly, gene abundance cannot be taken as an unequivocal reflection of activity levels of relevant functions in bacterial metabolism under different conditions. However, we suggest that genomic rearrangements represent responses to specific functional requirements. Genes may be retained or even propagate in a population if they enhance organismal fitness. Increasing abundances of bacterial strains bearing advantageous genes may also explain the observed phenomenon. As shown in the case of *pilB* gene, analysis of gene ranks derived from metagenomics analysis can complement expression studies.

### 3.4 The importance of unknown organisms in a functional overview of these metagenomes

The presence of unclassified taxa indicates an increase in abundance of unknown organisms upon inoculation into BES reactors. Our results (Fig. S3) do not align with other works (Ishii *et al*., 2014; Dennis *et al*., 2016), in which unclassified taxa in the inoculum contained comparable numbers of unclassified reads, but only several percent of unclassified organisms were identified in sampled electrodes. The discrepancy between the two studies may be due to the difference in sampling and sequencing methods. The former studies employed only 16S samples, whereas we compared Illumina MiSeq whole metagenomic reads. Such discrepancies were also reported in a later study by Ishii et al. (2018), where lower diversity was reported in the same samples when only 16S analysis was employed. The abundance of novel unidentified organisms suggests the existence of novel electrogenic microorganisms. Such organisms may not be as efficient in EET as *Geobacter spp*.; hence the term “weak electricigens” (Doyle and Marsili, 2018), but they may nonetheless provide useful insight into the divergence of EET mechanisms. However, since their increases do not follow electrode potential, the presence of so many unclassified organisms is perhaps more dependent on the inoculum than the reactor conditions.

## 4 EXPERIMENTAL PROCEDURES

### 4.1 Reactor setup and operating conditions

Four reactors (M1-4) were designed as follows: 1.2L chamber with an anode consisting of 6 carbon-fiber strips (Zoltec) 3×10 cm connected with titanium wire (Kojundo chemical laboratory), a cathode consisting of 6 carbon-fiber strips (Zoltec) 3×10 cm connected with titanium wire, and a reference electrode (Radiometer Analytical, Hach). Additionally, one control reactor (M5) consisted only of one set of 6 carbon-fiber strips (Zoltec) 3×10 cm connected with titanium wire and reference electrode (Radiometer Analytical, Hach), and was operated in open circuit mode (see Fig.S1 for schematic view). A four-channel potentiostat (UniChem) was connected to each reactor with stainless steel clips and potential differences of −50 mV, −150 mV, −250 mV, −350 mV vs Ag/AgCl (147 mV, 47 mV, −53 mV and - 153 mV vs SHE) were applied to anodes M 1-4A, respectively. Each reactor was inoculated with rice wash water (1.2L) and incubated for 2 weeks at room temperature (23°C), after which the liquid was replaced with an equal volume of the following medium: 0.05 M phosphate buffer (pH 6), 200 mg/L CaCl_2_ •2H_2_ O, 250 mg/L MgCl •6H_2_O, 500 mg/L NH_4_ Cl, sodium acetate 2g COD/L (Fedorovich *et al*., 2009). COD concentration was measured using a Hach COD kit (Hach, USA). The medium was replaced 6 times at 2-week intervals, yielding a total operating time of 12 weeks. Additionally, 50 mL of liquid fraction and one strip of each electrode (1x A and 1x C from M1-4 and 1x A from M5) were collected with every change of medium. These were used for DNA extraction and SEM analysis.

### 4.2 Microscopic imaging

Samples for microscopic imaging were taken simultaneously with the DNA samples and processed with osmium, as follows (Fischer *et al*., 2012): upon removal from the anode compartment, the samples were immediately cut by knife, and fixed by 1%Osmium diluted with 0.2M Cacodylate (Wako) buffer 30min. The samples were then washed three times with RQ water and dehydrated stepwise with a graded series of ethanol solutions (70, 80, 90, 95 and three times 100%). The electrode samples were finally critical-point dried with tert-butyl ethanol and sputter coated with a thin layer of gold. The samples were analyzed by a scanning electron microscopy (SEM) (JSM-7900F JEOL).

### 4.3 DNA extraction and library preparation

DNA was extracted using TRIzol (Life Technologies) and additional samples were subjected to Maxwell extraction (GMO purefood kit, Maxwell) using an automated RSC system (Promega). Samples with sufficient amounts of DNA were subjected to Illumina sequencing (48). Remaining samples (32) were subjected to 16S sequencing. DNA libraries were constructed using Nextera XT kit (Illumina) and sequencing was performed on MiSeq platform (Illumina, San-Diego, CA, USA). Samples were uploaded to MG-RAST (mgp81854 for 16S, mgp82844 for metagenomes). We were unable to collect data from 3 cathodal samples (see Table S1).

### 4.4 Bioinformatic analysis

Whole-genome sequences and 16S sequences were analyzed using a custom-developed pipeline, as described elsewhere (Orakov *et al*., 2017), which carried out taxonomic analysis using Kaiju (Menzel and Krogh, 2016), as well as functional analysis using PALADIN (only applicable to metagenomes) (Westbrook *et al*., 2017). Results of PALADIN analysis for anodes M1 and M4 can be found in the Supplementary material (Table S3). Compositional analysis of communities was performed in R version 1.4.0 (van den Boogaart *et al*., 2018) with package compositions (van den Boogaart and Tolosana-Delgado, 2008). Relative abundance was represented as composition with absolute geometry (rcomp). To combine 16S and metagenomic sequences, datasets from Kaiju and MG-RAST were manually curated (a more detailed description can be found at https://github.com/lptolik/ASAR). One-way ANOVA was conducted to determine the significance of differences in abundance of *Geobacter* between M1A, M2A and M4A for the period between week 8 and 12. For visualization purposes, the five most abundant genera in the inoculum and five most abundant genera in Week 12 were selected. All other genera were included in the “Other” group. The R script employed is described in Orakov *et al*. (2017). The analysis of metagenome diversity was carried out using R version 3.6.0 (need to add ref here and below). Multidimensional scaling was performed with the “dist” and “cmdscale” functions, and MDS (PCoA) plots were generated with ggplot2. For PERMANOVA we used the “adonis” function in the vegan package (Anderson, 2001).

## ACKNOWLEDGEMENTS

This research was supported by Okinawa Institute of Science and Technology Graduate University. We thank Dr. Larisa Kiseleva for collecting electrode and plankton samples and Dr. Toshio Sasaki for preparing SEM specimens.

