## Supplemental Figure S1-S2, Supplemental Table S1-S7 for "Evolutionary dynamics of microbial communities in bioelectrochemical systems"

### Supplementary material

**Figure S1.** Schematic diagram of M1-M4 **a)** side view, **b)** front view and **c)** M5 reactors.

**Figure S2.** Relative abundances of top genera collected from M3 a) working electrode b) counter electrode and c) plankton throughout the time of experiment. Colors represent according taxonomy groups, + - other organisms.

[1] Okinawa Institute of Science and Technology, Japan

[2] Institute of Cell Biophysics, Russia

[3] Institute of Physics and Technology, Russia

[4] University of Liverpool, United Kingdom

[5] The School of Informatics, University of Edinburgh, United Kingdom

[6] Tianjin Institute for Industrial Biotechnology, China

**Table S1.** Sequencing data for all samples.

**Table S2.** Taxonomy analysis of initial community.

**Table S3.** Functional overview of initial community (top 500 rank only).

**Table S4.** Functional overview of **a)** M1A **b)** M1C **c)** M1P communities (top 500 rank only) after 12-week reactor operation.

**Table S5.** Functional overview of **a)** M2A **b)** M2C **c)** M2P communities (top 500 rank only) after 12-week reactor operation.

**Table S6.** Functional overview of **a)** M4A **b)** M4C **c)** M4P communities (top 500 rank only) after 12-week reactor operation.

**Table S7.** Functional overview of M5A communities (top 500 rank only) after 12-week reactor operation.

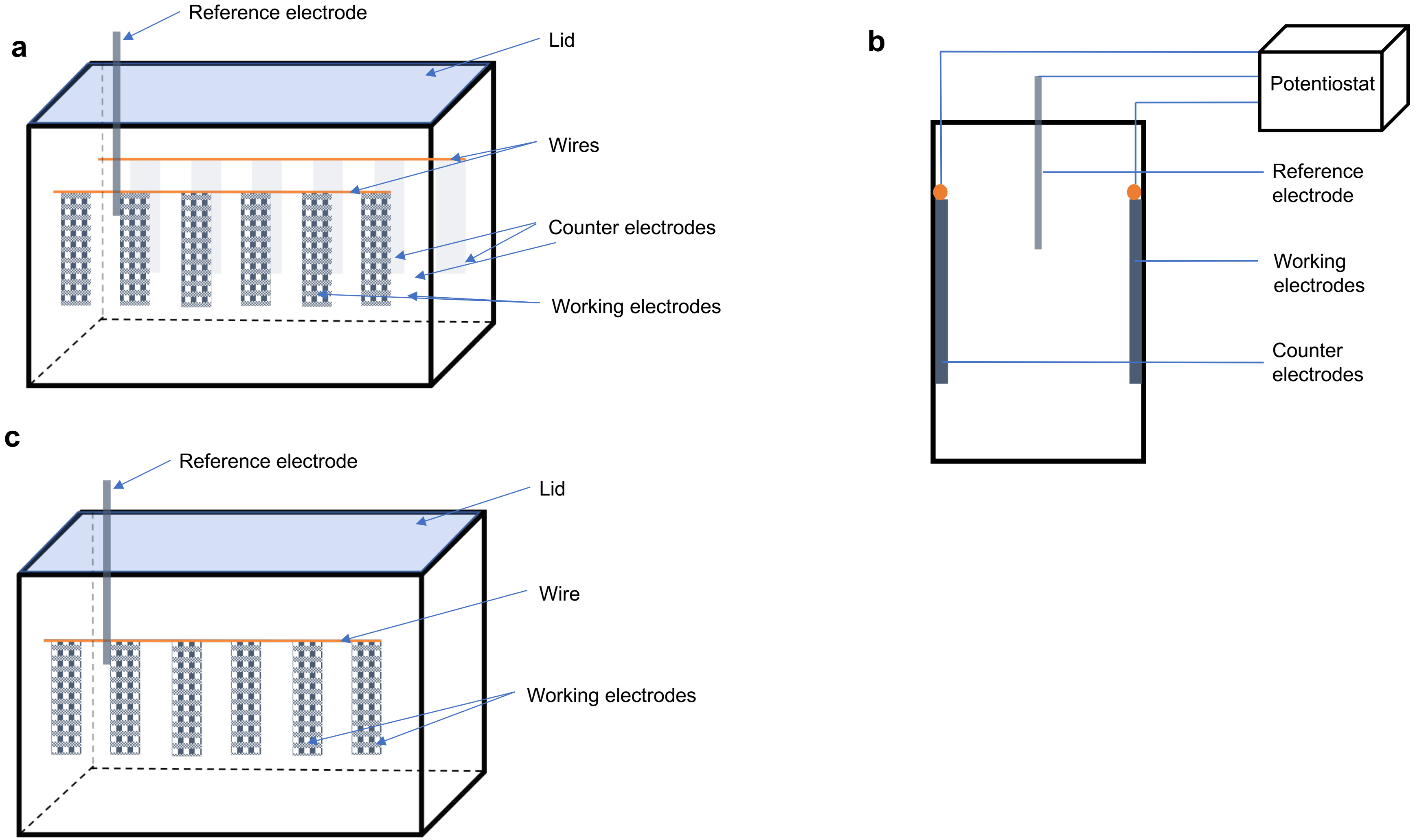


**Fig. S1.** Schematic diagram of M1-M4 **a)** side view, **b)** front view and **c)** M5 reactors.

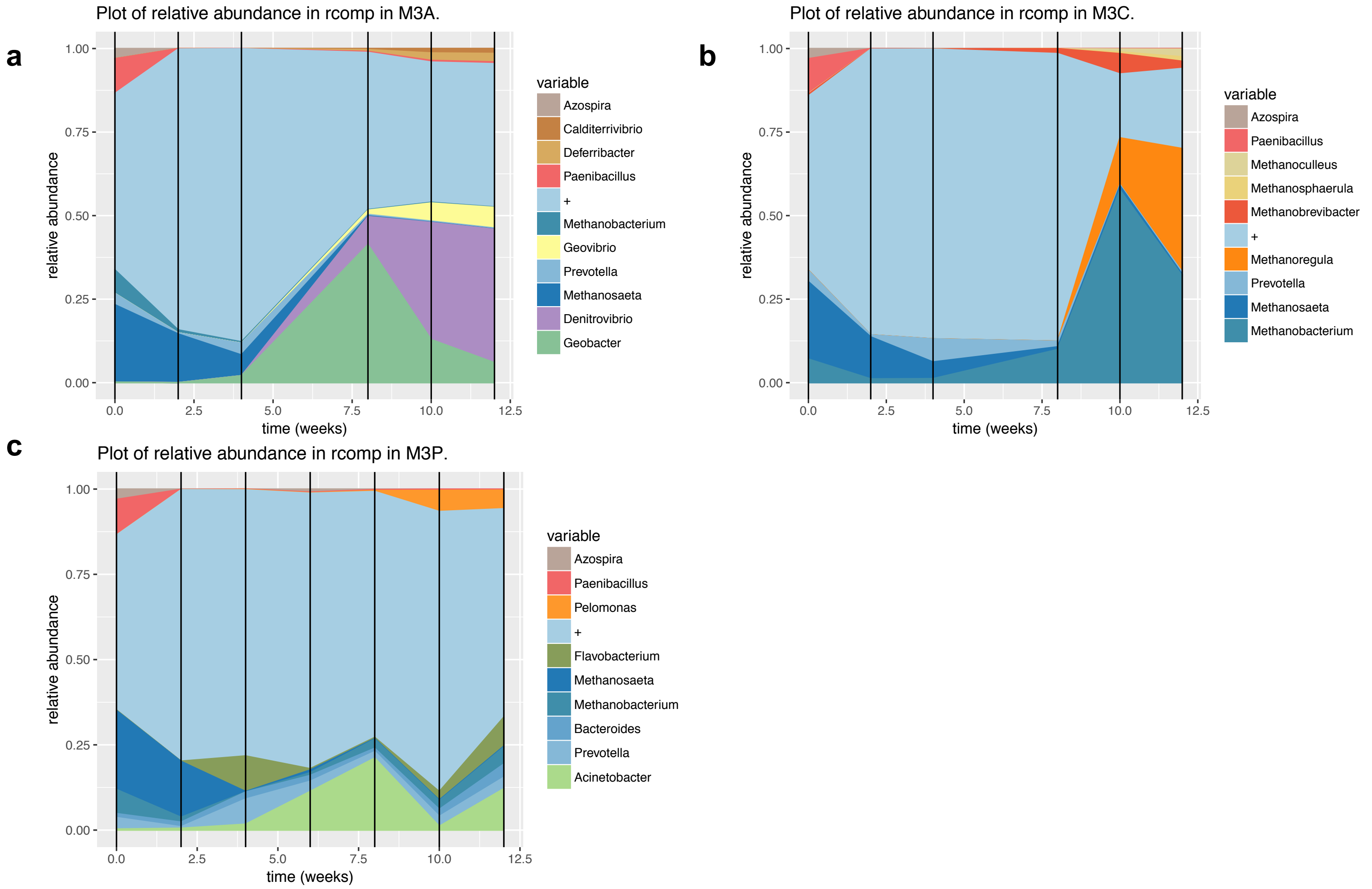


**Fig. S2.** Relative abundances of top genera collected from M3 a) working electrode b) counter electrode and c) plankton throughout the time of experiment. Colors represent according taxonomy groups, + - other organisms.

**Table S1.** Sample sequencing sheet. ND – no data.

| Sample | Sequencing type | Sample | Sequencing type | Sample | Sequencing type |
|---|---|---|---|---|---|
| Initial inoculum | Illumina | 6 weeks M1A | Illumina | 10 weeks M1A | Illumina |
| 2 weeks M1A | 16S | 6 weeks M1C | Illumina | 10 weeks M1C | Illumina |
| 2 weeks M1C | 16S | 6 weeks M1P | Illumina | 10 weeks M1P | Illumina |
| 2 weeks M1P | 16S | 6 weeks M2A | Illumina | 10 weeks M2A | 16S |
| 2 weeks M2A | 16S | 6 weeks M2C | 16S | 10 weeks M2C | Illumina |
| 2 weeks M2C | 16S | 6 weeks M2P | Illumina | 10 weeks M2P | Illumina |
| 2 weeks M2P | 16S | 6 weeks M3A | Illumina | 10 weeks M3A | Illumina |
| 2 weeks M3A | 16S | 6 weeks M3C | ND | 10 weeks M3C | Illumina |
| 2 weeks M3C | 16S | 6 weeks M3P | Illumina | 10 weeks M3P | Illumina |
| 2 weeks M3P | 16S | 6 weeks M4A | Illumina | 10 weeks M4A | Illumina |
| 2 weeks M4A | 16S | 6 weeks M4C | ND | 10 weeks M4C | Illumina |
| 2 weeks M4C | 16S | 6 weeks M4P | Illumina | 10 weeks M4P | Illumina |
| 2 weeks M4P | 16S | 6 weeks M5A | Illumina | 10 weeks M5A | Illumina |
| 2 weeks M5A | 16S | 6 weeks M5P | Illumina | 10 weeks M5P | Illumina |
| 2 weeks M5P | 16S | 8 weeks M1A | Illumina | 12 weeks M1A | Illumina |
| 4 weeks M1A | 16S | 8 weeks M1C | Illumina | 12 weeks M1C | Illumina |
| 4 weeks M1C | 16S | 8 weeks M1P | Illumina | 12 weeks M1P | Illumina |
| 4 weeks M1P | 16S | 8 weeks M2A | Illumina | 12 weeks M2A | Illumina |
| 4 weeks M2A | 16S | 8 weeks M2C | 16S | 12 weeks M2C | Illumina |
| 4 weeks M2C | 16S | 8 weeks M2P | Illumina | 12 weeks M2P | Illumina |
| 4 weeks M2P | 16S | 8 weeks M3A | Illumina | 12 weeks M3A | Illumina |

| | | | | | |
|---|---|---|---|---|---|
| 4 weeks M3A | 16S | 8 weeks M3C | 16S | 12 weeks M3C | Illumina |
| 4 weeks M3C | 16S | 8 weeks M3P | Illumina | 12 weeks M3P | Illumina |
| 4 weeks M3P | 16S | 8 weeks M4A | Illumina | 12 weeks M4A | Illumina |
| 4 weeks M4A | 16S | 8 weeks M4C | ND | 12 weeks M4C | Illumina |
| 4 weeks M4C | 16S | 8 weeks M4P | Illumina | 12 weeks M4P | Illumina |
| 4 weeks M4P | 16S | 8 weeks M5A | 16S | 12 weeks M5A | Illumina |
| 4 weeks M5A | 16S | 8 weeks M5P | Illumina | 12 weeks M5P | Illumina |
| 4 weeks M5P | 16S | | | | |

**Table S2.** Taxonomy analysis of initial community. Genera with less than 100 reads were not considered.

| % | reads | Genus | % | reads | Genus | % | reads | Genus |
|---|---|---|---|---|---|---|---|---|
| 12.59 | 922582 | *Methanosaeta* | 0.01 | 518 | *Acidiphilium* | 0.00 | 211 | *Rothia* |
| 5.67 | 415612 | *Paenibacillus* | 0.01 | 517 | *Haliangium* | 0.00 | 211 | *Kingella* |
| 3.84 | 281366 | *Methanobacterium* | 0.01 | 516 | *Nakamurella* | 0.00 | 210 | *Methanogenium* |
| 1.83 | 133887 | *Prevotella* | 0.01 | 513 | *Castellaniella* | 0.00 | 210 | *Chrysiogenes* |
| 1.65 | 121192 | *Azospira* | 0.01 | 511 | *Chthonomonas* | 0.00 | 210 | *Algibacter* |
| 0.87 | 63773 | *Desulfovibrio* | 0.01 | 510 | *Caldanaerobius* | 0.00 | 209 | *Thiorhodospira* |
| 0.73 | 53428 | *Mesotoga* | 0.01 | 509 | *Ectocarpus* | 0.00 | 208 | *Amphibacillus* |
| 0.70 | 51642 | *Megasphaera* | 0.01 | 509 | *Proteiniclasticum* | 0.00 | 208 | *Balneola* |
| 0.68 | 49705 | *Streptomyces* | 0.01 | 508 | *Absidia* | 0.00 | 208 | *Cesiribacter* |
| 0.66 | 48416 | *Candidatus Accumulibacter* | 0.01 | 508 | *Thiohalocapsa* | 0.00 | 207 | *Cyclospora* |
| 0.63 | 46281 | *Bacteroides* | 0.01 | 507 | *Pseudobutyrivibrio* | 0.00 | 207 | *Neonectria* |
| 0.60 | 44138 | *Clostridium* | 0.01 | 507 | *Alkaliflexus* | 0.00 | 207 | *Bhargavaea* |
| 0.58 | 42636 | *Lentimicrobium* | 0.01 | 507 | *Alkalitalea* | 0.00 | 207 | *Jatrophihabitans* |
| 0.43 | 31511 | *Pseudomonas* | 0.01 | 505 | *Pseudovibrio* | 0.00 | 206 | *Paxillus* |
| 0.39 | 28532 | *Azovibrio* | 0.01 | 502 | *Cladophialophora* | 0.00 | 206 | *Olivibacter* |
| 0.34 | 24617 | *Mycobacterium* | 0.01 | 500 | *Salegentibacter* | 0.00 | 206 | *Clostridiisalibacter* |
| 0.33 | 23953 | *Methanobrevibacter* | 0.01 | 499 | *Dickeya* | 0.00 | 206 | *Paramaledivibacter* |
| 0.32 | 23207 | *Methanosarcina* | 0.01 | 498 | *Oscillatoria* | 0.00 | 205 | *Thiocystis* |
| 0.27 | 19922 | *Oscillibacter* | 0.01 | 495 | *Thermophagus* | 0.00 | 205 | *Citromicrobium* |
| 0.27 | 19921 | *Dechloromonas* | 0.01 | 494 | *Heliobacterium* | 0.00 | 205 | *Plantibacter* |
| 0.25 | 18513 | *Parabacteroides* | 0.01 | 494 | *Cryobacterium* | 0.00 | 205 | *Stomatobaculum* |
| 0.23 | 16893 | *Bacillus* | 0.01 | 492 | *Vitrella* | 0.00 | 204 | *Stentor* |
| 0.22 | 15850 | *Burkholderia* | 0.01 | 490 | *Metarhizium* | 0.00 | 204 | *Emmonsia* |
| 0.21 | 15561 | *Methanothermobacter* | 0.01 | 490 | *Snodgrassella* | 0.00 | 204 | *Actinobaculum* |
| 0.20 | 14845 | *Thauera* | 0.01 | 488 | *Acidaminobacter* | 0.00 | 204 | *Thermoactinospora* |
| 0.20 | 14453 | *Rhodococcus* | 0.01 | 487 | *Pelotomaculum* | 0.00 | 204 | *Wenyingzhuangia* |
| 0.19 | 14140 | *Rhizobium* | 0.01 | 485 | *Lamprocystis* | 0.00 | 203 | *Thermomicrobium* |
| 0.18 | 13022 | *Methylocystis* | 0.01 | 485 | *Lampropedia* | 0.00 | 203 | *Phlebia* |
| 0.18 | 12861 | *Propionivibrio* | 0.01 | 485 | *Thermincola* | 0.00 | 203 | *Pleurotus* |
| 0.17 | 12729 | *Eubacterium* | 0.01 | 485 | *Candidatus Magnetoglobus* | 0.00 | 203 | *Auraticoccus* |
| 0.17 | 12318 | *Acinetobacter* | 0.01 | 484 | *Geoalkalibacter* | 0.00 | 202 | *Herbiconiux* |
| 0.16 | 11663 | *Azoarcus* | 0.01 | 483 | *Ferrimonas* | 0.00 | 202 | *Schleiferia* |
| 0.16 | 11530 | *Treponema* | 0.01 | 482 | *Senegalimassilia* | 0.00 | 202 | *Catalinimonas* |

| | | | | | | | | |
|---|---|---|---|---|---|---|---|---|
| 0.15 | 11239 | *Aminobacterium* | 0.01 | 479 | *Candidatus Entotheonella* | 0.00 | 201 | *Desulfurella* |
| 0.15 | 11036 | *Alistipes* | 0.01 | 478 | *Flammeovirga* | 0.00 | 201 | *Terasakiella* |
| 0.15 | 10993 | *Sphaerochaeta* | 0.01 | 478 | *Virgibacillus* | 0.00 | 201 | *Caldanaerobacter* |
| 0.14 | 10608 | *Smithella* | 0.01 | 472 | *Leminorella* | 0.00 | 201 | *Tamlana* |
| 0.14 | 10311 | *Syntrophorhabdus* | 0.01 | 472 | *Lutibacter* | 0.00 | 200 | *Candidatus Microthrix* |
| 0.14 | 10104 | *Azonexus* | 0.01 | 471 | *Bartonella* | 0.00 | 200 | *Roseateles* |
| 0.13 | 9504 | *Acidovorax* | 0.01 | 471 | *Planktothrix* | 0.00 | 200 | *Thermoleophilum* |
| 0.11 | 8249 | *Dehalobacter* | 0.01 | 468 | *Gonapodya* | 0.00 | 200 | *Geothermobacter* |
| 0.11 | 7973 | *Lactobacillus* | 0.01 | 467 | *Lentzea* | 0.00 | 199 | *Rubellimicrobium* |
| 0.11 | 7912 | *Zoogloea* | 0.01 | 465 | *Desulfobacula* | 0.00 | 199 | *Caenispirillum* |
| 0.10 | 7145 | *Syntrophobacter* | 0.01 | 465 | *Fuerstia* | 0.00 | 199 | *Candidatus Izimaplasma* |
| 0.10 | 7055 | *Ruminococcus* | 0.01 | 464 | *Succiniclasticum* | 0.00 | 198 | *Pestalotiopsis* |
| 0.10 | 7016 | *Kosmotoga* | 0.01 | 463 | *Giesbergeria* | 0.00 | 198 | *Granulicoccus* |
| 0.09 | 6924 | *Desulfomonile* | 0.01 | 463 | *Pararhodospirillum* | 0.00 | 198 | *Baudoinia* |
| 0.09 | 6529 | *Desulfotomaculum* | 0.01 | 462 | *Limnochorda* | 0.00 | 197 | *Tilletiaria* |
| 0.09 | 6523 | *Bradyrhizobium* | 0.01 | 461 | *Azorhizobium* | 0.00 | 197 | *Aestuariimicrobium* |
| 0.09 | 6491 | *Geobacter* | 0.01 | 460 | *Nonlabens* | 0.00 | 197 | *Caldibacillus* |
| 0.09 | 6278 | *Achromobacter* | 0.01 | 459 | *Bodo* | 0.00 | 197 | *Silvibacterium* |
| 0.09 | 6240 | *Actinomyces* | 0.01 | 459 | *Brachymonas* | 0.00 | 196 | *Grifola* |
| 0.08 | 6165 | *Methyloversatilis* | 0.01 | 459 | *Myroides* | 0.00 | 196 | *Algiphilus* |
| 0.08 | 6138 | *Flavobacterium* | 0.01 | 459 | *Asticcacaulis* | 0.00 | 196 | *Arcticibacter* |
| 0.08 | 6132 | *Sphingomonas* | 0.01 | 458 | *Planctomicrobium* | 0.00 | 195 | *Edwardsiella* |
| 0.08 | 6126 | *Anaerolinea* | 0.01 | 457 | *Herpetosiphon* | 0.00 | 195 | *Kytococcus* |
| 0.08 | 6022 | *Paenirhodobacter* | 0.01 | 457 | *Methanosalsum* | 0.00 | 195 | *Batrachochytrium* |
| 0.08 | 5933 | *Syntrophus* | 0.01 | 457 | *Luteibacter* | 0.00 | 195 | *Ornithinibacillus* |
| 0.08 | 5818 | *Microbacterium* | 0.01 | 457 | *Gorillibacterium* | 0.00 | 195 | *Nitrososphaera* |
| 0.07 | 5407 | *Nocardia* | 0.01 | 456 | *Crocinitomix* | 0.00 | 195 | *Halogranum* |
| 0.07 | 5406 | *Aminiphilus* | 0.01 | 456 | *Mahella* | 0.00 | 195 | *Croceicoccus* |
| 0.07 | 5348 | *Syntrophomonas* | 0.01 | 454 | *Proteiniborus* | 0.00 | 194 | *Helicosporidium* |
| 0.07 | 5285 | *Candidatus Cloacimonas* | 0.01 | 454 | *Roseimaritima* | 0.00 | 194 | *Serinicoccus* |
| 0.07 | 5270 | *Crenobacter* | 0.01 | 453 | *Leptothrix* | 0.00 | 194 | *Pseudoduganella* |
| 0.07 | 5162 | *Nitrosomonas* | 0.01 | 452 | *Anaerosporomusa* | 0.00 | 194 | *Phaeodactylibacter* |
| 0.07 | 5156 | *Micromonospora* | 0.01 | 451 | *Pseudanabaena* | 0.00 | 194 | *Massilibacterium* |
| 0.07 | 5151 | *Massilia* | 0.01 | 451 | *Candidatus Symbiothrix* | 0.00 | 193 | *Vagococcus* |
| 0.07 | 5067 | *Thiobacillus* | 0.01 | 451 | *Gallaecimonas* | 0.00 | 193 | *Piscicoccus* |
| 0.07 | 4973 | *Paraburkholderia* | 0.01 | 450 | *Altererythrobacter* | 0.00 | 193 | *Gynuella* |

| | | | | | | | | |
|---|---|---|---|---|---|---|---|---|
| 0.07 | 4832 | *Ruminiclostridium* | 0.01 | 449 | *Exidia* | 0.00 | 193 | *Phocea* |
| 0.07 | 4830 | *Flexilinea* | 0.01 | 449 | *Petrimonas* | 0.00 | 192 | *Phytomonas* |
| 0.06 | 4716 | *Dethiosulfovibrio* | 0.01 | 448 | *Psychroflexus* | 0.00 | 192 | *Desulfohalobium* |
| 0.06 | 4671 | *Methylococcus* | 0.01 | 447 | *Aphanomyces* | 0.00 | 192 | *Facklamia* |
| 0.06 | 4564 | *Herminiimonas* | 0.01 | 447 | *Minicystis* | 0.00 | 192 | *Pannonibacter* |
| 0.06 | 4522 | *Vibrio* | 0.01 | 446 | *Nitrosococcus* | 0.00 | 192 | *Salinarimonas* |
| 0.06 | 4502 | *Cupriavidus* | 0.01 | 446 | *Nevskia* | 0.00 | 192 | *Mageeibacillus* |
| 0.06 | 4499 | *Chromobacterium* | 0.01 | 444 | *Diplodia* | 0.00 | 192 | *Cephaloticoccus* |
| 0.06 | 4493 | *Olsenella* | 0.01 | 444 | *Idiomarina* | 0.00 | 190 | *Luteococcus* |
| 0.06 | 4487 | *Bordetella* | 0.01 | 444 | *Gemmobacter* | 0.00 | 190 | *Chroococcidiopsis* |
| 0.06 | 4393 | *Methanoculleus* | 0.01 | 444 | *Advenella* | 0.00 | 190 | *Tremella* |
| 0.06 | 4365 | *Herbaspirillum* | 0.01 | 442 | *Psychromonas* | 0.00 | 190 | *Kordia* |
| 0.06 | 4329 | *Sulfurovum* | 0.01 | 442 | *Hammondia* | 0.00 | 190 | *Simplicispira* |
| 0.06 | 4241 | *Pedobacter* | 0.01 | 440 | *Laccaria* | 0.00 | 190 | *Kiloniella* |
| 0.06 | 4231 | *Methylomonas* | 0.01 | 440 | *Rubrobacter* | 0.00 | 190 | *Candidatus Magnetoovum* |
| 0.06 | 4206 | *Desulfosporosinus* | 0.01 | 438 | *Mortierella* | 0.00 | 189 | *Bizionia* |
| 0.06 | 4183 | *Xanthomonas* | 0.01 | 438 | *Carboxydocella* | 0.00 | 189 | *Grosmannia* |
| 0.06 | 4091 | *Chryseobacterium* | 0.01 | 437 | *Pusillimonas* | 0.00 | 189 | *Aquincola* |
| 0.05 | 4012 | *Methylobacterium* | 0.01 | 436 | *Xenorhabdus* | 0.00 | 189 | *Aggregatibacter* |
| 0.05 | 3962 | *Aminomonas* | 0.01 | 436 | *Pelagibacterium* | 0.00 | 188 | *Borrelia* |
| 0.05 | 3939 | *Frankia* | 0.01 | 436 | *Desulfamplus* | 0.00 | 188 | *Catonella* |
| 0.05 | 3885 | *Lachnoclostridium* | 0.01 | 436 | *Candidatus Thiodiazotropha* | 0.00 | 188 | *Desulfonispora* |
| 0.05 | 3858 | *Ralstonia* | 0.01 | 436 | *Arabia* | 0.00 | 188 | *Rhizoclosmatium* |
| 0.05 | 3837 | *Synergistes* | 0.01 | 435 | *Rhodothermus* | 0.00 | 188 | *Caminicella* |
| 0.05 | 3821 | *Legionella* | 0.01 | 435 | *Gaeumannomyces* | 0.00 | 188 | *Proteocatella* |
| 0.05 | 3818 | *Corynebacterium* | 0.01 | 434 | *Cyanobium* | 0.00 | 188 | *Thalassobacillus* |
| 0.05 | 3734 | *Novosphingobium* | 0.01 | 433 | *Herbidospora* | 0.00 | 188 | *Moelleriella* |
| 0.05 | 3682 | *Mesorhizobium* | 0.01 | 432 | *Microlunatus* | 0.00 | 187 | *Flexibacter* |
| 0.05 | 3608 | *Uliginosibacterium* | 0.01 | 431 | *Leptospirillum* | 0.00 | 187 | *Schizophyllum* |
| 0.05 | 3545 | *Cloacibacillus* | 0.01 | 431 | *Dietzia* | 0.00 | 187 | *Microbotryum* |
| 0.05 | 3500 | *Arthrobacter* | 0.01 | 431 | *Glycomyces* | 0.00 | 187 | *Ferroglobus* |
| 0.05 | 3464 | *Thiomonas* | 0.01 | 430 | *Desulforhopalus* | 0.00 | 187 | *Adhaeribacter* |
| 0.05 | 3435 | *Levilinea* | 0.01 | 430 | *Hahella* | 0.00 | 187 | *Terrisporobacter* |
| 0.05 | 3426 | *Azospirillum* | 0.01 | 430 | *Methanotorris* | 0.00 | 186 | *Phyllobacterium* |
| 0.05 | 3410 | *Butyrivibrio* | 0.01 | 430 | *Orrella* | 0.00 | 186 | *Psychroserpens* |
| 0.04 | 3250 | *Variovorax* | 0.01 | 430 | *Kiritimatiella* | 0.00 | 186 | *Malassezia* |

| | | | | | | | | |
|---|---|---|---|---|---|---|---|---|
| 0.04 | 3231 | *Longilinea* | 0.01 | 429 | *Chlorella* | 0.00 | 186 | *Rhizophagus* |
| 0.04 | 3169 | *Porphyromonas* | 0.01 | 428 | *Polaribacter* | 0.00 | 186 | *Cutaneotrichosporon* |
| 0.04 | 3159 | *Eimeria* | 0.01 | 428 | *Alicycliphilus* | 0.00 | 185 | *Cycloclasticus* |
| 0.04 | 3140 | *Pseudonocardia* | 0.01 | 427 | *Hydrogenoanaerobacterium* | 0.00 | 185 | *Ustilaginoidea* |
| 0.04 | 3064 | *Dysgonomonas* | 0.01 | 427 | *Verticillium* | 0.00 | 185 | *Rubritepida* |
| 0.04 | 3062 | *Janthinobacterium* | 0.01 | 425 | *Mobilibacterium* | 0.00 | 185 | *Endomicrobium* |
| 0.04 | 3049 | *Comamonas* | 0.01 | 424 | *Enterovibrio* | 0.00 | 185 | *Wenzhouxiangella* |
| 0.04 | 3045 | *Chitinophaga* | 0.01 | 423 | *Sporocytophaga* | 0.00 | 184 | *Rikenella* |
| 0.04 | 3035 | *Aquaspirillum* | 0.01 | 421 | *Propionispira* | 0.00 | 184 | *Macrophomina* |
| 0.04 | 3002 | *Halomonas* | 0.01 | 421 | *Chthoniobacter* | 0.00 | 184 | *Desulfurispirillum* |
| 0.04 | 2998 | *Streptococcus* | 0.01 | 421 | *Lachancea* | 0.00 | 184 | *Lishizhenia* |
| 0.04 | 2984 | *Derxia* | 0.01 | 421 | *Saccharicrinis* | 0.00 | 183 | *Teredinibacter* |
| 0.04 | 2916 | *Curvibacter* | 0.01 | 420 | *Methanocorpusculum* | 0.00 | 183 | *Rhinocladiella* |
| 0.04 | 2900 | *Hyphomicrobium* | 0.01 | 420 | *Marinilabilia* | 0.00 | 183 | *Heterobasidion* |
| 0.04 | 2855 | *Methanoregula* | 0.01 | 420 | *Caldicoprobacter* | 0.00 | 183 | *Cyanobacterium* |
| 0.04 | 2808 | *Leptolinea* | 0.01 | 418 | *Bipolaris* | 0.00 | 183 | *Novispirillum* |
| 0.04 | 2771 | *Amycolatopsis* | 0.01 | 418 | *Prosthecobacter* | 0.00 | 183 | *Novibacillus* |
| 0.04 | 2723 | *Acidaminococcus* | 0.01 | 418 | *Candidatus Koribacter* | 0.00 | 182 | *Thermosediminibacter* |
| 0.04 | 2703 | *Methanocella* | 0.01 | 417 | *Halorubrum* | 0.00 | 182 | *Pseudosphingobacterium* |
| 0.04 | 2697 | *Stenotrophomonas* | 0.01 | 415 | *Pyrococcus* | 0.00 | 182 | *Schizopora* |
| 0.04 | 2677 | *Azotobacter* | 0.01 | 415 | *Fictibacillus* | 0.00 | 181 | *Albugo* |
| 0.04 | 2677 | *Rhodobacter* | 0.01 | 414 | *Fonsecaea* | 0.00 | 181 | *Coniophora* |
| 0.04 | 2669 | *Sulfuritalea* | 0.01 | 414 | *Kutzneria* | 0.00 | 181 | *Blastomyces* |
| 0.04 | 2659 | *Methanosphaera* | 0.01 | 414 | *Conexibacter* | 0.00 | 181 | *Geopsychrobacter* |
| 0.04 | 2625 | *Faecalibacterium* | 0.01 | 414 | *Gottschalkia* | 0.00 | 181 | *Pyrenochaeta* |
| 0.03 | 2559 | *Marinobacter* | 0.01 | 413 | *Providencia* | 0.00 | 181 | *Caldalkalibacillus* |
| 0.03 | 2528 | *Acetobacter* | 0.01 | 413 | *Bryobacter* | 0.00 | 180 | *Spirillospora* |
| 0.03 | 2516 | *Methanothermus* | 0.01 | 412 | *Geobacillus* | 0.00 | 180 | *Planktothricoides* |
| 0.03 | 2509 | *Spirochaeta* | 0.01 | 412 | *Kribbella* | 0.00 | 180 | *Actinocatenispora* |
| 0.03 | 2499 | *Ideonella* | 0.01 | 410 | *Kerstersia* | 0.00 | 179 | *Haladaptatus* |
| 0.03 | 2499 | *Paludibacter* | 0.01 | 410 | *Zavarzinella* | 0.00 | 178 | *Podospora* |
| 0.03 | 2489 | *Nocardioides* | 0.01 | 410 | *Paludisphaera* | 0.00 | 178 | *Riemerella* |
| 0.03 | 2445 | *Thermanaerovibrio* | 0.01 | 409 | *Acanthamoeba* | 0.00 | 178 | *Gluconacetobacter* |
| 0.03 | 2431 | *Hydrogenophaga* | 0.01 | 409 | *Gloeobacter* | 0.00 | 178 | *Madurella* |
| 0.03 | 2423 | *Cellulomonas* | 0.01 | 409 | *Trichococcus* | 0.00 | 178 | *Gallibacterium* |
| 0.03 | 2421 | *Bifidobacterium* | 0.01 | 409 | *Terriglobus* | 0.00 | 178 | *Peptoanaerobacter* |

| | | | | | | | | |
|---|---|---|---|---|---|---|---|---|
| 0.03 | 2408 | *Aspergillus* | 0.01 | 408 | *Gracilibacillus* | 0.00 | 177 | *Gemella* |
| 0.03 | 2393 | *Pandoraea* | 0.01 | 407 | *Actinoalloteichus* | 0.00 | 177 | *Fomitopsis* |
| 0.03 | 2385 | *Desulfacinum* | 0.01 | 406 | *Tolypothrix* | 0.00 | 177 | *Arthrospira* |
| 0.03 | 2372 | *Sphingobacterium* | 0.01 | 406 | *Anaerostipes* | 0.00 | 177 | *Daedalea* |
| 0.03 | 2371 | *Mucilaginibacter* | 0.01 | 406 | *Methyloceanibacter* | 0.00 | 177 | *Empedobacter* |
| 0.03 | 2362 | *Caulobacter* | 0.01 | 405 | *Rhodospirillum* | 0.00 | 177 | *Rivularia* |
| 0.03 | 2340 | *Papillibacter* | 0.01 | 404 | *Taylorella* | 0.00 | 177 | *Nitriliruptor* |
| 0.03 | 2339 | *Paracoccus* | 0.01 | 404 | *Phialocephala* | 0.00 | 176 | *Trichosporon* |
| 0.03 | 2329 | *Enterococcus* | 0.01 | 404 | *Fluviicola* | 0.00 | 176 | *Acidomonas* |
| 0.03 | 2325 | *Selenomonas* | 0.01 | 403 | *Psychrobacter* | 0.00 | 176 | *Saccharibacillus* |
| 0.03 | 2303 | *Nocardiopsis* | 0.01 | 403 | *Allochromatium* | 0.00 | 176 | *Lachnoanaerobaculum* |
| 0.03 | 2261 | *Methanolinea* | 0.01 | 403 | *Arenibacter* | 0.00 | 176 | *Fervidicella* |
| 0.03 | 2251 | *Deinococcus* | 0.01 | 402 | *Barnesiella* | 0.00 | 176 | *Apibacter* |
| 0.03 | 2249 | *Christensenella* | 0.01 | 402 | *Desulfopila* | 0.00 | 175 | *Giardia* |
| 0.03 | 2238 | *Acetomicrobium* | 0.01 | 401 | *Gimesia* | 0.00 | 175 | *Oerskovia* |
| 0.03 | 2233 | *Gordonia* | 0.01 | 400 | *Salpingoeca* | 0.00 | 175 | *Jaapia* |
| 0.03 | 2214 | *Tepidiphilus* | 0.01 | 400 | *Pectobacterium* | 0.00 | 175 | *Pseudodonghicola* |
| 0.03 | 2213 | *Rhodopseudomonas* | 0.01 | 399 | *Microcystis* | 0.00 | 175 | *Lunatimonas* |
| 0.03 | 2208 | *Porphyra* | 0.01 | 398 | *Shinella* | 0.00 | 175 | *Traorella* |
| 0.03 | 2195 | *Ornatilinea* | 0.01 | 397 | *Streptosporangium* | 0.00 | 174 | *Ascosphaera* |
| 0.03 | 2180 | *Arcobacter* | 0.01 | 397 | *Desulfonatronovibrio* | 0.00 | 174 | *Kinetoplastibacterium* |
| 0.03 | 2139 | *Desulfocurvus* | 0.01 | 397 | *Candidatus Contendobacter* | 0.00 | 174 | *Clavispora* |
| 0.03 | 2082 | *Desulfitobacterium* | 0.01 | 395 | *Candidatus Tenderia* | 0.00 | 174 | *Ketogulonicigenium* |
| 0.03 | 2081 | *Hymenobacter* | 0.01 | 393 | *Haemophilus* | 0.00 | 174 | *Anaeroarcus* |
| 0.03 | 2054 | *Shewanella* | 0.01 | 393 | *Syntrophobotulus* | 0.00 | 174 | *Ogataea* |
| 0.03 | 2037 | *Actinomadura* | 0.01 | 393 | *Desulfonauticus* | 0.00 | 173 | *Hanseniaspora* |
| 0.03 | 2014 | *Desulfobulbus* | 0.01 | 393 | *Sandaracinus* | 0.00 | 173 | *Magnaporthiopsis* |
| 0.03 | 2013 | *Thioalkalivibrio* | 0.01 | 391 | *Tissierella* | 0.00 | 173 | *Hugenholtzia* |
| 0.03 | 2006 | *Methylotenera* | 0.01 | 391 | *Domibacillus* | 0.00 | 172 | *Gelatoporia* |
| 0.03 | 1990 | *Alicyclobacillus* | 0.01 | 390 | *Phormidium* | 0.00 | 172 | *Pochonia* |
| 0.03 | 1984 | *Pseudogulbenkiania* | 0.01 | 389 | *Halolactibacillus* | 0.00 | 172 | *Edaphobacter* |
| 0.03 | 1979 | *Thioclava* | 0.01 | 386 | *Modestobacter* | 0.00 | 172 | *Sugiyamaella* |
| 0.03 | 1973 | *Leptospira* | 0.01 | 384 | *Chrysochromulina* | 0.00 | 172 | *Succinatimonas* |
| 0.03 | 1971 | *Actinoplanes* | 0.01 | 383 | *Salinisphaera* | 0.00 | 172 | *Sinosporangium* |
| 0.03 | 1968 | *Lautropia* | 0.01 | 382 | *Akkermansia* | 0.00 | 172 | *Hydrogenibacillus* |
| 0.03 | 1966 | *Nitrospira* | 0.01 | 381 | *Auricularia* | 0.00 | 171 | *Gloeophyllum* |

| | | | | | | | | |
|---|---|---|---|---|---|---|---|---|
| 0.03 | 1959 | *Eggerthella* | 0.01 | 380 | *Desulfospira* | 0.00 | 171 | *Postia* |
| 0.03 | 1958 | *Anaerotruncus* | 0.01 | 380 | *Kouleothrix* | 0.00 | 171 | *Pseudopropionibacterium* |
| 0.03 | 1956 | *Delftia* | 0.01 | 379 | *Leucosporidium* | 0.00 | 170 | *Catenuloplanes* |
| 0.03 | 1954 | *Methanospirillum* | 0.01 | 379 | *Guillardia* | 0.00 | 170 | *Aquiflexum* |
| 0.03 | 1953 | *Campylobacter* | 0.01 | 378 | *Saprolegnia* | 0.00 | 170 | *Obba* |
| 0.03 | 1940 | *Nitrosospira* | 0.01 | 378 | *Natronincola* | 0.00 | 170 | *Coniosporium* |
| 0.03 | 1887 | *Pelobacter* | 0.01 | 378 | *Actinophytocola* | 0.00 | 170 | *Ileibacterium* |
| 0.03 | 1884 | *Algoriphagus* | 0.01 | 377 | *Lutispora* | 0.00 | 169 | *Methanomicrobium* |
| 0.03 | 1880 | *Sulfuricella* | 0.01 | 375 | *Skermanella* | 0.00 | 169 | *Amanita* |
| 0.03 | 1876 | *Kitasatospora* | 0.01 | 373 | *Thermoactinomyces* | 0.00 | 169 | *Pyronema* |
| 0.03 | 1875 | *Devosia* | 0.01 | 373 | *Tindallia* | 0.00 | 169 | *Ferroplasma* |
| 0.03 | 1870 | *Collinsella* | 0.01 | 373 | *Deefgea* | 0.00 | 169 | *Acremonium* |
| 0.03 | 1866 | *Neisseria* | 0.01 | 372 | *Ustilago* | 0.00 | 169 | *Pelagibaca* |
| 0.03 | 1851 | *Pirellula* | 0.01 | 372 | *Gilliamella* | 0.00 | 169 | *Ceraceosorus* |
| 0.03 | 1849 | *Rhodopirellula* | 0.01 | 372 | *Mariniblastus* | 0.00 | 169 | *Terribacillus* |
| 0.03 | 1844 | *Nonomuraea* | 0.01 | 370 | *Coriobacterium* | 0.00 | 168 | *Methanocalculus* |
| 0.03 | 1841 | *Desulfuromonas* | 0.01 | 369 | *Sulfolobus* | 0.00 | 168 | *Asanoa* |
| 0.03 | 1835 | *Sphingobium* | 0.01 | 369 | *Salinicoccus* | 0.00 | 168 | *Allisonella* |
| 0.03 | 1833 | *Candidatus Methanoperedens* | 0.01 | 369 | *Thiorhodovibrio* | 0.00 | 168 | *Enhydrobacter* |
| 0.02 | 1828 | *Polaromonas* | 0.01 | 368 | *Fournierella* | 0.00 | 168 | *Thermodesulfobium* |
| 0.02 | 1817 | *Synechococcus* | 0.01 | 367 | *Flavisolibacter* | 0.00 | 168 | *Candidatus Kuenenia* |
| 0.02 | 1814 | *Thermotoga* | 0.01 | 367 | *Intestinimonas* | 0.00 | 168 | *Sharpea* |
| 0.02 | 1810 | *Thermovirga* | 0.00 | 366 | *Bulleidia* | 0.00 | 167 | *Pichia* |
| 0.02 | 1803 | *Blautia* | 0.00 | 365 | *Desulfovermiculus* | 0.00 | 167 | *Ilyobacter* |
| 0.02 | 1802 | *Roseburia* | 0.00 | 363 | *Pelodictyon* | 0.00 | 167 | *Silanimonas* |
| 0.02 | 1787 | *Leucobacter* | 0.00 | 363 | *Neocallimastix* | 0.00 | 167 | *Marssonina* |
| 0.02 | 1780 | *Sporomusa* | 0.00 | 362 | *Candidatus Competibacter* | 0.00 | 167 | *Hoyosella* |
| 0.02 | 1769 | *Pseudoflavonifractor* | 0.00 | 362 | *Candidatus Methylopumilus* | 0.00 | 166 | *Kluyveromyces* |
| 0.02 | 1765 | *Methanolobus* | 0.00 | 361 | *Rhodocista* | 0.00 | 166 | *Tepidimicrobium* |
| 0.02 | 1763 | *Candidatus Aegiribacteria* | 0.00 | 361 | *Fusibacter* | 0.00 | 166 | *Phaeobacter* |
| 0.02 | 1751 | *Bosea* | 0.00 | 361 | *Aequorivita* | 0.00 | 166 | *Cutibacterium* |
| 0.02 | 1750 | *Methylobacillus* | 0.00 | 360 | *Peptostreptococcus* | 0.00 | 165 | *Thermomonospora* |
| 0.02 | 1748 | *Aquitalea* | 0.00 | 360 | *Methanohalobium* | 0.00 | 165 | *Acetohalobium* |
| 0.02 | 1741 | *Desulfococcus* | 0.00 | 360 | *Thermanaeromonas* | 0.00 | 165 | *Succinivibrio* |
| 0.02 | 1739 | *Agrobacterium* | 0.00 | 358 | *Reticulomyxa* | 0.00 | 164 | *Terrabacter* |

| | | | | | | | | |
|---|---|---|---|---|---|---|---|---|
| 0.02 | 1724 | *Duganella* | 0.00 | 358 | *Neglecta* | 0.00 | 164 | *Phlebiopsis* |
| 0.02 | 1718 | *Helicobacter* | 0.00 | 357 | *Muricauda* | 0.00 | 164 | *Truepera* |
| 0.02 | 1713 | *Chitinimonas* | 0.00 | 357 | *Herbinix* | 0.00 | 164 | *Fontibacillus* |
| 0.02 | 1707 | *Aeromonas* | 0.00 | 356 | *Porphyrobacter* | 0.00 | 164 | *Hamadaea* |
| 0.02 | 1698 | *Rugosibacter* | 0.00 | 356 | *Acetonema* | 0.00 | 164 | *Catenovulum* |
| 0.02 | 1697 | *Plasmodium* | 0.00 | 356 | *Frateuria* | 0.00 | 163 | *Deferribacter* |
| 0.02 | 1696 | *Lysobacter* | 0.00 | 355 | *Haliscomenobacter* | 0.00 | 163 | *Parvibaculum* |
| 0.02 | 1695 | *Rhodoferax* | 0.00 | 355 | *Cordyceps* | 0.00 | 163 | *Kordiimonas* |
| 0.02 | 1693 | *Penicillium* | 0.00 | 355 | *Halobacillus* | 0.00 | 163 | *Phycisphaera* |
| 0.02 | 1689 | *Colletotrichum* | 0.00 | 355 | *Anaerovibrio* | 0.00 | 163 | *Coleofasciculus* |
| 0.02 | 1688 | *Polynucleobacter* | 0.00 | 355 | *Inquilinus* | 0.00 | 163 | *Caldisalinibacter* |
| 0.02 | 1680 | *Desulfobacterium* | 0.00 | 351 | *Symbiobacterium* | 0.00 | 163 | *Arsenicibacter* |
| 0.02 | 1670 | *Acidithiobacillus* | 0.00 | 351 | *Calocera* | 0.00 | 162 | *Pimelobacter* |
| 0.02 | 1662 | *Phenylobacterium* | 0.00 | 351 | *Moorea* | 0.00 | 162 | *Methanopyrus* |
| 0.02 | 1660 | *Anaerovorax* | 0.00 | 349 | *Sulfobacillus* | 0.00 | 162 | *Schwartzia* |
| 0.02 | 1651 | *Bellilinea* | 0.00 | 347 | *Peniophora* | 0.00 | 162 | *Humibacter* |
| 0.02 | 1646 | *Sorangium* | 0.00 | 347 | *Congregibacter* | 0.00 | 162 | *Aliivibrio* |
| 0.02 | 1632 | *Collimonas* | 0.00 | 347 | *Desulfuribacillus* | 0.00 | 162 | *Verruconis* |
| 0.02 | 1602 | *Pseudoalteromonas* | 0.00 | 346 | *Anabaena* | 0.00 | 162 | *Hathewaya* |
| 0.02 | 1568 | *Atopobium* | 0.00 | 346 | *Thermoanaerobaculum* | 0.00 | 161 | *Polysphondylium* |
| 0.02 | 1558 | *Marinitoga* | 0.00 | 344 | *Planctopirus* | 0.00 | 161 | *Methylopila* |
| 0.02 | 1557 | *Andreprevotia* | 0.00 | 343 | *Blastocystis* | 0.00 | 161 | *Starkeya* |
| 0.02 | 1556 | *Denitrobacterium* | 0.00 | 342 | *Bacteriovorax* | 0.00 | 161 | *Calditerrivibrio* |
| 0.02 | 1555 | *Slackia* | 0.00 | 342 | *Thermovenabulum* | 0.00 | 161 | *Afifella* |
| 0.02 | 1515 | *Tannerella* | 0.00 | 342 | *Candidatus Desulfofervidus* | 0.00 | 161 | *Paenisporosarcina* |
| 0.02 | 1515 | *Pyramidobacter* | 0.00 | 341 | *Dictyoglomus* | 0.00 | 161 | *Massilioclostridium* |
| 0.02 | 1511 | *Sunxiuqinia* | 0.00 | 341 | *Salinispira* | 0.00 | 160 | *Achlya* |
| 0.02 | 1507 | *Brevundimonas* | 0.00 | 340 | *Roseivivax* | 0.00 | 160 | *Arcanobacterium* |
| 0.02 | 1481 | *Coprobacillus* | 0.00 | 340 | *Jannaschia* | 0.00 | 160 | *Rhodovibrio* |
| 0.02 | 1477 | *Microvirgula* | 0.00 | 340 | *Thiohalospira* | 0.00 | 160 | *Turneriella* |
| 0.02 | 1472 | *Desulfatibacillum* | 0.00 | 339 | *Coxiella* | 0.00 | 159 | *Agaricus* |
| 0.02 | 1468 | *Sulfuricurvum* | 0.00 | 339 | *Tritrichomonas* | 0.00 | 159 | *Candidatus Arthromitus* |
| 0.02 | 1465 | *Photobacterium* | 0.00 | 339 | *Microcoleus* | 0.00 | 159 | *Halogeometricum* |
| 0.02 | 1461 | *Rhodanobacter* | 0.00 | 339 | *Frigoribacterium* | 0.00 | 159 | *Propionimicrobium* |
| 0.02 | 1457 | *Dyadobacter* | 0.00 | 339 | *Caloranaerobacter* | 0.00 | 159 | *Litoreibacter* |
| 0.02 | 1448 | *Chitinibacter* | 0.00 | 339 | *Fimbriimonas* | 0.00 | 159 | *Kandleria* |

| | | | | | | | | |
|---|---|---|---|---|---|---|---|---|
| 0.02 | 1442 | *Staphylococcus* | 0.00 | 338 | *Flexithrix* | 0.00 | 159 | *Sinomicrobium* |
| 0.02 | 1436 | *Thiothrix* | 0.00 | 338 | *Megamonas* | 0.00 | 158 | *Thermonema* |
| 0.02 | 1434 | *Acetobacterium* | 0.00 | 338 | *Serendipita* | 0.00 | 158 | *Geovibrio* |
| 0.02 | 1433 | *Brevibacillus* | 0.00 | 337 | *Gymnopus* | 0.00 | 158 | *Desulfotalea* |
| 0.02 | 1432 | *Roseomonas* | 0.00 | 337 | *Oceanibaculum* | 0.00 | 158 | *Thiohalomonas* |
| 0.02 | 1430 | *Spirosoma* | 0.00 | 337 | *Candida* | 0.00 | 158 | *Geminicoccus* |
| 0.02 | 1425 | *Fusobacterium* | 0.00 | 337 | *Arachidicoccus* | 0.00 | 158 | *Salisediminibacterium* |
| 0.02 | 1424 | *Rubrivivax* | 0.00 | 336 | *Defluviimonas* | 0.00 | 157 | *Spizellomyces* |
| 0.02 | 1419 | *Desulfobacca* | 0.00 | 334 | *Nannochloropsis* | 0.00 | 157 | *Palaeococcus* |
| 0.02 | 1417 | *Dehalococcoides* | 0.00 | 334 | *Sanguibacter* | 0.00 | 157 | *Halorhodospira* |
| 0.02 | 1406 | *Fusarium* | 0.00 | 334 | *Cryptosporangium* | 0.00 | 157 | *Pseudaminobacter* |
| 0.02 | 1404 | *Methylophilus* | 0.00 | 334 | *Hippea* | 0.00 | 157 | *Cylindrobasidium* |
| 0.02 | 1404 | *Desulfocarbo* | 0.00 | 334 | *Roseivirga* | 0.00 | 157 | *Halioglobus* |
| 0.02 | 1403 | *Thermococcus* | 0.00 | 334 | *Candidatus Jettenia* | 0.00 | 156 | *Limnothrix* |
| 0.02 | 1396 | *Butyricicoccus* | 0.00 | 334 | *Methylomarinum* | 0.00 | 156 | *Lobosporangium* |
| 0.02 | 1378 | *Phascolarctobacterium* | 0.00 | 332 | *Pseudorhodobacter* | 0.00 | 156 | *Tuberibacillus* |
| 0.02 | 1373 | *Candidatus Vecturithrix* | 0.00 | 332 | *Martelella* | 0.00 | 156 | *Rubritalea* |
| 0.02 | 1371 | *Magnetospirillum* | 0.00 | 332 | *Gracilibacter* | 0.00 | 156 | *Pseudobacteriovorax* |
| 0.02 | 1370 | *Sporobacter* | 0.00 | 332 | *Granulicella* | 0.00 | 155 | *Arhodomonas* |
| 0.02 | 1367 | *Geodermatophilus* | 0.00 | 331 | *Rugamonas* | 0.00 | 155 | *Bauldia* |
| 0.02 | 1363 | *Sphingopyxis* | 0.00 | 331 | *Fabibacter* | 0.00 | 155 | *Effusibacillus* |
| 0.02 | 1359 | *Ferriphaselus* | 0.00 | 330 | *Gillisia* | 0.00 | 154 | *Mycetocola* |
| 0.02 | 1358 | *Vogesella* | 0.00 | 329 | *Cellulophaga* | 0.00 | 154 | *Jeotgalibacillus* |
| 0.02 | 1347 | *Leptolyngbya* | 0.00 | 328 | *Scytonema* | 0.00 | 154 | *Salimicrobium* |
| 0.02 | 1346 | *Mycoplasma* | 0.00 | 328 | *Lechevalieria* | 0.00 | 153 | *Phaeodactylum* |
| 0.02 | 1335 | *Dialister* | 0.00 | 328 | *Sediminibacterium* | 0.00 | 153 | *Paracoccidioides* |
| 0.02 | 1330 | *Anaeromyxobacter* | 0.00 | 326 | *Rachicladosporium* | 0.00 | 153 | *Drechmeria* |
| 0.02 | 1324 | *Methylibium* | 0.00 | 326 | *Solitalea* | 0.00 | 153 | *Actinospica* |
| 0.02 | 1321 | *Desulfatitalea* | 0.00 | 325 | *Thermobifida* | 0.00 | 153 | *Methylovulum* |
| 0.02 | 1318 | *Escherichia* | 0.00 | 325 | *Candidatus Methylomirabilis* | 0.00 | 153 | *Marininema* |
| 0.02 | 1302 | *Moorella* | 0.00 | 324 | *Oceanospirillum* | 0.00 | 153 | *Ruthenibacterium* |
| 0.02 | 1301 | *Sterolibacterium* | 0.00 | 323 | *Haloferax* | 0.00 | 153 | *Ndongobacter* |
| 0.02 | 1301 | *Anaerocolumna* | 0.00 | 323 | *Hydrocarboniphaga* | 0.00 | 153 | *Phoenicibacter* |
| 0.02 | 1296 | *Sideroxydans* | 0.00 | 323 | *Sedimenticola* | 0.00 | 152 | *Lentinula* |
| 0.02 | 1284 | *Gemmatimonas* | 0.00 | 323 | *Sinomonas* | 0.00 | 152 | *Frondihabitans* |

| | | | | | | | | |
|---|---|---|---|---|---|---|---|---|
| 0.02 | 1279 | *Geotoga* | 0.00 | 322 | *Parasporobacterium* | 0.00 | 152 | *Lepidopterella* |
| 0.02 | 1273 | *Pseudogymnoascus* | 0.00 | 321 | *Chryseolinea* | 0.00 | 152 | *Rudaea* |
| 0.02 | 1269 | *Tyzzerella* | 0.00 | 320 | *Arthrobotrys* | 0.00 | 152 | *Oceanibacterium* |
| 0.02 | 1249 | *Methanofollis* | 0.00 | 319 | *Sporolactobacillus* | 0.00 | 152 | *Pseudarthrobacter* |
| 0.02 | 1245 | *Singulisphaera* | 0.00 | 319 | *Sphaerobolus* | 0.00 | 151 | *Coccidioides* |
| 0.02 | 1239 | *Exophiala* | 0.00 | 319 | *Cetobacterium* | 0.00 | 151 | *Salinivibrio* |
| 0.02 | 1232 | *Chlorobium* | 0.00 | 319 | *Auxenochlorella* | 0.00 | 151 | *Aliifodinibius* |
| 0.02 | 1228 | *Myxococcus* | 0.00 | 318 | *Mycena* | 0.00 | 151 | *Xanthophyllomyces* |
| 0.02 | 1221 | *Sediminispirochaeta* | 0.00 | 318 | *Methylosarcina* | 0.00 | 150 | *Pycnoporus* |
| 0.02 | 1219 | *Oxalobacter* | 0.00 | 318 | *Methylocapsa* | 0.00 | 150 | *Rhodoplanes* |
| 0.02 | 1215 | *Methylovorus* | 0.00 | 318 | *Candidatus Soleaferrea* | 0.00 | 150 | *Hebeloma* |
| 0.02 | 1204 | *Solimonas* | 0.00 | 318 | *Gemmatirosa* | 0.00 | 150 | *Thermobacillus* |
| 0.02 | 1202 | *Methanohalophilus* | 0.00 | 317 | *Nitrobacter* | 0.00 | 150 | *Serpula* |
| 0.02 | 1200 | *Streptacidiphilus* | 0.00 | 317 | *Williamsia* | 0.00 | 150 | *Oceanithermus* |
| 0.02 | 1192 | *Candidatus Marispirochaeta* | 0.00 | 317 | *Oligella* | 0.00 | 150 | *Thalassotalea* |
| 0.02 | 1191 | *Trypanosoma* | 0.00 | 317 | *Acetoanaerobium* | 0.00 | 149 | *Sporidiobolus* |
| 0.02 | 1187 | *Solobacterium* | 0.00 | 316 | *Acidocella* | 0.00 | 149 | *Verrucosispora* |
| 0.02 | 1180 | *Ignavibacterium* | 0.00 | 315 | *Xenophilus* | 0.00 | 149 | *Thermosinus* |
| 0.02 | 1171 | *Methanococcoides* | 0.00 | 315 | *Verminephrobacter* | 0.00 | 149 | *Pseudopedobacter* |
| 0.02 | 1158 | *Pantoea* | 0.00 | 315 | *Desulfovirgula* | 0.00 | 149 | *Rodentibacter* |
| 0.02 | 1154 | *Methanococcus* | 0.00 | 315 | *Mangrovimonas* | 0.00 | 148 | *Flectobacillus* |
| 0.02 | 1153 | *Sporanaerobacter* | 0.00 | 314 | *Chaetomium* | 0.00 | 148 | *Kineococcus* |
| 0.02 | 1150 | *Pseudoxanthomonas* | 0.00 | 313 | *Formosa* | 0.00 | 148 | *Malonomonas* |
| 0.02 | 1149 | *Thermanaerothrix* | 0.00 | 312 | *Glaciecola* | 0.00 | 148 | *Oceanicaulis* |
| 0.02 | 1144 | *Curtobacterium* | 0.00 | 312 | *Reichenbachiella* | 0.00 | 147 | *Weeksella* |
| 0.02 | 1143 | *Fretibacterium* | 0.00 | 312 | *Actinotalea* | 0.00 | 147 | *Pelistega* |
| 0.02 | 1140 | *Marinobacterium* | 0.00 | 312 | *Maledivibacter* | 0.00 | 147 | *Dactylellina* |
| 0.02 | 1133 | *Symbiodinium* | 0.00 | 312 | *Raineyella* | 0.00 | 147 | *Melaminivora* |
| 0.02 | 1127 | *Xanthobacter* | 0.00 | 311 | *Trichomonas* | 0.00 | 147 | *Rubeoparvulum* |
| 0.02 | 1123 | *Methylobacter* | 0.00 | 311 | *Marmoricola* | 0.00 | 146 | *Maricaulis* |
| 0.02 | 1121 | *Capnocytophaga* | 0.00 | 311 | *Candidimonas* | 0.00 | 146 | *Terracidiphilus* |
| 0.02 | 1117 | *Leifsonia* | 0.00 | 311 | *Anaerosalibacter* | 0.00 | 146 | *Methylobrevis* |
| 0.02 | 1115 | *Rhizobacter* | 0.00 | 310 | *Entamoeba* | 0.00 | 146 | *Merdibacter* |
| 0.02 | 1113 | *Pontibacter* | 0.00 | 310 | *Hypholoma* | 0.00 | 145 | *Brucella* |
| 0.02 | 1109 | *Klebsiella* | 0.00 | 310 | *Fodinicurvata* | 0.00 | 145 | *Rhynchosporium* |
| 0.02 | 1107 | *Paludibacterium* | 0.00 | 309 | *Pyrobaculum* | 0.00 | 145 | *Halobacteroides* |

| | | | | | | | | |
|---|---|---|---|---|---|---|---|---|
| 0.02 | 1107 | *Acidihalobacter* | 0.00 | 309 | *Corallococcus* | 0.00 | 145 | *Eutypa* |
| 0.02 | 1100 | *Phaeospirillum* | 0.00 | 309 | *Hylemonella* | 0.00 | 145 | *Haloterrigena* |
| 0.01 | 1092 | *Tessaracoccus* | 0.00 | 308 | *Persicobacter* | 0.00 | 145 | *Dacryopinax* |
| 0.01 | 1088 | *Halanaerobium* | 0.00 | 308 | *Leeuwenhoekiella* | 0.00 | 145 | *Magnaporthe* |
| 0.01 | 1086 | *Leeia* | 0.00 | 308 | *Candidatus Omnitrophus* | 0.00 | 145 | *Jeotgalibaca* |
| 0.01 | 1085 | *Prolixibacter* | 0.00 | 307 | *Tsukamurella* | 0.00 | 144 | *Chondrus* |
| 0.01 | 1083 | *Cohnella* | 0.00 | 307 | *Pontibacillus* | 0.00 | 144 | *Schizosaccharomyces* |
| 0.01 | 1073 | *Caldithrix* | 0.00 | 306 | *Parvularcula* | 0.00 | 144 | *Histoplasma* |
| 0.01 | 1073 | *Draconibacterium* | 0.00 | 306 | *Dethiosulfatibacter* | 0.00 | 144 | *Leucoagaricus* |
| 0.01 | 1071 | *Dictyostelium* | 0.00 | 305 | *Alcaligenes* | 0.00 | 144 | *Gloeocapsa* |
| 0.01 | 1070 | *Fibrobacter* | 0.00 | 305 | *Catenaria* | 0.00 | 144 | *Zunongwangia* |
| 0.01 | 1063 | *Fervidobacterium* | 0.00 | 305 | *Anoxybacillus* | 0.00 | 144 | *Fervidicola* |
| 0.01 | 1061 | *Thermosipho* | 0.00 | 304 | *Vitreoscilla* | 0.00 | 144 | *Tepidibacillus* |
| 0.01 | 1055 | *Desulfonatronum* | 0.00 | 304 | *Orenia* | 0.00 | 144 | *Candidatus Altiarchaeum* |
| 0.01 | 1055 | *Alkaliphilus* | 0.00 | 304 | *Hyalangium* | 0.00 | 143 | *Thielavia* |
| 0.01 | 1053 | *Petrotoga* | 0.00 | 304 | *Provencibacterium* | 0.00 | 143 | *Neolentinus* |
| 0.01 | 1049 | *Cryptobacterium* | 0.00 | 303 | *Cytophaga* | 0.00 | 143 | *Promicromonospora* |
| 0.01 | 1047 | *Gordonibacter* | 0.00 | 302 | *Beijerinckia* | 0.00 | 143 | *Yangia* |
| 0.01 | 1046 | *Pedosphaera* | 0.00 | 302 | *Micrococcus* | 0.00 | 143 | *Elioraea* |
| 0.01 | 1038 | *Peptoniphilus* | 0.00 | 302 | *Sporisorium* | 0.00 | 143 | *Glonium* |
| 0.01 | 1038 | *Aureimonas* | 0.00 | 302 | *Lentibacillus* | 0.00 | 143 | *Nafulsella* |
| 0.01 | 1037 | *Thiorhodococcus* | 0.00 | 302 | *Gramella* | 0.00 | 142 | *Kluyvera* |
| 0.01 | 1032 | *Lysinibacillus* | 0.00 | 301 | *Vulgatibacter* | 0.00 | 142 | *Zobellia* |
| 0.01 | 1029 | *Erwinia* | 0.00 | 301 | *Desnuesiella* | 0.00 | 142 | *Tateyamaria* |
| 0.01 | 1025 | *Planctomyces* | 0.00 | 299 | *Anthracocystis* | 0.00 | 142 | *Chelativorans* |
| 0.01 | 1025 | *Blastopirellula* | 0.00 | 298 | *Prosthecochloris* | 0.00 | 142 | *Niveispirillum* |
| 0.01 | 1023 | *Caballeronia* | 0.00 | 298 | *Acidisphaera* | 0.00 | 141 | *Haloarcula* |
| 0.01 | 1015 | *Serratia* | 0.00 | 298 | *Monosiga* | 0.00 | 141 | *Acidianus* |
| 0.01 | 1015 | *Archaeoglobus* | 0.00 | 297 | *Anaerobranca* | 0.00 | 141 | *Acetitomaculum* |
| 0.01 | 1015 | *Desulfosarcina* | 0.00 | 297 | *Chloroherpeton* | 0.00 | 141 | *Bathycoccus* |
| 0.01 | 1013 | *Odoribacter* | 0.00 | 296 | *Citrobacter* | 0.00 | 141 | *Bryocella* |
| 0.01 | 1012 | *Ardenticatena* | 0.00 | 296 | *Agrococcus* | 0.00 | 141 | *Cecembia* |
| 0.01 | 1008 | *Butyricimonas* | 0.00 | 296 | *Moesziomyces* | 0.00 | 141 | *Chitinivibrio* |
| 0.01 | 1007 | *Thermus* | 0.00 | 296 | *Alkalibacterium* | 0.00 | 141 | *Thermorudis* |
| 0.01 | 1006 | *Methylocaldum* | 0.00 | 295 | *Halothermothrix* | 0.00 | 140 | *Melanopsichium* |
| 0.01 | 1004 | *Oribacterium* | 0.00 | 295 | *Stachybotrys* | 0.00 | 140 | *Magnetococcus* |

| | | | | | | | | |
|---|---|---|---|---|---|---|---|---|
| 0.01 | 1003 | *Chloroflexus* | 0.00 | 295 | *Anaeromusa* | 0.00 | 140 | *Garciella* |
| 0.01 | 1000 | *Gallionella* | 0.00 | 295 | *Sulfurifustis* | 0.00 | 140 | *Henriciella* |
| 0.01 | 995 | *Mariniphaga* | 0.00 | 294 | *Scedosporium* | 0.00 | 139 | *Thermofilum* |
| 0.01 | 994 | *Halothiobacillus* | 0.00 | 292 | *Nitrincola* | 0.00 | 139 | *Tolypocladium* |
| 0.01 | 988 | *Propionibacterium* | 0.00 | 291 | *Candidatus Nitrosoglobus* | 0.00 | 139 | *Soonwooa* |
| 0.01 | 974 | *Pelosinus* | 0.00 | 290 | *Marichromatium* | 0.00 | 139 | *Sanghuangporus* |
| 0.01 | 972 | *Thermodesulfovibrio* | 0.00 | 290 | *Methanothermococcus* | 0.00 | 139 | *Nannizzia* |
| 0.01 | 971 | *Toxoplasma* | 0.00 | 290 | *Niameybacter* | 0.00 | 139 | *Lacunisphaera* |
| 0.01 | 970 | *Chitinilyticum* | 0.00 | 289 | *Spiroplasma* | 0.00 | 138 | *Saprospira* |
| 0.01 | 965 | *Dethiosulfatarculus* | 0.00 | 289 | *Pisolithus* | 0.00 | 138 | *Gregarina* |
| 0.01 | 957 | *Ottowia* | 0.00 | 289 | *Thermocrinis* | 0.00 | 138 | *Thalassomonas* |
| 0.01 | 952 | *Enterobacter* | 0.00 | 288 | *Tulasnella* | 0.00 | 138 | *Candidatus Marithrix* |
| 0.01 | 952 | *Caloramator* | 0.00 | 288 | *Oscillochloris* | 0.00 | 138 | *Pseudohongiella* |
| 0.01 | 952 | *Microbulbifer* | 0.00 | 287 | *Xylella* | 0.00 | 138 | *Faecalibaculum* |
| 0.01 | 950 | *Kocuria* | 0.00 | 287 | *Nitrospina* | 0.00 | 138 | *Pseudomassariella* |
| 0.01 | 949 | *Microvirga* | 0.00 | 287 | *Alkalilimnicola* | 0.00 | 137 | *Wolinella* |
| 0.01 | 948 | *Leishmania* | 0.00 | 287 | *Tistrella* | 0.00 | 137 | *Sebaldella* |
| 0.01 | 948 | *Methanocaldococcus* | 0.00 | 287 | *Lentisphaera* | 0.00 | 137 | *Blattabacterium* |
| 0.01 | 942 | *Sinorhizobium* | 0.00 | 287 | *Methyloglobulus* | 0.00 | 137 | *Gulosibacter* |
| 0.01 | 939 | *Proteiniphilum* | 0.00 | 287 | *Magnetofaba* | 0.00 | 137 | *Roseisalinus* |
| 0.01 | 938 | *Alcanivorax* | 0.00 | 286 | *Dactylosporangium* | 0.00 | 137 | *Mobilicoccus* |
| 0.01 | 935 | *Laribacter* | 0.00 | 285 | *Sarcina* | 0.00 | 137 | *Kallipyga* |
| 0.01 | 934 | *Rhodotorula* | 0.00 | 285 | *Coccomyxa* | 0.00 | 136 | *Penicilliopsis* |
| 0.01 | 929 | *Desulfatirhabdium* | 0.00 | 285 | *Aciduliprofundum* | 0.00 | 136 | *Labrys* |
| 0.01 | 927 | *Sulfurospirillum* | 0.00 | 285 | *Chitinispirillum* | 0.00 | 136 | *Amphritea* |
| 0.01 | 920 | *Pseudothermotoga* | 0.00 | 284 | *Mitsuokella* | 0.00 | 136 | *Vermiphilus* |
| 0.01 | 916 | *Roseiflexus* | 0.00 | 284 | *Echinicola* | 0.00 | 135 | *Acidimicrobium* |
| 0.01 | 912 | *Macellibacteroides* | 0.00 | 284 | *Thermoflexibacter* | 0.00 | 135 | *Filomicrobium* |
| 0.01 | 911 | *Saccharothrix* | 0.00 | 283 | *Pseudoramibacter* | 0.00 | 135 | *Ornithinimicrobium* |
| 0.01 | 910 | *Azohydromonas* | 0.00 | 283 | *Albimonas* | 0.00 | 135 | *Finegoldia* |
| 0.01 | 908 | *Desulfomicrobium* | 0.00 | 282 | *Actinopolyspora* | 0.00 | 135 | *Pseudozobellia* |
| 0.01 | 904 | *Acetanaerobacterium* | 0.00 | 282 | *Fibularhizoctonia* | 0.00 | 135 | *Salisaeta* |
| 0.01 | 901 | *Cellvibrio* | 0.00 | 281 | *Nitratifractor* | 0.00 | 135 | *Fibrisoma* |
| 0.01 | 899 | *Dehalogenimonas* | 0.00 | 281 | *Salinicola* | 0.00 | 134 | *Mobiluncus* |
| 0.01 | 896 | *Aquimarina* | 0.00 | 280 | *Photorhabdus* | 0.00 | 134 | *Termitomyces* |
| 0.01 | 894 | *Pelomonas* | 0.00 | 280 | *Ostreococcus* | 0.00 | 134 | *Anaeroglobus* |

| | | | | | | | | |
|---|---|---|---|---|---|---|---|---|
| 0.01 | 893 | *Gulbenkiania* | 0.00 | 280 | *Steroidobacter* | 0.00 | 134 | *Hydrogenivirga* |
| 0.01 | 888 | *Candidatus Brocadia* | 0.00 | 280 | *Anaerosporobacter* | 0.00 | 134 | *Nitrosopumilus* |
| 0.01 | 881 | *Methanolacinia* | 0.00 | 279 | *Stereum* | 0.00 | 133 | *Piscirickettsia* |
| 0.01 | 875 | *Chlamydia* | 0.00 | 278 | *Tetrahymena* | 0.00 | 133 | *Couchioplanes* |
| 0.01 | 875 | *Methanosphaerula* | 0.00 | 278 | *Fulvivirga* | 0.00 | 133 | *Alishewanella* |
| 0.01 | 870 | *Desulfarculus* | 0.00 | 277 | *Xylophilus* | 0.00 | 133 | *Crenothrix* |
| 0.01 | 864 | *Acholeplasma* | 0.00 | 275 | *Caenimonas* | 0.00 | 133 | *Ectothiorhodosinus* |
| 0.01 | 862 | *Dorea* | 0.00 | 275 | *Komagataeibacter* | 0.00 | 133 | *Aestuariibacter* |
| 0.01 | 854 | *Geothrix* | 0.00 | 274 | *Moritella* | 0.00 | 133 | *Gracilimonas* |
| 0.01 | 854 | *Hungatella* | 0.00 | 274 | *Candidatus Pelagibacter* | 0.00 | 133 | *Candidatus Saccharimonas* |
| 0.01 | 851 | *Candidatus Solibacter* | 0.00 | 274 | *Pleomorphomonas* | 0.00 | 133 | *Bernardetia* |
| 0.01 | 845 | *Erythrobacter* | 0.00 | 274 | *Dethiobacter* | 0.00 | 132 | *Hirsutella* |
| 0.01 | 844 | *Salinispora* | 0.00 | 274 | *Thermogemmatispora* | 0.00 | 132 | *Aliiglaciecola* |
| 0.01 | 844 | *Desulfatiglans* | 0.00 | 273 | *Valsa* | 0.00 | 131 | *Leptosphaeria* |
| 0.01 | 843 | *Beggiatoa* | 0.00 | 273 | *Pseudooceanicola* | 0.00 | 131 | *Pyrenophora* |
| 0.01 | 840 | *Coprococcus* | 0.00 | 272 | *Theileria* | 0.00 | 131 | *Claviceps* |
| 0.01 | 835 | *Melioribacter* | 0.00 | 272 | *Thermobaculum* | 0.00 | 131 | *Nitrococcus* |
| 0.01 | 833 | *Thermosyntropha* | 0.00 | 271 | *Ammonifex* | 0.00 | 131 | *Haloplasma* |
| 0.01 | 833 | *Formivibrio* | 0.00 | 271 | *Coprinopsis* | 0.00 | 131 | *Mariniradius* |
| 0.01 | 833 | *Ktedonobacter* | 0.00 | 271 | *Geoglobus* | 0.00 | 131 | *Moheibacter* |
| 0.01 | 832 | *Phytophthora* | 0.00 | 271 | *Leisingera* | 0.00 | 130 | *Byssochlamys* |
| 0.01 | 831 | *Veillonella* | 0.00 | 271 | *Coraliomargarita* | 0.00 | 130 | *Phanerochaete* |
| 0.01 | 831 | *Thalassiosira* | 0.00 | 271 | *Candidatus Symbiobacter* | 0.00 | 130 | *Planobispora* |
| 0.01 | 825 | *Candidatus Syntrophoarchaeum* | 0.00 | 271 | *Mesoaciditoga* | 0.00 | 130 | *Borreliella* |
| 0.01 | 824 | *Roseovarius* | 0.00 | 271 | *Pyrinomonas* | 0.00 | 130 | *Glarea* |
| 0.01 | 821 | *Sutterella* | 0.00 | 270 | *Nannocystis* | 0.00 | 130 | *Escovopsis* |
| 0.01 | 817 | *Alteromonas* | 0.00 | 270 | *Thermomonas* | 0.00 | 130 | *Eisenibacter* |
| 0.01 | 810 | *Emiliania* | 0.00 | 270 | *Moniliophthora* | 0.00 | 129 | *Halomicrobium* |
| 0.01 | 808 | *Gemmata* | 0.00 | 269 | *Lyngbya* | 0.00 | 129 | *Marinimicrobium* |
| 0.01 | 807 | *Propionicicella* | 0.00 | 269 | *Fragilariopsis* | 0.00 | 129 | *Elstera* |
| 0.01 | 803 | *Loktanella* | 0.00 | 269 | *Parafilimonas* | 0.00 | 128 | *Pasteurella* |
| 0.01 | 802 | *Holdemania* | 0.00 | 268 | *Raoultella* | 0.00 | 128 | *Perkinsus* |
| 0.01 | 800 | *Aromatoleum* | 0.00 | 268 | *Leadbetterella* | 0.00 | 128 | *Lipomyces* |
| 0.01 | 793 | *Aneurinibacillus* | 0.00 | 268 | *Lactonifactor* | 0.00 | 128 | *Streptobacillus* |

| | | | | | | | | |
|---|---|---|---|---|---|---|---|---|
| 0.01 | 793 | *Ferrovum* | 0.00 | 267 | *Galerina* | 0.00 | 128 | *Scleroderma* |
| 0.01 | 793 | *Kwoniella* | 0.00 | 267 | *Endocarpon* | 0.00 | 128 | *Roseibacterium* |
| 0.01 | 790 | *Jonquetella* | 0.00 | 267 | *Siphonobacter* | 0.00 | 128 | *Candidatus Amoebophilus* |
| 0.01 | 790 | *Labrenzia* | 0.00 | 267 | *Phormidesmis* | 0.00 | 128 | *Haematobacter* |
| 0.01 | 787 | *Puccinia* | 0.00 | 266 | *Spirulina* | 0.00 | 128 | *Mastigocoleus* |
| 0.01 | 786 | *Epulopiscium* | 0.00 | 266 | *Winogradskyella* | 0.00 | 128 | *Candidatus Berkiella* |
| 0.01 | 784 | *Holophaga* | 0.00 | 266 | *Kushneria* | 0.00 | 127 | *Lawsonia* |
| 0.01 | 783 | *Methylomicrobium* | 0.00 | 265 | *Roseobacter* | 0.00 | 127 | *Rosellinia* |
| 0.01 | 774 | *Thalassospira* | 0.00 | 265 | *Hortaea* | 0.00 | 127 | *Selenihalanaerobacter* |
| 0.01 | 774 | *Amantichitinum* | 0.00 | 265 | *Spongiibacter* | 0.00 | 127 | *Mesonia* |
| 0.01 | 771 | *Sporothrix* | 0.00 | 264 | *Hypoxylon* | 0.00 | 126 | *Ureaplasma* |
| 0.01 | 771 | *Niabella* | 0.00 | 264 | *Denitrovibrio* | 0.00 | 126 | *Sphaerotilus* |
| 0.01 | 768 | *Paraprevotella* | 0.00 | 264 | *Plicaturopsis* | 0.00 | 126 | *Oidiodendron* |
| 0.01 | 767 | *Jeongeupia* | 0.00 | 264 | *Varibaculum* | 0.00 | 126 | *Haliea* |
| 0.01 | 763 | *Desulfitibacter* | 0.00 | 263 | *Paucibacter* | 0.00 | 126 | *Monashia* |
| 0.01 | 761 | *Candidatus Scalindua* | 0.00 | 263 | *Reyranella* | 0.00 | 126 | *Oblitimonas* |
| 0.01 | 759 | *Marinomonas* | 0.00 | 263 | *Rubinisphaera* | 0.00 | 125 | *Helcococcus* |
| 0.01 | 758 | *Cellulosilyticum* | 0.00 | 261 | *Erysipelothrix* | 0.00 | 125 | *Botrytis* |
| 0.01 | 752 | *Thermithiobacillus* | 0.00 | 261 | *Sphaerobacter* | 0.00 | 125 | *Halonatronum* |
| 0.01 | 752 | *Celeribacter* | 0.00 | 261 | *Candidatus Kapabacteria* | 0.00 | 125 | *Vulcanibacillus* |
| 0.01 | 745 | *Anaerophaga* | 0.00 | 260 | *Haloferula* | 0.00 | 125 | *Ruania* |
| 0.01 | 743 | *Flavonifractor* | 0.00 | 259 | *Halococcus* | 0.00 | 125 | *Krasilnikoviella* |
| 0.01 | 742 | *Nitrosovibrio* | 0.00 | 259 | *Nitrolancea* | 0.00 | 124 | *Fistulina* |
| 0.01 | 740 | *Brachyspira* | 0.00 | 258 | *Paraphaeosphaeria* | 0.00 | 124 | *Thielaviopsis* |
| 0.01 | 738 | *Acidipropionibacterium* | 0.00 | 258 | *Citreicella* | 0.00 | 124 | *Kazachstania* |
| 0.01 | 737 | *Acetivibrio* | 0.00 | 257 | *Synechocystis* | 0.00 | 124 | *Xylona* |
| 0.01 | 734 | *Candidatus Moduliflexus* | 0.00 | 257 | *Methylohalobius* | 0.00 | 124 | *Kalmanozyma* |
| 0.01 | 733 | *Talaromyces* | 0.00 | 257 | *Fibrella* | 0.00 | 124 | *Parageobacillus* |
| 0.01 | 733 | *Thermodesulforhabdus* | 0.00 | 257 | *Mailhella* | 0.00 | 123 | *Streptoalloteichus* |
| 0.01 | 728 | *Clostridioides* | 0.00 | 256 | *Diaporthe* | 0.00 | 123 | *Natronobacterium* |
| 0.01 | 727 | *Chloracidobacterium* | 0.00 | 255 | *Francisella* | 0.00 | 123 | *Gardnerella* |
| 0.01 | 719 | *Dyella* | 0.00 | 255 | *Microscilla* | 0.00 | 123 | *Dichomitus* |
| 0.01 | 719 | *Parapedobacter* | 0.00 | 255 | *Thermoplasma* | 0.00 | 123 | *Segniliparus* |
| 0.01 | 718 | *Runella* | 0.00 | 255 | *Desulfocapsa* | 0.00 | 123 | *Caldimicrobium* |
| 0.01 | 718 | *Rhizoctonia* | 0.00 | 255 | *Turicibacter* | 0.00 | 123 | *Indibacter* |
| 0.01 | 715 | *Saccharomonospora* | 0.00 | 255 | *Gilvimarinus* | 0.00 | 122 | *Wolbachia* |

| | | | | | | | | |
|---|---|---|---|---|---|---|---|---|
| 0.01 | 713 | *Tolumonas* | 0.00 | 255 | *Dissulfuribacter* | 0.00 | 122 | *Flexistipes* |
| 0.01 | 712 | *Methylosinus* | 0.00 | 255 | *Acidibacillus* | 0.00 | 122 | *Sordaria* |
| 0.01 | 712 | *Libanicoccus* | 0.00 | 255 | *Halofilum* | 0.00 | 122 | *Suillus* |
| 0.01 | 711 | *Ramlibacter* | 0.00 | 254 | *Ancylobacter* | 0.00 | 122 | *Saitoella* |
| 0.01 | 710 | *Afipia* | 0.00 | 254 | *Rhizopus* | 0.00 | 122 | *Metallosphaera* |
| 0.01 | 709 | *Cyanothece* | 0.00 | 254 | *Neurospora* | 0.00 | 122 | *Xenococcus* |
| 0.01 | 705 | *Rhodomicrobium* | 0.00 | 254 | *Plesiocystis* | 0.00 | 122 | *Stanieria* |
| 0.01 | 705 | *Tangfeifania* | 0.00 | 254 | *Natranaerobius* | 0.00 | 122 | *Mucispirillum* |
| 0.01 | 697 | *Demequina* | 0.00 | 254 | *Adlercreutzia* | 0.00 | 122 | *Planifilum* |
| 0.01 | 691 | *Niastella* | 0.00 | 254 | *Defluviitalea* | 0.00 | 122 | *Candidatus Azobacteroides* |
| 0.01 | 690 | *Catabacter* | 0.00 | 254 | *Mizugakiibacter* | 0.00 | 121 | *Phycomyces* |
| 0.01 | 690 | *Pseudacidovorax* | 0.00 | 253 | *Leuconostoc* | 0.00 | 121 | *Kockovaella* |
| 0.01 | 684 | *Caldilinea* | 0.00 | 253 | *Alkalibacter* | 0.00 | 121 | *Laetiporus* |
| 0.01 | 681 | *Colwellia* | 0.00 | 253 | *Emticicia* | 0.00 | 121 | *Candidatus Phytoplasma* |
| 0.01 | 681 | *Ethanoligenens* | 0.00 | 253 | *Alloactinosynnema* | 0.00 | 121 | *Oceanisphaera* |
| 0.01 | 679 | *Syntrophothermus* | 0.00 | 253 | *Desertifilum* | 0.00 | 121 | *Ulvibacter* |
| 0.01 | 677 | *Neospora* | 0.00 | 252 | *Acytostelium* | 0.00 | 121 | *Humibacillus* |
| 0.01 | 676 | *Ectothiorhodospira* | 0.00 | 252 | *Vulcanisaeta* | 0.00 | 121 | *Fibroporia* |
| 0.01 | 676 | *Microbispora* | 0.00 | 251 | *Aerococcus* | 0.00 | 121 | *Cyberlindnera* |
| 0.01 | 676 | *Nitratireductor* | 0.00 | 251 | *Halobacterium* | 0.00 | 121 | *Trueperella* |
| 0.01 | 676 | *Alloprevotella* | 0.00 | 251 | *Trichophyton* | 0.00 | 121 | *Allokutzneria* |
| 0.01 | 674 | *Verrucomicrobium* | 0.00 | 250 | *Blastochloris* | 0.00 | 121 | *Thiolapillus* |
| 0.01 | 670 | *Cyclobacterium* | 0.00 | 250 | *Oceanicola* | 0.00 | 121 | *Candidatus Chrysopegis* |
| 0.01 | 669 | *Gonium* | 0.00 | 250 | *Luteipulveratus* | 0.00 | 120 | *Proteus* |
| 0.01 | 668 | *Salinivirga* | 0.00 | 250 | *Marivirga* | 0.00 | 120 | *Ascochyta* |
| 0.01 | 667 | *Listeria* | 0.00 | 250 | *Erysipelatoclostridium* | 0.00 | 120 | *Dermacoccus* |
| 0.01 | 666 | *Desulfobacter* | 0.00 | 249 | *Kibdelosporangium* | 0.00 | 120 | *Maritimibacter* |
| 0.01 | 665 | *Thiocapsa* | 0.00 | 249 | *Coniochaeta* | 0.00 | 120 | *Elusimicrobium* |
| 0.01 | 664 | *Agromyces* | 0.00 | 249 | *Tumebacillus* | 0.00 | 120 | *Cohaesibacter* |
| 0.01 | 663 | *Ensifer* | 0.00 | 249 | *Candidatus Desulforudis* | 0.00 | 120 | *Tersicoccus* |
| 0.01 | 660 | *Syntrophaceticus* | 0.00 | 249 | *Halarsenatibacter* | 0.00 | 120 | *Pricia* |
| 0.01 | 657 | *Arenimonas* | 0.00 | 248 | *Clavibacter* | 0.00 | 120 | *Sulfuricaulis* |
| 0.01 | 656 | *Allomyces* | 0.00 | 248 | *Pseudoruegeria* | 0.00 | 120 | *Candidatus Kryptobacter* |
| 0.01 | 655 | *Trichoderma* | 0.00 | 248 | *Stenoxybacter* | 0.00 | 119 | *Dothistroma* |
| 0.01 | 652 | *Cystobacter* | 0.00 | 248 | *Rubripirellula* | 0.00 | 119 | *Paenalcaligenes* |

| | | | | | | | | |
|---|---|---|---|---|---|---|---|---|
| 0.01 | 650 | *Moraxella* | 0.00 | 247 | *Methylacidiphilum* | 0.00 | 119 | *Lihuaxuella* |
| 0.01 | 650 | *Monoraphidium* | 0.00 | 247 | *Zymoseptoria* | 0.00 | 119 | *Rhodohalobacter* |
| 0.01 | 650 | *Janibacter* | 0.00 | 247 | *Millionella* | 0.00 | 118 | *Melampsora* |
| 0.01 | 650 | *Ruegeria* | 0.00 | 245 | *Thiomargarita* | 0.00 | 118 | *Tuber* |
| 0.01 | 648 | *Tilletia* | 0.00 | 245 | *Halodesulfovibrio* | 0.00 | 118 | *Natronomonas* |
| 0.01 | 647 | *Labilibacter* | 0.00 | 244 | *Piloderma* | 0.00 | 118 | *Beutenbergia* |
| 0.01 | 645 | *Blastococcus* | 0.00 | 244 | *Desulfoplanes* | 0.00 | 118 | *Epicoccum* |
| 0.01 | 645 | *Oxobacter* | 0.00 | 243 | *Desulfurobacterium* | 0.00 | 118 | *Thermobrachium* |
| 0.01 | 644 | *Chondromyces* | 0.00 | 243 | *Mogibacterium* | 0.00 | 118 | *Komagataella* |
| 0.01 | 644 | *Acidiferrobacter* | 0.00 | 243 | *Lokiarchaeum* | 0.00 | 118 | *Thalassobaculum* |
| 0.01 | 643 | *Kaistia* | 0.00 | 242 | *Saccharomyces* | 0.00 | 118 | *Alteribacillus* |
| 0.01 | 643 | *Sulfurimonas* | 0.00 | 242 | *Cryptosporidium* | 0.00 | 117 | *Halopiger* |
| 0.01 | 641 | *Volvox* | 0.00 | 242 | *Achromatium* | 0.00 | 117 | *Beduini* |
| 0.01 | 640 | *Exiguobacterium* | 0.00 | 242 | *Marinospirillum* | 0.00 | 116 | *Piromyces* |
| 0.01 | 640 | *Cellulosimicrobium* | 0.00 | 242 | *Balneatrix* | 0.00 | 116 | *Syncephalastrum* |
| 0.01 | 639 | *Yersinia* | 0.00 | 242 | *Punctularia* | 0.00 | 116 | *Acidilobus* |
| 0.01 | 639 | *Propionispora* | 0.00 | 242 | *Pseudoclavibacter* | 0.00 | 116 | *Thermobispora* |
| 0.01 | 636 | *Tepidanaerobacter* | 0.00 | 242 | *Filimonas* | 0.00 | 116 | *Zooshikella* |
| 0.01 | 631 | *Tenacibaculum* | 0.00 | 242 | *Geminocystis* | 0.00 | 116 | *Kineosphaera* |
| 0.01 | 630 | *Lewinella* | 0.00 | 241 | *Oleiphilus* | 0.00 | 116 | *Hydrogenimonas* |
| 0.01 | 630 | *Solirubrobacter* | 0.00 | 241 | *Tepidibacter* | 0.00 | 116 | *Cyphellophora* |
| 0.01 | 626 | *Leptonema* | 0.00 | 241 | *Candidatus Stoquefichus* | 0.00 | 116 | *Estrella* |
| 0.01 | 625 | *Jiangella* | 0.00 | 240 | *Phialophora* | 0.00 | 116 | *Wickerhamomyces* |
| 0.01 | 624 | *Aureococcus* | 0.00 | 240 | *Sulfurihydrogenibium* | 0.00 | 116 | *Sporolituus* |
| 0.01 | 623 | *Geosporobacter* | 0.00 | 240 | *Catenulispora* | 0.00 | 116 | *Halorientalis* |
| 0.01 | 622 | *Stigmatella* | 0.00 | 239 | *Intrasporangium* | 0.00 | 116 | *Pseudorhodoplanes* |
| 0.01 | 621 | *Rheinheimera* | 0.00 | 239 | *Lichtheimia* | 0.00 | 116 | *Criibacterium* |
| 0.01 | 618 | *Saccharopolyspora* | 0.00 | 238 | *Owenweeksia* | 0.00 | 115 | *Thermocrispum* |
| 0.01 | 618 | *Desulfofustis* | 0.00 | 238 | *Crenotalea* | 0.00 | 115 | *Cyanidioschyzon* |
| 0.01 | 618 | *Desulfotignum* | 0.00 | 238 | *Romboutsia* | 0.00 | 115 | *Choanephora* |
| 0.01 | 616 | *Ochrobactrum* | 0.00 | 237 | *Microdochium* | 0.00 | 115 | *Halorhabdus* |
| 0.01 | 614 | *Flavihumibacter* | 0.00 | 237 | *Thioflavicoccus* | 0.00 | 114 | *Peptococcus* |
| 0.01 | 613 | *Methylophaga* | 0.00 | 237 | *Thermosulfurimonas* | 0.00 | 114 | *Plasmodiophora* |
| 0.01 | 611 | *Aquabacterium* | 0.00 | 237 | *Nitrospirillum* | 0.00 | 114 | *Salinibacter* |
| 0.01 | 610 | *Sulfitobacter* | 0.00 | 237 | *Cnuella* | 0.00 | 114 | *Granulibacter* |
| 0.01 | 610 | *Pseudorhodoferax* | 0.00 | 236 | *Pediococcus* | 0.00 | 114 | *Hydnomerulius* |

| | | | | | | | | |
|---|---|---|---|---|---|---|---|---|
| 0.01 | 610 | *Anaeromassilibacillus* | 0.00 | 236 | *Neptunomonas* | 0.00 | 114 | *Actinomycetospora* |
| 0.01 | 609 | *Endozoicomonas* | 0.00 | 236 | *Thalassobius* | 0.00 | 114 | *Rasamsonia* |
| 0.01 | 608 | *Actinokineospora* | 0.00 | 236 | *Pseudoxanthobacter* | 0.00 | 114 | *Psychrobacillus* |
| 0.01 | 608 | *Maribacter* | 0.00 | 236 | *Halovenus* | 0.00 | 114 | *Motiliproteus* |
| 0.01 | 608 | *Candidatus Magnetomorum* | 0.00 | 236 | *Candidatus Kryptonium* | 0.00 | 114 | *Colibacter* |
| 0.01 | 607 | *Meiothermus* | 0.00 | 235 | *Acidobacterium* | 0.00 | 113 | *Cardiobacterium* |
| 0.01 | 607 | *Oceanobacillus* | 0.00 | 235 | *Candidatus Glomeribacter* | 0.00 | 113 | *Tetragenococcus* |
| 0.01 | 606 | *Rhodovulum* | 0.00 | 235 | *Anaerofustis* | 0.00 | 113 | *Thermosynechococcus* |
| 0.01 | 606 | *Belnapia* | 0.00 | 235 | *Thermosulfidibacter* | 0.00 | 113 | *Planomicrobium* |
| 0.01 | 604 | *Aurantimonas* | 0.00 | 234 | *Planomonospora* | 0.00 | 113 | *Psychrilyobacter* |
| 0.01 | 604 | *Defluviitoga* | 0.00 | 233 | *Tistlia* | 0.00 | 113 | *Pleomorphobacterium* |
| 0.01 | 603 | *Brevibacterium* | 0.00 | 233 | *Pseudarcicella* | 0.00 | 113 | *Naematelia* |
| 0.01 | 603 | *Noviherbaspirillum* | 0.00 | 232 | *Rhizopogon* | 0.00 | 112 | *Pyrodictium* |
| 0.01 | 601 | *Trametes* | 0.00 | 232 | *Naegleria* | 0.00 | 112 | *Eremothecium* |
| 0.01 | 599 | *Microtetraspora* | 0.00 | 232 | *Kyrpidia* | 0.00 | 112 | *Sporichthya* |
| 0.01 | 599 | *Anaerobium* | 0.00 | 231 | *Trichodesmium* | 0.00 | 112 | *Isaria* |
| 0.01 | 598 | *Chitiniphilus* | 0.00 | 231 | *Stappia* | 0.00 | 112 | *Succinimonas* |
| 0.01 | 596 | *Chlorobaculum* | 0.00 | 231 | *Inediibacterium* | 0.00 | 112 | *Maribius* |
| 0.01 | 595 | *Candidatus Magnetobacterium* | 0.00 | 230 | *Paramecium* | 0.00 | 112 | *Caldanaerovirga* |
| 0.01 | 595 | *Brachybacterium* | 0.00 | 230 | *Geitlerinema* | 0.00 | 112 | *Faecalicoccus* |
| 0.01 | 595 | *Methanomassiliicoccus* | 0.00 | 230 | *Desulfurivibrio* | 0.00 | 111 | *Ahrensia* |
| 0.01 | 595 | *Pseudobacteroides* | 0.00 | 229 | *Agreia* | 0.00 | 111 | *Moellerella* |
| 0.01 | 593 | *Limnohabitans* | 0.00 | 229 | *Kangiella* | 0.00 | 111 | *Conchiformibius* |
| 0.01 | 592 | *Rufibacter* | 0.00 | 229 | *Ohtaekwangia* | 0.00 | 111 | *Wenxinia* |
| 0.01 | 591 | *Micromonas* | 0.00 | 229 | *Massilimaliae* | 0.00 | 111 | *Spiribacter* |
| 0.01 | 588 | *Planococcus* | 0.00 | 228 | *Chelatococcus* | 0.00 | 111 | *Terrimicrobium* |
| 0.01 | 588 | *Thermodesulfatator* | 0.00 | 228 | *Aminobacter* | 0.00 | 111 | *Dubosiella* |
| 0.01 | 588 | *Enterorhabdus* | 0.00 | 228 | *Nesterenkonia* | 0.00 | 110 | *Pseudozyma* |
| 0.01 | 586 | *Hespellia* | 0.00 | 228 | *Parachlamydia* | 0.00 | 110 | *Fulvimarina* |
| 0.01 | 583 | *Thiomicrospira* | 0.00 | 228 | *Mariprofundus* | 0.00 | 110 | *Grimontia* |
| 0.01 | 583 | *Cryptococcus* | 0.00 | 228 | *Magnetospira* | 0.00 | 110 | *Torrubiella* |
| 0.01 | 583 | *Peptoclostridium* | 0.00 | 228 | *Paeniclostridium* | 0.00 | 110 | *Tropicibacter* |
| 0.01 | 582 | *Limnobacter* | 0.00 | 227 | *Babesia* | 0.00 | 110 | *Oceanicoccus* |
| 0.01 | 580 | *Geofilum* | 0.00 | 227 | *Cobetia* | 0.00 | 110 | *Candidatus Babela* |
| 0.01 | 579 | *Elizabethkingia* | 0.00 | 227 | *Dokdonella* | 0.00 | 109 | *Ornithobacterium* |

| | | | | | | | | |
|---|---|---|---|---|---|---|---|---|
| 0.01 | 578 | *Prosthecomicrobium* | 0.00 | 227 | *Clohesyomyces* | 0.00 | 109 | *Daldinia* |
| 0.01 | 577 | *Thermoanaerobacterium* | 0.00 | 227 | *Basilea* | 0.00 | 109 | *Streptomonospora* |
| 0.01 | 574 | *Youngiibacter* | 0.00 | 226 | *Acaryochloris* | 0.00 | 109 | *Lachnobacterium* |
| 0.01 | 573 | *Methanoplanus* | 0.00 | 226 | *Persephonella* | 0.00 | 109 | *Polymorphum* |
| 0.01 | 571 | *Sedimentibacter* | 0.00 | 226 | *Marvinbryantia* | 0.00 | 109 | *Murdochiella* |
| 0.01 | 570 | *Schlesneria* | 0.00 | 226 | *Desulfonatronospira* | 0.00 | 109 | *Parastagonospora* |
| 0.01 | 567 | *Opitutus* | 0.00 | 225 | *Smittium* | 0.00 | 108 | *Beauveria* |
| 0.01 | 563 | *Gluconobacter* | 0.00 | 225 | *Labilithrix* | 0.00 | 108 | *Sphaerulina* |
| 0.01 | 563 | *Patulibacter* | 0.00 | 224 | *Isoptericola* | 0.00 | 107 | *Plasmopara* |
| 0.01 | 561 | *Aeromicrobium* | 0.00 | 224 | *Ophiocordyceps* | 0.00 | 107 | *Leucothrix* |
| 0.01 | 561 | *Robinsoniella* | 0.00 | 223 | *Fischerella* | 0.00 | 107 | *Gelidibacter* |
| 0.01 | 561 | *Marinifilum* | 0.00 | 223 | *Zygosaccharomyces* | 0.00 | 107 | *Natrinema* |
| 0.01 | 560 | *Bdellovibrio* | 0.00 | 223 | *Salinimicrobium* | 0.00 | 107 | *Asaia* |
| 0.01 | 559 | *Thermoanaerobacter* | 0.00 | 223 | *Thermotalea* | 0.00 | 107 | *Halomicronema* |
| 0.01 | 559 | *Luteimonas* | 0.00 | 222 | *Lachnospira* | 0.00 | 107 | *Robiginitalea* |
| 0.01 | 558 | *Methanomethylovorans* | 0.00 | 222 | *Capronia* | 0.00 | 107 | *Viridibacillus* |
| 0.01 | 558 | *Dendrosporobacter* | 0.00 | 222 | *Nitratiruptor* | 0.00 | 107 | *Polycyclovorans* |
| 0.01 | 557 | *Halolamina* | 0.00 | 222 | *Cronobacter* | 0.00 | 107 | *Candidatus Acetothermum* |
| 0.01 | 555 | *Mitsuaria* | 0.00 | 222 | *Neorhizobium* | 0.00 | 106 | *Ichthyophthirius* |
| 0.01 | 554 | *Bilophila* | 0.00 | 221 | *Knoellia* | 0.00 | 106 | *Mixia* |
| 0.01 | 554 | *Emergencia* | 0.00 | 221 | *Sistotremastrum* | 0.00 | 106 | *Mannheimia* |
| 0.01 | 553 | *Actinopolymorpha* | 0.00 | 221 | *Belliella* | 0.00 | 106 | *Prauserella* |
| 0.01 | 550 | *Carnobacterium* | 0.00 | 221 | *Sciscionella* | 0.00 | 106 | *Thermovibrio* |
| 0.01 | 548 | *Insolitispirillum* | 0.00 | 221 | *Ilumatobacter* | 0.00 | 106 | *Caldimonas* |
| 0.01 | 546 | *Leptomonas* | 0.00 | 220 | *Kurthia* | 0.00 | 105 | *Linderina* |
| 0.01 | 545 | *Caldicellulosiruptor* | 0.00 | 220 | *Oxytricha* | 0.00 | 105 | *Pleurocapsa* |
| 0.01 | 545 | *Thermacetogenium* | 0.00 | 220 | *Stylonychia* | 0.00 | 105 | *Stagonospora* |
| 0.01 | 545 | *Thecamonas* | 0.00 | 220 | *Arsenicicoccus* | 0.00 | 105 | *Sodalis* |
| 0.01 | 544 | *Leptotrichia* | 0.00 | 220 | *Segetibacter* | 0.00 | 105 | *Parvimonas* |
| 0.01 | 542 | *Prochlorococcus* | 0.00 | 220 | *Purpureocillium* | 0.00 | 104 | *Roseospirillum* |
| 0.01 | 541 | *Sporosarcina* | 0.00 | 219 | *Rickettsia* | 0.00 | 104 | *Natronorubrum* |
| 0.01 | 538 | *Calothrix* | 0.00 | 219 | *Hypsizygus* | 0.00 | 104 | *Crinalium* |
| 0.01 | 538 | *Lysinimicrobium* | 0.00 | 219 | *Actinosynnema* | 0.00 | 104 | *Acidomyces* |
| 0.01 | 537 | *Chlamydomonas* | 0.00 | 219 | *Coprothermobacter* | 0.00 | 104 | *Halalkalicoccus* |
| 0.01 | 537 | *Deferrisoma* | 0.00 | 219 | *Thioalkalimicrobium* | 0.00 | 104 | *Millerozyma* |

| | | | | | | | | |
|---|---|---|---|---|---|---|---|---|
| 0.01 | 536 | *Thermodesulfobacterium* | 0.00 | 219 | *Pseudospirillum* | 0.00 | 104 | *Faecalicatena* |
| 0.01 | 535 | *Mucinivorans* | 0.00 | 219 | *Perlucidibaca* | 0.00 | 103 | *Pseudocohnilembus* |
| 0.01 | 533 | *Anaerorhabdus* | 0.00 | 219 | *Methyloprofundus* | 0.00 | 103 | *Rubrimonas* |
| 0.01 | 533 | *Pseudolabrys* | 0.00 | 218 | *Botryobasidium* | 0.00 | 103 | *Longispora* |
| 0.01 | 533 | *Angelakisella* | 0.00 | 218 | *Lacinutrix* | 0.00 | 103 | *Meganema* |
| 0.01 | 531 | *Anaerobacillus* | 0.00 | 218 | *Neptuniibacter* | 0.00 | 103 | *Cribrihabitans* |
| 0.01 | 530 | *Friedmanniella* | 0.00 | 218 | *Wohlfahrtiimonas* | 0.00 | 103 | *Paramesorhizobium* |
| 0.01 | 530 | *Hoeflea* | 0.00 | 217 | *Eikenella* | 0.00 | 103 | *Numidum* |
| 0.01 | 528 | *Tetrasphaera* | 0.00 | 217 | *Thermothelomyces* | 0.00 | 103 | *Immundisolibacter* |
| 0.01 | 528 | *Anaerococcus* | 0.00 | 216 | *Succinispira* | 0.00 | 102 | *Caryophanon* |
| 0.01 | 527 | *Archangium* | 0.00 | 216 | *Methylogaea* | 0.00 | 102 | *Jeotgalicoccus* |
| 0.01 | 526 | *Lactococcus* | 0.00 | 215 | *Ophiostoma* | 0.00 | 102 | *Saccharospirillum* |
| 0.01 | 525 | *Desulforegula* | 0.00 | 215 | *Desulfuromusa* | 0.00 | 102 | *Candidatus Protochlamydia* |
| 0.01 | 525 | *Desulfoluna* | 0.00 | 215 | *Anaeromyces* | 0.00 | 102 | *Flaviramulus* |
| 0.01 | 525 | *Caldisericum* | 0.00 | 215 | *Enhygromyxa* | 0.00 | 102 | *Marinactinospora* |
| 0.01 | 524 | *Subdoligranulum* | 0.00 | 215 | *Aureispira* | 0.00 | 101 | *Basidiobolus* |
| 0.01 | 522 | *Nostoc* | 0.00 | 215 | *Strigomonas* | 0.00 | 101 | *Johnsonella* |
| 0.01 | 522 | *Tepidimonas* | 0.00 | 215 | *Paraglaciecola* | 0.00 | 101 | *Oenococcus* |
| 0.01 | 521 | *Carboxydothermus* | 0.00 | 214 | *Thioploca* | 0.00 | 101 | *Tetrapisispora* |
| 0.01 | 519 | *Salmonella* | 0.00 | 214 | *Weissella* | 0.00 | 101 | *Quadrisphaera* |
| 0.01 | 519 | *Mucor* | 0.00 | 214 | *Natrialba* | 0.00 | 101 | *Rudanella* |
| 0.01 | 519 | *Aureobasidium* | 0.00 | 214 | *Pseudocercospora* | 0.00 | 101 | *Catelliglobosispora* |
| 0.01 | 519 | *Rathayibacter* | 0.00 | 213 | *Hassallia* | 0.00 | 101 | *Dongia* |
| 0.01 | 519 | *Thermaerobacter* | 0.00 | 213 | *Bariatricus* | 0.00 | 101 | *Candidatus Paracaedibacter* |
| 0.01 | 519 | *Phycicoccus* | 0.00 | 212 | *Sclerotinia* | 0.00 | 100 | *Actinotignum* |
| 0.01 | 518 | *Hyphomonas* | 0.00 | 212 | *Desulfurispora* | 0.00 | 100 | *Massiliomicrobiota* |
| 0.01 | 518 | *Isosphaera* | 0.00 | 212 | *Hydrotalea* | | | |

**Table S3.** Functional overview of initial community (top 500 rank only).

| Rank | Count | Abundance | ID | Organism | Protein Names |
|---|---|---|---|---|---|
| 1 | 4376 | 0.14 | F4BUN6 | *Methanosaeta concilii* | Transposase, IS116/IS110/IS902 family |
| 2 | 3604 | 0.12 | F4BZD0 | *Methanosaeta concilii* | Transposase, putative |
| 3 | 2697 | 0.09 | F4BY05 | *Methanosaeta concilii* | Transposase, IS116/IS110/IS902 family |
| 4 | 1997 | 0.07 | F4BY15 | *Methanosaeta concilii* | Transposase, IS116/IS110/IS902 family |
| 5 | 1995 | 0.07 | I9R3U4 | *Bacteroides cellulosilyticus* | Uncharacterized protein |
| 6 | 1977 | 0.07 | F4BSV9 | *Methanosaeta concilii* | Transposase, IS4 family |
| 7 | 1641 | 0.05 | F4BZ63 | *Methanosaeta concilii* | Transposase, IS4 family |
| 8 | 1506 | 0.05 | F4BSV1 | *Methanosaeta concilii* | Transposase, IS4 family, putative |
| 9 | 1289 | 0.04 | F4BV05 | *Methanosaeta concilii* | Transposase, Rhodopirellula family protein |
| 10 | 1247 | 0.04 | F4BZW8 | *Methanosaeta concilii* | Transposase, IS605 OrfB family |
| 11 | 1232 | 0.04 | F4BX44 | *Methanosaeta concilii* | Transposase, IS605 OrfB family |
| 12 | 1164 | 0.04 | N8ZVC9 | *Acinetobacter gerneri* | Uncharacterized protein |
| 13 | 1132 | 0.04 | F3ZQL8 | *Bacteroides coprosuis* | Transposase IS4 family protein |
| 14 | 1125 | 0.04 | F4BYN0 | *Methanosaeta concilii* | Transposase, IS605 OrfB family |
| 15 | 1124 | 0.04 | N8ZNX3 | *Acinetobacter gerneri* | Uncharacterized protein |
| 16 | 1122 | 0.04 | F4BXC1 | *Methanosaeta concilii* | Transposase, IS4 |
| 17 | 981 | 0.03 | F4BV83 | *Methanosaeta concilii* | Uncharacterized protein |
| 18 | 916 | 0.03 | F4BUD0 | *Methanosaeta concilii* | Conserved domain transposase IS605 family |
| 19 | 872 | 0.03 | F4BVK0 | *Methanosaeta concilii* | Transposase, IS116/IS110/IS902 family |
| 20 | 803 | 0.03 | | | |
| 21 | 796 | 0.03 | F4C073 | *Methanosaeta concilii* | Transposase, IS605 OrfB family |
| 22 | 780 | 0.03 | A0A0R3QEE4 | *Brugia timori* | Uncharacterized protein |
| 23 | 738 | 0.02 | F4BZ72 | *Methanosaeta concilii* | Uncharacterized protein |
| 24 | 729 | 0.02 | F4BTW8 | *Methanosaeta concilii* | Transposase IS3 family protein, putative integrase |
| 25 | 716 | 0.02 | F4BUB6 | *Methanosaeta concilii* | Transposase, IS605 OrfB family |
| 26 | 661 | 0.02 | F4BUL3 | *Methanosaeta concilii* | Transposase |
| 27 | 624 | 0.02 | H8MU17 | *Corallococcus coralloides* | Long-chain-fatty-acid--CoA ligase |
| 28 | 608 | 0.02 | | | |
| 29 | 607 | 0.02 | A0A077XLW5 | *Sphinx1.76-related DNA* | Replication protein |
| 30 | 592 | 0.02 | F4C0V0 | *Methanosaeta concilii* | Uncharacterized protein |
| 31 | 586 | 0.02 | F4BTV1 | *Methanosaeta concilii* | Glycosyltransferase family 36 protein |
| 32 | 578 | 0.02 | A0A1Q2CUJ4 | *Tessaracoccus flavescens* | ISL3 family transposase |
| 33 | 570 | 0.02 | F4BZ82 | *Methanosaeta concilii* | Cna B domain protein |

| | | | | | |
|---|---|---|---|---|---|
| 34 | 555 | 0.02 | F4BV12 | *Methanosaeta concilii* | Transposase, Mutator family |
| 35 | 549 | 0.02 | F4BTX0 | *Methanosaeta concilii* | Conserved domain protein |
| 36 | 536 | 0.02 | A0A0T5YH86 | *Acinetobacter baumannii* | Mobilization protein A |
| 37 | 534 | 0.02 | A0A1Q3QH52 | *Bacteroidia bacterium* | Uncharacterized protein |
| 38 | 521 | 0.02 | F4BW50 | *Methanosaeta concilii* | Transposase, IS605 OrfB family domain protein |
| 39 | 520 | 0.02 | F4BT92 | *Methanosaeta concilii* | Transposase, Mutator family |
| 40 | 513 | 0.02 | A0A077XNB6 | *Sphinx1.76-related DNA* | Replication protein |
| 41 | 505 | 0.02 | A0A1Q3QCL0 | *Bacteroidia bacterium* | Uncharacterized protein |
| 42 | 504 | 0.02 | G9FB24 | *uncultured bacterium* | IS21 family transposition protein |
| 43 | 501 | 0.02 | H8MYX7 | *Corallococcus coralloides* | Non-ribosomal peptide synthetase/polyketide synthase |
| 44 | 487 | 0.02 | | | |
| 45 | 483 | 0.02 | A0A1Q3QJR2 | *Bacteroidia bacterium* | Cell surface protein SprA |
| 46 | 482 | 0.02 | | | |
| 47 | 481 | 0.02 | | | |
| 48 | 478 | 0.02 | F4BW44 | *Methanosaeta concilii* | Transposase, IS605 OrfB family |
| 49 | 469 | 0.02 | A0A178FWP5 | *Acinetobacter sp. SFC* | Transposase |
| 50 | 466 | 0.02 | F4BVI0 | *Methanosaeta concilii* | Methyltransferase/glycosyl transferase fusion protein |
| 51 | 437 | 0.01 | F4BXD7 | *Methanosaeta concilii* | Transposase, IS4 (Transposase, IS4 family) |
| 52 | 426 | 0.01 | A0A1M7BMK5 | *Fibrobacter sp. UWEL* | Transposase InsO and inactivated derivatives |
| 53 | 423 | 0.01 | | | |
| 54 | 418 | 0.01 | | | |
| 55 | 417 | 0.01 | N8QHE7 | *Acinetobacter bohemicus* | Uncharacterized protein |
| 56 | 414 | 0.01 | A0A1Q3QC75 | *Bacteroidia bacterium* | Uncharacterized protein |
| 57 | 409 | 0.01 | | *Bacteroidia bacterium* | |
| 58 | 408 | 0.01 | | | |
| 59 | 405 | 0.01 | U6RUP9 | *Bacteroides sp.* | Uncharacterized protein |
| 60 | 405 | 0.01 | | | |
| 61 | 402 | 0.01 | | | |
| 62 | 401 | 0.01 | | | |
| 63 | 399 | 0.01 | F4BUA5 | *Methanosaeta concilii* | Transposase, IS605 OrfB family |
| 64 | 393 | 0.01 | | | |
| 65 | 384 | 0.01 | A0A1Q3QBX8 | *Bacteroidia bacterium* | Uncharacterized protein |
| 66 | 383 | 0.01 | F4BUD1 | *Methanosaeta concilii* | Transposase, putative |
| 67 | 380 | 0.01 | | | |
| 68 | 379 | 0.01 | A0A1Q3QGY6 | *Bacteroidia bacterium* | Hybrid sensor histidine kinase/response regulator |
| 69 | 374 | 0.01 | I2F7Y4 | *Mesotoga prima* | Retron-type reverse transcriptase |

| | | | | | |
|---|---|---|---|---|---|
| 70 | 370 | 0.01 | T5L8A5 | *Microbacterium maritypicum* | Uncharacterized protein |
| 71 | 361 | 0.01 | H8MN58 | *Corallococcus coralloides* | Long-chain-fatty-acid--CoA ligase |
| 72 | 356 | 0.01 | A0A1Q3QI02 | *Bacteroidia bacterium* | Uncharacterized protein |
| 73 | 355 | 0.01 | F4BXC2 | *Methanosaeta concilii* | Transposase IS200-like protein (Transposase, IS200-like, putative MITE) |
| 74 | 355 | 0.01 | F4BY39 | *Methanosaeta concilii* | Transposase, IS605 OrfB family |
| 75 | 351 | 0.01 | A0A0C5RZ24 | *Pseudomonas putida* | Adhesin |
| 76 | 351 | 0.01 | | | |
| 77 | 348 | 0.01 | F3ZQK5 | *Bacteroides coprosuis* | Transposase IS4 family protein |
| 78 | 348 | 0.01 | F4BYV3 | *Methanosaeta concilii* | Riboflavin biosynthesis protein RibD C-domain protein |
| 79 | 346 | 0.01 | | | |
| 80 | 345 | 0.01 | G9FB23 | *uncultured bacterium* | IS21 family transposition protein |
| 81 | 345 | 0.01 | | | |
| 82 | 342 | 0.01 | F4BZ39 | *Methanosaeta concilii* | Transposase, IS116/IS110/IS902 family |
| 83 | 340 | 0.01 | A0A1M6CDY9 | *Tessaracoccus bendigoensis* | Cd2+/Zn2+-exporting ATPase |
| 84 | 336 | 0.01 | F4BXE4 | *Methanosaeta concilii* | Conserved domain, transposase IS3 family protein, putative integrase |
| 85 | 336 | 0.01 | | | |
| 86 | 332 | 0.01 | | | |
| 87 | 331 | 0.01 | N8P3X5 | *Acinetobacter bohemicus* | Uncharacterized protein |
| 88 | 330 | 0.01 | | | |
| 89 | 329 | 0.01 | F4BXF6 | *Methanosaeta concilii* | Uncharacterized protein |
| 90 | 327 | 0.01 | A0A1Q3QHS9 | *Bacteroidia bacterium* | Uncharacterized protein |
| 91 | 322 | 0.01 | H8MSC6 | *Corallococcus coralloides* | Non-ribosomal peptide synthetase |
| 92 | 321 | 0.01 | | | |
| 93 | 320 | 0.01 | | | |
| 94 | 314 | 0.01 | F4BVL8 | *Methanosaeta concilii* | Uncharacterized protein |
| 95 | 313 | 0.01 | | | |
| 96 | 311 | 0.01 | F4BZ62 | *Methanosaeta concilii* | Conserved domain protein |
| 97 | 307 | 0.01 | A0A1Q3QCN8 | *Bacteroidia bacterium* | Uncharacterized protein |
| 98 | 307 | 0.01 | A0A1Q3QIV6 | *Bacteroidia bacterium* | Uncharacterized protein |
| 99 | 307 | 0.01 | N8YFI4 | *Acinetobacter gerneri* | Uncharacterized protein |
| 100 | 305 | 0.01 | | | |
| 101 | 304 | 0.01 | A0A1Q3QBE7 | *Bacteroidia bacterium* | Uncharacterized protein |
| 102 | 302 | 0.01 | F4BVN9 | *Methanosaeta concilii* | Conserved repeat domain protein |
| 103 | 301 | 0.01 | N9GC16 | *Acinetobacter lwoffii* | Uncharacterized protein |
| 104 | 300 | 0.01 | F4BTW7 | *Methanosaeta concilii* | Conserved domain protein |
| 105 | 297 | 0.01 | A0A1Q3QDS6 | *Bacteroidia bacterium* | Uncharacterized protein |

| | | | | | |
|---|---|---|---|---|---|
| 106 | 297 | 0.01 | F4BUE8 | *Methanosaeta concilii* | Transposase, predicted |
| 107 | 291 | 0.01 | A0A1Q3QHX6 | *Bacteroidia bacterium* | Multidrug transporter AcrB |
| 108 | 290 | 0.01 | | | |
| 109 | 289 | 0.01 | | | |
| 110 | 288 | 0.01 | A0A1Q3QI54 | *Bacteroidia bacterium* | SusC/RagA family TonB-linked outer membrane protein |
| 111 | 287 | 0.01 | F4BXK5 | *Methanosaeta concilii* | Transposase |
| 112 | 284 | 0.01 | F4BX50 | *Methanosaeta concilii* | Type III restriction enzyme, res subunit |
| 113 | 284 | 0.01 | | | |
| 114 | 283 | 0.01 | D2E402 | *Kernia pachypleura* | Ribosomal protein 3 |
| 115 | 283 | 0.01 | | | |
| 116 | 282 | 0.01 | | | |
| 117 | 281 | 0.01 | A0A0W8FAI0 | *hydrocarbon metagenome* | Uncharacterized protein |
| 118 | 281 | 0.01 | A0A1Q3QIY4 | *Bacteroidia bacterium* | SusC/RagA family protein |
| 119 | 279 | 0.01 | A0A1Q3QGJ2 | *Bacteroidia bacterium* | Uncharacterized protein |
| 120 | 278 | 0.01 | | | |
| 121 | 277 | 0.01 | | | |
| 122 | 276 | 0.01 | N8YF58 | *Acinetobacter gerneri* | Uncharacterized protein |
| 123 | 276 | 0.01 | | | |
| 124 | 276 | 0.01 | | | |
| 125 | 275 | 0.01 | A0A1Q3QB47 | *Bacteroidia bacterium* | Deoxyribonuclease HsdR |
| 126 | 274 | 0.01 | F4BU67 | *Methanosaeta concilii* | Multi-sensor hybrid histidine kinase |
| 127 | 272 | 0.01 | F4BT93 | *Methanosaeta concilii* | Uncharacterized protein |
| 128 | 272 | 0.01 | | | |
| 129 | 271 | 0.01 | | | |
| 130 | 271 | 0.01 | | | |
| 131 | 269 | 0.01 | | | |
| 132 | 268 | 0.01 | | | |
| 133 | 267 | 0.01 | | | |
| 134 | 267 | 0.01 | | | |
| 135 | 266 | 0.01 | F4BXN8 | *Methanosaeta concilii* | CobN/magnesium chelatase domain protein |
| 136 | 266 | 0.01 | | | |
| 137 | 265 | 0.01 | | | |
| 138 | 263 | 0.01 | A0A1Q3QF20 | *Bacteroidia bacterium* | SusC/RagA family protein |
| 139 | 263 | 0.01 | A0A1Q3QI16 | *Bacteroidia bacterium* | SusC/RagA family TonB-linked outer membrane protein |
| 140 | 262 | 0.01 | A0A1Q3QH94 | *Bacteroidia bacterium* | DNA-directed DNA polymerase (EC 2.7.7.7) |
| 141 | 262 | 0.01 | | | |

| | | | | | |
|---|---|---|---|---|---|
| 142 | 262 | 0.01 | | | |
| 143 | 262 | 0.01 | | | |
| 144 | 261 | 0.01 | | | |
| 145 | 260 | 0.01 | F4BXH3 | *Methanosaeta concilii* | Cobaltochelatase, CobN subunit |
| 146 | 259 | 0.01 | A0A1Q3QIK1 | *Bacteroidia bacterium* | Phosphoribosylformylglycinamidine synthase (FGAM synthase) (FGAMS) (EC 6.3.5.3) (Formylglycinamide ribonucleotide amidotransferase) (FGAR amidotransferase) (FGAR-AT) |
| 147 | 257 | 0.01 | A0A1Q3Q9Q2 | *Bacteroidia bacterium* | SusC/RagA family protein |
| 148 | 257 | 0.01 | F4BT87 | *Methanosaeta concilii* | TPR-repeat protein |
| 149 | 257 | 0.01 | F4BXG6 | *Methanosaeta concilii* | Cobaltochelatase, CobN subunit (EC 6.6.1.2) |
| 150 | 256 | 0.01 | | | |
| 151 | 253 | 0.01 | A0A1Q3QAN2 | *Bacteroidia bacterium* | SusC/RagA family TonB-linked outer membrane protein |
| 152 | 251 | 0.01 | A0A1Q3Q9V3 | *Bacteroidia bacterium* | Peptidase S9 |
| 153 | 249 | 0.01 | A0A1M6CCR6 | *Tessaracoccus bendigoensis* | Uncharacterized protein |
| 154 | 249 | 0.01 | A0A1Q3QA74 | *Bacteroidia bacterium* | TonB-dependent receptor |
| 155 | 248 | 0.01 | H8MS99 | *Corallococcus coralloides* | Adventurous gliding motility protein AgmK |
| 156 | 247 | 0.01 | A0A1Q3QBC6 | *Bacteroidia bacterium* | Uncharacterized protein |
| 157 | 247 | 0.01 | A0A1Q3QE63 | *Bacteroidia bacterium* | Uncharacterized protein |
| 158 | 245 | 0.01 | A0A077XLZ7 | *Sphinx1.76-related DNA* | Replication protein |
| 159 | 245 | 0.01 | A0A1Q3QF86 | *Bacteroidia bacterium* | Multidrug transporter AcrB |
| 160 | 244 | 0.01 | F4C043 | *Methanosaeta concilii* | Membrane protein, putative |
| 161 | 243 | 0.01 | A0A1G4G9S9 | *Petrimonas mucosa* | TonB-dependent receptor SusC |
| 162 | 243 | 0.01 | A0A1Q3QGM8 | *Bacteroidia bacterium* | SusC/RagA family TonB-linked outer membrane protein |
| 163 | 243 | 0.01 | F4BV15 | *Methanosaeta concilii* | Transposase, IS605 OrfB family |
| 164 | 243 | 0.01 | | | |
| 165 | 242 | 0.01 | A0A1Q3QAH7 | *Bacteroidia bacterium* | DNA polymerase I (EC 2.7.7.7) |
| 166 | 242 | 0.01 | A0A1Q3QDC9 | *Bacteroidia bacterium* | SusC/RagA family TonB-linked outer membrane protein |
| 167 | 242 | 0.01 | | | |
| 168 | 241 | 0.01 | F4BWZ9 | *Methanosaeta concilii* | Two-component hybrid sensor and regulator |
| 169 | 240 | 0.01 | A0A1Q3QBL0 | *Bacteroidia bacterium* | Acriflavine resistance protein B |
| 170 | 240 | 0.01 | A0A1Q3QD12 | *Bacteroidia bacterium* | SusC/RagA family TonB-linked outer membrane protein |
| 171 | 240 | 0.01 | | | |
| 172 | 239 | 0.01 | A0A1Q3QFV8 | *Bacteroidia bacterium* | DNA helicase (EC 3.6.4.12) |
| 173 | 239 | 0.01 | F4BUA4 | *Methanosaeta concilii* | Transposase, IS605 OrfB family |
| 174 | 238 | 0.01 | F4BTN1 | *Methanosaeta concilii* | Two-component hybrid sensor and regulator |
| 175 | 238 | 0.01 | | | |
| 176 | 238 | 0.01 | | | |

| | | | | | |
|---|---|---|---|---|---|
| 177 | 237 | 0.01 | A0A1R3TE89 | *Proteiniphilum saccharofermentans* | Uncharacterized protein |
| 178 | 237 | 0.01 | F4C0L1 | *Methanosaeta concilii* | TPR-repeat protein |
| 179 | 236 | 0.01 | | | |
| 180 | 235 | 0.01 | A0A0N1L384 | *Novosphingobium sp. AAP1* | Glutamate dehydrogenase |
| 181 | 235 | 0.01 | A0A1Q3QG05 | *Bacteroidia bacterium* | Multifunctional fusion protein [Includes: Protein translocase subunit SecD; Protein-export membrane protein SecF] |
| 182 | 234 | 0.01 | A0A1M6CDN2 | *Tessaracoccus bendigoensis* | Multicopper oxidase with three cupredoxin domains (Includes cell division protein FtsP and spore coat protein CotA) |
| 183 | 234 | 0.01 | A0A1Q3QCP4 | *Bacteroidia bacterium* | SusC/RagA family TonB-linked outer membrane protein |
| 184 | 234 | 0.01 | | | |
| 185 | 233 | 0.01 | F4BY67 | *Methanosaeta concilii* | Uncharacterized protein |
| 186 | 232 | 0.01 | | | |
| 187 | 231 | 0.01 | F4BXB3 | *Methanosaeta concilii* | Transposase, IS605 family protein, putative MITE |
| 188 | 231 | 0.01 | | | |
| 189 | 231 | 0.01 | | | |
| 190 | 230 | 0.01 | F4BWV9 | *Methanosaeta concilii* | S-layer-related duplication domain protein |
| 191 | 230 | 0.01 | | | |
| 192 | 228 | 0.01 | A0A1Q3QGP0 | *Bacteroidia bacterium* | Glycosyl hydrolase family 35 |
| 193 | 228 | 0.01 | | | |
| 194 | 228 | 0.01 | | | |
| 195 | 226 | 0.01 | F4BVL9 | *Methanosaeta concilii* | Uncharacterized protein |
| 196 | 226 | 0.01 | | | |
| 197 | 225 | 0.01 | | | |
| 198 | 225 | 0.01 | | | |
| 199 | 224 | 0.01 | | | |
| 200 | 224 | 0.01 | | | |
| 201 | 223 | 0.01 | A0A1Q3QI74 | *Bacteroidia bacterium* | Peptidase M16 |
| 202 | 223 | 0.01 | A0A1Q3QIQ0 | *Bacteroidia bacterium* | Uncharacterized protein |
| 203 | 223 | 0.01 | | | |
| 204 | 222 | 0.01 | A0A1Q3QBF6 | *Bacteroidia bacterium* | SusC/RagA family protein |
| 205 | 222 | 0.01 | A0A1Q3QDK9 | *Bacteroidia bacterium* | Uncharacterized protein |
| 206 | 222 | 0.01 | A0A1Q3QJE6 | *Bacteroidia bacterium* | Cation transporter |
| 207 | 222 | 0.01 | | | |
| 208 | 222 | 0.01 | | | |
| 209 | 221 | 0.01 | A0A1Q3QD11 | *Bacteroidia bacterium* | SusC/RagA family protein |
| 210 | 220 | 0.01 | A0A1Q3QGR5 | *Bacteroidia bacterium* | SusC/RagA family TonB-linked outer membrane protein |

| | | | | | |
|---|---|---|---|---|---|
| 211 | 220 | 0.01 | F1Z681 | *Novosphingobium nitrogenifigens* | Type I site-specific deoxyribonuclease, HsdR family |
| 212 | 220 | 0.01 | F4BSZ2 | *Methanosaeta concilii* | Oligosaccharyl transferase STT3 subunit |
| 213 | 220 | 0.01 | F4BVN7 | *Methanosaeta concilii* | Pentapeptide repeat protein |
| 214 | 220 | 0.01 | F4C050 | *Methanosaeta concilii* | Asparagine synthase (Glutamine-hydrolyzing) (EC 6.3.5.4) |
| 215 | 220 | 0.01 | | | |
| 216 | 219 | 0.01 | A0A1Q3Q9V2 | *Bacteroidia bacterium* | Phosphate transporter |
| 217 | 219 | 0.01 | A0A1Q3QFQ4 | *Bacteroidia bacterium* | Uncharacterized protein |
| 218 | 219 | 0.01 | F4BVI9 | *Methanosaeta concilii* | IS3/IS66 family element, Orf1 protein |
| 219 | 219 | 0.01 | | | |
| 220 | 218 | 0.01 | A0A0W8FC74 | *hydrocarbon metagenome* | Sensor histidine kinase |
| 221 | 218 | 0.01 | F1ZDD0 | *Novosphingobium nitrogenifigens* | Penicillin amidase family protein |
| 222 | 218 | 0.01 | F4BZ35 | *Methanosaeta concilii* | Uncharacterized protein |
| 223 | 218 | 0.01 | | | |
| 224 | 218 | 0.01 | | | |
| 225 | 218 | 0.01 | | | |
| 226 | 218 | 0.01 | | | |
| 227 | 217 | 0.01 | A0A1Q3QAJ6 | *Bacteroidia bacterium* | Uncharacterized protein |
| 228 | 217 | 0.01 | F4BSW7 | *Methanosaeta concilii* | Multi-sensor hybrid histidine kinase |
| 229 | 217 | 0.01 | F4BW20 | *Methanosaeta concilii* | Glycosyl transferase group 2/group 1 fusion protein |
| 230 | 217 | 0.01 | | | |
| 231 | 216 | 0.01 | A0A1Q3QBY0 | *Bacteroidia bacterium* | Phosphoenolpyruvate synthase |
| 232 | 216 | 0.01 | A0A1Q3QCR5 | *Bacteroidia bacterium* | Isoleucine--tRNA ligase (EC 6.1.1.5) (Isoleucyl-tRNA synthetase) (IleRS) |
| 233 | 216 | 0.01 | F4BXZ6 | *Methanosaeta concilii* | S-layer-related duplication domain protein |
| 234 | 216 | 0.01 | | | |
| 235 | 216 | 0.01 | | | |
| 236 | 216 | 0.01 | | | |
| 237 | 215 | 0.01 | A0A0F0LYK2 | *Microbacterium ginsengisoli* | Putative cadmium-transporting ATPase (EC 3.6.3.3) |
| 238 | 215 | 0.01 | A0A1Q3QJ62 | *Bacteroidia bacterium* | Uncharacterized protein (Fragment) |
| 239 | 215 | 0.01 | | | |
| 240 | 214 | 0.01 | A0A1Q3QAN0 | *Bacteroidia bacterium* | Uncharacterized protein |
| 241 | 214 | 0.01 | | | |
| 242 | 214 | 0.01 | | | |
| 243 | 214 | 0.01 | | | |
| 244 | 213 | 0.01 | A0A1Q3Q9W7 | *Bacteroidia bacterium* | Uncharacterized protein |
| 245 | 213 | 0.01 | A0A1Q3QED9 | *Bacteroidia bacterium* | Uncharacterized protein |
| 246 | 213 | 0.01 | A0A1Q3QGE6 | *Bacteroidia bacterium* | Uncharacterized protein |

| | | | | | |
|---|---|---|---|---|---|
| 247 | 213 | 0.01 | A0A1Q3QGG2 | *Bacteroidia bacterium* | SusC/RagA family protein |
| 248 | 213 | 0.01 | | | |
| 249 | 212 | 0.01 | A0A1Q3QDG3 | *Bacteroidia bacterium* | SusC/RagA family TonB-linked outer membrane protein |
| 250 | 212 | 0.01 | F0M3U0 | *Pseudarthrobacter phenanthrenivorans* | Phospholipase/Carboxylesterase |
| 251 | 212 | 0.01 | | | |
| 252 | 211 | 0.01 | A0A1Q3QC05 | *Bacteroidia bacterium* | Uncharacterized protein |
| 253 | 211 | 0.01 | A0A1Q3QES6 | *Bacteroidia bacterium* | Uncharacterized protein |
| 254 | 211 | 0.01 | F4BXL6 | *Methanosaeta concilii* | Conserved domain protein |
| 255 | 211 | 0.01 | | | |
| 256 | 211 | 0.01 | | | |
| 257 | 211 | 0.01 | | | |
| 258 | 210 | 0.01 | A0A0Q7S8L4 | *Pelomonas sp.* | Transposase |
| 259 | 210 | 0.01 | A0A1Q3QCH6 | *Bacteroidia bacterium* | Nitrate reductase subunit alpha |
| 260 | 210 | 0.01 | A0A1Q3QFU6 | *Bacteroidia bacterium* | TonB-dependent receptor |
| 261 | 210 | 0.01 | | | |
| 262 | 210 | 0.01 | | | |
| 263 | 209 | 0.01 | | | |
| 264 | 209 | 0.01 | | | |
| 265 | 208 | 0.01 | A0A0N1ANW5 | *Novosphingobium sp. AAP1* | Cellulose synthase |
| 266 | 208 | 0.01 | A0A1Q3QAN4 | *Bacteroidia bacterium* | Uncharacterized protein |
| 267 | 208 | 0.01 | F4BXF7 | *Methanosaeta concilii* | Transposase, putative integrase |
| 268 | 208 | 0.01 | | | |
| 269 | 207 | 0.01 | A0A1Q3QIQ9 | *Bacteroidia bacterium* | Uncharacterized protein |
| 270 | 207 | 0.01 | F4BYM4 | *Methanosaeta concilii* | Chromosome partition protein Smc |
| 271 | 207 | 0.01 | | | |
| 272 | 207 | 0.01 | | | |
| 273 | 206 | 0.01 | A0A1Q3QIR5 | *Bacteroidia bacterium* | Alpha-glucan phosphorylase |
| 274 | 206 | 0.01 | A0A1Q3QJ29 | *Bacteroidia bacterium* | DNA mismatch repair protein MutS |
| 275 | 205 | 0.01 | H8MFD7 | *Corallococcus coralloides* | Long-chain-fatty-acid--CoA ligase |
| 276 | 205 | 0.01 | L7LGQ0 | *Gordonia sihwensis* | Putative non-ribosomal peptide synthetase |
| 277 | 205 | 0.01 | | | |
| 278 | 205 | 0.01 | | | |
| 279 | 204 | 0.01 | | | |
| 280 | 204 | 0.01 | | | |
| 281 | 203 | 0.01 | A0A1G4GAX8 | *Petrimonas mucosa* | Uncharacterized protein |
| 282 | 203 | 0.01 | A0A1Q3QDL1 | *Bacteroidia bacterium* | Methionine synthase |

| | | | | | |
|---|---|---|---|---|---|
| 283 | 203 | 0.01 | P11408 | *Moraxella sp.* | Modification methylase MspI (M.MspI) (EC 2.1.1.37) (Cytosine-specific methyltransferase MspI) |
| 284 | 203 | 0.01 | | | |
| 285 | 202 | 0.01 | | | |
| 286 | 202 | 0.01 | | | |
| 287 | 201 | 0.01 | F4BY50 | *Methanosaeta concilii* | ATP-dependent RNA helicase, putative |
| 288 | 201 | 0.01 | | | |
| 289 | 201 | 0.01 | | | |
| 290 | 201 | 0.01 | | | |
| 291 | 200 | 0.01 | F4BWJ6 | *Methanosaeta concilii* | Peptidase families S8 and S53 protein |
| 292 | 200 | 0.01 | F4BWT2 | *Methanosaeta concilii* | DNA polymerase (EC 2.7.7.7) |
| 293 | 200 | 0.01 | F5XPC2 | *Microlunatus phosphovorus* | Uncharacterized protein |
| 294 | 200 | 0.01 | | | |
| 295 | 199 | 0.01 | A0A1Q3QB44 | *Bacteroidia bacterium* | Uncharacterized protein |
| 296 | 199 | 0.01 | A0A1Q3QBJ9 | *Bacteroidia bacterium* | Uncharacterized protein |
| 297 | 199 | 0.01 | A0A1R3T512 | *Proteiniphilum saccharofermentans* | TonB-linked outer membrane protein, SusC/RagA family |
| 298 | 199 | 0.01 | D8L7S1 | *Paramecium caudatum* | NADH-ubiquinone oxidoreductase chain 5 (EC 1.6.5.3) |
| 299 | 198 | 0.01 | A0A0R3R918 | *Brugia timori* | Uncharacterized protein |
| 300 | 198 | 0.01 | F4BZ75 | *Methanosaeta concilii* | Carbamoyl-phosphate synthase large chain (EC 6.3.5.5) (Carbamoyl-phosphate synthetase ammonia chain) |
| 301 | 198 | 0.01 | S2WV93 | *Delftia acidovorans* | Uncharacterized protein |
| 302 | 198 | 0.01 | | | |
| 303 | 197 | 0.01 | | | |
| 304 | 197 | 0.01 | | | |
| 305 | 196 | 0.01 | F4BU15 | *Methanosaeta concilii* | DNA mismatch repair protein MutS |
| 306 | 196 | 0.01 | F5XPA1 | *Microlunatus phosphovorus* | Uncharacterized protein |
| 307 | 196 | 0.01 | | | |
| 308 | 195 | 0.01 | A0A1Q3Q9V6 | *Bacteroidia bacterium* | Multidrug transporter AcrB |
| 309 | 195 | 0.01 | A0A1Q3QF64 | *Bacteroidia bacterium* | Alpha-rhamnosidase |
| 310 | 195 | 0.01 | U6RCL0 | *Bacteroides sp. HPS0048* | Uncharacterized protein |
| 311 | 195 | 0.01 | | | |
| 312 | 194 | 0.01 | A0A1Q3QED5 | *Bacteroidia bacterium* | Multidrug transporter AcrB |
| 313 | 194 | 0.01 | A0A1Q3QF91 | *Bacteroidia bacterium* | Alpha-rhamnosidase |
| 314 | 194 | 0.01 | A0A1Q3QJP8 | *Bacteroidia bacterium* | Cytochrome C biogenesis protein |
| 315 | 194 | 0.01 | | | |
| 316 | 193 | 0.01 | A0A1Q3QGG1 | *Bacteroidia bacterium* | Uncharacterized protein |

| | | | | | |
|---|---|---|---|---|---|
| 317 | 193 | 0.01 | A0A1Q3QIS8 | *Bacteroidia bacterium* | NADH dehydrogenase |
| 318 | 193 | 0.01 | F4BVD8 | *Methanosaeta concilii* | Cation transporting P-type ATPase |
| 319 | 193 | 0.01 | F4BVJ1 | *Methanosaeta concilii* | Beta-propeller repeat protein |
| 320 | 193 | 0.01 | | | |
| 321 | 192 | 0.01 | A0A1Q3QFG4 | *Bacteroidia bacterium* | Alpha-galactosidase |
| 322 | 192 | 0.01 | A0A1Q3QFT2 | *Bacteroidia bacterium* | Uncharacterized protein |
| 323 | 192 | 0.01 | A0A1Q3QI00 | *Bacteroidia bacterium* | Tricorn protease homolog (EC 3.4.21.-) |
| 324 | 192 | 0.01 | | | |
| 325 | 192 | 0.01 | | | |
| 326 | 192 | 0.01 | | | |
| 327 | 191 | 0.01 | F4BVJ0 | *Methanosaeta concilii* | Uncharacterized protein |
| 328 | 191 | 0.01 | H8MSC8 | *Corallococcus coralloides* | Tyrocidine synthase |
| 329 | 191 | 0.01 | | | |
| 330 | 191 | 0.01 | | | |
| 331 | 190 | 0.01 | A0A154DNE7 | *Acinetobacter baumannii* | mRNA endoribonuclease LS (EC 3.1.-.-) |
| 332 | 190 | 0.01 | A0A1Q3QFK3 | *Bacteroidia bacterium* | Alpha-rhamnosidase |
| 333 | 190 | 0.01 | A0A1Q3QHQ6 | *Bacteroidia bacterium* | SusC/RagA family protein |
| 334 | 190 | 0.01 | A0A1Q3QIM7 | *Bacteroidia bacterium* | Uncharacterized protein |
| 335 | 190 | 0.01 | | | |
| 336 | 189 | 0.01 | A0A0E3T6K6 | *Gordonia phage Gmala1* | Tape measure protein |
| 337 | 189 | 0.01 | A0A1Q3Q9V8 | *Bacteroidia bacterium* | TonB-dependent receptor |
| 338 | 189 | 0.01 | | | |
| 339 | 189 | 0.01 | | | |
| 340 | 189 | 0.01 | | | |
| 341 | 189 | 0.01 | | | |
| 342 | 189 | 0.01 | | | |
| 343 | 188 | 0.01 | A0A1Q3QA64 | *Bacteroidia bacterium* | Uncharacterized protein |
| 344 | 188 | 0.01 | A0A1Q3QD21 | *Bacteroidia bacterium* | Beta-galactosidase (EC 3.2.1.23) (Lactase) |
| 345 | 188 | 0.01 | A0A1Q3QHZ1 | *Bacteroidia bacterium* | Peptidase M10A/M12B |
| 346 | 188 | 0.01 | A0A1Q3QI69 | *Bacteroidia bacterium* | Uncharacterized protein |
| 347 | 188 | 0.01 | A0A1Q3QIW6 | *Bacteroidia bacterium* | Uncharacterized protein |
| 348 | 188 | 0.01 | F4BZ38 | *Methanosaeta concilii* | Transposase, IS116/IS110/IS902 family |
| 349 | 188 | 0.01 | | | |
| 350 | 187 | 0.01 | A0A1Q3QD59 | *Bacteroidia bacterium* | Translation initiation factor IF-2 |
| 351 | 187 | 0.01 | F4BXQ5 | *Methanosaeta concilii* | Two-component hybrid sensor and regulator |
| 352 | 187 | 0.01 | F4BY09 | *Methanosaeta concilii* | DNA polymerase II large subunit (Pol II) (EC 2.7.7.7) |

| | | | | | |
|---|---|---|---|---|---|
| 353 | 187 | 0.01 | | | |
| 354 | 187 | 0.01 | | | |
| 355 | 187 | 0.01 | | | |
| 356 | 187 | 0.01 | | | |
| 357 | 187 | 0.01 | | | |
| 358 | 186 | 0.01 | A0A151BT14 | *Branchiibius sp.* | Mercuric reductase |
| 359 | 186 | 0.01 | A0A1Q3QCA7 | *Bacteroidia bacterium* | Glycosyl hydrolase family 31 |
| 360 | 186 | 0.01 | A0A1Q3QGC9 | *Bacteroidia bacterium* | Peptidase M16 |
| 361 | 186 | 0.01 | F4BU80 | *Methanosaeta concilii* | Transposase, IS605 OrfB family |
| 362 | 186 | 0.01 | | | |
| 363 | 185 | 0.01 | A0A1M3FLT2 | *Bacteroidia bacterium* | Uncharacterized protein |
| 364 | 185 | 0.01 | A0A1M6WBE7 | *Fibrobacter sp.* | Transposase (Or an inactivated derivative) |
| 365 | 185 | 0.01 | A0A1Q3QA25 | *Bacteroidia bacterium* | UvrABC system protein C (Protein UvrC) (Excinuclease ABC subunit C) |
| 366 | 185 | 0.01 | A0A1Q3QD60 | *Bacteroidia bacterium* | Alanine racemase (EC 5.1.1.1) |
| 367 | 185 | 0.01 | A0A1Q3QFA7 | *Bacteroidia bacterium* | Glycoside hydrolase family 2 |
| 368 | 185 | 0.01 | A0A1Q3QG11 | *Bacteroidia bacterium* | Capsular biosynthesis protein |
| 369 | 185 | 0.01 | A1W5C2 | *Acidovorax sp.* | IstB domain protein ATP-binding protein |
| 370 | 184 | 0.01 | A0A1G4G461 | *Petrimonas mucosa* | Outer membrane protein assembly factor BamA |
| 371 | 184 | 0.01 | A0A1Q3QAU4 | *Bacteroidia bacterium* | Uncharacterized protein |
| 372 | 184 | 0.01 | A0A1Q3QB09 | *Bacteroidia bacterium* | Glycoside hydrolase |
| 373 | 184 | 0.01 | A0A1Q3QGV1 | *Bacteroidia bacterium* | Uncharacterized protein |
| 374 | 184 | 0.01 | A1W6Q0 | *Acidovorax sp.* | Integrase, catalytic region |
| 375 | 184 | 0.01 | | | |
| 376 | 184 | 0.01 | | | |
| 377 | 184 | 0.01 | | | |
| 378 | 184 | 0.01 | | | |
| 379 | 183 | 0.01 | A0A0N0JFZ2 | *Novosphingobium sp.* | Glutamate synthase |
| 380 | 183 | 0.01 | A0A1Q3QI34 | *Bacteroidia bacterium* | Uncharacterized protein |
| 381 | 183 | 0.01 | | | |
| 382 | 183 | 0.01 | | | |
| 383 | 182 | 0.01 | A0A1Q3QBL5 | *Bacteroidia bacterium* | Uncharacterized protein |
| 384 | 182 | 0.01 | A0A1Q3QF81 | *Bacteroidia bacterium* | Uncharacterized protein |
| 385 | 182 | 0.01 | A0A1Q3QIV4 | *Bacteroidia bacterium* | Uncharacterized protein |
| 386 | 182 | 0.01 | F0M493 | *Pseudarthrobacter phenanthrenivorans* | Uncharacterized protein |
| 387 | 182 | 0.01 | F4BWG4 | *Methanosaeta concilii* | Uncharacterized protein |
| 388 | 182 | 0.01 | | | |

| | | | | | |
|---|---|---|---|---|---|
| 389 | 182 | 0.01 | | | |
| 390 | 181 | 0.01 | A0A1Q3QBI7 | *Bacteroidia bacterium* | Uncharacterized protein |
| 391 | 181 | 0.01 | A0A1Q3QIS1 | *Bacteroidia bacterium* | Uncharacterized protein |
| 392 | 181 | 0.01 | F4BW92 | *Methanosaeta concilii* | Cation-transporting P-type ATPase |
| 393 | 181 | 0.01 | | | |
| 394 | 180 | 0.01 | C5CEK1 | *Kosmotoga olearia* | IstB domain protein ATP-binding protein |
| 395 | 180 | 0.01 | F4BWV3 | *Methanosaeta concilii* | Alanine--tRNA ligase (EC 6.1.1.7) (Alanyl-tRNA synthetase) (AlaRS) |
| 396 | 180 | 0.01 | F4BXD3 | *Methanosaeta concilii* | S-layer-related duplication domain protein |
| 397 | 180 | 0.01 | | | |
| 398 | 179 | 0.01 | A0A1Q3QBV2 | *Bacteroidia bacterium* | Serine protease |
| 399 | 179 | 0.01 | A0A1Q3QF68 | *Bacteroidia bacterium* | Glycoside hydrolase |
| 400 | 179 | 0.01 | | | |
| 401 | 179 | 0.01 | | | |
| 402 | 178 | 0.01 | A0A1Q3QAC3 | *Bacteroidia bacterium* | Leucine--tRNA ligase (Fragment) |
| 403 | 178 | 0.01 | A0A1Q3QF16 | *Bacteroidia bacterium* | Beta-galactosidase (EC 3.2.1.23) (Lactase) |
| 404 | 178 | 0.01 | A0A1Q3QFY4 | *Bacteroidia bacterium* | Uncharacterized protein |
| 405 | 178 | 0.01 | A0A1Q3QIH3 | *Bacteroidia bacterium* | Uncharacterized protein |
| 406 | 178 | 0.01 | H8MH05 | *Corallococcus coralloides* | Long-chain-fatty-acid--CoA ligase |
| 407 | 178 | 0.01 | | | |
| 408 | 178 | 0.01 | | | |
| 409 | 177 | 0.01 | A0A1Q3QAJ9 | *Bacteroidia bacterium* | Uncharacterized protein |
| 410 | 177 | 0.01 | A0A1Q3QGA2 | *Bacteroidia bacterium* | Ferrous iron transport protein B |
| 411 | 177 | 0.01 | A0A1Q3QIL1 | *Bacteroidia bacterium* | Uncharacterized protein |
| 412 | 177 | 0.01 | A0A1Q3QIP6 | *Bacteroidia bacterium* | Transcription-repair-coupling factor (TRCF) (EC 3.6.4.-) |
| 413 | 177 | 0.01 | A7AI47 | *Parabacteroides merdae* | Uncharacterized protein |
| 414 | 177 | 0.01 | F4BYK4 | *Methanosaeta concilii* | Leucine--tRNA ligase (EC 6.1.1.4) (Leucyl-tRNA synthetase) (LeuRS) |
| 415 | 177 | 0.01 | P15689 | *Paramecium tetraurelia* | NADH-ubiquinone oxidoreductase 49 kDa subunit (EC 1.6.5.3) (EC 1.6.99.3) (NADH dehydrogenase subunit 7) |
| 416 | 177 | 0.01 | P11405 | *Moraxella sp.* | Type-2 restriction enzyme MspI (R.MspI) (EC 3.1.21.4) (Endonuclease MspI) (Type II restriction enzyme MspI) |
| 417 | 177 | 0.01 | | | |
| 418 | 177 | 0.01 | | | |
| 419 | 177 | 0.01 | | | |
| 420 | 177 | 0.01 | | | |
| 421 | 177 | 0.01 | | | |
| 422 | 176 | 0.01 | A0A1G4GAZ3 | *Petrimonas mucosa* | Uncharacterized protein |

| | | | | | |
|---|---|---|---|---|---|
| 423 | 176 | 0.01 | A0A1M6CEL6 | *Tessaracoccus bendigoensis* | Uncharacterized protein |
| 424 | 176 | 0.01 | A0A1Q3QAR6 | *Bacteroidia bacterium* | Mannose-6-phosphate isomerase |
| 425 | 176 | 0.01 | | | |
| 426 | 176 | 0.01 | | | |
| 427 | 175 | 0.01 | A0A1Q2CK56 | *Tessaracoccus sp. NSG39* | Uncharacterized protein |
| 428 | 175 | 0.01 | A0A1Q3QF30 | *Bacteroidia bacterium* | Hydrolase |
| 429 | 175 | 0.01 | A0A1Q3QGE3 | *Bacteroidia bacterium* | Uncharacterized protein |
| 430 | 175 | 0.01 | | | |
| 431 | 175 | 0.01 | | | |
| 432 | 175 | 0.01 | | | |
| 433 | 174 | 0.01 | F4BVH9 | *Methanosaeta concilii* | Glycosyl transferase, group 2 family protein |
| 434 | 174 | 0.01 | N1JKD7 | *Mesotoga infera* | RNA-directed DNA polymerase (Reverse transcriptase) |
| 435 | 174 | 0.01 | | | |
| 436 | 173 | 0.01 | A0A1Q3QBX1 | *Bacteroidia bacterium* | Polysaccharide biosynthesis protein |
| 437 | 173 | 0.01 | A0A1Q3QGI1 | *Bacteroidia bacterium* | Uncharacterized protein |
| 438 | 173 | 0.01 | A0A1Q3QHE9 | *Bacteroidia bacterium* | Glycosyl transferase |
| 439 | 173 | 0.01 | F4BTH0 | *Methanosaeta concilii* | Glycosyl transferase, group 2 family protein |
| 440 | 173 | 0.01 | F4BTY4 | *Methanosaeta concilii* | SNF2 family helicase (EC 3.6.1.-) |
| 441 | 173 | 0.01 | | | |
| 442 | 173 | 0.01 | | | |
| 443 | 172 | 0.01 | A0A180EWA0 | *Bacteroidales bacterium* | Replication protein |
| 444 | 172 | 0.01 | A0A1Q3QH96 | *Bacteroidia bacterium* | PAS domain-containing sensor histidine kinase |
| 445 | 172 | 0.01 | F0M490 | *Pseudarthrobacter phenanthrenivorans* | DNA primase/polymerase-like protein |
| 446 | 172 | 0.01 | F4BSY5 | *Methanosaeta concilii* | Uncharacterized protein |
| 447 | 172 | 0.01 | | | |
| 448 | 172 | 0.01 | | | |
| 449 | 171 | 0.01 | A0A1Q3Q9R3 | *Bacteroidia bacterium* | DUF5107 domain-containing protein |
| 450 | 171 | 0.01 | A0A1Q3QGP8 | *Bacteroidia bacterium* | Glycoside hydrolase |
| 451 | 171 | 0.01 | F4BUE7 | *Methanosaeta concilii* | Uncharacterized protein |
| 452 | 171 | 0.01 | | | |
| 453 | 171 | 0.01 | | | |
| 454 | 171 | 0.01 | | | |
| 455 | 170 | 0.01 | A0A1Q3QEW4 | *Bacteroidia bacterium* | Alanine--tRNA ligase (EC 6.1.1.7) (Alanyl-tRNA synthetase) (AlaRS) |
| 456 | 170 | 0.01 | A0A1Q3QGC5 | *Bacteroidia bacterium* | Uncharacterized protein |
| 457 | 170 | 0.01 | A0A1Q3QGL6 | *Bacteroidia bacterium* | Copper-translocating P-type ATPase |
| 458 | 170 | 0.01 | A0A1Q3QID2 | *Bacteroidia bacterium* | Uncharacterized protein |

| | | | | | |
|---|---|---|---|---|---|
| 459 | 170 | 0.01 | D8L7R1 | *Paramecium caudatum* | NADH-ubiquinone oxidoreductase chain 4 (EC 1.6.5.3) |
| 460 | 170 | 0.01 | F4BY71 | *Methanosaeta concilii* | WD40-like Beta Propeller Repeat protein |
| 461 | 170 | 0.01 | I4ZNR1 | *Acinetobacter sp. HA* | Uncharacterized protein (Fragment) |
| 462 | 170 | 0.01 | I8W058 | *Bacteroides cellulosilyticus* | Uncharacterized protein |
| 463 | 169 | 0.01 | A0A098BYW1 | *Fermentimonas caenicola* | Transposase |
| 464 | 169 | 0.01 | A0A1G4G8N7 | *Petrimonas mucosa* | ATP-dependent DNA helicase RecQ (EC 3.6.4.12) |
| 465 | 169 | 0.01 | A0A1Q3QBS3 | *Bacteroidia bacterium* | Uncharacterized protein |
| 466 | 169 | 0.01 | A0A1Q3QFW3 | *Bacteroidia bacterium* | Alpha-mannosidase |
| 467 | 169 | 0.01 | A0A1Q3QIT0 | *Bacteroidia bacterium* | Polyphosphate kinase (EC 2.7.4.1) (ATP-polyphosphate phosphotransferase) (Polyphosphoric acid kinase) |
| 468 | 169 | 0.01 | | | |
| 469 | 169 | 0.01 | V2USD3 | *Acinetobacter brisouii* | Uncharacterized protein |
| 470 | 169 | 0.01 | W9GKA2 | *Intrasporangium chromatireducens* | DNA primase (EC 2.7.7.-) |
| 471 | 168 | 0.01 | A0A023UP33 | *Magnusiomyces ingens* | Uncharacterized protein (Fragment) |
| 472 | 168 | 0.01 | A0A1Q3Q9X7 | *Bacteroidia bacterium* | Uncharacterized protein |
| 473 | 168 | 0.01 | A0A1Q3QFZ4 | *Bacteroidia bacterium* | Uncharacterized protein |
| 474 | 168 | 0.01 | F1Z7R9 | *Novosphingobium nitrogenifigens* | Type-IV secretion system protein TraC |
| 475 | 168 | 0.01 | F4BTS2 | *Methanosaeta concilii* | M6 family metalloprotease domain protein |
| 476 | 168 | 0.01 | F4BVQ0 | *Methanosaeta concilii* | Uncharacterized protein |
| 477 | 168 | 0.01 | | | |
| 478 | 168 | 0.01 | | | |
| 479 | 168 | 0.01 | | | |
| 480 | 167 | 0.01 | A0A0M3AFY3 | *Acinetobacter sp.* | Cytochrome C peroxidase |
| 481 | 167 | 0.01 | A0A1Q3QAM8 | *Bacteroidia bacterium* | Uncharacterized protein |
| 482 | 167 | 0.01 | A0A1Q3QC30 | *Bacteroidia bacterium* | 4-alpha-glucanotransferase |
| 483 | 167 | 0.01 | A0A1Q3QC51 | *Bacteroidia bacterium* | Valine--tRNA ligase (EC 6.1.1.9) (Valyl-tRNA synthetase) (ValRS) |
| 484 | 167 | 0.01 | A0A1Q3QCU6 | *Bacteroidia bacterium* | Uncharacterized protein |
| 485 | 167 | 0.01 | F4BXE8 | *Methanosaeta concilii* | Conserved domain protein |
| 486 | 167 | 0.01 | F4BZQ7 | *Methanosaeta concilii* | S-layer-related duplication domain protein |
| 487 | 167 | 0.01 | A6LE80 | *Parabacteroides distasonis* | DNA-directed RNA polymerase subunit beta' (RNAP subunit beta') (EC 2.7.7.6) (RNA polymerase subunit beta') (Transcriptase subunit beta') |
| 488 | 167 | 0.01 | | | |
| 489 | 167 | 0.01 | | | |
| 490 | 166 | 0.01 | A0A1Q3QC45 | *Bacteroidia bacterium* | Uncharacterized protein |
| 491 | 166 | 0.01 | A0A1Q3QFI3 | *Bacteroidia bacterium* | DNA topoisomerase IV |
| 492 | 166 | 0.01 | F4BU69 | *Methanosaeta concilii* | Hydrophobe/amphiphile efflux-3 (HAE3) family transporter |

| | | | | | |
|---|---|---|---|---|---|
| 493 | 166 | 0.01 | F4BXP6 | *Methanosaeta concilii* | IS66 family element, transposase |
| 494 | 166 | 0.01 | F4C0R1 | *Methanosaeta concilii* | Uncharacterized protein |
| 495 | 166 | 0.01 | H8MK59 | *Corallococcus coralloides* | Linear gramicidin synthase subunit C |
| 496 | 166 | 0.01 | | | |
| 497 | 165 | 0.01 | A0A1Q3QDR2 | *Bacteroidia bacterium* | Uncharacterized protein |
| 498 | 165 | 0.01 | A0A1Q3QEA7 | *Bacteroidia bacterium* | Acriflavine resistance protein B |
| 499 | 165 | 0.01 | A0A1Q3QGG3 | *Bacteroidia bacterium* | Hydrolase |
| 500 | 165 | 0.01 | F4BWB4 | *Methanosaeta concilii* | TPR-repeat protein |

**Table S4a.** Functional overview of M1A community (top 500 rank only).

| Rank | Count | Abundance | ID | Organism | Protein Names |
|---|---|---|---|---|---|
| 1 | 3584 | 0.12 | A0A0D5N7W1 | *Geobacter sulfurreducens* | ATPase |
| 2 | 3399 | 0.12 | | *NA* | NA |
| 3 | 3122 | 0.11 | Q39Y12 | *Geobacter metallireducens* | DNA-directed RNA polymerase subunit beta' (RNAP subunit beta') (EC 2.7.7.6) (RNA polymerase subunit beta') (Transcriptase subunit beta') |
| 4 | 3076 | 0.11 | Q39RM9 | *Geobacter metallireducens* | Membrane protein, putative |
| 5 | 3014 | 0.10 | Q74AK2 | *Geobacter sulfurreducens* | Sodium/solute symporter family protein |
| 6 | 2908 | 0.10 | Q748Y6 | *Geobacter sulfurreducens* | DNA-directed RNA polymerase subunit beta (RNAP subunit beta) (EC 2.7.7.6) (RNA polymerase subunit beta) (Transcriptase subunit beta) |
| 7 | 2778 | 0.10 | A0A0L8ETQ7 | *Achromobacter sp.* | Uncharacterized protein |
| 8 | 2592 | 0.09 | Q747C7 | *Geobacter sulfurreducens* | 60 kDa chaperonin (GroEL protein) (Protein Cpn60) |
| 9 | 2557 | 0.09 | | | |
| 10 | 2544 | 0.09 | Q748X8 | *Geobacter sulfurreducens* | Elongation factor Tu (EF-Tu) |
| 11 | 2541 | 0.09 | E8WU24 | *Geobacter sp.* | Helicase domain protein |
| 12 | 2507 | 0.09 | A5GAB4 | *Geobacter uraniireducens* | Putative GAF sensor protein |
| 13 | 2448 | 0.08 | | | |
| 14 | 2424 | 0.08 | A0A1F9PUL2 | *Desulfuromonadales bacterium* | Restriction endonuclease |
| 15 | 2416 | 0.08 | A0A0D5N8M0 | *Geobacter sulfurreducens* | MgtE integral membrane protein |
| 16 | 2342 | 0.08 | A0A0B5BF68 | *Geobacter pickeringii* | CRISPR-associated protein Cas3 |
| 17 | 2334 | 0.08 | Q748Y8 | *Geobacter sulfurreducens* | Elongation factor G 2 (EF-G 2) |
| 18 | 2274 | 0.08 | Q749A7 | *Geobacter sulfurreducens* | Protein translocase subunit SecY |
| 19 | 2266 | 0.08 | A0A0B4XMH5 | *Alcanivorax pacificus* | Adenine-specific DNA methylase containing a Zn-ribbon |
| 20 | 2243 | 0.08 | A0A1G0M9J2 | *Geobacteraceae bacterium* | Uncharacterized protein |
| 21 | 2241 | 0.08 | | | |
| 22 | 2043 | 0.07 | Q74FF1 | *Geobacter sulfurreducens* | Chaperone protein ClpB |
| 23 | 2015 | 0.07 | A0A0C5IZ87 | *Rugosibacter aromaticivorans* | Uncharacterized protein |
| 24 | 1982 | 0.07 | A0A0B5BJ81 | *Geobacter pickeringii* | NADH-quinone oxidoreductase subunit D (EC 1.6.5.11) (NADH dehydrogenase I subunit D) (NDH-1 subunit D) |

| | | | | | |
|---|---|---|---|---|---|
| 25 | 1958 | 0.07 | A0A0B5BF21 | *Geobacter pickeringii* | Glycosyl transferase family 1 |
| 26 | 1954 | 0.07 | A1AKN8 | *Pelobacter propionicus* | Resolvase, N-terminal domain protein |
| 27 | 1953 | 0.07 | Q39QB9 | *Geobacter metallireducens* | NADH dehydrogenase I, M subunit |
| 28 | 1926 | 0.07 | A0A0B5B7H3 | *Geobacter pickeringii* | Cytochrome C |
| 29 | 1904 | 0.07 | A0A0D5N8Q4 | *Geobacter sulfurreducens* | Transposase |
| 30 | 1875 | 0.06 | Q39Q43 | *Geobacter metallireducens* | Pyruvate-flavodoxin oxidoreductase (EC 1.2.7.-) |
| 31 | 1869 | 0.06 | Q747X9 | *Geobacter sulfurreducens* | Isoleucine--tRNA ligase (EC 6.1.1.5) (Isoleucyl-tRNA synthetase) (IleRS) |
| 32 | 1857 | 0.06 | S0G1I5 | *Desulfotignum phosphitoxidans* | Type I restriction enzyme HindVIIP/EcoKI protein HsdR (EC 3.1.21.3) |
| 33 | 1854 | 0.06 | Q74FK9 | *Geobacter sulfurreducens* | Sigma-54-dependent transcriptional response regulator |
| 34 | 1851 | 0.06 | Q39U60 | *Geobacter metallireducens* | Translation elongation factor G |
| 35 | 1843 | 0.06 | A0A1S0XXD8 | *Rhodanobacter sp.* | Histidine kinase |
| 36 | 1835 | 0.06 | A0A0A8WNU3 | *Geobacter sp.* | Protein TraC |
| 37 | 1797 | 0.06 | Q39WU2 | *Geobacter metallireducens* | Phosphate-selective outer membrane channel |
| 38 | 1776 | 0.06 | Q74C76 | *Geobacter sulfurreducens* | (R)-citramalate synthase (EC 2.3.1.182) (Citramalate synthase) |
| 39 | 1745 | 0.06 | Q39QB2 | *Geobacter metallireducens* | NADH dehydrogenase I, F subunit |
| 40 | 1743 | 0.06 | Q747H5 | *Geobacter sulfurreducens* | K(+)-insensitive pyrophosphate-energized proton pump (EC 3.6.1.1) (Membrane-bound proton-translocating pyrophosphatase) (Pyrophosphate-energized inorganic pyrophosphatase) (H(+)-PPase) |
| 41 | 1736 | 0.06 | A0A0B5BJ46 | *Geobacter pickeringii* | Uncharacterized protein |
| 42 | 1727 | 0.06 | A0A0B5BB65 | *Geobacter pickeringii* | Uncharacterized protein |
| 43 | 1718 | 0.06 | A0A0B5BCT5 | *Geobacter pickeringii* | NADH dehydrogenase |
| 44 | 1687 | 0.06 | Q39ZC8 | *Geobacter metallireducens* | Ferredoxin-dependent glutamate synthase |
| 45 | 1683 | 0.06 | A0A0D5NE26 | *Geobacter sulfurreducens* | Multidrug resistance protein B |
| 46 | 1653 | 0.06 | A5GFJ7 | *Geobacter uraniireducens* | Erythronolide synthase (EC 2.3.1.94) |
| 47 | 1650 | 0.06 | Q3A5L6 | *Pelobacter carbinolicus* | Uncharacterized protein |
| 48 | 1635 | 0.06 | A0A0L8EU90 | *Achromobacter sp.* | Uncharacterized protein |
| 49 | 1635 | 0.06 | Q74FM6 | *Geobacter sulfurreducens* | Pyruvate, phosphate dikinase (EC 2.7.9.1) |
| 50 | 1634 | 0.06 | | | |

| | | | | | |
|---|---|---|---|---|---|
| 51 | 1625 | 0.06 | A0A0D5NDJ8 | *Geobacter sulfurreducens* | Stage II sporulation protein E |
| 52 | 1621 | 0.06 | | | |
| 53 | 1603 | 0.06 | Q39RM8 | *Geobacter metallireducens* | Glycosyltransferase |
| 54 | 1589 | 0.05 | Q39YK5 | *Geobacter metallireducens* | Flagellar biogenesis protein FlhA |
| 55 | 1587 | 0.05 | A0A0D5N2P4 | *Geobacter sulfurreducens* | Uncharacterized protein |
| 56 | 1585 | 0.05 | A0A0B5BK17 | *Geobacter pickeringii* | Diguanylate cyclase |
| 57 | 1563 | 0.05 | | | |
| 58 | 1559 | 0.05 | A0A0D5N8G6 | *Geobacter sulfurreducens* | Cation transporter |
| 59 | 1552 | 0.05 | | | |
| 60 | 1548 | 0.05 | A0A1T4RHG2 | *Geobacter thiogenes* | Nif-specific regulatory protein |
| 61 | 1538 | 0.05 | A0A0B5BDJ5 | *Geobacter pickeringii* | NADH-quinone oxidoreductase subunit H (EC 1.6.5.11) (NADH dehydrogenase I subunit H) (NDH-1 subunit H) |
| 62 | 1515 | 0.05 | A0A0B5B8A8 | *Geobacter pickeringii* | Valine--tRNA ligase (EC 6.1.1.9) (Valyl-tRNA synthetase) (ValRS) |
| 63 | 1514 | 0.05 | Q39Y76 | *Geobacter metallireducens* | Sigma-54-dependent transcriptional response regulator |
| 64 | 1507 | 0.05 | A0A0D5N8Q2 | *Geobacter sulfurreducens* | Uncharacterized protein |
| 65 | 1503 | 0.05 | A0A0D5N8I5 | *Geobacter sulfurreducens* | Signal recognition particle protein (Fifty-four homolog) |
| 66 | 1490 | 0.05 | Q748Z4 | *Geobacter sulfurreducens* | 30S ribosomal protein S3 |
| 67 | 1479 | 0.05 | A0A0B5BJ26 | *Geobacter pickeringii* | Uncharacterized protein |
| 68 | 1472 | 0.05 | | | |
| 69 | 1466 | 0.05 | | | |
| 70 | 1458 | 0.05 | A1AKK8 | *Pelobacter propionicus* | Uncharacterized protein |
| 71 | 1450 | 0.05 | Q746X3 | *Geobacter sulfurreducens* | Bifunctional protein PutA |
| 72 | 1448 | 0.05 | Q39QB8 | *Geobacter metallireducens* | NADH dehydrogenase I, L subunit |
| 73 | 1445 | 0.05 | A0A0B5BI89 | *Geobacter pickeringii* | Type II secretion system protein |
| 74 | 1445 | 0.05 | Q74H82 | *Geobacter sulfurreducens* | Flagellar biogenesis master response receiver sensor histidine kinase, PAS and GAF domain-containing |
| 75 | 1433 | 0.05 | Q747M6 | *Geobacter sulfurreducens* | Ribonuclease G |
| 76 | 1432 | 0.05 | Q74H58 | *Geobacter sulfurreducens* | Chaperone protein DnaJ |

| | | | | | |
|---|---|---|---|---|---|
| 77 | 1431 | 0.05 | A0A0D5N8S3 | *Geobacter sulfurreducens* | NADH dehydrogenase |
| 78 | 1421 | 0.05 | A0A0D5N8E2 | *Geobacter sulfurreducens* | ATP synthase (EC 3.6.3.14) |
| 79 | 1417 | 0.05 | Q39RL8 | *Geobacter metallireducens* | Undecaprenyl-diphospho-oligosaccharide flippase |
| 80 | 1404 | 0.05 | Q39XL9 | *Geobacter metallireducens* | Sensor sigma-54-dependent transcriptional regulator, GAF and GAF domain-containing |
| 81 | 1397 | 0.05 | A5GB28 | *Geobacter uraniireducens* | Plasmid maintenance system antidote protein, XRE family |
| 82 | 1397 | 0.05 | B3E9R4 | *Geobacter lovleyi* | Isocitrate dehydrogenase, NADP-dependent (EC 1.1.1.42) |
| 83 | 1392 | 0.05 | A0A0D5NAK5 | *Geobacter sulfurreducens* | Bifunctional uridylyltransferase/uridylyl-removing enzyme (UTase/UR) (Bifunctional [protein-PII] modification enzyme) (Bifunctional nitrogen sensor protein) [Includes: [Protein-PII]-UMP uridylyl-removing enzyme (UR) (EC 3.1.4.-); [Protein-PII] uridylyltransferase (PII uridylyltransferase) (UTase) (EC 2.7.7.59)] |
| 84 | 1388 | 0.05 | A0A0D5N712 | *Geobacter sulfurreducens* | Phosphoglucomutase |
| 85 | 1386 | 0.05 | Q39ZH1 | *Geobacter metallireducens* | Glutamine--fructose-6-phosphate aminotransferase [isomerizing] (EC 2.6.1.16) (D-fructose-6-phosphate amidotransferase) (GFAT) (Glucosamine-6-phosphate synthase) (Hexosephosphate aminotransferase) (L-glutamine--D-fructose-6-phosphate amidotransferase) |
| 86 | 1381 | 0.05 | A0A0D5ND20 | *Geobacter sulfurreducens* | Amino acid dehydrogenase |
| 87 | 1379 | 0.05 | A0A1G0M577 | *Geobacteraceae bacterium* | Uncharacterized protein |
| 88 | 1374 | 0.05 | Q74AK3 | *Geobacter sulfurreducens* | Cation-translocating P-type ATPase |
| 89 | 1373 | 0.05 | Q39ZK4 | *Geobacter metallireducens* | Bifunctional glutamine synthetase adenylyltransferase/adenylyl-removing enzyme (ATP:glutamine synthetase adenylyltransferase) (ATase) [Includes: Glutamine synthetase adenylyl-L-tyrosine phosphorylase (EC 2.7.7.89) (Adenylyl removase) (AR) (AT-N); Glutamine synthetase adenylyl transferase (EC 2.7.7.42) (Adenylyl transferase) (AT) (AT-C)] |
| 90 | 1372 | 0.05 | A0A0B5BIS9 | *Geobacter pickeringii* | Multidrug transporter AcrB |
| 91 | 1369 | 0.05 | A0A0B5BH93 | *Geobacter pickeringii* | C-and b-type cytochrome L |
| 92 | 1369 | 0.05 | A0A0D5N5P7 | *Geobacter sulfurreducens* | Chemotaxis protein |
| 93 | 1357 | 0.05 | B5EGM4 | *Geobacter bemidjiensis* | Glycerol/1,2-propanediol dehydratase |
| 94 | 1349 | 0.05 | B3EAX1 | *Geobacter lovleyi* | Protein CyaE |
| 95 | 1335 | 0.05 | A0A1G0MAK9 | *Geobacteraceae bacterium* | Sodium:proton exchanger |
| 96 | 1333 | 0.05 | Q74FD5 | *Geobacter sulfurreducens* | Hydroxylamine reductase (EC 1.7.99.1) (Hybrid-cluster protein) (HCP) (Prismane protein) |
| 97 | 1329 | 0.05 | B3EA87 | *Geobacter lovleyi* | RNA polymerase sigma factor |
| 98 | 1325 | 0.05 | Q39ZE7 | *Geobacter metallireducens* | UvrABC system protein A (UvrA protein) (Excinuclease ABC subunit A) |

| | | | | | |
|---|---|---|---|---|---|
| 99 | 1322 | 0.05 | Q39T74 | *Geobacter metallireducens* | GTP/GDP 3'-pyrophosphokinase and (P)ppGpp 3'-pyrophosphohydrolase |
| 100 | 1318 | 0.05 | A0A0D5ND99 | *Geobacter sulfurreducens* | ATPase AAA |
| 101 | 1302 | 0.05 | Q39QD0 | *Geobacter metallireducens* | Periplasmically oriented, membrane-bound [NiFe]-hydrogenase, large subunit |
| 102 | 1297 | 0.04 | Q39Q36 | *Geobacter metallireducens* | Heterodisulfide oxidoreductase, FAD-binding and iron-sulfur cluster-binding subunit A |
| 103 | 1285 | 0.04 | A0A0D5N7R7 | *Geobacter sulfurreducens* | GntR family transcriptional regulator |
| 104 | 1281 | 0.04 | A0A0B5BDE2 | *Geobacter pickeringii* | ATP phosphoribosyltransferase regulatory subunit |
| 105 | 1275 | 0.04 | B3EAX6 | *Geobacter lovleyi* | Uncharacterized protein |
| 106 | 1272 | 0.04 | A0A0C1U608 | *Geobacter soli* | Chemotaxis protein |
| 107 | 1268 | 0.04 | A5GFJ6 | *Geobacter uraniireducens* | Beta-hydroxyacyl-(Acyl-carrier-protein) dehydratase, FabA/FabZ |
| 108 | 1265 | 0.04 | A0A0B5BAC1 | *Geobacter pickeringii* | Dihydrolipoyl dehydrogenase (EC 1.8.1.4) |
| 109 | 1263 | 0.04 | Q39V52 | *Geobacter metallireducens* | GAF sensor methyl-accepting chemotaxis sensory transducer, class 40H |
| 110 | 1255 | 0.04 | A5GAY1 | *Geobacter uraniireducens* | DNA-directed RNA polymerase subunit beta (RNAP subunit beta) (EC 2.7.7.6) (RNA polymerase subunit beta) (Transcriptase subunit beta) |
| 111 | 1243 | 0.04 | B5EFV6 | *Geobacter bemidjiensis* | Bifunctional protein PutA |
| 112 | 1241 | 0.04 | A0A0B5BIF0 | *Geobacter pickeringii* | NADH-ubiquinone oxidoreductase subunit 3 |
| 113 | 1237 | 0.04 | A0A0D5NDD6 | *Geobacter sulfurreducens* | Type II secretion system protein E |
| 114 | 1236 | 0.04 | Q39QJ5 | *Geobacter metallireducens* | Sensor histidine kinase CheA associated with MCPs of classes 40H and 40+24H |
| 115 | 1236 | 0.04 | Q39RM6 | *Geobacter metallireducens* | Glycosyltransferase |
| 116 | 1234 | 0.04 | Q39QJ8 | *Geobacter metallireducens* | Flagellar biogenesis master sigma-54-dependent transcriptional response regulator |
| 117 | 1232 | 0.04 | A0A0C1TL30 | *Geobacter soli* | Flagellar basal body stator protein MotB |
| 118 | 1231 | 0.04 | | | |
| 119 | 1224 | 0.04 | A0A0B5BC86 | *Geobacter pickeringii* | Multicopper oxidase |
| 120 | 1224 | 0.04 | A0A0D5NE46 | *Geobacter sulfurreducens* | Molecular chaperone DnaJ |
| 121 | 1221 | 0.04 | Q39SI5 | *Geobacter metallireducens* | Fumarate hydratase class I (EC 4.2.1.2) |
| 122 | 1217 | 0.04 | Q39R25 | *Geobacter metallireducens* | Sensor diguanylate cyclase/phosphodiesterase, PAS, PAS and PAS domain-containing |
| 123 | 1203 | 0.04 | Q39UR9 | *Geobacter metallireducens* | Carbamoyl-phosphate synthase large chain (EC 6.3.5.5) (Carbamoyl-phosphate synthetase ammonia chain) |
| 124 | 1202 | 0.04 | A0A0B5BD43 | *Geobacter pickeringii* | ABC transporter permease |

| | | | | | |
|---|---|---|---|---|---|
| 125 | 1197 | 0.04 | A0A0B5BDG1 | *Geobacter pickeringii* | tRNA uridine 5-carboxymethylaminomethyl modification enzyme MnmG (Glucose-inhibited division protein A) |
| 126 | 1194 | 0.04 | A0A0B5BIQ7 | *Geobacter pickeringii* | Flagellar basal-body rod protein FlgF (Flagellar hook protein FlgE) |
| 127 | 1193 | 0.04 | P61679 | *Geobacter sulfurreducens* | UDP-N-acetylmuramate--L-alanine ligase (EC 6.3.2.8) (UDP-N-acetylmuramoyl-L-alanine synthetase) |
| 128 | 1192 | 0.04 | A0A0B5BCL9 | *Geobacter pickeringii* | Uncharacterized protein |
| 129 | 1191 | 0.04 | A0A0B5BH05 | *Geobacter pickeringii* | Malic enzyme (EC 1.1.1.40) |
| 130 | 1187 | 0.04 | A0A0B5BK89 | *Geobacter pickeringii* | FAD-binding protein |
| 131 | 1186 | 0.04 | A0A0B5BHV9 | *Geobacter pickeringii* | ATPase AAA |
| 132 | 1186 | 0.04 | A0A0B5BK56 | *Geobacter pickeringii* | Lon protease (EC 3.4.21.53) (ATP-dependent protease La) |
| 133 | 1186 | 0.04 | Q39WG2 | *Geobacter metallireducens* | Chromosome partition protein Smc |
| 134 | 1186 | 0.04 | A5GAY2 | *Geobacter uraniireducens* | DNA-directed RNA polymerase subunit beta' (RNAP subunit beta') (EC 2.7.7.6) (RNA polymerase subunit beta') (Transcriptase subunit beta') |
| 135 | 1183 | 0.04 | A0A0B5BIA1 | *Geobacter pickeringii* | DNA-directed DNA polymerase (EC 2.7.7.7) |
| 136 | 1178 | 0.04 | A5GF11 | *Geobacter uraniireducens* | Acetyl-CoA carboxylase carboxyltransferase subunit alpha (EC 6.3.4.14) |
| 137 | 1176 | 0.04 | Q39SJ2 | *Geobacter metallireducens* | Nitrogen fixation master sigma-54-dependent transcriptional response regulator |
| 138 | 1175 | 0.04 | A0A0B5BCZ8 | *Geobacter pickeringii* | Succinate dehydrogenase (EC 1.3.5.1) |
| 139 | 1175 | 0.04 | Q74F57 | *Geobacter sulfurreducens* | Membrane protein, major facilitator superfamily |
| 140 | 1172 | 0.04 | A0A0A8WPS5 | *Geobacter sp.* | Efflux pump membrane transporter |
| 141 | 1171 | 0.04 | Q749N5 | *Geobacter sulfurreducens* | Acetate kinase (EC 2.7.2.1) (Acetokinase) |
| 142 | 1168 | 0.04 | A0A0B5B668 | *Geobacter pickeringii* | Peptidase U32 |
| 143 | 1168 | 0.04 | Q39WR2 | *Geobacter metallireducens* | CRISPR-associated endonuclease Cas1 (EC 3.1.-.-) |
| 144 | 1167 | 0.04 | A0A0D5N400 | *Geobacter sulfurreducens* | Uncharacterized protein |
| 145 | 1164 | 0.04 | Q74GY2 | *Geobacter sulfurreducens* | ATP synthase subunit alpha (EC 3.6.3.14) (ATP synthase F1 sector subunit alpha) (F-ATPase subunit alpha) |
| 146 | 1164 | 0.04 | A0A4W3 | *Geobacter sulfurreducens* | Biosynthetic arginine decarboxylase (ADC) (EC 4.1.1.19) |
| 147 | 1164 | 0.04 | | | |
| 148 | 1163 | 0.04 | A0A0B5BBC9 | *Geobacter pickeringii* | Type I citrate synthase (EC 2.3.3.1) |
| 149 | 1162 | 0.04 | Q74FW2 | *Geobacter sulfurreducens* | Succinyl:acetate coenzyme A transferase |
| 150 | 1160 | 0.04 | Q39QU2 | *Geobacter metallireducens* | Phosphoenolpyruvate carboxykinase (ATP) (PCK) (PEP carboxykinase) (PEPCK) (EC 4.1.1.49) |

| | | | | | |
|---|---|---|---|---|---|
| 151 | 1153 | 0.04 | A0A0D5N7I1 | *Geobacter sulfurreducens* | NADH-quinone oxidoreductase subunit N (EC 1.6.5.11) (NADH dehydrogenase I subunit N) (NDH-1 subunit N) |
| 152 | 1150 | 0.04 | A0A0B5B9H5 | *Geobacter pickeringii* | Uncharacterized protein |
| 153 | 1150 | 0.04 | Q748I6 | *Geobacter sulfurreducens* | Metalloprotease domain protein, M6 family |
| 154 | 1149 | 0.04 | Q74CR5 | *Geobacter sulfurreducens* | Serine hydroxymethyltransferase (SHMT) (Serine methylase) (EC 2.1.2.1) |
| 155 | 1149 | 0.04 | P61667 | *Geobacter sulfurreducens* | DNA mismatch repair protein MutS |
| 156 | 1145 | 0.04 | Q39U54 | *Geobacter metallireducens* | Sensor histidine kinase cyclic nucleotide phosphodiesterase, GAF, GAF and HD-GYP-related domain-containing |
| 157 | 1144 | 0.04 | A5GDX4 | *Geobacter uraniireducens* | DNA gyrase subunit B (EC 5.99.1.3) |
| 158 | 1143 | 0.04 | Q747G2 | *Geobacter sulfurreducens* | Outer membrane channel OmpJ |
| 159 | 1140 | 0.04 | Q39QP7 | *Geobacter metallireducens* | Lon protease (EC 3.4.21.53) (ATP-dependent protease La) |
| 160 | 1138 | 0.04 | A0A0D5NCW1 | *Geobacter sulfurreducens* | Histidine kinase |
| 161 | 1138 | 0.04 | Q39ZQ2 | *Geobacter metallireducens* | Uncharacterized protein |
| 162 | 1136 | 0.04 | Q74GY0 | *Geobacter sulfurreducens* | ATP synthase subunit beta (EC 3.6.3.14) (ATP synthase F1 sector subunit beta) (F-ATPase subunit beta) |
| 163 | 1135 | 0.04 | A0A0B5BE38 | *Geobacter pickeringii* | Transcription-repair-coupling factor (TRCF) (EC 3.6.4.-) |
| 164 | 1128 | 0.04 | B3EAX2 | *Geobacter lovleyi* | Efflux transporter, RND family, MFP subunit |
| 165 | 1127 | 0.04 | A0A0D5N6Q1 | *Geobacter sulfurreducens* | FeS-binding protein |
| 166 | 1126 | 0.04 | E1QK39 | *Desulfarculus baarsii* | Helicase domain protein |
| 167 | 1126 | 0.04 | Q39PS2 | *Geobacter metallireducens* | Transcription-repair-coupling factor (TRCF) (EC 3.6.4.-) |
| 168 | 1123 | 0.04 | Q39X91 | *Geobacter metallireducens* | DNA topoisomerase 1 (EC 5.99.1.2) (DNA topoisomerase I) |
| 169 | 1122 | 0.04 | Q74GG6 | *Geobacter sulfurreducens* | Chromosomal replication initiator protein DnaA |
| 170 | 1121 | 0.04 | A0A0D5NB12 | *Geobacter sulfurreducens* | Aldehyde dehydrogenase |
| 171 | 1116 | 0.04 | A5GCN0 | *Geobacter uraniireducens* | Short chain amide porin |
| 172 | 1114 | 0.04 | Q39RQ5 | *Geobacter metallireducens* | Chaperone protein ClpB |
| 173 | 1111 | 0.04 | Q39X99 | *Geobacter metallireducens* | UDP-glucuronate 4-dehydrogenase, decarboxylating |
| 174 | 1109 | 0.04 | Q39W73 | *Geobacter metallireducens* | 2-isopropylmalate synthase (EC 2.3.3.13) (Alpha-IPM synthase) (Alpha-isopropylmalate synthase) |
| 175 | 1108 | 0.04 | A0LQ17 | *Syntrophobacter fumaroxidans* | Helicase domain protein |
| 176 | 1107 | 0.04 | | | |

| | | | | | |
|---|---|---|---|---|---|
| 177 | 1103 | 0.04 | B9M1Q4 | *Geobacter daltonii* | Biotin-dependent acyl-CoA carboxylase, biotin carboxylase subunit |
| 178 | 1102 | 0.04 | Q74C83 | *Geobacter sulfurreducens* | ATP-dependent Clp protease ATP-binding subunit ClpX |
| 179 | 1101 | 0.04 | A0A0D5N7N0 | *Geobacter sulfurreducens* | Glutamate synthase |
| 180 | 1100 | 0.04 | A5G8I3 | *Geobacter uraniireducens* | Chaperone protein ClpB |
| 181 | 1099 | 0.04 | A0A1F9PUK4 | *Desulfuromonadales bacterium* | Methyltransferase (EC 2.1.1.-) |
| 182 | 1099 | 0.04 | B9M5J7 | *Geobacter daltonii* | CRISPR-associated protein Csd2 |
| 183 | 1098 | 0.04 | A0A1G0LEY2 | *Geobacteraceae bacterium* | Uncharacterized protein |
| 184 | 1097 | 0.04 | A0A0C1QTJ3 | *Geobacter soli* | Hemolysin D |
| 185 | 1097 | 0.04 | Q74D18 | *Geobacter sulfurreducens* | ABC transporter, ATP-binding protein |
| 186 | 1092 | 0.04 | A0A0D5N435 | *Geobacter sulfurreducens* | Multidrug transporter |
| 187 | 1088 | 0.04 | A0A0A8WNZ4 | *Geobacter sp.* | DNA gyrase subunit A (EC 5.99.1.3) |
| 188 | 1087 | 0.04 | B3EA98 | *Geobacter lovleyi* | Transcriptional regulator, XRE family |
| 189 | 1085 | 0.04 | A0A0B5BCH1 | *Geobacter pickeringii* | Hydrogenase 2 large subunit (EC 1.12.99.6) |
| 190 | 1081 | 0.04 | A0A0B5BB89 | *Geobacter pickeringii* | Chromosome partition protein Smc |
| 191 | 1081 | 0.04 | A0A0D5N952 | *Geobacter sulfurreducens* | Flagellar motor switch protein FliG |
| 192 | 1077 | 0.04 | A1AN79 | *Pelobacter propionicus* | Adenosylmethionine-8-amino-7-oxononanoate aminotransferase (EC 2.6.1.62) (7,8-diamino-pelargonic acid aminotransferase) (DAPA AT) (DAPA aminotransferase) (7,8-diaminononanoate synthase) (DANS) (Diaminopelargonic acid synthase) |
| 193 | 1073 | 0.04 | Q39QK7 | *Geobacter metallireducens* | Outer membrane channel OmpJ |
| 194 | 1073 | 0.04 | Q39YR4 | *Geobacter metallireducens* | FtsK/SpoIIIE domain protein |
| 195 | 1072 | 0.04 | | | |
| 196 | 1064 | 0.04 | Q39RM7 | *Geobacter metallireducens* | Glycosyltransferase |
| 197 | 1064 | 0.04 | | | |
| 198 | 1061 | 0.04 | A0A0D5N8X4 | *Geobacter sulfurreducens* | Flagellar M-ring protein |
| 199 | 1060 | 0.04 | A0A0C1TL33 | *Geobacter soli* | Uncharacterized protein |
| 200 | 1057 | 0.04 | Q39X64 | *Geobacter metallireducens* | Cell shape-determining protein MreB |
| 201 | 1051 | 0.04 | Q74EQ2 | *Geobacter sulfurreducens* | Aldehyde:ferredoxin oxidoreductase, tungsten-containing |
| 202 | 1049 | 0.04 | A0A0D5N7M4 | *Geobacter sulfurreducens* | Chemotaxis protein |

| | | | | | |
|---|---|---|---|---|---|
| 203 | 1049 | 0.04 | A5G9A7 | *Geobacter uraniireducens* | NADH dehydrogenase subunit M (EC 1.6.5.3) |
| 204 | 1049 | 0.04 | Q39XG6 | *Geobacter metallireducens* | Pyruvate carboxylase (EC 6.4.1.1) |
| 205 | 1044 | 0.04 | A0A0B5B7G6 | *Geobacter pickeringii* | Cytochrome C |
| 206 | 1041 | 0.04 | P60789 | *Geobacter sulfurreducens* | Elongation factor 4 (EF-4) (EC 3.6.5.n1) (Ribosomal back-translocase LepA) |
| 207 | 1040 | 0.04 | A0R7X1 | *Pelobacter propionicus* | Type III restriction enzyme, res subunit |
| 208 | 1036 | 0.04 | Q74EP9 | *Geobacter sulfurreducens* | DNA-binding ATPase Uup |
| 209 | 1035 | 0.04 | A0A0B5B8F2 | *Geobacter pickeringii* | Sodium transporter |
| 210 | 1032 | 0.04 | Q39Z79 | *Geobacter metallireducens* | Twitching motility pilus retraction ATPase |
| 211 | 1032 | 0.04 | Q74BT1 | *Geobacter sulfurreducens* | Repeat-containing protein |
| 212 | 1030 | 0.04 | Q39YH3 | *Geobacter metallireducens* | UDP-N-acetylglucosamine 4,6-dehydratase and UDP-2-acetamido-2,6-dideoxy-alpha-D-xylo-4-hexulose 5-epimerase |
| 213 | 1029 | 0.04 | A0A0B5BDE7 | *Geobacter pickeringii* | Fis family transcriptional regulator |
| 214 | 1026 | 0.04 | Q39RI3 | *Geobacter metallireducens* | Sensor histidine kinase, HAMP domain-containing, 2 heme-binding sites |
| 215 | 1018 | 0.04 | A0A0B5BGI9 | *Geobacter pickeringii* | Lon protease (EC 3.4.21.53) (ATP-dependent protease La) |
| 216 | 1017 | 0.04 | A0A0B5BKE7 | *Geobacter pickeringii* | NADH-quinone oxidoreductase subunit N (EC 1.6.5.11) (NADH dehydrogenase I subunit N) (NDH-1 subunit N) |
| 217 | 1016 | 0.04 | Q74GT3 | *Geobacter sulfurreducens* | LL-diaminopimelate aminotransferase (DAP-AT) (DAP-aminotransferase) (LL-DAP-aminotransferase) (EC 2.6.1.83) |
| 218 | 1015 | 0.04 | A5GAC6 | *Geobacter uraniireducens* | FMN-dependent NADH-azoreductase (EC 1.7.-.-) (Azo-dye reductase) (FMN-dependent NADH-azo compound oxidoreductase) |
| 219 | 1014 | 0.04 | A0A0B5BGV8 | *Geobacter pickeringii* | Zinc protease |
| 220 | 1014 | 0.04 | Q74C70 | *Geobacter sulfurreducens* | Phosphoglucosamine mutase (EC 5.4.2.10) |
| 221 | 1013 | 0.04 | Q39W76 | *Geobacter metallireducens* | Ketol-acid reductoisomerase (NADP(+)) (KARI) (EC 1.1.1.86) (Acetohydroxy-acid isomeroreductase) (AHIR) (Alpha-keto-beta-hydroxylacyl reductoisomerase) (Ketol-acid reductoisomerase type 1) (Ketol-acid reductoisomerase type I) |
| 222 | 1011 | 0.03 | A0A0A8WRD3 | *Geobacter sp.* | Putative ABC transporter ATP-binding protein YkpA |
| 223 | 1009 | 0.03 | A0A0B5BDW0 | *Geobacter pickeringii* | Leucine--tRNA ligase (EC 6.1.1.4) (Leucyl-tRNA synthetase) (LeuRS) |
| 224 | 1008 | 0.03 | A0A1T4RGQ7 | *Geobacter thiogenes* | NAD+---dinitrogen-reductase ADP-D-ribosyltransferase |
| 225 | 1007 | 0.03 | A0A0B5BGY7 | *Geobacter pickeringii* | Phosphoglucomutase |
| 226 | 1006 | 0.03 | Q74DL9 | *Geobacter sulfurreducens* | Transposase of ISGsu2, IS5 family |

| | | | | | |
|---|---|---|---|---|---|
| 227 | 1005 | 0.03 | A0A0D5NAW4 | *Geobacter sulfurreducens* | Cation transporter |
| 228 | 1001 | 0.03 | A0A0B5BG42 | *Geobacter pickeringii* | Vitamin B12-dependent ribonucleotide reductase (EC 1.17.4.1) |
| 229 | 1000 | 0.03 | Q39QB3 | *Geobacter metallireducens* | NADH dehydrogenase I, G subunit |
| 230 | 998 | 0.03 | A0A0D5N949 | *Geobacter sulfurreducens* | Response regulator |
| 231 | 995 | 0.03 | A0A0B5BB11 | *Geobacter pickeringii* | Pyruvate carboxylase (EC 6.4.1.1) |
| 232 | 994 | 0.03 | Q39QA3 | *Geobacter metallireducens* | ATP synthase subunit a (ATP synthase F0 sector subunit a) (F-ATPase subunit 6) |
| 233 | 990 | 0.03 | A0A0B5BCB2 | *Geobacter pickeringii* | Nitrogen fixation protein NifE |
| 234 | 984 | 0.03 | A0A0B5BBZ4 | *Geobacter pickeringii* | DNA polymerase I (EC 2.7.7.7) |
| 235 | 983 | 0.03 | A0A0B5BE21 | *Geobacter pickeringii* | Histidine kinase |
| 236 | 983 | 0.03 | Q39QY3 | *Geobacter metallireducens* | Efflux pump, RND family, inner membrane protein |
| 237 | 982 | 0.03 | Q39ZL2 | *Geobacter metallireducens* | TrkA domain protein |
| 238 | 980 | 0.03 | A0A1F9PXI1 | *Desulfuromonadales bacterium* | Cytosine-specific methyltransferase (EC 2.1.1.37) |
| 239 | 979 | 0.03 | Q74AJ5 | *Geobacter sulfurreducens* | Glycoside hydrolase, family 57, DUF3536 domain-containing |
| 240 | 978 | 0.03 | | | |
| 241 | 977 | 0.03 | Q39UA2 | *Geobacter metallireducens* | Adenylosuccinate lyase (ASL) (EC 4.3.2.2) (Adenylosuccinase) |
| 242 | 971 | 0.03 | A0A0B5B8W1 | *Geobacter pickeringii* | ATP-dependent zinc metalloprotease FtsH (EC 3.4.24.-) |
| 243 | 971 | 0.03 | A1AM25 | *Pelobacter propionicus* | Squalene-hopene cyclase |
| 244 | 970 | 0.03 | A0A0B5B7K6 | *Geobacter pickeringii* | Cytochrome C biogenesis protein ResC |
| 245 | 970 | 0.03 | A0A1T4RHG8 | *Geobacter thiogenes* | Iron complex outermembrane recepter protein |
| 246 | 968 | 0.03 | A0A0C1TQJ7 | *Geobacter soli* | Uncharacterized protein |
| 247 | 967 | 0.03 | B3EAB9 | *Geobacter lovleyi* | 4Fe-4S ferredoxin, iron-sulfur binding domain protein |
| 248 | 966 | 0.03 | A0A0D5N5Y8 | *Geobacter sulfurreducens* | Cell division protein FtsA |
| 249 | 965 | 0.03 | Q39ZI6 | *Geobacter metallireducens* | DNA helicase (EC 3.6.4.12) |
| 250 | 963 | 0.03 | Q746T2 | *Geobacter sulfurreducens* | NADH-quinone oxidoreductase subunit H 2 (EC 1.6.5.11) (NADH dehydrogenase I subunit H 2) (NDH-1 subunit H 2) |
| 251 | 960 | 0.03 | A0A0B5BDP4 | *Geobacter pickeringii* | Polya polymerase |
| 252 | 959 | 0.03 | A0A0B5B8J3 | *Geobacter pickeringii* | Rod shape-determining protein RodA |

| | | | | | |
|---|---|---|---|---|---|
| 253 | 953 | 0.03 | Q74BT7 | *Geobacter sulfurreducens* | Sensor histidine kinase cyclic nucleotide phosphodiesterase, GAF, GAF and HD-GYP-related domain-containing |
| 254 | 953 | 0.03 | | | |
| 255 | 949 | 0.03 | P61946 | *Geobacter sulfurreducens* | S-adenosylmethionine synthase (AdoMet synthase) (EC 2.5.1.6) (MAT) (Methionine adenosyltransferase) |
| 256 | 949 | 0.03 | Q74AF7 | *Geobacter sulfurreducens* | Methyl viologen-reducing hydrogenase, large subunit |
| 257 | 948 | 0.03 | A0A0D5N8K2 | *Geobacter sulfurreducens* | RND transporter |
| 258 | 948 | 0.03 | Q39QT7 | *Geobacter metallireducens* | Cytochrome c |
| 259 | 947 | 0.03 | Q39Y82 | *Geobacter metallireducens* | 5-methyltetrahydrofolate--homocysteine S-methyltransferase, cobalamin-dependent |
| 260 | 941 | 0.03 | A1ANK6 | *Pelobacter propionicus* | Peptidase C1A, papain |
| 261 | 938 | 0.03 | A0A1T4RTR1 | *Geobacter thiogenes* | Iron complex outermembrane recepter protein |
| 262 | 938 | 0.03 | Q39VY0 | *Geobacter metallireducens* | Malate dehydrogenase (EC 1.1.1.37) |
| 263 | 933 | 0.03 | A0A0B5BC52 | *Geobacter pickeringii* | General secretion pathway protein GspD |
| 264 | 932 | 0.03 | A0A0B5BH19 | *Geobacter pickeringii* | Uncharacterized protein |
| 265 | 932 | 0.03 | B5EHY9 | *Geobacter bemidjiensis* | Uncharacterized protein |
| 266 | 931 | 0.03 | A0A0D5NAV3 | *Geobacter sulfurreducens* | Peptidase S41 |
| 267 | 931 | 0.03 | B9LZR8 | *Geobacter daltonii* | Succinyl:acetate coenzyme A transferase |
| 268 | 930 | 0.03 | A0A0A8WV23 | *Geobacter sp.* | Protein translocase subunit SecD |
| 269 | 930 | 0.03 | Q74AT0 | *Geobacter sulfurreducens* | Lysine--tRNA ligase (EC 6.1.1.6) (Lysyl-tRNA synthetase) (LysRS) |
| 270 | 928 | 0.03 | A0A1G0LYB2 | *Geobacteraceae bacterium* | Uncharacterized protein |
| 271 | 925 | 0.03 | A0A0B5B5Z5 | *Geobacter pickeringii* | DNA polymerase III subunit beta (EC 2.7.7.7) |
| 272 | 925 | 0.03 | A0A0B5BIX5 | *Geobacter pickeringii* | DNA polymerase II |
| 273 | 924 | 0.03 | Q748B3 | *Geobacter sulfurreducens* | UDP-N-acetylglucosamine 1-carboxyvinyltransferase (EC 2.5.1.7) (Enoylpyruvate transferase) (UDP-N-acetylglucosamine enolpyruvyl transferase) (EPT) |
| 274 | 923 | 0.03 | A5GC25 | *Geobacter uraniireducens* | Propionyl-CoA carboxylase (EC 6.4.1.3) |
| 275 | 923 | 0.03 | Q74F32 | *Geobacter sulfurreducens* | Periplasmically oriented, membrane-bound formate dehydrogenase, major subunit, selenocysteine-containing |
| 276 | 921 | 0.03 | A0A0B5B822 | *Geobacter pickeringii* | Formate dehydrogenase |
| 277 | 921 | 0.03 | A0A0B5BHJ9 | *Geobacter pickeringii* | Chaperone protein HtpG (Heat shock protein HtpG) (High temperature protein G) |

| | | | | | |
|---|---|---|---|---|---|
| 278 | 921 | 0.03 | | | |
| 279 | 920 | 0.03 | A0A0B5BGU8 | *Geobacter pickeringii* | Chemotaxis protein |
| 280 | 919 | 0.03 | A0A0B5B7W4 | *Geobacter pickeringii* | Transporter |
| 281 | 919 | 0.03 | A0A0B5BFT0 | *Geobacter pickeringii* | Phosphoribosylformylglycinamidine synthase subunit PurL (FGAM synthase) (EC 6.3.5.3) (Formylglycinamide ribonucleotide amidotransferase subunit II) (FGAR amidotransferase II) (FGAR-AT II) (Glutamine amidotransferase PurL) (Phosphoribosylformylglycinamidine synthase subunit II) |
| 282 | 918 | 0.03 | A0A0B5BJV3 | *Geobacter pickeringii* | ABC transporter substrate-binding protein |
| 283 | 917 | 0.03 | A0A0B5BFH3 | *Geobacter pickeringii* | Polyribonucleotide nucleotidyltransferase (EC 2.7.7.8) (Polynucleotide phosphorylase) (PNPase) |
| 284 | 917 | 0.03 | E1QK40 | *Desulfarculus baarsii* | DNA-cytosine methyltransferase |
| 285 | 917 | 0.03 | Q74D56 | *Geobacter sulfurreducens* | Aspartate--tRNA(Asp/Asn) ligase (EC 6.1.1.23) (Aspartyl-tRNA synthetase) (AspRS) (Non-discriminating aspartyl-tRNA synthetase) (ND-AspRS) |
| 286 | 912 | 0.03 | B0L930 | *uncultured Geobacter sp.* | Tuf (Fragment) |
| 287 | 911 | 0.03 | A0A0D5N419 | *Geobacter sulfurreducens* | Uncharacterized protein |
| 288 | 911 | 0.03 | Q74E27 | *Geobacter sulfurreducens* | Ribonuclease Y (RNase Y) (EC 3.1.-.-) |
| 289 | 909 | 0.03 | Q74GU4 | *Geobacter sulfurreducens* | Acetylglutamate kinase (EC 2.7.2.8) (N-acetyl-L-glutamate 5-phosphotransferase) (NAG kinase) (NAGK) |
| 290 | 906 | 0.03 | Q747L0 | *Geobacter sulfurreducens* | Glycoside hydrolase, family 57 |
| 291 | 905 | 0.03 | A0A0B5BGX4 | *Geobacter pickeringii* | Stage II sporulation protein E |
| 292 | 903 | 0.03 | A5GE75 | *Geobacter uraniireducens* | Heavy metal translocating P-type ATPase |
| 293 | 903 | 0.03 | | | |
| 294 | 902 | 0.03 | A0A0D5N645 | *Geobacter sulfurreducens* | Penicillin-binding protein |
| 295 | 900 | 0.03 | A0A0B5B6N9 | *Geobacter pickeringii* | Alanine--tRNA ligase (EC 6.1.1.7) (Alanyl-tRNA synthetase) (AlaRS) |
| 296 | 899 | 0.03 | A0A1G0N1F9 | *Geobacteraceae bacterium* | Tellurium resistance protein TerC |
| 297 | 899 | 0.03 | Q749D7 | *Geobacter sulfurreducens* | Nitrogenase molybdenum-iron cofactor biosynthesis protein NifEN |
| 298 | 899 | 0.03 | Q748Y3 | *Geobacter sulfurreducens* | 50S ribosomal protein L1 |
| 299 | 897 | 0.03 | Q746S4 | *Geobacter sulfurreducens* | NADH-quinone oxidoreductase subunit B/C/D (EC 1.6.5.11) (NADH dehydrogenase I subunit B/C/D) (NDH-1 subunit B/C/D) |
| 300 | 895 | 0.03 | A0A1G0MT57 | *Geobacteraceae bacterium* | Uncharacterized protein |
| 301 | 892 | 0.03 | Q39Y18 | *Geobacter metallireducens* | Transcription termination/antitermination protein NusG |

| | | | | | |
|---|---|---|---|---|---|
| 302 | 891 | 0.03 | A0A0B5B942 | *Geobacter pickeringii* | Bifunctional uridylyltransferase/uridylyl-removing enzyme (UTase/UR) (Bifunctional [protein-PII] modification enzyme) (Bifunctional nitrogen sensor protein) [Includes: [Protein-PII]-UMP uridylyl-removing enzyme (UR) (EC 3.1.4.-); [Protein-PII] uridylyltransferase (PII uridylyltransferase) (UTase) (EC 2.7.7.59)] |
| 303 | 891 | 0.03 | A0A0D5N4Y8 | *Geobacter sulfurreducens* | Pilus assembly protein PilB |
| 304 | 890 | 0.03 | A5GDX5 | *Geobacter uraniireducens* | DNA gyrase subunit A (EC 5.99.1.3) |
| 305 | 890 | 0.03 | Q39YP6 | *Geobacter metallireducens* | Histidinol-phosphate aminotransferase (EC 2.6.1.9) (Imidazole acetol-phosphate transaminase) |
| 306 | 890 | 0.03 | Q74H61 | *Geobacter sulfurreducens* | Heat-inducible transcription repressor HrcA |
| 307 | 889 | 0.03 | | | |
| 308 | 888 | 0.03 | A0A0B5BBF5 | *Geobacter pickeringii* | Twin-arginine translocation pathway signal protein |
| 309 | 888 | 0.03 | A0A0B5BJ17 | *Geobacter pickeringii* | UvrABC system protein C (Protein UvrC) (Excinuclease ABC subunit C) |
| 310 | 888 | 0.03 | Q74AR7 | *Geobacter sulfurreducens* | Membrane-associated metal-dependent phosphohydrolase, HDc domain-containing |
| 311 | 886 | 0.03 | A0A0B5B792 | *Geobacter pickeringii* | Flagellar biosynthesis protein FlhF |
| 312 | 886 | 0.03 | Q747D6 | *Geobacter sulfurreducens* | Pyruvate kinase (EC 2.7.1.40) |
| 313 | 884 | 0.03 | Q74D04 | *Geobacter sulfurreducens* | Threonine--tRNA ligase (EC 6.1.1.3) (Threonyl-tRNA synthetase) (ThrRS) |
| 314 | 883 | 0.03 | A0A0B5BFC0 | *Geobacter pickeringii* | Ni/Fe hydrogenase |
| 315 | 882 | 0.03 | A0A0B5BJA2 | *Geobacter pickeringii* | RND transporter |
| 316 | 879 | 0.03 | A0A0B5BI14 | *Geobacter pickeringii* | GTPase Der (GTP-binding protein EngA) |
| 317 | 878 | 0.03 | A0A0A8WYA5 | *Geobacter sp.* | DNA-directed RNA polymerase subunit beta' (RNAP subunit beta') (EC 2.7.7.6) (RNA polymerase subunit beta') (Transcriptase subunit beta') |
| 318 | 876 | 0.03 | | | |
| 319 | 875 | 0.03 | B9M149 | *Geobacter daltonii* | Translation-regulating membrane GTPase TypA |
| 320 | 875 | 0.03 | B9M4C2 | *Geobacter daltonii* | Squalene cyclase |
| 321 | 875 | 0.03 | Q39RM2 | *Geobacter metallireducens* | Glycosyltransferase |
| 322 | 875 | 0.03 | Q39UG5 | *Geobacter metallireducens* | Aspartokinase (EC 2.7.2.4) |
| 323 | 872 | 0.03 | A0A0D5N7Y6 | *Geobacter sulfurreducens* | Chemotaxis protein CheA |
| 324 | 870 | 0.03 | A5G3L3 | *Geobacter uraniireducens* | Isocitrate dehydrogenase, NADP-dependent (EC 1.1.1.42) |
| 325 | 870 | 0.03 | Q39WZ1 | *Geobacter metallireducens* | Nitrogen fixation protein NifU |

| | | | | | |
|---|---|---|---|---|---|
| 326 | 868 | 0.03 | A0A0B5B8X9 | *Geobacter pickeringii* | Phosphoenolpyruvate-protein phosphotransferase (EC 2.7.3.9) (Phosphotransferase system, enzyme I) |
| 327 | 868 | 0.03 | Q39UH4 | *Geobacter metallireducens* | Lon protease (EC 3.4.21.53) (ATP-dependent protease La) |
| 328 | 868 | 0.03 | Q39VA8 | *Geobacter metallireducens* | Transcription termination/antitermination protein NusA |
| 329 | 866 | 0.03 | A0A0B5BJM3 | *Geobacter pickeringii* | NADH-ubiquinone oxidoreductase subunit 3 |
| 330 | 866 | 0.03 | Q39VY8 | *Geobacter metallireducens* | Glutamine synthetase (EC 6.3.1.2) |
| 331 | 865 | 0.03 | A0A0B5B6P6 | *Geobacter pickeringii* | Histidine kinase |
| 332 | 865 | 0.03 | A0A0D5N8X6 | *Geobacter sulfurreducens* | Chemotaxis protein |
| 333 | 864 | 0.03 | A0A0B5BB86 | *Geobacter pickeringii* | Uncharacterized protein |
| 334 | 864 | 0.03 | A0A0D5N2Y9 | *Geobacter sulfurreducens* | Diguanylate cyclase |
| 335 | 864 | 0.03 | Q39Q79 | *Geobacter metallireducens* | Winged-helix transcriptional response regulator |
| 336 | 864 | 0.03 | Q39ZP3 | *Geobacter metallireducens* | VWFA superfamily protein |
| 337 | 862 | 0.03 | A0A0B5BIU4 | *Geobacter pickeringii* | Probable glycine dehydrogenase (decarboxylating) subunit 1 (EC 1.4.4.2) (Glycine cleavage system P-protein subunit 1) (Glycine decarboxylase subunit 1) (Glycine dehydrogenase (aminomethyl-transferring) subunit 1) |
| 338 | 859 | 0.03 | Q74GF2 | *Geobacter sulfurreducens* | Metal-dependent phosphohydrolase, HDOD domain-containing |
| 339 | 858 | 0.03 | A0A0D5N3S3 | *Geobacter sulfurreducens* | Chemotaxis protein CheY |
| 340 | 858 | 0.03 | Q39RK5 | *Geobacter metallireducens* | ResB-like family cytochrome c |
| 341 | 855 | 0.03 | A0A0B5BIW8 | *Geobacter pickeringii* | Aminotransferase |
| 342 | 854 | 0.03 | B3EAZ5 | *Geobacter lovleyi* | TraG family protein |
| 343 | 853 | 0.03 | Q74D76 | *Geobacter sulfurreducens* | Histidine kinase (EC 2.7.13.3) |
| 344 | 848 | 0.03 | A0A1G1FB69 | *Nitrospirae bacterium* | Uncharacterized protein |
| 345 | 848 | 0.03 | Q39VR4 | *Geobacter metallireducens* | Bifunctional uridylyltransferase/uridylyl-removing enzyme (UTase/UR) (Bifunctional [protein-PII] modification enzyme) (Bifunctional nitrogen sensor protein) [Includes: [Protein-PII]-UMP uridylyl-removing enzyme (UR) (EC 3.1.4.-); [Protein-PII] uridylyltransferase (PII uridylyltransferase) (UTase) (EC 2.7.7.59)] |
| 346 | 846 | 0.03 | B3E8C0 | *Geobacter lovleyi* | Iron-containing alcohol dehydrogenase |
| 347 | 846 | 0.03 | Q74D19 | *Geobacter sulfurreducens* | Uncharacterized protein |
| 348 | 845 | 0.03 | A0A1G0N2A8 | *Geobacteraceae bacterium* | Thiol:disulfide interchange protein |
| 349 | 845 | 0.03 | A5G8W9 | *Geobacter uraniireducens* | Flagellar hook-associated protein 2 (HAP2) (Flagellar cap protein) |

| | | | | | |
|---|---|---|---|---|---|
| 350 | 843 | 0.03 | Q39VA6 | *Geobacter metallireducens* | Translation initiation factor IF-2 |
| 351 | 843 | 0.03 | Q30VS6 | *Desulfovibrio alaskensis* | Site-specific DNA-methyltransferase (Adenine-specific) (EC 2.1.1.72) |
| 352 | 842 | 0.03 | A0A0B5BHW3 | *Geobacter pickeringii* | Ammonium transporter |
| 353 | 841 | 0.03 | A0A0D5N2S0 | *Geobacter sulfurreducens* | Uncharacterized protein |
| 354 | 841 | 0.03 | A0A1F9P4Q4 | *Desulfuromonadales bacterium* | Uncharacterized protein |
| 355 | 841 | 0.03 | Q39XC1 | *Geobacter metallireducens* | PilB/PulE/GspE family ATPase |
| 356 | 841 | 0.03 | Q39Z54 | *Geobacter metallireducens* | Membrane protein, putative |
| 357 | 840 | 0.03 | A0A0B5BCR8 | *Geobacter pickeringii* | Alpha-1,4 glucan phosphorylase (EC 2.4.1.1) |
| 358 | 839 | 0.03 | Q39XL2 | *Geobacter metallireducens* | Phosphoenolpyruvate synthase (PEP synthase) (EC 2.7.9.2) (Pyruvate, water dikinase) |
| 359 | 838 | 0.03 | A0A0B5BK21 | *Geobacter pickeringii* | Inner membrane cytochrome H |
| 360 | 837 | 0.03 | A0A0D5NDV9 | *Geobacter sulfurreducens* | HD family phosphohydrolase |
| 361 | 837 | 0.03 | Q74BQ5 | *Geobacter sulfurreducens* | Undecaprenyl-phosphate glycosylphosphotransferase |
| 362 | 836 | 0.03 | Q39X31 | *Geobacter metallireducens* | Protein translocase subunit SecA |
| 363 | 835 | 0.03 | Q747G4 | *Geobacter sulfurreducens* | (R)-methylmalonyl-CoA mutase, isobutyryl-CoA mutase-like catalytic subunit |
| 364 | 834 | 0.03 | A0A0B5BIH3 | *Geobacter pickeringii* | Uncharacterized protein |
| 365 | 834 | 0.03 | Q74A32 | *Geobacter sulfurreducens* | PppGpp 5'-phosphohydrolase and exopolyphosphatase, HD domain-containing |
| 366 | 833 | 0.03 | A0A0B5B672 | *Geobacter pickeringii* | FeS-binding protein |
| 367 | 833 | 0.03 | C6DZ95 | *Geobacter sp.* | Lytic transglycosylase catalytic |
| 368 | 829 | 0.03 | B9M5J8 | *Geobacter daltonii* | CRISPR-associated protein Csd1 |
| 369 | 828 | 0.03 | Q39X71 | *Geobacter metallireducens* | Uncharacterized protein |
| 370 | 828 | 0.03 | Q74FM8 | *Geobacter sulfurreducens* | Glycine--tRNA ligase alpha subunit (EC 6.1.1.14) (Glycyl-tRNA synthetase alpha subunit) (GlyRS) |
| 371 | 827 | 0.03 | P60401 | *Geobacter sulfurreducens* | 50S ribosomal protein L2 |
| 372 | 826 | 0.03 | A0A0B5BBV1 | *Geobacter pickeringii* | Cytochrome C |
| 373 | 825 | 0.03 | A0A0B5B7V2 | *Geobacter pickeringii* | Ribosome-binding ATPase YchF |
| 374 | 824 | 0.03 | P61523 | *Geobacter sulfurreducens* | Argininosuccinate synthase (EC 6.3.4.5) (Citrulline--aspartate ligase) |
| 375 | 823 | 0.03 | Q74D63 | *Geobacter sulfurreducens* | Uncharacterized protein |
| 376 | 821 | 0.03 | A0A0B5BD26 | *Geobacter pickeringii* | CDP-diacylglycerol--serine O-phosphatidyltransferase |

| | | | | | |
|---|---|---|---|---|---|
| 377 | 821 | 0.03 | A0A0B5BDG8 | *Geobacter pickeringii* | ABC transporter ATP-binding protein |
| 378 | 820 | 0.03 | Q748T0 | *Geobacter sulfurreducens* | 5-methyltetrahydrofolate--homocysteine S-methyltransferase, cobalamin-dependent |
| 379 | 815 | 0.03 | A0A0C1TS26 | *Geobacter soli* | L-aspartate oxidase (EC 1.4.3.16) |
| 380 | 815 | 0.03 | Q748X2 | *Geobacter sulfurreducens* | 3-isopropylmalate dehydrogenase (EC 1.1.1.85) (3-IPM-DH) (Beta-IPM dehydrogenase) (IMDH) |
| 381 | 814 | 0.03 | A0A0B5BEL7 | *Geobacter pickeringii* | Cytochrome C |
| 382 | 814 | 0.03 | A0A0B5BK07 | *Geobacter pickeringii* | Histidine kinase |
| 383 | 814 | 0.03 | A0A0D5N8L0 | *Geobacter sulfurreducens* | ATP-dependent protease |
| 384 | 814 | 0.03 | A5G9T7 | *Geobacter uraniireducens* | Peptide chain release factor 3 (RF-3) |
| 385 | 813 | 0.03 | A0A174BLS6 | *Mitsuokella jalaludinii* | DNA topoisomerase 3 (EC 5.99.1.2) |
| 386 | 813 | 0.03 | Q39PY9 | *Geobacter metallireducens* | Histidine kinase (EC 2.7.13.3) |
| 387 | 811 | 0.03 | A0A0A8WQH6 | *Geobacter sp.* | Uncharacterized protein |
| 388 | 810 | 0.03 | B9M7P3 | *Geobacter daltonii* | UvrABC system protein A (UvrA protein) (Excinuclease ABC subunit A) |
| 389 | 807 | 0.03 | A0A0D5N5H4 | *Geobacter sulfurreducens* | Cell division protein FtsK |
| 390 | 805 | 0.03 | A0A0B5B9N3 | *Geobacter pickeringii* | 3-oxoacyl-[acyl-carrier-protein] synthase 2 (EC 2.3.1.179) |
| 391 | 805 | 0.03 | A0A0D5N8I2 | *Geobacter sulfurreducens* | Histidine kinase |
| 392 | 803 | 0.03 | B3EA94 | *Geobacter lovleyi* | Uncharacterized protein |
| 393 | 803 | 0.03 | Q748A7 | *Geobacter sulfurreducens* | Transcription termination factor Rho (EC 3.6.4.-) (ATP-dependent helicase Rho) |
| 394 | 803 | 0.03 | Q74CQ3 | *Geobacter sulfurreducens* | Uncharacterized protein |
| 395 | 802 | 0.03 | A0A0D5N4L7 | *Geobacter sulfurreducens* | Outer membrane protein assembly factor BamA |
| 396 | 801 | 0.03 | A0A0B5BBX6 | *Geobacter pickeringii* | Ribonuclease E (RNase E) (EC 3.1.26.12) |
| 397 | 800 | 0.03 | A0A0B5B9Q9 | *Geobacter pickeringii* | Glycosyl transferase |
| 398 | 800 | 0.03 | A0A0S8I0Q8 | *bacterium SM23_31* | Uncharacterized protein |
| 399 | 799 | 0.03 | P60859 | *Geobacter sulfurreducens* | Histidinol dehydrogenase (HDH) (EC 1.1.1.23) |
| 400 | 798 | 0.03 | A0A0B5BBH7 | *Geobacter pickeringii* | Endopeptidase La (EC 3.4.21.53) |
| 401 | 798 | 0.03 | Q39YS7 | *Geobacter metallireducens* | Peptidoglycan transglycosylase and transpeptidase MrcA |
| 402 | 796 | 0.03 | A0A0B5BLL1 | *Geobacter pickeringii* | Pyridine nucleotide-disulfide oxidoreductase |
| 403 | 796 | 0.03 | A0A0C1U294 | *Geobacter soli* | Pilus assembly protein PilQ |

| | | | | | |
|---|---|---|---|---|---|
| 404 | 795 | 0.03 | Q39UW3 | *Geobacter metallireducens* | Succinyl:acetate coenzyme A transferase |
| 405 | 794 | 0.03 | Q39VU9 | *Geobacter metallireducens* | Polyprenyl-phospho-glycoside--protein O-glycosyltransferase DUF2723 membrane protein, putative |
| 406 | 792 | 0.03 | A0A0D5N732 | *Geobacter sulfurreducens* | Transcription-repair-coupling factor (TRCF) (EC 3.6.4.-) |
| 407 | 792 | 0.03 | A0A1G0LE89 | *Geobacteraceae bacterium* | Aconitate hydratase (Aconitase) (EC 4.2.1.3) |
| 408 | 789 | 0.03 | Q74GK2 | *Geobacter sulfurreducens* | Radical SAM domain iron-sulfur cluster-binding oxidoreductase |
| 409 | 787 | 0.03 | A0A0B5BE23 | *Geobacter pickeringii* | Uncharacterized protein |
| 410 | 786 | 0.03 | A0A0B5BIT5 | *Geobacter pickeringii* | Diadenylate cyclase (DAC) (EC 2.7.7.85) (Cyclic-di-AMP synthase) (c-di-AMP synthase) |
| 411 | 786 | 0.03 | Q39U98 | *Geobacter metallireducens* | Phosphoglycerate kinase (EC 2.7.2.3) |
| 412 | 786 | 0.03 | Q39T98 | *Geobacter metallireducens* | Cytochrome c |
| 413 | 785 | 0.03 | A0A0B5B8X8 | *Geobacter pickeringii* | Pilus assembly protein PilB |
| 414 | 785 | 0.03 | B5E8R5 | *Geobacter bemidjiensis* | Glutamine synthetase (EC 6.3.1.2) |
| 415 | 784 | 0.03 | A0A0D5N9G3 | *Geobacter sulfurreducens* | Multidrug ABC transporter substrate-binding protein |
| 416 | 781 | 0.03 | A0A0B5B7W3 | *Geobacter pickeringii* | Squalene cyclase |
| 417 | 781 | 0.03 | Q39RT7 | *Geobacter metallireducens* | Efflux pump, RND superfamily, and rRNA methyltransferase, putative |
| 418 | 780 | 0.03 | Q39QN4 | *Geobacter metallireducens* | Pentapeptide repeat protein |
| 419 | 780 | 0.03 | Q74GB8 | *Geobacter sulfurreducens* | Type II secretion system ATPase GspE |
| 420 | 779 | 0.03 | A0A0C1TRW2 | *Geobacter soli* | Fibronectin |
| 421 | 779 | 0.03 | Q748T9 | *Geobacter sulfurreducens* | Cytochrome c |
| 422 | 778 | 0.03 | A0A0B5BE51 | *Geobacter pickeringii* | DNA replication and repair protein RecF |
| 423 | 778 | 0.03 | Q39W43 | *Geobacter metallireducens* | Oligoendopeptidase F |
| 424 | 778 | 0.03 | | | |
| 425 | 777 | 0.03 | A0A0D5N4I0 | *Geobacter sulfurreducens* | Chemotaxis protein |
| 426 | 777 | 0.03 | Q747K3 | *Geobacter sulfurreducens* | UvrABC system protein B (Protein UvrB) (Excinuclease ABC subunit B) |
| 427 | 776 | 0.03 | A5GC26 | *Geobacter uraniireducens* | Biotin/lipoyl attachment domain-containing protein |
| 428 | 776 | 0.03 | Q39XB5 | *Geobacter metallireducens* | 30S ribosomal protein S1 |
| 429 | 776 | 0.03 | Q74BT6 | *Geobacter sulfurreducens* | Sigma-54-dependent transcriptional response regulator |

| | | | | | |
|---|---|---|---|---|---|
| 430 | 776 | 0.03 | W6LX92 | *Candidatus Contendobacter odensis* | Putative Lipoprotein |
| 431 | 774 | 0.03 | A0A0B5BCX9 | *Geobacter pickeringii* | Flagellar motor protein MotA |
| 432 | 772 | 0.03 | A0A0B5BCP9 | *Geobacter pickeringii* | Uncharacterized protein |
| 433 | 772 | 0.03 | A0A0B5BI05 | *Geobacter pickeringii* | Dihydrolipoamide acetyltransferase component of pyruvate dehydrogenase complex (EC 2.3.1.-) |
| 434 | 772 | 0.03 | Q39VY3 | *Geobacter metallireducens* | Aspartate--tRNA(Asp/Asn) ligase (EC 6.1.1.23) (Aspartyl-tRNA synthetase) (AspRS) (Non-discriminating aspartyl-tRNA synthetase) (ND-AspRS) |
| 435 | 771 | 0.03 | A0A0A8WN28 | *Geobacter sp.* | Protein TraN |
| 436 | 771 | 0.03 | Q74BG6 | *Geobacter sulfurreducens* | Serine protease, subtilase family |
| 437 | 769 | 0.03 | A0A0B5BFN3 | *Geobacter pickeringii* | Bifunctional protein HldE [Includes: D-beta-D-heptose 1-phosphate adenylyltransferase (EC 2.7.7.70) (D-glycero-beta-D-manno-heptose 1-phosphate adenylyltransferase); D-beta-D-heptose 7-phosphate kinase (EC 2.7.1.167) (D-beta-D-heptose 7-phosphotransferase) (D-glycero-beta-D-manno-heptose-7-phosphate kinase)] |
| 438 | 769 | 0.03 | B3EB58 | *Geobacter lovleyi* | Uncharacterized protein |
| 439 | 767 | 0.03 | Q74H55 | *Geobacter sulfurreducens* | Serine--tRNA ligase (EC 6.1.1.11) (Seryl-tRNA synthetase) (SerRS) (Seryl-tRNA(Ser/Sec) synthetase) |
| 440 | 766 | 0.03 | A0A133XMR2 | *Dechloromonas denitrificans* | Type I restriction endonuclease subunit M |
| 441 | 766 | 0.03 | | | |
| 442 | 765 | 0.03 | Q39US2 | *Geobacter metallireducens* | Carbamoyl-phosphate synthase small chain (EC 6.3.5.5) (Carbamoyl-phosphate synthetase glutamine chain) |
| 443 | 764 | 0.03 | A0A0D5N7J7 | *Geobacter sulfurreducens* | Alpha-glucan phosphorylase |
| 444 | 763 | 0.03 | A0A0B5BDH6 | *Geobacter pickeringii* | Membrane protein |
| 445 | 763 | 0.03 | A0A0B5BJ50 | *Geobacter pickeringii* | Chemotaxis protein CheA |
| 446 | 763 | 0.03 | Q39VA1 | *Geobacter metallireducens* | Polyribonucleotide nucleotidyltransferase (EC 2.7.7.8) (Polynucleotide phosphorylase) (PNPase) |
| 447 | 763 | 0.03 | Q39QK5 | *Geobacter metallireducens* | Uncharacterized protein |
| 448 | 762 | 0.03 | A0A0B5BD51 | *Geobacter pickeringii* | Uncharacterized protein |
| 449 | 761 | 0.03 | A0A0D5NCI9 | *Geobacter sulfurreducens* | GTPase-activating protein |
| 450 | 759 | 0.03 | Q747E4 | *Geobacter sulfurreducens* | Polyphosphate kinase (EC 2.7.4.1) (ATP-polyphosphate phosphotransferase) (Polyphosphoric acid kinase) |
| 451 | 757 | 0.03 | A0A0A8WMQ9 | *Geobacter sp.* | Uncharacterized protein |
| 452 | 757 | 0.03 | A5G661 | *Geobacter uraniireducens* | Uncharacterized protein |
| 453 | 756 | 0.03 | Q748Y7 | *Geobacter sulfurreducens* | 30S ribosomal protein S7 |

| | | | | | |
|---|---|---|---|---|---|
| 454 | 754 | 0.03 | A0A0D5N2Y8 | *Geobacter sulfurreducens* | Pyridine nucleotide-disulfide oxidoreductase |
| 455 | 753 | 0.03 | A0A0D5N9L3 | *Geobacter sulfurreducens* | Uncharacterized protein |
| 456 | 753 | 0.03 | A0A1F9PKR5 | *Desulfuromonadales bacterium* | Uncharacterized protein |
| 457 | 752 | 0.03 | Q39XT7 | *Geobacter metallireducens* | Ammonium transporter |
| 458 | 751 | 0.03 | A0A0D5N627 | *Geobacter sulfurreducens* | SpoVR family protein |
| 459 | 751 | 0.03 | A0A1G0N3T4 | *Geobacteraceae bacterium* | Hydroxylamine reductase (EC 1.7.99.1) (Hybrid-cluster protein) (HCP) (Prismane protein) |
| 460 | 751 | 0.03 | Q39WW8 | *Geobacter metallireducens* | Periplasmic serine endoprotease DegP-like (EC 3.4.21.107) |
| 461 | 749 | 0.03 | Q39Q96 | *Geobacter metallireducens* | Type II secretion system inner membrane protein GspF |
| 462 | 749 | 0.03 | Q748Q9 | *Geobacter sulfurreducens* | Methyl-accepting chemotaxis sensory transducer, class 40H |
| 463 | 748 | 0.03 | A0A0A8WW30 | *Geobacter sp.* | DNA-directed RNA polymerase subunit beta (RNAP subunit beta) (EC 2.7.7.6) (RNA polymerase subunit beta) (Transcriptase subunit beta) |
| 464 | 748 | 0.03 | Q74BF0 | *Geobacter sulfurreducens* | Phosphoribosylaminoimidazole-succinocarboxamide synthase (EC 6.3.2.6) (SAICAR synthetase) |
| 465 | 748 | 0.03 | Q39T78 | *Geobacter metallireducens* | Methionine--tRNA ligase (EC 6.1.1.10) (Methionyl-tRNA synthetase) (MetRS) |
| 466 | 748 | 0.03 | Q74F34 | *Geobacter sulfurreducens* | Sensor histidine kinase, HAMP domain-containing |
| 467 | 747 | 0.03 | Q74DB6 | *Geobacter sulfurreducens* | DNA-directed DNA polymerase (EC 2.7.7.7) |
| 468 | 746 | 0.03 | K1YAK3 | *groundwater metagenome* | Uncharacterized protein (Fragment) |
| 469 | 746 | 0.03 | | | |
| 470 | 745 | 0.03 | A0A0B5BEJ5 | *Geobacter pickeringii* | Diguanylate phosphodiesterase |
| 471 | 745 | 0.03 | B5EAA1 | *Geobacter bemidjiensis* | Undecaprenyl-diphospho-oligosaccharide flippase |
| 472 | 745 | 0.03 | Q39Q46 | *Geobacter metallireducens* | ABC transporter, ATP-binding/membrane protein |
| 473 | 745 | 0.03 | Q39YT5 | *Geobacter metallireducens* | 3-deoxy-D-arabino-heptulosonate 7-phosphate synthase |
| 474 | 744 | 0.03 | Q74CS2 | *Geobacter sulfurreducens* | Phosphate acyltransferase (EC 2.3.1.n2) (Acyl-ACP phosphotransacylase) (Acyl-[acyl-carrier-protein]--phosphate acyltransferase) (Phosphate-acyl-ACP acyltransferase) |
| 475 | 743 | 0.03 | A0A0B5BAN8 | *Geobacter pickeringii* | Branched-chain amino acid ABC transporter permease |
| 476 | 743 | 0.03 | Q39TA6 | *Geobacter metallireducens* | Inosine-5'-monophosphate dehydrogenase (IMP dehydrogenase) (IMPD) (IMPDH) (EC 1.1.1.205) |
| 477 | 741 | 0.03 | A0A0B5BDF2 | *Geobacter pickeringii* | Multifunctional fusion protein [Includes: ADP-dependent (S)-NAD(P)H-hydrate dehydratase (EC 4.2.1.136) (ADP-dependent NAD(P)HX dehydratase); NAD(P)H-hydrate epimerase (EC 5.1.99.6) (NAD(P)HX epimerase)] |
| 478 | 741 | 0.03 | Q39R24 | *Geobacter metallireducens* | Helix-turn-helix transcriptional regulator, putative |

| | | | | | |
|---|---|---|---|---|---|
| 479 | 740 | 0.03 | A0A0B5BHF1 | *Geobacter pickeringii* | Antibiotic ABC transporter ATP-binding protein |
| 480 | 740 | 0.03 | A0A0D5N930 | *Geobacter sulfurreducens* | Cytochrome C |
| 481 | 740 | 0.03 | Q39RQ8 | *Geobacter metallireducens* | Ribose-phosphate pyrophosphokinase (RPPK) (EC 2.7.6.1) (5-phospho-D-ribosyl alpha-1-diphosphate) (Phosphoribosyl diphosphate synthase) (Phosphoribosyl pyrophosphate synthase) (P-Rib-PP synthase) (PRPP synthase) (PRPPase) |
| 482 | 740 | 0.03 | Q39ZM0 | *Geobacter metallireducens* | Iron-sulfur cluster-binding sigma-54-dependent transcriptional regulator, FehydlgC and FeS domain-containing |
| 483 | 739 | 0.03 | Q39RF8 | *Geobacter metallireducens* | Carbon starvation protein CstA |
| 484 | 738 | 0.03 | A0A0B5B759 | *Geobacter pickeringii* | Probable peptidoglycan glycosyltransferase FtsW (PGT) (EC 2.4.1.129) (Cell division protein FtsW) (Cell wall polymerase) (Peptidoglycan polymerase) (PG polymerase) |
| 485 | 738 | 0.03 | A0A1G0M889 | *Geobacteraceae bacterium* | Uncharacterized protein |
| 486 | 738 | 0.03 | A5G6K7 | *Geobacter uraniireducens* | Diguanylate cyclase |
| 487 | 738 | 0.03 | P14243 | *Citrobacter freundii* | Modification methylase Cfr9I (M.Cfr9I) (EC 2.1.1.113) (N(4)- cytosine-specific methyltransferase Cfr9I) |
| 488 | 735 | 0.03 | Q39ZF6 | *Geobacter metallireducens* | Carbamoyltransferase HypF (EC 2.1.3.-) |
| 489 | 734 | 0.03 | A0A0B5BCY4 | *Geobacter pickeringii* | Aminodeoxyfutalosine synthase (AFL synthase) (Aminofutalosine synthase) (EC 2.5.1.120) (Menaquinone biosynthetic enzyme MqnE) |
| 490 | 734 | 0.03 | Q74BW2 | *Geobacter sulfurreducens* | Uridylate kinase (UK) (EC 2.7.4.22) (Uridine monophosphate kinase) (UMP kinase) (UMPK) |
| 491 | 734 | 0.03 | Q39RR7 | *Geobacter metallireducens* | Cytochrome c |
| 492 | 734 | 0.03 | | | |
| 493 | 732 | 0.03 | A0A0B5BHV2 | *Geobacter pickeringii* | Cytochrome C |
| 494 | 731 | 0.03 | A0A0D5N7A2 | *Geobacter sulfurreducens* | Chemotaxis protein CheY |
| 495 | 731 | 0.03 | Q39XX9 | *Geobacter metallireducens* | DNA-directed RNA polymerase subunit alpha (RNAP subunit alpha) (EC 2.7.7.6) (RNA polymerase subunit alpha) (Transcriptase subunit alpha) |
| 496 | 730 | 0.03 | A0A0D5N6L1 | *Geobacter sulfurreducens* | Histidine kinase |
| 497 | 729 | 0.03 | A0A0B5BJG9 | *Geobacter pickeringii* | GntR family transcriptional regulator |
| 498 | 729 | 0.03 | A0A0D5N4L3 | *Geobacter sulfurreducens* | UDP-N-acetyl-D-glucosamine dehydrogenase |
| 499 | 728 | 0.03 | A0A0B5BAQ7 | *Geobacter pickeringii* | HD family phosphohydrolase |
| 500 | 728 | 0.03 | A0A1E4LQS0 | *Comamonadaceae bacterium* | Histidine kinase |

**Table S4b.** Functional overview of M1C community (top 500 rank only).

| Rank | Count | Abundance | ID | Organism | Protein Names |
|---|---|---|---|---|---|
| 1 | 6173 | 0.19 | A0A090I137 | *Methanobacterium formicicum* | Transposase |
| 2 | 4631 | 0.14 | A0A1D3L4G2 | *Methanobacterium congolense* | Uncharacterized protein |
| 3 | 2833 | 0.09 | A0A090JUS9 | *Methanobacterium formicicum* | Putative secreted protein |
| 4 | 2664 | 0.08 | A0A090I3J2 | *Methanobacterium formicicum* | Uncharacterized protein |
| 5 | 2508 | 0.08 | | | |
| 6 | 2495 | 0.08 | A0A1D3L5C3 | *Methanobacterium congolense* | Uncharacterized protein |
| 7 | 2469 | 0.07 | A7I7H9 | *Methanoregula boonei* | TPR repeat-containing protein |
| 8 | 2078 | 0.06 | A0A1D3L0E4 | *Methanobacterium congolense* | WD40 domain-containing protein |
| 9 | 2071 | 0.06 | A7I4I9 | *Methanoregula boonei* | DNA polymerase II large subunit (Pol II) (EC 2.7.7.7) |
| 10 | 2008 | 0.06 | A0A1D3L0V2 | *Methanobacterium congolense* | Putative ATP-dependent helicase MG140 (EC 3.6.4.-) |
| 11 | 1964 | 0.06 | A0A1D3L5A7 | *Methanobacterium congolense* | Uncharacterized protein |
| 12 | 1923 | 0.06 | A7IAM2 | *Methanoregula boonei* | Isoleucine--tRNA ligase (EC 6.1.1.5) (Isoleucyl-tRNA synthetase) (IleRS) |
| 13 | 1869 | 0.06 | A0A1D3L3T8 | *Methanobacterium congolense* | Type IIS restriction enzyme Eco57I (EC 3.1.21.4) |
| 14 | 1856 | 0.06 | P06131 | *Methanobacterium formicicum* | Formate dehydrogenase subunit alpha (EC 1.2.1.2) |
| 15 | 1821 | 0.05 | A0A1D3L508 | *Methanobacterium congolense* | WD40 domain-containing protein |
| 16 | 1770 | 0.05 | A7IAV3 | *Methanoregula boonei* | V-type ATP synthase subunit I |
| 17 | 1762 | 0.05 | A0A101HE03 | *Methanobacterium sp.* | WD40 domain protein beta Propeller |
| 18 | 1652 | 0.05 | A0A1D3KZI4 | *Methanobacterium congolense* | DNA polymerase II large subunit (Pol II) (EC 2.7.7.7) |
| 19 | 1645 | 0.05 | A0A0S4FV74 | *Methanobacterium formicicum* | Uncharacterized protein |
| 20 | 1618 | 0.05 | A0A1D3L5B8 | *Methanobacterium congolense* | Isoleucine--tRNA ligase (EC 6.1.1.5) (Isoleucyl-tRNA synthetase) (IleRS) |
| 21 | 1604 | 0.05 | A7I486 | *Methanoregula boonei* | K(+)-insensitive pyrophosphate-energized proton pump (EC 3.6.1.1) (Membrane-bound proton-translocating pyrophosphatase) (Pyrophosphate-energized inorganic pyrophosphatase) (H(+)-PPase) |
| 22 | 1589 | 0.05 | A7IAD8 | *Methanoregula boonei* | Chromosome partition protein Smc |
| 23 | 1587 | 0.05 | A7I9N6 | *Methanoregula boonei* | DNA-directed RNA polymerase subunit (EC 2.7.7.6) |
| 24 | 1555 | 0.05 | A0A089ZCJ0 | *Methanobacterium formicicum* | Cobalamin biosynthesis protein N (EC 6.6.1.2) (Cobaltochelatase CobN2) |

| | | | | | |
|---|---|---|---|---|---|
| 25 | 1547 | 0.05 | A0A1D3KZI9 | *Methanobacterium congolense* | Leucine--tRNA ligase (EC 6.1.1.4) (Leucyl-tRNA synthetase) (LeuRS) |
| 26 | 1545 | 0.05 | A7I927 | *Methanoregula boonei* | Valine--tRNA ligase (EC 6.1.1.9) (Valyl-tRNA synthetase) (ValRS) |
| 27 | 1511 | 0.05 | A0A1D3L5M9 | *Methanobacterium congolense* | Putative ATP-dependent RNA helicase MJ1505 (EC 3.6.4.13) |
| 28 | 1500 | 0.05 | A0A1D3L5A8 | *Methanobacterium congolense* | Long-chain-fatty-acid-CoA ligase FadD15 (EC 6.2.1.3) |
| 29 | 1499 | 0.05 | A7I656 | *Methanoregula boonei* | Elongation factor 1-alpha (EF-1-alpha) (Elongation factor Tu) (EF-Tu) |
| 30 | 1451 | 0.04 | A7I4X4 | *Methanoregula boonei* | Elongation factor 2 (EF-2) |
| 31 | 1448 | 0.04 | U6ED34 | *Methanobacterium sp.* | Protoporphyrin IX magnesium chelatase |
| 32 | 1443 | 0.04 | A0A1D3L027 | *Methanobacterium congolense* | Carbamoyl-phosphate synthase large chain (EC 6.3.5.5) (Carbamoyl-phosphate synthetase ammonia chain) |
| 33 | 1440 | 0.04 | A7I9M8 | *Methanoregula boonei* | Leucine--tRNA ligase (EC 6.1.1.4) (Leucyl-tRNA synthetase) (LeuRS) |
| 34 | 1439 | 0.04 | A7I5I6 | *Methanoregula boonei* | Coenzyme F420 hydrogenase (EC 1.12.98.1) |
| 35 | 1425 | 0.04 | A0A1D3L429 | *Methanobacterium congolense* | Uncharacterized protein |
| 36 | 1400 | 0.04 | A7I5U5 | *Methanoregula boonei* | 4Fe-4S ferredoxin, iron-sulfur binding domain protein |
| 37 | 1391 | 0.04 | A7I4W0 | *Methanoregula boonei* | Oligosaccharyl transferase, STT3 subunit |
| 38 | 1389 | 0.04 | A7I6J1 | *Methanoregula boonei* | Carbamoyl-phosphate synthase large chain (EC 6.3.5.5) (Carbamoyl-phosphate synthetase ammonia chain) |
| 39 | 1374 | 0.04 | A0A1D3L3R4 | *Methanobacterium congolense* | Putative membrane protein MJ1525 |
| 40 | 1352 | 0.04 | A0A1D3KZ93 | *Methanobacterium congolense* | Sodium/potassium-transporting ATPase subunit alpha-2 (EC 3.6.3.9) |
| 41 | 1345 | 0.04 | A0A1D3L3L7 | *Methanobacterium congolense* | Metabotropic glutamate receptor-like protein R |
| 42 | 1337 | 0.04 | A0A1D3L118 | *Methanobacterium congolense* | Putative thiamine pyrophosphate-containing protein YdaP |
| 43 | 1332 | 0.04 | A7I766 | *Methanoregula boonei* | Efflux transporter, putative, hydrophobe/amphiphile efflux-3 (HAE3) family |
| 44 | 1320 | 0.04 | U6EA67 | *Methanobacterium sp.* | Uncharacterized protein |
| 45 | 1310 | 0.04 | A7I6S1 | *Methanoregula boonei* | CoA-binding domain protein |
| 46 | 1306 | 0.04 | A7IAQ1 | *Methanoregula boonei* | Uncharacterized protein |
| 47 | 1277 | 0.04 | A0A1D3L4J7 | *Methanobacterium congolense* | Transglutaminase domain-containing protein |
| 48 | 1273 | 0.04 | A7I4R1 | *Methanoregula boonei* | Thermosome |
| 49 | 1264 | 0.04 | A0A1D3L5E6 | *Methanobacterium congolense* | Replication factor A |
| 50 | 1261 | 0.04 | A7I4K7 | *Methanoregula boonei* | Lysine decarboxylase (EC 4.1.1.18) |

| | | | | | |
|---|---|---|---|---|---|
| 51 | 1260 | 0.04 | A0A1D3L2M2 | *Methanobacterium congolense* | Valine--tRNA ligase (EC 6.1.1.9) (Valyl-tRNA synthetase) (ValRS) |
| 52 | 1258 | 0.04 | A0A1D3L412 | *Methanobacterium congolense* | Putative MFS-type transporter YusP |
| 53 | 1246 | 0.04 | A7I6H7 | *Methanoregula boonei* | Methionine--tRNA ligase (EC 6.1.1.10) (Methionyl-tRNA synthetase) (MetRS) |
| 54 | 1243 | 0.04 | A7I5I3 | *Methanoregula boonei* | Coenzyme F420 hydrogenase/dehydrogenase beta subunit domain protein |
| 55 | 1232 | 0.04 | A0A089ZBK8 | *Methanobacterium formicicum* | Adhesin-like protein |
| 56 | 1232 | 0.04 | A0A1D3L5Q7 | *Methanobacterium congolense* | PAS domain S-box |
| 57 | 1231 | 0.04 | A0A1D3L0B4 | *Methanobacterium congolense* | Putative ATP-dependent helicase MJ1401 (EC 3.6.4.-) |
| 58 | 1226 | 0.04 | A7I791 | *Methanoregula boonei* | UvrABC system protein A (UvrA protein) (Excinuclease ABC subunit A) |
| 59 | 1224 | 0.04 | A0A089ZEB0 | *Methanobacterium formicicum* | WD40 domain-containing cell surface protein |
| 60 | 1222 | 0.04 | A0A1D3KZH1 | *Methanobacterium congolense* | Copper-exporting P-type ATPase A (EC 3.6.3.54) |
| 61 | 1221 | 0.04 | A7I9M7 | *Methanoregula boonei* | PilT protein domain protein |
| 62 | 1218 | 0.04 | L0HAI4 | *Methanoregula formicica* | Formate dehydrogenase, alpha subunit, archaeal-type |
| 63 | 1214 | 0.04 | A0A1D3L0D2 | *Methanobacterium congolense* | Copper-exporting P-type ATPase B (EC 3.6.3.54) |
| 64 | 1212 | 0.04 | A7IAB8 | *Methanoregula boonei* | Phosphoenolpyruvate synthase (PEP synthase) (EC 2.7.9.2) (Pyruvate, water dikinase) |
| 65 | 1205 | 0.04 | | | |
| 66 | 1200 | 0.04 | A0A1D3L4E7 | *Methanobacterium congolense* | Phosphoenolpyruvate synthase (PEP synthase) (EC 2.7.9.2) (Pyruvate, water dikinase) |
| 67 | 1184 | 0.04 | A0A1D3KZY6 | *Methanobacterium congolense* | Elongation factor 2 (EF-2) |
| 68 | 1175 | 0.04 | A0A1D3L3Z9 | *Methanobacterium congolense* | Uncharacterized protein |
| 69 | 1172 | 0.04 | L0HG76 | *Methanoregula formicica* | CRISPR-associated helicase Cas3/CRISPR-associated endonuclease Cas3-HD |
| 70 | 1161 | 0.03 | E1RD00 | *Methanolacinia petrolearia* | Pyrrolo-quinoline quinone |
| 71 | 1155 | 0.03 | A0A1D3L4X9 | *Methanobacterium congolense* | Putative transcriptional regulatory protein pdtaR |
| 72 | 1155 | 0.03 | A7I485 | *Methanoregula boonei* | Vitamin B12-dependent ribonucleotide reductase (EC 1.17.4.1) |
| 73 | 1146 | 0.03 | A7I7B5 | *Methanoregula boonei* | DNA polymerase (EC 2.7.7.7) |
| 74 | 1143 | 0.03 | A0A1D3L4R2 | *Methanobacterium congolense* | Cyanobacterial phytochrome A (EC 2.7.13.3) |
| 75 | 1137 | 0.03 | A0A1D3L0F2 | *Methanobacterium congolense* | Archaeal Lon protease (EC 3.4.21.-) |
| 76 | 1136 | 0.03 | A7I4Y9 | *Methanoregula boonei* | DEAD/DEAH box helicase domain protein |
| 77 | 1128 | 0.03 | A7I7Z8 | *Methanoregula boonei* | Type II secretion system protein |

| | | | | | |
|---|---|---|---|---|---|
| 78 | 1123 | 0.03 | A7I5N0 | *Methanoregula boonei* | AMP phosphorylase (AMPpase) (EC 2.4.2.57) (Nucleoside monophosphate phosphorylase) (NMP phosphorylase) |
| 79 | 1119 | 0.03 | A7I9L7 | *Methanoregula boonei* | 4-vinyl reductase, 4VR |
| 80 | 1107 | 0.03 | A0A090I3X0 | *Methanobacterium formicicum* | Nitrogenase protein alpha chain (EC 1.18.6.1) |
| 81 | 1102 | 0.03 | A7I7A6 | *Methanoregula boonei* | Methyl-accepting chemotaxis sensory transducer |
| 82 | 1098 | 0.03 | A0A1D3L5G6 | *Methanobacterium congolense* | DNA polymerase II small subunit (Pol II) (EC 2.7.7.7) |
| 83 | 1097 | 0.03 | A7IB61 | *Methanoregula boonei* | ATP-dependent DNA helicase Hel308 (EC 3.6.4.12) |
| 84 | 1097 | 0.03 | A7I7S4 | *Methanoregula boonei* | Alanine--tRNA ligase (EC 6.1.1.7) (Alanyl-tRNA synthetase) (AlaRS) |
| 85 | 1096 | 0.03 | A0A1D3KZE7 | *Methanobacterium congolense* | Putative sensor histidine kinase pdtaS (EC 2.7.13.3) |
| 86 | 1096 | 0.03 | A0A1D3L3K7 | *Methanobacterium congolense* | Uncharacterised protein family (UPF0240) |
| 87 | 1085 | 0.03 | A0A1D3L0D3 | *Methanobacterium congolense* | Calcium-transporting ATPase 1 (EC 3.6.3.8) |
| 88 | 1084 | 0.03 | A0A1D3L4A4 | *Methanobacterium congolense* | Putative protease YrrO (EC 3.4.-.-) |
| 89 | 1082 | 0.03 | A0A1D3L2A8 | *Methanobacterium congolense* | Hydroxylamine reductase (EC 1.7.99.1) (Hybrid-cluster protein) (HCP) (Prismane protein) |
| 90 | 1078 | 0.03 | A0A1D3L276 | *Methanobacterium congolense* | UvrABC system protein A (UvrA protein) (Excinuclease ABC subunit A) |
| 91 | 1070 | 0.03 | A0A1D3L596 | *Methanobacterium congolense* | CoB-CoM heterodisulfide reductase iron-sulfur subunit A (EC 1.8.98.1) |
| 92 | 1067 | 0.03 | | | |
| 93 | 1066 | 0.03 | A0A1D3L1V6 | *Methanobacterium congolense* | Uncharacterized protein |
| 94 | 1054 | 0.03 | A0A1D3L506 | *Methanobacterium congolense* | Carbamoyltransferase (EC 2.1.3.-) |
| 95 | 1053 | 0.03 | A0A1D3L4H2 | *Methanobacterium congolense* | Inner membrane protein YccS |
| 96 | 1049 | 0.03 | E1RH75 | *Methanolacinia petrolearia* | Putative PAS/PAC sensor protein |
| 97 | 1044 | 0.03 | A0A089ZDV6 | *Methanobacterium formicicum* | Transposase |
| 98 | 1043 | 0.03 | A0A090I5G5 | *Methanobacterium formicicum* | Uncharacterized protein |
| 99 | 1043 | 0.03 | A7I7Z9 | *Methanoregula boonei* | Type II secretion system protein E |
| 100 | 1040 | 0.03 | A0A089ZVJ1 | *Methanobacterium formicicum* | Cobaltochelatase CobN3 (Putative secreted protein) |
| 101 | 1039 | 0.03 | A0A1D3L1G9 | *Methanobacterium congolense* | DNA helicase II related protein |
| 102 | 1038 | 0.03 | A7I9Q1 | *Methanoregula boonei* | Glutamine synthetase (EC 6.3.1.2) |
| 103 | 1030 | 0.03 | A0A1D3L3V1 | *Methanobacterium congolense* | Cyanobacterial phytochrome A (EC 2.7.13.3) |

| | | | | | |
|---|---|---|---|---|---|
| 104 | 1011 | 0.03 | A0A1D3L129 | *Methanobacterium congolense* | Putative ATP-dependent helicase MTH_1802 (EC 3.6.4.-) |
| 105 | 1011 | 0.03 | A7I4V8 | *Methanoregula boonei* | Translation initiation factor 2 subunit gamma (aIF2-gamma) (eIF-2-gamma) |
| 106 | 1010 | 0.03 | A7IAF8 | *Methanoregula boonei* | ABC-1 domain protein |
| 107 | 1009 | 0.03 | A7I6S2 | *Methanoregula boonei* | Acetate--CoA ligase |
| 108 | 1007 | 0.03 | A0A1D3KZR9 | *Methanobacterium congolense* | Putative PAS/PAC sensor protein |
| 109 | 1005 | 0.03 | A7I5R2 | *Methanoregula boonei* | Protein translocase subunit SecY (Protein transport protein SEC61 subunit alpha homolog) |
| 110 | 1004 | 0.03 | A0A090JW16 | *Methanobacterium formicicum* | Uncharacterized protein |
| 111 | 1002 | 0.03 | A7IAZ8 | *Methanoregula boonei* | Chaperonin Cpn60/TCP-1 |
| 112 | 1000 | 0.03 | A7I6T1 | *Methanoregula boonei* | Coenzyme F420 hydrogenase/dehydrogenase beta subunit domain protein |
| 113 | 999 | 0.03 | A7I6P0 | *Methanoregula boonei* | Threonine--tRNA ligase (EC 6.1.1.3) (Threonyl-tRNA synthetase) (ThrRS) |
| 114 | 998 | 0.03 | A0A1D3L584 | *Methanobacterium congolense* | Uncharacterized protein |
| 115 | 992 | 0.03 | A0A1D3L132 | *Methanobacterium congolense* | Putative pseudomurein-binding protein |
| 116 | 988 | 0.03 | A7I5S6 | *Methanoregula boonei* | Tetrahydromethanopterin S-methyltransferase subunit A (EC 2.1.1.86) (N5-methyltetrahydromethanopterin--coenzyme M methyltransferase subunit A) |
| 117 | 987 | 0.03 | A0A1D3L2H5 | *Methanobacterium congolense* | tRNA-guanine(15) transglycosylase (EC 2.4.2.48) (7-cyano-7-deazaguanine tRNA-ribosyltransferase) (Archaeal tRNA-guanine transglycosylase) |
| 118 | 987 | 0.03 | A0A1D3L5K7 | *Methanobacterium congolense* | Threonine--tRNA ligase (EC 6.1.1.3) (Threonyl-tRNA synthetase) (ThrRS) |
| 119 | 987 | 0.03 | A7I8Y5 | *Methanoregula boonei* | DEAD/DEAH box helicase domain protein |
| 120 | 987 | 0.03 | U6E9M3 | *Methanobacterium sp.* | DNA-directed RNA polymerase subunit (EC 2.7.7.6) |
| 121 | 986 | 0.03 | A0A1D3L0Y3 | *Methanobacterium congolense* | Cell division cycle protein 48 homolog MJ1156 |
| 122 | 983 | 0.03 | A0A1D3L007 | *Methanobacterium congolense* | Alanine--tRNA ligase (EC 6.1.1.7) (Alanyl-tRNA synthetase) (AlaRS) |
| 123 | 983 | 0.03 | A0A1D3L4D8 | *Methanobacterium congolense* | Pyruvate carboxylase subunit B (EC 6.4.1.1) |
| 124 | 982 | 0.03 | A0A1D3L3Z6 | *Methanobacterium congolense* | DNA replication licensing factor MCM7 (EC 3.6.4.12) |
| 125 | 982 | 0.03 | A0A1D3L579 | *Methanobacterium congolense* | Phosphoribosylformylglycinamidine synthase subunit PurL (FGAM synthase) (EC 6.3.5.3) (Formylglycinamide ribonucleotide amidotransferase subunit II) (FGAR amidotransferase II) (FGAR-AT II) (Glutamine amidotransferase PurL) (Phosphoribosylformylglycinamidine synthase subunit II) |
| 126 | 980 | 0.03 | A7I4F8 | *Methanoregula boonei* | Amidophosphoribosyltransferase (ATase) (EC 2.4.2.14) (Glutamine phosphoribosylpyrophosphate amidotransferase) (GPATase) |
| 127 | 978 | 0.03 | A0A1D3L379 | *Methanobacterium congolense* | Calcium-transporting ATPase 1 (EC 3.6.3.8) |

| | | | | | |
|---|---|---|---|---|---|
| 128 | 976 | 0.03 | A0A1D3KZ84 | *Methanobacterium congolense* | Uncharacterized protein |
| 129 | 975 | 0.03 | A0A1D3L1K6 | *Methanobacterium congolense* | Putative ATP-dependent helicase YprA (EC 3.6.4.-) |
| 130 | 974 | 0.03 | A0A1D3L1R0 | *Methanobacterium congolense* | Acetate-CoA ligase [ADP-forming] I (EC 6.2.1.13) |
| 131 | 973 | 0.03 | L0HHJ6 | *Methanoregula formicica (strain DSM 22288 / NBRC 105244 / SMSP)* | Tubulin-like protein CetZ |
| 132 | 972 | 0.03 | A0A1D3KZW8 | *Methanobacterium congolense* | Uncharacterized protein |
| 133 | 970 | 0.03 | A0A1D3L3P1 | *Methanobacterium congolense* | Probable translation initiation factor IF-2 |
| 134 | 964 | 0.03 | A0A1D3L0U2 | *Methanobacterium congolense* | Archaeal glutamate synthase [NADPH] (EC 1.4.1.13) |
| 135 | 961 | 0.03 | A7I667 | *Methanoregula boonei* | Phenylacetate--CoA ligase |
| 136 | 956 | 0.03 | A7I8X9 | *Methanoregula boonei* | MCM family protein |
| 137 | 955 | 0.03 | A0A1D3L3F6 | *Methanobacterium congolense* | Minor teichoic acid biosynthesis protein GgaB |
| 138 | 954 | 0.03 | L0HCK9 | *Methanoregula formicica* | Formylmethanofuran dehydrogenase subunit A |
| 139 | 953 | 0.03 | A0A1D3L355 | *Methanobacterium congolense* | Lon protease (EC 3.4.21.53) (ATP-dependent protease La) |
| 140 | 952 | 0.03 | A0A1D3L1R3 | *Methanobacterium congolense* | Putative sensor histidine kinase pdtaS (EC 2.7.13.3) |
| 141 | 947 | 0.03 | A0A1D3KZX3 | *Methanobacterium congolense* | DNA-directed RNA polymerase subunit beta (EC 2.7.7.6) |
| 142 | 947 | 0.03 | A7I9N8 | *Methanoregula boonei* | DNA-directed RNA polymerase (EC 2.7.7.6) |
| 143 | 945 | 0.03 | A0A1D3L5J6 | *Methanobacterium congolense* | Acetolactate synthase (EC 2.2.1.6) |
| 144 | 942 | 0.03 | A0A1D3L4X3 | *Methanobacterium congolense* | Archaeal glutamate synthase [NADPH] (EC 1.4.1.13) |
| 145 | 938 | 0.03 | A0A1D3L3V9 | *Methanobacterium congolense* | Putative KH and PIN-domain containing protein MJ1533 |
| 146 | 936 | 0.03 | A7I8P2 | *Methanoregula boonei* | Phosphoesterase, RecJ domain protein |
| 147 | 935 | 0.03 | A7I7Z3 | *Methanoregula boonei* | CheA signal transduction histidine kinase |
| 148 | 934 | 0.03 | A7I6H1 | *Methanoregula boonei* | Glycyl-tRNA synthetase |
| 149 | 930 | 0.03 | A0A1D3L430 | *Methanobacterium congolense* | Cell division cycle protein 48 homolog MJ1156 |
| 150 | 928 | 0.03 | A7I8Y8 | *Methanoregula boonei* | AMP-dependent synthetase and ligase |
| 151 | 926 | 0.03 | A0A1D3L2S6 | *Methanobacterium congolense* | Putative sensor histidine kinase pdtaS (EC 2.7.13.3) |
| 152 | 920 | 0.03 | A0A1D3L3E8 | *Methanobacterium congolense* | Indolepyruvate oxidoreductase subunit IorA (IOR) (EC 1.2.7.8) (Indolepyruvate ferredoxin oxidoreductase subunit alpha) |
| 153 | 919 | 0.03 | A0A089ZFB5 | *Methanobacterium formicicum* | Adhesin-like protein |

| | | | | | |
|---|---|---|---|---|---|
| 154 | 916 | 0.03 | A7IAP7 | *Methanoregula boonei* | Probable translation initiation factor IF-2 |
| 155 | 915 | 0.03 | A7I9N7 | *Methanoregula boonei* | DNA-directed RNA polymerase subunit beta (EC 2.7.7.6) |
| 156 | 912 | 0.03 | A7IB37 | *Methanoregula boonei* | ABC transporter related |
| 157 | 911 | 0.03 | A7IAJ6 | *Methanoregula boonei* | DEAD/DEAH box helicase domain protein |
| 158 | 909 | 0.03 | A7IAY8 | *Methanoregula boonei* | Type II secretion system protein |
| 159 | 908 | 0.03 | A0A1D3L5K4 | *Methanobacterium congolense* | Uncharacterized protein |
| 160 | 907 | 0.03 | A7IB30 | *Methanoregula boonei* | AAA family ATPase, CDC48 subfamily (EC 3.6.4.6) |
| 161 | 906 | 0.03 | A0A1D3L5S0 | *Methanobacterium congolense* | Arginine--tRNA ligase (EC 6.1.1.19) (Arginyl-tRNA synthetase) (ArgRS) |
| 162 | 906 | 0.03 | L0HHJ9 | *Methanoregula formicica* | DNA gyrase subunit A (EC 5.99.1.3) |
| 163 | 904 | 0.03 | A0A1D3L452 | *Methanobacterium congolense* | DNA topoisomerase 1 (EC 5.99.1.2) (DNA topoisomerase I) |
| 164 | 903 | 0.03 | U6EDJ2 | *Methanobacterium sp.* | Uncharacterized protein |
| 165 | 900 | 0.03 | P06130 | *Methanobacterium formicicum* | Formate dehydrogenase subunit beta (EC 1.2.1.2) |
| 166 | 897 | 0.03 | A7IAE0 | *Methanoregula boonei* | Serine hydroxymethyltransferase (SHMT) (Serine methylase) (EC 2.1.2.1) |
| 167 | 895 | 0.03 | A0A1D3L5X1 | *Methanobacterium congolense* | Dihydroxy-acid dehydratase (DAD) (EC 4.2.1.9) |
| 168 | 892 | 0.03 | A0A1D3L2D6 | *Methanobacterium congolense* | Putative cadmium-transporting ATPase (EC 3.6.3.3) |
| 169 | 892 | 0.03 | A0A1D3L4H3 | *Methanobacterium congolense* | V-type ATP synthase subunit I |
| 170 | 892 | 0.03 | A0A1D3L5R3 | *Methanobacterium congolense* | Probable bifunctional tRNA threonylcarbamoyladenosine biosynthesis protein [Includes: Serine/threonine-protein kinase Bud32 (EC 2.7.11.1); tRNA N6-adenosine threonylcarbamoyltransferase (EC 2.3.1.234) (tRNA threonylcarbamoyladenosine biosynthesis protein Kae1) (t(6)A37 threonylcarbamoyladenosine biosynthesis protein Kae1)] |
| 171 | 891 | 0.03 | A0A1D3KZS9 | *Methanobacterium congolense* | Uncharacterized protein |
| 172 | 889 | 0.03 | A0A1D3L170 | *Methanobacterium congolense* | ATP-dependent DNA helicase Hel308 (EC 3.6.4.12) |
| 173 | 889 | 0.03 | A7I7B3 | *Methanoregula boonei* | UvrABC system protein B (Protein UvrB) (Excinuclease ABC subunit B) |
| 174 | 885 | 0.03 | A7I6K0 | *Methanoregula boonei* | Radical SAM domain protein |
| 175 | 884 | 0.03 | A0A1D3L0K8 | *Methanobacterium congolense* | Glutamine--fructose-6-phosphate aminotransferase [isomerizing] (EC 2.6.1.16) (D-fructose-6-phosphate amidotransferase) (GFAT) (Glucosamine-6-phosphate synthase) (Hexosephosphate aminotransferase) (L-glutamine--D-fructose-6-phosphate amidotransferase) |
| 176 | 882 | 0.03 | A0A1D3L3E2 | *Methanobacterium congolense* | Putative sensor histidine kinase pdtaS (EC 2.7.13.3) |

| | | | | | |
|---|---|---|---|---|---|
| 177 | 882 | 0.03 | | | |
| 178 | 881 | 0.03 | A7I859 | *Methanoregula boonei* | Uncharacterized protein |
| 179 | 881 | 0.03 | A7IAU7 | *Methanoregula boonei* | V-type ATP synthase beta chain (V-ATPase subunit B) |
| 180 | 880 | 0.03 | A0A1D3L5K9 | *Methanobacterium congolense* | Uncharacterized protein |
| 181 | 879 | 0.03 | A0A1D3L094 | *Methanobacterium congolense* | Magnesium-chelatase subunit ChlI homolog |
| 182 | 876 | 0.03 | A0A1D3L4H1 | *Methanobacterium congolense* | Acetyl-CoA decarbonylase/synthase complex subunit alpha (ACDS complex subunit alpha) (EC 1.2.7.4) (ACDS complex carbon monoxide dehydrogenase subunit alpha) (ACDS CODH subunit alpha) |
| 183 | 872 | 0.03 | A0A1D3L223 | *Methanobacterium congolense* | CoB-CoM heterodisulfide reductase 1 iron-sulfur subunit A (EC 1.8.98.1) |
| 184 | 872 | 0.03 | A7I4W3 | *Methanoregula boonei* | Glutamine--fructose-6-phosphate aminotransferase [isomerizing] (EC 2.6.1.16) (D-fructose-6-phosphate amidotransferase) (GFAT) (Glucosamine-6-phosphate synthase) (Hexosephosphate aminotransferase) (L-glutamine--D-fructose-6-phosphate amidotransferase) |
| 185 | 870 | 0.03 | A0A089ZH30 | *Methanobacterium formicicum* | Signal transduction histidine kinase |
| 186 | 870 | 0.03 | A7I798 | *Methanoregula boonei* | 60 kDa chaperonin (GroEL protein) (Protein Cpn60) |
| 187 | 868 | 0.03 | A7I6L6 | *Methanoregula boonei* | DNA topoisomerase 1 (EC 5.99.1.2) (DNA topoisomerase I) |
| 188 | 867 | 0.03 | A0A1D3L407 | *Methanobacterium congolense* | Putative actinorhodin transporter |
| 189 | 867 | 0.03 | A0A1D3L551 | *Methanobacterium congolense* | Uncharacterized protein |
| 190 | 866 | 0.03 | A0A0E4GC68 | *Syntrophomonas zehnderi OL-4* | Pectin lyase fold/virulence factor (Fragment) |
| 191 | 866 | 0.03 | A7IAQ3 | *Methanoregula boonei* | GCN5-related N-acetyltransferase |
| 192 | 865 | 0.03 | A0A1D3L5B6 | *Methanobacterium congolense* | Helicase c2 |
| 193 | 855 | 0.03 | A0A1D3L1W6 | *Methanobacterium congolense* | Methionine--tRNA ligase (EC 6.1.1.10) (Methionyl-tRNA synthetase) (MetRS) |
| 194 | 855 | 0.03 | A7I8Z2 | *Methanoregula boonei* | Ammonium transporter |
| 195 | 853 | 0.03 | A0A1D3L0K7 | *Methanobacterium congolense* | Digeranylgeranylglycerophospholipid reductase (DGGGPL reductase) (EC 1.3.7.11) (2,3-bis-O-geranylgeranylglyceryl phosphate reductase) (Geranylgeranyl reductase) (GGR) |
| 196 | 852 | 0.03 | A0A1D3L4V1 | *Methanobacterium congolense* | Uncharacterized protein |
| 197 | 852 | 0.03 | A7I988 | *Methanoregula boonei* | Acetyl-CoA carboxylase, biotin carboxylase |
| 198 | 852 | 0.03 | | | |
| 199 | 849 | 0.03 | A0A1D3L3Q6 | *Methanobacterium congolense* | Pyridoxal 5'-phosphate synthase subunit PdxS (PLP synthase subunit PdxS) (EC 4.3.3.6) (Pdx1) |

| | | | | | |
|---|---|---|---|---|---|
| 200 | 848 | 0.03 | A7I8B1 | *Methanoregula boonei* | Polyphosphate kinase (EC 2.7.4.1) (ATP-polyphosphate phosphotransferase) (Polyphosphoric acid kinase) |
| 201 | 848 | 0.03 | E1RH76 | *Methanolacinia petrolearia* | F420-dependent oxidoreductase, G6PDH family |
| 202 | 847 | 0.03 | A7I4G8 | *Methanoregula boonei* | Aspartate--tRNA(Asp/Asn) ligase (EC 6.1.1.23) (Aspartyl-tRNA synthetase) (AspRS) (Non-discriminating aspartyl-tRNA synthetase) (ND-AspRS) |
| 203 | 843 | 0.03 | A0A1D3L3N4 | *Methanobacterium congolense* | Acetyl-coenzyme A synthetase (EC 6.2.1.1) |
| 204 | 841 | 0.03 | A0A1D3L4J0 | *Methanobacterium congolense* | Phage infection protein |
| 205 | 840 | 0.03 | A7I822 | *Methanoregula boonei* | K+ potassium transporter |
| 206 | 839 | 0.03 | A7I464 | *Methanoregula boonei* | ORC1-type DNA replication protein |
| 207 | 839 | 0.03 | L0HEU5 | *Methanoregula formicica* | DNA gyrase subunit B (EC 5.99.1.3) |
| 208 | 838 | 0.03 | A0A1D3L0A6 | *Methanobacterium congolense* | Putative acyl-CoA synthetase YngI (EC 6.2.1.-) |
| 209 | 838 | 0.03 | A7I9N5 | *Methanoregula boonei* | DNA-directed RNA polymerase subunit A'' (EC 2.7.7.6) |
| 210 | 838 | 0.03 | A7I4Y2 | *Methanoregula boonei* | Arginine--tRNA ligase (EC 6.1.1.19) (Arginyl-tRNA synthetase) (ArgRS) |
| 211 | 832 | 0.03 | K2R0T8 | *Methanobacterium formicicum* | DNA-directed RNA polymerase subunit (EC 2.7.7.6) |
| 212 | 829 | 0.02 | A7I4C9 | *Methanoregula boonei* | Acetate--CoA ligase |
| 213 | 827 | 0.02 | A0A1D3L2G6 | *Methanobacterium congolense* | Methyl sulfide methyltransferase-associated sensor (EC 2.7.13.3) |
| 214 | 826 | 0.02 | A7I5U3 | *Methanoregula boonei* | Methyl-coenzyme M reductase, beta subunit |
| 215 | 825 | 0.02 | A0A1D3L5S7 | *Methanobacterium congolense* | Uncharacterized protein |
| 216 | 825 | 0.02 | A7I679 | *Methanoregula boonei* | Tyrosine recombinase XerA |
| 217 | 824 | 0.02 | A0A1D3KZQ9 | *Methanobacterium congolense* | Putative (R)-citramalate synthase CimA (EC 2.3.1.182) |
| 218 | 822 | 0.02 | A0A1D3L3Z3 | *Methanobacterium congolense* | Putative ABC transporter ATP-binding protein MJ0719 |
| 219 | 822 | 0.02 | A0A1D3L537 | *Methanobacterium congolense* | Uncharacterized protein |
| 220 | 821 | 0.02 | A7IA93 | *Methanoregula boonei* | CTP synthase (EC 6.3.4.2) (Cytidine 5'-triphosphate synthase) (Cytidine triphosphate synthetase) (CTP synthetase) (CTPS) (UTP--ammonia ligase) |
| 221 | 820 | 0.02 | A0A090I571 | *Methanobacterium formicicum* | Carbamoyl-phosphate synthase large chain (EC 6.3.5.5) (Carbamoyl-phosphate synthetase ammonia chain) |
| 222 | 819 | 0.02 | A7I4B9 | *Methanoregula boonei* | Aminotransferase (EC 2.6.1.-) |
| 223 | 817 | 0.02 | A0A089ZDS9 | *Methanobacterium formicicum* | DNA polymerase II large subunit (Pol II) (EC 2.7.7.7) |
| 224 | 817 | 0.02 | A0A1D3L0U0 | *Methanobacterium congolense* | Insertion element ISM1 uncharacterized 48,3 kDa protein |

| | | | | | |
|---|---|---|---|---|---|
| 225 | 815 | 0.02 | A0A1D3L0X2 | *Methanobacterium congolense* | Vitamin B12-dependent ribonucleotide reductase (EC 1.17.4.1) |
| 226 | 814 | 0.02 | A0A1D3L4L3 | *Methanobacterium congolense* | Trk system potassium uptake protein TrkH |
| 227 | 814 | 0.02 | A7I666 | *Methanoregula boonei* | Phenylacetate--CoA ligase (EC 6.2.1.30) |
| 228 | 814 | 0.02 | A7IAM8 | *Methanoregula boonei* | TrkA-N domain protein |
| 229 | 813 | 0.02 | A0A1D3L203 | *Methanobacterium congolense* | Uncharacterized protein |
| 230 | 813 | 0.02 | A0A1D3L3U3 | *Methanobacterium congolense* | DNA-binding protein MutS2 |
| 231 | 810 | 0.02 | A0A1D3L3M7 | *Methanobacterium congolense* | Putative glycosyltransferase YfnE (EC 2.4.-.-) |
| 232 | 810 | 0.02 | A0A1D3L5J2 | *Methanobacterium congolense* | Putative sensor histidine kinase pdtaS (EC 2.7.13.3) |
| 233 | 809 | 0.02 | A0A1D3L2F4 | *Methanobacterium congolense* | Inosine-5'-monophosphate dehydrogenase (IMP dehydrogenase) (IMPD) (IMPDH) (EC 1.1.1.205) |
| 234 | 806 | 0.02 | A7IAZ4 | *Methanoregula boonei* | Peptidase S16, Lon-like protease (EC 3.4.21.53) |
| 235 | 805 | 0.02 | A7IAB7 | *Methanoregula boonei* | D-3-phosphoglycerate dehydrogenase (EC 1.1.1.95) |
| 236 | 805 | 0.02 | A7I5I7 | *Methanoregula boonei* | GTP cyclohydrolase MptA (EC 3.5.4.39) (GTP cyclohydrolase IV) |
| 237 | 800 | 0.02 | A7I931 | *Methanoregula boonei* | GTP-binding protein, HSR1-related |
| 238 | 797 | 0.02 | A0A1D3KZR8 | *Methanobacterium congolense* | DNA ligase (EC 6.5.1.1) (Polydeoxyribonucleotide synthase [ATP]) |
| 239 | 795 | 0.02 | L0HI79 | *Methanoregula formicica* | CRISPR-associated protein Cas8c/Csd1, subtype I-C/DVULG |
| 240 | 795 | 0.02 | Q8VQM8 | *Prevotella intermedia* | Mobilization protein |
| 241 | 792 | 0.02 | E1RF47 | *Methanolacinia petrolearia* | Ammonium transporter |
| 242 | 791 | 0.02 | A0A1D3L598 | *Methanobacterium congolense* | Chaperone protein DnaK (HSP70) (Heat shock 70 kDa protein) (Heat shock protein 70) |
| 243 | 791 | 0.02 | A7I4I5 | *Methanoregula boonei* | UbiD family decarboxylase |
| 244 | 791 | 0.02 | A7I7L4 | *Methanoregula boonei* | Chaperone protein DnaK (HSP70) (Heat shock 70 kDa protein) (Heat shock protein 70) |
| 245 | 789 | 0.02 | A7IAG9 | *Methanoregula boonei* | Enolase (EC 4.2.1.11) (2-phospho-D-glycerate hydro-lyase) (2-phosphoglycerate dehydratase) |
| 246 | 788 | 0.02 | A0A1D3L4I8 | *Methanobacterium congolense* | Polyferredoxin protein MvhB |
| 247 | 787 | 0.02 | A7I648 | *Methanoregula boonei* | Uncharacterized protein |
| 248 | 787 | 0.02 | A7I800 | *Methanoregula boonei* | Putative flagellar protein FlaH |
| 249 | 786 | 0.02 | A7IB03 | *Methanoregula boonei* | Phosphate uptake regulator, PhoU |

| | | | | | |
|---|---|---|---|---|---|
| 250 | 785 | 0.02 | A0A1D3KZ36 | *Methanobacterium congolense* | O-phosphoserine--tRNA(Cys) ligase (O-phosphoserine--tRNA ligase) (EC 6.1.1.27) (Non-canonical O-phosphoseryl-tRNA(Cys) synthetase) (O-phosphoseryl-tRNA(Cys) synthetase) (SepRS) |
| 251 | 785 | 0.02 | A0A1D3L078 | *Methanobacterium congolense* | DEAD-box ATP-dependent RNA helicase CshA (EC 3.6.4.13) |
| 252 | 785 | 0.02 | A7IAV0 | *Methanoregula boonei* | V-type ATP synthase subunit C (V-ATPase subunit C) |
| 253 | 782 | 0.02 | A0A1D3KZT8 | *Methanobacterium congolense* | Acyl-coenzyme A synthetase ACSM3, mitochondrial (EC 6.2.1.2) |
| 254 | 781 | 0.02 | L0HD61 | *Methanoregula formicica* | Coenzyme F420-reducing hydrogenase, beta subunit |
| 255 | 780 | 0.02 | A7I9L0 | *Methanoregula boonei* | Type II secretion system protein E |
| 256 | 777 | 0.02 | F3ZQL8 | *Bacteroides coprosuis* | Transposase IS4 family protein |
| 257 | 776 | 0.02 | A0A1D3L4R1 | *Methanobacterium congolense* | V-type ATP synthase alpha chain (EC 3.6.3.14) (V-ATPase subunit A) |
| 258 | 775 | 0.02 | A0A1D3L2V1 | *Methanobacterium congolense* | Uncharacterized protein |
| 259 | 775 | 0.02 | A7I989 | *Methanoregula boonei* | Oxaloacetate decarboxylase alpha subunit |
| 260 | 775 | 0.02 | A7IAT4 | *Methanoregula boonei* | Lysine--tRNA ligase (EC 6.1.1.6) (Lysyl-tRNA synthetase) (LysRS) |
| 261 | 774 | 0.02 | A0A1D3L0L8 | *Methanobacterium congolense* | Signal transduction histidine kinase |
| 262 | 772 | 0.02 | A0A089ZG44 | *Methanobacterium formicicum* | Histidine kinase/GAF/PAS domain-containing protein |
| 263 | 772 | 0.02 | A0A1D3L1A5 | *Methanobacterium congolense* | Acetyl-coenzyme A synthetase (EC 6.2.1.1) |
| 264 | 772 | 0.02 | A7I6E6 | *Methanoregula boonei* | Heavy metal translocating P-type ATPase |
| 265 | 770 | 0.02 | A0A1D3L3V6 | *Methanobacterium congolense* | Methanogenesis regulatory histidine kinase FilI (EC 2.7.13.3) |
| 266 | 769 | 0.02 | A0A1D3L5A3 | *Methanobacterium congolense* | Coenzyme F420 hydrogenase subunit alpha (EC 1.12.98.1) |
| 267 | 769 | 0.02 | | | |
| 268 | 767 | 0.02 | A0A1D3KZT0 | *Methanobacterium congolense* | Adenine deaminase (Adenase) (Adenine aminase) (EC 3.5.4.2) |
| 269 | 767 | 0.02 | A7I842 | *Methanoregula boonei* | Beta-lactamase domain protein |
| 270 | 765 | 0.02 | A0A1D3KZN6 | *Methanobacterium congolense* | Global nitrogen regulator NrpR |
| 271 | 765 | 0.02 | A7I498 | *Methanoregula boonei* | DNA polymerase II small subunit (Pol II) (EC 2.7.7.7) |
| 272 | 765 | 0.02 | A7I6N3 | *Methanoregula boonei* | Amidohydrolase 3 |
| 273 | 765 | 0.02 | A7I941 | *Methanoregula boonei* | Type 2 DNA topoisomerase 6 subunit B (EC 5.99.1.3) (Type II DNA topoisomerase VI subunit B) (TopoVI-B) |
| 274 | 763 | 0.02 | A0A1D3KZQ6 | *Methanobacterium congolense* | Type 2 DNA topoisomerase 6 subunit B (EC 5.99.1.3) (Type II DNA topoisomerase VI subunit B) (TopoVI-B) |

| | | | | | |
|---|---|---|---|---|---|
| 275 | 761 | 0.02 | | | |
| 276 | 760 | 0.02 | A0A1D3L5T8 | *Methanobacterium congolense* | Putative transporter YebQ |
| 277 | 760 | 0.02 | A7IAJ9 | *Methanoregula boonei* | Carbamoyltransferase (EC 2.1.3.-) |
| 278 | 759 | 0.02 | A0A1D3L514 | *Methanobacterium congolense* | GTP-binding protein 1 |
| 279 | 758 | 0.02 | A0A1D3KZW0 | *Methanobacterium congolense* | Putative ABC transporter ATP-binding protein MJ1242 |
| 280 | 758 | 0.02 | A7I654 | *Methanoregula boonei* | Glycosyl transferase, family 39 |
| 281 | 756 | 0.02 | A0A1D3L0A1 | *Methanobacterium congolense* | UDP-N-acetylmuramoyl-L-alanyl-D-glutamate-2,6-diaminopimelate ligase (EC 6.3.2.13) |
| 282 | 756 | 0.02 | A0A1D3L5F2 | *Methanobacterium congolense* | Diaminopimelate decarboxylase (DAP decarboxylase) (DAPDC) (EC 4.1.1.20) |
| 283 | 753 | 0.02 | F6D827 | *Methanobacterium paludis* | Formylmethanofuran dehydrogenase subunit B (EC 1.2.99.5) |
| 284 | 752 | 0.02 | A0A1D3L014 | *Methanobacterium congolense* | GAF domain protein |
| 285 | 752 | 0.02 | A0A1D3L5F7 | *Methanobacterium congolense* | Putative transporter YebQ |
| 286 | 751 | 0.02 | A0A1D3L2V9 | *Methanobacterium congolense* | Uncharacterized protein |
| 287 | 750 | 0.02 | A0A1D3L095 | *Methanobacterium congolense* | Putative membrane protein |
| 288 | 749 | 0.02 | A0A089ZV91 | *Methanobacterium formicicum* | Histidine kinase/PAS domain-containing protein |
| 289 | 749 | 0.02 | A0A1D3L4K9 | *Methanobacterium congolense* | Uncharacterized protein |
| 290 | 748 | 0.02 | A7IAG1 | *Methanoregula boonei* | Glutamate--tRNA ligase (EC 6.1.1.17) (Glutamyl-tRNA synthetase) (GluRS) |
| 291 | 747 | 0.02 | A0A1D3L3I1 | *Methanobacterium congolense* | Glycine-tRNA ligase (EC 6.1.1.14) |
| 292 | 747 | 0.02 | A7I8Q2 | *Methanoregula boonei* | AAA family ATPase, CDC48 subfamily (EC 3.6.4.6) |
| 293 | 747 | 0.02 | A7I9C0 | *Methanoregula boonei* | Pyruvate carboxyltransferase |
| 294 | 746 | 0.02 | A7I487 | *Methanoregula boonei* | NADH dehydrogenase (Quinone) (EC 1.6.99.5) |
| 295 | 745 | 0.02 | A0A1D3L0W5 | *Methanobacterium congolense* | Putative membrane protein |
| 296 | 744 | 0.02 | A0A1D3L5L2 | *Methanobacterium congolense* | Uncharacterized protein |
| 297 | 744 | 0.02 | A7I4R9 | *Methanoregula boonei* | Sodium/hydrogen exchanger |
| 298 | 743 | 0.02 | A0A1D3L4I5 | *Methanobacterium congolense* | Methyl-coenzyme M reductase I subunit beta (EC 2.8.4.1) |
| 299 | 743 | 0.02 | A7I7Y9 | *Methanoregula boonei* | ABC transporter related |
| 300 | 740 | 0.02 | A0A1D3L1B1 | *Methanobacterium congolense* | Putative LRR receptor-like serine/threonine-protein kinase At1g06840 (EC 2.7.11.1) |
| 301 | 740 | 0.02 | A0A1D3L4J5 | *Methanobacterium congolense* | Uncharacterized protein |

| | | | | | |
|---|---|---|---|---|---|
| 302 | 740 | 0.02 | A0A1D3L4T1 | *Methanobacterium congolense* | Homoserine dehydrogenase (EC 1.1.1.3) |
| 303 | 739 | 0.02 | A0A1D3KZV6 | *Methanobacterium congolense* | Uncharacterized protein |
| 304 | 739 | 0.02 | A0A1D3L0W3 | *Methanobacterium congolense* | AAA ATPase |
| 305 | 739 | 0.02 | A0A1D3L554 | *Methanobacterium congolense* | Divalent metal cation transporter MntH |
| 306 | 737 | 0.02 | A0A1D3KZ40 | *Methanobacterium congolense* | Fumarate reductase (CoM/CoB) subunit A |
| 307 | 737 | 0.02 | A0A1D3L5L1 | *Methanobacterium congolense* | Multidrug resistance protein Stp |
| 308 | 737 | 0.02 | U6ECW1 | *Methanobacterium sp.* | Uncharacterized protein |
| 309 | 736 | 0.02 | A7I4F0 | *Methanoregula boonei* | Aspartokinase (EC 2.7.2.4) |
| 310 | 735 | 0.02 | A7I6C7 | *Methanoregula boonei* | AMP-dependent synthetase and ligase |
| 311 | 733 | 0.02 | A0A1D3KZ91 | *Methanobacterium congolense* | Putative glutamine amidotransferase-like protein MJ0138 |
| 312 | 732 | 0.02 | A0A1D3L220 | *Methanobacterium congolense* | Uncharacterized protein |
| 313 | 732 | 0.02 | A7I9A2 | *Methanoregula boonei* | Protein-export membrane protein SecD |
| 314 | 731 | 0.02 | A0A1D3L102 | *Methanobacterium congolense* | Putative membrane protein YabM |
| 315 | 731 | 0.02 | F6D828 | *Methanobacterium paludis* | Formylmethanofuran dehydrogenase subunit A |
| 316 | 730 | 0.02 | A0A1D3L4V5 | *Methanobacterium congolense* | Putative cation transporter MJ1485 |
| 317 | 730 | 0.02 | A7I7G0 | *Methanoregula boonei* | Beta-lactamase domain protein |
| 318 | 729 | 0.02 | A0A1D3L317 | *Methanobacterium congolense* | Uncharacterized protein |
| 319 | 728 | 0.02 | A0A1D3L068 | *Methanobacterium congolense* | Multidrug export protein EmrB |
| 320 | 728 | 0.02 | A0A1D3L3X0 | *Methanobacterium congolense* | Acetolactate synthase large subunit (EC 2.2.1.6) |
| 321 | 728 | 0.02 | A0A1D3L504 | *Methanobacterium congolense* | DNA polymerase (EC 2.7.7.7) |
| 322 | 728 | 0.02 | A7IAU5 | *Methanoregula boonei* | Phosphoglycerate kinase (EC 2.7.2.3) |
| 323 | 727 | 0.02 | A0A1D3L4A8 | *Methanobacterium congolense* | Uncharacterized protein |
| 324 | 726 | 0.02 | A0A1D3L1D7 | *Methanobacterium congolense* | Uncharacterized protein |
| 325 | 726 | 0.02 | A0A1D3L369 | *Methanobacterium congolense* | Transposase |
| 326 | 726 | 0.02 | A0A1D3L5B2 | *Methanobacterium congolense* | Putative polyferredoxin-like protein MJ1303 |
| 327 | 726 | 0.02 | A7I763 | *Methanoregula boonei* | Conserved TM helix repeat-containing protein |
| 328 | 726 | 0.02 | A7IAF0 | *Methanoregula boonei* | Major facilitator superfamily MFS_1 |

| | | | | | |
|---|---|---|---|---|---|
| 329 | 725 | 0.02 | A0A1D3L0S3 | *Methanobacterium congolense* | Single-stranded-DNA-specific exonuclease RecJ (EC 3.1.-.-) |
| 330 | 723 | 0.02 | A0A1D3L153 | *Methanobacterium congolense* | Uncharacterized protein |
| 331 | 721 | 0.02 | A7I7F3 | *Methanoregula boonei* | Nitrogenase molybdenum-iron protein beta chain (EC 1.18.6.1) |
| 332 | 720 | 0.02 | A0A1D3KZ76 | *Methanobacterium congolense* | Putative 2-isopropylmalate synthase (EC 2.3.3.13) |
| 333 | 720 | 0.02 | A0A1D3L0L3 | *Methanobacterium congolense* | Amidophosphoribosyltransferase (ATase) (EC 2.4.2.14) (Glutamine phosphoribosylpyrophosphate amidotransferase) (GPATase) |
| 334 | 720 | 0.02 | A0A1D3L243 | *Methanobacterium congolense* | CTP synthase (EC 6.3.4.2) (Cytidine 5'-triphosphate synthase) (Cytidine triphosphate synthetase) (CTP synthetase) (CTPS) (UTP--ammonia ligase) |
| 335 | 720 | 0.02 | A0A1D3L4L0 | *Methanobacterium congolense* | Uncharacterized protein |
| 336 | 720 | 0.02 | A0A1D3L5D6 | *Methanobacterium congolense* | Uncharacterized protein |
| 337 | 717 | 0.02 | A0A1D3KZL5 | *Methanobacterium congolense* | Uncharacterized protein |
| 338 | 716 | 0.02 | A7I4Y1 | *Methanoregula boonei* | Peptide chain release factor subunit 1 (Translation termination factor aRF1) |
| 339 | 715 | 0.02 | A7I754 | *Methanoregula boonei* | Homospermidine synthase (EC 2.5.1.44) |
| 340 | 714 | 0.02 | A0A1D3L546 | *Methanobacterium congolense* | Multifunctional fusion protein [Includes: ADP-dependent (S)-NAD(P)H-hydrate dehydratase (EC 4.2.1.136) (ADP-dependent NAD(P)HX dehydratase); NAD(P)H-hydrate epimerase (EC 5.1.99.6) (NAD(P)HX epimerase)] |
| 341 | 714 | 0.02 | A7I4K8 | *Methanoregula boonei* | Amino acid permease-associated region |
| 342 | 713 | 0.02 | A0A090I147 | *Methanobacterium formicicum* | DNA double-strand break repair Rad50 ATPase |
| 343 | 713 | 0.02 | A0A1D3L4C7 | *Methanobacterium congolense* | Transglutaminase domain-containing protein |
| 344 | 713 | 0.02 | A7I8Z5 | *Methanoregula boonei* | Glutamine synthetase (EC 6.3.1.2) |
| 345 | 712 | 0.02 | A0A1D3KZK0 | *Methanobacterium congolense* | Glutamyl-tRNA(Gln) amidotransferase subunit E (Glu-ADT subunit E) (EC 6.3.5.-) |
| 346 | 712 | 0.02 | A0A1D3L103 | *Methanobacterium congolense* | Ferrous iron transport protein B |
| 347 | 711 | 0.02 | A7I4C2 | *Methanoregula boonei* | Uncharacterized protein |
| 348 | 710 | 0.02 | B8GJD9 | *Methanosphaerula palustris* | Ferrous iron transport protein B |
| 349 | 709 | 0.02 | A0A1D3L4N1 | *Methanobacterium congolense* | Type-2 serine--tRNA ligase (EC 6.1.1.11) (Seryl-tRNA synthetase) (SerRS) (Seryl-tRNA(Ser/Sec) synthetase) |
| 350 | 707 | 0.02 | A0A1D3L0E2 | *Methanobacterium congolense* | Uncharacterized protein |
| 351 | 707 | 0.02 | A0A1D3L1P3 | *Methanobacterium congolense* | Putative MFS-type transporter EfpA |
| 352 | 707 | 0.02 | A0A1D3L252 | *Methanobacterium congolense* | Putative ABC transporter ATP-binding protein MJ1662 |
| 353 | 706 | 0.02 | A0A1D3L0B1 | *Methanobacterium congolense* | Probable cobyric acid synthase |

| | | | | | |
|---|---|---|---|---|---|
| 354 | 706 | 0.02 | A7I5H9 | *Methanoregula boonei* | Phosphoribosylformylglycinamidine synthase subunit PurL (FGAM synthase) (EC 6.3.5.3) (Formylglycinamide ribonucleotide amidotransferase subunit II) (FGAR amidotransferase II) (FGAR-AT II) (Glutamine amidotransferase PurL) (Phosphoribosylformylglycinamidine synthase subunit II) |
| 355 | 706 | 0.02 | A7I8Y7 | *Methanoregula boonei* | AMP-dependent synthetase and ligase |
| 356 | 705 | 0.02 | A7I860 | *Methanoregula boonei* | Putative phosphoserine phosphatase |
| 357 | 705 | 0.02 | A7I8N6 | *Methanoregula boonei* | Methyl-accepting chemotaxis sensory transducer |
| 358 | 704 | 0.02 | A0A1D3L4N6 | *Methanobacterium congolense* | Inositol-3-phosphate synthase (EC 5.5.1.4) |
| 359 | 704 | 0.02 | A0A1D3L5C5 | *Methanobacterium congolense* | Magnesium transporter MgtE |
| 360 | 703 | 0.02 | A0A1D3L4B7 | *Methanobacterium congolense* | Uncharacterized protein |
| 361 | 702 | 0.02 | A0A1D3KZK6 | *Methanobacterium congolense* | Glutamate decarboxylase (EC 4.1.1.15) |
| 362 | 702 | 0.02 | A0A1D3L3W8 | *Methanobacterium congolense* | Cation diffusion facilitator family transporter |
| 363 | 702 | 0.02 | A0A1D3L5T3 | *Methanobacterium congolense* | Phosphoribosylamine--glycine ligase (EC 6.3.4.13) (GARS) (Glycinamide ribonucleotide synthetase) (Phosphoribosylglycinamide synthetase) |
| 364 | 702 | 0.02 | A7I4W1 | *Methanoregula boonei* | Glycosyl transferase, group 1 |
| 365 | 702 | 0.02 | A7I7F2 | *Methanoregula boonei* | Nitrogenase protein alpha chain (EC 1.18.6.1) |
| 366 | 701 | 0.02 | A7I8Z7 | *Methanoregula boonei* | Archaeal glutamate synthase [NADPH] (EC 1.4.1.13) |
| 367 | 700 | 0.02 | A7I6J4 | *Methanoregula boonei* | Uncharacterized protein |
| 368 | 699 | 0.02 | A0A1D3L065 | *Methanobacterium congolense* | Multidrug export protein EmrB |
| 369 | 699 | 0.02 | A0A1D3L0C2 | *Methanobacterium congolense* | Putative lon protease homolog (EC 3.4.21.-) |
| 370 | 699 | 0.02 | A0A1D3L117 | *Methanobacterium congolense* | Uncharacterized protein |
| 371 | 698 | 0.02 | A0A1D3L1Y8 | *Methanobacterium congolense* | Putative O-antigen transporter |
| 372 | 698 | 0.02 | A0A1D3L2K6 | *Methanobacterium congolense* | Uncharacterized protein |
| 373 | 698 | 0.02 | A7I9L6 | *Methanoregula boonei* | Response regulator receiver protein |
| 374 | 698 | 0.02 | A7IA07 | *Methanoregula boonei* | Aminotransferase, class V |
| 375 | 698 | 0.02 | U6ED26 | *Methanobacterium sp.* | ORC1-type DNA replication protein |
| 376 | 697 | 0.02 | A0A1D3L230 | *Methanobacterium congolense* | UvrABC system protein B (Protein UvrB) (Excinuclease ABC subunit B) |
| 377 | 697 | 0.02 | A7I890 | *Methanoregula boonei* | Serine--tRNA ligase (EC 6.1.1.11) (Seryl-tRNA synthetase) (SerRS) (Seryl-tRNA(Ser/Sec) synthetase) |

| | | | | | |
|---|---|---|---|---|---|
| 378 | 696 | 0.02 | A0A1D3KZI2 | *Methanobacterium congolense* | Probable tRNA pseudouridine synthase D (EC 5.4.99.27) (tRNA pseudouridine(13) synthase) (tRNA pseudouridylate synthase D) (tRNA-uridine isomerase D) |
| 379 | 695 | 0.02 | A0A1D3L064 | *Methanobacterium congolense* | UDP-N-acetylmuramate-L-alanine ligase (EC 6.3.2.8) |
| 380 | 695 | 0.02 | A0A1D3L434 | *Methanobacterium congolense* | Putative sensor histidine kinase pdtaS (EC 2.7.13.3) |
| 381 | 695 | 0.02 | A7I4X0 | *Methanoregula boonei* | SufBD protein |
| 382 | 695 | 0.02 | A7I5L0 | *Methanoregula boonei* | Indolepyruvate oxidoreductase subunit IorA (IOR) (EC 1.2.7.8) (Indolepyruvate ferredoxin oxidoreductase subunit alpha) |
| 383 | 695 | 0.02 | A7I617 | *Methanoregula boonei* | Adenosylhomocysteinase (EC 3.3.1.1) |
| 384 | 693 | 0.02 | A0A089Z8E6 | *Methanobacterium formicicum* | Histidine kinase/PAS domain-containing protein |
| 385 | 693 | 0.02 | A0A1D3L510 | *Methanobacterium congolense* | Uncharacterized protein |
| 386 | 692 | 0.02 | A0A1D3KZC3 | *Methanobacterium congolense* | Glutamine synthetase (EC 6.3.1.2) |
| 387 | 691 | 0.02 | A0A1D3L1I3 | *Methanobacterium congolense* | Uncharacterized protein |
| 388 | 690 | 0.02 | A7I696 | *Methanoregula boonei* | Argininosuccinate lyase (ASAL) (EC 4.3.2.1) (Arginosuccinase) |
| 389 | 690 | 0.02 | A7I6J3 | *Methanoregula boonei* | Argininosuccinate synthase (EC 6.3.4.5) (Citrulline--aspartate ligase) |
| 390 | 686 | 0.02 | A0A1D3KZC4 | *Methanobacterium congolense* | Inner membrane transport permease YbhR |
| 391 | 686 | 0.02 | A7I8Z4 | *Methanoregula boonei* | Ammonium transporter |
| 392 | 685 | 0.02 | A0A090I3S9 | *Methanobacterium formicicum* | Putative copper-exporting P-type ATPase A |
| 393 | 685 | 0.02 | A0A1D3KZ99 | *Methanobacterium congolense* | Lysine--tRNA ligase (EC 6.1.1.6) (Lysyl-tRNA synthetase) (LysRS) |
| 394 | 685 | 0.02 | A0A1D3L1N4 | *Methanobacterium congolense* | Signal transduction histidine kinase |
| 395 | 685 | 0.02 | A7I6P1 | *Methanoregula boonei* | Carbon monoxide dehydrogenase (EC 1.2.7.4) |
| 396 | 685 | 0.02 | A7I6C9 | *Methanoregula boonei* | Bifunctional enzyme Fae/Hps [Includes: 5,6,7,8-tetrahydromethanopterin hydro-lyase (EC 4.2.1.147) (Formaldehyde-activating enzyme) (Fae); 3-hexulose-6-phosphate synthase (HPS) (EC 4.1.2.43) (D-arabino-3-hexulose-6-phosphate formaldehyde lyase)] |
| 397 | 685 | 0.02 | L0HFP5 | *Methanoregula formicica* | Magnesium chelatase, H subunit |
| 398 | 684 | 0.02 | A0A1D3L3D7 | *Methanobacterium congolense* | Sodium/pantothenate symporter |
| 399 | 684 | 0.02 | A7I861 | *Methanoregula boonei* | Cell division protein FtsZ |
| 400 | 683 | 0.02 | A0A1D3KZ61 | *Methanobacterium congolense* | 5'-deoxyadenosine deaminase (5'-dA deaminase) (EC 3.5.4.41) |
| 401 | 683 | 0.02 | A0A1D3KZT9 | *Methanobacterium congolense* | Mur ligase middle domain protein |
| 402 | 682 | 0.02 | A0A1D3L0F1 | *Methanobacterium congolense* | Uncharacterized protein |

| | | | | | |
|---|---|---|---|---|---|
| 403 | 681 | 0.02 | L0HG11 | *Methanoregula formicica* | Uncharacterized protein |
| 404 | 681 | 0.02 | | | |
| 405 | 680 | 0.02 | A0A089ZVQ3 | *Methanobacterium formicicum* | Anti-sigma factor antagonist |
| 406 | 680 | 0.02 | A0A1D3L3Q7 | *Methanobacterium congolense* | Putative methyltransferase PH0819 (EC 2.1.1.-) |
| 407 | 679 | 0.02 | A0A1D3L062 | *Methanobacterium congolense* | Putative transporter HI_0608 |
| 408 | 678 | 0.02 | A0A1D3L5I9 | *Methanobacterium congolense* | tRNA pseudouridine synthase Pus10 (EC 5.4.99.25) (tRNA pseudouridine 54/55 synthase) (Psi54/55 synthase) |
| 409 | 677 | 0.02 | A0A1D3KZ70 | *Methanobacterium congolense* | Outer membrane protein assembly factor BamB |
| 410 | 677 | 0.02 | A7IA49 | *Methanoregula boonei* | Aminotransferase, class I and II |
| 411 | 676 | 0.02 | A0A1D3L490 | *Methanobacterium congolense* | Lactaldehyde dehydrogenase (EC 1.2.1.22) |
| 412 | 675 | 0.02 | A7I9F2 | *Methanoregula boonei* | Small GTP-binding protein |
| 413 | 674 | 0.02 | A0A090JY84 | *Methanobacterium formicicum* | Transposase |
| 414 | 674 | 0.02 | A7I6B3 | *Methanoregula boonei* | ABC transporter related |
| 415 | 673 | 0.02 | A0A1D3KZ75 | *Methanobacterium congolense* | Cobyrinate a,c-diamide synthase (EC 6.3.5.11) (Cobyrinic acid a,c-diamide synthetase) |
| 416 | 673 | 0.02 | A7I7B2 | *Methanoregula boonei* | Ribulose bisphosphate carboxylase (RuBisCO) (EC 4.1.1.39) |
| 417 | 672 | 0.02 | A7I5N4 | *Methanoregula boonei* | Cell division GTPase-like protein |
| 418 | 671 | 0.02 | A7I630 | *Methanoregula boonei* | Pyruvate flavodoxin/ferredoxin oxidoreductase domain protein |
| 419 | 670 | 0.02 | A7I6E9 | *Methanoregula boonei* | Uncharacterized protein |
| 420 | 669 | 0.02 | A0A1D3KYZ1 | *Methanobacterium congolense* | HAD superfamily hydrolase |
| 421 | 669 | 0.02 | A0A1D3KZ06 | *Methanobacterium congolense* | Tryptophan synthase beta chain (EC 4.2.1.20) |
| 422 | 669 | 0.02 | A0A1D3L563 | *Methanobacterium congolense* | Putative cysteine desulfurase (EC 2.8.1.7) |
| 423 | 668 | 0.02 | A0A089Z880 | *Methanobacterium formicicum* | TrmB family transcriptional regulator |
| 424 | 668 | 0.02 | A0A1D3L3U4 | *Methanobacterium congolense* | Integrase |
| 425 | 668 | 0.02 | A7I803 | *Methanoregula boonei* | Uncharacterized protein |
| 426 | 668 | 0.02 | A7I9I1 | *Methanoregula boonei* | Signal recognition particle 54 kDa protein (SRP54) |
| 427 | 668 | 0.02 | A7IAK3 | *Methanoregula boonei* | 50S ribosomal protein L10 (Acidic ribosomal protein P0 homolog) |
| 428 | 667 | 0.02 | A0A1D3L472 | *Methanobacterium congolense* | D-3-phosphoglycerate dehydrogenase (EC 1.1.1.95) |

| | | | | | |
|---|---|---|---|---|---|
| 429 | 667 | 0.02 | A7I7F4 | *Methanoregula boonei* | Nitrogenase MoFe cofactor biosynthesis protein NifE (EC 1.18.6.1) |
| 430 | 666 | 0.02 | A7I5V0 | *Methanoregula boonei* | Formylmethanofuran--tetrahydromethanopterin formyltransferase (EC 2.3.1.101) (H4MPT formyltransferase) |
| 431 | 666 | 0.02 | A7IB58 | *Methanoregula boonei* | Tyrosine--tRNA ligase (EC 6.1.1.1) (Tyrosyl-tRNA synthetase) (TyrRS) |
| 432 | 665 | 0.02 | A0A090I4C4 | *Methanobacterium formicicum* | Uncharacterized protein |
| 433 | 665 | 0.02 | A0A1D3KZ83 | *Methanobacterium congolense* | Adenylosuccinate lyase (ASL) (EC 4.3.2.2) (Adenylosuccinase) |
| 434 | 665 | 0.02 | A0A1D3L2M9 | *Methanobacterium congolense* | Phenylalanine--tRNA ligase beta subunit (EC 6.1.1.20) (Phenylalanyl-tRNA synthetase beta subunit) (PheRS) |
| 435 | 665 | 0.02 | U6EDF8 | *Methanobacterium sp.* | UvrABC system protein A (UvrA protein) (Excinuclease ABC subunit A) |
| 436 | 664 | 0.02 | A0A089ZAN7 | *Methanobacterium formicicum* | Phosphoenolpyruvate synthase/pyruvate phosphate dikinase |
| 437 | 664 | 0.02 | A0A089ZV30 | *Methanobacterium formicicum* | Phosphoenolpyruvate synthase (PEP synthase) (EC 2.7.9.2) (Pyruvate, water dikinase) |
| 438 | 663 | 0.02 | U6EE51 | *Methanobacterium sp.* | Acetyl-CoA decarbonylase/synthase complex subunit alpha (ACDS complex subunit alpha) (EC 1.2.7.4) (ACDS complex carbon monoxide dehydrogenase subunit alpha) (ACDS CODH subunit alpha) |
| 439 | 660 | 0.02 | A0A089ZVC2 | *Methanobacterium formicicum* | CoB--CoM heterodisulfide reductase subunit A HdrA1 (Methyl-viologen-reducing hydrogenase subunit delta) |
| 440 | 660 | 0.02 | A7I7M3 | *Methanoregula boonei* | DNA mismatch repair protein MutS |
| 441 | 659 | 0.02 | A0A1D3KZ63 | *Methanobacterium congolense* | Bacterial Ig-like domain (Group 3) |
| 442 | 659 | 0.02 | A0A1D3L3L2 | *Methanobacterium congolense* | Putative iron-sulfur protein MMP1067 |
| 443 | 659 | 0.02 | A0A1D3L4E2 | *Methanobacterium congolense* | RNA-metabolising metallo-beta-lactamase |
| 444 | 659 | 0.02 | U6EFC2 | *Methanobacterium sp.* | Helicase |
| 445 | 658 | 0.02 | O27035 | *Methanothermobacter thermautotrophicus* | V-type ATP synthase beta chain (V-ATPase subunit B) |
| 446 | 657 | 0.02 | A0A090JTH0 | *Methanobacterium formicicum* | Uncharacterized protein |
| 447 | 657 | 0.02 | A0A1D3L3Q2 | *Methanobacterium congolense* | Argininosuccinate lyase (ASAL) (EC 4.3.2.1) (Arginosuccinase) |
| 448 | 657 | 0.02 | A7I4B6 | *Methanoregula boonei* | Aminotransferase, class V |
| 449 | 656 | 0.02 | A0A1D3KZD9 | *Methanobacterium congolense* | Uncharacterized protein |
| 450 | 655 | 0.02 | A0A1D3L3K2 | *Methanobacterium congolense* | F420-non-reducing hydrogenase vhc subunit A (EC 1.12.99.-) |
| 451 | 655 | 0.02 | A0A1D3L4Y2 | *Methanobacterium congolense* | Multidrug export protein EmrB |
| 452 | 654 | 0.02 | L0HF89 | *Methanoregula formicica* | Formylmethanofuran dehydrogenase subunit B |

| | | | | | |
|---|---|---|---|---|---|
| 453 | 653 | 0.02 | A0A1D3L5K3 | *Methanobacterium congolense* | Putative NADH oxidase (EC 1.6.99.3) |
| 454 | 652 | 0.02 | A0A1D3L2P8 | *Methanobacterium congolense* | Polysaccharide biosynthesis protein |
| 455 | 651 | 0.02 | A0A1D3KZS3 | *Methanobacterium congolense* | Ketoisovalerate oxidoreductase subunit VorA (EC 1.-.-.-) |
| 456 | 651 | 0.02 | A0A1D3L3Y3 | *Methanobacterium congolense* | Signal recognition particle receptor FtsY (SRP receptor) |
| 457 | 650 | 0.02 | A0A1D3L131 | *Methanobacterium congolense* | 7,8-dihydro-6-hydroxymethylpterin dimethyltransferase (EC 2.1.1.-) |
| 458 | 650 | 0.02 | A0A1D3L1Q0 | *Methanobacterium congolense* | Thermosome subunit alpha |
| 459 | 648 | 0.02 | A0A1D3L0Z3 | *Methanobacterium congolense* | Thermosome subunit beta |
| 460 | 648 | 0.02 | A7I922 | *Methanoregula boonei* | Inosine-5'-monophosphate dehydrogenase (IMP dehydrogenase) (IMPD) (IMPDH) (EC 1.1.1.205) |
| 461 | 648 | 0.02 | A7I942 | *Methanoregula boonei* | Type 2 DNA topoisomerase 6 subunit A (EC 5.99.1.3) (Type II DNA topoisomerase VI subunit A) |
| 462 | 647 | 0.02 | A0A1D3L414 | *Methanobacterium congolense* | Uncharacterized protein |
| 463 | 646 | 0.02 | A7I8B3 | *Methanoregula boonei* | Ppx/GppA phosphatase |
| 464 | 646 | 0.02 | | | |
| 465 | 645 | 0.02 | A0A1D2WDD6 | *Methanobacterium sp.* | Nitrogenase iron-molybdenum cofactor biosynthesis protein NifN |
| 466 | 645 | 0.02 | A7I4T9 | *Methanoregula boonei* | Anthranilate synthase component 1 (EC 4.1.3.27) |
| 467 | 644 | 0.02 | A7IAL8 | *Methanoregula boonei* | Cell division protein FtsZ |
| 468 | 643 | 0.02 | A0A090I3K9 | *Methanobacterium formicicum* | Nitrogenase beta subunit NifK (Nitrogenase molybdenum-iron protein beta chain) (EC 1.18.6.1) |
| 469 | 642 | 0.02 | A7I5S3 | *Methanoregula boonei* | Tetrahydromethanopterin S-methyltransferase, MtrH subunit |
| 470 | 642 | 0.02 | A7I928 | *Methanoregula boonei* | Uncharacterized protein |
| 471 | 641 | 0.02 | A0A1D3KZK7 | *Methanobacterium congolense* | 2-oxoglutarate synthase subunit KorA (EC 1.2.7.3) |
| 472 | 640 | 0.02 | A7I4L8 | *Methanoregula boonei* | Thiolase |
| 473 | 639 | 0.02 | A0A1D3L1H8 | *Methanobacterium congolense* | Uncharacterized protein |
| 474 | 639 | 0.02 | A0A1D3L4R5 | *Methanobacterium congolense* | Putative sensor histidine kinase pdtaS (EC 2.7.13.3) |
| 475 | 638 | 0.02 | A0A090I199 | *Methanobacterium formicicum* | Uncharacterized protein |
| 476 | 636 | 0.02 | A7I4V1 | *Methanoregula boonei* | Probable bifunctional tRNA threonylcarbamoyladenosine biosynthesis protein [Includes: Serine/threonine-protein kinase Bud32 (EC 2.7.11.1); tRNA N6-adenosine threonylcarbamoyltransferase (EC 2.3.1.234) (tRNA threonylcarbamoyladenosine |

| | | | | | |
|---|---|---|---|---|---|
| | | | | | biosynthesis protein Kae1) (t(6)A37 threonylcarbamoyladenosine biosynthesis protein Kae1)] |
| 477 | 635 | 0.02 | A0A090I2N5 | *Methanobacterium formicicum* | Putative secreted protein |
| 478 | 635 | 0.02 | A0A1D3L0L5 | *Methanobacterium congolense* | Multidrug resistance protein Stp |
| 479 | 635 | 0.02 | A0A1D3L526 | *Methanobacterium congolense* | Uncharacterized protein |
| 480 | 635 | 0.02 | A7I5X5 | *Methanoregula boonei* | Uncharacterized protein |
| 481 | 635 | 0.02 | A7I8W3 | *Methanoregula boonei* | Uncharacterized protein |
| 482 | 634 | 0.02 | A0A1D3KZA6 | *Methanobacterium congolense* | Phenylacetate-coenzyme A ligase (EC 6.2.1.30) |
| 483 | 634 | 0.02 | A0A1D3L4S7 | *Methanobacterium congolense* | Glucose-methanol-choline oxidoreductase |
| 484 | 634 | 0.02 | A7IB51 | *Methanoregula boonei* | Elongation factor Tu, domain 2 protein |
| 485 | 634 | 0.02 | F6D5H0 | *Methanobacterium paludis* | DNA repair and recombination protein RadA |
| 486 | 633 | 0.02 | A0A1D3KZ90 | *Methanobacterium congolense* | Phosphomethylpyrimidine synthase (EC 4.1.99.17) (Hydroxymethylpyrimidine phosphate synthase) (HMP-P synthase) (HMP-phosphate synthase) (HMPP synthase) (Thiamine biosynthesis protein ThiC) |
| 487 | 633 | 0.02 | A0A1D3L3N9 | *Methanobacterium congolense* | Putative methyltransferase PH0819 (EC 2.1.1.-) |
| 488 | 633 | 0.02 | A0A1D3L571 | *Methanobacterium congolense* | Serine hydroxymethyltransferase (SHMT) (Serine methylase) (EC 2.1.2.-) |
| 489 | 632 | 0.02 | A0A089ZUV6 | *Methanobacterium formicicum* | Uncharacterized protein |
| 490 | 632 | 0.02 | A0A090I3G7 | *Methanobacterium formicicum* | CoB--CoM heterodisulfide reductase subunit A HdrA2 (CoB-CoM heterodisulfide reductase iron-sulfur subunit A) (EC 1.8.98.1) |
| 491 | 632 | 0.02 | A0A1D3L152 | *Methanobacterium congolense* | Putative threonylcarbamoyladenosine tRNA methylthiotransferase (EC 2.8.4.5) |
| 492 | 631 | 0.02 | A7I5J3 | *Methanoregula boonei* | AAA ATPase, central domain protein |
| 493 | 630 | 0.02 | A0A1D3L0F7 | *Methanobacterium congolense* | DNA double-strand break repair Rad50 ATPase |
| 494 | 630 | 0.02 | A0A1D3L3I9 | *Methanobacterium congolense* | Aminotransferase (EC 2.6.1.-) |
| 495 | 630 | 0.02 | A0A1D3L3J9 | *Methanobacterium congolense* | Replication factor C large subunit (RFC large subunit) (Clamp loader large subunit) |
| 496 | 630 | 0.02 | A0A1D3L557 | *Methanobacterium congolense* | Polysaccharide pyruvyl transferase |
| 497 | 629 | 0.02 | A0A1D3L4Y5 | *Methanobacterium congolense* | Putative NOP5 family protein MJ0694 |
| 498 | 628 | 0.02 | L0HKB8 | *Methanoregula formicica* | Uncharacterized protein |
| 499 | 627 | 0.02 | A0A1D3L0I6 | *Methanobacterium congolense* | Putative methyltransferase PH0819 (EC 2.1.1.-) |
| 500 | 627 | 0.02 | A0A1D3L0M7 | *Methanobacterium congolense* | Putative MFS-type transporter YusP |

**Table S4c.** Functional overview of M1P community (top 500 rank only).

| Rank | Count | Abundance | ID | Organism | Protein Names |
|---|---|---|---|---|---|
| 1 | 2377 | 0.09 | A0A023UP33 | *Magnusiomyces ingens* | Uncharacterized protein (Fragment) |
| 2 | 1524 | 0.06 | A0A023UP27 | *Magnusiomyces ingens* | Uncharacterized protein |
| 3 | 1277 | 0.05 | A0A1Q3M6H6 | *Bacteroidales bacterium* | CRISPR-associated endonuclease Cas9 (EC 3.1.-.-) |
| 4 | 957 | 0.03 | E4T3F9 | *Paludibacter propionicigenes* | Glutamate synthase (NADH) large subunit (EC 1.4.1.14) (EC 1.4.7.1) |
| 5 | 942 | 0.03 | I9R3U4 | *Bacteroides cellulosilyticus* | Uncharacterized protein |
| 6 | 926 | 0.03 | A0A023UMT3 | *Magnusiomyces ingens* | Uncharacterized protein (Fragment) |
| 7 | 913 | 0.03 | Q8VQM8 | *Prevotella intermedia* | Mobilization protein |
| 8 | 858 | 0.03 | A0A023UMS7 | *Magnusiomyces ingens* | NADH-ubiquinone oxidoreductase chain 5 (EC 1.6.5.3) |
| 9 | 815 | 0.03 | A0A1J5AN19 | *Bacteroidetes bacterium* | IS110 family transposase |
| 10 | 790 | 0.03 | K9L3G6 | *Magnusiomyces magnusii* | Cytochrome b |
| 11 | 765 | 0.03 | E4T8J6 | *Paludibacter propionicigenes* | Cell surface receptor IPT/TIG domain protein |
| 12 | 680 | 0.02 | E4T5D1 | *Paludibacter propionicigenes* | Methionine--tRNA ligase (EC 6.1.1.10) (Methionyl-tRNA synthetase) (MetRS) |
| 13 | 661 | 0.02 | A0A023UP41 | *Magnusiomyces ingens* | Cytochrome c oxidase subunit 1 (EC 1.9.3.1) |
| 14 | 651 | 0.02 | A0A023UPJ4 | *Magnusiomyces ingens* | Uncharacterized protein (Fragment) |
| 15 | 644 | 0.02 | G0IXG6 | *Cyclobacterium marinum* | Transposase IS116/IS110/IS902 family protein |
| 16 | 644 | 0.02 | | | |
| 17 | 630 | 0.02 | A0A1L9GVN0 | *Cand. Micrarchaeum acidiphilum* | Uncharacterized protein |
| 18 | 616 | 0.02 | F0FB90 | *Prevotella multiformis* | Initiator RepB protein (Fragment) |
| 19 | 611 | 0.02 | A0A1T5HUH3 | *Alkalitalea saponilacus* | Uncharacterized protein |
| 20 | 606 | 0.02 | A0A023UM92 | *Magnusiomyces ingens* | Uncharacterized protein |
| 21 | 600 | 0.02 | F3ZQL8 | *Bacteroides coprosuis* | Transposase IS4 family protein |
| 22 | 598 | 0.02 | A0A1M3DNL6 | *Bacteroidales bacterium* | IS4 family transposase |
| 23 | 581 | 0.02 | A0A0D2WNJ4 | *Capsaspora owczarzaki* | Paramyosin |
| 24 | 581 | 0.02 | A0A1Q3M3N2 | *Bacteroidales bacterium* | Uncharacterized protein (Fragment) |
| 25 | 577 | 0.02 | | | |

| | | | | | |
|---|---|---|---|---|---|
| 26 | 561 | 0.02 | F0FB88 | *Prevotella multiformis* | Putative plasmid recombination enzyme |
| 27 | 549 | 0.02 | A0A060RAF8 | *Mucinivorans hirudinis* | Mobile element protein |
| 28 | 539 | 0.02 | A0A0D8Y6E2 | *Dictyocaulus viviparus* | Uncharacterized protein |
| 29 | 538 | 0.02 | A0A0U1CQ28 | *Chlamydia trachomatis* | Replication protein |
| 30 | 509 | 0.02 | | | |
| 31 | 502 | 0.02 | A0A1J5HM76 | *Porphyromonadaceae bacterium* | Multidrug transporter AcrB |
| 32 | 498 | 0.02 | E4T5D5 | *Paludibacter propionicigenes* | DNA-directed DNA polymerase (EC 2.7.7.7) |
| 33 | 490 | 0.02 | A0A023UP13 | *Magnusiomyces ingens* | NADH-ubiquinone oxidoreductase chain 4 (EC 1.6.5.3) |
| 34 | 471 | 0.02 | A0A023UM78 | *Magnusiomyces ingens* | NADH dehydrogenase subunit 2 (EC 1.6.5.3) |
| 35 | 463 | 0.02 | A0A1J5HWN1 | *Porphyromonadaceae bacterium* | Uncharacterized protein |
| 36 | 452 | 0.02 | E4T5Y5 | *Paludibacter propionicigenes* | Multifunctional fusion protein [Includes: Protein translocase subunit SecD; Protein-export membrane protein SecF] |
| 37 | 448 | 0.02 | E4T072 | *Paludibacter propionicigenes* | Pyruvate, phosphate dikinase (EC 2.7.9.1) |
| 38 | 436 | 0.02 | A0A1Q3M3K5 | *Bacteroidales bacterium* | Alpha-mannosidase |
| 39 | 431 | 0.02 | A0A0L8VCE1 | *Sunxiuqinia dokdonensis* | Multidrug transporter AcrB |
| 40 | 430 | 0.02 | A0A023UNG0 | *Magnusiomyces ingens* | Uncharacterized protein (Fragment) |
| 41 | 425 | 0.02 | A0A1M3MCY6 | *Paludibacter sp.* | Glycosyl hydrolase |
| 42 | 423 | 0.02 | A0A1J5HGG6 | *Porphyromonadaceae bacterium* | DNA-directed RNA polymerase subunit beta (RNAP subunit beta) (EC 2.7.7.6) (RNA polymerase subunit beta) (Transcriptase subunit beta) |
| 43 | 420 | 0.02 | A0A1Q3LCT2 | *Bacteroidales bacterium* | Cell filamentation protein Fic |
| 44 | 420 | 0.02 | W0JMT5 | *Desulfurella acetivorans* | Uncharacterized protein |
| 45 | 419 | 0.02 | A0A142L3A0 | *Bacteroidetes bacterium* | DEAD/DEAH box helicase |
| 46 | 419 | 0.02 | A0A1M3MBI7 | *Paludibacter sp.* | Cell surface protein SprA |
| 47 | 418 | 0.02 | Q8VQM9 | *Prevotella intermedia* | Replication protein |
| 48 | 415 | 0.02 | A0A0U1CQ01 | *Chlamydia trachomatis* | Uncharacterised protein |
| 49 | 413 | 0.01 | | | |
| 50 | 405 | 0.01 | F3PIP2 | *Bacteroides clarus* | TonB-dependent receptor plug domain protein |
| 51 | 403 | 0.01 | A0A1Q3M633 | *Bacteroidales bacterium* | ATP-dependent helicase |

| | | | | | |
|---|---|---|---|---|---|
| 52 | 401 | 0.01 | E4T570 | *Paludibacter propionicigenes* | DNA-directed RNA polymerase subunit beta (RNAP subunit beta) (EC 2.7.7.6) (RNA polymerase subunit beta) (Transcriptase subunit beta) |
| 53 | 400 | 0.01 | A0A0T5VPL4 | *Pedobacter ginsenosidimutans* | Uncharacterized protein |
| 54 | 399 | 0.01 | A0A1Q3M5E8 | *Bacteroidales bacterium* | Uncharacterized protein (Fragment) |
| 55 | 398 | 0.01 | E4T450 | *Paludibacter propionicigenes* | UvrABC system protein A |
| 56 | 398 | 0.01 | N8ZVC9 | *Acinetobacter gerneri* | Uncharacterized protein |
| 57 | 395 | 0.01 | U6RAE1 | *Bacteroides sp.* | Uncharacterized protein |
| 58 | 394 | 0.01 | E4T0X8 | *Paludibacter propionicigenes* | DNA helicase (EC 3.6.4.12) |
| 59 | 390 | 0.01 | A0A1M3DJX9 | *Bacteroidales bacterium* | Beta-galactosidase (EC 3.2.1.23) (Lactase) |
| 60 | 390 | 0.01 | A0A1Q3M609 | *Bacteroidales bacterium* | Uncharacterized protein |
| 61 | 388 | 0.01 | A0A0Q7S8L4 | *Pelomonas sp.* | Transposase |
| 62 | 387 | 0.01 | E4T2R0 | *Paludibacter propionicigenes* | ATPase AAA-2 domain protein |
| 63 | 384 | 0.01 | A0A1J5HA84 | *Porphyromonadaceae bacterium* | Potassium transporter |
| 64 | 383 | 0.01 | A0A1M3DC14 | *Bacteroidales bacterium* | Formate C-acetyltransferase |
| 65 | 382 | 0.01 | A0A023UPL4 | *Magnusiomyces ingens* | Uncharacterized protein (Fragment) |
| 66 | 379 | 0.01 | A0A016UCK1 | *Ancylostoma ceylanicum* | Uncharacterized protein |
| 67 | 378 | 0.01 | A0A0T5VPN0 | *Pedobacter ginsenosidimutans* | SusC/RagA family TonB-linked outer membrane protein |
| 68 | 377 | 0.01 | E4T3F8 | *Paludibacter propionicigenes* | L-glutamine synthetase (EC 6.3.1.2) |
| 69 | 375 | 0.01 | E4T1A7 | *Paludibacter propionicigenes* | Pyruvate phosphate dikinase PEP/pyruvate-binding protein |
| 70 | 364 | 0.01 | E4T5C1 | *Paludibacter propionicigenes* | DNA topoisomerase IV subunit A (EC 5.99.1.3) |
| 71 | 363 | 0.01 | A0A1R3T9L6 | *Proteiniphilum saccharofermentans* | Transposase IS4 family protein |
| 72 | 360 | 0.01 | A0A1J5HML6 | *Porphyromonadaceae bacterium* | Penicillin-binding protein |
| 73 | 360 | 0.01 | E4T3R4 | *Paludibacter propionicigenes* | AMP-dependent synthetase and ligase |
| 74 | 359 | 0.01 | E4T571 | *Paludibacter propionicigenes* | DNA-directed RNA polymerase subunit beta' (RNAP subunit beta') (EC 2.7.7.6) (RNA polymerase subunit beta') (Transcriptase subunit beta') |
| 75 | 358 | 0.01 | A0A1M3MAE9 | *Paludibacter sp.* | DNA-directed RNA polymerase subunit beta' (RNAP subunit beta') (EC 2.7.7.6) (RNA polymerase subunit beta') (Transcriptase subunit beta') |
| 76 | 358 | 0.01 | W1I7F0 | *uncultured bacterium* | Uncultured bacterium extrachromosomal DNA RGI01790 |

| | | | | | |
|---|---|---|---|---|---|
| 77 | 357 | 0.01 | E4T8D5 | *Paludibacter propionicigenes* | Orn/DAP/Arg decarboxylase 2 |
| 78 | 355 | 0.01 | A0A1E4ELA1 | *Paludibacter sp.* | Dihydrodipicolinate synthase family protein |
| 79 | 352 | 0.01 | E4T3X9 | *Paludibacter propionicigenes* | DNA topoisomerase III (EC 5.99.1.2) |
| 80 | 351 | 0.01 | A0A1J5HT60 | *Porphyromonadaceae bacterium* | DNA polymerase I (EC 2.7.7.7) |
| 81 | 351 | 0.01 | A0A1M3DNS5 | *Bacteroidales bacterium* | Putative K(+)-stimulated pyrophosphate-energized sodium pump (EC 3.6.1.1) (Membrane-bound sodium-translocating pyrophosphatase) (Pyrophosphate-energized inorganic pyrophosphatase) (Na(+)-PPase) |
| 82 | 350 | 0.01 | R6DEG5 | *Bacteroides sp.* | RagB/SusD domain-containing protein |
| 83 | 349 | 0.01 | A0A1Q3LBC0 | *Bacteroidales bacterium* | Uncharacterized protein |
| 84 | 347 | 0.01 | E4T653 | *Paludibacter propionicigenes* | Peptidoglycan glycosyltransferase (EC 2.4.1.129) |
| 85 | 342 | 0.01 | E4T2N1 | *Paludibacter propionicigenes* | DNA gyrase subunit A (EC 5.99.1.3) |
| 86 | 341 | 0.01 | K9L3F4 | *Magnusiomyces magnusii* | NADH-ubiquinone oxidoreductase chain 1 (EC 1.6.5.3) |
| 87 | 340 | 0.01 | E4T1U6 | *Paludibacter propionicigenes* | Carbamoyl-phosphate synthase L chain ATP-binding protein (EC 6.4.1.1) |
| 88 | 339 | 0.01 | E4T5N8 | *Paludibacter propionicigenes* | UvrABC system protein A |
| 89 | 339 | 0.01 | F4GG80 | *Alicycliphilus denitrificans* | Transposase IS4 family protein |
| 90 | 338 | 0.01 | E4T2H4 | *Paludibacter propionicigenes* | Phosphoribosylformylglycinamidine synthase (FGAM synthase) (FGAMS) (EC 6.3.5.3) (Formylglycinamide ribonucleotide amidotransferase) (FGAR amidotransferase) (FGAR-AT) |
| 91 | 336 | 0.01 | A0A023UMR4 | *Magnusiomyces ingens* | NADH-ubiquinone oxidoreductase chain 3 (EC 1.6.5.3) |
| 92 | 335 | 0.01 | A0A1M3M9Z9 | *Paludibacter sp.* | Uncharacterized protein |
| 93 | 333 | 0.01 | A0A023UP21 | *Magnusiomyces ingens* | Ribosomal protein S3 |
| 94 | 327 | 0.01 | A0A1J5H765 | *Porphyromonadaceae bacterium* | Multifunctional fusion protein [Includes: Protein translocase subunit SecD; Protein-export membrane protein SecF] |
| 95 | 327 | 0.01 | A0A1J5HIK6 | *Porphyromonadaceae bacterium* | UDP-N-acetylglucosamine 2-epimerase |
| 96 | 327 | 0.01 | E4T7X4 | *Paludibacter propionicigenes* | Alanine--tRNA ligase (EC 6.1.1.7) (Alanyl-tRNA synthetase) (AlaRS) |
| 97 | 326 | 0.01 | A0A1J5HBV2 | *Porphyromonadaceae bacterium* | Uncharacterized protein |
| 98 | 325 | 0.01 | A0A0E9G9S8 | *Chlamydia trachomatis* | Uncharacterised protein |
| 99 | 322 | 0.01 | E4T682 | *Paludibacter propionicigenes* | UvrABC system protein C (Protein UvrC) (Excinuclease ABC subunit C) |
| 100 | 318 | 0.01 | | | |

| | | | | | |
|---|---|---|---|---|---|
| 101 | 316 | 0.01 | E4T2W8 | *Paludibacter propionicigenes* | Pyruvate carboxylase (EC 6.4.1.1) |
| 102 | 315 | 0.01 | A0A085EMB6 | *Flavobacterium gilvum* | Uncharacterized protein |
| 103 | 315 | 0.01 | A0A1J5HRY1 | *Porphyromonadaceae bacterium* | Cell surface protein SprA |
| 104 | 315 | 0.01 | A0A1J5I0W6 | *Porphyromonadaceae bacterium* | DNA mismatch repair protein MutS |
| 105 | 315 | 0.01 | E4T5Y6 | *Paludibacter propionicigenes* | Two component regulator three Y domain-containing protein |
| 106 | 314 | 0.01 | A0A1F3LXP2 | *Bacteroidetes bacterium* | Lon protease (EC 3.4.21.53) (ATP-dependent protease La) |
| 107 | 314 | 0.01 | E4T3R5 | *Paludibacter propionicigenes* | Phenylalanine--tRNA ligase beta subunit (EC 6.1.1.20) (Phenylalanyl-tRNA synthetase beta subunit) (PheRS) |
| 108 | 314 | 0.01 | E4T409 | *Paludibacter propionicigenes* | Mg chelatase, subunit ChlI |
| 109 | 314 | 0.01 | E4T5Q6 | *Paludibacter propionicigenes* | Uncharacterized protein |
| 110 | 312 | 0.01 | A0A133PWA7 | *Prevotella corporis* | Transposase |
| 111 | 312 | 0.01 | A0A161LEB7 | *Paludibacter jiangxiensis* | RHH-type transcriptional regulator |
| 112 | 312 | 0.01 | E4T592 | *Paludibacter propionicigenes* | Methionine synthase (B12-dependent) (EC 2.1.1.13) |
| 113 | 310 | 0.01 | A0A1J5HSL1 | *Porphyromonadaceae bacterium* | Pyruvate-flavodoxin oxidoreductase (EC 1.2.7.-) |
| 114 | 310 | 0.01 | K9L3A6 | *Porphyromonadaceae bacterium* | NADH-ubiquinone oxidoreductase chain 5 (EC 1.6.5.3) |
| 115 | 308 | 0.01 | E4T4N5 | *Paludibacter propionicigenes* | 2-succinyl-5-enolpyruvyl-6-hydroxy-3-cyclohexene-1-carboxylate synthase (SEPHCHC synthase) (EC 2.2.1.9) (Menaquinone biosynthesis protein MenD) |
| 116 | 306 | 0.01 | A0A1B1ICB8 | *Prevotella scopos* | Uncharacterized protein |
| 117 | 300 | 0.01 | A0A1Q3M5Y6 | *Bacteroidales bacterium* | Uncharacterized protein |
| 118 | 300 | 0.01 | E4T1S7 | *Paludibacter propionicigenes* | Carbamoyl-phosphate synthase small chain (EC 6.3.5.5) (Carbamoyl-phosphate synthetase glutamine chain) |
| 119 | 300 | 0.01 | | | |
| 120 | 298 | 0.01 | A0A1J5HVD9 | *Porphyromonadaceae bacterium* | Citrate synthase |
| 121 | 297 | 0.01 | A0A1J5HFB3 | *Porphyromonadaceae bacterium* | L-fucose isomerase (FucIase) (EC 5.3.1.25) (6-deoxy-L-galactose isomerase) |
| 122 | 295 | 0.01 | K9L3A9 | *Magnusiomyces magnusii* | ATP synthase subunit a |
| 123 | 291 | 0.01 | A7AI47 | *Parabacteroides merdae* | Uncharacterized protein |
| 124 | 290 | 0.01 | A0A1J5H7V2 | *Porphyromonadaceae bacterium* | Collagen-binding protein (Fragment) |
| 125 | 290 | 0.01 | U6RCL0 | *Bacteroides sp.* | Uncharacterized protein |

| | | | | | |
|---|---|---|---|---|---|
| 126 | 289 | 0.01 | E4T347 | *Paludibacter propionicigenes* | Uncharacterized protein |
| 127 | 288 | 0.01 | E4T676 | *Paludibacter propionicigenes* | Collagenase |
| 128 | 288 | 0.01 | | | |
| 129 | 286 | 0.01 | E4T386 | *Paludibacter propionicigenes* | Heterodimeric methylmalonyl-CoA mutase large subunit (EC 5.4.99.2) |
| 130 | 286 | 0.01 | N8ZNX3 | *Acinetobacter gerneri* | Uncharacterized protein |
| 131 | 285 | 0.01 | A0A1M3MAC5 | *Paludibacter sp.* | Pyruvate kinase (EC 2.7.1.40) |
| 132 | 284 | 0.01 | A0A1J5HP94 | *Porphyromonadaceae bacterium* | UPF0313 protein AUK44_04705 |
| 133 | 284 | 0.01 | C9MTU8 | *Prevotella veroralis* | Initiator RepB protein |
| 134 | 284 | 0.01 | | | |
| 135 | 283 | 0.01 | A0A1Q3LZZ0 | *Bacteroidales bacterium* | Valine--tRNA ligase (EC 6.1.1.9) (Valyl-tRNA synthetase) (ValRS) |
| 136 | 283 | 0.01 | E4T683 | *Paludibacter propionicigenes* | tRNA uridine 5-carboxymethylaminomethyl modification enzyme MnmG (Glucose-inhibited division protein A) |
| 137 | 283 | 0.01 | | | |
| 138 | 277 | 0.01 | A0A1J5HUJ2 | *Porphyromonadaceae bacterium* | Cytochrome C oxidase assembly protein |
| 139 | 277 | 0.01 | | | |
| 140 | 276 | 0.01 | A0A1T4JPM0 | *Sediminibacterium ginsengisoli* | DGQHR domain-containing protein |
| 141 | 276 | 0.01 | E4T225 | *Paludibacter propionicigenes* | Allosteric NADP-dependent malic enzyme (EC 1.1.1.40) |
| 142 | 274 | 0.01 | E4T4E1 | *Paludibacter propionicigenes* | DNA-binding protein Fis |
| 143 | 273 | 0.01 | E4T2V1 | *Paludibacter propionicigenes* | 4-alpha-glucanotransferase (EC 2.4.1.25) |
| 144 | 273 | 0.01 | R6D4I0 | *Bacteroides sp.* | Uncharacterized protein |
| 145 | 273 | 0.01 | | | |
| 146 | 272 | 0.01 | A0A1J5HLR1 | *Porphyromonadaceae bacterium* | Carbamoyl-phosphate synthase (glutamine-hydrolyzing) (EC 6.3.5.5) |
| 147 | 272 | 0.01 | E4T4E6 | *Paludibacter propionicigenes* | Outer membrane protein assembly complex, YaeT protein |
| 148 | 270 | 0.01 | A0A0P0F9C5 | *Bacteroides thetaiotaomicron* | 2-dehydro-3-deoxygluconokinase (EC 2.7.1.45) (Kinase, PfkB family) (PfkB domain protein) (EC 2.7.1.92) |
| 149 | 269 | 0.01 | A0A179DEU4 | *Pedobacter sp.* | DNA methyltransferase |
| 150 | 269 | 0.01 | A0A1J5HHU5 | *Porphyromonadaceae bacterium* | Rod shape-determining protein RodA |
| 151 | 269 | 0.01 | E4T4K6 | *Paludibacter propionicigenes* | Peptidoglycan glycosyltransferase (EC 2.4.1.129) |

| | | | | | |
|---|---|---|---|---|---|
| 152 | 268 | 0.01 | E4T3B7 | *Paludibacter propionicigenes* | DNA ligase (EC 6.5.1.2) (Polydeoxyribonucleotide synthase [NAD(+)]) |
| 153 | 268 | 0.01 | Q45FH5 | *Adineta vaga* | Transposase |
| 154 | 267 | 0.01 | A0A1Q3KSP0 | *Bacteroidales bacterium* | Uncharacterized protein (Fragment) |
| 155 | 267 | 0.01 | A0A1Q3QBX8 | *Bacteroidia bacterium* | Uncharacterized protein |
| 156 | 267 | 0.01 | I8Y0K1 | *Bacteroides salyersiae* | SusC/RagA family TonB-linked outer membrane protein |
| 157 | 266 | 0.01 | A0A1F3ME11 | *Bacteroidetes bacterium* | Cadmium-translocating P-type ATPase |
| 158 | 265 | 0.01 | A0A1E4EKR5 | *Paludibacter sp.* | L-fucose:H+ symporter permease |
| 159 | 265 | 0.01 | A0A1J5HNC6 | *Porphyromonadaceae bacterium* | Cation/H(+) antiporter |
| 160 | 265 | 0.01 | A0A1M3M9F4 | *Paludibacter sp.* | Uncharacterized protein |
| 161 | 265 | 0.01 | E4T1V0 | *Paludibacter propionicigenes* | Alanine racemase (EC 5.1.1.1) |
| 162 | 264 | 0.01 | A0A170Y7T4 | *Paludibacter jiangxiensis* | Formate-dependent phosphoribosylglycinamide formyltransferase (5'-phosphoribosylglycinamide transformylase 2) (Formate-dependent GAR transformylase) (EC 2.1.2.-) (GAR transformylase 2) (GART 2) (Non-folate glycinamide ribonucleotide transformylase) (Phosphoribosylglycinamide formyltransferase 2) |
| 163 | 263 | 0.01 | A0A1J5HSK7 | *Porphyromonadaceae bacterium* | Peptidase S41 |
| 164 | 263 | 0.01 | E4T6Q8 | *Paludibacter propionicigenes* | Major facilitator superfamily MFS_1 |
| 165 | 263 | 0.01 | E4T6Z1 | *Paludibacter propionicigenes* | Exonuclease RecJ (EC 3.1.-.-) |
| 166 | 262 | 0.01 | E4T5B8 | *Paludibacter propionicigenes* | Translation elongation factor 2 (EF-2/EF-G) |
| 167 | 260 | 0.01 | C7X5L7 | *Parabacteroides sp.* | Glycosyl hydrolase, family 31 (EC 3.2.1.-) |
| 168 | 258 | 0.01 | A0A1M3M7V7 | *Paludibacter sp.* | Carbamoyl-phosphate synthase (glutamine-hydrolyzing) (EC 6.3.5.5) |
| 169 | 258 | 0.01 | E4T6D1 | *Paludibacter propionicigenes* | Galactokinase (EC 2.7.1.6) (Galactose kinase) |
| 170 | 253 | 0.01 | A0A1J5HP18 | *Porphyromonadaceae bacterium* | Cytochrome ubiquinol oxidase subunit I |
| 171 | 253 | 0.01 | E4T529 | *Paludibacter propionicigenes* | DNA gyrase subunit B (EC 5.99.1.3) |
| 172 | 253 | 0.01 | E4T5N7 | *Paludibacter propionicigenes* | Primosomal protein N' (EC 3.6.4.-) (ATP-dependent helicase PriA) |
| 173 | 253 | 0.01 | E4T8H6 | *Paludibacter propionicigenes* | Anthranilate synthase, component I (EC 4.1.3.27) |
| 174 | 253 | 0.01 | K9L3Q8 | *Magnusiomyces magnusii* | NADH-ubiquinone oxidoreductase chain 4 (EC 1.6.5.3) |
| 175 | 253 | 0.01 | | | |
| 176 | 253 | 0.01 | | | |

| | | | | | |
|---|---|---|---|---|---|
| 177 | 252 | 0.01 | E4T587 | *Paludibacter propionicigenes* | DNA mismatch repair protein MutS |
| 178 | 251 | 0.01 | E4T2L0 | *Paludibacter propionicigenes* | Glutamine--fructose-6-phosphate aminotransferase [isomerizing] (EC 2.6.1.16) (D-fructose-6-phosphate amidotransferase) (GFAT) (Glucosamine-6-phosphate synthase) (Hexosephosphate aminotransferase) (L-glutamine--D-fructose-6-phosphate amidotransferase) |
| 179 | 249 | 0.01 | E4T3Y7 | *Paludibacter propionicigenes* | 4-alpha-glucanotransferase (EC 2.4.1.25) |
| 180 | 248 | 0.01 | A0A1M3DPZ9 | *Bacteroidales bacterium* | DUF4954 domain-containing protein |
| 181 | 248 | 0.01 | A0A1M3FLT2 | *Bacteroidia bacterium* | Uncharacterized protein |
| 182 | 248 | 0.01 | E4T1U5 | *Paludibacter propionicigenes* | Propionyl-CoA carboxylase carboxyltransferase subunit (EC 6.4.1.3) |
| 183 | 248 | 0.01 | E4T526 | *Paludibacter propionicigenes* | Heat shock protein Hsp90 |
| 184 | 248 | 0.01 | | | |
| 185 | 247 | 0.01 | A0A1F3LNZ9 | *Bacteroidetes bacterium* | Glucarate dehydratase |
| 186 | 247 | 0.01 | E4T522 | *Paludibacter propionicigenes* | Isoleucine--tRNA ligase (EC 6.1.1.5) (Isoleucyl-tRNA synthetase) (IleRS) |
| 187 | 246 | 0.01 | E4T139 | *Paludibacter propionicigenes* | Ribonuclease, Rne/Rng family |
| 188 | 245 | 0.01 | A0A1J5HQ06 | *Porphyromonadaceae bacterium* | Protein translocase subunit SecA |
| 189 | 245 | 0.01 | A0A1J5HWA7 | *Porphyromonadaceae bacterium* | ATP-dependent DNA helicase RecG (EC 3.6.4.12) |
| 190 | 245 | 0.01 | E4T0F4 | *Paludibacter propionicigenes* | Alpha-glucan phosphorylase (EC 2.4.1.1) |
| 191 | 245 | 0.01 | | | |
| 192 | 244 | 0.01 | E4T4K3 | *Paludibacter propionicigenes* | UDP-N-acetylmuramoylalanine--D-glutamate ligase (EC 6.3.2.9) (D-glutamic acid-adding enzyme) (UDP-N-acetylmuramoyl-L-alanyl-D-glutamate synthetase) |
| 193 | 244 | 0.01 | K5ZYH0 | *Parabacteroides johnsonii* | Uncharacterized protein |
| 194 | 244 | 0.01 | | | |
| 195 | 241 | 0.01 | A0A1E4EHY6 | *Paludibacter sp.* | DNA polymerase I (EC 2.7.7.7) |
| 196 | 241 | 0.01 | A0A1M3M9P3 | *Paludibacter sp.* | Outer membrane protein assembly factor BamA |
| 197 | 241 | 0.01 | E4T2N4 | *Paludibacter propionicigenes* | Xanthosine permease |
| 198 | 241 | 0.01 | E4T4D3 | *Paludibacter propionicigenes* | Nitrogenase (EC 1.18.6.1) |
| 199 | 240 | 0.01 | E4T2Q7 | *Paludibacter propionicigenes* | Pullulanase, type I |
| 200 | 240 | 0.01 | E4T5N0 | *Paludibacter propionicigenes* | (P)ppGpp synthetase I, SpoT/RelA (EC 2.7.6.5) |
| 201 | 240 | 0.01 | E4T5Y8 | *Paludibacter propionicigenes* | Xanthine/uracil/vitamin C permease |

| | | | | | |
|---|---|---|---|---|---|
| 202 | 239 | 0.01 | A0A023UMS1 | *Magnusiomyces ingens* | NADH-ubiquinone oxidoreductase chain 1 (EC 1.6.5.3) |
| 203 | 239 | 0.01 | A0A1F3H411 | *Bacteroidetes bacterium* | Uncharacterized protein |
| 204 | 239 | 0.01 | A0A1J5HAI3 | *Porphyromonadaceae bacterium* | Uncharacterized protein |
| 205 | 239 | 0.01 | A0A1J5HW56 | *Porphyromonadaceae bacterium* | Succinate--CoA ligase [ADP-forming] subunit beta (EC 6.2.1.5) (Succinyl-CoA synthetase subunit beta) (SCS-beta) |
| 206 | 239 | 0.01 | E4T8K8 | *Paludibacter propionicigenes* | Uncharacterized protein |
| 207 | 238 | 0.01 | A0A1J5H9M4 | *Porphyromonadaceae bacterium* | Cysteine--tRNA ligase (EC 6.1.1.16) (Cysteinyl-tRNA synthetase) (CysRS) |
| 208 | 238 | 0.01 | A0A1J5HJU7 | *Porphyromonadaceae bacterium* | Anaerobic sulfatase maturase |
| 209 | 238 | 0.01 | A0A1Q3QJR2 | *Bacteroidia bacterium* | Cell surface protein SprA |
| 210 | 237 | 0.01 | A0A1J5HMM6 | *Porphyromonadaceae bacterium* | Glutaconyl-CoA decarboxylase subunit beta |
| 211 | 237 | 0.01 | E4T8H5 | *Paludibacter propionicigenes* | Tryptophan synthase beta chain (EC 4.2.1.20) |
| 212 | 236 | 0.01 | E4T069 | *Paludibacter propionicigenes* | Glycine dehydrogenase (decarboxylating) (EC 1.4.4.2) (Glycine cleavage system P-protein) (Glycine decarboxylase) (Glycine dehydrogenase (aminomethyl-transferring)) |
| 213 | 236 | 0.01 | E4T141 | *Paludibacter propionicigenes* | SSU ribosomal protein S1P |
| 214 | 235 | 0.01 | G4KVF2 | *Oscillibacter valericigenes* | Uncharacterized protein |
| 215 | 234 | 0.01 | A0A0P0GD61 | *Bacteroides cellulosilyticus* | Periplasmic beta-glucosidase (EC 3.2.1.21) |
| 216 | 234 | 0.01 | E4T4J0 | *Paludibacter propionicigenes* | Alpha-N-acetylglucosaminidase (EC 3.2.1.50) |
| 217 | 234 | 0.01 | E4T5C0 | *Paludibacter propionicigenes* | Glycine--tRNA ligase (EC 6.1.1.14) (Glycyl-tRNA synthetase) (GlyRS) |
| 218 | 234 | 0.01 | E4T6K7 | *Paludibacter propionicigenes* | Sodium/hydrogen exchanger |
| 219 | 234 | 0.01 | | | |
| 220 | 233 | 0.01 | A0A0Q7S7D9 | *Pelomonas sp.* | Uncharacterized protein |
| 221 | 233 | 0.01 | A0A1E4EKS6 | *Paludibacter sp.* | SusC/RagA family protein |
| 222 | 233 | 0.01 | A0A1J5HKW0 | *Porphyromonadaceae bacterium* | Phosphate transporter |
| 223 | 233 | 0.01 | E4T353 | *Paludibacter propionicigenes* | Uncharacterized protein |
| 224 | 233 | 0.01 | E4T3C6 | *Paludibacter propionicigenes* | Major facilitator superfamily MFS_1 |
| 225 | 233 | 0.01 | E4T3W0 | *Paludibacter propionicigenes* | Alpha-2-macroglobulin domain protein |
| 226 | 232 | 0.01 | A0A171AE46 | *Paludibacter jiangxiensis* | Signal transduction histidine kinase |
| 227 | 232 | 0.01 | E4T0P6 | *Paludibacter propionicigenes* | Succinate dehydrogenase subunit A (EC 1.3.5.1) (EC 1.3.99.1) |

| | | | | | |
|---|---|---|---|---|---|
| 228 | 232 | 0.01 | E4T115 | *Paludibacter propionicigenes* | Aspartokinase (EC 2.7.2.4) |
| 229 | 231 | 0.01 | A0A1J5HH65 | *Porphyromonadaceae bacterium* | Uncharacterized protein |
| 230 | 231 | 0.01 | E4T4D2 | *Paludibacter propionicigenes* | Nitrogenase MoFe cofactor biosynthesis protein NifE |
| 231 | 230 | 0.01 | A0A1Q3QC75 | *Bacteroidia bacterium* | Uncharacterized protein |
| 232 | 230 | 0.01 | E4T2K9 | *Paludibacter propionicigenes* | Amidophosphoribosyltransferase (EC 2.4.2.14) |
| 233 | 230 | 0.01 | E4T8I3 | *Paludibacter propionicigenes* | Transcription termination/antitermination protein NusA |
| 234 | 229 | 0.01 | E4T417 | *Paludibacter propionicigenes* | Leucine--tRNA ligase (EC 6.1.1.4) (Leucyl-tRNA synthetase) (LeuRS) |
| 235 | 228 | 0.01 | A0A1Q3QH52 | *Bacteroidia bacterium* | Uncharacterized protein |
| 236 | 227 | 0.01 | A0A1M3DMW5 | *Bacteroidales bacterium* | Long-chain fatty acid--CoA ligase |
| 237 | 227 | 0.01 | A0A1M3DPI5 | *Bacteroidales bacterium* | Uncharacterized protein |
| 238 | 226 | 0.01 | A0A1G4G387 | *Petrimonas mucosa* | Uncharacterized protein |
| 239 | 226 | 0.01 | A0A1M3DNS9 | *Bacteroidales bacterium* | Cell surface protein SprA |
| 240 | 226 | 0.01 | E4T0E5 | *Paludibacter propionicigenes* | DNA polymerase III subunit gamma/tau (EC 2.7.7.7) |
| 241 | 225 | 0.01 | A0A0K0WVI4 | *Litoditis aff. marina PmIII* | NADH dehydrogenase subunit 5 |
| 242 | 225 | 0.01 | A0A1E4ELV7 | *Paludibacter sp.* | Dehydrogenase |
| 243 | 224 | 0.01 | A0A1J5HQC1 | *Porphyromonadaceae bacterium* | Fe-S cluster assembly protein SufB |
| 244 | 224 | 0.01 | A0A1J5HR75 | *Porphyromonadaceae bacterium* | Phytoene desaturase |
| 245 | 224 | 0.01 | A0A1J5HS20 | *Porphyromonadaceae bacterium* | Flagellar motor protein MotA |
| 246 | 224 | 0.01 | A0A1M5CJ15 | *Bacteroides luti* | Glycosyltransferase, catalytic subunit of cellulose synthase and poly-beta-1,6-N-acetylglucosamine synthase |
| 247 | 224 | 0.01 | | | |
| 248 | 223 | 0.01 | D0TGP7 | *Bacteroides sp.* | Helicase C-terminal domain protein |
| 249 | 223 | 0.01 | E4T2R2 | *Paludibacter propionicigenes* | Phosphoglucomutase/phosphomannomutase alpha/beta/alpha domain I |
| 250 | 222 | 0.01 | A0A1J5HBP7 | *Porphyromonadaceae bacterium* | Uncharacterized protein |
| 251 | 222 | 0.01 | A0A1J5HII7 | *Porphyromonadaceae bacterium* | Methionine synthase |
| 252 | 221 | 0.01 | A0A0P6BFP2 | *Daphnia magna* | Uncharacterized protein |
| 253 | 221 | 0.01 | A0A1J5HRQ6 | *Porphyromonadaceae bacterium* | Uncharacterized protein |

| | | | | | |
|---|---|---|---|---|---|
| 254 | 221 | 0.01 | E9I741 | *Daphnia pulex* | Putative uncharacterized protein (Fragment) |
| 255 | 220 | 0.01 | A0A161LF68 | *Paludibacter jiangxiensis* | Two-component system, NtrC family |
| 256 | 220 | 0.01 | E4T650 | *Paludibacter propionicigenes* | Cell shape determining protein, MreB/Mrl family |
| 257 | 220 | 0.01 | E4T793 | *Paludibacter propionicigenes* | Elongation factor G (EF-G) |
| 258 | 219 | 0.01 | A0A1J5HIC5 | *Porphyromonadaceae bacterium* | ABC transporter permease |
| 259 | 219 | 0.01 | A0A1J5I218 | *Porphyromonadaceae bacterium* | Uncharacterized protein |
| 260 | 219 | 0.01 | E4T825 | *Paludibacter propionicigenes* | Chaperone protein DnaK (HSP70) (Heat shock 70 kDa protein) (Heat shock protein 70) |
| 261 | 219 | 0.01 | E4T8K7 | *Paludibacter propionicigenes* | L-aspartate oxidase (EC 1.4.3.16) |
| 262 | 219 | 0.01 | W2UCH0 | *Gammaproteobacteria bacterium* | Uncharacterized protein |
| 263 | 218 | 0.01 | A0A023UPK8 | *Magnusiomyces ingens* | Cytochrome c oxidase subunit 2 |
| 264 | 218 | 0.01 | A0A1M3MA44 | *Paludibacter sp.* | Uncharacterized protein |
| 265 | 218 | 0.01 | D8L7S1 | *Paramecium caudatum* | NADH-ubiquinone oxidoreductase chain 5 (EC 1.6.5.3) |
| 266 | 218 | 0.01 | E4T544 | *Paludibacter propionicigenes* | Response regulator receiver protein |
| 267 | 218 | 0.01 | G4KNZ0 | *Oscillibacter valericigenes* | Pyruvate carboxylase (EC 6.4.1.1) |
| 268 | 217 | 0.01 | E4T4C8 | *Paludibacter propionicigenes* | Nitrogenase protein alpha chain (EC 1.18.6.1) |
| 269 | 217 | 0.01 | E4T7D6 | *Paludibacter propionicigenes* | AMP-dependent synthetase and ligase |
| 270 | 216 | 0.01 | A0A1E4EAJ0 | *Paludibacter sp.* | Signal recognition particle protein (Fifty-four homolog) |
| 271 | 216 | 0.01 | A0A1J5HKA8 | *Porphyromonadaceae bacterium* | 4Fe-4S ferredoxin |
| 272 | 216 | 0.01 | A0A1J5HPN8 | *Porphyromonadaceae bacterium* | Chloride channel protein |
| 273 | 216 | 0.01 | E4T0A9 | *Paludibacter propionicigenes* | Chromosomal replication initiator protein DnaA |
| 274 | 215 | 0.01 | E4T604 | *Paludibacter propionicigenes* | AMP-dependent synthetase and ligase |
| 275 | 215 | 0.01 | E4T8A8 | *Paludibacter propionicigenes* | Carboxynorspermidine decarboxylase (EC 4.1.1.-) |
| 276 | 215 | 0.01 | A6LE80 | *Parabacteroides distasonis* | DNA-directed RNA polymerase subunit beta' (RNAP subunit beta') (EC 2.7.7.6) (RNA polymerase subunit beta') (Transcriptase subunit beta') |
| 277 | 214 | 0.01 | A0A1J5H964 | *Porphyromonadaceae bacterium* | Multifunctional fusion protein [Includes: UDP-3-O-acyl-N-acetylglucosamine deacetylase (UDP-3-O-acyl-GlcNAc deacetylase) (EC 3.5.1.108) (UDP-3-O-[R-3-hydroxymyristoyl]-N-acetylglucosamine deacetylase); 3-hydroxyacyl-[acyl-carrier-protein] dehydratase FabZ (EC 4.2.1.59) ((3R)-hydroxymyristoyl-[acyl-carrier-protein] dehydratase) (Beta-hydroxyacyl-ACP dehydratase) ((3R)-hydroxymyristoyl-ACP dehydrase)] |

| | | | | | |
|---|---|---|---|---|---|
| 278 | 214 | 0.01 | A0A1J5HM03 | *Porphyromonadaceae bacterium* | TonB-dependent receptor |
| 279 | 213 | 0.01 | E4T3G0 | *Paludibacter propionicigenes* | Glutamate synthase (NADH) small subunit (EC 1.4.1.14) |
| 280 | 213 | 0.01 | | | |
| 281 | 212 | 0.01 | A0A1E4EGA4 | *Paludibacter sp.* | Elongation factor 4 (EF-4) (EC 3.6.5.n1) (Ribosomal back-translocase LepA) |
| 282 | 212 | 0.01 | A0A1E4EKQ5 | *Paludibacter sp.* | DUF4914 domain-containing protein |
| 283 | 212 | 0.01 | A0A1J5HYA6 | *Porphyromonadaceae bacterium* | Amidophosphoribosyltransferase |
| 284 | 212 | 0.01 | E4T144 | *Paludibacter propionicigenes* | Drug resistance transporter, EmrB/QacA subfamily |
| 285 | 212 | 0.01 | E4T8G7 | *Paludibacter propionicigenes* | Phosphate transporter |
| 286 | 211 | 0.01 | E4T5H4 | *Paludibacter propionicigenes* | Uncharacterized protein |
| 287 | 210 | 0.01 | A0A0R3QEE4 | *Brugia timori* | Uncharacterized protein |
| 288 | 210 | 0.01 | A0A1Q3KXL3 | *Bacteroidales bacterium* | Glucosamine-6-phosphate deaminase (EC 3.5.99.6) |
| 289 | 210 | 0.01 | | | |
| 290 | 209 | 0.01 | A0A0S8JYW7 | *Bacteroides sp.* | Oxalate:formate antiporter |
| 291 | 209 | 0.01 | A0A1F3LP63 | *Bacteroidetes bacterium* | Phosphoenolpyruvate carboxykinase (ATP) (PCK) (PEP carboxykinase) (PEPCK) (EC 4.1.1.49) |
| 292 | 209 | 0.01 | A0A1J5HEC6 | *Porphyromonadaceae bacterium* | DNA topoisomerase IV |
| 293 | 209 | 0.01 | A0A1J5HF64 | *Porphyromonadaceae bacterium* | L-rhamnose isomerase (EC 5.3.1.14) |
| 294 | 209 | 0.01 | A0A1M3MDN4 | *Paludibacter sp.* | Tryptophanase |
| 295 | 209 | 0.01 | E4T1T8 | *Paludibacter propionicigenes* | Uncharacterized protein |
| 296 | 209 | 0.01 | E4T5S3 | *Paludibacter propionicigenes* | 2-oxoglutarate synthase (EC 1.2.7.3) |
| 297 | 209 | 0.01 | W0ER83 | *Barnesiella viscericola* | Alpha-xylosidase |
| 298 | 208 | 0.01 | A0A1J5HY73 | *Porphyromonadaceae bacterium* | DNA-directed DNA polymerase (EC 2.7.7.7) |
| 299 | 208 | 0.01 | E4T0I0 | *Paludibacter propionicigenes* | Dipeptidyl-peptidase IV (EC 3.4.14.5) |
| 300 | 208 | 0.01 | E4T3A0 | *Paludibacter propionicigenes* | RNA polymerase, sigma 70 subunit, RpoD subfamily |
| 301 | 207 | 0.01 | A0A0Q7RZB1 | *Pelomonas sp.* | Uncharacterized protein |
| 302 | 207 | 0.01 | E4T2M4 | *Paludibacter propionicigenes* | Transcription-repair-coupling factor (TRCF) (EC 3.6.4.-) |
| 303 | 207 | 0.01 | E4T3Y3 | *Paludibacter propionicigenes* | Seryl-tRNA synthetase (EC 6.1.1.11) |
| 304 | 207 | 0.01 | E4T4E2 | *Paludibacter propionicigenes* | Nucleoside recognition domain protein |

| | | | | | |
|---|---|---|---|---|---|
| 305 | 207 | 0.01 | E4T4G7 | *Paludibacter propionicigenes* | Ferrous iron transport protein B |
| 306 | 207 | 0.01 | E4T7Y7 | *Paludibacter propionicigenes* | Major facilitator superfamily MFS_1 |
| 307 | 207 | 0.01 | E4T872 | *Paludibacter propionicigenes* | Serine palmitoyltransferase (EC 2.3.1.29) (EC 2.3.1.50) |
| 308 | 207 | 0.01 | E4T8M9 | *Paludibacter propionicigenes* | Alpha amylase catalytic region |
| 309 | 204 | 0.01 | A0A1J5H7Y6 | *Porphyromonadaceae bacterium* | CTP synthase (EC 6.3.4.2) (Cytidine 5'-triphosphate synthase) (Cytidine triphosphate synthetase) (CTP synthetase) (CTPS) (UTP--ammonia ligase) |
| 310 | 204 | 0.01 | A0A1J5HDM2 | *Porphyromonadaceae bacterium* | Transglycosylase (Fragment) |
| 311 | 204 | 0.01 | A0A1J5HFV4 | *Porphyromonadaceae bacterium* | tRNA-2-methylthio-N(6)-dimethylallyladenosine synthase (EC 2.8.4.3) ((Dimethylallyl)adenosine tRNA methylthiotransferase MiaB) (tRNA-i(6)A37 methylthiotransferase) |
| 312 | 204 | 0.01 | A0A1J5HX74 | *Porphyromonadaceae bacterium* | CoA-disulfide reductase |
| 313 | 204 | 0.01 | I9FDN5 | *Bacteroides cellulosilyticus* | Uncharacterized protein |
| 314 | 204 | 0.01 | | | |
| 315 | 203 | 0.01 | A0A1J5HLC9 | *Porphyromonadaceae bacterium* | Uncharacterized protein |
| 316 | 203 | 0.01 | D8L7R1 | *Paramecium caudatum* | NADH-ubiquinone oxidoreductase chain 4 (EC 1.6.5.3) |
| 317 | 203 | 0.01 | E4T0G8 | *Paludibacter propionicigenes* | Aspartate kinase homoserine dehydrogenase (EC 1.1.1.3) (EC 2.7.2.4) |
| 318 | 203 | 0.01 | E4T0X5 | *Paludibacter propionicigenes* | Peptidase U32 |
| 319 | 203 | 0.01 | E4T7C2 | *Paludibacter propionicigenes* | DNA-directed RNA polymerase subunit alpha (RNAP subunit alpha) (EC 2.7.7.6) (RNA polymerase subunit alpha) (Transcriptase subunit alpha) |
| 320 | 203 | 0.01 | | | |
| 321 | 202 | 0.01 | A0A0K3A869 | *Xanthomonas translucens pv. poae* | Riboflavin biosynthesis protein (EC 2.7.7.2) |
| 322 | 202 | 0.01 | E4T727 | *Paludibacter propionicigenes* | ATP-dependent DNA helicase RecG (EC 3.6.4.12) |
| 323 | 202 | 0.01 | | | |
| 324 | 201 | 0.01 | A0A1J5SFW2 | *mine drainage metagenome* | Cyclic pyranopterin monophosphate synthase (EC 4.1.99.22) |
| 325 | 201 | 0.01 | A0A1Q3MTC6 | *Bacteroidetes bacterium* | DNA helicase (EC 3.6.4.12) |
| 326 | 201 | 0.01 | E4T299 | *Paludibacter propionicigenes* | Malonyl CoA-acyl carrier protein transacylase (EC 2.3.1.39) |
| 327 | 201 | 0.01 | E4T5K0 | *Paludibacter propionicigenes* | Peptidase C1A papain |
| 328 | 201 | 0.01 | E4T5M4 | *Paludibacter propionicigenes* | Fumarate hydratase class I (EC 4.2.1.2) |
| 329 | 201 | 0.01 | K5ZY38 | *Parabacteroides johnsonii* | Uncharacterized protein |

| | | | | | |
|---|---|---|---|---|---|
| 330 | 200 | 0.01 | A0A0S7C3I4 | *Lentimicrobium saccharophilum* | Uncharacterized protein |
| 331 | 200 | 0.01 | K9L4A0 | *Magnusiomyces magnusii* | Cytochrome c oxidase subunit 1 (EC 1.9.3.1) |
| 332 | 199 | 0.01 | A0A1J5HHU1 | *Porphyromonadaceae bacterium* | Magnesium transport protein CorA |
| 333 | 199 | 0.01 | E4T576 | *Paludibacter propionicigenes* | Integral membrane sensor signal transduction histidine kinase |
| 334 | 199 | 0.01 | E4T6H1 | *Paludibacter propionicigenes* | AAA ATPase |
| 335 | 199 | 0.01 | G4KT17 | *Oscillibacter valericigenes* | Putative helicase |
| 336 | 199 | 0.01 | K6AJ45 | *Parabacteroides distasonis* | SusC/RagA family TonB-linked outer membrane protein |
| 337 | 198 | 0.01 | A0A0P0GH26 | *Bacteroides cellulosilyticus* | Isomalto-dextranase (EC 3.2.1.94) |
| 338 | 198 | 0.01 | A0A1J5HKH3 | *Porphyromonadaceae bacterium* | Methylmalonyl-CoA mutase small subunit |
| 339 | 198 | 0.01 | A0A1J5HNH1 | *Porphyromonadaceae bacterium* | Transketolase |
| 340 | 198 | 0.01 | A0A1M3MEQ1 | *Paludibacter sp.* | Phosphoribosylformylglycinamidine synthase (FGAM synthase) (FGAMS) (EC 6.3.5.3) (Formylglycinamide ribonucleotide amidotransferase) (FGAR amidotransferase) (FGAR-AT) |
| 341 | 198 | 0.01 | A6LIH5 | *Parabacteroides distasonis* | Phosphoenolpyruvate synthase/pyruvate phosphate dikinase |
| 342 | 198 | 0.01 | B0MX71 | *Alistipes putredinis* | Integrase core domain protein |
| 343 | 198 | 0.01 | E4T3P8 | *Paludibacter propionicigenes* | 60 kDa chaperonin (GroEL protein) (Protein Cpn60) |
| 344 | 198 | 0.01 | | | |
| 345 | 197 | 0.01 | A0A150XAA9 | *Roseivirga spongicola* | Transposase |
| 346 | 197 | 0.01 | A0A1J5HJ85 | *Porphyromonadaceae bacterium* | DEAD/DEAH box helicase |
| 347 | 197 | 0.01 | A0A1J5HRP9 | *Porphyromonadaceae bacterium* | Uncharacterized protein |
| 348 | 197 | 0.01 | E4T3A4 | *Paludibacter propionicigenes* | Endonuclease MutS2 (EC 3.1.-.-) |
| 349 | 197 | 0.01 | E4T5K5 | *Paludibacter propionicigenes* | ATP-dependent DNA helicase RecQ (EC 3.6.1.-) |
| 350 | 197 | 0.01 | E4T766 | *Paludibacter propionicigenes* | Glutamine--tRNA ligase (EC 6.1.1.18) |
| 351 | 196 | 0.01 | A0A1E4EB10 | *Paludibacter sp.* | Uncharacterized protein |
| 352 | 196 | 0.01 | A0A1E4EHT2 | *Paludibacter sp.* | Uncharacterized protein |
| 353 | 196 | 0.01 | A0A1J5HF31 | *Porphyromonadaceae bacterium* | Cell division protein FtsK |
| 354 | 196 | 0.01 | E4T428 | *Paludibacter propionicigenes* | Glutamate--tRNA ligase (EC 6.1.1.17) (Glutamyl-tRNA synthetase) (GluRS) |
| 355 | 196 | 0.01 | F8X245 | *Dysgonomonas mossii* | UDP-glucose 4-epimerase |

| | | | | | |
|---|---|---|---|---|---|
| 356 | 196 | 0.01 | | | |
| 357 | 195 | 0.01 | A0A1M3MAV7 | *Paludibacter sp. 47-17* | Uncharacterized protein |
| 358 | 195 | 0.01 | E4T4F1 | *Paludibacter propionicigenes* | Lytic transglycosylase catalytic |
| 359 | 195 | 0.01 | E4T4H8 | *Paludibacter propionicigenes* | RelA/SpoT family protein (EC 2.7.6.5) |
| 360 | 195 | 0.01 | E4T4P1 | *Paludibacter propionicigenes* | Peptidase U34 dipeptidase |
| 361 | 195 | 0.01 | E4T5T2 | *Paludibacter propionicigenes* | Holliday junction ATP-dependent DNA helicase RuvB (EC 3.6.4.12) |
| 362 | 195 | 0.01 | | | |
| 363 | 195 | 0.01 | | | |
| 364 | 194 | 0.01 | A0A0N7IFQ6 | *Bacteroides cellulosilyticus* | Periplasmic beta-glucosidase (EC 3.2.1.21) |
| 365 | 194 | 0.01 | A0A1F3JYD9 | *Bacteroidetes bacterium* | Uncharacterized protein |
| 366 | 194 | 0.01 | A0A1J5HD63 | *Porphyromonadaceae bacterium* | RelA/SpoT family protein |
| 367 | 194 | 0.01 | E4T2Y0 | *Paludibacter propionicigenes* | UDP-N-acetylglucosamine 1-carboxyvinyltransferase (EC 2.5.1.7) (Enoylpyruvate transferase) (UDP-N-acetylglucosamine enolpyruvyl transferase) (EPT) |
| 368 | 194 | 0.01 | E4T4K5 | *Paludibacter propionicigenes* | UDP-N-acetylmuramoyl-L-alanyl-D-glutamate--2,6-diaminopimelate ligase (EC 6.3.2.13) (Meso-A2pm-adding enzyme) (Meso-diaminopimelate-adding enzyme) (UDP-MurNAc-L-Ala-D-Glu:meso-diaminopimelate ligase) (UDP-MurNAc-tripeptide synthetase) (UDP-N-acetylmuramyl-tripeptide synthetase) |
| 369 | 193 | 0.01 | A0A076NZ72 | *Flavobacterium psychrophilum* | Uncharacterized protein |
| 370 | 193 | 0.01 | E4T6L4 | *Paludibacter propionicigenes* | Succinylglutamate desuccinylase/aspartoacylase |
| 371 | 192 | 0.01 | A0A1J5HBA6 | *Porphyromonadaceae bacterium* | DNA primase (EC 2.7.7.-) |
| 372 | 192 | 0.01 | E4T1E6 | *Paludibacter propionicigenes* | Protein translocase subunit SecA |
| 373 | 192 | 0.01 | E4T451 | *Paludibacter propionicigenes* | Histidine biosynthesis bifunctional protein HisB [Includes: Imidazoleglycerol-phosphate dehydratase (IGPD) (EC 4.2.1.19); Histidinol-phosphatase (EC 3.1.3.15)] |
| 374 | 192 | 0.01 | E4T591 | *Paludibacter propionicigenes* | 2,3-bisphosphoglycerate-independent phosphoglycerate mutase (BPG-independent PGAM) (Phosphoglyceromutase) (iPGM) (EC 5.4.2.12) |
| 375 | 192 | 0.01 | E4T8R6 | *Paludibacter propionicigenes* | Lysine--tRNA ligase (EC 6.1.1.6) (Lysyl-tRNA synthetase) (LysRS) |
| 376 | 191 | 0.01 | A0A1J5HBR6 | *Porphyromonadaceae bacterium* | Protein translocase subunit SecY |
| 377 | 190 | 0.01 | A0A1Q3M3M9 | *Bacteroidales bacterium* | Metal-independent alpha-mannosidase |
| 378 | 190 | 0.01 | E4T3C0 | *Paludibacter propionicigenes* | Aminotransferase class I and II |

| | | | | | |
|---|---|---|---|---|---|
| 379 | 190 | 0.01 | E4T775 | *Paludibacter propionicigenes* | Ribonuclease R (RNase R) (EC 3.1.13.1) |
| 380 | 190 | 0.01 | F5UKT4 | *Microcoleus vaginatus* | Polymorphic outer membrane protein |
| 381 | 189 | 0.01 | A0A0D5N7W1 | *Geobacter sulfurreducens* | ATPase |
| 382 | 189 | 0.01 | A0A1E4EG73 | *Paludibacter sp.* | Acetate kinase (EC 2.7.2.1) (Acetokinase) |
| 383 | 189 | 0.01 | A0A1J5HEW8 | *Porphyromonadaceae bacterium* | ABC transporter |
| 384 | 189 | 0.01 | A0A1J5HFI3 | *Porphyromonadaceae bacterium* | Glutamine--fructose-6-phosphate aminotransferase [isomerizing] (EC 2.6.1.16) (D-fructose-6-phosphate amidotransferase) (GFAT) (Glucosamine-6-phosphate synthase) (Hexosephosphate aminotransferase) (L-glutamine--D-fructose-6-phosphate amidotransferase) |
| 385 | 189 | 0.01 | A0A1M3MEB3 | *Paludibacter sp.* | Uncharacterized protein |
| 386 | 189 | 0.01 | A0A1M3MF51 | *Paludibacter sp.* | Translation initiation factor IF-2 |
| 387 | 189 | 0.01 | A0A1Q3LR33 | *Bacteroidales bacterium* | Inosine 5-monophosphate dehydrogenase |
| 388 | 188 | 0.01 | A0A015TUI8 | *Bacteroides fragilis* | AAA domain protein |
| 389 | 188 | 0.01 | A0A1M3DK57 | *Bacteroidales bacterium* | Outer membrane protein assembly factor BamA |
| 390 | 188 | 0.01 | A0A1M3MFN9 | *Paludibacter sp.* | Ribulokinase (EC 2.7.1.16) |
| 391 | 188 | 0.01 | A0A1Q3QIV6 | *Bacteroidia bacterium* | Uncharacterized protein |
| 392 | 188 | 0.01 | E4T3Y1 | *Paludibacter propionicigenes* | Glutamine-dependent NAD(+) synthetase (EC 6.3.5.1) (NAD(+) synthase [glutamine-hydrolyzing]) |
| 393 | 188 | 0.01 | E4T841 | *Paludibacter propionicigenes* | Peptidyl-dipeptidase Dcp (EC 3.4.15.5) |
| 394 | 187 | 0.01 | A0A1E4EG51 | *Paludibacter sp.* | Symporter |
| 395 | 187 | 0.01 | A0A1J5HWJ7 | *Porphyromonadaceae bacterium* | UvrABC system protein B (Protein UvrB) (Excinuclease ABC subunit B) |
| 396 | 187 | 0.01 | A0A1M3DH56 | *Bacteroidales bacterium* | Bifunctional purine biosynthesis protein PurH [Includes: IMP cyclohydrolase (EC 3.5.4.10) (IMP synthase) (Inosinicase) (ATIC); Phosphoribosylaminoimidazolecarboxamide formyltransferase (EC 2.1.2.3) (AICAR transformylase)] |
| 397 | 187 | 0.01 | B7B780 | *Parabacteroides johnsonii* | Glycosyl hydrolase family 2, sugar binding domain protein (Fragment) |
| 398 | 187 | 0.01 | E4T1D9 | *Paludibacter propionicigenes* | RNA-metabolising metallo-beta-lactamase |
| 399 | 187 | 0.01 | E4T405 | *Paludibacter propionicigenes* | Glycogen synthase (UDP-glucose) (EC 2.4.1.21) |
| 400 | 187 | 0.01 | E4T418 | *Paludibacter propionicigenes* | Uncharacterized protein |
| 401 | 187 | 0.01 | E4T618 | *Paludibacter propionicigenes* | DNA topoisomerase (ATP-hydrolyzing) (EC 5.99.1.3) |

| | | | | | |
|---|---|---|---|---|---|
| 402 | 187 | 0.01 | E4T6E7 | *Paludibacter propionicigenes* | 3-oxoacyl-[acyl-carrier-protein] synthase 2 (EC 2.3.1.179) |
| 403 | 186 | 0.01 | A0A023UNE7 | *Magnusiomyces ingens* | ATP synthase subunit a |
| 404 | 186 | 0.01 | A0A1J5B2X9 | *Bacteroidetes bacterium* | Transglycosylase |
| 405 | 186 | 0.01 | A0A1J5HIH9 | *Porphyromonadaceae bacterium* | Polysaccharide biosynthesis protein |
| 406 | 186 | 0.01 | A0A1J5HJ31 | *Porphyromonadaceae bacterium* | DNA repair protein RadA |
| 407 | 186 | 0.01 | E4T3B8 | *Paludibacter propionicigenes* | Uncharacterized protein |
| 408 | 186 | 0.01 | E4T543 | *Paludibacter propionicigenes* | N-acetylornithine carbamoyltransferase (EC 2.1.3.9) |
| 409 | 185 | 0.01 | A0A1F3MS16 | *Bacteroidetes bacterium* | Uncharacterized protein (Fragment) |
| 410 | 185 | 0.01 | A0A1J5HBM9 | *Porphyromonadaceae bacterium* | Uncharacterized protein |
| 411 | 185 | 0.01 | A0A1M3FTD4 | *Bacteroidia bacterium* | Beta-galactosidase (EC 3.2.1.23) (Lactase) |
| 412 | 185 | 0.01 | A0A1M3M8G5 | *Paludibacter sp.* | 1,4-alpha-glucan branching enzyme (EC 2.4.1.18) |
| 413 | 185 | 0.01 | E4T772 | *Paludibacter propionicigenes* | DNA topoisomerase 1 (EC 5.99.1.2) (DNA topoisomerase I) |
| 414 | 185 | 0.01 | K9L486 | *Magnusiomyces magnusii* | NADH-ubiquinone oxidoreductase chain 4L (EC 1.6.5.3) |
| 415 | 184 | 0.01 | A0A1M3MCQ7 | *Paludibacter sp.* | 30S ribosomal protein S1 |
| 416 | 184 | 0.01 | E4T658 | *Paludibacter propionicigenes* | 3-hydroxyacyl-CoA dehydrogenase NAD-binding protein |
| 417 | 184 | 0.01 | E4T7N5 | *Paludibacter propionicigenes* | Histidine--tRNA ligase (EC 6.1.1.21) (Histidyl-tRNA synthetase) (HisRS) |
| 418 | 184 | 0.01 | E4T8A5 | *Paludibacter propionicigenes* | Agmatine deiminase (EC 3.5.3.12) |
| 419 | 184 | 0.01 | E4T8A7 | *Paludibacter propionicigenes* | Carboxynorspermidine dehydrogenase |
| 420 | 183 | 0.01 | A0A0D5N8M0 | *Geobacter sulfurreducens* | MgtE integral membrane protein |
| 421 | 183 | 0.01 | A0A1J5HIR8 | *Porphyromonadaceae bacterium* | Ammonium transporter |
| 422 | 183 | 0.01 | A0A1J5HK82 | *Porphyromonadaceae bacterium* | 1-deoxy-D-xylulose-5-phosphate synthase (EC 2.2.1.7) (1-deoxyxylulose-5-phosphate synthase) (DXP synthase) (DXPS) |
| 423 | 183 | 0.01 | A0A1M3M8L6 | *Paludibacter sp.* | ATP-dependent DNA helicase RecQ |
| 424 | 183 | 0.01 | E4T130 | *Paludibacter propionicigenes* | GDP-mannose 4,6-dehydratase (EC 4.2.1.47) (GDP-D-mannose dehydratase) |
| 425 | 183 | 0.01 | E4T6F0 | *Paludibacter propionicigenes* | Ribosome-binding ATPase YchF |
| 426 | 183 | 0.01 | G4KSU6 | *Oscillibacter valericigenes* | DNA-directed RNA polymerase subunit beta (RNAP subunit beta) (EC 2.7.7.6) (RNA polymerase subunit beta) (Transcriptase subunit beta) |
| 427 | 183 | 0.01 | | | |

| | | | | | |
|---|---|---|---|---|---|
| 428 | 182 | 0.01 | A0A161LUJ1 | *Paludibacter jiangxiensis* | L-arabinose isomerase (EC 5.3.1.4) |
| 429 | 182 | 0.01 | A0A1J5HLA6 | *Porphyromonadaceae bacterium* | Peptidylprolyl isomerase (EC 5.2.1.8) |
| 430 | 182 | 0.01 | A0A1Q3LMQ1 | *Bacteroidales bacterium* | Acriflavin resistance protein |
| 431 | 182 | 0.01 | A0A1Q3QB76 | *Bacteroidia bacterium* | Transaldolase (EC 2.2.1.2) |
| 432 | 182 | 0.01 | E4T197 | *Paludibacter propionicigenes* | Uncharacterized protein |
| 433 | 182 | 0.01 | E4T300 | *Paludibacter propionicigenes* | ABC transporter related protein |
| 434 | 181 | 0.01 | A0A1J5HH39 | *Porphyromonadaceae bacterium* | Fe-S cluster assembly protein SufD |
| 435 | 181 | 0.01 | A0A1J5HIJ9 | *Porphyromonadaceae bacterium* | Aerotolerance regulator BatA |
| 436 | 181 | 0.01 | E4T3L2 | *Paludibacter propionicigenes* | Imidazole glycerol phosphate synthase subunit HisF (EC 4.1.3.-) (IGP synthase cyclase subunit) (IGP synthase subunit HisF) (ImGP synthase subunit HisF) (IGPS subunit HisF) |
| 437 | 181 | 0.01 | | | |
| 438 | 180 | 0.01 | A0A023UM83 | *Magnusiomyces ingens* | NADH-ubiquinone oxidoreductase chain 6 (EC 1.6.5.3) |
| 439 | 180 | 0.01 | A0A0F5J3W9 | *Parabacteroides sp.* | SusC/RagA family TonB-linked outer membrane protein |
| 440 | 180 | 0.01 | A0A1M3DKS7 | *Bacteroidales bacterium* | Uncharacterized protein |
| 441 | 180 | 0.01 | A0A1Q3QDG3 | *Bacteroidia bacterium* | SusC/RagA family TonB-linked outer membrane protein |
| 442 | 180 | 0.01 | E4T2P1 | *Paludibacter propionicigenes* | 3-methyl-2-oxobutanoate hydroxymethyltransferase (EC 2.1.2.11) (Ketopantoate hydroxymethyltransferase) (KPHMT) |
| 443 | 180 | 0.01 | E4T3C5 | *Paludibacter propionicigenes* | Periplasmic binding protein |
| 444 | 180 | 0.01 | E4T414 | *Paludibacter propionicigenes* | Uncharacterized protein |
| 445 | 180 | 0.01 | E4T7C5 | *Paludibacter propionicigenes* | Uridylate kinase (UK) (EC 2.7.4.22) (Uridine monophosphate kinase) (UMP kinase) (UMPK) |
| 446 | 180 | 0.01 | E4T836 | *Paludibacter propionicigenes* | Phosphoglycerate mutase |
| 447 | 179 | 0.01 | A0A171A7A4 | *Paludibacter jiangxiensis* | Methionine--tRNA ligase (EC 6.1.1.10) (Methionyl-tRNA synthetase) (MetRS) |
| 448 | 179 | 0.01 | A0A1J5HVJ6 | *Porphyromonadaceae bacterium* | MATE family efflux transporter |
| 449 | 179 | 0.01 | A0A1M5KAB4 | *Dysgonomonas macrotermitis* | Transposase InsO and inactivated derivatives |
| 450 | 179 | 0.01 | E4T073 | *Paludibacter propionicigenes* | Chaperone protein ClpB |
| 451 | 179 | 0.01 | E4T0C8 | *Paludibacter propionicigenes* | Fmu (Sun) domain protein |
| 452 | 179 | 0.01 | E4T293 | *Paludibacter propionicigenes* | Tryptophan synthase beta chain (EC 4.2.1.20) |
| 453 | 179 | 0.01 | E4T3S1 | *Paludibacter propionicigenes* | Glucose-1-phosphate thymidylyltransferase (EC 2.7.7.24) |

| | | | | | |
|---|---|---|---|---|---|
| 454 | 179 | 0.01 | E4T665 | *Paludibacter propionicigenes* | D-alanine--D-alanine ligase (EC 6.3.2.4) (D-Ala-D-Ala ligase) (D-alanylalanine synthetase) |
| 455 | 178 | 0.01 | A0A1M3DLX8 | *Bacteroidales bacterium* | Uncharacterized protein |
| 456 | 178 | 0.01 | E4T3A2 | *Paludibacter propionicigenes* | Enolase (EC 4.2.1.11) (2-phospho-D-glycerate hydro-lyase) (2-phosphoglycerate dehydratase) |
| 457 | 178 | 0.01 | E4T429 | *Paludibacter propionicigenes* | Elongation factor Ts (EF-Ts) |
| 458 | 178 | 0.01 | E4T5S2 | *Paludibacter propionicigenes* | Pyruvate flavodoxin/ferredoxin oxidoreductase domain protein |
| 459 | 177 | 0.01 | A0A133Y220 | *Bacteroidales bacterium* | Response regulator receiver domain protein |
| 460 | 177 | 0.01 | A0A1J5HJ83 | *Porphyromonadaceae bacterium* | Threonine synthase |
| 461 | 177 | 0.01 | A0A1M3MDT6 | *Paludibacter sp.* | tRNA uridine 5-carboxymethylaminomethyl modification enzyme MnmG (Glucose-inhibited division protein A) |
| 462 | 177 | 0.01 | A0A1Q3QCA7 | *Bacteroidia bacterium* | Glycosyl hydrolase family 31 |
| 463 | 177 | 0.01 | E4T088 | *Paludibacter propionicigenes* | Amylo-alpha-16-glucosidase |
| 464 | 177 | 0.01 | E4T3V0 | *Paludibacter propionicigenes* | Cysteine desulfurase (EC 2.8.1.7) |
| 465 | 177 | 0.01 | E4T5J8 | *Paludibacter propionicigenes* | Aminoacyl-histidine dipeptidase |
| 466 | 177 | 0.01 | E4T7Z2 | *Paludibacter propionicigenes* | Glycosyl transferase family 2 |
| 467 | 176 | 0.01 | A0A1M3MDP9 | *Paludibacter sp.* | DNA-directed DNA polymerase (EC 2.7.7.7) |
| 468 | 176 | 0.01 | E4T575 | *Paludibacter propionicigenes* | Two component transcriptional regulator, winged helix family |
| 469 | 175 | 0.01 | A0A1J5HL29 | *Porphyromonadaceae bacterium* | Peptidase S9 |
| 470 | 175 | 0.01 | A0A1M3MBJ3 | *Paludibacter sp.* | Thiol:disulfide interchange protein |
| 471 | 175 | 0.01 | A0A1M3MF71 | *Paludibacter sp.* | Oxidoreductase |
| 472 | 175 | 0.01 | E4T2L1 | *Paludibacter propionicigenes* | LL-diaminopimelate aminotransferase (DAP-AT) (DAP-aminotransferase) (LL-DAP-aminotransferase) (EC 2.6.1.83) |
| 473 | 175 | 0.01 | E4T3A7 | *Paludibacter propionicigenes* | ABC transporter related protein |
| 474 | 175 | 0.01 | E4T7X5 | *Paludibacter propionicigenes* | Peptidase M23 |
| 475 | 175 | 0.01 | K5ZFA7 | *Parabacteroides merdae* | Pyruvate formate-lyase |
| 476 | 175 | 0.01 | S3C5M7 | *Ophiostoma piceae* | Uncharacterized protein |
| 477 | 175 | 0.01 | U6RUP9 | *Bacteroides sp.* | Uncharacterized protein |
| 478 | 174 | 0.01 | A0A1F3H388 | *Bacteroidetes bacterium* | Uncharacterized protein |

| | | | | | |
|---|---|---|---|---|---|
| 479 | 174 | 0.01 | A0A1Q3QI16 | *Bacteroidia bacterium* | SusC/RagA family TonB-linked outer membrane protein |
| 480 | 174 | 0.01 | E4T338 | *Paludibacter propionicigenes* | Carboxyl transferase (EC 6.4.1.3) |
| 481 | 174 | 0.01 | E4T4P4 | *Paludibacter propionicigenes* | TonB-dependent receptor |
| 482 | 173 | 0.01 | A0A095ZHM4 | *Prevotella buccalis* | Gliding motility protein |
| 483 | 173 | 0.01 | A0A1F3M9H0 | *Bacteroidetes bacterium* | GMP synthase [glutamine-hydrolyzing] (EC 6.3.5.2) (Fragment) |
| 484 | 173 | 0.01 | E4T0H2 | *Paludibacter propionicigenes* | Uncharacterized protein |
| 485 | 172 | 0.01 | A0A077XLW5 | *Sphinx1.76-related DNA* | Replication protein |
| 486 | 172 | 0.01 | A0A174TJ83 | *Parabacteroides distasonis* | Methylmalonyl-CoA mutase large subunit (EC 5.4.99.2) |
| 487 | 172 | 0.01 | A0A1J5HPZ5 | *Porphyromonadaceae bacterium* | Bifunctional aspartate kinase/homoserine dehydrogenase I |
| 488 | 172 | 0.01 | A0A1M3MEH0 | *Paludibacter sp.* | SusC/RagA family TonB-linked outer membrane protein |
| 489 | 172 | 0.01 | E4T1J0 | *Paludibacter propionicigenes* | Purine or other phosphorylase family 1 |
| 490 | 171 | 0.01 | A0A1M3DCQ2 | *Bacteroidales bacterium* | DNA-directed RNA polymerase subunit beta (RNAP subunit beta) (EC 2.7.7.6) (RNA polymerase subunit beta) (Transcriptase subunit beta) |
| 491 | 171 | 0.01 | R7LEV7 | *Prevotella sp.* | Type III restriction protein res subunit |
| 492 | 171 | 0.01 | | | |
| 493 | 170 | 0.01 | A0A0J6FCE6 | *Parabacteroides goldsteinii* | Uncharacterized protein |
| 494 | 170 | 0.01 | A0A1J5HZK4 | *Porphyromonadaceae bacterium* | NADH-quinone oxidoreductase subunit C |
| 495 | 170 | 0.01 | A0A1M3M9Y9 | *Paludibacter sp.* | Uncharacterized protein |
| 496 | 170 | 0.01 | E4T335 | *Paludibacter propionicigenes* | Nucleotide sugar dehydrogenase |
| 497 | 170 | 0.01 | E4T396 | *Paludibacter propionicigenes* | Aconitase (EC 4.2.1.3) |
| 498 | 170 | 0.01 | E4T8I1 | *Paludibacter propionicigenes* | Tryptophan synthase alpha chain (EC 4.2.1.20) |
| 499 | 169 | 0.01 | A0A170Z6B9 | *Paludibacter jiangxiensis* | Elongation factor G (EF-G) |
| 500 | 169 | 0.01 | A0A1E4EGX9 | *Paludibacter sp.* | Uncharacterized protein |

**Table S5a.** Functional overview of M2A community (top 500 rank only).

| Rank | Count | Abundance | ID | Organism | Protein Names |
|---|---|---|---|---|---|
| 1 | 5150 | 0.16 | A0A0D5N8Q4 | *Geobacter sulfurreducens* | Transposase |
| 2 | 3422 | 0.10 | E1QK39 | *Desulfarculus baarsii* | Helicase domain protein |
| 3 | 3198 | 0.10 | A0A0A8WNU3 | *Geobacter sp.* | Protein TraC |
| 4 | 3107 | 0.09 | A0LQ17 | *Syntrophobacter fumaroxidans* | Helicase domain protein |
| 5 | 2629 | 0.08 | Q39Y12 | *Geobacter metallireducens* | DNA-directed RNA polymerase subunit beta' (RNAP subunit beta') (EC 2.7.7.6) (RNA polymerase subunit beta') (Transcriptase subunit beta') |
| 6 | 2512 | 0.08 | Q748X8 | *Geobacter sulfurreducens* | Elongation factor Tu (EF-Tu) |
| 7 | 2494 | 0.08 | Q39RM9 | *Geobacter metallireducens* | Membrane protein, putative |
| 8 | 2485 | 0.08 | A1AKK8 | *Pelobacter propionicus* | Uncharacterized protein |
| 9 | 2398 | 0.07 | | | |
| 10 | 2396 | 0.07 | A0A1G0LE89 | *Geobacteraceae bacterium* | Aconitate hydratase (Aconitase) (EC 4.2.1.3) |
| 11 | 2391 | 0.07 | Q748Y6 | *Geobacter sulfurreducens* | DNA-directed RNA polymerase subunit beta (RNAP subunit beta) (EC 2.7.7.6) (RNA polymerase subunit beta) (Transcriptase subunit beta) |
| 12 | 2284 | 0.07 | | | |
| 13 | 2234 | 0.07 | A0A0D5N7W1 | *Geobacter sulfurreducens* | ATPase |
| 14 | 2234 | 0.07 | B3EB58 | *Geobacter lovleyi* | Uncharacterized protein |
| 15 | 2228 | 0.07 | | | |
| 16 | 2184 | 0.07 | E1QK33 | *Desulfarculus baarsii* | ATP-binding region ATPase domain protein |
| 17 | 2061 | 0.06 | Q747C7 | *Geobacter sulfurreducens* | 60 kDa chaperonin (GroEL protein) (Protein Cpn60) |
| 18 | 2017 | 0.06 | A5GAB4 | *Geobacter sulfurreducens* | Putative GAF sensor protein |
| 19 | 2007 | 0.06 | Q74BB9 | *Geobacter sulfurreducens* | TraG family protein |
| 20 | 1993 | 0.06 | Q748Y8 | *Geobacter sulfurreducens* | Elongation factor G 2 (EF-G 2) |
| 21 | 1979 | 0.06 | Q74AK2 | *Geobacter sulfurreducens* | Sodium/solute symporter family protein |
| 22 | 1935 | 0.06 | A0A1G0M577 | *Geobacteraceae bacterium* | Uncharacterized protein |
| 23 | 1928 | 0.06 | Q74BB8 | *Geobacter sulfurreducens* | Metal-dependent phosphohydrolase |
| 24 | 1851 | 0.06 | Q39QB9 | *Geobacter metallireducens* | NADH dehydrogenase I, M subunit |
| 25 | 1851 | 0.06 | Q74BC2 | *Geobacter sulfurreducens* | Integrative genetic element Gsu56, integrase |
| 26 | 1832 | 0.06 | Q749A7 | *Geobacter sulfurreducens* | Protein translocase subunit SecY |
| 27 | 1768 | 0.05 | A0A0D5N8M0 | *Geobacter sulfurreducens* | MgtE integral membrane protein |
| 28 | 1755 | 0.05 | Q74B77 | *Geobacter sulfurreducens* | Uncharacterized protein |
| 29 | 1738 | 0.05 | A1AKN8 | *Pelobacter propionicus* | Resolvase, N-terminal domain protein |
| 30 | 1723 | 0.05 | A0A0B5BJ81 | *Geobacter pickeringii* | NADH-quinone oxidoreductase subunit D (EC 1.6.5.11) (NADH dehydrogenase I subunit D) (NDH-1 subunit D) |

| | | | | | |
|---|---|---|---|---|---|
| 31 | 1715 | 0.05 | | | |
| 32 | 1661 | 0.05 | | | |
| 33 | 1624 | 0.05 | Q39Q43 | *Geobacter metallireducens* | Pyruvate-flavodoxin oxidoreductase (EC 1.2.7.-) |
| 34 | 1616 | 0.05 | A0A0B5BCT5 | *Geobacter pickeringii* | NADH dehydrogenase |
| 35 | 1612 | 0.05 | A0A0B5BF68 | *Geobacter pickeringii* | CRISPR-associated protein Cas3 |
| 36 | 1611 | 0.05 | A0A1T4LT91 | *Geobacter thiogenes* | Putative transposase |
| 37 | 1610 | 0.05 | Q39QB2 | *Geobacter metallireducens* | NADH dehydrogenase I, F subunit |
| 38 | 1606 | 0.05 | A0A1G0M9J2 | *Geobacteraceae bacterium* | Uncharacterized protein |
| 39 | 1601 | 0.05 | A0A1T4RHG2 | *Geobacter thiogenes* | Nif-specific regulatory protein |
| 40 | 1576 | 0.05 | E1QK40 | *Desulfarculus baarsii* | DNA-cytosine methyltransferase |
| 41 | 1545 | 0.05 | Q74FK9 | *Geobacter sulfurreducens* | Sigma-54-dependent transcriptional response regulator |
| 42 | 1543 | 0.05 | B9M4C2 | *Geobacter daltonii* | Squalene cyclase |
| 43 | 1523 | 0.05 | A0A1S0XXD8 | *Rhodanobacter sp.* | Histidine kinase |
| 44 | 1491 | 0.05 | A0A1G0MUH0 | *Geobacteraceae bacterium* | Integrase |
| 45 | 1486 | 0.05 | Q747H5 | *Geobacter sulfurreducens* | K(+)-insensitive pyrophosphate-energized proton pump (EC 3.6.1.1) (Membrane-bound proton-translocating pyrophosphatase) (Pyrophosphate-energized inorganic pyrophosphatase) (H(+)-PPase) |
| 46 | 1480 | 0.04 | A0A0A8WLB0 | *Geobacter sp.* | Protein TraH |
| 47 | 1465 | 0.04 | B3EB49 | *Geobacter lovleyi* | Uncharacterized protein |
| 48 | 1458 | 0.04 | A0A1F8XVG3 | *Deltaproteobacteria bacterium* | Helicase |
| 49 | 1456 | 0.04 | A0A0B5BDJ5 | *Geobacter pickeringii* | NADH-quinone oxidoreductase subunit H (EC 1.6.5.11) (NADH dehydrogenase I subunit H) (NDH-1 subunit H) |
| 50 | 1455 | 0.04 | Q747X9 | *Geobacter sulfurreducens* | Isoleucine--tRNA ligase (EC 6.1.1.5) (Isoleucyl-tRNA synthetase) (IleRS) |
| 51 | 1450 | 0.04 | A0A0D5N8S3 | *Geobacter sulfurreducens* | NADH dehydrogenase |
| 52 | 1449 | 0.04 | Q74B80 | *Geobacter sulfurreducens* | Uncharacterized protein |
| 53 | 1448 | 0.04 | A0A0B5BF21 | *Geobacter pickeringii* | Glycosyl transferase family 1 |
| 54 | 1436 | 0.04 | Q39QB8 | *Geobacter metallireducens* | NADH dehydrogenase I, L subunit |
| 55 | 1426 | 0.04 | B3EA87 | *Geobacter lovleyi* | RNA polymerase sigma factor |
| 56 | 1402 | 0.04 | A0A1G0MAK9 | *Geobacteraceae bacterium* | Sodium:proton exchanger |
| 57 | 1347 | 0.04 | Q39U60 | *Geobacter metallireducens* | Translation elongation factor G |
| 58 | 1346 | 0.04 | A0A0D5N712 | *Geobacter sulfurreducens* | Phosphoglucomutase |
| 59 | 1344 | 0.04 | | | |
| 60 | 1339 | 0.04 | Q746X3 | *Geobacter sulfurreducens* | Bifunctional protein PutA |
| 61 | 1339 | 0.04 | | | |
| 62 | 1333 | 0.04 | A0A0D5N2S0 | *Geobacter sulfurreducens* | Uncharacterized protein |

| | | | | | |
|---|---|---|---|---|---|
| 63 | 1328 | 0.04 | Q39RM8 | *Geobacter metallireducens* | Glycosyltransferase |
| 64 | 1314 | 0.04 | A0A0A8WNV3 | *Geobacter sp. OR-1* | Uncharacterized protein |
| 65 | 1313 | 0.04 | Q74C76 | *Geobacter sulfurreducens* | (R)-citramalate synthase (EC 2.3.1.182) (Citramalate synthase) |
| 66 | 1306 | 0.04 | Q74FF1 | *Geobacter sulfurreducens* | Chaperone protein ClpB |
| 67 | 1304 | 0.04 | | | |
| 68 | 1294 | 0.04 | | | |
| 69 | 1290 | 0.04 | Q74AK3 | *Geobacter sulfurreducens* | Cation-translocating P-type ATPase |
| 70 | 1288 | 0.04 | Q39QJ8 | *Geobacter metallireducens* | Flagellar biogenesis master sigma-54-dependent transcriptional response regulator |
| 71 | 1266 | 0.04 | Q39ZC8 | *Geobacter metallireducens* | Ferredoxin-dependent glutamate synthase |
| 72 | 1261 | 0.04 | C6E159 | *Geobacter sp. (s* | Uncharacterized protein |
| 73 | 1256 | 0.04 | Q748Z4 | *Geobacter sulfurreducens* | 30S ribosomal protein S3 |
| 74 | 1251 | 0.04 | A0A0B5BB65 | *Geobacter pickeringii* | Uncharacterized protein |
| 75 | 1249 | 0.04 | A0A0W8FRW6 | *hydrocarbon metagenome* | Uncharacterized protein |
| 76 | 1242 | 0.04 | A0A0B5BJ46 | *Geobacter pickeringii* | Uncharacterized protein |
| 77 | 1240 | 0.04 | A0A0B5BJ26 | *Geobacter pickeringii* | Uncharacterized protein |
| 78 | 1232 | 0.04 | Q74H58 | *Geobacter sulfurreducens* | Chaperone protein DnaJ |
| 79 | 1229 | 0.04 | Q74FM6 | *Geobacter sulfurreducens* | Pyruvate, phosphate dikinase (EC 2.7.9.1) |
| 80 | 1225 | 0.04 | Q39YK5 | *Geobacter metallireducens* | Flagellar biogenesis protein FlhA |
| 81 | 1218 | 0.04 | A0A0B5BH93 | *Geobacter pickeringii* | C-and b-type cytochrome L |
| 82 | 1212 | 0.04 | A0A0C1TL30 | *Geobacter soli* | Flagellar basal body stator protein MotB |
| 83 | 1211 | 0.04 | | | |
| 84 | 1210 | 0.04 | A0A0D5NE26 | *Geobacter sulfurreducens* | Multidrug resistance protein B |
| 85 | 1209 | 0.04 | A0A0D5N8I5 | *Geobacter sulfurreducens* | Signal recognition particle protein (Fifty-four homolog) |
| 86 | 1201 | 0.04 | E1QK35 | *Desulfarculus baarsii* | Uncharacterized protein |
| 87 | 1198 | 0.04 | Q39Y76 | *Geobacter metallireducens* | Sigma-54-dependent transcriptional response regulator |
| 88 | 1195 | 0.04 | A1AM25 | *Pelobacter propionicus* | Squalene-hopene cyclase |
| 89 | 1191 | 0.04 | Q74H82 | *Geobacter sulfurreducens* | Flagellar biogenesis master response receiver sensor histidine kinase, PAS and GAF domain-containing |
| 90 | 1180 | 0.04 | Q74B66 | *Geobacter sulfurreducens* | TrfA family protein |
| 91 | 1179 | 0.04 | A0A0B5BDE2 | *Geobacter pickeringii* | ATP phosphoribosyltransferase regulatory subunit |
| 92 | 1179 | 0.04 | A0A0D5ND99 | *Geobacter sulfurreducens* | ATPase AAA |
| 93 | 1179 | 0.04 | A0A0D5NDJ8 | *Geobacter sulfurreducens* | Stage II sporulation protein E |
| 94 | 1172 | 0.04 | B9M1Q4 | *Geobacter daltonii* | Biotin-dependent acyl-CoA carboxylase, biotin carboxylase subunit |
| 95 | 1164 | 0.04 | | | |
| 96 | 1161 | 0.04 | A0A0D5NE46 | *Geobacter sulfurreducens* | Molecular chaperone DnaJ |
| 97 | 1160 | 0.04 | A0A0D5N8E2 | *Geobacter sulfurreducens* | ATP synthase (EC 3.6.3.14) |

| | | | | | |
|---|---|---|---|---|---|
| 98 | 1159 | 0.04 | A0A0B5B7H3 | *Geobacter pickeringii* | Cytochrome C |
| 99 | 1159 | 0.04 | Q74B89 | *Geobacter sulfurreducens* | Uncharacterized protein |
| 100 | 1152 | 0.03 | Q39WU2 | *Geobacter metallireducens* | Phosphate-selective outer membrane channel |
| 101 | 1147 | 0.03 | A0A0C1TPB6 | *Geobacter soli* | Nitric-oxide reductase |
| 102 | 1140 | 0.03 | Q39ZE7 | *Geobacter metallireducens* | UvrABC system protein A (UvrA protein) (Excinuclease ABC subunit A) |
| 103 | 1137 | 0.03 | A0A0D5N7I1 | *Geobacter sulfurreducens* | NADH-quinone oxidoreductase subunit N (EC 1.6.5.11) (NADH dehydrogenase I subunit N) (NDH-1 subunit N) |
| 104 | 1132 | 0.03 | Q74GG6 | *Geobacter sulfurreducens* | Chromosomal replication initiator protein DnaA |
| 105 | 1129 | 0.03 | B3EAB4 | *Geobacter lovleyi* | Cytochrome c-552 (EC 1.7.2.2) |
| 106 | 1127 | 0.03 | A0A0D5N7R7 | *Geobacter sulfurreducens* | GntR family transcriptional regulator |
| 107 | 1127 | 0.03 | | | |
| 108 | 1122 | 0.03 | A0A0B5BK17 | *Geobacter pickeringii* | Diguanylate cyclase |
| 109 | 1121 | 0.03 | A0A1F8XUV7 | *Deltaproteobacteria bacterium* | Uncharacterized protein |
| 110 | 1121 | 0.03 | Q747M6 | *Geobacter sulfurreducens* | Ribonuclease G |
| 111 | 1119 | 0.03 | A0A0A8WQV9 | *Geobacter sp.* | Protein TraU |
| 112 | 1114 | 0.03 | A0A0D5N5P7 | *Geobacter sulfurreducens* | Chemotaxis protein |
| 113 | 1112 | 0.03 | A0A0A8WQR9 | *Geobacter sp.* | Helix-turn-helix domain protein |
| 114 | 1110 | 0.03 | A0A0B5BI89 | *Geobacter pickeringii* | Type II secretion system protein |
| 115 | 1110 | 0.03 | A0LQ14 | *Syntrophobacter fumaroxidans* | Uncharacterized protein |
| 116 | 1100 | 0.03 | A5G9A7 | *Geobacter uraniireducens* | NADH dehydrogenase subunit M (EC 1.6.5.3) |
| 117 | 1092 | 0.03 | Q74GY0 | *Geobacter sulfurreducens* | ATP synthase subunit beta (EC 3.6.3.14) (ATP synthase F1 sector subunit beta) (F-ATPase subunit beta) |
| 118 | 1086 | 0.03 | B5EFV6 | *Geobacter bemidjiensis* | Bifunctional protein PutA |
| 119 | 1081 | 0.03 | A0A0B5BD43 | *Geobacter pickeringii* | ABC transporter permease |
| 120 | 1081 | 0.03 | Q74FW2 | *Geobacter sulfurreducens* | Succinyl:acetate coenzyme A transferase |
| 121 | 1075 | 0.03 | A0A0B5BHV9 | *Geobacter pickeringii* | ATPase AAA |
| 122 | 1068 | 0.03 | A0A0D5N7M4 | *Geobacter sulfurreducens* | Chemotaxis protein |
| 123 | 1062 | 0.03 | Q39ZK4 | *Geobacter metallireducens* | Bifunctional glutamine synthetase adenylyltransferase/adenylyl-removing enzyme (ATP:glutamine synthetase adenylyltransferase) (ATase) [Includes: Glutamine synthetase adenylyl-L-tyrosine phosphorylase (EC 2.7.7.89) (Adenylyl removase) (AR) (AT-N); Glutamine synthetase adenylyl transferase (EC 2.7.7.42) (Adenylyl transferase) (AT) (AT-C)] |
| 124 | 1059 | 0.03 | A0A0B5B8A8 | *Geobacter pickeringii* | Valine--tRNA ligase (EC 6.1.1.9) (Valyl-tRNA synthetase) (ValRS) |
| 125 | 1058 | 0.03 | B3E9R4 | *Geobacter lovleyi* | Isocitrate dehydrogenase, NADP-dependent (EC 1.1.1.42) |
| 126 | 1055 | 0.03 | A0A0B5BCL9 | *Geobacter pickeringii* | Uncharacterized protein |

| | | | | | |
|---|---|---|---|---|---|
| 127 | 1054 | 0.03 | Q74GY2 | *Geobacter sulfurreducens* | ATP synthase subunit alpha (EC 3.6.3.14) (ATP synthase F1 sector subunit alpha) (F-ATPase subunit alpha) |
| 128 | 1050 | 0.03 | Q74FD5 | *Geobacter sulfurreducens* | Hydroxylamine reductase (EC 1.7.99.1) (Hybrid-cluster protein) (HCP) (Prismane protein) |
| 129 | 1047 | 0.03 | Q39QJ5 | *Geobacter metallireducens* | Sensor histidine kinase CheA associated with MCPs of classes 40H and 40+24H |
| 130 | 1043 | 0.03 | | | |
| 131 | 1038 | 0.03 | A0A0A8WPS5 | *Geobacter sp.* | Efflux pump membrane transporter |
| 132 | 1035 | 0.03 | A5GB28 | *Geobacter uraniireducens* | Plasmid maintenance system antidote protein, XRE family |
| 133 | 1033 | 0.03 | Q39QU2 | *Geobacter metallireducens* | Phosphoenolpyruvate carboxykinase (ATP) (PCK) (PEP carboxykinase) (PEPCK) (EC 4.1.1.49) |
| 134 | 1032 | 0.03 | A0A0A8WN28 | *Geobacter sp.* | Protein TraN |
| 135 | 1028 | 0.03 | A0A1T4LST4 | *Geobacter thiogenes* | Multidrug efflux pump subunit AcrB |
| 136 | 1025 | 0.03 | A0A1F9PMW1 | *Desulfuromonadales bacterium* | DNA-directed RNA polymerase subunit beta (RNAP subunit beta) (EC 2.7.7.6) (RNA polymerase subunit beta) (Transcriptase subunit beta) |
| 137 | 1025 | 0.03 | Q39QP7 | *Geobacter metallireducens* | Lon protease (EC 3.4.21.53) (ATP-dependent protease La) |
| 138 | 1022 | 0.03 | D4H382 | *Denitrovibrio acetiphilus* | DNA-directed RNA polymerase subunit beta' (RNAP subunit beta') (EC 2.7.7.6) (RNA polymerase subunit beta') (Transcriptase subunit beta') |
| 139 | 1019 | 0.03 | A5GDX4 | *Geobacter uraniireducens* | DNA gyrase subunit B (EC 5.99.1.3) |
| 140 | 1018 | 0.03 | A0A0B5BK56 | *Geobacter pickeringii* | Lon protease (EC 3.4.21.53) (ATP-dependent protease La) |
| 141 | 1018 | 0.03 | D4H381 | *Denitrovibrio acetiphilus* | DNA-directed RNA polymerase subunit beta (RNAP subunit beta) (EC 2.7.7.6) (RNA polymerase subunit beta) (Transcriptase subunit beta) |
| 142 | 1017 | 0.03 | Q39Q36 | *Geobacter metallireducens* | Heterodisulfide oxidoreductase, FAD-binding and iron-sulfur cluster-binding subunit A |
| 143 | 1014 | 0.03 | A0A0D5N8Q2 | *Geobacter sulfurreducens* | Uncharacterized protein |
| 144 | 1009 | 0.03 | A5GFJ7 | *Geobacter sulfurreducens* | Erythronolide synthase (EC 2.3.1.94) |
| 145 | 997 | 0.03 | A5GCN0 | *Geobacter sulfurreducens* | Short chain amide porin |
| 146 | 984 | 0.03 | A0A4W3 | *Geobacter sulfurreducens* | Biosynthetic arginine decarboxylase (ADC) (EC 4.1.1.19) |
| 147 | 977 | 0.03 | A0A0D5N8X4 | *Geobacter sulfurreducens* | Flagellar M-ring protein |
| 148 | 976 | 0.03 | Q39PS2 | *Geobacter metallireducens* | Transcription-repair-coupling factor (TRCF) (EC 3.6.4.-) |
| 149 | 976 | 0.03 | Q39ZH1 | *Geobacter metallireducens* | Glutamine--fructose-6-phosphate aminotransferase [isomerizing] (EC 2.6.1.16) (D-fructose-6-phosphate amidotransferase) (GFAT) (Glucosamine-6-phosphate synthase) (Hexosephosphate aminotransferase) (L-glutamine--D-fructose-6-phosphate amidotransferase) |
| 150 | 974 | 0.03 | A0A0B5BIQ7 | *Geobacter pickeringii* | Flagellar basal-body rod protein FlgF (Flagellar hook protein FlgE) |
| 151 | 974 | 0.03 | A5GAY2 | *Geobacter uraniireducens* | DNA-directed RNA polymerase subunit beta' (RNAP subunit beta') (EC 2.7.7.6) (RNA polymerase subunit beta') (Transcriptase subunit beta') |
| 152 | 972 | 0.03 | A0A0B5BIS9 | *Geobacter pickeringii* | Multidrug transporter AcrB |
| 153 | 969 | 0.03 | | | |
| 154 | 960 | 0.03 | Q39T74 | *Geobacter metallireducens* | GTP/GDP 3'-pyrophosphokinase and (P)ppGpp 3'-pyrophosphohydrolase |

| | | | | | |
|---|---|---|---|---|---|
| 155 | 960 | 0.03 | A5GAY1 | *Geobacter uraniireducens* | DNA-directed RNA polymerase subunit beta (RNAP subunit beta) (EC 2.7.7.6) (RNA polymerase subunit beta) (Transcriptase subunit beta) |
| 156 | 958 | 0.03 | A0A1T4RGQ7 | *Geobacter thiogenes* | NAD+---dinitrogen-reductase ADP-D-ribosyltransferase |
| 157 | 958 | 0.03 | Q39QA3 | *Geobacter metallireducens* | ATP synthase subunit a (ATP synthase F0 sector subunit a) (F-ATPase subunit 6) |
| 158 | 957 | 0.03 | A0A0D5N952 | *Geobacter sulfurreducens* | Flagellar motor switch protein FliG |
| 159 | 957 | 0.03 | P61679 | *Geobacter sulfurreducens* | UDP-N-acetylmuramate--L-alanine ligase (EC 6.3.2.8) (UDP-N-acetylmuramoyl-L-alanine synthetase) |
| 160 | 956 | 0.03 | A0A0D5N8G6 | *Geobacter sulfurreducens* | Cation transporter |
| 161 | 953 | 0.03 | Q39R25 | *Geobacter metallireducens* | Sensor diguanylate cyclase/phosphodiesterase, PAS, PAS and PAS domain-containing |
| 162 | 952 | 0.03 | A0A0D5NAK5 | *Geobacter sulfurreducens* | Bifunctional uridylyltransferase/uridylyl-removing enzyme (UTase/UR) (Bifunctional [protein-PII] modification enzyme) (Bifunctional nitrogen sensor protein) [Includes: [Protein-PII]-UMP uridylyl-removing enzyme (UR) (EC 3.1.4.-); [Protein-PII] uridylyltransferase (PII uridylyltransferase) (UTase) (EC 2.7.7.59)] |
| 163 | 950 | 0.03 | | | |
| 164 | 947 | 0.03 | A0A0B5BE38 | *Geobacter pickeringii* | Transcription-repair-coupling factor (TRCF) (EC 3.6.4.-) |
| 165 | 946 | 0.03 | Q74B81 | *Geobacter sulfurreducens* | Uncharacterized protein |
| 166 | 945 | 0.03 | Q747G2 | *Geobacter sulfurreducens* | Outer membrane channel OmpJ |
| 167 | 942 | 0.03 | Q39QT7 | *Geobacter metallireducens* | Cytochrome c |
| 168 | 941 | 0.03 | A0A0A8WMX3 | *Geobacter sp.* | Type III restriction enzyme, res subunit |
| 169 | 939 | 0.03 | A0A0D5NCW1 | *Geobacter sulfurreducens* | Histidine kinase |
| 170 | 932 | 0.03 | A0A0B5BH05 | *Geobacter pickeringii* | Malic enzyme (EC 1.1.1.40) |
| 171 | 929 | 0.03 | A0A0B5BIF0 | *Geobacter pickeringii* | NADH-ubiquinone oxidoreductase subunit 3 |
| 172 | 929 | 0.03 | Q39RL8 | *Geobacter metallireducens* | Undecaprenyl-diphospho-oligosaccharide flippase |
| 173 | 925 | 0.03 | A0A0D5NCN6 | *Geobacter sulfurreducens* | Thiol:disulfide interchange protein |
| 174 | 922 | 0.03 | A0A0D5NDD6 | *Geobacter sulfurreducens* | Type II secretion system protein E |
| 175 | 918 | 0.03 | A0A0D5ND20 | *Geobacter sulfurreducens* | Amino acid dehydrogenase |
| 176 | 913 | 0.03 | A0A0B5B5Z5 | *Geobacter pickeringii* | DNA polymerase III subunit beta (EC 2.7.7.7) |
| 177 | 913 | 0.03 | Q39QK7 | *Geobacter metallireducens* | Outer membrane channel OmpJ |
| 178 | 911 | 0.03 | A0A0B5BDE7 | *Geobacter pickeringii* | Fis family transcriptional regulator |
| 179 | 905 | 0.03 | A0A0B5BDG1 | *Geobacter pickeringii* | tRNA uridine 5-carboxymethylaminomethyl modification enzyme MnmG (Glucose-inhibited division protein A) |
| 180 | 904 | 0.03 | Q39QD0 | *Geobacter metallireducens* | Periplasmically oriented, membrane-bound [NiFe]-hydrogenase, large subunit |
| 181 | 900 | 0.03 | Q748I6 | *Geobacter sulfurreducens* | Metalloprotease domain protein, M6 family |
| 182 | 897 | 0.03 | A0A0B5BKE7 | *Geobacter pickeringii* | NADH-quinone oxidoreductase subunit N (EC 1.6.5.11) (NADH dehydrogenase I subunit N) (NDH-1 subunit N) |

| | | | | | |
|---|---|---|---|---|---|
| 183 | 895 | 0.03 | Q74GT3 | *Geobacter sulfurreducens* | LL-diaminopimelate aminotransferase (DAP-AT) (DAP-aminotransferase) (LL-DAP-aminotransferase) (EC 2.6.1.83) |
| 184 | 894 | 0.03 | Q39QB3 | *Geobacter metallireducens* | NADH dehydrogenase I, G subunit |
| 185 | 889 | 0.03 | A0A0A8WNZ4 | *Geobacter sp.* | DNA gyrase subunit A (EC 5.99.1.3) |
| 186 | 889 | 0.03 | B5EGM4 | *Geobacter bemidjiensis* | Glycerol/1,2-propanediol dehydratase |
| 187 | 887 | 0.03 | A0R7X1 | *Pelobacter propionicus* | Type III restriction enzyme, res subunit |
| 188 | 882 | 0.03 | Q749N5 | *Geobacter sulfurreducens* | Acetate kinase (EC 2.7.2.1) (Acetokinase) |
| 189 | 879 | 0.03 | Q39UR9 | *Geobacter metallireducens* | Carbamoyl-phosphate synthase large chain (EC 6.3.5.5) (Carbamoyl-phosphate synthetase ammonia chain) |
| 190 | 877 | 0.03 | A0A0D5N3I5 | *Geobacter sulfurreducens* | Uncharacterized protein |
| 191 | 877 | 0.03 | Q74C83 | *Geobacter sulfurreducens* | ATP-dependent Clp protease ATP-binding subunit ClpX |
| 192 | 875 | 0.03 | A0A0D5N400 | *Geobacter sulfurreducens* | Uncharacterized protein |
| 193 | 874 | 0.03 | A0A0D5N6Q1 | *Geobacter sulfurreducens* | FeS-binding protein |
| 194 | 870 | 0.03 | I7EF12 | *Geobacter sulfurreducens* | Uncharacterized protein |
| 195 | 869 | 0.03 | A0A0B5BK89 | *Geobacter pickeringii* | FAD-binding protein |
| 196 | 868 | 0.03 | B3EA93 | *Geobacter lovleyi* | 2-alkenal reductase (EC 1.3.1.74) |
| 197 | 867 | 0.03 | Q39SJ2 | *Geobacter metallireducens* | Nitrogen fixation master sigma-54-dependent transcriptional response regulator |
| 198 | 866 | 0.03 | A0A0A8WQW8 | *Geobacter sp.* | Protein TraE |
| 199 | 866 | 0.03 | | | |
| 200 | 861 | 0.03 | Q39V52 | *Geobacter metallireducens* | GAF sensor methyl-accepting chemotaxis sensory transducer, class 40H |
| 201 | 858 | 0.03 | A0A0B5B668 | *Geobacter pickeringii* | Peptidase U32 |
| 202 | 858 | 0.03 | A0A1T4R5F2 | *Geobacter thiogenes* | Putative transposase |
| 203 | 858 | 0.03 | Q39XL9 | *Geobacter metallireducens* | Sensor sigma-54-dependent transcriptional regulator, GAF and GAF domain-containing |
| 204 | 850 | 0.03 | | | |
| 205 | 848 | 0.03 | A0A0B5BJV3 | *Geobacter pickeringii* | ABC transporter substrate-binding protein |
| 206 | 848 | 0.03 | Q39RI3 | *Geobacter metallireducens* | Sensor histidine kinase, HAMP domain-containing, 2 heme-binding sites |
| 207 | 840 | 0.03 | Q74CR5 | *Geobacter sulfurreducens* | Serine hydroxymethyltransferase (SHMT) (Serine methylase) (EC 2.1.2.1) |
| 208 | 840 | 0.03 | Q39X91 | *Geobacter metallireducens* | DNA topoisomerase 1 (EC 5.99.1.2) (DNA topoisomerase I) |
| 209 | 840 | 0.03 | Q39X99 | *Geobacter metallireducens* | UDP-glucuronate 4-dehydrogenase, decarboxylating |
| 210 | 839 | 0.03 | Q74C70 | *Geobacter sulfurreducens* | Phosphoglucosamine mutase (EC 5.4.2.10) |
| 211 | 838 | 0.03 | Q39QY3 | *Geobacter metallireducens* | Efflux pump, RND family, inner membrane protein |
| 212 | 837 | 0.03 | A5GF11 | *Geobacter uraniireducens* | Acetyl-CoA carboxylase carboxyltransferase subunit alpha (EC 6.3.4.14) |
| 213 | 837 | 0.03 | Q39U54 | *Geobacter metallireducens* | Sensor histidine kinase cyclic nucleotide phosphodiesterase, GAF, GAF and HD-GYP-related domain-containing |
| 214 | 837 | 0.03 | Q74B83 | *Geobacter sulfurreducens* | ATPase, AAA family, CbbQ_C domain-containing |
| 215 | 836 | 0.03 | A0A0B5BJ17 | *Geobacter pickeringii* | UvrABC system protein C (Protein UvrC) (Excinuclease ABC subunit C) |

| | | | | | |
|---|---|---|---|---|---|
| 216 | 835 | 0.03 | Q39VY8 | *Geobacter metallireducens* | Glutamine synthetase (EC 6.3.1.2) |
| 217 | 833 | 0.03 | A0A0B5BK21 | *Geobacter pickeringii* | Inner membrane cytochrome H |
| 218 | 833 | 0.03 | A0A0D5N949 | *Geobacter sulfurreducens* | Response regulator |
| 219 | 832 | 0.03 | A0A0B5BAC1 | *Geobacter pickeringii* | Dihydrolipoyl dehydrogenase (EC 1.8.1.4) |
| 220 | 831 | 0.03 | B9M474 | *Geobacter daltonii* | Uncharacterized protein |
| 221 | 830 | 0.03 | A5GC25 | *Geobacter uraniireducens* | Propionyl-CoA carboxylase (EC 6.4.1.3) |
| 222 | 829 | 0.03 | A0A0B5BC52 | *Geobacter pickeringii* | General secretion pathway protein GspD |
| 223 | 821 | 0.02 | Q39VY0 | *Geobacter metallireducens* | Malate dehydrogenase (EC 1.1.1.37) |
| 224 | 821 | 0.02 | Q74GF2 | *Geobacter sulfurreducens* | Metal-dependent phosphohydrolase, HDOD domain-containing |
| 225 | 816 | 0.02 | A0A0D5NB12 | *Geobacter sulfurreducens* | Aldehyde dehydrogenase |
| 226 | 815 | 0.02 | A0A0A8WV23 | *Geobacter sp.* | Protein translocase subunit SecD |
| 227 | 814 | 0.02 | A1ANK6 | *Pelobacter propionicus* | Peptidase C1A, papain |
| 228 | 813 | 0.02 | | | |
| 229 | 808 | 0.02 | Q39RM6 | *Geobacter metallireducens* | Glycosyltransferase |
| 230 | 808 | 0.02 | A1ALT5 | *Pelobacter propionicus* | DNA-directed RNA polymerase subunit beta' (RNAP subunit beta') (EC 2.7.7.6) (RNA polymerase subunit beta') (Transcriptase subunit beta') |
| 231 | 806 | 0.02 | Q74H61 | *Geobacter sulfurreducens* | Heat-inducible transcription repressor HrcA |
| 232 | 804 | 0.02 | A0A0B5BH19 | *Geobacter pickeringii* | Uncharacterized protein |
| 233 | 803 | 0.02 | P61667 | *Geobacter sulfurreducens* | DNA mismatch repair protein MutS |
| 234 | 802 | 0.02 | A0A0B5BC86 | *Geobacter pickeringii* | Multicopper oxidase |
| 235 | 799 | 0.02 | B9M5J7 | *Geobacter daltonii* | CRISPR-associated protein Csd2 |
| 236 | 798 | 0.02 | Q39UW3 | *Geobacter metallireducens* | Succinyl:acetate coenzyme A transferase |
| 237 | 796 | 0.02 | Q39Q79 | *Geobacter metallireducens* | Winged-helix transcriptional response regulator |
| 238 | 795 | 0.02 | Q39ZL2 | *Geobacter metallireducens* | TrkA domain protein |
| 239 | 791 | 0.02 | Q747G4 | *Geobacter sulfurreducens* | (R)-methylmalonyl-CoA mutase, isobutyryl-CoA mutase-like catalytic subunit |
| 240 | 790 | 0.02 | P60401 | *Geobacter sulfurreducens* | 50S ribosomal protein L2 |
| 241 | 789 | 0.02 | A0A0B5BIH3 | *Geobacter pickeringii* | Uncharacterized protein |
| 242 | 789 | 0.02 | N8ZVC9 | *Acinetobacter gerneri* | Uncharacterized protein |
| 243 | 789 | 0.02 | Q39YH3 | *Geobacter metallireducens* | UDP-N-acetylglucosamine 4,6-dehydratase and UDP-2-acetamido-2,6-dideoxy-alpha-D-xylo-4-hexulose 5-epimerase |
| 244 | 788 | 0.02 | A0A0B5BCZ8 | *Geobacter pickeringii* | Succinate dehydrogenase (EC 1.3.5.1) |
| 245 | 788 | 0.02 | A0A1F9PXI1 | *Desulfuromonadales bacterium* | Cytosine-specific methyltransferase (EC 2.1.1.37) |
| 246 | 787 | 0.02 | B0L930 | *uncultured Geobacter sp.* | Tuf (Fragment) |
| 247 | 787 | 0.02 | Q39Z79 | *Geobacter metallireducens* | Twitching motility pilus retraction ATPase |
| 248 | 785 | 0.02 | A0A0D5N732 | *Geobacter sulfurreducens* | Transcription-repair-coupling factor (TRCF) (EC 3.6.4.-) |

| | | | | | |
|---|---|---|---|---|---|
| 249 | 785 | 0.02 | Q39W73 | *Geobacter metallireducens* | 2-isopropylmalate synthase (EC 2.3.3.13) (Alpha-IPM synthase) (Alpha-isopropylmalate synthase) |
| 250 | 783 | 0.02 | A0A0B5BB89 | *Geobacter pickeringii* | Chromosome partition protein Smc |
| 251 | 781 | 0.02 | A0A0A8WLL1 | *Geobacter sp.* | Uncharacterized protein |
| 252 | 781 | 0.02 | Q39ZQ2 | *Geobacter metallireducens* | Uncharacterized protein |
| 253 | 775 | 0.02 | A0A0B5BBC9 | *Geobacter pickeringii* | Type I citrate synthase (EC 2.3.3.1) |
| 254 | 775 | 0.02 | A0A0D5NAW4 | *Geobacter sulfurreducens* | Cation transporter |
| 255 | 773 | 0.02 | | | |
| 256 | 772 | 0.02 | A0A0D5N7Y6 | *Geobacter sulfurreducens* | Chemotaxis protein CheA |
| 257 | 767 | 0.02 | Q747K3 | *Geobacter sulfurreducens* | UvrABC system protein B (Protein UvrB) (Excinuclease ABC subunit B) |
| 258 | 766 | 0.02 | A0A0A8WRD3 | *Geobacter sp.* | Putative ABC transporter ATP-binding protein YkpA |
| 259 | 766 | 0.02 | A0A0D5NAV3 | *Geobacter sulfurreducens* | Peptidase S41 |
| 260 | 765 | 0.02 | B5E7W0 | *Geobacter bemidjiensis* | Lipoprotein, putative |
| 261 | 765 | 0.02 | Q74B64 | *Geobacter sulfurreducens* | RNA polymerase sigma factor |
| 262 | 764 | 0.02 | A0A0B5BJT6 | *Geobacter pickeringii* | Membrane protein insertase YidC (Foldase YidC) (Membrane integrase YidC) (Membrane protein YidC) |
| 263 | 763 | 0.02 | A0A0D5N7N0 | *Geobacter sulfurreducens* | Glutamate synthase |
| 264 | 763 | 0.02 | Q39SI5 | *Geobacter metallireducens* | Fumarate hydratase class I (EC 4.2.1.2) |
| 265 | 763 | 0.02 | Q39WG2 | *Geobacter metallireducens* | Chromosome partition protein Smc |
| 266 | 760 | 0.02 | Q39RM7 | *Geobacter metallireducens* | Glycosyltransferase |
| 267 | 759 | 0.02 | A0A0A8WYA5 | *Geobacter sp.* | DNA-directed RNA polymerase subunit beta' (RNAP subunit beta') (EC 2.7.7.6) (RNA polymerase subunit beta') (Transcriptase subunit beta') |
| 268 | 759 | 0.02 | A0A0B5BGY7 | *Geobacter pickeringii* | Phosphoglucomutase |
| 269 | 759 | 0.02 | Q74B87 | *Geobacter sulfurreducens* | Uncharacterized protein |
| 270 | 757 | 0.02 | P60789 | *Geobacter sulfurreducens* | Elongation factor 4 (EF-4) (EC 3.6.5.n1) (Ribosomal back-translocase LepA) |
| 271 | 756 | 0.02 | A0A0B5BGI9 | *Geobacter pickeringii* | Lon protease (EC 3.4.21.53) (ATP-dependent protease La) |
| 272 | 756 | 0.02 | A0A0C1TQJ7 | *Geobacter soli* | Uncharacterized protein |
| 273 | 750 | 0.02 | A1AN79 | *Pelobacter propionicus* | Adenosylmethionine-8-amino-7-oxononanoate aminotransferase (EC 2.6.1.62) (7,8-diamino-pelargonic acid aminotransferase) (DAPA AT) (DAPA aminotransferase) (7,8-diaminononanoate synthase) (DANS) (Diaminopelargonic acid synthase) |
| 274 | 750 | 0.02 | B3EA94 | *Geobacter lovleyi* | Uncharacterized protein |
| 275 | 747 | 0.02 | A0A0C1QTJ3 | *Geobacter soli* | Hemolysin D |
| 276 | 744 | 0.02 | A0A0B5BGU8 | *Geobacter pickeringii* | Chemotaxis protein |
| 277 | 744 | 0.02 | Q74H55 | *Geobacter sulfurreducens* | Serine--tRNA ligase (EC 6.1.1.11) (Seryl-tRNA synthetase) (SerRS) (Seryl-tRNA(Ser/Sec) synthetase) |
| 278 | 743 | 0.02 | Q747L0 | *Geobacter sulfurreducens* | Glycoside hydrolase, family 57 |

| | | | | | |
|---|---|---|---|---|---|
| 279 | 742 | 0.02 | A0A0B5BGV8 | *Geobacter pickeringii* | Zinc protease |
| 280 | 742 | 0.02 | A0A0B5BIW8 | *Geobacter pickeringii* | Aminotransferase |
| 281 | 742 | 0.02 | A0A0D5N848 | *Geobacter sulfurreducens* | Chemotaxis protein methyltransferase (EC 2.1.1.80) |
| 282 | 742 | 0.02 | Q39X64 | *Geobacter metallireducens* | Cell shape-determining protein MreB |
| 283 | 740 | 0.02 | A0A0B5B8F2 | *Geobacter pickeringii* | Sodium transporter |
| 284 | 740 | 0.02 | | | |
| 285 | 739 | 0.02 | A0A0B5BKQ2 | *Geobacter pickeringii* | Branched-chain amino acid ABC transporter substrate-binding protein |
| 286 | 739 | 0.02 | Q39QK5 | *Geobacter metallireducens* | Uncharacterized protein |
| 287 | 736 | 0.02 | A0A0B5B672 | *Geobacter pickeringii* | FeS-binding protein |
| 288 | 736 | 0.02 | A0A0B5B6P6 | *Geobacter pickeringii* | Histidine kinase |
| 289 | 736 | 0.02 | B3EAA0 | *Geobacter lovleyi* | Uncharacterized protein |
| 290 | 735 | 0.02 | Q74BT7 | *Geobacter sulfurreducens* | Sensor histidine kinase cyclic nucleotide phosphodiesterase, GAF, GAF and HD-GYP-related domain-containing |
| 291 | 735 | 0.02 | Q74F57 | *Geobacter sulfurreducens* | Membrane protein, major facilitator superfamily |
| 292 | 734 | 0.02 | Q39WR2 | *Geobacter metallireducens* | CRISPR-associated endonuclease Cas1 (EC 3.1.-.-) |
| 293 | 730 | 0.02 | A5GFJ6 | *Geobacter uraniireducens* | Beta-hydroxyacyl-(Acyl-carrier-protein) dehydratase, FabA/FabZ |
| 294 | 728 | 0.02 | A0A0C1U608 | *Geobacter soli* | Chemotaxis protein |
| 295 | 725 | 0.02 | Q39Y18 | *Geobacter metallireducens* | Transcription termination/antitermination protein NusG |
| 296 | 724 | 0.02 | Q39UA2 | *Geobacter metallireducens* | Adenylosuccinate lyase (ASL) (EC 4.3.2.2) (Adenylosuccinase) |
| 297 | 723 | 0.02 | A0A0B5BE21 | *Geobacter pickeringii* | Histidine kinase |
| 298 | 723 | 0.02 | Q39PY9 | *Geobacter metallireducens* | Histidine kinase (EC 2.7.13.3) |
| 299 | 719 | 0.02 | A0A1F9P4Q4 | *Desulfuromonadales bacterium* | Uncharacterized protein |
| 300 | 719 | 0.02 | A0A1G0MT57 | *Geobacteraceae bacterium* | Uncharacterized protein |
| 301 | 717 | 0.02 | A0A0D5N8K2 | *Geobacter sulfurreducens* | RND transporter |
| 302 | 717 | 0.02 | B3EB59 | *Geobacter lovleyi* | Uncharacterized protein |
| 303 | 716 | 0.02 | A0A0D5N5Y8 | *Geobacter sulfurreducens* | Cell division protein FtsA |
| 304 | 716 | 0.02 | A0A0D5N8X6 | *Geobacter sulfurreducens* | Chemotaxis protein |
| 305 | 715 | 0.02 | A0A0D5N627 | *Geobacter sulfurreducens* | SpoVR family protein |
| 306 | 714 | 0.02 | A0A0D5N4E2 | *Geobacter sulfurreducens* | Conjugal transfer protein TraB |
| 307 | 713 | 0.02 | A5GDX5 | *Geobacter uraniireducens* | DNA gyrase subunit A (EC 5.99.1.3) |
| 308 | 710 | 0.02 | Q39WW8 | *Geobacter metallireducens* | Periplasmic serine endoprotease DegP-like (EC 3.4.21.107) |
| 309 | 709 | 0.02 | A0A0B5BCB2 | *Geobacter pickeringii* | Nitrogen fixation protein NifE |
| 310 | 708 | 0.02 | Q74GU4 | *Geobacter sulfurreducens* | Acetylglutamate kinase (EC 2.7.2.8) (N-acetyl-L-glutamate 5-phosphotransferase) (NAG kinase) (NAGK) |
| 311 | 707 | 0.02 | A0A0B5B9H5 | *Geobacter pickeringii* | Uncharacterized protein |

| | | | | | |
|---|---|---|---|---|---|
| 312 | 707 | 0.02 | A5GC26 | *Geobacter uraniireducens* | Biotin/lipoyl attachment domain-containing protein |
| 313 | 707 | 0.02 | Q39QM2 | *Geobacter metallireducens* | K(+)-insensitive pyrophosphate-energized proton pump (EC 3.6.1.1) (Membrane-bound proton-translocating pyrophosphatase) (Pyrophosphate-energized inorganic pyrophosphatase) (H(+)-PPase) |
| 314 | 706 | 0.02 | A5G3L3 | *Geobacter uraniireducens* | Isocitrate dehydrogenase, NADP-dependent (EC 1.1.1.42) |
| 315 | 705 | 0.02 | A0A0D5N3E1 | *Geobacter sulfurreducens* | Conjugal transfer protein TrbC |
| 316 | 705 | 0.02 | Q39YR4 | *Geobacter metallireducens* | FtsK/SpoIIIE domain protein |
| 317 | 705 | 0.02 | Q748Y3 | *Geobacter sulfurreducens* | 50S ribosomal protein L1 |
| 318 | 704 | 0.02 | A0A0D5NCM1 | *Geobacter sulfurreducens* | Transposase |
| 319 | 700 | 0.02 | A0A0B5BBZ4 | *Geobacter pickeringii* | DNA polymerase I (EC 2.7.7.7) |
| 320 | 700 | 0.02 | Q39QN4 | *Geobacter metallireducens* | Pentapeptide repeat protein |
| 321 | 699 | 0.02 | Q746S4 | *Geobacter sulfurreducens* | NADH-quinone oxidoreductase subunit B/C/D (EC 1.6.5.11) (NADH dehydrogenase I subunit B/C/D) (NDH-1 subunit B/C/D) |
| 322 | 698 | 0.02 | A0A0B5B7K6 | *Geobacter pickeringii* | Cytochrome C biogenesis protein ResC |
| 323 | 696 | 0.02 | A0A0B5BK07 | *Geobacter pickeringii* | Histidine kinase |
| 324 | 696 | 0.02 | B9M149 | *Geobacter daltonii* | Translation-regulating membrane GTPase TypA |
| 325 | 695 | 0.02 | A0A0B5BE51 | *Geobacter pickeringii* | DNA replication and repair protein RecF |
| 326 | 695 | 0.02 | A0A1G0LRW7 | *Geobacteraceae bacterium* | Uncharacterized protein |
| 327 | 692 | 0.02 | P61946 | *Geobacter sulfurreducens* | S-adenosylmethionine synthase (AdoMet synthase) (EC 2.5.1.6) (MAT) (Methionine adenosyltransferase) |
| 328 | 690 | 0.02 | A0A0B5BCP9 | *Geobacter pickeringii* | Uncharacterized protein |
| 329 | 689 | 0.02 | A0A0B5BCH1 | *Geobacter pickeringii* | Hydrogenase 2 large subunit (EC 1.12.99.6) |
| 330 | 689 | 0.02 | A0A0B5BG42 | *Geobacter pickeringii* | Vitamin B12-dependent ribonucleotide reductase (EC 1.17.4.1) |
| 331 | 689 | 0.02 | Q74AR7 | *Geobacter sulfurreducens* | Membrane-associated metal-dependent phosphohydrolase, HDc domain-containing |
| 332 | 688 | 0.02 | B9M6V1 | *Geobacter daltonii* | DNA-directed RNA polymerase subunit beta' (RNAP subunit beta') (EC 2.7.7.6) (RNA polymerase subunit beta') (Transcriptase subunit beta') |
| 333 | 686 | 0.02 | A0A0B5B7G6 | *Geobacter pickeringii* | Cytochrome C |
| 334 | 685 | 0.02 | A0A0D5N930 | *Geobacter sulfurreducens* | Cytochrome C |
| 335 | 685 | 0.02 | Q746T2 | *Geobacter sulfurreducens* | NADH-quinone oxidoreductase subunit H 2 (EC 1.6.5.11) (NADH dehydrogenase I subunit H 2) (NDH-1 subunit H 2) |
| 336 | 685 | 0.02 | Q39RQ5 | *Geobacter metallireducens* | Chaperone protein ClpB |
| 337 | 683 | 0.02 | A0A0D5N6L1 | *Geobacter sulfurreducens* | Histidine kinase |
| 338 | 682 | 0.02 | A0A0B5BDP4 | *Geobacter pickeringii* | Polya polymerase |
| 339 | 682 | 0.02 | Q74D56 | *Geobacter sulfurreducens* | Aspartate--tRNA(Asp/Asn) ligase (EC 6.1.1.23) (Aspartyl-tRNA synthetase) (AspRS) (Non-discriminating aspartyl-tRNA synthetase) (ND-AspRS) |

| | | | | | |
|---|---|---|---|---|---|
| 340 | 681 | 0.02 | Q39ZM0 | *Geobacter metallireducens* | Iron-sulfur cluster-binding sigma-54-dependent transcriptional regulator, FehydlgC and FeS domain-containing |
| 341 | 681 | 0.02 | Q74D04 | *Geobacter sulfurreducens* | Threonine--tRNA ligase (EC 6.1.1.3) (Threonyl-tRNA synthetase) (ThrRS) |
| 342 | 680 | 0.02 | A0A0C1TL33 | *Geobacter soli* | Uncharacterized protein |
| 343 | 680 | 0.02 | A0A0D5N2Y9 | *Geobacter sulfurreducens* | Diguanylate cyclase |
| 344 | 680 | 0.02 | A0A0D5N645 | *Geobacter sulfurreducens* | Penicillin-binding protein |
| 345 | 680 | 0.02 | | | |
| 346 | 679 | 0.02 | A5G8W9 | *Geobacter uraniireducens* | Flagellar hook-associated protein 2 (HAP2) (Flagellar cap protein) |
| 347 | 679 | 0.02 | B3E8E5 | *Geobacter lovleyi* | Cell surface receptor IPT/TIG domain protein |
| 348 | 679 | 0.02 | Q39QJ4 | *Geobacter metallireducens* | Chemotaxis protein methyltransferase (EC 2.1.1.80) |
| 349 | 678 | 0.02 | A0A0D5N435 | *Geobacter sulfurreducens* | Multidrug transporter |
| 350 | 678 | 0.02 | Q39XG6 | *Geobacter metallireducens* | Pyruvate carboxylase (EC 6.4.1.1) |
| 351 | 677 | 0.02 | A1AKK9 | *Pelobacter propionicus* | Conserved hypothetical membrane protein |
| 352 | 676 | 0.02 | Q74GH2 | *Geobacter sulfurreducens* | Cytochrome c, and cytochrome b |
| 353 | 675 | 0.02 | A0A0M3QFU7 | *Desulfuromonas soudanensis* | Transposase, IS630 family |
| 354 | 675 | 0.02 | Q74D18 | *Geobacter sulfurreducens* | ABC transporter, ATP-binding protein |
| 355 | 673 | 0.02 | A0A0D5NCU1 | *Geobacter sulfurreducens* | Nucleotidyl transferase, PF08843 family protein |
| 356 | 672 | 0.02 | D4H180 | *Denitrovibrio acetiphilus* | Proton-translocating NADH-quinone oxidoreductase, chain L |
| 357 | 672 | 0.02 | Q74E27 | *Geobacter sulfurreducens* | Ribonuclease Y (RNase Y) (EC 3.1.-.-) |
| 358 | 671 | 0.02 | Q39YP6 | *Geobacter metallireducens* | Histidinol-phosphate aminotransferase (EC 2.6.1.9) (Imidazole acetol-phosphate transaminase) |
| 359 | 671 | 0.02 | Q748A7 | *Geobacter sulfurreducens* | Transcription termination factor Rho (EC 3.6.4.-) (ATP-dependent helicase Rho) |
| 360 | 670 | 0.02 | A0A0B5BFC0 | *Geobacter pickeringii* | Ni/Fe hydrogenase |
| 361 | 670 | 0.02 | B9M5J8 | *Geobacter daltonii* | CRISPR-associated protein Csd1 |
| 362 | 669 | 0.02 | A0A0B5BJ50 | *Geobacter pickeringii* | Chemotaxis protein CheA |
| 363 | 669 | 0.02 | | | |
| 364 | 668 | 0.02 | Q74B63 | *Geobacter sulfurreducens* | Uncharacterized protein |
| 365 | 668 | 0.02 | Q74F32 | *Geobacter sulfurreducens* | Periplasmically oriented, membrane-bound formate dehydrogenase, major subunit, selenocysteine-containing |
| 366 | 664 | 0.02 | A0A0D5N7A2 | *Geobacter sulfurreducens* | Chemotaxis protein CheY |
| 367 | 664 | 0.02 | Q39W76 | *Geobacter metallireducens* | Ketol-acid reductoisomerase (NADP(+)) (KARI) (EC 1.1.1.86) (Acetohydroxy-acid isomeroreductase) (AHIR) (Alpha-keto-beta-hydroxylacyl reductoisomerase) (Ketol-acid reductoisomerase type 1) (Ketol-acid reductoisomerase type I) |
| 368 | 661 | 0.02 | A0A0D5N4Y8 | *Geobacter sulfurreducens* | Pilus assembly protein PilB |
| 369 | 660 | 0.02 | A0A1G1FB69 | *Nitrospirae bacterium* | Uncharacterized protein |
| 370 | 659 | 0.02 | A0A0B5BB11 | *Geobacter pickeringii* | Pyruvate carboxylase (EC 6.4.1.1) |
| 371 | 657 | 0.02 | A0A0B5B6N9 | *Geobacter pickeringii* | Alanine--tRNA ligase (EC 6.1.1.7) (Alanyl-tRNA synthetase) (AlaRS) |

| | | | | | |
|---|---|---|---|---|---|
| 372 | 657 | 0.02 | B9LZR8 | *Geobacter daltonii* | Succinyl:acetate coenzyme A transferase |
| 373 | 656 | 0.02 | Q74H59 | *Geobacter sulfurreducens* | Chaperone protein DnaK (HSP70) (Heat shock 70 kDa protein) (Heat shock protein 70) |
| 374 | 655 | 0.02 | A0A0A8WQT9 | *Geobacter sp.* | Dyslexia-associated protein |
| 375 | 655 | 0.02 | P61343 | *Geobacter sulfurreducens* | Aspartyl/glutamyl-tRNA(Asn/Gln) amidotransferase subunit B (Asp/Glu-ADT subunit B) (EC 6.3.5.-) |
| 376 | 653 | 0.02 | Q747D6 | *Geobacter sulfurreducens* | Pyruvate kinase (EC 2.7.1.40) |
| 377 | 651 | 0.02 | Q39ZP3 | *Geobacter metallireducens* | VWFA superfamily protein |
| 378 | 650 | 0.02 | Q39VA8 | *Geobacter metallireducens* | Transcription termination/antitermination protein NusA |
| 379 | 649 | 0.02 | A0A0D5N6S5 | *Geobacter sulfurreducens* | Magnesium transport protein CorA |
| 380 | 649 | 0.02 | P61523 | *Geobacter sulfurreducens* | Argininosuccinate synthase (EC 6.3.4.5) (Citrulline--aspartate ligase) |
| 381 | 649 | 0.02 | D4H844 | *Denitrovibrio acetiphilus* | Carbamoyl-phosphate synthase large chain (EC 6.3.5.5) (Carbamoyl-phosphate synthetase ammonia chain) |
| 382 | 648 | 0.02 | Q39QK1 | *Geobacter metallireducens* | Adenylosuccinate synthetase (AMPSase) (AdSS) (EC 6.3.4.4) (IMP--aspartate ligase) |
| 383 | 644 | 0.02 | A0A0D5N4L7 | *Geobacter sulfurreducens* | Outer membrane protein assembly factor BamA |
| 384 | 644 | 0.02 | I7EP93 | *Geobacter sulfurreducens* | Uncharacterized protein |
| 385 | 644 | 0.02 | Q748B3 | *Geobacter sulfurreducens* | UDP-N-acetylglucosamine 1-carboxyvinyltransferase (EC 2.5.1.7) (Enoylpyruvate transferase) (UDP-N-acetylglucosamine enolpyruvyl transferase) (EPT) |
| 386 | 644 | 0.02 | Q74B69 | *Geobacter sulfurreducens* | Peptidoglycan-binding protein, OmpA family |
| 387 | 644 | 0.02 | Q74D76 | *Geobacter sulfurreducens* | Histidine kinase (EC 2.7.13.3) |
| 388 | 644 | 0.02 | Q74EP9 | *Geobacter sulfurreducens* | DNA-binding ATPase Uup |
| 389 | 641 | 0.02 | A0A0B5B8J3 | *Geobacter pickeringii* | Rod shape-determining protein RodA |
| 390 | 641 | 0.02 | Q39X71 | *Geobacter metallireducens* | Uncharacterized protein |
| 391 | 640 | 0.02 | A5GAC6 | *Geobacter uraniireducens* | FMN-dependent NADH-azoreductase (EC 1.7.-.-) (Azo-dye reductase) (FMN-dependent NADH-azo compound oxidoreductase) |
| 392 | 638 | 0.02 | Q74AT0 | *Geobacter sulfurreducens* | Lysine--tRNA ligase (EC 6.1.1.6) (Lysyl-tRNA synthetase) (LysRS) |
| 393 | 637 | 0.02 | Q74FM8 | *Geobacter sulfurreducens* | Glycine--tRNA ligase alpha subunit (EC 6.1.1.14) (Glycyl-tRNA synthetase alpha subunit) (GlyRS) |
| 394 | 635 | 0.02 | B9M7P3 | *Geobacter daltonii* | UvrABC system protein A (UvrA protein) (Excinuclease ABC subunit A) |
| 395 | 634 | 0.02 | A0A0B5B792 | *Geobacter pickeringii* | Flagellar biosynthesis protein FlhF |
| 396 | 633 | 0.02 | A0A0B5BIA1 | *Geobacter pickeringii* | DNA-directed DNA polymerase (EC 2.7.7.7) |
| 397 | 631 | 0.02 | A0A1G0LEY2 | *Geobacteraceae bacterium* | Uncharacterized protein |
| 398 | 631 | 0.02 | B3EAZ5 | *Geobacter lovleyi* | TraG family protein |
| 399 | 631 | 0.02 | Q74GK2 | *Geobacter sulfurreducens* | Radical SAM domain iron-sulfur cluster-binding oxidoreductase |
| 400 | 630 | 0.02 | Q39RK5 | *Geobacter metallireducens* | ResB-like family cytochrome c |
| 401 | 628 | 0.02 | A0A0A8WMQ9 | *Geobacter sp.* | Uncharacterized protein |
| 402 | 628 | 0.02 | A5GBP1 | *Geobacter uraniireducens* | Uncharacterized protein |
| 403 | 628 | 0.02 | | | |

| | | | | | |
|---|---|---|---|---|---|
| 404 | 627 | 0.02 | A0A1G0M4P0 | *Geobacteraceae bacterium* | Isocitrate dehydrogenase (NADP(+)) |
| 405 | 627 | 0.02 | Q39ZH5 | *Geobacter metallireducens* | Cytochrome c/b |
| 406 | 627 | 0.02 | Q748T9 | *Geobacter sulfurreducens* | Cytochrome c |
| 407 | 626 | 0.02 | A0A0B5BHU5 | *Geobacter pickeringii* | Peptidylprolyl isomerase (EC 5.2.1.8) |
| 408 | 626 | 0.02 | A0A0D5N9G3 | *Geobacter sulfurreducens* | Multidrug ABC transporter substrate-binding protein |
| 409 | 626 | 0.02 | | | |
| 410 | 625 | 0.02 | A0A1G0LUU9 | *Geobacteraceae bacterium* | NADH dehydrogenase |
| 411 | 625 | 0.02 | Q747E4 | *Geobacter sulfurreducens* | Polyphosphate kinase (EC 2.7.4.1) (ATP-polyphosphate phosphotransferase) (Polyphosphoric acid kinase) |
| 412 | 624 | 0.02 | A0A0C1U294 | *Geobacter soli* | Pilus assembly protein PilQ |
| 413 | 623 | 0.02 | A0A0C1TS26 | *Geobacter soli* | L-aspartate oxidase (EC 1.4.3.16) |
| 414 | 623 | 0.02 | Q39UH4 | *Geobacter metallireducens* | Lon protease (EC 3.4.21.53) (ATP-dependent protease La) |
| 415 | 622 | 0.02 | A0A0A8WQH6 | *Geobacter sp.* | Uncharacterized protein |
| 416 | 622 | 0.02 | A0A0B5BJM3 | *Geobacter pickeringii* | NADH-ubiquinone oxidoreductase subunit 3 |
| 417 | 622 | 0.02 | A0A0D5NDV9 | *Geobacter sulfurreducens* | HD family phosphohydrolase |
| 418 | 622 | 0.02 | A0A1F9PTX0 | *Desulfuromonadales bacterium* | Pyruvate-flavodoxin oxidoreductase (EC 1.2.7.-) |
| 419 | 621 | 0.02 | Q74BT1 | *Geobacter sulfurreducens* | Repeat-containing protein |
| 420 | 619 | 0.02 | Q746X5 | *Geobacter sulfurreducens* | Branched-chain amino acid ABC transporter, membrane protein |
| 421 | 616 | 0.02 | | | |
| 422 | 615 | 0.02 | A0A0B5BFH3 | *Geobacter pickeringii* | Polyribonucleotide nucleotidyltransferase (EC 2.7.7.8) (Polynucleotide phosphorylase) (PNPase) |
| 423 | 614 | 0.02 | A0A0A8WT91 | *Geobacter sp.* | Protein-export membrane protein SecF |
| 424 | 614 | 0.02 | A0A0B5BLL1 | *Geobacter pickeringii* | Pyridine nucleotide-disulfide oxidoreductase |
| 425 | 614 | 0.02 | A0A0D5N717 | *Geobacter sulfurreducens* | DNA polymerase III subunit beta (EC 2.7.7.7) |
| 426 | 614 | 0.02 | A5G9T7 | *Geobacter uraniireducens* | Peptide chain release factor 3 (RF-3) |
| 427 | 613 | 0.02 | A0A0D5N6Q2 | *Geobacter sulfurreducens* | Lactamase |
| 428 | 613 | 0.02 | Q74A32 | *Geobacter sulfurreducens* | PppGpp 5'-phosphohydrolase and exopolyphosphatase, HD domain-containing |
| 429 | 612 | 0.02 | A0A0B5BJG9 | *Geobacter pickeringii* | GntR family transcriptional regulator |
| 430 | 612 | 0.02 | A5G8I3 | *Geobacter uraniireducens* | Chaperone protein ClpB |
| 431 | 612 | 0.02 | Q74AF7 | *Geobacter sulfurreducens* | Methyl viologen-reducing hydrogenase, large subunit |
| 432 | 610 | 0.02 | A0A0B5BJ79 | *Geobacter pickeringii* | NADH dehydrogenase |
| 433 | 609 | 0.02 | Q39PS5 | *Geobacter metallireducens* | Quinolinate synthase A (EC 2.5.1.72) |
| 434 | 609 | 0.02 | Q39XX9 | *Geobacter metallireducens* | DNA-directed RNA polymerase subunit alpha (RNAP subunit alpha) (EC 2.7.7.6) (RNA polymerase subunit alpha) (Transcriptase subunit alpha) |
| 435 | 608 | 0.02 | A0A0D5N5H4 | *Geobacter sulfurreducens* | Cell division protein FtsK |
| 436 | 607 | 0.02 | Q748Y7 | *Geobacter sulfurreducens* | 30S ribosomal protein S7 |

| | | | | | |
|---|---|---|---|---|---|
| 437 | 606 | 0.02 | A0A0B5B7V2 | *Geobacter pickeringii* | Ribosome-binding ATPase YchF |
| 438 | 606 | 0.02 | A0A0B5BBV1 | *Geobacter pickeringii* | Cytochrome C |
| 439 | 606 | 0.02 | Q39RQ8 | *Geobacter metallireducens* | Ribose-phosphate pyrophosphokinase (RPPK) (EC 2.7.6.1) (5-phospho-D-ribosyl alpha-1-diphosphate) (Phosphoribosyl diphosphate synthase) (Phosphoribosyl pyrophosphate synthase) (P-Rib-PP synthase) (PRPP synthase) (PRPPase) |
| 440 | 606 | 0.02 | Q74BB7 | *Geobacter sulfurreducens* | Thioredoxin-related domain protein |
| 441 | 605 | 0.02 | A0A0B5BB86 | *Geobacter pickeringii* | Uncharacterized protein |
| 442 | 605 | 0.02 | A0A0D5N3S3 | *Geobacter sulfurreducens* | Chemotaxis protein CheY |
| 443 | 604 | 0.02 | A0A0B5B8W1 | *Geobacter pickeringii* | ATP-dependent zinc metalloprotease FtsH (EC 3.4.24.-) |
| 444 | 604 | 0.02 | A0A0C1TWG6 | *Geobacter soli* | Histidine kinase |
| 445 | 604 | 0.02 | Q748X2 | *Geobacter sulfurreducens* | 3-isopropylmalate dehydrogenase (EC 1.1.1.85) (3-IPM-DH) (Beta-IPM dehydrogenase) (IMDH) |
| 446 | 603 | 0.02 | A0A0B5BJA2 | *Geobacter pickeringii* | RND transporter |
| 447 | 603 | 0.02 | Q39UG5 | *Geobacter metallireducens* | Aspartokinase (EC 2.7.2.4) |
| 448 | 603 | 0.02 | Q74GB8 | *Geobacter sulfurreducens* | Type II secretion system ATPase GspE |
| 449 | 602 | 0.02 | A0A0B5B822 | *Geobacter pickeringii* | Formate dehydrogenase |
| 450 | 601 | 0.02 | A0A0B5BBP3 | *Geobacter pickeringii* | Flagellar basal body stator protein MotB |
| 451 | 599 | 0.02 | A0A0B5BIX5 | *Geobacter pickeringii* | DNA polymerase II |
| 452 | 599 | 0.02 | B5E8R5 | *Geobacter bemidjiensis* | Glutamine synthetase (EC 6.3.1.2) |
| 453 | 597 | 0.02 | A0A0D5N7J7 | *Geobacter sulfurreducens* | Alpha-glucan phosphorylase |
| 454 | 597 | 0.02 | A0A1F9PKR5 | *Desulfuromonadales bacterium* | Uncharacterized protein |
| 455 | 597 | 0.02 | Q39QB1 | *Geobacter metallireducens* | NADH dehydrogenase I, E subunit |
| 456 | 597 | 0.02 | Q39RM2 | *Geobacter metallireducens* | Glycosyltransferase |
| 457 | 596 | 0.02 | A0A0B5B8X9 | *Geobacter pickeringii* | Phosphoenolpyruvate-protein phosphotransferase (EC 2.7.3.9) (Phosphotransferase system, enzyme I) |
| 458 | 596 | 0.02 | Q39WZ1 | *Geobacter metallireducens* | Nitrogen fixation protein NifU |
| 459 | 596 | 0.02 | Q39ZI6 | *Geobacter metallireducens* | DNA helicase (EC 3.6.4.12) |
| 460 | 595 | 0.02 | Q74G55 | *Geobacter sulfurreducens* | Efflux pump, RND family, inner membrane protein |
| 461 | 594 | 0.02 | A0A0B5BD26 | *Geobacter pickeringii* | CDP-diacylglycerol--serine O-phosphatidyltransferase |
| 462 | 594 | 0.02 | Q39X31 | *Geobacter metallireducens* | Protein translocase subunit SecA |
| 463 | 592 | 0.02 | A0A0D5N7G7 | *Geobacter sulfurreducens* | Periplasmic serine endoprotease DegP-like (EC 3.4.21.107) |
| 464 | 592 | 0.02 | D4H7W9 | *Denitrovibrio acetiphilus* | Translation initiation factor IF-2 |
| 465 | 591 | 0.02 | Q748Z0 | *Geobacter sulfurreducens* | 30S ribosomal protein S10 |
| 466 | 590 | 0.02 | A0A0A8WW30 | *Geobacter sp.* | DNA-directed RNA polymerase subunit beta (RNAP subunit beta) (EC 2.7.7.6) (RNA polymerase subunit beta) (Transcriptase subunit beta) |
| 467 | 590 | 0.02 | A0A0D5N2Q7 | *Geobacter sulfurreducens* | DNA-binding protein |

| | | | | | |
|---|---|---|---|---|---|
| 468 | 589 | 0.02 | A0A0B5BBX6 | *Geobacter pickeringii* | Ribonuclease E (RNase E) (EC 3.1.26.12) |
| 469 | 589 | 0.02 | A0A0B5BDV3 | *Geobacter pickeringii* | L-threonine aldolase (EC 4.1.2.48) |
| 470 | 589 | 0.02 | A0A0B5BHJ9 | *Geobacter pickeringii* | Chaperone protein HtpG (Heat shock protein HtpG) (High temperature protein G) |
| 471 | 589 | 0.02 | Q74EQ2 | *Geobacter sulfurreducens* | Aldehyde:ferredoxin oxidoreductase, tungsten-containing |
| 472 | 588 | 0.02 | A0A0B5B9Q9 | *Geobacter pickeringii* | Glycosyl transferase |
| 473 | 587 | 0.02 | A0A0B5BEL7 | *Geobacter pickeringii* | Cytochrome C |
| 474 | 587 | 0.02 | A0A0C1TRJ5 | *Geobacter soli* | Uncharacterized protein |
| 475 | 585 | 0.02 | Q746W5 | *Geobacter sulfurreducens* | Outer membrane channel, OmpJ-related protein |
| 476 | 585 | 0.02 | Q747I8 | *Geobacter sulfurreducens* | Pentapeptide repeat protein |
| 477 | 585 | 0.02 | W6LX92 | *Cand. Contendobacter odensis* | Putative Lipoprotein |
| 478 | 584 | 0.02 | Q39Z54 | *Geobacter metallireducens* | Membrane protein, putative |
| 479 | 583 | 0.02 | A0A0B5BCX9 | *Geobacter pickeringii* | Flagellar motor protein MotA |
| 480 | 583 | 0.02 | B5EE12 | *Geobacter bemidjiensis* | ABC transporter, membrane protein |
| 481 | 582 | 0.02 | Q39XB5 | *Geobacter metallireducens* | 30S ribosomal protein S1 |
| 482 | 581 | 0.02 | A0A1E4LQS0 | *Comamonadaceae bacterium* | Histidine kinase |
| 483 | 580 | 0.02 | A0A0B5BCD2 | *Geobacter pickeringii* | Flagellar biosynthetic protein FliR |
| 484 | 580 | 0.02 | A0LQ19 | *Syntrophobacter fumaroxidans* | Phospholipase D/Transphosphatidylase |
| 485 | 580 | 0.02 | Q39RQ2 | *Geobacter metallireducens* | RNA polymerase sigma factor |
| 486 | 580 | 0.02 | Q39YM5 | *Geobacter metallireducens* | Peptidoglycan transglycosylase and transpeptidase FtsI |
| 487 | 579 | 0.02 | A0A0D5N653 | *Geobacter sulfurreducens* | Histidine kinase |
| 488 | 579 | 0.02 | A0A1G0LYB2 | *Geobacteraceae bacterium* | Uncharacterized protein |
| 489 | 579 | 0.02 | Q39U98 | *Geobacter metallireducens* | Phosphoglycerate kinase (EC 2.7.2.3) |
| 490 | 579 | 0.02 | Q39Y82 | *Geobacter metallireducens* | 5-methyltetrahydrofolate--homocysteine S-methyltransferase, cobalamin-dependent |
| 491 | 578 | 0.02 | A0A0D5N9L3 | *Geobacter sulfurreducens* | Uncharacterized protein |
| 492 | 577 | 0.02 | A0A0B5BHF1 | *Geobacter pickeringii* | Antibiotic ABC transporter ATP-binding protein |
| 493 | 577 | 0.02 | A0A0D5N8L5 | *Geobacter sulfurreducens* | Uncharacterized protein |
| 494 | 577 | 0.02 | A0A1T4RTR1 | *Geobacter thiogenes* | Iron complex outermembrane recepter protein |
| 495 | 576 | 0.02 | A0A0B5BDF3 | *Geobacter pickeringii* | Peptidylprolyl isomerase (EC 5.2.1.8) |
| 496 | 576 | 0.02 | A0A0D5N5F3 | *Geobacter sulfurreducens* | Nitrogenase iron protein (EC 1.18.6.1) (Nitrogenase Fe protein) (Nitrogenase component II) (Nitrogenase reductase) |
| 497 | 576 | 0.02 | Q749D7 | *Geobacter sulfurreducens* | Nitrogenase molybdenum-iron cofactor biosynthesis protein NifEN |
| 498 | 575 | 0.02 | A0A0B5BJS3 | *Geobacter pickeringii* | Integrase |
| 499 | 575 | 0.02 | | | |
| 500 | 574 | 0.02 | A0A0B5B8X8 | *Geobacter pickeringii* | Pilus assembly protein PilB |

**Table S5b**. Functional overview of M2C community (top 500 rank only).

| Rank | Count | Abundance | ID | Organism | Protein Names |
|---|---|---|---|---|---|
| 1 | 3767 | 0.11 | A0A1D3L4G2 | *Methanobacterium congolense* | Uncharacterized protein |
| 2 | 2594 | 0.08 | A7I7H9 | *Methanoregula boonei* | TPR repeat-containing protein |
| 3 | 2359 | 0.07 | A7I4I9 | *Methanoregula boonei* | DNA polymerase II large subunit (Pol II) (EC 2.7.7.7) |
| 4 | 2284 | 0.07 | A0A1D3L5C3 | *Methanobacterium congolense* | Uncharacterized protein |
| 5 | 2163 | 0.07 | A0A090I137 | *Methanobacterium formicicum* | Transposase |
| 6 | 1978 | 0.06 | A7IAM2 | *Methanoregula boonei* | Isoleucine--tRNA ligase (EC 6.1.1.5) (Isoleucyl-tRNA synthetase) (IleRS) |
| 7 | 1951 | 0.06 | A0A1D3L0E4 | *Methanobacterium congolense* | WD40 domain-containing protein |
| 8 | 1882 | 0.06 | | | |
| 9 | 1819 | 0.05 | A0A1D3L0V2 | *Methanobacterium congolense* | Putative ATP-dependent helicase MG140 (EC 3.6.4.-) |
| 10 | 1761 | 0.05 | A0A1D3L429 | *Methanobacterium congolense* | Uncharacterized protein |
| 11 | 1761 | 0.05 | A7IAV3 | *Methanoregula boonei* | V-type ATP synthase subunit I |
| 12 | 1721 | 0.05 | A7I486 | *Methanoregula boonei* | K(+)-insensitive pyrophosphate-energized proton pump (EC 3.6.1.1) (Membrane-bound proton-translocating pyrophosphatase) (Pyrophosphate-energized inorganic pyrophosphatase) (H(+)-PPase) |
| 13 | 1712 | 0.05 | A7IAD8 | *Methanoregula boonei* | Chromosome partition protein Smc |
| 14 | 1692 | 0.05 | A7I9N6 | *Methanoregula boonei* | DNA-directed RNA polymerase subunit (EC 2.7.7.6) |
| 15 | 1668 | 0.05 | A0A1D3KZI4 | *Methanobacterium congolense* | DNA polymerase II large subunit (Pol II) (EC 2.7.7.7) |
| 16 | 1653 | 0.05 | A0A1D3L508 | *Methanobacterium congolense* | WD40 domain-containing protein |
| 17 | 1584 | 0.05 | A0A1D3L5B8 | *Methanobacterium congolense* | Isoleucine--tRNA ligase (EC 6.1.1.5) (Isoleucyl-tRNA synthetase) (IleRS) |
| 18 | 1573 | 0.05 | A7I9M8 | *Methanoregula boonei* | Leucine--tRNA ligase (EC 6.1.1.4) (Leucyl-tRNA synthetase) (LeuRS) |
| 19 | 1559 | 0.05 | A7I4X4 | *Methanoregula boonei* | Elongation factor 2 (EF-2) |
| 20 | 1504 | 0.05 | A7I927 | *Methanoregula boonei* | Valine--tRNA ligase (EC 6.1.1.9) (Valyl-tRNA synthetase) (ValRS) |
| 21 | 1493 | 0.05 | A7I6J1 | *Methanoregula boonei* | Carbamoyl-phosphate synthase large chain (EC 6.3.5.5) (Carbamoyl-phosphate synthetase ammonia chain) |
| 22 | 1449 | 0.04 | A0A1D3KZI9 | *Methanobacterium congolense* | Leucine--tRNA ligase (EC 6.1.1.4) (Leucyl-tRNA synthetase) (LeuRS) |
| 23 | 1379 | 0.04 | A7I485 | *Methanoregula boonei* | Vitamin B12-dependent ribonucleotide reductase (EC 1.17.4.1) |
| 24 | 1369 | 0.04 | P06131 | *Methanobacterium formicicum* | Formate dehydrogenase subunit alpha (EC 1.2.1.2) |
| 25 | 1363 | 0.04 | A7IAB8 | *Methanoregula boonei* | Phosphoenolpyruvate synthase (PEP synthase) (EC 2.7.9.2) (Pyruvate, water dikinase) |
| 26 | 1331 | 0.04 | A0A1D3L027 | *Methanobacterium congolense* | Carbamoyl-phosphate synthase large chain (EC 6.3.5.5) (Carbamoyl-phosphate synthetase ammonia chain) |
| 27 | 1309 | 0.04 | F4BUN6 | *Methanosaeta concilii* | Transposase, IS116/IS110/IS902 family |
| 28 | 1301 | 0.04 | A7I4K7 | *Methanoregula boonei* | Lysine decarboxylase (EC 4.1.1.18) |
| 29 | 1299 | 0.04 | A7I766 | *Methanoregula boonei* | Efflux transporter, putative, hydrophobe/amphiphile efflux-3 (HAE3) family |
| 30 | 1288 | 0.04 | A7I791 | *Methanoregula boonei* | UvrABC system protein A (UvrA protein) (Excinuclease ABC subunit A) |

| | | | | | |
|---|---|---|---|---|---|
| 31 | 1283 | 0.04 | A0A1D3L5A7 | *Methanobacterium congolense* | Uncharacterized protein |
| 32 | 1258 | 0.04 | A7I4R1 | *Methanoregula boonei* | Thermosome |
| 33 | 1258 | 0.04 | A7I5U5 | *Methanoregula boonei* | 4Fe-4S ferredoxin, iron-sulfur binding domain protein |
| 34 | 1252 | 0.04 | A0A1D3L5A8 | *Methanobacterium congolense* | Long-chain-fatty-acid-CoA ligase FadD15 (EC 6.2.1.3) |
| 35 | 1248 | 0.04 | A0A1D3L3Z9 | *Methanobacterium congolense* | Uncharacterized protein |
| 36 | 1235 | 0.04 | A0A1D3KZ93 | *Methanobacterium congolense* | Sodium/potassium-transporting ATPase subunit alpha-2 (EC 3.6.3.9) |
| 37 | 1234 | 0.04 | A7I5I6 | *Methanoregula boonei* | Coenzyme F420 hydrogenase (EC 1.12.98.1) |
| 38 | 1232 | 0.04 | A7I6S1 | *Methanoregula boonei* | CoA-binding domain protein |
| 39 | 1231 | 0.04 | A7I9M7 | *Methanoregula boonei* | PilT protein domain protein |
| 40 | 1223 | 0.04 | A0A1D3L3R4 | *Methanobacterium congolense* | Putative membrane protein MJ1525 |
| 41 | 1217 | 0.04 | A7IB61 | *Methanoregula boonei* | ATP-dependent DNA helicase Hel308 (EC 3.6.4.12) |
| 42 | 1215 | 0.04 | A7I7B5 | *Methanoregula boonei* | DNA polymerase (EC 2.7.7.7) |
| 43 | 1212 | 0.04 | L0HAI4 | *Methanoregula formicica* | Formate dehydrogenase, alpha subunit, archaeal-type |
| 44 | 1194 | 0.04 | A7I5I3 | *Methanoregula boonei* | Coenzyme F420 hydrogenase/dehydrogenase beta subunit domain protein |
| 45 | 1187 | 0.04 | A0A090JUS9 | *Methanobacterium formicicum* | Putative secreted protein |
| 46 | 1184 | 0.04 | A0A1D3L2M2 | *Methanobacterium congolense* | Valine--tRNA ligase (EC 6.1.1.9) (Valyl-tRNA synthetase) (ValRS) |
| 47 | 1182 | 0.04 | A7IAF8 | *Methanoregula boonei* | ABC-1 domain protein |
| 48 | 1179 | 0.04 | A7I4Y9 | *Methanoregula boonei* | DEAD/DEAH box helicase domain protein |
| 49 | 1170 | 0.04 | A7I7S4 | *Methanoregula boonei* | Alanine--tRNA ligase (EC 6.1.1.7) (Alanyl-tRNA synthetase) (AlaRS) |
| 50 | 1157 | 0.03 | A7I6H7 | *Methanoregula boonei* | Methionine--tRNA ligase (EC 6.1.1.10) (Methionyl-tRNA synthetase) (MetRS) |
| 51 | 1153 | 0.03 | A0A1D3L0B4 | *Methanobacterium congolense* | Putative ATP-dependent helicase MJ1401 (EC 3.6.4.-) |
| 52 | 1151 | 0.03 | A0A1D3L4E7 | *Methanobacterium congolense* | Phosphoenolpyruvate synthase (PEP synthase) (EC 2.7.9.2) (Pyruvate, water dikinase) |
| 53 | 1141 | 0.03 | A0A1D3KZH1 | *Methanobacterium congolense* | Copper-exporting P-type ATPase A (EC 3.6.3.54) |
| 54 | 1127 | 0.03 | A7I656 | *Methanoregula boonei* | Elongation factor 1-alpha (EF-1-alpha) (Elongation factor Tu) (EF-Tu) |
| 55 | 1109 | 0.03 | A7I8Y5 | *Methanoregula boonei* | DEAD/DEAH box helicase domain protein |
| 56 | 1102 | 0.03 | A0A1D3L596 | *Methanobacterium congolense* | CoB-CoM heterodisulfide reductase iron-sulfur subunit A (EC 1.8.98.1) |
| 57 | 1095 | 0.03 | A0A1D3L3T8 | *Methanobacterium congolense* | Type IIS restriction enzyme Eco57I (EC 3.1.21.4) |
| 58 | 1092 | 0.03 | A0A1D3L506 | *Methanobacterium congolense* | Carbamoyltransferase (EC 2.1.3.-) |
| 59 | 1087 | 0.03 | A0A1D3L379 | *Methanobacterium congolense* | Calcium-transporting ATPase 1 (EC 3.6.3.8) |
| 60 | 1087 | 0.03 | A7IB37 | *Methanoregula boonei* | ABC transporter related |
| 61 | 1083 | 0.03 | A0A1D3L4H1 | *Methanobacterium congolense* | Acetyl-CoA decarbonylase/synthase complex subunit alpha (ACDS complex subunit alpha) (EC 1.2.7.4) (ACDS complex carbon monoxide dehydrogenase subunit alpha) (ACDS CODH subunit alpha) |
| 62 | 1080 | 0.03 | A7I4W0 | *Methanoregula boonei* | Oligosaccharyl transferase, STT3 subunit |
| 63 | 1079 | 0.03 | A7I8X9 | *Methanoregula boonei* | MCM family protein |
| 64 | 1078 | 0.03 | A0A1D3L276 | *Methanobacterium congolense* | UvrABC system protein A (UvrA protein) (Excinuclease ABC subunit A) |

| | | | | | |
|---|---|---|---|---|---|
| 65 | 1078 | 0.03 | A7IAZ8 | *Methanoregula boonei* | Chaperonin Cpn60/TCP-1 |
| 66 | 1077 | 0.03 | A0A1D3KZY6 | *Methanobacterium congolense* | Elongation factor 2 (EF-2) |
| 67 | 1077 | 0.03 | A0A1D3L355 | *Methanobacterium congolense* | Lon protease (EC 3.4.21.53) (ATP-dependent protease La) |
| 68 | 1074 | 0.03 | A0A1D3L007 | *Methanobacterium congolense* | Alanine--tRNA ligase (EC 6.1.1.7) (Alanyl-tRNA synthetase) (AlaRS) |
| 69 | 1049 | 0.03 | E1RH75 | *Methanolacinia petrolearia* | Putative PAS/PAC sensor protein |
| 70 | 1046 | 0.03 | A0A1D3L118 | *Methanobacterium congolense* | Putative thiamine pyrophosphate-containing protein YdaP |
| 71 | 1040 | 0.03 | A7I6P0 | *Methanoregula boonei* | Threonine--tRNA ligase (EC 6.1.1.3) (Threonyl-tRNA synthetase) (ThrRS) |
| 72 | 1037 | 0.03 | A7I4F8 | *Methanoregula boonei* | Amidophosphoribosyltransferase (ATase) (EC 2.4.2.14) (Glutamine phosphoribosylpyrophosphate amidotransferase) (GPATase) |
| 73 | 1032 | 0.03 | A0A1D3L5M9 | *Methanobacterium congolense* | Putative ATP-dependent RNA helicase MJ1505 (EC 3.6.4.13) |
| 74 | 1029 | 0.03 | A0A1D3L129 | *Methanobacterium congolense* | Putative ATP-dependent helicase MTH_1802 (EC 3.6.4.-) |
| 75 | 1027 | 0.03 | A0A1D3L0D3 | *Methanobacterium congolense* | Calcium-transporting ATPase 1 (EC 3.6.3.8) |
| 76 | 1018 | 0.03 | A0A1D3L0D2 | *Methanobacterium congolense* | Copper-exporting P-type ATPase B (EC 3.6.3.54) |
| 77 | 1013 | 0.03 | A0A1D3KZ84 | *Methanobacterium congolense* | Uncharacterized protein |
| 78 | 1013 | 0.03 | A0A1D3L3L7 | *Methanobacterium congolense* | Metabotropic glutamate receptor-like protein R |
| 79 | 1013 | 0.03 | A0A1D3L4X9 | *Methanobacterium congolense* | Putative transcriptional regulatory protein pdtaR |
| 80 | 1013 | 0.03 | A7IB30 | *Methanoregula boonei* | AAA family ATPase, CDC48 subfamily (EC 3.6.4.6) |
| 81 | 1010 | 0.03 | A7I9Q1 | *Methanoregula boonei* | Glutamine synthetase (EC 6.3.1.2) |
| 82 | 1008 | 0.03 | A7I5R2 | *Methanoregula boonei* | Protein translocase subunit SecY (Protein transport protein SEC61 subunit alpha homolog) |
| 83 | 1002 | 0.03 | A0A1D3L4J7 | *Methanobacterium congolense* | Transglutaminase domain-containing protein |
| 84 | 1002 | 0.03 | L0HCK9 | *Methanoregula formicica* | Formylmethanofuran dehydrogenase subunit A |
| 85 | 989 | 0.03 | A7I6L6 | *Methanoregula boonei* | DNA topoisomerase 1 (EC 5.99.1.2) (DNA topoisomerase I) |
| 86 | 981 | 0.03 | A7IAP7 | *Methanoregula boonei* | Probable translation initiation factor IF-2 |
| 87 | 981 | 0.03 | A7IA93 | *Methanoregula boonei* | CTP synthase (EC 6.3.4.2) (Cytidine 5'-triphosphate synthase) (Cytidine triphosphate synthetase) (CTP synthetase) (CTPS) (UTP--ammonia ligase) |
| 88 | 978 | 0.03 | A7I6T1 | *Methanoregula boonei* | Coenzyme F420 hydrogenase/dehydrogenase beta subunit domain protein |
| 89 | 976 | 0.03 | A0A1D3L0Y3 | *Methanobacterium congolense* | Cell division cycle protein 48 homolog MJ1156 |
| 90 | 964 | 0.03 | A7IAU7 | *Methanoregula boonei* | V-type ATP synthase beta chain (V-ATPase subunit B) |
| 91 | 963 | 0.03 | A0A1D3L579 | *Methanobacterium congolense* | Phosphoribosylformylglycinamidine synthase subunit PurL (FGAM synthase) (EC 6.3.5.3) (Formylglycinamide ribonucleotide amidotransferase subunit II) (FGAR amidotransferase II) (FGAR-AT II) (Glutamine amidotransferase PurL) (Phosphoribosylformylglycinamidine synthase subunit II) |
| 92 | 963 | 0.03 | A0A1D3L5J6 | *Methanobacterium congolense* | Acetolactate synthase (EC 2.2.1.6) |
| 93 | 963 | 0.03 | A7I6H1 | *Methanoregula boonei* | Glycyl-tRNA synthetase |
| 94 | 961 | 0.03 | A7I9N8 | *Methanoregula boonei* | DNA-directed RNA polymerase (EC 2.7.7.6) |
| 95 | 961 | 0.03 | L0HG76 | *Methanoregula formicica* | CRISPR-associated helicase Cas3/CRISPR-associated endonuclease Cas3-HD |

| | | | | | |
|---|---|---|---|---|---|
| 96 | 959 | 0.03 | A0A1D3L2A8 | *Methanobacterium congolense* | Hydroxylamine reductase (EC 1.7.99.1) (Hybrid-cluster protein) (HCP) (Prismane protein) |
| 97 | 957 | 0.03 | A7I4V8 | *Methanoregula boonei* | Translation initiation factor 2 subunit gamma (aIF2-gamma) (eIF-2-gamma) |
| 98 | 955 | 0.03 | A7I7A6 | *Methanoregula boonei* | Methyl-accepting chemotaxis sensory transducer |
| 99 | 954 | 0.03 | A0A1D3L584 | *Methanobacterium congolense* | Uncharacterized protein |
| 100 | 953 | 0.03 | A7I5N0 | *Methanoregula boonei* | AMP phosphorylase (AMPpase) (EC 2.4.2.57) (Nucleoside monophosphate phosphorylase) (NMP phosphorylase) |
| 101 | 949 | 0.03 | A7I6S2 | *Methanoregula boonei* | Acetate--CoA ligase |
| 102 | 947 | 0.03 | A7I4G8 | *Methanoregula boonei* | Aspartate--tRNA(Asp/Asn) ligase (EC 6.1.1.23) (Aspartyl-tRNA synthetase) (AspRS) (Non-discriminating aspartyl-tRNA synthetase) (ND-AspRS) |
| 103 | 938 | 0.03 | A7IAZ4 | *Methanoregula boonei* | Peptidase S16, Lon-like protease (EC 3.4.21.53) |
| 104 | 937 | 0.03 | A0A1D3L0F2 | *Methanobacterium congolense* | Archaeal Lon protease (EC 3.4.21.-) |
| 105 | 933 | 0.03 | A7I9N7 | *Methanoregula boonei* | DNA-directed RNA polymerase subunit beta (EC 2.7.7.6) |
| 106 | 929 | 0.03 | A7I8P2 | *Methanoregula boonei* | Phosphoesterase, RecJ domain protein |
| 107 | 924 | 0.03 | A7I798 | *Methanoregula boonei* | 60 kDa chaperonin (GroEL protein) (Protein Cpn60) |
| 108 | 921 | 0.03 | A0A1D3L5E6 | *Methanobacterium congolense* | Replication factor A |
| 109 | 919 | 0.03 | A7I7Z3 | *Methanoregula boonei* | CheA signal transduction histidine kinase |
| 110 | 913 | 0.03 | A7I5S6 | *Methanoregula boonei* | Tetrahydromethanopterin S-methyltransferase subunit A (EC 2.1.1.86) (N5-methyltetrahydromethanopterin--coenzyme M methyltransferase subunit A) |
| 111 | 911 | 0.03 | A7IAJ6 | *Methanoregula boonei* | DEAD/DEAH box helicase domain protein |
| 112 | 906 | 0.03 | A0A1D3L5K7 | *Methanobacterium congolense* | Threonine--tRNA ligase (EC 6.1.1.3) (Threonyl-tRNA synthetase) (ThrRS) |
| 113 | 903 | 0.03 | A7IAU5 | *Methanoregula boonei* | Phosphoglycerate kinase (EC 2.7.2.3) |
| 114 | 901 | 0.03 | A0A1D3L1G9 | *Methanobacterium congolense* | DNA helicase II related protein |
| 115 | 901 | 0.03 | A0A1D3L4A4 | *Methanobacterium congolense* | Putative protease YrrO (EC 3.4.-.-) |
| 116 | 899 | 0.03 | A0A1D3L1K6 | *Methanobacterium congolense* | Putative ATP-dependent helicase YprA (EC 3.6.4.-) |
| 117 | 899 | 0.03 | A7I7Z8 | *Methanoregula boonei* | Type II secretion system protein |
| 118 | 898 | 0.03 | A7IAG9 | *Methanoregula boonei* | Enolase (EC 4.2.1.11) (2-phospho-D-glycerate hydro-lyase) (2-phosphoglycerate dehydratase) |
| 119 | 894 | 0.03 | A0A1D3L4X3 | *Methanobacterium congolense* | Archaeal glutamate synthase [NADPH] (EC 1.4.1.13) |
| 120 | 891 | 0.03 | A0A1D3L4D8 | *Methanobacterium congolense* | Pyruvate carboxylase subunit B (EC 6.4.1.1) |
| 121 | 891 | 0.03 | A7I7B3 | *Methanoregula boonei* | UvrABC system protein B (Protein UvrB) (Excinuclease ABC subunit B) |
| 122 | 891 | 0.03 | A7I8Y8 | *Methanoregula boonei* | AMP-dependent synthetase and ligase |
| 123 | 889 | 0.03 | A0A1D3L598 | *Methanobacterium congolense* | Chaperone protein DnaK (HSP70) (Heat shock 70 kDa protein) (Heat shock protein 70) |
| 124 | 887 | 0.03 | E1RD00 | *Methanolacinia petrolearia* | Pyrrolo-quinoline quinone |
| 125 | 884 | 0.03 | A7I464 | *Methanoregula boonei* | ORC1-type DNA replication protein |
| 126 | 883 | 0.03 | A0A1D3L430 | *Methanobacterium congolense* | Cell division cycle protein 48 homolog MJ1156 |
| 127 | 882 | 0.03 | A7I8Z2 | *Methanoregula boonei* | Ammonium transporter |
| 128 | 881 | 0.03 | A7I4Y2 | *Methanoregula boonei* | Arginine--tRNA ligase (EC 6.1.1.19) (Arginyl-tRNA synthetase) (ArgRS) |

| | | | | | |
|---|---|---|---|---|---|
| 129 | 880 | 0.03 | A0A1D3L4R2 | *Methanobacterium congolense* | Cyanobacterial phytochrome A (EC 2.7.13.3) |
| 130 | 880 | 0.03 | A0A1D3L5X1 | *Methanobacterium congolense* | Dihydroxy-acid dehydratase (DAD) (EC 4.2.1.9) |
| 131 | 874 | 0.03 | A7I8Q2 | *Methanoregula boonei* | AAA family ATPase, CDC48 subfamily (EC 3.6.4.6) |
| 132 | 869 | 0.03 | A7I8B1 | *Methanoregula boonei* | Polyphosphate kinase (EC 2.7.4.1) (ATP-polyphosphate phosphotransferase) (Polyphosphoric acid kinase) |
| 133 | 866 | 0.03 | L0HHJ9 | *Methanoregula formicica* | DNA gyrase subunit A (EC 5.99.1.3) |
| 134 | 864 | 0.03 | A7I988 | *Methanoregula boonei* | Acetyl-CoA carboxylase, biotin carboxylase |
| 135 | 864 | 0.03 | E1RH76 | *Methanolacinia petrolearia* | F420-dependent oxidoreductase, G6PDH family |
| 136 | 863 | 0.03 | A0A1D3L1R0 | *Methanobacterium congolense* | Acetate-CoA ligase [ADP-forming] I (EC 6.2.1.13) |
| 137 | 860 | 0.03 | A0A1D3L170 | *Methanobacterium congolense* | ATP-dependent DNA helicase Hel308 (EC 3.6.4.12) |
| 138 | 858 | 0.03 | A7I6E6 | *Methanoregula boonei* | Heavy metal translocating P-type ATPase |
| 139 | 857 | 0.03 | A0A1D3L3V9 | *Methanobacterium congolense* | Putative KH and PIN-domain containing protein MJ1533 |
| 140 | 854 | 0.03 | A0A1D3L407 | *Methanobacterium congolense* | Putative actinorhodin transporter |
| 141 | 851 | 0.03 | A7IAE0 | *Methanoregula boonei* | Serine hydroxymethyltransferase (SHMT) (Serine methylase) (EC 2.1.2.1) |
| 142 | 851 | 0.03 | A7IAJ9 | *Methanoregula boonei* | Carbamoyltransferase (EC 2.1.3.-) |
| 143 | 850 | 0.03 | A0A1D3KZ36 | *Methanobacterium congolense* | O-phosphoserine--tRNA(Cys) ligase (O-phosphoserine--tRNA ligase) (EC 6.1.1.27) (Non-canonical O-phosphoseryl-tRNA(Cys) synthetase) (O-phosphoseryl-tRNA(Cys) synthetase) (SepRS) |
| 144 | 849 | 0.03 | A7I4C9 | *Methanoregula boonei* | Acetate--CoA ligase |
| 145 | 849 | 0.03 | A7I6K0 | *Methanoregula boonei* | Radical SAM domain protein |
| 146 | 846 | 0.03 | A0A1D3L3N4 | *Methanobacterium congolense* | Acetyl-coenzyme A synthetase (EC 6.2.1.1) |
| 147 | 843 | 0.03 | A7IAY8 | *Methanoregula boonei* | Type II secretion system protein |
| 148 | 843 | 0.03 | F4BZD0 | *Methanosaeta concilii* | Transposase, putative |
| 149 | 840 | 0.03 | U6E9M3 | *Methanobacterium sp.* | DNA-directed RNA polymerase subunit (EC 2.7.7.6) |
| 150 | 839 | 0.03 | A0A101HE03 | *Methanobacterium sp.* | WD40 domain protein beta Propeller |
| 151 | 836 | 0.03 | A0A090I3J2 | *Methanobacterium formicicum* | Uncharacterized protein |
| 152 | 836 | 0.03 | A7IAM8 | *Methanoregula boonei* | TrkA-N domain protein |
| 153 | 836 | 0.03 | E1RF47 | *Methanolacinia petrolearia* | Ammonium transporter |
| 154 | 835 | 0.03 | A7IAB7 | *Methanoregula boonei* | D-3-phosphoglycerate dehydrogenase (EC 1.1.1.95) |
| 155 | 834 | 0.03 | A7I4R9 | *Methanoregula boonei* | Sodium/hydrogen exchanger |
| 156 | 833 | 0.03 | A7IA07 | *Methanoregula boonei* | Aminotransferase, class V |
| 157 | 832 | 0.03 | A0A1D3L551 | *Methanobacterium congolense* | Uncharacterized protein |
| 158 | 830 | 0.03 | A0A1D3L132 | *Methanobacterium congolense* | Putative pseudomurein-binding protein |
| 159 | 830 | 0.03 | A7IAT4 | *Methanoregula boonei* | Lysine--tRNA ligase (EC 6.1.1.6) (Lysyl-tRNA synthetase) (LysRS) |
| 160 | 829 | 0.03 | A7I842 | *Methanoregula boonei* | Beta-lactamase domain protein |
| 161 | 825 | 0.02 | A0A1D3L3Z6 | *Methanobacterium congolense* | DNA replication licensing factor MCM7 (EC 3.6.4.12) |
| 162 | 823 | 0.02 | A0A1D3L4H2 | *Methanobacterium congolense* | Inner membrane protein YccS |

| | | | | | |
|---|---|---|---|---|---|
| 163 | 820 | 0.02 | A7I696 | *Methanoregula boonei* | Argininosuccinate lyase (ASAL) (EC 4.3.2.1) (Arginosuccinase) |
| 164 | 819 | 0.02 | A0A089ZCJ0 | *Methanobacterium formicicum* | Cobalamin biosynthesis protein N (EC 6.6.1.2) (Cobaltochelatase CobN2) |
| 165 | 815 | 0.02 | A7I4I5 | *Methanoregula boonei* | UbiD family decarboxylase |
| 166 | 815 | 0.02 | A7I941 | *Methanoregula boonei* | Type 2 DNA topoisomerase 6 subunit B (EC 5.99.1.3) (Type II DNA topoisomerase VI subunit B) (TopoVI-B) |
| 167 | 814 | 0.02 | A7I498 | *Methanoregula boonei* | DNA polymerase II small subunit (Pol II) (EC 2.7.7.7) |
| 168 | 813 | 0.02 | A0A1D3KZK0 | *Methanobacterium congolense* | Glutamyl-tRNA(Gln) amidotransferase subunit E (Glu-ADT subunit E) (EC 6.3.5.-) |
| 169 | 812 | 0.02 | A0A1D3KZE7 | *Methanobacterium congolense* | Putative sensor histidine kinase pdtaS (EC 2.7.13.3) |
| 170 | 812 | 0.02 | A7I487 | *Methanoregula boonei* | NADH dehydrogenase (Quinone) (EC 1.6.99.5) |
| 171 | 810 | 0.02 | A0A1D3L3Q6 | *Methanobacterium congolense* | Pyridoxal 5'-phosphate synthase subunit PdxS (PLP synthase subunit PdxS) (EC 4.3.3.6) (Pdx1) |
| 172 | 806 | 0.02 | A0A1D3L1W6 | *Methanobacterium congolense* | Methionine--tRNA ligase (EC 6.1.1.10) (Methionyl-tRNA synthetase) (MetRS) |
| 173 | 806 | 0.02 | A7IAF0 | *Methanoregula boonei* | Major facilitator superfamily MFS_1 |
| 174 | 806 | 0.02 | A7IAV0 | *Methanoregula boonei* | V-type ATP synthase subunit C (V-ATPase subunit C) |
| 175 | 805 | 0.02 | A0A1D3KZX3 | *Methanobacterium congolense* | DNA-directed RNA polymerase subunit beta (EC 2.7.7.6) |
| 176 | 803 | 0.02 | A0A1D3L3E8 | *Methanobacterium congolense* | Indolepyruvate oxidoreductase subunit IorA (IOR) (EC 1.2.7.8) (Indolepyruvate ferredoxin oxidoreductase subunit alpha) |
| 177 | 803 | 0.02 | A0A1D3L5R3 | *Methanobacterium congolense* | Probable bifunctional tRNA threonylcarbamoyladenosine biosynthesis protein [Includes: Serine/threonine-protein kinase Bud32 (EC 2.7.11.1); tRNA N6-adenosine threonylcarbamoyltransferase (EC 2.3.1.234) (tRNA threonylcarbamoyladenosine biosynthesis protein Kae1) (t(6)A37 threonylcarbamoyladenosine biosynthesis protein Kae1)] |
| 178 | 799 | 0.02 | A0A1D3KZ40 | *Methanobacterium congolense* | Fumarate reductase (CoM/CoB) subunit A |
| 179 | 798 | 0.02 | A0A1D3L5Q7 | *Methanobacterium congolense* | PAS domain S-box |
| 180 | 798 | 0.02 | A7I7Z9 | *Methanoregula boonei* | Type II secretion system protein E |
| 181 | 797 | 0.02 | A0A1D3KZQ9 | *Methanobacterium congolense* | Putative (R)-citramalate synthase CimA (EC 2.3.1.182) |
| 182 | 795 | 0.02 | A0A1D3L4I5 | *Methanobacterium congolense* | Methyl-coenzyme M reductase I subunit beta (EC 2.8.4.1) |
| 183 | 794 | 0.02 | A0A1D3L203 | *Methanobacterium congolense* | Uncharacterized protein |
| 184 | 791 | 0.02 | A7I6C7 | *Methanoregula boonei* | AMP-dependent synthetase and ligase |
| 185 | 790 | 0.02 | A0A1D3L2D6 | *Methanobacterium congolense* | Putative cadmium-transporting ATPase (EC 3.6.3.3) |
| 186 | 790 | 0.02 | A0A1D3L2H5 | *Methanobacterium congolense* | tRNA-guanine(15) transglycosylase (EC 2.4.2.48) (7-cyano-7-deazaguanine tRNA-ribosyltransferase) (Archaeal tRNA-guanine transglycosylase) |
| 187 | 790 | 0.02 | A7I7L4 | *Methanoregula boonei* | Chaperone protein DnaK (HSP70) (Heat shock 70 kDa protein) (Heat shock protein 70) |
| 188 | 787 | 0.02 | A7I989 | *Methanoregula boonei* | Oxaloacetate decarboxylase alpha subunit |
| 189 | 785 | 0.02 | A7IAG1 | *Methanoregula boonei* | Glutamate--tRNA ligase (EC 6.1.1.17) (Glutamyl-tRNA synthetase) (GluRS) |
| 190 | 784 | 0.02 | A0A1D3L5S7 | *Methanobacterium congolense* | Uncharacterized protein |
| 191 | 783 | 0.02 | A7I931 | *Methanoregula boonei* | GTP-binding protein, HSR1-related |
| 192 | 779 | 0.02 | A0A1D3L1R3 | *Methanobacterium congolense* | Putative sensor histidine kinase pdtaS (EC 2.7.13.3) |

| | | | | | |
|---|---|---|---|---|---|
| 193 | 775 | 0.02 | A0A1D3L2V9 | *Methanobacterium congolense* | Uncharacterized protein |
| 194 | 775 | 0.02 | A7I9L7 | *Methanoregula boonei* | 4-vinyl reductase, 4VR |
| 195 | 774 | 0.02 | A7I5U3 | *Methanoregula boonei* | Methyl-coenzyme M reductase, beta subunit |
| 196 | 773 | 0.02 | A0A1D3L5B6 | *Methanobacterium congolense* | Helicase c2 |
| 197 | 773 | 0.02 | A7I4F0 | *Methanoregula boonei* | Aspartokinase (EC 2.7.2.4) |
| 198 | 773 | 0.02 | A7I9N5 | *Methanoregula boonei* | DNA-directed RNA polymerase subunit A'' (EC 2.7.7.6) |
| 199 | 771 | 0.02 | A0A1D3L452 | *Methanobacterium congolense* | DNA topoisomerase 1 (EC 5.99.1.2) (DNA topoisomerase I) |
| 200 | 769 | 0.02 | A0A1D3L412 | *Methanobacterium congolense* | Putative MFS-type transporter YusP |
| 201 | 768 | 0.02 | A0A090I3X0 | *Methanobacterium formicicum* | Nitrogenase protein alpha chain (EC 1.18.6.1) |
| 202 | 768 | 0.02 | A0A1D3KZR8 | *Methanobacterium congolense* | DNA ligase (EC 6.5.1.1) (Polydeoxyribonucleotide synthase [ATP]) |
| 203 | 765 | 0.02 | A0A1D3L5K9 | *Methanobacterium congolense* | Uncharacterized protein |
| 204 | 764 | 0.02 | A0A1D3L0A6 | *Methanobacterium congolense* | Putative acyl-CoA synthetase YngI (EC 6.2.1.-) |
| 205 | 763 | 0.02 | A0A1D3L4R1 | *Methanobacterium congolense* | V-type ATP synthase alpha chain (EC 3.6.3.14) (V-ATPase subunit A) |
| 206 | 762 | 0.02 | A7I822 | *Methanoregula boonei* | K+ potassium transporter |
| 207 | 762 | 0.02 | A7I915 | *Methanoregula boonei* | Anaerobic ribonucleoside-triphosphate reductase |
| 208 | 758 | 0.02 | A7I648 | *Methanoregula boonei* | Uncharacterized protein |
| 209 | 757 | 0.02 | A0A1D3L223 | *Methanobacterium congolense* | CoB-CoM heterodisulfide reductase 1 iron-sulfur subunit A (EC 1.8.98.1) |
| 210 | 756 | 0.02 | A7I4W3 | *Methanoregula boonei* | Glutamine--fructose-6-phosphate aminotransferase [isomerizing] (EC 2.6.1.16) (D-fructose-6-phosphate amidotransferase) (GFAT) (Glucosamine-6-phosphate synthase) (Hexosephosphate aminotransferase) (L-glutamine--D-fructose-6-phosphate amidotransferase) |
| 211 | 755 | 0.02 | A7I9C0 | *Methanoregula boonei* | Pyruvate carboxyltransferase |
| 212 | 752 | 0.02 | A0A1D3L4A8 | *Methanobacterium congolense* | Uncharacterized protein |
| 213 | 752 | 0.02 | F6D827 | *Methanobacterium paludis* | Formylmethanofuran dehydrogenase subunit B (EC 1.2.99.5) |
| 214 | 751 | 0.02 | A7I4K8 | *Methanoregula boonei* | Amino acid permease-associated region |
| 215 | 750 | 0.02 | A0A1D3L1A5 | *Methanobacterium congolense* | Acetyl-coenzyme A synthetase (EC 6.2.1.1) |
| 216 | 750 | 0.02 | A0A1D3L4J0 | *Methanobacterium congolense* | Phage infection protein |
| 217 | 749 | 0.02 | A0A1D3L2M9 | *Methanobacterium congolense* | Phenylalanine--tRNA ligase beta subunit (EC 6.1.1.20) (Phenylalanyl-tRNA synthetase beta subunit) (PheRS) |
| 218 | 749 | 0.02 | A0A1D3L5K4 | *Methanobacterium congolense* | Uncharacterized protein |
| 219 | 746 | 0.02 | A0A1D3L0R9 | *Methanobacterium congolense* | Catalase-peroxidase (CP) (EC 1.11.1.21) (Peroxidase/catalase) |
| 220 | 746 | 0.02 | A0A1D3L369 | *Methanobacterium congolense* | Transposase |
| 221 | 746 | 0.02 | F4BY05 | *Methanosaeta concilii* | Transposase, IS116/IS110/IS902 family |
| 222 | 744 | 0.02 | A7I4Y1 | *Methanoregula boonei* | Peptide chain release factor subunit 1 (Translation termination factor aRF1) |
| 223 | 743 | 0.02 | A7I617 | *Methanoregula boonei* | Adenosylhomocysteinase (EC 3.3.1.1) |
| 224 | 742 | 0.02 | A7IB41 | *Methanoregula boonei* | Putative CoA-substrate-specific enzyme activase |
| 225 | 741 | 0.02 | K2R0T8 | *Methanobacterium formicicum* | DNA-directed RNA polymerase subunit (EC 2.7.7.6) |

| | | | | | |
|---|---|---|---|---|---|
| 226 | 740 | 0.02 | A0A1D3KZW0 | *Methanobacterium congolense* | Putative ABC transporter ATP-binding protein MJ1242 |
| 227 | 740 | 0.02 | A0A1D3L5S0 | *Methanobacterium congolense* | Arginine--tRNA ligase (EC 6.1.1.19) (Arginyl-tRNA synthetase) (ArgRS) |
| 228 | 740 | 0.02 | A7I667 | *Methanoregula boonei* | Phenylacetate--CoA ligase |
| 229 | 739 | 0.02 | A7I4B9 | *Methanoregula boonei* | Aminotransferase (EC 2.6.1.-) |
| 230 | 738 | 0.02 | A0A1D3L153 | *Methanobacterium congolense* | Uncharacterized protein |
| 231 | 738 | 0.02 | A7I7Y9 | *Methanoregula boonei* | ABC transporter related |
| 232 | 737 | 0.02 | L0HFP5 | *Methanoregula formicica* | Magnesium chelatase, H subunit |
| 233 | 736 | 0.02 | U6EA67 | *Methanobacterium sp.* | Uncharacterized protein |
| 234 | 735 | 0.02 | A7I5L0 | *Methanoregula boonei* | Indolepyruvate oxidoreductase subunit IorA (IOR) (EC 1.2.7.8) (Indolepyruvate ferredoxin oxidoreductase subunit alpha) |
| 235 | 735 | 0.02 | A7I859 | *Methanoregula boonei* | Uncharacterized protein |
| 236 | 734 | 0.02 | A0A1D3L2F4 | *Methanobacterium congolense* | Inosine-5'-monophosphate dehydrogenase (IMP dehydrogenase) (IMPD) (IMPDH) (EC 1.1.1.205) |
| 237 | 733 | 0.02 | A0A1D3L103 | *Methanobacterium congolense* | Ferrous iron transport protein B |
| 238 | 730 | 0.02 | A7I5H9 | *Methanoregula boonei* | Phosphoribosylformylglycinamidine synthase subunit PurL (FGAM synthase) (EC 6.3.5.3) (Formylglycinamide ribonucleotide amidotransferase subunit II) (FGAR amidotransferase II) (FGAR-AT II) (Glutamine amidotransferase PurL) (Phosphoribosylformylglycinamidine synthase subunit II) |
| 239 | 728 | 0.02 | A0A1D3L094 | *Methanobacterium congolense* | Magnesium-chelatase subunit ChlI homolog |
| 240 | 728 | 0.02 | A0A1D3L3V1 | *Methanobacterium congolense* | Cyanobacterial phytochrome A (EC 2.7.13.3) |
| 241 | 727 | 0.02 | A0A1D3L0U2 | *Methanobacterium congolense* | Archaeal glutamate synthase [NADPH] (EC 1.4.1.13) |
| 242 | 726 | 0.02 | A0A1D3L4N1 | *Methanobacterium congolense* | Type-2 serine--tRNA ligase (EC 6.1.1.11) (Seryl-tRNA synthetase) (SerRS) (Seryl-tRNA(Ser/Sec) synthetase) |
| 243 | 725 | 0.02 | A0A1D3L0Z3 | *Methanobacterium congolense* | Thermosome subunit beta |
| 244 | 725 | 0.02 | A0A1D3L2G6 | *Methanobacterium congolense* | Methyl sulfide methyltransferase-associated sensor (EC 2.7.13.3) |
| 245 | 724 | 0.02 | A7I666 | *Methanoregula boonei* | Phenylacetate--CoA ligase (EC 6.2.1.30) |
| 246 | 723 | 0.02 | A0A1D3L504 | *Methanobacterium congolense* | DNA polymerase (EC 2.7.7.7) |
| 247 | 722 | 0.02 | A0A1D3KZW8 | *Methanobacterium congolense* | Uncharacterized protein |
| 248 | 722 | 0.02 | A0A1D3L0K8 | *Methanobacterium congolense* | Glutamine--fructose-6-phosphate aminotransferase [isomerizing] (EC 2.6.1.16) (D-fructose-6-phosphate amidotransferase) (GFAT) (Glucosamine-6-phosphate synthase) (Hexosephosphate aminotransferase) (L-glutamine--D-fructose-6-phosphate amidotransferase) |
| 249 | 719 | 0.02 | A0A1D3L3Z3 | *Methanobacterium congolense* | Putative ABC transporter ATP-binding protein MJ0719 |
| 250 | 718 | 0.02 | A0A1D3L2V1 | *Methanobacterium congolense* | Uncharacterized protein |
| 251 | 717 | 0.02 | A7I8Y7 | *Methanoregula boonei* | AMP-dependent synthetase and ligase |
| 252 | 716 | 0.02 | A0A1D3KZT8 | *Methanobacterium congolense* | Acyl-coenzyme A synthetase ACSM3, mitochondrial (EC 6.2.1.2) |
| 253 | 716 | 0.02 | A0A1D3L1D7 | *Methanobacterium congolense* | Uncharacterized protein |
| 254 | 714 | 0.02 | A0A1D3L1V6 | *Methanobacterium congolense* | Uncharacterized protein |

| | | | | | |
|---|---|---|---|---|---|
| 255 | 714 | 0.02 | A7I4G3 | *Methanoregula boonei* | AMP-dependent synthetase and ligase |
| 256 | 713 | 0.02 | A7I8Z7 | *Methanoregula boonei* | Archaeal glutamate synthase [NADPH] (EC 1.4.1.13) |
| 257 | 712 | 0.02 | A0A1D3L3S1 | *Methanobacterium congolense* | Archaeal glutamate synthase [NADPH] (EC 1.4.1.13) |
| 258 | 712 | 0.02 | L0HEU5 | *Methanoregula formicica* | DNA gyrase subunit B (EC 5.99.1.3) |
| 259 | 711 | 0.02 | A0A1D3L117 | *Methanobacterium congolense* | Uncharacterized protein |
| 260 | 710 | 0.02 | A7I5J3 | *Methanoregula boonei* | AAA ATPase, central domain protein |
| 261 | 709 | 0.02 | A0A1D3L5G6 | *Methanobacterium congolense* | DNA polymerase II small subunit (Pol II) (EC 2.7.7.7) |
| 262 | 709 | 0.02 | A7I6C9 | *Methanoregula boonei* | Bifunctional enzyme Fae/Hps [Includes: 5,6,7,8-tetrahydromethanopterin hydro-lyase (EC 4.2.1.147) (Formaldehyde-activating enzyme) (Fae); 3-hexulose-6-phosphate synthase (HPS) (EC 4.1.2.43) (D-arabino-3-hexulose-6-phosphate formaldehyde lyase)] |
| 263 | 709 | 0.02 | A7I5I7 | *Methanoregula boonei* | GTP cyclohydrolase MptA (EC 3.5.4.39) (GTP cyclohydrolase IV) |
| 264 | 708 | 0.02 | A0A1D3L230 | *Methanobacterium congolense* | UvrABC system protein B (Protein UvrB) (Excinuclease ABC subunit B) |
| 265 | 708 | 0.02 | A7I6J4 | *Methanoregula boonei* | Uncharacterized protein |
| 266 | 707 | 0.02 | A0A1D3L4H3 | *Methanobacterium congolense* | V-type ATP synthase subunit I |
| 267 | 707 | 0.02 | A0A1D3L537 | *Methanobacterium congolense* | Uncharacterized protein |
| 268 | 707 | 0.02 | A7I4B6 | *Methanoregula boonei* | Aminotransferase, class V |
| 269 | 707 | 0.02 | A7I4R6 | *Methanoregula boonei* | Pyruvate flavodoxin/ferredoxin oxidoreductase domain protein |
| 270 | 705 | 0.02 | L0HD61 | *Methanoregula formicica* | Coenzyme F420-reducing hydrogenase, beta subunit |
| 271 | 704 | 0.02 | A7I922 | *Methanoregula boonei* | Inosine-5'-monophosphate dehydrogenase (IMP dehydrogenase) (IMPD) (IMPDH) (EC 1.1.1.205) |
| 272 | 702 | 0.02 | A0A1D3KZ91 | *Methanobacterium congolense* | Putative glutamine amidotransferase-like protein MJ0138 |
| 273 | 701 | 0.02 | A7I7B2 | *Methanoregula boonei* | Ribulose bisphosphate carboxylase (RuBisCO) (EC 4.1.1.39) |
| 274 | 700 | 0.02 | A0A1D3L514 | *Methanobacterium congolense* | GTP-binding protein 1 |
| 275 | 699 | 0.02 | A0A1D3L3P1 | *Methanobacterium congolense* | Probable translation initiation factor IF-2 |
| 276 | 698 | 0.02 | A7I9L0 | *Methanoregula boonei* | Type II secretion system protein E |
| 277 | 696 | 0.02 | A0A1D3L078 | *Methanobacterium congolense* | DEAD-box ATP-dependent RNA helicase CshA (EC 3.6.4.13) |
| 278 | 696 | 0.02 | A7I7F4 | *Methanoregula boonei* | Nitrogenase MoFe cofactor biosynthesis protein NifE (EC 1.18.6.1) |
| 279 | 696 | 0.02 | A7IB58 | *Methanoregula boonei* | Tyrosine--tRNA ligase (EC 6.1.1.1) (Tyrosyl-tRNA synthetase) (TyrRS) |
| 280 | 693 | 0.02 | A0A1D3L5L1 | *Methanobacterium congolense* | Multidrug resistance protein Stp |
| 281 | 693 | 0.02 | B8GJD9 | *Methanosphaerula palustris* | Ferrous iron transport protein B |
| 282 | 693 | 0.02 | F6D828 | *Methanobacterium paludis* | Formylmethanofuran dehydrogenase subunit A |
| 283 | 692 | 0.02 | A7I5T0 | *Methanoregula boonei* | Tetrahydromethanopterin S-methyltransferase subunit E (EC 2.1.1.86) (N5-methyltetrahydromethanopterin--coenzyme M methyltransferase subunit E) |
| 284 | 692 | 0.02 | L0HHJ6 | *Methanoregula formicica* | Tubulin-like protein CetZ |
| 285 | 692 | 0.02 | A7I5N9 | *Methanoregula boonei* | 50S ribosomal protein L3 |
| 286 | 690 | 0.02 | A0A1D3KZ99 | *Methanobacterium congolense* | Lysine--tRNA ligase (EC 6.1.1.6) (Lysyl-tRNA synthetase) (LysRS) |
| 287 | 690 | 0.02 | A0A1D3L4T1 | *Methanobacterium congolense* | Homoserine dehydrogenase (EC 1.1.1.3) |

| | | | | | |
|---|---|---|---|---|---|
| 288 | 689 | 0.02 | A0A1D3KZT0 | *Methanobacterium congolense* | Adenine deaminase (Adenase) (Adenine aminase) (EC 3.5.4.2) |
| 289 | 689 | 0.02 | A7I6G3 | *Methanoregula boonei* | Sodium/hydrogen exchanger |
| 290 | 689 | 0.02 | P06130 | *Methanobacterium formicicum* | Formate dehydrogenase subunit beta (EC 1.2.1.2) |
| 291 | 688 | 0.02 | A7I940 | *Methanoregula boonei* | Histidine--tRNA ligase (EC 6.1.1.21) (Histidyl-tRNA synthetase) (HisRS) |
| 292 | 687 | 0.02 | A0A1D3L220 | *Methanobacterium congolense* | Uncharacterized protein |
| 293 | 687 | 0.02 | A7I654 | *Methanoregula boonei* | Glycosyl transferase, family 39 |
| 294 | 686 | 0.02 | A0A1D3L2Q0 | *Methanobacterium congolense* | Glutamate--tRNA ligase (EC 6.1.1.17) (Glutamyl-tRNA synthetase) (GluRS) |
| 295 | 686 | 0.02 | L0HF89 | *Methanoregula formicica* | Formylmethanofuran dehydrogenase subunit B |
| 296 | 685 | 0.02 | A7I4T9 | *Methanoregula boonei* | Anthranilate synthase component 1 (EC 4.1.3.27) |
| 297 | 684 | 0.02 | A0A1D3KZR9 | *Methanobacterium congolense* | Putative PAS/PAC sensor protein |
| 298 | 684 | 0.02 | A0A1D3KZS3 | *Methanobacterium congolense* | Ketoisovalerate oxidoreductase subunit VorA (EC 1.-.-.-) |
| 299 | 684 | 0.02 | A7IA49 | *Methanoregula boonei* | Aminotransferase, class I and II |
| 300 | 684 | 0.02 | A7IAK3 | *Methanoregula boonei* | 50S ribosomal protein L10 (Acidic ribosomal protein P0 homolog) |
| 301 | 683 | 0.02 | A7IA50 | *Methanoregula boonei* | Diaminopimelate decarboxylase (DAP decarboxylase) (DAPDC) (EC 4.1.1.20) |
| 302 | 681 | 0.02 | A0A1D3L1B1 | *Methanobacterium congolense* | Putative LRR receptor-like serine/threonine-protein kinase At1g06840 (EC 2.7.11.1) |
| 303 | 678 | 0.02 | A7I890 | *Methanoregula boonei* | Serine--tRNA ligase (EC 6.1.1.11) (Seryl-tRNA synthetase) (SerRS) (Seryl-tRNA(Ser/Sec) synthetase) |
| 304 | 676 | 0.02 | A0A1D3L1Q0 | *Methanobacterium congolense* | Thermosome subunit alpha |
| 305 | 676 | 0.02 | A0A1D3L5T8 | *Methanobacterium congolense* | Putative transporter YebQ |
| 306 | 674 | 0.02 | A7I754 | *Methanoregula boonei* | Homospermidine synthase (EC 2.5.1.44) |
| 307 | 673 | 0.02 | A0A1D3L3L2 | *Methanobacterium congolense* | Putative iron-sulfur protein MMP1067 |
| 308 | 673 | 0.02 | A7I861 | *Methanoregula boonei* | Cell division protein FtsZ |
| 309 | 673 | 0.02 | A7IAL8 | *Methanoregula boonei* | Cell division protein FtsZ |
| 310 | 671 | 0.02 | A0A1D3L4J5 | *Methanobacterium congolense* | Uncharacterized protein |
| 311 | 670 | 0.02 | A0A1D3L4L0 | *Methanobacterium congolense* | Uncharacterized protein |
| 312 | 670 | 0.02 | A7I9I1 | *Methanoregula boonei* | Signal recognition particle 54 kDa protein (SRP54) |
| 313 | 669 | 0.02 | A7I7K0 | *Methanoregula boonei* | Ferredoxin |
| 314 | 669 | 0.02 | A7I9F2 | *Methanoregula boonei* | Small GTP-binding protein |
| 315 | 667 | 0.02 | A0A1D3L3I1 | *Methanobacterium congolense* | Glycine-tRNA ligase (EC 6.1.1.14) |
| 316 | 664 | 0.02 | A0A1D3L2K6 | *Methanobacterium congolense* | Uncharacterized protein |
| 317 | 664 | 0.02 | A0A1D3L4V1 | *Methanobacterium congolense* | Uncharacterized protein |
| 318 | 664 | 0.02 | A7I7F3 | *Methanoregula boonei* | Nitrogenase molybdenum-iron protein beta chain (EC 1.18.6.1) |
| 319 | 663 | 0.02 | A0A1D3L3K7 | *Methanobacterium congolense* | Uncharacterised protein family (UPF0240) |
| 320 | 663 | 0.02 | L0HI79 | *Methanoregula formicica* | CRISPR-associated protein Cas8c/Csd1, subtype I-C/DVULG |
| 321 | 663 | 0.02 | L0HKB8 | *Methanoregula formicica* | Uncharacterized protein |
| 322 | 660 | 0.02 | A0A1D3KZ83 | *Methanobacterium congolense* | Adenylosuccinate lyase (ASL) (EC 4.3.2.2) (Adenylosuccinase) |

| | | | | | |
|---|---|---|---|---|---|
| 323 | 659 | 0.02 | A0A1D3KZD9 | *Methanobacterium congolense* | Uncharacterized protein |
| 324 | 659 | 0.02 | A7I6T4 | *Methanoregula boonei* | Plasma-membrane proton-efflux P-type ATPase |
| 325 | 659 | 0.02 | A7I9A2 | *Methanoregula boonei* | Protein-export membrane protein SecD |
| 326 | 658 | 0.02 | A0A1D3L546 | *Methanobacterium congolense* | Multifunctional fusion protein [Includes: ADP-dependent (S)-NAD(P)H-hydrate dehydratase (EC 4.2.1.136) (ADP-dependent NAD(P)HX dehydratase); NAD(P)H-hydrate epimerase (EC 5.1.99.6) (NAD(P)HX epimerase)] |
| 327 | 657 | 0.02 | A0A1D3L0K7 | *Methanobacterium congolense* | Digeranylgeranylglycerophospholipid reductase (DGGGPL reductase) (EC 1.3.7.11) (2,3-bis-O-geranylgeranylglyceryl phosphate reductase) (Geranylgeranyl reductase) (GGR) |
| 328 | 657 | 0.02 | A0A1D3L5T3 | *Methanobacterium congolense* | Phosphoribosylamine--glycine ligase (EC 6.3.4.13) (GARS) (Glycinamide ribonucleotide synthetase) (Phosphoribosylglycinamide synthetase) |
| 329 | 657 | 0.02 | A7I8Z5 | *Methanoregula boonei* | Glutamine synthetase (EC 6.3.1.2) |
| 330 | 655 | 0.02 | A0A1D3L5A3 | *Methanobacterium congolense* | Coenzyme F420 hydrogenase subunit alpha (EC 1.12.98.1) |
| 331 | 654 | 0.02 | A7I630 | *Methanoregula boonei* | Pyruvate flavodoxin/ferredoxin oxidoreductase domain protein |
| 332 | 653 | 0.02 | A7I5V0 | *Methanoregula boonei* | Formylmethanofuran--tetrahydromethanopterin formyltransferase (EC 2.3.1.101) (H4MPT formyltransferase) |
| 333 | 652 | 0.02 | A0A1D3L2W2 | *Methanobacterium congolense* | Ribonuclease J (RNase J) (EC 3.1.-.-) |
| 334 | 650 | 0.02 | A0A1D3KZN6 | *Methanobacterium congolense* | Global nitrogen regulator NrpR |
| 335 | 650 | 0.02 | A0A1D3L065 | *Methanobacterium congolense* | Multidrug export protein EmrB |
| 336 | 649 | 0.02 | A0A1D3L563 | *Methanobacterium congolense* | Putative cysteine desulfurase (EC 2.8.1.7) |
| 337 | 649 | 0.02 | A0A1D3L5D6 | *Methanobacterium congolense* | Uncharacterized protein |
| 338 | 647 | 0.02 | A0A1D3L4I8 | *Methanobacterium congolense* | Polyferredoxin protein MvhB |
| 339 | 646 | 0.02 | A0A1D3KZI2 | *Methanobacterium congolense* | Probable tRNA pseudouridine synthase D (EC 5.4.99.27) (tRNA pseudouridine(13) synthase) (tRNA pseudouridylate synthase D) (tRNA-uridine isomerase D) |
| 340 | 645 | 0.02 | A0A1D3KZS6 | *Methanobacterium congolense* | O-phospho-L-seryl-tRNA:Cys-tRNA synthase (EC 2.5.1.73) (Sep-tRNA:Cys-tRNA synthase) (SepCysS) |
| 341 | 645 | 0.02 | A0A1D3L5K3 | *Methanobacterium congolense* | Putative NADH oxidase (EC 1.6.99.3) |
| 342 | 645 | 0.02 | A7I4I6 | *Methanoregula boonei* | Uncharacterized protein |
| 343 | 645 | 0.02 | A7I5Z2 | *Methanoregula boonei* | Uncharacterized protein |
| 344 | 644 | 0.02 | A0A1D3L0E2 | *Methanobacterium congolense* | Uncharacterized protein |
| 345 | 644 | 0.02 | A0A1D3L3U3 | *Methanobacterium congolense* | DNA-binding protein MutS2 |
| 346 | 644 | 0.02 | A7I5X5 | *Methanoregula boonei* | Uncharacterized protein |
| 347 | 644 | 0.02 | A7I7G8 | *Methanoregula boonei* | Anti-sigma-factor antagonist |
| 348 | 643 | 0.02 | A0A1D3KZ76 | *Methanobacterium congolense* | Putative 2-isopropylmalate synthase (EC 2.3.3.13) |
| 349 | 643 | 0.02 | A7I8A0 | *Methanoregula boonei* | Acetolactate synthase (EC 2.2.1.6) |
| 350 | 642 | 0.02 | A0A1D3KZK6 | *Methanobacterium congolense* | Glutamate decarboxylase (EC 4.1.1.15) |

| | | | | | |
|---|---|---|---|---|---|
| 351 | 640 | 0.02 | A7I6L5 | *Methanoregula boonei* | Aspartyl/glutamyl-tRNA(Asn/Gln) amidotransferase subunit B (Asp/Glu-ADT subunit B) (EC 6.3.5.-) |
| 352 | 639 | 0.02 | A0A1D3L102 | *Methanobacterium congolense* | Putative membrane protein YabM |
| 353 | 639 | 0.02 | A0A1D3L5F2 | *Methanobacterium congolense* | Diaminopimelate decarboxylase (DAP decarboxylase) (DAPDC) (EC 4.1.1.20) |
| 354 | 638 | 0.02 | A0A1D3KZ90 | *Methanobacterium congolense* | Phosphomethylpyrimidine synthase (EC 4.1.99.17) (Hydroxymethylpyrimidine phosphate synthase) (HMP-P synthase) (HMP-phosphate synthase) (HMPP synthase) (Thiamine biosynthesis protein ThiC) |
| 355 | 638 | 0.02 | A0A1D3KZA6 | *Methanobacterium congolense* | Phenylacetate-coenzyme A ligase (EC 6.2.1.30) |
| 356 | 638 | 0.02 | A0A1D3L3E2 | *Methanobacterium congolense* | Putative sensor histidine kinase pdtaS (EC 2.7.13.3) |
| 357 | 637 | 0.02 | A0A1D3L0A1 | *Methanobacterium congolense* | UDP-N-acetylmuramoyl-L-alanyl-D-glutamate-2,6-diaminopimelate ligase (EC 6.3.2.13) |
| 358 | 637 | 0.02 | A7I6N3 | *Methanoregula boonei* | Amidohydrolase 3 |
| 359 | 636 | 0.02 | A7I4C2 | *Methanoregula boonei* | Uncharacterized protein |
| 360 | 636 | 0.02 | A7IAT1 | *Methanoregula boonei* | Uncharacterized protein |
| 361 | 635 | 0.02 | A0A1D3L3M7 | *Methanobacterium congolense* | Putative glycosyltransferase YfnE (EC 2.4.-.-) |
| 362 | 635 | 0.02 | A0A1D3L571 | *Methanobacterium congolense* | Serine hydroxymethyltransferase (SHMT) (Serine methylase) (EC 2.1.2.-) |
| 363 | 635 | 0.02 | A0A1D3L5B2 | *Methanobacterium congolense* | Putative polyferredoxin-like protein MJ1303 |
| 364 | 634 | 0.02 | A0A1D3L1I3 | *Methanobacterium congolense* | Uncharacterized protein |
| 365 | 633 | 0.02 | A0A1D3L3D7 | *Methanobacterium congolense* | Sodium/pantothenate symporter |
| 366 | 633 | 0.02 | A0A1D3L4L3 | *Methanobacterium congolense* | Trk system potassium uptake protein TrkH |
| 367 | 633 | 0.02 | L0HK93 | *Methanoregula formicica* | Major Facilitator Superfamily transporter |
| 368 | 632 | 0.02 | A7I7K3 | *Methanoregula boonei* | CO dehydrogenase/acetyl-CoA synthase complex, beta subunit (EC 2.3.1.169) |
| 369 | 631 | 0.02 | A0A1D3L2S6 | *Methanobacterium congolense* | Putative sensor histidine kinase pdtaS (EC 2.7.13.3) |
| 370 | 631 | 0.02 | A7I7M3 | *Methanoregula boonei* | DNA mismatch repair protein MutS |
| 371 | 631 | 0.02 | L0HDG0 | *Methanoregula formicica* | SSS sodium solute transporter |
| 372 | 630 | 0.02 | A0A1D3L4N6 | *Methanobacterium congolense* | Inositol-3-phosphate synthase (EC 5.5.1.4) |
| 373 | 630 | 0.02 | A7I4X0 | *Methanoregula boonei* | SufBD protein |
| 374 | 629 | 0.02 | A0A1D3L243 | *Methanobacterium congolense* | CTP synthase (EC 6.3.4.2) (Cytidine 5'-triphosphate synthase) (Cytidine triphosphate synthetase) (CTP synthetase) (CTPS) (UTP--ammonia ligase) |
| 375 | 629 | 0.02 | A7I7L2 | *Methanoregula boonei* | Phosphoribosylaminoimidazolecarboxamide formyltransferase/IMP cyclohydrolase (EC 2.1.2.3) |
| 376 | 627 | 0.02 | A0A1D3L329 | *Methanobacterium congolense* | Uncharacterized protein |
| 377 | 627 | 0.02 | A7I942 | *Methanoregula boonei* | Type 2 DNA topoisomerase 6 subunit A (EC 5.99.1.3) (Type II DNA topoisomerase VI subunit A) |
| 378 | 626 | 0.02 | A7IAJ2 | *Methanoregula boonei* | Uncharacterized protein |
| 379 | 626 | 0.02 | N8ZVC9 | *Acinetobacter gerneri* | Uncharacterized protein |
| 380 | 625 | 0.02 | A7I860 | *Methanoregula boonei* | Putative phosphoserine phosphatase |
| 381 | 622 | 0.02 | A0A1D3KZC3 | *Methanobacterium congolense* | Glutamine synthetase (EC 6.3.1.2) |
| 382 | 622 | 0.02 | A0A1D3L4K2 | *Methanobacterium congolense* | Methyl-coenzyme M reductase subunit alpha (EC 2.8.4.1) |

| | | | | | |
|---|---|---|---|---|---|
| 383 | 622 | 0.02 | A7I615 | *Methanoregula boonei* | Phenylalanine--tRNA ligase beta subunit (EC 6.1.1.20) (Phenylalanyl-tRNA synthetase beta subunit) (PheRS) |
| 384 | 621 | 0.02 | A0A1D3L3Q2 | *Methanobacterium congolense* | Argininosuccinate lyase (ASAL) (EC 4.3.2.1) (Arginosuccinase) |
| 385 | 621 | 0.02 | A7I8N6 | *Methanoregula boonei* | Methyl-accepting chemotaxis sensory transducer |
| 386 | 621 | 0.02 | A7I8W3 | *Methanoregula boonei* | Uncharacterized protein |
| 387 | 620 | 0.02 | A7I698 | *Methanoregula boonei* | Glutamyl-tRNA(Gln) amidotransferase subunit E (Glu-ADT subunit E) (EC 6.3.5.-) |
| 388 | 617 | 0.02 | A7IA28 | *Methanoregula boonei* | 3-isopropylmalate dehydratase large subunit (EC 4.2.1.33) (Alpha-IPM isomerase) (IPMI) (Isopropylmalate isomerase) |
| 389 | 617 | 0.02 | Q8VQM8 | *Prevotella intermedia* | Mobilization protein |
| 390 | 616 | 0.02 | A0A1D3L0X2 | *Methanobacterium congolense* | Vitamin B12-dependent ribonucleotide reductase (EC 1.17.4.1) |
| 391 | 616 | 0.02 | A0A1D3L0Z7 | *Methanobacterium congolense* | Putative transporter MJ0709 |
| 392 | 615 | 0.02 | A7I6G2 | *Methanoregula boonei* | Sodium/hydrogen exchanger |
| 393 | 614 | 0.02 | A0A1D3L0C2 | *Methanobacterium congolense* | Putative lon protease homolog (EC 3.4.21.-) |
| 394 | 614 | 0.02 | A7I4T4 | *Methanoregula boonei* | Tryptophan synthase beta chain (EC 4.2.1.20) |
| 395 | 614 | 0.02 | A7IA56 | *Methanoregula boonei* | Small GTP-binding protein |
| 396 | 613 | 0.02 | A7I7Y7 | *Methanoregula boonei* | Uncharacterized protein |
| 397 | 613 | 0.02 | A7IAS3 | *Methanoregula boonei* | Uncharacterized protein |
| 398 | 612 | 0.02 | A0A1D3L0B1 | *Methanobacterium congolense* | Probable cobyric acid synthase |
| 399 | 612 | 0.02 | A7I6E9 | *Methanoregula boonei* | Uncharacterized protein |
| 400 | 612 | 0.02 | A7I7G0 | *Methanoregula boonei* | Beta-lactamase domain protein |
| 401 | 611 | 0.02 | A0A1D3L1P3 | *Methanobacterium congolense* | Putative MFS-type transporter EfpA |
| 402 | 611 | 0.02 | A0A1D3L472 | *Methanobacterium congolense* | D-3-phosphoglycerate dehydrogenase (EC 1.1.1.95) |
| 403 | 611 | 0.02 | A7I6J3 | *Methanoregula boonei* | Argininosuccinate synthase (EC 6.3.4.5) (Citrulline--aspartate ligase) |
| 404 | 610 | 0.02 | A0A1D3L4V5 | *Methanobacterium congolense* | Putative cation transporter MJ1485 |
| 405 | 610 | 0.02 | A7I4A1 | *Methanoregula boonei* | Radical SAM domain protein |
| 406 | 609 | 0.02 | A0A1D3L0F1 | *Methanobacterium congolense* | Uncharacterized protein |
| 407 | 609 | 0.02 | A7IA52 | *Methanoregula boonei* | DNA repair and recombination protein RadA |
| 408 | 609 | 0.02 | L0HBR8 | *Methanoregula formicica* | PAS domain S-box |
| 409 | 606 | 0.02 | A0A1D3KZ61 | *Methanobacterium congolense* | 5'-deoxyadenosine deaminase (5'-dA deaminase) (EC 3.5.4.41) |
| 410 | 605 | 0.02 | A0A1D3L2V0 | *Methanobacterium congolense* | Pyruvate carboxylase subunit A (EC 6.4.1.1) |
| 411 | 605 | 0.02 | A0A1D3L490 | *Methanobacterium congolense* | Lactaldehyde dehydrogenase (EC 1.2.1.22) |
| 412 | 605 | 0.02 | A0A1D3L5F7 | *Methanobacterium congolense* | Putative transporter YebQ |
| 413 | 605 | 0.02 | A7I5N4 | *Methanoregula boonei* | Cell division GTPase-like protein |
| 414 | 605 | 0.02 | A7I9R0 | *Methanoregula boonei* | Histone acetyltransferase, ELP3 family |
| 415 | 605 | 0.02 | A7IA39 | *Methanoregula boonei* | Beta-lactamase domain protein |
| 416 | 604 | 0.02 | A0A1D3L090 | *Methanobacterium congolense* | Probable phosphoglucosamine mutase (EC 5.4.2.10) |

| | | | | | |
|---|---|---|---|---|---|
| 417 | 604 | 0.02 | A0A1D3L4E2 | *Methanobacterium congolense* | RNA-metabolising metallo-beta-lactamase |
| 418 | 603 | 0.02 | A0A1D3KZ75 | *Methanobacterium congolense* | Cobyrinate a,c-diamide synthase (EC 6.3.5.11) (Cobyrinic acid a,c-diamide synthetase) |
| 419 | 603 | 0.02 | A7I8S2 | *Methanoregula boonei* | Diadenylate cyclase (DAC) (EC 2.7.7.85) (Cyclic-di-AMP synthase) (c-di-AMP synthase) |
| 420 | 602 | 0.02 | A0A1D3L4B7 | *Methanobacterium congolense* | Uncharacterized protein |
| 421 | 602 | 0.02 | A0A1D3L554 | *Methanobacterium congolense* | Divalent metal cation transporter MntH |
| 422 | 602 | 0.02 | A7I5S3 | *Methanoregula boonei* | Tetrahydromethanopterin S-methyltransferase, MtrH subunit |
| 423 | 602 | 0.02 | A7I8B3 | *Methanoregula boonei* | Ppx/GppA phosphatase |
| 424 | 601 | 0.02 | A0A1D3KZL5 | *Methanobacterium congolense* | Uncharacterized protein |
| 425 | 600 | 0.02 | A0A1D2WDD6 | *Methanobacterium sp.* | Nitrogenase iron-molybdenum cofactor biosynthesis protein NifN |
| 426 | 600 | 0.02 | A7I4I1 | *Methanoregula boonei* | Metal dependent phosphohydrolase |
| 427 | 599 | 0.02 | A0A1D3L3F6 | *Methanobacterium congolense* | Minor teichoic acid biosynthesis protein GgaB |
| 428 | 599 | 0.02 | A7I4L8 | *Methanoregula boonei* | Thiolase |
| 429 | 599 | 0.02 | A7IB45 | *Methanoregula boonei* | Asparagine synthase (Glutamine-hydrolyzing) (EC 6.3.5.4) |
| 430 | 598 | 0.02 | A7I5S8 | *Methanoregula boonei* | Tetrahydromethanopterin S-methyltransferase subunit C (EC 2.1.1.86) (N5-methyltetrahydromethanopterin--coenzyme M methyltransferase subunit C) |
| 431 | 598 | 0.02 | A7I763 | *Methanoregula boonei* | Conserved TM helix repeat-containing protein |
| 432 | 598 | 0.02 | A7I8Z4 | *Methanoregula boonei* | Ammonium transporter |
| 433 | 598 | 0.02 | A7I9D3 | *Methanoregula boonei* | Phosphoesterase, RecJ domain protein |
| 434 | 597 | 0.02 | A0A1D3L3J9 | *Methanobacterium congolense* | Replication factor C large subunit (RFC large subunit) (Clamp loader large subunit) |
| 435 | 597 | 0.02 | O27035 | *Methanothermobacter thermautotrophicus* | V-type ATP synthase beta chain (V-ATPase subunit B) |
| 436 | 596 | 0.02 | A7I865 | *Methanoregula boonei* | LOR/SDH bifunctional enzyme conserved domain protein |
| 437 | 596 | 0.02 | U6EDJ2 | *Methanobacterium sp.* | Uncharacterized protein |
| 438 | 595 | 0.02 | A0A1D3KZ06 | *Methanobacterium congolense* | Tryptophan synthase beta chain (EC 4.2.1.20) |
| 439 | 595 | 0.02 | A0A1D3KZP4 | *Methanobacterium congolense* | Phosphomethylpyrimidine synthase (EC 4.1.99.17) (Hydroxymethylpyrimidine phosphate synthase) (HMP-P synthase) (HMP-phosphate synthase) (HMPP synthase) (Thiamine biosynthesis protein ThiC) |
| 440 | 595 | 0.02 | A0A1D3L131 | *Methanobacterium congolense* | 7,8-dihydro-6-hydroxymethylpterin dimethyltransferase (EC 2.1.1.-) |
| 441 | 594 | 0.02 | A0A1D3L3K2 | *Methanobacterium congolense* | F420-non-reducing hydrogenase vhc subunit A (EC 1.12.99.-) |
| 442 | 594 | 0.02 | A0A1D3L3N9 | *Methanobacterium congolense* | Putative methyltransferase PH0819 (EC 2.1.1.-) |
| 443 | 594 | 0.02 | A7I7A0 | *Methanoregula boonei* | UvrABC system protein C (Protein UvrC) (Excinuclease ABC subunit C) |
| 444 | 593 | 0.02 | A7I6B2 | *Methanoregula boonei* | Uncharacterized protein |
| 445 | 591 | 0.02 | A7I928 | *Methanoregula boonei* | Uncharacterized protein |
| 446 | 590 | 0.02 | A0A1D3L252 | *Methanobacterium congolense* | Putative ABC transporter ATP-binding protein MJ1662 |
| 447 | 590 | 0.02 | A7I800 | *Methanoregula boonei* | Putative flagellar protein FlaH |
| 448 | 589 | 0.02 | A7I466 | *Methanoregula boonei* | Aminotransferase (EC 2.6.1.-) |

| | | | | | |
|---|---|---|---|---|---|
| 449 | 589 | 0.02 | A7I6P1 | *Methanoregula boonei* | Carbon monoxide dehydrogenase (EC 1.2.7.4) |
| 450 | 589 | 0.02 | A7IAA8 | *Methanoregula boonei* | Uncharacterized protein |
| 451 | 588 | 0.02 | A0A1D3L3V6 | *Methanobacterium congolense* | Methanogenesis regulatory histidine kinase FilI (EC 2.7.13.3) |
| 452 | 588 | 0.02 | A0A1D3L3V8 | *Methanobacterium congolense* | tRNA-splicing ligase RtcB (EC 6.5.1.-) |
| 453 | 587 | 0.02 | A0A1D3KZV6 | *Methanobacterium congolense* | Uncharacterized protein |
| 454 | 586 | 0.02 | A7IAG3 | *Methanoregula boonei* | Ribonuclease J (RNase J) (EC 3.1.-.-) |
| 455 | 585 | 0.02 | A0A1D3L5K1 | *Methanobacterium congolense* | Putative Rieske 2Fe-2S iron-sulfur protein YhfW (EC 1.-.-.-) |
| 456 | 585 | 0.02 | A7I4W6 | *Methanoregula boonei* | Phosphoglucomutase/phosphomannomutase alpha/beta/alpha domain I |
| 457 | 585 | 0.02 | A7I7F2 | *Methanoregula boonei* | Nitrogenase protein alpha chain (EC 1.18.6.1) |
| 458 | 585 | 0.02 | A7I9R7 | *Methanoregula boonei* | Proline--tRNA ligase (EC 6.1.1.15) (Prolyl-tRNA synthetase) (ProRS) |
| 459 | 584 | 0.02 | A0A1D3L0W3 | *Methanobacterium congolense* | AAA ATPase |
| 460 | 584 | 0.02 | A7IA30 | *Methanoregula boonei* | 3-isopropylmalate dehydrogenase (EC 1.1.1.85) |
| 461 | 583 | 0.02 | A0A1D3L581 | *Methanobacterium congolense* | Coenzyme A biosynthesis bifunctional protein CoaBC (EC 4.1.1.36) |
| 462 | 582 | 0.02 | A0A1D3L414 | *Methanobacterium congolense* | Uncharacterized protein |
| 463 | 582 | 0.02 | A7I4F1 | *Methanoregula boonei* | Phosphoribosylformylglycinamidine cyclo-ligase (EC 6.3.3.1) (AIR synthase) (AIRS) (Phosphoribosyl-aminoimidazole synthetase) |
| 464 | 581 | 0.02 | A0A1D3KZ63 | *Methanobacterium congolense* | Bacterial Ig-like domain (Group 3) |
| 465 | 581 | 0.02 | A0A1D3L3A3 | *Methanobacterium congolense* | ABC-2 type transporter |
| 466 | 581 | 0.02 | A7I4A7 | *Methanoregula boonei* | Thiamine biosynthesis protein ThiC |
| 467 | 581 | 0.02 | A7I606 | *Methanoregula boonei* | DEAD/DEAH box helicase domain protein |
| 468 | 581 | 0.02 | A7I899 | *Methanoregula boonei* | Putative (R)-citramalate synthase CimA (EC 2.3.1.182) |
| 469 | 579 | 0.02 | A7IAM6 | *Methanoregula boonei* | Radical SAM domain protein |
| 470 | 578 | 0.02 | A0A1D3L014 | *Methanobacterium congolense* | GAF domain protein |
| 471 | 578 | 0.02 | A0A1D3L364 | *Methanobacterium congolense* | Enolase (EC 4.2.1.11) (2-phospho-D-glycerate hydro-lyase) (2-phosphoglycerate dehydratase) |
| 472 | 578 | 0.02 | A0A1D3L5L2 | *Methanobacterium congolense* | Uncharacterized protein |
| 473 | 578 | 0.02 | A7I5Z9 | *Methanoregula boonei* | Uncharacterized protein |
| 474 | 578 | 0.02 | A7I6B3 | *Methanoregula boonei* | ABC transporter related |
| 475 | 578 | 0.02 | A7IB43 | *Methanoregula boonei* | Uncharacterized protein |
| 476 | 578 | 0.02 | L0HFI7 | *Methanoregula formicica* | Type I restriction-modification system methyltransferase subunit |
| 477 | 577 | 0.02 | A0A1D3L3X0 | *Methanobacterium congolense* | Acetolactate synthase large subunit (EC 2.2.1.6) |
| 478 | 577 | 0.02 | A0A1D3L5H4 | *Methanobacterium congolense* | Uncharacterized protein |
| 479 | 576 | 0.02 | A0A1D3KZC4 | *Methanobacterium congolense* | Inner membrane transport permease YbhR |
| 480 | 576 | 0.02 | A0A1D3L5J0 | *Methanobacterium congolense* | Type A flavoprotein FprA (EC 1.-.-.-) |
| 481 | 576 | 0.02 | A7IA57 | *Methanoregula boonei* | Uncharacterized protein |
| 482 | 575 | 0.02 | A0A1D3L4S7 | *Methanobacterium congolense* | Glucose-methanol-choline oxidoreductase |
| 483 | 575 | 0.02 | | | |

| | | | | | |
|---|---|---|---|---|---|
| 484 | 574 | 0.02 | A7IAQ1 | *Methanoregula boonei* | Uncharacterized protein |
| 485 | 574 | 0.02 | A7I7L0 | *Methanoregula boonei* | Dihydroxy-acid dehydratase (DAD) (EC 4.2.1.9) |
| 486 | 574 | 0.02 | A7I771 | *Methanoregula boonei* | S-adenosylmethionine synthase (AdoMet synthase) (EC 2.5.1.6) (Methionine adenosyltransferase) |
| 487 | 573 | 0.02 | A0A089ZFB5 | *Methanobacterium formicicum* | Adhesin-like protein |
| 488 | 573 | 0.02 | A0A1D3L0L3 | *Methanobacterium congolense* | Amidophosphoribosyltransferase (ATase) (EC 2.4.2.14) (Glutamine phosphoribosylpyrophosphate amidotransferase) (GPATase) |
| 489 | 572 | 0.02 | A0A1D3KZ70 | *Methanobacterium congolense* | Outer membrane protein assembly factor BamB |
| 490 | 572 | 0.02 | A0A1D3L586 | *Methanobacterium congolense* | S-adenosylmethionine synthase (AdoMet synthase) (EC 2.5.1.6) (Methionine adenosyltransferase) |
| 491 | 572 | 0.02 | A7I4P1 | *Methanoregula boonei* | Nucleic acid binding, OB-fold, tRNA/helicase-type |
| 492 | 572 | 0.02 | A7I4Q6 | *Methanoregula boonei* | tRNA(Ile2) 2-agmatinylcytidine synthetase TiaS (tRNA(Ile2)-agm2C synthetase) (EC 6.3.4.22) (tRNA(Ile2) agmatidine synthetase) |
| 493 | 571 | 0.02 | A0A1D3KZS9 | *Methanobacterium congolense* | Uncharacterized protein |
| 494 | 571 | 0.02 | A0A1D3L0S3 | *Methanobacterium congolense* | Single-stranded-DNA-specific exonuclease RecJ (EC 3.1.-.-) |
| 495 | 571 | 0.02 | A7I4L9 | *Methanoregula boonei* | UPF0219 protein Mboo_0157 |
| 496 | 571 | 0.02 | A7I7W3 | *Methanoregula boonei* | Molybdenum cofactor synthesis domain |
| 497 | 570 | 0.02 | A7I7C4 | *Methanoregula boonei* | DEAD_2 domain protein |
| 498 | 570 | 0.02 | A7I9H9 | *Methanoregula boonei* | tRNA pseudouridine synthase Pus10 (EC 5.4.99.25) (tRNA pseudouridine 54/55 synthase) (Psi54/55 synthase) |
| 499 | 569 | 0.02 | A0A1D3L3L1 | *Methanobacterium congolense* | Ammonium transporter |
| 500 | 569 | 0.02 | A0A1D3L510 | *Methanobacterium congolense* | Uncharacterized protein |

**Table S5c.** Functional overview of M2P community (top 500 rank only).

| Rank | Count | Abundance | ID | Organism | Protein Names |
|---|---|---|---|---|---|
| 1 | 3587 | 0.11 | N8ZVC9 | *Acinetobacter gerneri* | Uncharacterized protein |
| 2 | 3329 | 0.10 | Q8VQM8 | *Prevotella intermedia* | Mobilization protein |
| 3 | 2921 | 0.09 | A0A0A2LWZ9 | *Flavobacterium suncheonense* | Transposase |
| 4 | 2578 | 0.08 | N8ZNX3 | *Acinetobacter gerneri* | Uncharacterized protein |
| 5 | 2045 | 0.06 | F0FB90 | *Prevotella multiformis* | Initiator RepB protein (Fragment) |
| 6 | 2007 | 0.06 | A0A0R3QEE4 | *Brugia timori* | Uncharacterized protein |
| 7 | 1817 | 0.06 | A0A0U1CQ28 | *Chlamydia trachomatis* | Replication protein |
| 8 | 1814 | 0.06 | F0FB88 | *Prevotella multiformis* | Putative plasmid recombination enzyme |
| 9 | 1491 | 0.05 | A0A0U1CQ01 | *Chlamydia trachomatis* | Uncharacterised protein |
| 10 | 1470 | 0.05 | A0A0Q7S8L4 | *Pelomonas sp.* | Transposase |
| 11 | 1443 | 0.04 | A0A077XLW5 | *Sphinx1.76-related DNA* | Replication protein |
| 12 | 1443 | 0.04 | | | |
| 13 | 1422 | 0.04 | A0A1Q3RZJ5 | *Flavobacterium sp.* | Uncharacterized protein |
| 14 | 1339 | 0.04 | F4GG80 | *Alicycliphilus denitrificans* | Transposase IS4 family protein |
| 15 | 1331 | 0.04 | Q8VQM9 | *Prevotella intermedia* | Replication protein |
| 16 | 1310 | 0.04 | A0A077XNB6 | *Sphinx1.76-related DNA* | Replication protein |
| 17 | 1240 | 0.04 | A0A1Q3SCF5 | *Flavobacterium sp.* | Uncharacterized protein |
| 18 | 1094 | 0.03 | A0A1E4A455 | *Chryseobacterium sp.* | Cell surface protein SprA |
| 19 | 1076 | 0.03 | A0A1Q3S858 | *Flavobacterium sp.* | Uncharacterized protein |
| 20 | 1070 | 0.03 | | | |
| 21 | 1055 | 0.03 | A0A1Q3RWC6 | *Flavobacterium sp.* | Uncharacterized protein |
| 22 | 1051 | 0.03 | A0A0E9G9S8 | *Chlamydia trachomatis* | Uncharacterised protein |
| 23 | 957 | 0.03 | A0A1B1ICB8 | *Prevotella scopos* | Uncharacterized protein |
| 24 | 943 | 0.03 | N8YFI4 | *Acinetobacter gerneri* | Uncharacterized protein |
| 25 | 942 | 0.03 | A0A1E3ZQ34 | *Chryseobacterium sp.* | Cell envelope biogenesis protein OmpA |
| 26 | 895 | 0.03 | C9MTU8 | *Prevotella veroralis* | Initiator RepB protein |
| 27 | 880 | 0.03 | A0A023UP33 | *Magnusiomyces ingens* | Uncharacterized protein (Fragment) |
| 28 | 848 | 0.03 | A0A0Q8GZB4 | *Pelomonas sp.* | UvrABC system protein A |
| 29 | 833 | 0.03 | A0A1Q3S9F2 | *Flavobacterium sp.* | Uncharacterized protein |
| 30 | 824 | 0.03 | A0A1Q3S0Y5 | *Flavobacterium sp.* | Uncharacterized protein |
| 31 | 803 | 0.02 | G9FB24 | *uncultured bacterium* | IS21 family transposition protein |
| 32 | 765 | 0.02 | A0A0Q7SV37 | *Pelomonas sp.* | Glutamate synthase |
| 33 | 745 | 0.02 | A0A0Q7S7D9 | *Pelomonas sp.* | Uncharacterized protein |

| | | | | | |
|---|---|---|---|---|---|
| 34 | 741 | 0.02 | A0A0Q7RZB1 | *Pelomonas sp.* | Uncharacterized protein |
| 35 | 730 | 0.02 | A0A023UP27 | *Magnusiomyces ingens* | Uncharacterized protein |
| 36 | 726 | 0.02 | A0A1Q3M6H6 | *Bacteroidales bacterium* | CRISPR-associated endonuclease Cas9 (EC 3.1.-.-) |
| 37 | 719 | 0.02 | A0A0Q6WNI0 | *Pelomonas sp.* | DNA-directed RNA polymerase subunit beta' (RNAP subunit beta') (EC 2.7.7.6) (RNA polymerase subunit beta') (Transcriptase subunit beta') |
| 38 | 709 | 0.02 | A0A0T5YH86 | *Acinetobacter baumannii* | Mobilization protein A |
| 39 | 670 | 0.02 | A0A1K2IH81 | *Chryseobacterium limigenitum* | Uncharacterized protein |
| 40 | 661 | 0.02 | A0A0Q7RZF8 | *Pelomonas sp.* | Phosphoribosylformylglycinamidine synthase (FGAM synthase) (FGAMS) (EC 6.3.5.3) (Formylglycinamide ribonucleotide amidotransferase) (FGAR amidotransferase) (FGAR-AT) |
| 41 | 655 | 0.02 | E4T3F9 | *Paludibacter propionicigenes* | Glutamate synthase (NADH) large subunit (EC 1.4.1.14) (EC 1.4.7.1) |
| 42 | 650 | 0.02 | A0A1Q3S812 | *Flavobacterium sp.* | Uncharacterized protein |
| 43 | 647 | 0.02 | A0A0Q7SS31 | *Pelomonas sp.* | Transposase |
| 44 | 647 | 0.02 | A0A1E4A9F0 | *Chryseobacterium sp.* | Uncharacterized protein |
| 45 | 623 | 0.02 | A0A0Q7S6W6 | *Pelomonas sp.* | Indolepyruvate ferredoxin oxidoreductase (EC 1.2.7.8) |
| 46 | 621 | 0.02 | A0A0M8MCD5 | *Flavobacterium akiainvivens* | Integrase |
| 47 | 611 | 0.02 | A0A0Q6UG42 | *Pelomonas sp.* | Pyruvate ferredoxin oxidoreductase |
| 48 | 610 | 0.02 | A0A1L9GVN0 | *Cand. Micrarchaeum acidiphilum* | Uncharacterized protein |
| 49 | 608 | 0.02 | A0A0Q7SCT5 | *Pelomonas sp.* | Chromosome partition protein Smc |
| 50 | 597 | 0.02 | A0A0Q7SRI3 | *Pelomonas sp.* | Efflux pump membrane transporter |
| 51 | 596 | 0.02 | A0A0Q6WK82 | *Pelomonas sp.* | RND transporter |
| 52 | 593 | 0.02 | A0A0D2WNJ4 | *Capsaspora owczarzaki* | Paramyosin |
| 53 | 591 | 0.02 | V6RWN6 | *Flavobacterium cauense* | Uncharacterized protein |
| 54 | 590 | 0.02 | A0A1J5AN19 | *Bacteroidetes bacterium* | IS110 family transposase |
| 55 | 586 | 0.02 | A0A1Q3RUL1 | *Flavobacterium sp.* | Uncharacterized protein |
| 56 | 583 | 0.02 | A0A0Q7S7G4 | *Pelomonas sp.* | Efflux pump membrane transporter |
| 57 | 572 | 0.02 | A0A0Q7SMD8 | *Pelomonas sp.* | Acriflavine resistance protein B |
| 58 | 571 | 0.02 | A0A0Q7SKX9 | *Pelomonas sp.* | ATP-dependent helicase |
| 59 | 564 | 0.02 | A0A0R0LGW3 | *Ferrovum sp.* | ATP-dependent RNA helicase RhlE (EC 3.6.4.13) |
| 60 | 559 | 0.02 | A0A0Q8GR06 | *Pelomonas sp.* | Alpha-2-macroglobulin |
| 61 | 550 | 0.02 | A0A0Q7SFB9 | *Pelomonas sp.* | Multidrug transporter |
| 62 | 541 | 0.02 | A0A0Q7SBL7 | *Pelomonas sp.* | Efflux pump membrane transporter |
| 63 | 535 | 0.02 | A0A0Q6UIJ2 | *Pelomonas sp.* | Nitrate reductase |
| 64 | 534 | 0.02 | A0A0D8Y6E2 | *Dictyocaulus viviparus* | Uncharacterized protein |
| 65 | 531 | 0.02 | A0A0Q7S584 | *Pelomonas sp.* | DNA-directed DNA polymerase (EC 2.7.7.7) |

| | | | | | |
|---|---|---|---|---|---|
| 66 | 525 | 0.02 | A0A0Q7SJP7 | *Pelomonas sp.* | Phosphoenolpyruvate carboxylase (PEPC) (PEPCase) (EC 4.1.1.31) |
| 67 | 524 | 0.02 | A0A1Q3S0Z6 | *Flavobacterium sp.* | Uncharacterized protein |
| 68 | 522 | 0.02 | A0A0Q7RZ30 | *Pelomonas sp.* | Acriflavin resistance protein |
| 69 | 521 | 0.02 | A0A0Q7SL82 | *Pelomonas sp.* | Multidrug transporter AcrB |
| 70 | 520 | 0.02 | A0A1E3ZXL4 | *Chryseobacterium sp.* | Uncharacterized protein |
| 71 | 512 | 0.02 | A0A1Q3RUG7 | *Flavobacterium sp.* | Uncharacterized protein |
| 72 | 512 | 0.02 | A0A1Q3S3S6 | *Flavobacterium sp.* | Uncharacterized protein |
| 73 | 511 | 0.02 | A0A1Q3S3X4 | *Flavobacterium sp.* | Uncharacterized protein |
| 74 | 509 | 0.02 | E4T8J6 | *Paludibacter propionicigenes* | Cell surface receptor IPT/TIG domain protein |
| 75 | 507 | 0.02 | A0A167ZLW1 | *Flavobacterium fryxellicola* | Uncharacterized protein |
| 76 | 507 | 0.02 | A0A1Q3RUG1 | *Flavobacterium sp.* | Uncharacterized protein |
| 77 | 505 | 0.02 | A0A0Q6U523 | *Pelomonas sp.* | Nitric oxide reductase large subunit |
| 78 | 505 | 0.02 | G0IXG6 | *Cyclobacterium marinum* | Transposase IS116/IS110/IS902 family protein |
| 79 | 504 | 0.02 | A0A1E4ARQ1 | *Chryseobacterium sp.* | Uncharacterized protein |
| 80 | 498 | 0.02 | A0A127MR59 | *Pseudomonas citronellolis* | Uncharacterized protein |
| 81 | 493 | 0.02 | A0A1E4ABZ6 | *Chryseobacterium sp.* | Multifunctional fusion protein [Includes: Protein translocase subunit SecD; Protein-export membrane protein SecF] |
| 82 | 492 | 0.02 | A0A1Q3RVF6 | *Flavobacterium sp.* | Multidrug transporter AcrB |
| 83 | 491 | 0.02 | A0A1T5HUH3 | *Alkalitalea saponilacus* | Uncharacterized protein |
| 84 | 490 | 0.02 | A0A0R3R918 | *Brugia timori* | Uncharacterized protein |
| 85 | 487 | 0.02 | | | |
| 86 | 486 | 0.02 | A0A1Q3S042 | *Flavobacterium sp.* | Peptidase C25 |
| 87 | 483 | 0.01 | A0A1E4AD77 | *Chryseobacterium sp.* | Uncharacterized protein |
| 88 | 482 | 0.01 | A0A1E4AA02 | *Chryseobacterium sp.* | TonB-dependent receptor |
| 89 | 478 | 0.01 | A0A1Q3S6M2 | *Flavobacterium sp.* | Uncharacterized protein |
| 90 | 477 | 0.01 | A0A0A2M931 | *Flavobacterium suncheonense* | Uncharacterized protein (Fragment) |
| 91 | 477 | 0.01 | G9FB23 | *uncultured bacterium* | IS21 family transposition protein |
| 92 | 474 | 0.01 | A0A0Q6T5H8 | *Pelomonas sp.* | Nodulation protein NolG |
| 93 | 473 | 0.01 | A0A1E4A330 | *Chryseobacterium sp.* | Acriflavin resistance protein |
| 94 | 470 | 0.01 | A0A0Q7T1P6 | *Pelomonas sp.* | 3-hydroxyacyl-CoA dehydrogenase |
| 95 | 470 | 0.01 | A0A1E4ADX1 | *Chryseobacterium sp.* | Uncharacterized protein |
| 96 | 469 | 0.01 | A0A0Q7SAQ8 | *Pelomonas sp.* | Bifunctional uridylyltransferase/uridylyl-removing enzyme (UTase/UR) (Bifunctional [protein-PII] modification enzyme) (Bifunctional nitrogen sensor protein) [Includes: [Protein-PII]-UMP uridylyl-removing enzyme (UR) (EC 3.1.4.-); [Protein-PII] uridylyltransferase (PII uridylyltransferase) (UTase) (EC 2.7.7.59)] |
| 97 | 469 | 0.01 | A0A0Q7SNJ9 | *Pelomonas sp.* | Cyanophycin synthetase |

| | | | | | |
|---|---|---|---|---|---|
| 98 | 465 | 0.01 | A0A1E4AFR9 | *Chryseobacterium sp.* | Uncharacterized protein |
| 99 | 464 | 0.01 | A0A177LQD3 | *Chryseobacterium sp.* | Uncharacterized protein |
| 100 | 464 | 0.01 | A0A1Q3RVI5 | *Flavobacterium sp.* | Uncharacterized protein |
| 101 | 463 | 0.01 | A0A0Q7SA64 | *Pelomonas sp.* | Transcription-repair-coupling factor (TRCF) (EC 3.6.4.-) |
| 102 | 462 | 0.01 | A0A1Q3S6H8 | *Flavobacterium sp.* | Uncharacterized protein |
| 103 | 461 | 0.01 | A0A0Q7SE50 | *Pelomonas sp.* | Bifunctional glutamine synthetase adenylyltransferase/adenylyl-removing enzyme (ATP:glutamine synthetase adenylyltransferase) (ATase) [Includes: Glutamine synthetase adenylyl-L-tyrosine phosphorylase (EC 2.7.7.89) (Adenylyl removase) (AR) (AT-N); Glutamine synthetase adenylyl transferase (EC 2.7.7.42) (Adenylyl transferase) (AT) (AT-C)] |
| 104 | 460 | 0.01 | A0A023UMT3 | *Magnusiomyces ingens* | Uncharacterized protein (Fragment) |
| 105 | 459 | 0.01 | A0A023UMS7 | *Magnusiomyces ingens* | NADH-ubiquinone oxidoreductase chain 5 (EC 1.6.5.3) |
| 106 | 459 | 0.01 | | | |
| 107 | 454 | 0.01 | A0A0Q8H5F4 | *Pelomonas sp.* | TonB-dependent receptor |
| 108 | 453 | 0.01 | A0A0Q6WN69 | *Pelomonas sp.* | Histidine kinase (EC 2.7.13.3) |
| 109 | 453 | 0.01 | A0A0Q7SL61 | *Pelomonas sp.* | 2-oxoglutarate dehydrogenase subunit E1 (EC 1.2.4.2) |
| 110 | 451 | 0.01 | A0A0Q7S182 | *Pelomonas sp.* | Phenylalanine--tRNA ligase beta subunit (EC 6.1.1.20) (Phenylalanyl-tRNA synthetase beta subunit) (PheRS) |
| 111 | 451 | 0.01 | A0A1Q3S3Z3 | *Flavobacterium sp.* | DNA methylase |
| 112 | 449 | 0.01 | A0A0Q6T477 | *Pelomonas sp.* | Uncharacterized protein |
| 113 | 448 | 0.01 | A0A1E4AD73 | *Chryseobacterium sp.* | Outer membrane protein assembly factor BamA |
| 114 | 446 | 0.01 | A0A177LRY5 | *Chryseobacterium sp.* | Uncharacterized protein |
| 115 | 445 | 0.01 | E4T5D1 | *Paludibacter propionicigenes* | Methionine--tRNA ligase (EC 6.1.1.10) (Methionyl-tRNA synthetase) (MetRS) |
| 116 | 443 | 0.01 | A0A0Q7SUY8 | *Pelomonas sp.* | Enoyl-CoA hydratase |
| 117 | 441 | 0.01 | A0A1E4A2B9 | *Chryseobacterium sp.* | Quinol:cytochrome C oxidoreductase |
| 118 | 439 | 0.01 | A0A1Q3S2G4 | *Flavobacterium sp.* | Protein translocase subunit SecA |
| 119 | 438 | 0.01 | A0A0Q7S238 | *Pelomonas sp.* | Histidine kinase |
| 120 | 438 | 0.01 | A0A0Q7S7S5 | *Pelomonas sp.* | Valine--tRNA ligase (EC 6.1.1.9) (Valyl-tRNA synthetase) (ValRS) |
| 121 | 438 | 0.01 | | | |
| 122 | 435 | 0.01 | A0A1E4A8D6 | *Chryseobacterium sp.* | Metallophosphoesterase |
| 123 | 434 | 0.01 | A0A0Q6TFR1 | *Pelomonas sp.* | Uncharacterized protein |
| 124 | 434 | 0.01 | A0A0Q7SAX3 | *Pelomonas sp.* | Glycine dehydrogenase (decarboxylating) (EC 1.4.4.2) (Glycine cleavage system P-protein) (Glycine decarboxylase) (Glycine dehydrogenase (aminomethyl-transferring)) |
| 125 | 430 | 0.01 | A0A1Q3RYV8 | *Flavobacterium sp.* | 2-oxoglutarate dehydrogenase E1 component |
| 126 | 429 | 0.01 | A0A0Q7SY69 | *Flavobacterium sp.* | DNA helicase (EC 3.6.4.12) |
| 127 | 429 | 0.01 | A0A0Q8H242 | *Pelomonas sp.* | PAS domain-containing sensor histidine kinase |
| 128 | 427 | 0.01 | A0A1E4AHR4 | *Chryseobacterium sp.* | Uncharacterized protein |

| | | | | | |
|---|---|---|---|---|---|
| 129 | 427 | 0.01 | A0A1Q3S919 | *Flavobacterium sp.* | SusC/RagA family protein |
| 130 | 426 | 0.01 | A0A0Q7SED5 | *Pelomonas sp.* | Alanine--tRNA ligase (EC 6.1.1.7) (Alanyl-tRNA synthetase) (AlaRS) |
| 131 | 424 | 0.01 | A0A0Q6TP10 | *Pelomonas sp.* | Uncharacterized protein |
| 132 | 424 | 0.01 | A0A0Q7SEF7 | *Pelomonas sp.* | Translation initiation factor IF-2 |
| 133 | 424 | 0.01 | A0A0Q7SN01 | *Pelomonas sp.* | ATPase |
| 134 | 424 | 0.01 | A0A0Q7SSX8 | *Pelomonas sp.* | Fused isobutyryl-CoA mutase [Includes: Isobutyryl-CoA mutase (ICM) (EC 5.4.99.13); P-loop GTPase (EC 3.6.5.-) (G-protein chaperone)] |
| 135 | 423 | 0.01 | A0A0Q7SRF6 | *Pelomonas sp.* | DNA topoisomerase III |
| 136 | 422 | 0.01 | A0A0Q7SL07 | *Pelomonas sp.* | DNA polymerase I (EC 2.7.7.7) |
| 137 | 421 | 0.01 | A0A0Q7SDY4 | *Pelomonas sp.* | GCN5 family acetyltransferase |
| 138 | 421 | 0.01 | A0A1Q3S993 | *Flavobacterium sp.* | Uncharacterized protein |
| 139 | 420 | 0.01 | A0A0Q6W7U3 | *Pelomonas sp.* | Uncharacterized protein |
| 140 | 420 | 0.01 | A0A0Q7SXZ5 | *Pelomonas sp.* | Aminopeptidase |
| 141 | 419 | 0.01 | A0A1Q3RUH2 | *Flavobacterium sp.* | Uncharacterized protein |
| 142 | 417 | 0.01 | A0A0Q8GKD4 | *Pelomonas sp.* | Uncharacterized protein |
| 143 | 416 | 0.01 | A0A1G4VJQ3 | *Flavobacterium saliperosum* | Gliding motility-associated C-terminal domain-containing protein |
| 144 | 416 | 0.01 | A0A1Q3S8X4 | *Flavobacterium sp.* | Cation transporter |
| 145 | 416 | 0.01 | A0A1Q3S9A2 | *Flavobacterium sp.* | Uncharacterized protein |
| 146 | 415 | 0.01 | A0A0Q6WDT0 | *Pelomonas sp.* | Copper-transporting ATPase |
| 147 | 415 | 0.01 | A0A0Q7SCD2 | *Pelomonas sp.* | Helicase |
| 148 | 414 | 0.01 | A0A0Q7S4W2 | *Pelomonas sp.* | FAD-linked oxidase |
| 149 | 414 | 0.01 | A0A0Q7SD92 | *Pelomonas sp.* | Uncharacterized protein |
| 150 | 414 | 0.01 | A0A0Q7SL81 | *Pelomonas sp.* | Uncharacterized protein |
| 151 | 413 | 0.01 | A0A0Q7S0G4 | *Pelomonas sp.* | AAA family ATPase |
| 152 | 412 | 0.01 | | | |
| 153 | 411 | 0.01 | A0A0Q6SZK0 | *Pelomonas sp.* | Protein translocase subunit SecA |
| 154 | 411 | 0.01 | A0A0Q7SJ43 | *Pelomonas sp.* | Transposase |
| 155 | 410 | 0.01 | A0A1Q3S1D7 | *Flavobacterium sp.* | TonB-dependent receptor |
| 156 | 408 | 0.01 | A0A0Q7SLN7 | *Pelomonas sp.* | UvrABC system protein A (UvrA protein) (Excinuclease ABC subunit A) |
| 157 | 408 | 0.01 | A0A0Q7T373 | *Pelomonas sp.* | Ribonuclease R (RNase R) (EC 3.1.13.1) |
| 158 | 408 | 0.01 | A0A1E4AC15 | *Chryseobacterium sp.* | Uncharacterized protein |
| 159 | 407 | 0.01 | A0A1Q3S5R1 | *Flavobacterium sp.* | Uncharacterized protein |
| 160 | 406 | 0.01 | A0A1Q3SAB4 | *Flavobacterium sp.* | TonB-dependent receptor |
| 161 | 405 | 0.01 | A0A0Q7RYH2 | *Pelomonas sp.* | 4-deoxy-4-formamido-L-arabinose-phospho-UDP deformylase |
| 162 | 404 | 0.01 | A0A0Q7SEL2 | *Pelomonas sp.* | Uncharacterized protein |
| 163 | 403 | 0.01 | A0A0Q7SCD0 | *Pelomonas sp.* | NADPH-dependent 2,4-dienoyl-CoA reductase (EC 1.3.1.34) |

| | | | | | |
|---|---|---|---|---|---|
| 164 | 403 | 0.01 | A0A1Q3S7F1 | *Flavobacterium sp.* | Beta-N-acetylglucosaminidase |
| 165 | 402 | 0.01 | A0A0Q8GQS0 | *Pelomonas sp.* | DNA mismatch repair protein MutS |
| 166 | 400 | 0.01 | A0A0Q7S059 | *Pelomonas sp.* | DNA topoisomerase 4 subunit A (EC 5.99.1.3) (Topoisomerase IV subunit A) |
| 167 | 399 | 0.01 | A0A1Q3RXA0 | *Flavobacterium sp.* | Uncharacterized protein |
| 168 | 399 | 0.01 | A0A1Q3RYP3 | *Flavobacterium sp.* | DNA-directed DNA polymerase (EC 2.7.7.7) |
| 169 | 398 | 0.01 | A0A0Q6U1A9 | *Pelomonas sp.* | Uncharacterized protein |
| 170 | 398 | 0.01 | A0A0Q6WQC8 | *Pelomonas sp.* | Uncharacterized protein |
| 171 | 398 | 0.01 | A0A0Q7T172 | *Pelomonas sp.* | Ribonucleoside-diphosphate reductase (EC 1.17.4.1) |
| 172 | 397 | 0.01 | A0A0Q7SLM8 | *Pelomonas sp.* | Tat pathway signal sequence domain protein |
| 173 | 397 | 0.01 | A0A0Q7SNK2 | *Pelomonas sp.* | Short-chain dehydrogenase |
| 174 | 397 | 0.01 | A0A0Q8GMD4 | *Pelomonas sp.* | Uncharacterized protein |
| 175 | 396 | 0.01 | A0A060RAF8 | *Mucinivorans hirudinis* | Mobile element protein |
| 176 | 396 | 0.01 | A0A0Q7SDQ5 | *Pelomonas sp.* | Efflux pump membrane transporter |
| 177 | 396 | 0.01 | A0A1E4A6N2 | *Chryseobacterium sp.* | Alkaline phosphatase |
| 178 | 395 | 0.01 | A0A1Q3S7G4 | *Flavobacterium sp.* | Acriflavin resistance protein |
| 179 | 394 | 0.01 | A0A1E4A9M1 | *Chryseobacterium sp.* | Cation transporter |
| 180 | 393 | 0.01 | A0A1E4AHQ8 | *Chryseobacterium sp.* | TonB-dependent receptor |
| 181 | 391 | 0.01 | A0A0Q6U3Y0 | *Pelomonas sp.* | Uncharacterized protein |
| 182 | 390 | 0.01 | A0A0Q7SDJ3 | *Pelomonas sp.* | Iron permease |
| 183 | 390 | 0.01 | A0A0Q7SRK7 | *Pelomonas sp.* | Leucine--tRNA ligase (EC 6.1.1.4) (Leucyl-tRNA synthetase) (LeuRS) |
| 184 | 390 | 0.01 | | | |
| 185 | 389 | 0.01 | A0A1Q3M3N2 | *Bacteroidales bacterium* | Uncharacterized protein (Fragment) |
| 186 | 389 | 0.01 | A0A1Q3SAM2 | *Flavobacterium sp.* | Uncharacterized protein |
| 187 | 388 | 0.01 | A0A1E4AAS0 | *Chryseobacterium sp.* | Uncharacterized protein |
| 188 | 387 | 0.01 | A0A0Q7ST55 | *Pelomonas sp.* | Guanosine-3',5'-bis(Diphosphate) 3'-pyrophosphohydrolase |
| 189 | 387 | 0.01 | A0A1Q3RXG1 | *Flavobacterium sp.* | Uncharacterized protein |
| 190 | 386 | 0.01 | A0A0Q7SWA7 | *Pelomonas sp.* | Aldehyde oxidase |
| 191 | 386 | 0.01 | A0A0Q8H4A2 | *Pelomonas sp.* | Uncharacterized protein |
| 192 | 386 | 0.01 | A0A1M3DNL6 | *Bacteroidales bacterium* | IS4 family transposase |
| 193 | 384 | 0.01 | A0A0Q7T0L4 | *Pelomonas sp.* | Primosomal protein N' (EC 3.6.4.-) (ATP-dependent helicase PriA) |
| 194 | 378 | 0.01 | F3ZQL8 | *Bacteroides coprosuis* | Transposase IS4 family protein |
| 195 | 378 | 0.01 | | | |
| 196 | 377 | 0.01 | A0A0Q6UGX4 | *Pelomonas sp.* | Nitrate reductase |
| 197 | 377 | 0.01 | A0A1E4AFY9 | *Chryseobacterium sp.* | Uncharacterized protein |
| 198 | 376 | 0.01 | A0A0Q6WDL9 | *Pelomonas sp.* | Serine/threonine protein kinase |
| 199 | 375 | 0.01 | A0A1B2YV26 | *uncultured bacterium* | Conserved repeat domain-containing protein |

| | | | | | |
|---|---|---|---|---|---|
| 200 | 374 | 0.01 | A0A0Q6X783 | *Pelomonas sp.* | Peptidase M13 |
| 201 | 374 | 0.01 | A0A0Q7SH01 | *Pelomonas sp.* | Lysine decarboxylase |
| 202 | 374 | 0.01 | A0A1Q3S3T4 | *Flavobacterium sp.* | TonB-dependent receptor |
| 203 | 372 | 0.01 | A0A077XLZ7 | *Sphinx1.76-related DNA* | Replication protein |
| 204 | 372 | 0.01 | A0A0Q7S0S4 | *Pelomonas sp.* | GTP pyrophosphokinase |
| 205 | 372 | 0.01 | A0A0Q8H074 | *Pelomonas sp.* | Uncharacterized protein |
| 206 | 369 | 0.01 | A0A1Q3RWE1 | *Flavobacterium sp.* | Penicillin-binding protein |
| 207 | 368 | 0.01 | A0A1Q3S2W6 | *Flavobacterium sp.* | Cytochrome C biogenesis protein |
| 208 | 368 | 0.01 | | | |
| 209 | 367 | 0.01 | A0A0Q7T112 | *Pelomonas sp.* | ATP-dependent DNA helicase Rep (EC 3.6.4.12) |
| 210 | 367 | 0.01 | A0A1E3ZYA8 | *Chryseobacterium sp.* | Uncharacterized protein |
| 211 | 367 | 0.01 | A0A1E4A2A9 | *Chryseobacterium sp.* | Organic solvent tolerance protein OstA |
| 212 | 367 | 0.01 | A0A1E4A7M8 | *Chryseobacterium sp.* | Uncharacterized protein |
| 213 | 366 | 0.01 | A0A0Q6T0C3 | *Pelomonas sp.* | DNA repair protein RecN (Recombination protein N) |
| 214 | 365 | 0.01 | A0A016UCK1 | *Ancylostoma ceylanicum* | Uncharacterized protein |
| 215 | 365 | 0.01 | A0A0Q6UIQ9 | *Pelomonas sp.* | Histidine kinase (EC 2.7.13.3) |
| 216 | 365 | 0.01 | A0A0Q7S7L6 | *Pelomonas sp.* | Protease/lipase ABC transporter permease/ATP-binding protein |
| 217 | 365 | 0.01 | A0A1Q3RXE2 | *Flavobacterium sp.* | Uncharacterized protein |
| 218 | 363 | 0.01 | | | |
| 219 | 363 | 0.01 | | | |
| 220 | 362 | 0.01 | A0A0Q7SKP8 | *Pelomonas sp.* | Outer membrane protein assembly factor BamA |
| 221 | 362 | 0.01 | A0A0Q7SQT9 | *Pelomonas sp.* | Chemotaxis protein |
| 222 | 362 | 0.01 | | | |
| 223 | 361 | 0.01 | A0A0Q7SW91 | *Pelomonas sp.* | Helicase |
| 224 | 361 | 0.01 | A0A0Q7T0C2 | *Pelomonas sp.* | Cell division protein |
| 225 | 361 | 0.01 | A0A177LPI8 | *Chryseobacterium sp.* | Uncharacterized protein |
| 226 | 361 | 0.01 | A0A1Q3S2Z4 | *Flavobacterium sp.* | Aminopeptidase |
| 227 | 361 | 0.01 | | | |
| 228 | 359 | 0.01 | A0A0Q7SYC6 | *Pelomonas sp.* | Chaperone protein ClpB |
| 229 | 358 | 0.01 | A0A0Q7S820 | *Pelomonas sp.* | Asparagine synthase |
| 230 | 356 | 0.01 | A0A1Q3S9B3 | *Flavobacterium sp.* | Uncharacterized protein |
| 231 | 355 | 0.01 | A0A0Q6THH1 | *Pelomonas sp.* | Uncharacterized protein |
| 232 | 355 | 0.01 | A0A0Q7S1U2 | *Pelomonas sp.* | Potassium transporter |
| 233 | 355 | 0.01 | A0A1Q3S511 | *Flavobacterium sp.* | Heat-shock protein Hsp70 |
| 234 | 354 | 0.01 | A0A1Q3S549 | *Flavobacterium sp.* | Phenylalanine--tRNA ligase beta subunit (EC 6.1.1.20) (Phenylalanyl-tRNA synthetase beta subunit) (PheRS) |

| | | | | | |
|---|---|---|---|---|---|
| 235 | 354 | 0.01 | | | |
| 236 | 353 | 0.01 | A0A127MVT9 | *Pseudomonas citronellolis* | Uncharacterized protein |
| 237 | 353 | 0.01 | A0A1Q3S8S4 | *Flavobacterium sp.* | Sodium:proline symporter |
| 238 | 352 | 0.01 | A0A0Q7S4H3 | *Pelomonas sp.* | Carbon starvation protein A |
| 239 | 352 | 0.01 | A0A0Q8GZT2 | *Pelomonas sp.* | Penicillin-binding protein |
| 240 | 352 | 0.01 | A0A1E3ZRN6 | *Chryseobacterium sp.* | Uncharacterized protein |
| 241 | 352 | 0.01 | A0A1Q3S9F5 | *Flavobacterium sp.* | DNA mismatch repair protein MutS |
| 242 | 351 | 0.01 | A0A1E3ZQQ1 | *Chryseobacterium sp.* | Uncharacterized protein |
| 243 | 351 | 0.01 | N8YF58 | *Acinetobacter gerneri* | Uncharacterized protein |
| 244 | 350 | 0.01 | A0A0Q7SU04 | *Pelomonas sp.* | K(+)-insensitive pyrophosphate-energized proton pump (EC 3.6.1.1) (Membrane-bound proton-translocating pyrophosphatase) (Pyrophosphate-energized inorganic pyrophosphatase) (H(+)-PPase) |
| 245 | 349 | 0.01 | A0A0Q6WQM3 | *Pelomonas sp.* | Carbon monoxide dehydrogenase |
| 246 | 349 | 0.01 | A0A0Q7T0F9 | *Pelomonas sp.* | Lon protease (EC 3.4.21.53) (ATP-dependent protease La) |
| 247 | 349 | 0.01 | A0A0R0LL75 | *Ferrovum sp.* | N-6 DNA methylase |
| 248 | 348 | 0.01 | A0A0Q7T6T8 | *Pelomonas sp.* | Histidine kinase |
| 249 | 347 | 0.01 | A0A1E3ZTP9 | *Chryseobacterium sp.* | Hybrid sensor histidine kinase/response regulator |
| 250 | 345 | 0.01 | A0A1E4ACQ6 | *Chryseobacterium sp.* | LmbE family protein |
| 251 | 345 | 0.01 | A0A1J5HM76 | *Porphyromonadaceae bacterium* | Multidrug transporter AcrB |
| 252 | 344 | 0.01 | A0A0K0WVI4 | *Litoditis aff. marina PmIII* | NADH dehydrogenase subunit 5 |
| 253 | 344 | 0.01 | A0A0Q7S0Z7 | *Pelomonas sp.* | RNA-binding transcriptional accessory protein |
| 254 | 344 | 0.01 | A0A0Q7SHH7 | *Pelomonas sp.* | Aldehyde dehydrogenase |
| 255 | 342 | 0.01 | A0A0Q7S5A2 | *Pelomonas sp.* | Glutamate-1-semialdehyde 2,1-aminomutase |
| 256 | 342 | 0.01 | A0A0Q7T2M8 | *Pelomonas sp.* | Uncharacterized protein |
| 257 | 342 | 0.01 | A0A1E3ZPZ7 | *Chryseobacterium sp.* | Peptidase S46 |
| 258 | 341 | 0.01 | A0A0Q6UF08 | *Pelomonas sp.* | NnrS family protein |
| 259 | 340 | 0.01 | A0A0Q6WL84 | *Pelomonas sp.* | Uncharacterized protein |
| 260 | 340 | 0.01 | A0A0Q6X5F3 | *Pelomonas sp.* | Alpha-2-macroglobulin |
| 261 | 340 | 0.01 | A0A0Q7RYJ8 | *Pelomonas sp.* | 5-methyltetrahydropteroyltriglutamate--homocysteine methyltransferase (EC 2.1.1.14) (Cobalamin-independent methionine synthase) (Methionine synthase, vitamin-B12 independent isozyme) |
| 262 | 340 | 0.01 | A0A0Q7SFG6 | *Pelomonas sp.* | Chaperone protein HtpG (Heat shock protein HtpG) (High temperature protein G) |
| 263 | 340 | 0.01 | A0A0Q7SP22 | *Pelomonas sp.* | UPF0313 protein ASD35_21020 |
| 264 | 339 | 0.01 | A0A1E4AD61 | *Chryseobacterium sp.* | Uncharacterized protein |
| 265 | 339 | 0.01 | A0A1Q3S9B2 | *Flavobacterium sp.* | Uncharacterized protein |

| | | | | | |
|---|---|---|---|---|---|
| 266 | 339 | 0.01 | | | |
| 267 | 336 | 0.01 | A0A0Q7SSA6 | *Pelomonas sp.* | CbbBc protein |
| 268 | 335 | 0.01 | S2WV93 | *Delftia acidovorans* | Uncharacterized protein |
| 269 | 334 | 0.01 | A0A0Q6T720 | *Pelomonas sp.* | DNA gyrase subunit A (EC 5.99.1.3) |
| 270 | 334 | 0.01 | A0A0Q6TI43 | *Pelomonas sp.* | Elongation factor G |
| 271 | 334 | 0.01 | A0A0Q8H948 | *Pelomonas sp.* | Tat pathway signal sequence domain protein |
| 272 | 334 | 0.01 | A0A1Q3S143 | *Flavobacterium sp.* | Glycine dehydrogenase (decarboxylating) (EC 1.4.4.2) (Glycine cleavage system P-protein) (Glycine decarboxylase) (Glycine dehydrogenase (aminomethyl-transferring)) |
| 273 | 333 | 0.01 | A0A0Q7S3D7 | *Pelomonas sp.* | ABC transporter |
| 274 | 333 | 0.01 | A0A0Q7SBS2 | *Pelomonas sp.* | Uncharacterized protein |
| 275 | 333 | 0.01 | A0A0Q7SGC9 | *Pelomonas sp.* | 3-hydroxyacyl-CoA dehydrogenase |
| 276 | 333 | 0.01 | A0A0Q8GG85 | *Pelomonas sp.* | Amidohydrolase |
| 277 | 332 | 0.01 | A0A0M9VJW9 | *Flavobacterium akiainvivens* | Transposase |
| 278 | 332 | 0.01 | A0A0Q6U057 | *Pelomonas sp.* | Multidrug transporter AcrB |
| 279 | 332 | 0.01 | A0A0Q8H0M9 | *Pelomonas sp.* | Uncharacterized protein |
| 280 | 332 | 0.01 | A0A180EWA0 | *Bacteroidales bacterium* | Replication protein |
| 281 | 331 | 0.01 | A0A1E4AAQ4 | *Chryseobacterium sp.* | Copper oxidase |
| 282 | 330 | 0.01 | A0A0Q6WEF8 | *Pelomonas sp.* | Uncharacterized protein |
| 283 | 329 | 0.01 | A0A0Q8GWX9 | *Pelomonas sp.* | TonB-dependent receptor |
| 284 | 329 | 0.01 | A0A174XH04 | *Bacteroides caccae* | Addiction module toxin, Txe/YoeB family |
| 285 | 329 | 0.01 | A0A1E4ADZ5 | *Chryseobacterium sp.* | ABC transporter substrate-binding protein |
| 286 | 329 | 0.01 | E4T5D5 | *Paludibacter propionicigenes* | DNA-directed DNA polymerase (EC 2.7.7.7) |
| 287 | 328 | 0.01 | A0A0Q6T9Y1 | *Pelomonas sp.* | Uncharacterized protein |
| 288 | 328 | 0.01 | A0A0Q7SNN2 | *Pelomonas sp.* | Chemotaxis protein |
| 289 | 328 | 0.01 | A0A0Q8GYI4 | *Pelomonas sp.* | Uncharacterized protein |
| 290 | 328 | 0.01 | A0A1Q3RVH4 | *Flavobacterium sp.* | Metallophosphoesterase |
| 291 | 328 | 0.01 | A0A1Q3S878 | *Flavobacterium sp.* | Patatin |
| 292 | 327 | 0.01 | A1W6Q0 | *Acidovorax sp.* | Integrase, catalytic region |
| 293 | 326 | 0.01 | A0A0Q6TWC3 | *Pelomonas sp.* | Uncharacterized protein |
| 294 | 325 | 0.01 | A0A0Q6TCI2 | *Pelomonas sp.* | Uncharacterized protein |
| 295 | 324 | 0.01 | A0A0Q7SCB4 | *Pelomonas sp.* | tRNA uridine 5-carboxymethylaminomethyl modification enzyme MnmG (Glucose-inhibited division protein A) |
| 296 | 324 | 0.01 | A0A0Q7SF49 | *Pelomonas sp.* | Phosphogluconate dehydratase |
| 297 | 324 | 0.01 | A0A0Q7SQV3 | *Pelomonas sp.* | Radical SAM protein |
| 298 | 324 | 0.01 | A0A0Q7SWN5 | *Pelomonas sp.* | Dihydrolipoyl dehydrogenase (EC 1.8.1.4) |
| 299 | 323 | 0.01 | A0A0Q6WW26 | *Pelomonas sp.* | Uncharacterized protein |

| | | | | | |
|---|---|---|---|---|---|
| 300 | 322 | 0.01 | A0A0Q6UDH8 | *Pelomonas sp.* | Tricorn protease homolog (EC 3.4.21.-) |
| 301 | 322 | 0.01 | A0A1E3ZY95 | *Chryseobacterium sp.* | Uncharacterized protein |
| 302 | 322 | 0.01 | A0A1Q3S7K6 | *Flavobacterium sp.* | Uncharacterized protein |
| 303 | 321 | 0.01 | A0A0Q7SBP1 | *Pelomonas sp.* | Histidine kinase |
| 304 | 320 | 0.01 | A0A0Q7SSL2 | *Pelomonas sp.* | Glycerol acyltransferase |
| 305 | 320 | 0.01 | A0A0Q7T1G9 | *Pelomonas sp.* | Transketolase (EC 2.2.1.1) |
| 306 | 320 | 0.01 | A0A1E3ZX97 | *Chryseobacterium sp.* | Gliding motility protein |
| 307 | 320 | 0.01 | A0A1Q3S6Q0 | *Flavobacterium sp.* | DNA polymerase I (EC 2.7.7.7) |
| 308 | 319 | 0.01 | A0A0Q6W9P5 | *Pelomonas sp.* | Uncharacterized protein |
| 309 | 318 | 0.01 | A0A0Q6TGJ4 | *Pelomonas sp.* | LPS-assembly protein LptD |
| 310 | 318 | 0.01 | A0A1T5GFF7 | *Sphingobacterium nematocida* | Gliding motility-associated C-terminal domain-containing protein |
| 311 | 317 | 0.01 | A0A0Q6U1R3 | *Pelomonas sp.* | Alkaline phosphatase |
| 312 | 317 | 0.01 | A0A0Q7SCU6 | *Pelomonas sp.* | ATP-dependent DNA helicase RecG (EC 3.6.4.12) |
| 313 | 317 | 0.01 | A0A0Q7SDP1 | *Pelomonas sp.* | Cyanophycin synthetase |
| 314 | 317 | 0.01 | A0A1Q3RWQ7 | *Flavobacterium sp.* | Cation/H(+) antiporter |
| 315 | 316 | 0.01 | A0A0Q7SDI5 | *Pelomonas sp.* | Malic enzyme (EC 1.1.1.40) |
| 316 | 316 | 0.01 | A0A0Q7SFN6 | *Pelomonas sp.* | Type II secretion system protein E |
| 317 | 316 | 0.01 | A0A1Q3RZB2 | *Flavobacterium sp.* | Uncharacterized protein |
| 318 | 315 | 0.01 | A0A0Q7S434 | *Pelomonas sp.* | Multifunctional fusion protein [Includes: 3-phosphoshikimate 1-carboxyvinyltransferase (EC 2.5.1.19) (5-enolpyruvylshikimate-3-phosphate synthase) (EPSP synthase) (EPSPS); Cytidylate kinase (CK) (EC 2.7.4.25) (Cytidine monophosphate kinase) (CMP kinase)] |
| 319 | 315 | 0.01 | A0A1E3ZRQ0 | *Chryseobacterium sp.* | Uncharacterized protein |
| 320 | 314 | 0.01 | A0A0Q7SKK0 | *Pelomonas sp.* | Uncharacterized protein |
| 321 | 314 | 0.01 | A0A0Q7SKQ6 | *Pelomonas sp.* | ATP-dependent DNA helicase RecQ |
| 322 | 314 | 0.01 | A0A1E4A395 | *Chryseobacterium sp.* | Peptidase S46 |
| 323 | 313 | 0.01 | A0A1Q3RZD7 | *Flavobacterium sp.* | ATP-dependent helicase |
| 324 | 313 | 0.01 | A6GYU0 | *Flavobacterium psychrophilum* | DNA-directed RNA polymerase subunit beta (RNAP subunit beta) (EC 2.7.7.6) (RNA polymerase subunit beta) (Transcriptase subunit beta) |
| 325 | 313 | 0.01 | | | |
| 326 | 312 | 0.01 | K9L3G6 | *Magnusiomyces magnusii* | Cytochrome b |
| 327 | 311 | 0.01 | A0A0Q7T0V8 | *Pelomonas sp.* | Sodium:solute symporter |
| 328 | 311 | 0.01 | A0A0Q7T1K0 | *Pelomonas sp.* | Carboxypeptidase |
| 329 | 311 | 0.01 | A0A1E4ABE8 | *Chryseobacterium sp.* | Methionine synthase |
| 330 | 311 | 0.01 | A0A1Q3S6W5 | *Flavobacterium sp.* | Peptidase M16 |
| 331 | 310 | 0.01 | A0A0L8VCE1 | *Sunxiuqinia dokdonensis* | Multidrug transporter AcrB |

| | | | | | |
|---|---|---|---|---|---|
| 332 | 310 | 0.01 | A0A0P6IS02 | *Daphnia magna* | Uncharacterized protein |
| 333 | 310 | 0.01 | A0A1E4ABW2 | *Chryseobacterium sp.* | Peptidase M1 |
| 334 | 309 | 0.01 | A0A0Q6TH52 | *Pelomonas sp.* | Uncharacterized protein |
| 335 | 309 | 0.01 | A0A0Q6U105 | *Pelomonas sp.* | Uncharacterized protein |
| 336 | 309 | 0.01 | A0A0Q7SYJ9 | *Pelomonas sp.* | Dihydroxy-acid dehydratase (EC 4.2.1.9) |
| 337 | 309 | 0.01 | A0A0Q7T6K8 | *Pelomonas sp.* | ABC transporter ATP-binding protein |
| 338 | 309 | 0.01 | A0A0Q8GEG5 | *Pelomonas sp.* | Uncharacterized protein |
| 339 | 308 | 0.01 | A0A1Q3RT16 | *Flavobacterium sp.* | S9 family peptidase |
| 340 | 308 | 0.01 | E4T072 | *Paludibacter propionicigenes* | Pyruvate, phosphate dikinase (EC 2.7.9.1) |
| 341 | 307 | 0.01 | A0A0Q7SMZ0 | *Pelomonas sp.* | TonB-dependent receptor |
| 342 | 307 | 0.01 | A0A0Q7ST48 | *Pelomonas sp.* | ABC transporter |
| 343 | 307 | 0.01 | A0A0Q7T1B8 | *Pelomonas sp.* | Glucose-6-phosphate isomerase (GPI) (EC 5.3.1.9) (Phosphoglucose isomerase) (PGI) (Phosphohexose isomerase) (PHI) |
| 344 | 307 | 0.01 | A0A1E4AC98 | *Chryseobacterium sp.* | Probable potassium transport system protein kup |
| 345 | 306 | 0.01 | A0A0Q7RWI0 | *Pelomonas sp.* | Glucokinase (EC 2.7.1.2) (Glucose kinase) |
| 346 | 306 | 0.01 | A0A0Q7SCJ9 | *Pelomonas sp.* | Heme ABC transporter ATP-binding protein |
| 347 | 306 | 0.01 | A0A1E4A600 | *Chryseobacterium sp.* | Recombinase |
| 348 | 306 | 0.01 | A0A1Q3LCT2 | *Bacteroidales bacterium* | Cell filamentation protein Fic |
| 349 | 306 | 0.01 | A0A1Q3SAJ2 | *Flavobacterium sp.* | Tail-specific protease |
| 350 | 305 | 0.01 | A0A0Q8GEL5 | *Pelomonas sp.* | Uncharacterized protein |
| 351 | 305 | 0.01 | A0A1Q3S7E9 | *Flavobacterium sp.* | Formate acetyltransferase |
| 352 | 304 | 0.01 | A0A0Q7RZY0 | *Pelomonas sp.* | DNA topoisomerase 4 subunit B (EC 5.99.1.3) (Topoisomerase IV subunit B) |
| 353 | 304 | 0.01 | A0A1Q3S1W4 | *Flavobacterium sp.* | Sulfatase |
| 354 | 303 | 0.01 | A0A0Q7SB38 | *Pelomonas sp.* | NADH-quinone oxidoreductase (EC 1.6.5.11) |
| 355 | 303 | 0.01 | A0A0Q7SBQ3 | *Pelomonas sp.* | Anthranilate synthase component 1 (EC 4.1.3.27) |
| 356 | 303 | 0.01 | A0A0Q7SJM3 | *Pelomonas sp.* | tRNA modification GTPase MnmE (EC 3.6.-.-) |
| 357 | 303 | 0.01 | A0A1E4AD98 | *Chryseobacterium sp.* | Ribonuclease R (RNase R) (EC 3.1.13.1) |
| 358 | 302 | 0.01 | H8KUZ8 | *Solitalea canadensis* | Uncharacterized protein |
| 359 | 301 | 0.01 | A0A0Q7S7F4 | *Pelomonas sp.* | Phosphoenolpyruvate-protein phosphotransferase (EC 2.7.3.9) (Phosphotransferase system, enzyme I) |
| 360 | 301 | 0.01 | D8L7S1 | *Paramecium caudatum* | NADH-ubiquinone oxidoreductase chain 5 (EC 1.6.5.3) |
| 361 | 300 | 0.01 | A0A0Q7S0T9 | *Pelomonas sp.* | MFS transporter |
| 362 | 300 | 0.01 | A0A0Q7S6Y8 | *Pelomonas sp.* | Uncharacterized protein |
| 363 | 300 | 0.01 | A0A1Q3SAW0 | *Flavobacterium sp.* | Sugar hydrolase |
| 364 | 300 | 0.01 | I9R3U4 | *Bacteroides cellulosilyticus* | Uncharacterized protein |
| 365 | 300 | 0.01 | | | |

| | | | | | |
|---|---|---|---|---|---|
| 366 | 299 | 0.01 | A0A023UM92 | *Magnusiomyces ingens* | Uncharacterized protein |
| 367 | 299 | 0.01 | A0A0Q7SKQ9 | *Pelomonas sp.* | L-aspartate oxidase (EC 1.4.3.16) |
| 368 | 299 | 0.01 | A0A1Q3SAL9 | *Flavobacterium sp.* | Peptidase M3 |
| 369 | 299 | 0.01 | | | |
| 370 | 298 | 0.01 | A0A0D6TL26 | *Flavobacterium sp.* | Metalloprotease |
| 371 | 298 | 0.01 | A0A0Q7SZN7 | *Pelomonas sp.* | Urease subunit alpha (EC 3.5.1.5) (Urea amidohydrolase subunit alpha) |
| 372 | 298 | 0.01 | A0A0Q8G4H5 | *Pelomonas sp.* | Elongation factor Tu (EF-Tu) |
| 373 | 298 | 0.01 | | | |
| 374 | 297 | 0.01 | A0A0Q7SKW7 | *Pelomonas sp.* | Long-chain fatty acid--CoA ligase |
| 375 | 297 | 0.01 | A0A1E4A183 | *Chryseobacterium sp.* | Tyrosine protein kinase |
| 376 | 297 | 0.01 | A0A1E4ACZ5 | *Chryseobacterium sp.* | TonB-dependent receptor |
| 377 | 296 | 0.01 | A0A0Q6THL3 | *Pelomonas sp.* | Uncharacterized protein |
| 378 | 296 | 0.01 | A0A0Q6TIL7 | *Pelomonas sp.* | Uncharacterized protein |
| 379 | 296 | 0.01 | A0A0Q6UFQ4 | *Pelomonas sp.* | Dihydroxy-acid dehydratase (EC 4.2.1.9) |
| 380 | 296 | 0.01 | A0A0Q7RWA8 | *Pelomonas sp.* | DNA mismatch repair protein MutL |
| 381 | 296 | 0.01 | A0A0Q7S6B9 | *Pelomonas sp.* | Uncharacterized protein |
| 382 | 296 | 0.01 | A0A0Q7SVE2 | *Pelomonas sp.* | Metal ABC transporter permease |
| 383 | 296 | 0.01 | A0A0Q8GMN6 | *Pelomonas sp.* | Uncharacterized protein |
| 384 | 295 | 0.01 | A0A0Q6U486 | *Pelomonas sp.* | Glucose-methanol-choline oxidoreductase |
| 385 | 295 | 0.01 | A0A1Q3S1U2 | *Flavobacterium sp.* | Phosphoribosylformylglycinamidine synthase (FGAM synthase) (FGAMS) (EC 6.3.5.3) (Formylglycinamide ribonucleotide amidotransferase) (FGAR amidotransferase) (FGAR-AT) |
| 386 | 294 | 0.01 | A0A0Q6SWR0 | *Pelomonas sp.* | Uncharacterized protein |
| 387 | 294 | 0.01 | A0A0Q6TDG6 | *Pelomonas sp.* | Uncharacterized protein |
| 388 | 294 | 0.01 | A0A0Q7SWY9 | *Pelomonas sp.* | Uncharacterized protein |
| 389 | 294 | 0.01 | A0A0Q7SXS7 | *Pelomonas sp.* | Uncharacterized protein |
| 390 | 294 | 0.01 | A0A0Q7SZT2 | *Pelomonas sp.* | Type IV-A pilus assembly ATPase PilB |
| 391 | 294 | 0.01 | A0A1Q3M3K5 | *Bacteroidales bacterium* | Alpha-mannosidase |
| 392 | 294 | 0.01 | A0A1Q3S478 | *Flavobacterium sp.* | Conjugal transfer protein TraG |
| 393 | 293 | 0.01 | A0A0Q6T928 | *Pelomonas sp.* | Beta-galactosidase |
| 394 | 293 | 0.01 | A0A0Q7S098 | *Pelomonas sp.* | Transcription termination factor Rho (EC 3.6.4.-) (ATP-dependent helicase Rho) |
| 395 | 293 | 0.01 | A0A0Q7SF85 | *Pelomonas sp.* | Aldehyde dehydrogenase |
| 396 | 293 | 0.01 | A0A0Q7SL43 | *Pelomonas sp.* | Polyphosphate kinase (EC 2.7.4.1) (ATP-polyphosphate phosphotransferase) (Polyphosphoric acid kinase) |
| 397 | 292 | 0.01 | A0A023UPJ4 | *Magnusiomyces ingens* | Uncharacterized protein (Fragment) |
| 398 | 292 | 0.01 | A0A0Q6TH04 | *Pelomonas sp.* | Uncharacterized protein |
| 399 | 292 | 0.01 | A0A0Q7SJQ2 | *Pelomonas sp.* | Dihydroxy-acid dehydratase (EC 4.2.1.9) |

| | | | | | |
|---|---|---|---|---|---|
| 400 | 292 | 0.01 | A0A0Q8G5N2 | *Pelomonas sp.* | Uncharacterized protein |
| 401 | 292 | 0.01 | A0A1M3DC14 | *Bacteroidales bacterium* | Formate C-acetyltransferase |
| 402 | 292 | 0.01 | A0A1Q3RWZ9 | *Flavobacterium sp.* | Uncharacterized protein |
| 403 | 292 | 0.01 | A0A1Q3SB52 | *Flavobacterium sp.* | Acriflavin resistance protein |
| 404 | 291 | 0.01 | A0A0Q6UA04 | *Pelomonas sp.* | Uncharacterized protein |
| 405 | 291 | 0.01 | A0A0Q6XBI6 | *Pelomonas sp.* | Penicillin-binding protein 1C |
| 406 | 291 | 0.01 | | | |
| 407 | 291 | 0.01 | | | |
| 408 | 290 | 0.01 | A0A0Q6TBZ3 | *Pelomonas sp.* | Multidrug transporter |
| 409 | 290 | 0.01 | A0A1M3MBI7 | *Paludibacter sp.* | Cell surface protein SprA |
| 410 | 290 | 0.01 | | | |
| 411 | 289 | 0.01 | A0A0P0UB73 | *Flavobacterium psychrophilum* | Multidrug transporter AcrB |
| 412 | 289 | 0.01 | A0A0Q6TGN2 | *Pelomonas sp.* | DNA helicase (EC 3.6.4.12) |
| 413 | 289 | 0.01 | A0A1E4AGS9 | *Chryseobacterium sp.* | TonB-dependent receptor |
| 414 | 288 | 0.01 | A0A0Q7SWP5 | *Pelomonas sp.* | Xylulose kinase |
| 415 | 288 | 0.01 | A0A1Q3S4I8 | *Flavobacterium sp.* | Uncharacterized protein |
| 416 | 287 | 0.01 | A0A023UP41 | *Magnusiomyces ingens* | Cytochrome c oxidase subunit 1 (EC 1.9.3.1) |
| 417 | 287 | 0.01 | A0A0Q7SUD0 | *Pelomonas sp.* | AMP-binding protein |
| 418 | 287 | 0.01 | A0A1E4A8B6 | *Chryseobacterium sp.* | Type II and III secretion system protein |
| 419 | 287 | 0.01 | A0A1E4ACD4 | *Chryseobacterium sp.* | Gliding motility lipoprotein GldJ |
| 420 | 287 | 0.01 | A0A1Q3S8N5 | *Flavobacterium sp.* | Uncharacterized protein |
| 421 | 287 | 0.01 | A0A1Q3S9I3 | *Flavobacterium sp.* | Type IV secretion protein Rhs |
| 422 | 286 | 0.01 | A0A0Q6T4E7 | *Pelomonas sp.* | TonB-dependent receptor |
| 423 | 286 | 0.01 | A0A0Q7SDP6 | *Pelomonas sp.* | Uncharacterized protein |
| 424 | 286 | 0.01 | A0A0Q7SFQ5 | *Pelomonas sp.* | Uncharacterized protein |
| 425 | 286 | 0.01 | A0A0Q7SN74 | *Pelomonas sp.* | Bifunctional purine biosynthesis protein PurH [Includes: IMP cyclohydrolase (EC 3.5.4.10) (IMP synthase) (Inosinicase) (ATIC); Phosphoribosylaminoimidazolecarboxamide formyltransferase (EC 2.1.2.3) (AICAR transformylase)] |
| 426 | 286 | 0.01 | A0A0Q7SZX8 | *Pelomonas sp.* | Malate dehydrogenase |
| 427 | 286 | 0.01 | A0A0Q8GY27 | *Pelomonas sp.* | Uncharacterized protein |
| 428 | 286 | 0.01 | A0A1Q3S8C6 | *Flavobacterium sp.* | Uncharacterized protein |
| 429 | 286 | 0.01 | H8MU17 | *Corallococcus coralloides* | Long-chain-fatty-acid--CoA ligase |
| 430 | 285 | 0.01 | A0A0Q7SGQ0 | *Pelomonas sp.* | ATP-dependent RNA helicase |
| 431 | 285 | 0.01 | A0A0Q7T3S1 | *Pelomonas sp.* | Radical SAM protein |
| 432 | 285 | 0.01 | A0A1Q3S0W7 | *Flavobacterium sp.* | Uncharacterized protein |

| | | | | | |
|---|---|---|---|---|---|
| 433 | 284 | 0.01 | A0A0Q6UEX1 | *Pelomonas sp.* | Glutamine--tRNA ligase (EC 6.1.1.18) (Glutaminyl-tRNA synthetase) (GlnRS) |
| 434 | 284 | 0.01 | A0A0Q7SMM0 | *Pelomonas sp.* | Aldehyde dehydrogenase |
| 435 | 283 | 0.01 | A0A0Q6U912 | *Pelomonas sp.* | Uncharacterized protein |
| 436 | 283 | 0.01 | A0A1E4A7R6 | *Chryseobacterium sp.* | Uncharacterized protein |
| 437 | 283 | 0.01 | A0A1Q3RW11 | *Flavobacterium sp.* | Magnesium-translocating P-type ATPase |
| 438 | 283 | 0.01 | A0A1Q3S955 | *Flavobacterium sp.* | Polyphosphate kinase (EC 2.7.4.1) (ATP-polyphosphate phosphotransferase) (Polyphosphoric acid kinase) |
| 439 | 282 | 0.01 | A0A023UM78 | *Magnusiomyces ingens* | NADH dehydrogenase subunit 2 (EC 1.6.5.3) |
| 440 | 282 | 0.01 | A0A0Q6TEV6 | *Pelomonas sp.* | Multidrug transporter |
| 441 | 282 | 0.01 | A0A0Q8GXY5 | *Pelomonas sp.* | Uncharacterized protein |
| 442 | 280 | 0.01 | A0A0Q7SF22 | *Pelomonas sp.* | Amidotransferase |
| 443 | 280 | 0.01 | A0A0Q8GHK7 | *Pelomonas sp.* | Uncharacterized protein |
| 444 | 280 | 0.01 | A0A0Q8GZU0 | *Pelomonas sp.* | Uncharacterized protein |
| 445 | 279 | 0.01 | A0A0Q6TJ53 | *Pelomonas sp.* | Uncharacterized protein |
| 446 | 279 | 0.01 | A0A0Q6U119 | *Pelomonas sp.* | Histidine kinase (EC 2.7.13.3) |
| 447 | 279 | 0.01 | A0A0Q7SSI3 | *Pelomonas sp.* | Methylmalonate-semialdehyde dehydrogenase |
| 448 | 279 | 0.01 | A0A0Q7T0Q3 | *Pelomonas sp.* | Alkyl hydroperoxide reductase subunit F |
| 449 | 279 | 0.01 | A0A1Q3S8T7 | *Flavobacterium sp.* | Nucleotidyltransferase |
| 450 | 278 | 0.01 | A0A0Q7SFI7 | *Pelomonas sp.* | Multidrug transporter |
| 451 | 278 | 0.01 | A0A1Q3RX80 | *Flavobacterium sp.* | Uncharacterized protein |
| 452 | 277 | 0.01 | A0A0Q7S282 | *Pelomonas sp.* | UvrABC system protein B (Protein UvrB) (Excinuclease ABC subunit B) |
| 453 | 277 | 0.01 | A0A1E3ZYM1 | *Chryseobacterium sp.* | Peptidylprolyl isomerase (EC 5.2.1.8) |
| 454 | 277 | 0.01 | A0A1Q3RY34 | *Flavobacterium sp.* | TonB-dependent receptor |
| 455 | 277 | 0.01 | A1W5C2 | *Acidovorax sp.* | IstB domain protein ATP-binding protein |
| 456 | 276 | 0.01 | A0A0Q7SGT2 | *Pelomonas sp.* | Gamma-glutamyltransferase |
| 457 | 275 | 0.01 | A0A0Q7SLU5 | *Pelomonas sp.* | Glutamate synthase (EC 1.4.1.13) |
| 458 | 275 | 0.01 | A0A140JXE2 | *Escherichia coli* | Uncharacterized protein |
| 459 | 275 | 0.01 | A0A1E4AEN9 | *Chryseobacterium sp.* | Peptidase S9 |
| 460 | 274 | 0.01 | A0A0Q6TC22 | *Pelomonas sp.* | Uncharacterized protein |
| 461 | 274 | 0.01 | A0A0Q7S1G7 | *Pelomonas sp.* | Thiamine pyrophosphate-binding protein |
| 462 | 274 | 0.01 | A0A1Q3S421 | *Flavobacterium sp.* | Uncharacterized protein (Fragment) |
| 463 | 273 | 0.01 | A0A077XLZ0 | *Sphinx1.76-related DNA* | Replication protein |
| 464 | 273 | 0.01 | A0A0Q7RU29 | *Pelomonas sp.* | Isoleucine--tRNA ligase (EC 6.1.1.5) (Isoleucyl-tRNA synthetase) (IleRS) |
| 465 | 273 | 0.01 | A0A0Q7SDH4 | *Pelomonas sp.* | Long-chain fatty acid--CoA ligase |
| 466 | 273 | 0.01 | A0A0Q7SQJ3 | *Pelomonas sp.* | Molybdopterin oxidoreductase |
| 467 | 272 | 0.01 | A0A0Q6THN9 | *Pelomonas sp.* | Uncharacterized protein |

| | | | | | |
|---|---|---|---|---|---|
| 468 | 272 | 0.01 | A0A0Q7S074 | *Pelomonas sp.* | Thymidine phosphorylase (EC 2.4.2.4) (TdRPase) |
| 469 | 272 | 0.01 | A0A0Q7SPJ8 | *Pelomonas sp.* | ABC transporter permease |
| 470 | 272 | 0.01 | A0A0Q7ST88 | *Pelomonas sp.* | Peptidase |
| 471 | 272 | 0.01 | A0A1E4A2H0 | *Chryseobacterium sp.* | Peptidylprolyl isomerase |
| 472 | 272 | 0.01 | A0A1E4A473 | *Chryseobacterium sp.* | Penicillin-binding protein 2 |
| 473 | 272 | 0.01 | A0A1Q3S481 | *Flavobacterium sp.* | Uncharacterized protein |
| 474 | 272 | 0.01 | | | |
| 475 | 271 | 0.01 | A0A0Q7S368 | *Pelomonas sp.* | MSHA biogenesis protein MshE |
| 476 | 271 | 0.01 | A0A0Q7SZ55 | *Pelomonas sp.* | 1-deoxy-D-xylulose-5-phosphate synthase (EC 2.2.1.7) (1-deoxyxylulose-5-phosphate synthase) (DXP synthase) (DXPS) |
| 477 | 271 | 0.01 | A0A1Q3RZ46 | *Flavobacterium sp.* | Uncharacterized protein |
| 478 | 271 | 0.01 | A0A1Q3SA66 | *Flavobacterium sp.* | Polyribonucleotide nucleotidyltransferase (EC 2.7.7.8) (Polynucleotide phosphorylase) (PNPase) |
| 479 | 271 | 0.01 | E4T2R0 | *Paludibacter propionicigenes* | ATPase AAA-2 domain protein |
| 480 | 270 | 0.01 | A0A0Q6TNS7 | *Pelomonas sp.* | Fis family transcriptional regulator |
| 481 | 270 | 0.01 | A0A0Q6WLQ9 | *Pelomonas sp.* | Cell division protein FtsZ |
| 482 | 270 | 0.01 | A0A0Q7S5E8 | *Pelomonas sp.* | Cysteine desulfurase |
| 483 | 270 | 0.01 | A0A0Q7SSA4 | *Pelomonas sp.* | PAS domain-containing sensor histidine kinase |
| 484 | 270 | 0.01 | A0A1Q3M5E8 | *Bacteroidales bacterium* | Uncharacterized protein (Fragment) |
| 485 | 270 | 0.01 | A0A1Q3M633 | *Bacteroidales bacterium* | ATP-dependent helicase |
| 486 | 270 | 0.01 | A0A1Q3RY20 | *Flavobacterium sp.* | Collagen-binding protein |
| 487 | 270 | 0.01 | A0A1Q3S3S5 | *Flavobacterium sp.* | Uncharacterized protein |
| 488 | 269 | 0.01 | A0A0Q7S8W9 | *Pelomonas sp.* | Peptidase M3 |
| 489 | 269 | 0.01 | A0A0Q7SK57 | *Pelomonas sp.* | 2-hydroxy-acid oxidase |
| 490 | 269 | 0.01 | A0A0Q7SQY5 | *Pelomonas sp.* | Uncharacterized protein |
| 491 | 269 | 0.01 | A0A0Q7SV39 | *Pelomonas sp.* | Acyl-CoA dehydrogenase |
| 492 | 269 | 0.01 | A0A0Q8H4B7 | *Pelomonas sp.* | Phenylacetic acid degradation protein |
| 493 | 268 | 0.01 | A0A0Q6T2K7 | *Pelomonas sp.* | 3-isopropylmalate dehydratase large subunit (EC 4.2.1.33) (Alpha-IPM isomerase) (IPMI) (Isopropylmalate isomerase) |
| 494 | 268 | 0.01 | A0A0Q7S8B3 | *Pelomonas sp.* | 23S rRNA (uracil(1939)-C(5))-methyltransferase RlmD (EC 2.1.1.190) (23S rRNA(m5U1939)-methyltransferase) |
| 495 | 268 | 0.01 | A0A0Q7SQ46 | *Pelomonas sp.* | Ferredoxin oxidoreductase |
| 496 | 268 | 0.01 | A0A0Q7SWZ8 | *Pelomonas sp.* | Ribonucleoside-diphosphate reductase (EC 1.17.4.1) |
| 497 | 268 | 0.01 | A0A1E4A3S3 | *Chryseobacterium sp.* | Glycosyl transferase |
| 498 | 268 | 0.01 | A0A1E4A9N2 | *Chryseobacterium sp.* | Uncharacterized protein |
| 499 | 268 | 0.01 | A0A1E4AGS6 | *Chryseobacterium sp.* | ATPase |
| 500 | 268 | 0.01 | A0A1I1TKT3 | *Pseudomonas citronellolis* | Uncharacterized protein |

**Table S6a.** Functional overview of M4A community (top 500 rank only).

| Rank | Count | Abundance | ID | Organism | Protein Names |
|---|---|---|---|---|---|
| 1 | 5032 | 0.17 | A0A0D5N8Q4 | *Geobacter sulfurreducens* | Transposase |
| 2 | 3549 | 0.12 | Q749Z8 | *Geobacter sulfurreducens* | Transposase of ISGsu6, IS21 family |
| 3 | 2853 | 0.10 | | | |
| 4 | 2145 | 0.07 | A0A1F9PUL2 | *Desulfuromonadales bacterium* | Restriction endonuclease |
| 5 | 2112 | 0.07 | Q749Z7 | *Geobacter sulfurreducens* | ATP/GTP-binding transposition helper of ISGsu6, IS21 family |
| 6 | 1828 | 0.06 | Q39Y12 | *Geobacter metallireducens* | DNA-directed RNA polymerase subunit beta' (RNAP subunit beta') (EC 2.7.7.6) (RNA polymerase subunit beta') (Transcriptase subunit beta') |
| 7 | 1611 | 0.06 | Q748Y6 | *Geobacter sulfurreducens* | DNA-directed RNA polymerase subunit beta (RNAP subunit beta) (EC 2.7.7.6) (RNA polymerase subunit beta) (Transcriptase subunit beta) |
| 8 | 1485 | 0.05 | A0A0D5N7W1 | *Geobacter sulfurreducens* | ATPase |
| 9 | 1402 | 0.05 | E8WQ80 | *Geobacter sp.* | Integrase catalytic region |
| 10 | 1390 | 0.05 | Q748X8 | *Geobacter sulfurreducens* | Elongation factor Tu (EF-Tu) |
| 11 | 1364 | 0.05 | J7LWF3 | *Geobacter metallireducens* | Transposase, IS256 family |
| 12 | 1289 | 0.04 | Q74AK2 | *Geobacter sulfurreducens* | Sodium/solute symporter family protein |
| 13 | 1266 | 0.04 | A0A0A8WNU3 | *Geobacter sp.* | Protein TraC |
| 14 | 1245 | 0.04 | Q39Q43 | *Geobacter metallireducens* | Pyruvate-flavodoxin oxidoreductase (EC 1.2.7.-) |
| 15 | 1182 | 0.04 | N8ZVC9 | *Acinetobacter gerneri* | Uncharacterized protein |
| 16 | 1180 | 0.04 | A0A0D5N8M0 | *Geobacter sulfurreducens* | MgtE integral membrane protein |
| 17 | 1165 | 0.04 | Q747C7 | *Geobacter sulfurreducens* | 60 kDa chaperonin (GroEL protein) (Protein Cpn60) |
| 18 | 1164 | 0.04 | A0A0L8ETQ7 | *Achromobacter sp.* | Uncharacterized protein |
| 19 | 1115 | 0.04 | Q39RM9 | *Geobacter metallireducens* | Membrane protein, putative |
| 20 | 1111 | 0.04 | | | |
| 21 | 1080 | 0.04 | E8WU24 | *Geobacter sp.* | Helicase domain protein |
| 22 | 1079 | 0.04 | A1AKK8 | *Pelobacter propionicus* | Uncharacterized protein |
| 23 | 1076 | 0.04 | Q748Y8 | *Geobacter sulfurreducens* | Elongation factor G 2 (EF-G 2) |
| 24 | 1073 | 0.04 | | | |
| 25 | 1072 | 0.04 | A0A1F9PUK4 | *Desulfuromonadales bacterium* | Methyltransferase (EC 2.1.1.-) |
| 26 | 1056 | 0.04 | Q39QB9 | *Geobacter metallireducens* | NADH dehydrogenase I, M subunit |
| 27 | 1025 | 0.04 | Q747X9 | *Geobacter sulfurreducens* | Isoleucine--tRNA ligase (EC 6.1.1.5) (Isoleucyl-tRNA synthetase) (IleRS) |
| 28 | 995 | 0.03 | A5GAB4 | *Geobacter uraniireducens* | Putative GAF sensor protein |
| 29 | 990 | 0.03 | Q39U60 | *Geobacter metallireducens* | Translation elongation factor G |
| 30 | 973 | 0.03 | Q39QB2 | *Geobacter metallireducens* | NADH dehydrogenase I, F subunit |
| 31 | 943 | 0.03 | Q39QB8 | *Geobacter metallireducens* | NADH dehydrogenase I, L subunit |
| 32 | 943 | 0.03 | | | |

| | | | | | |
|---|---|---|---|---|---|
| 33 | 934 | 0.03 | A0A0B4XMH5 | *Alcanivorax pacificus* | Adenine-specific DNA methylase containing a Zn-ribbon |
| 34 | 933 | 0.03 | Q39ZC8 | *Geobacter metallireducens* | Ferredoxin-dependent glutamate synthase |
| 35 | 928 | 0.03 | | | |
| 36 | 922 | 0.03 | A0A0B5BF68 | *Geobacter pickeringii* | CRISPR-associated protein Cas3 |
| 37 | 911 | 0.03 | I9R3U4 | *Bacteroides cellulosilyticus* | Uncharacterized protein |
| 38 | 908 | 0.03 | Q39YK5 | *Geobacter metallireducens* | Flagellar biogenesis protein FlhA |
| 39 | 907 | 0.03 | A0A1G0M9J2 | *Geobacteraceae bacterium* | Uncharacterized protein |
| 40 | 885 | 0.03 | Q74FK9 | *Geobacter sulfurreducens* | Sigma-54-dependent transcriptional response regulator |
| 41 | 884 | 0.03 | B3EB58 | *Geobacter lovleyi* | Uncharacterized protein |
| 42 | 879 | 0.03 | Q749A7 | *Geobacter sulfurreducens* | Protein translocase subunit SecY |
| 43 | 873 | 0.03 | A0A0B5BJ81 | *Geobacter pickeringii* | NADH-quinone oxidoreductase subunit D (EC 1.6.5.11) (NADH dehydrogenase I subunit D) (NDH-1 subunit D) |
| 44 | 853 | 0.03 | Q39ZK4 | *Geobacter metallireducens* | Bifunctional glutamine synthetase adenylyltransferase/adenylyl-removing enzyme (ATP:glutamine synthetase adenylyltransferase) (ATase) [Includes: Glutamine synthetase adenylyl-L-tyrosine phosphorylase (EC 2.7.7.89) (Adenylyl removase) (AR) (AT-N); Glutamine synthetase adenylyl transferase (EC 2.7.7.42) (Adenylyl transferase) (AT) (AT-C)] |
| 45 | 845 | 0.03 | A0A0B5BF21 | *Geobacter pickeringii* | Glycosyl transferase family 1 |
| 46 | 842 | 0.03 | Q39XL9 | *Geobacter metallireducens* | Sensor sigma-54-dependent transcriptional regulator, GAF and GAF domain-containing |
| 47 | 837 | 0.03 | N8ZNX3 | *Acinetobacter gerneri* | Uncharacterized protein |
| 48 | 823 | 0.03 | A0A4W3 | *Geobacter sulfurreducens* | Biosynthetic arginine decarboxylase (ADC) (EC 4.1.1.19) |
| 49 | 815 | 0.03 | S0G1I5 | *Desulfotignum phosphitoxidans* | Type I restriction enzyme HindVIIP/EcoKI protein HsdR (EC 3.1.21.3) |
| 50 | 809 | 0.03 | Q39T74 | *Geobacter metallireducens* | GTP/GDP 3'-pyrophosphokinase and (P)ppGpp 3'-pyrophosphohydrolase |
| 51 | 808 | 0.03 | Q74FM6 | *Geobacter sulfurreducens* | Pyruvate, phosphate dikinase (EC 2.7.9.1) |
| 52 | 796 | 0.03 | A0A0D5N448 | *Geobacter sulfurreducens* | Transposase |
| 53 | 794 | 0.03 | Q74FF1 | *Geobacter sulfurreducens* | Chaperone protein ClpB |
| 54 | 792 | 0.03 | Q39Y76 | *Geobacter metallireducens* | Sigma-54-dependent transcriptional response regulator |
| 55 | 790 | 0.03 | A0A1S0XXD8 | *Rhodanobacter sp.* | Histidine kinase |
| 56 | 790 | 0.03 | Q39V52 | *Geobacter metallireducens* | GAF sensor methyl-accepting chemotaxis sensory transducer, class 40H |
| 57 | 786 | 0.03 | A0A0B5BDJ5 | *Geobacter pickeringii* | NADH-quinone oxidoreductase subunit H (EC 1.6.5.11) (NADH dehydrogenase I subunit H) (NDH-1 subunit H) |
| 58 | 785 | 0.03 | Q39QJ8 | *Geobacter metallireducens* | Flagellar biogenesis master sigma-54-dependent transcriptional response regulator |
| 59 | 781 | 0.03 | A0A0S8I0Q8 | *bacterium SM23_31* | Uncharacterized protein |
| 60 | 775 | 0.03 | Q747H5 | *Geobacter sulfurreducens* | K(+)-insensitive pyrophosphate-energized proton pump (EC 3.6.1.1) (Membrane-bound proton-translocating pyrophosphatase) (Pyrophosphate-energized inorganic pyrophosphatase) (H(+)-PPase) |
| 61 | 768 | 0.03 | A0A0B5B8A8 | *Geobacter pickeringii* | Valine--tRNA ligase (EC 6.1.1.9) (Valyl-tRNA synthetase) (ValRS) |

| | | | | | |
|---|---|---|---|---|---|
| 62 | 765 | 0.03 | A0A0B5BCT5 | *Geobacter pickeringii* | NADH dehydrogenase |
| 63 | 756 | 0.03 | Q39UR9 | *Geobacter metallireducens* | Carbamoyl-phosphate synthase large chain (EC 6.3.5.5) (Carbamoyl-phosphate synthetase ammonia chain) |
| 64 | 756 | 0.03 | | | |
| 65 | 755 | 0.03 | | | |
| 66 | 739 | 0.03 | Q39PS2 | *Geobacter metallireducens* | Transcription-repair-coupling factor (TRCF) (EC 3.6.4.-) |
| 67 | 737 | 0.03 | Q39ZE7 | *Geobacter metallireducens* | UvrABC system protein A (UvrA protein) (Excinuclease ABC subunit A) |
| 68 | 736 | 0.03 | A0A0B5B7H3 | *Geobacter pickeringii* | Cytochrome C |
| 69 | 730 | 0.03 | Q74GY2 | *Geobacter sulfurreducens* | ATP synthase subunit alpha (EC 3.6.3.14) (ATP synthase F1 sector subunit alpha) (F-ATPase subunit alpha) |
| 70 | 729 | 0.03 | C6DZ95 | *Geobacter sp.* | Lytic transglycosylase catalytic |
| 71 | 726 | 0.03 | Q747M6 | *Geobacter sulfurreducens* | Ribonuclease G |
| 72 | 725 | 0.03 | A1AKN8 | *Pelobacter propionicus* | Resolvase, N-terminal domain protein |
| 73 | 722 | 0.02 | Q39U54 | *Geobacter metallireducens* | Sensor histidine kinase cyclic nucleotide phosphodiesterase, GAF, GAF and HD-GYP-related domain-containing |
| 74 | 719 | 0.02 | P14243 | *Citrobacter freundii* | Modification methylase Cfr9I (M.Cfr9I) (EC 2.1.1.113) (N(4)- cytosine-specific methyltransferase Cfr9I) |
| 75 | 718 | 0.02 | A0A0C5IZ87 | *Rugosibacter aromaticivorans* | Uncharacterized protein |
| 76 | 713 | 0.02 | A0A0D5ND20 | *Geobacter sulfurreducens* | Amino acid dehydrogenase |
| 77 | 712 | 0.02 | E1QK39 | *Desulfarculus baarsii* | Helicase domain protein |
| 78 | 704 | 0.02 | Q74C76 | *Geobacter sulfurreducens* | (R)-citramalate synthase (EC 2.3.1.182) (Citramalate synthase) |
| 79 | 703 | 0.02 | A0A0D5N8S3 | *Geobacter sulfurreducens* | NADH dehydrogenase |
| 80 | 700 | 0.02 | A0A0D5N8I5 | *Geobacter sulfurreducens* | Signal recognition particle protein (Fifty-four homolog) |
| 81 | 699 | 0.02 | Q74GY0 | *Geobacter sulfurreducens* | ATP synthase subunit beta (EC 3.6.3.14) (ATP synthase F1 sector subunit beta) (F-ATPase subunit beta) |
| 82 | 699 | 0.02 | Q74H58 | *Geobacter sulfurreducens* | Chaperone protein DnaJ |
| 83 | 696 | 0.02 | A0A0B5BH93 | *Geobacter pickeringii* | C-and b-type cytochrome L |
| 84 | 696 | 0.02 | Q39ST0 | *Geobacter metallireducens* | Fibronectin type III repeat, DUF2341 repeat and LamG-like domain protein |
| 85 | 695 | 0.02 | | | |
| 86 | 691 | 0.02 | Q746X3 | *Geobacter sulfurreducens* | Bifunctional protein PutA |
| 87 | 690 | 0.02 | | | |
| 88 | 687 | 0.02 | A0A0B5BJ46 | *Geobacter pickeringii* | Uncharacterized protein |
| 89 | 684 | 0.02 | A0A0B5BJ26 | *Geobacter pickeringii* | Uncharacterized protein |
| 90 | 684 | 0.02 | Q39WG2 | *Geobacter metallireducens* | Chromosome partition protein Smc |
| 91 | 681 | 0.02 | B4S9K8 | *Prosthecochloris aestuarii* | TRAG family protein |
| 92 | 681 | 0.02 | Q74AK3 | *Geobacter sulfurreducens* | Cation-translocating P-type ATPase |

| | | | | | |
|---|---|---|---|---|---|
| 93 | 672 | 0.02 | | | |
| 94 | 669 | 0.02 | Q39W73 | *Geobacter metallireducens* | 2-isopropylmalate synthase (EC 2.3.3.13) (Alpha-IPM synthase) (Alpha-isopropylmalate synthase) |
| 95 | 668 | 0.02 | Q39Q36 | *Geobacter metallireducens* | Heterodisulfide oxidoreductase, FAD-binding and iron-sulfur cluster-binding subunit A |
| 96 | 667 | 0.02 | Q39VR4 | *Geobacter metallireducens* | Bifunctional uridylyltransferase/uridylyl-removing enzyme (UTase/UR) (Bifunctional [protein-PII] modification enzyme) (Bifunctional nitrogen sensor protein) [Includes: [Protein-PII]-UMP uridylyl-removing enzyme (UR) (EC 3.1.4.-); [Protein-PII] uridylyltransferase (PII uridylyltransferase) (UTase) (EC 2.7.7.59)] |
| 97 | 666 | 0.02 | A0A0B5BB65 | *Geobacter pickeringii* | Uncharacterized protein |
| 98 | 664 | 0.02 | A0A0R3QEE4 | *Brugia timori* | Uncharacterized protein |
| 99 | 663 | 0.02 | A0A0D5NDD6 | *Geobacter sulfurreducens* | Type II secretion system protein E |
| 100 | 661 | 0.02 | Q39X91 | *Geobacter metallireducens* | DNA topoisomerase 1 (EC 5.99.1.2) (DNA topoisomerase I) |
| 101 | 660 | 0.02 | A0LQ17 | *Syntrophobacter fumaroxidans* | Helicase domain protein |
| 102 | 658 | 0.02 | Q74FW2 | *Geobacter sulfurreducens* | Succinyl:acetate coenzyme A transferase |
| 103 | 657 | 0.02 | Q74H82 | *Geobacter sulfurreducens* | Flagellar biogenesis master response receiver sensor histidine kinase, PAS and GAF domain-containing |
| 104 | 655 | 0.02 | Q39QB3 | *Geobacter metallireducens* | NADH dehydrogenase I, G subunit |
| 105 | 654 | 0.02 | A0A0D5NE26 | *Geobacter sulfurreducens* | Multidrug resistance protein B |
| 106 | 654 | 0.02 | Q39QU2 | *Geobacter metallireducens* | Phosphoenolpyruvate carboxykinase (ATP) (PCK) (PEP carboxykinase) (PEPCK) (EC 4.1.1.49) |
| 107 | 654 | 0.02 | Q39R25 | *Geobacter metallireducens* | Sensor diguanylate cyclase/phosphodiesterase, PAS, PAS and PAS domain-containing |
| 108 | 653 | 0.02 | G2J999 | *Cand. Glomeribacter gigasporarum* | TRAG family protein |
| 109 | 652 | 0.02 | | | |
| 110 | 651 | 0.02 | | | |
| 111 | 650 | 0.02 | A0A0D5N8G6 | *Geobacter sulfurreducens* | Cation transporter |
| 112 | 648 | 0.02 | A0A0B5BK17 | *Geobacter pickeringii* | Diguanylate cyclase |
| 113 | 647 | 0.02 | A0A0D5N712 | *Geobacter sulfurreducens* | Phosphoglucomutase |
| 114 | 643 | 0.02 | Q39QP7 | *Geobacter metallireducens* | Lon protease (EC 3.4.21.53) (ATP-dependent protease La) |
| 115 | 643 | 0.02 | U2TE33 | *Mitsuokella sp.* | DNA topoisomerase |
| 116 | 642 | 0.02 | A0A0D5NDJ8 | *Geobacter sulfurreducens* | Stage II sporulation protein E |
| 117 | 641 | 0.02 | Q74FD5 | *Geobacter sulfurreducens* | Hydroxylamine reductase (EC 1.7.99.1) (Hybrid-cluster protein) (HCP) (Prismane protein) |
| 118 | 640 | 0.02 | Q39RI3 | *Geobacter metallireducens* | Sensor histidine kinase, HAMP domain-containing, 2 heme-binding sites |
| 119 | 639 | 0.02 | B5EFV6 | *Geobacter bemidjiensis* | Bifunctional protein PutA |
| 120 | 637 | 0.02 | A0A0D5N2P4 | *Geobacter sulfurreducens* | Uncharacterized protein |
| 121 | 637 | 0.02 | A0A1T4RHG2 | *Geobacter thiogenes* | Nif-specific regulatory protein |
| 122 | 636 | 0.02 | | | |

| | | | | | |
|---|---|---|---|---|---|
| 123 | 631 | 0.02 | B3E9R4 | *Geobacter lovleyi* | Isocitrate dehydrogenase, NADP-dependent (EC 1.1.1.42) |
| 124 | 631 | 0.02 | Q39QJ5 | *Geobacter metallireducens* | Sensor histidine kinase CheA associated with MCPs of classes 40H and 40+24H |
| 125 | 630 | 0.02 | A0A0B5BH05 | *Geobacter pickeringii* | Malic enzyme (EC 1.1.1.40) |
| 126 | 624 | 0.02 | Q39UH4 | *Geobacter metallireducens* | Lon protease (EC 3.4.21.53) (ATP-dependent protease La) |
| 127 | 622 | 0.02 | A0A059GCP4 | *Hyphomonas oceanitis* | Conjugal transfer protein TraG |
| 128 | 622 | 0.02 | P61679 | *Geobacter sulfurreducens* | UDP-N-acetylmuramate--L-alanine ligase (EC 6.3.2.8) (UDP-N-acetylmuramoyl-L-alanine synthetase) |
| 129 | 619 | 0.02 | Q39WU2 | *Geobacter metallireducens* | Phosphate-selective outer membrane channel |
| 130 | 617 | 0.02 | A0A0B5BCZ8 | *Geobacter pickeringii* | Succinate dehydrogenase (EC 1.3.5.1) |
| 131 | 617 | 0.02 | A0A0B5BI89 | *Geobacter pickeringii* | Type II secretion system protein |
| 132 | 617 | 0.02 | A0A0L8EU90 | *Achromobacter sp.* | Uncharacterized protein |
| 133 | 616 | 0.02 | A0A0D5ND99 | *Geobacter sulfurreducens* | ATPase AAA |
| 134 | 616 | 0.02 | A0A174BLS6 | *Mitsuokella jalaludinii* | DNA topoisomerase 3 (EC 5.99.1.2) |
| 135 | 616 | 0.02 | Q749N5 | *Geobacter sulfurreducens* | Acetate kinase (EC 2.7.2.1) (Acetokinase) |
| 136 | 615 | 0.02 | Q748Z4 | *Geobacter sulfurreducens* | 30S ribosomal protein S3 |
| 137 | 607 | 0.02 | Q74GG6 | *Geobacter sulfurreducens* | Chromosomal replication initiator protein DnaA |
| 138 | 607 | 0.02 | Q39QY3 | *Geobacter metallireducens* | Efflux pump, RND family, inner membrane protein |
| 139 | 604 | 0.02 | A0A0A8WN28 | *Geobacter sp.* | Protein TraN |
| 140 | 602 | 0.02 | Q39SJ2 | *Geobacter metallireducens* | Nitrogen fixation master sigma-54-dependent transcriptional response regulator |
| 141 | 599 | 0.02 | A0A0D5N8Q2 | *Geobacter sulfurreducens* | Uncharacterized protein |
| 142 | 599 | 0.02 | Q39XG6 | *Geobacter metallireducens* | Pyruvate carboxylase (EC 6.4.1.1) |
| 143 | 599 | 0.02 | Q74DL9 | *Geobacter sulfurreducens* | Transposase of ISGsu2, IS5 family |
| 144 | 597 | 0.02 | Q39UA2 | *Geobacter metallireducens* | Adenylosuccinate lyase (ASL) (EC 4.3.2.2) (Adenylosuccinase) |
| 145 | 594 | 0.02 | A0A178FWP5 | *Acinetobacter sp.* | Transposase |
| 146 | 592 | 0.02 | A0A0B5BHV9 | *Geobacter pickeringii* | ATPase AAA |
| 147 | 590 | 0.02 | A0A0B5BDG1 | *Geobacter pickeringii* | tRNA uridine 5-carboxymethylaminomethyl modification enzyme MnmG (Glucose-inhibited division protein A) |
| 148 | 590 | 0.02 | A0A0D5NAK5 | *Geobacter sulfurreducens* | Bifunctional uridylyltransferase/uridylyl-removing enzyme (UTase/UR) (Bifunctional [protein-PII] modification enzyme) (Bifunctional nitrogen sensor protein) [Includes: [Protein-PII]-UMP uridylyl-removing enzyme (UR) (EC 3.1.4.-); [Protein-PII] uridylyltransferase (PII uridylyltransferase) (UTase) (EC 2.7.7.59)] |
| 149 | 589 | 0.02 | A0A0D5N7M4 | *Geobacter sulfurreducens* | Chemotaxis protein |
| 150 | 589 | 0.02 | Q39RQ5 | *Geobacter metallireducens* | Chaperone protein ClpB |
| 151 | 589 | 0.02 | Q74B77 | *Geobacter sulfurreducens* | Uncharacterized protein |
| 152 | 588 | 0.02 | A0A1G0M577 | *Geobacteraceae bacterium* | Uncharacterized protein |
| 153 | 586 | 0.02 | A0A0D5N7R7 | *Geobacter sulfurreducens* | GntR family transcriptional regulator |

| | | | | | |
|---|---|---|---|---|---|
| 154 | 584 | 0.02 | A0A0A8WPS5 | *Geobacter sp.* | Efflux pump membrane transporter |
| 155 | 582 | 0.02 | A0A0D5N8E2 | *Geobacter sulfurreducens* | ATP synthase (EC 3.6.3.14) |
| 156 | 581 | 0.02 | A0A0D5NCW1 | *Geobacter sulfurreducens* | Histidine kinase |
| 157 | 579 | 0.02 | A5GB28 | *Geobacter uraniireducens* | Plasmid maintenance system antidote protein, XRE family |
| 158 | 579 | 0.02 | Q39QN4 | *Geobacter metallireducens* | Pentapeptide repeat protein |
| 159 | 578 | 0.02 | Q749H1 | *Geobacter sulfurreducens* | Transposase of ISGsu3, IS5 family |
| 160 | 572 | 0.02 | Q39YR4 | *Geobacter metallireducens* | FtsK/SpoIIIE domain protein |
| 161 | 572 | 0.02 | | | |
| 162 | 571 | 0.02 | A0A0B5BE38 | *Geobacter pickeringii* | Transcription-repair-coupling factor (TRCF) (EC 3.6.4.-) |
| 163 | 570 | 0.02 | Q39ZQ2 | *Geobacter metallireducens* | Uncharacterized protein |
| 164 | 569 | 0.02 | Q39RM8 | *Geobacter metallireducens* | Glycosyltransferase |
| 165 | 568 | 0.02 | Q39SI5 | *Geobacter metallireducens* | Fumarate hydratase class I (EC 4.2.1.2) |
| 166 | 567 | 0.02 | A0A0B5BIS9 | *Geobacter pickeringii* | Multidrug transporter AcrB |
| 167 | 564 | 0.02 | Q746S4 | *Geobacter sulfurreducens* | NADH-quinone oxidoreductase subunit B/C/D (EC 1.6.5.11) (NADH dehydrogenase I subunit B/C/D) (NDH-1 subunit B/C/D) |
| 168 | 563 | 0.02 | P60789 | *Geobacter sulfurreducens* | Elongation factor 4 (EF-4) (EC 3.6.5.n1) (Ribosomal back-translocase LepA) |
| 169 | 562 | 0.02 | Q39ZH1 | *Geobacter metallireducens* | Glutamine--fructose-6-phosphate aminotransferase [isomerizing] (EC 2.6.1.16) (D-fructose-6-phosphate amidotransferase) (GFAT) (Glucosamine-6-phosphate synthase) (Hexosephosphate aminotransferase) (L-glutamine--D-fructose-6-phosphate amidotransferase) |
| 170 | 562 | 0.02 | Q3A5L6 | *Pelobacter carbinolicus* | Uncharacterized protein |
| 171 | 562 | 0.02 | Q74H49 | *Geobacter sulfurreducens* | Uncharacterized protein |
| 172 | 559 | 0.02 | Q39X49 | *Geobacter metallireducens* | Dystroglycan-type cadherin-like domain repeat protein |
| 173 | 557 | 0.02 | A0A0B5BDE2 | *Geobacter pickeringii* | ATP phosphoribosyltransferase regulatory subunit |
| 174 | 556 | 0.02 | Q39ZL2 | *Geobacter metallireducens* | TrkA domain protein |
| 175 | 555 | 0.02 | Q39VY8 | *Geobacter metallireducens* | Glutamine synthetase (EC 6.3.1.2) |
| 176 | 554 | 0.02 | A0A0D5N5P7 | *Geobacter sulfurreducens* | Chemotaxis protein |
| 177 | 554 | 0.02 | A0A1G0LHM5 | *Geobacteraceae bacterium* | Conjugal transfer protein TrbE |
| 178 | 552 | 0.02 | Q39XB5 | *Geobacter metallireducens* | 30S ribosomal protein S1 |
| 179 | 551 | 0.02 | B5EGM4 | *Geobacter bemidjiensis* | Glycerol/1,2-propanediol dehydratase |
| 180 | 550 | 0.02 | A5G9A7 | *Geobacter uraniireducens* | NADH dehydrogenase subunit M (EC 1.6.5.3) |
| 181 | 549 | 0.02 | Q39XC1 | *Geobacter metallireducens* | PilB/PulE/GspE family ATPase |
| 182 | 547 | 0.02 | Q39T42 | *Geobacter metallireducens* | Outer membrane protein assembly factor BamA |
| 183 | 546 | 0.02 | Q39QT7 | *Geobacter metallireducens* | Cytochrome c |
| 184 | 545 | 0.02 | Q39ZI6 | *Geobacter metallireducens* | DNA helicase (EC 3.6.4.12) |
| 185 | 544 | 0.02 | A0A0D5N7I1 | *Geobacter sulfurreducens* | NADH-quinone oxidoreductase subunit N (EC 1.6.5.11) (NADH dehydrogenase I subunit N) (NDH-1 subunit N) |

| | | | | | |
|---|---|---|---|---|---|
| 186 | 544 | 0.02 | A5GFJ7 | *Geobacter uraniireducens* | Erythronolide synthase (EC 2.3.1.94) |
| 187 | 544 | 0.02 | B3EB49 | *Geobacter lovleyi* | Uncharacterized protein |
| 188 | 544 | 0.02 | Q74C83 | *Geobacter sulfurreducens* | ATP-dependent Clp protease ATP-binding subunit ClpX |
| 189 | 542 | 0.02 | Q39UG5 | *Geobacter metallireducens* | Aspartokinase (EC 2.7.2.4) |
| 190 | 541 | 0.02 | Q74C70 | *Geobacter sulfurreducens* | Phosphoglucosamine mutase (EC 5.4.2.10) |
| 191 | 540 | 0.02 | A0A0D5N400 | *Geobacter sulfurreducens* | Uncharacterized protein |
| 192 | 540 | 0.02 | A0A0D5N7N0 | *Geobacter sulfurreducens* | Glutamate synthase |
| 193 | 540 | 0.02 | Q74CR5 | *Geobacter sulfurreducens* | Serine hydroxymethyltransferase (SHMT) (Serine methylase) (EC 2.1.2.1) |
| 194 | 540 | 0.02 | Q39WC2 | *Geobacter metallireducens* | DNA-directed DNA polymerase (EC 2.7.7.7) |
| 195 | 538 | 0.02 | A5GF11 | *Geobacter uraniireducens* | Acetyl-CoA carboxylase carboxyltransferase subunit alpha (EC 6.3.4.14) |
| 196 | 538 | 0.02 | A5GAY2 | *Geobacter uraniireducens* | DNA-directed RNA polymerase subunit beta' (RNAP subunit beta') (EC 2.7.7.6) (RNA polymerase subunit beta') (Transcriptase subunit beta') |
| 197 | 536 | 0.02 | A0A0B5BAC1 | *Geobacter pickeringii* | Dihydrolipoyl dehydrogenase (EC 1.8.1.4) |
| 198 | 536 | 0.02 | Q74GT3 | *Geobacter sulfurreducens* | LL-diaminopimelate aminotransferase (DAP-AT) (DAP-aminotransferase) (LL-DAP-aminotransferase) (EC 2.6.1.83) |
| 199 | 534 | 0.02 | Q39QD0 | *Geobacter metallireducens* | Periplasmically oriented, membrane-bound [NiFe]-hydrogenase, large subunit |
| 200 | 534 | 0.02 | Q74AJ5 | *Geobacter sulfurreducens* | Glycoside hydrolase, family 57, DUF3536 domain-containing |
| 201 | 533 | 0.02 | Q39Y82 | *Geobacter metallireducens* | 5-methyltetrahydrofolate--homocysteine S-methyltransferase, cobalamin-dependent |
| 202 | 532 | 0.02 | A0A0C1TL30 | *Geobacter soli* | Flagellar basal body stator protein MotB |
| 203 | 532 | 0.02 | A0A0D5N2S0 | *Geobacter sulfurreducens* | Uncharacterized protein |
| 204 | 531 | 0.02 | A5GAY1 | *Geobacter uraniireducens* | DNA-directed RNA polymerase subunit beta (RNAP subunit beta) (EC 2.7.7.6) (RNA polymerase subunit beta) (Transcriptase subunit beta) |
| 205 | 528 | 0.02 | Q39VY0 | *Geobacter metallireducens* | Malate dehydrogenase (EC 1.1.1.37) |
| 206 | 527 | 0.02 | A0A0B5BIQ7 | *Geobacter pickeringii* | Flagellar basal-body rod protein FlgF (Flagellar hook protein FlgE) |
| 207 | 526 | 0.02 | A0A0B5BD43 | *Geobacter pickeringii* | ABC transporter permease |
| 208 | 525 | 0.02 | A0A0B5BK56 | *Geobacter pickeringii* | Lon protease (EC 3.4.21.53) (ATP-dependent protease La) |
| 209 | 525 | 0.02 | A5GDX4 | *Geobacter uraniireducens* | DNA gyrase subunit B (EC 5.99.1.3) |
| 210 | 524 | 0.02 | | | |
| 211 | 523 | 0.02 | A0A1H6GN37 | *Selenomonas ruminantium* | DNA topoisomerase-3 |
| 212 | 523 | 0.02 | Q39QA3 | *Geobacter metallireducens* | ATP synthase subunit a (ATP synthase F0 sector subunit a) (F-ATPase subunit 6) |
| 213 | 523 | 0.02 | | | |
| 214 | 522 | 0.02 | Q39VA1 | *Geobacter metallireducens* | Polyribonucleotide nucleotidyltransferase (EC 2.7.7.8) (Polynucleotide phosphorylase) (PNPase) |
| 215 | 520 | 0.02 | A0A0C1U608 | *Geobacter soli* | Chemotaxis protein |
| 216 | 518 | 0.02 | Q39T98 | *Geobacter metallireducens* | Cytochrome c |
| 217 | 518 | 0.02 | Q39Z79 | *Geobacter metallireducens* | Twitching motility pilus retraction ATPase |
| 218 | 517 | 0.02 | A0A1G0MAK9 | *Geobacteraceae bacterium* | Sodium:proton exchanger |

| | | | | | |
|---|---|---|---|---|---|
| 219 | 517 | 0.02 | B3EAX1 | *Geobacter lovleyi* | Protein CyaE |
| 220 | 517 | 0.02 | Q39W76 | *Geobacter metallireducens* | Ketol-acid reductoisomerase (NADP(+)) (KARI) (EC 1.1.1.86) (Acetohydroxy-acid isomeroreductase) (AHIR) (Alpha-keto-beta-hydroxylacyl reductoisomerase) (Ketol-acid reductoisomerase type 1) (Ketol-acid reductoisomerase type I) |
| 221 | 515 | 0.02 | A0A0B5B5Z5 | *Geobacter pickeringii* | DNA polymerase III subunit beta (EC 2.7.7.7) |
| 222 | 515 | 0.02 | A0A0D5N952 | *Geobacter sulfurreducens* | Flagellar motor switch protein FliG |
| 223 | 515 | 0.02 | A0A0D5NAW4 | *Geobacter sulfurreducens* | Cation transporter |
| 224 | 514 | 0.02 | A0A0Q7AWL1 | *Nocardioides sp.* | Uncharacterized protein |
| 225 | 512 | 0.02 | A1AN79 | *Pelobacter propionicus* | Adenosylmethionine-8-amino-7-oxononanoate aminotransferase (EC 2.6.1.62) (7,8-diamino-pelargonic acid aminotransferase) (DAPA AT) (DAPA aminotransferase) (7,8-diaminononanoate synthase) (DANS) (Diaminopelargonic acid synthase) |
| 226 | 510 | 0.02 | Q39ZS3 | *Geobacter metallireducens* | Chromosomal replication initiator protein DnaA |
| 227 | 507 | 0.02 | A0A0B5BIA1 | *Geobacter pickeringii* | DNA-directed DNA polymerase (EC 2.7.7.7) |
| 228 | 507 | 0.02 | B9LZR8 | *Geobacter daltonii* | Succinyl:acetate coenzyme A transferase |
| 229 | 507 | 0.02 | Q749D7 | *Geobacter sulfurreducens* | Nitrogenase molybdenum-iron cofactor biosynthesis protein NifEN |
| 230 | 505 | 0.02 | A0A0B5BCH1 | *Geobacter pickeringii* | Hydrogenase 2 large subunit (EC 1.12.99.6) |
| 231 | 504 | 0.02 | Q39VA8 | *Geobacter metallireducens* | Transcription termination/antitermination protein NusA |
| 232 | 503 | 0.02 | A5GFJ6 | *Geobacter uraniireducens* | Beta-hydroxyacyl-(Acyl-carrier-protein) dehydratase, FabA/FabZ |
| 233 | 503 | 0.02 | | | |
| 234 | 501 | 0.02 | A0A0A8WNZ4 | *Geobacter sp.* | DNA gyrase subunit A (EC 5.99.1.3) |
| 235 | 501 | 0.02 | Q74BC2 | *Geobacter sulfurreducens* | Integrative genetic element Gsu56, integrase |
| 236 | 499 | 0.02 | A0A0D5NE46 | *Geobacter sulfurreducens* | Molecular chaperone DnaJ |
| 237 | 497 | 0.02 | A0A0A8WNV3 | *Geobacter sp.* | Uncharacterized protein |
| 238 | 495 | 0.02 | A0A0B5BCL9 | *Geobacter pickeringii* | Uncharacterized protein |
| 239 | 494 | 0.02 | Q39WR2 | *Geobacter metallireducens* | CRISPR-associated endonuclease Cas1 (EC 3.1.-.-) |
| 240 | 492 | 0.02 | P61667 | *Geobacter sulfurreducens* | DNA mismatch repair protein MutS |
| 241 | 492 | 0.02 | Q74BB9 | *Geobacter sulfurreducens* | TraG family protein |
| 242 | 491 | 0.02 | A0A0B5BB89 | *Geobacter pickeringii* | Chromosome partition protein Smc |
| 243 | 491 | 0.02 | A0A0B5BIF0 | *Geobacter pickeringii* | NADH-ubiquinone oxidoreductase subunit 3 |
| 244 | 491 | 0.02 | Q39RT7 | *Geobacter metallireducens* | Efflux pump, RND superfamily, and rRNA methyltransferase, putative |
| 245 | 490 | 0.02 | A0A0B5B8F2 | *Geobacter pickeringii* | Sodium transporter |
| 246 | 490 | 0.02 | Q748V3 | *Geobacter sulfurreducens* | Lipoprotein cytochrome c |
| 247 | 490 | 0.02 | Q74F57 | *Geobacter sulfurreducens* | Membrane protein, major facilitator superfamily |
| 248 | 488 | 0.02 | A0A0B5BB11 | *Geobacter pickeringii* | Pyruvate carboxylase (EC 6.4.1.1) |
| 249 | 488 | 0.02 | Q39ZM0 | *Geobacter metallireducens* | Iron-sulfur cluster-binding sigma-54-dependent transcriptional regulator, FehydlgC and FeS domain-containing |

| | | | | | |
|---|---|---|---|---|---|
| 250 | 487 | 0.02 | A0A096AIH0 | *Veillonella montpellierensis* | DNA topoisomerase |
| 251 | 487 | 0.02 | A0A0D5N6Q1 | *Geobacter sulfurreducens* | FeS-binding protein |
| 252 | 486 | 0.02 | A0A0B5B668 | *Geobacter pickeringii* | Peptidase U32 |
| 253 | 486 | 0.02 | A0A0B5BC86 | *Geobacter pickeringii* | Multicopper oxidase |
| 254 | 485 | 0.02 | A0R7X1 | *Pelobacter propionicus* | Type III restriction enzyme, res subunit |
| 255 | 485 | 0.02 | Q74GU4 | *Geobacter sulfurreducens* | Acetylglutamate kinase (EC 2.7.2.8) (N-acetyl-L-glutamate 5-phosphotransferase) (NAG kinase) (NAGK) |
| 256 | 484 | 0.02 | A0A0B5BK89 | *Geobacter pickeringii* | FAD-binding protein |
| 257 | 484 | 0.02 | A0A0D5N5Y8 | *Geobacter sulfurreducens* | Cell division protein FtsA |
| 258 | 484 | 0.02 | Q39QV2 | *Geobacter metallireducens* | Alpha-glucan phosphorylase |
| 259 | 482 | 0.02 | A0A0B5BDP4 | *Geobacter pickeringii* | Polya polymerase |
| 260 | 481 | 0.02 | A0A0D5NB12 | *Geobacter sulfurreducens* | Aldehyde dehydrogenase |
| 261 | 481 | 0.02 | | | |
| 262 | 480 | 0.02 | A0A0D5N8X4 | *Geobacter sulfurreducens* | Flagellar M-ring protein |
| 263 | 480 | 0.02 | B3EA87 | *Geobacter lovleyi* | RNA polymerase sigma factor |
| 264 | 480 | 0.02 | Q39XL2 | *Geobacter metallireducens* | Phosphoenolpyruvate synthase (PEP synthase) (EC 2.7.9.2) (Pyruvate, water dikinase) |
| 265 | 480 | 0.02 | Q748I6 | *Geobacter sulfurreducens* | Metalloprotease domain protein, M6 family |
| 266 | 480 | 0.02 | Q74AF7 | *Geobacter sulfurreducens* | Methyl viologen-reducing hydrogenase, large subunit |
| 267 | 478 | 0.02 | Q39Q80 | *Geobacter metallireducens* | Sensor histidine kinase, HAMP domain-containing |
| 268 | 478 | 0.02 | Q39X64 | *Geobacter metallireducens* | Cell shape-determining protein MreB |
| 269 | 477 | 0.02 | A0A0B5BBC9 | *Geobacter pickeringii* | Type I citrate synthase (EC 2.3.3.1) |
| 270 | 477 | 0.02 | A0A0C1QTJ3 | *Geobacter soli* | Hemolysin D |
| 271 | 477 | 0.02 | C6DZ85 | *Geobacter sp.* | Conjugal transfer protein |
| 272 | 476 | 0.02 | Q39QT3 | *Geobacter metallireducens* | Mannose-1-phosphate guanylyltransferase and mannose-6-phosphate isomerase-related protein |
| 273 | 475 | 0.02 | Q39QK7 | *Geobacter metallireducens* | Outer membrane channel OmpJ |
| 274 | 475 | 0.02 | Q74B66 | *Geobacter sulfurreducens* | TrfA family protein |
| 275 | 475 | 0.02 | | | |
| 276 | 474 | 0.02 | F3ZQL8 | *Bacteroides coprosuis* | Transposase IS4 family protein |
| 277 | 474 | 0.02 | Q39R87 | *Geobacter metallireducens* | NADPH oxidoreductase, alpha subunit |
| 278 | 473 | 0.02 | | | |
| 279 | 472 | 0.02 | A0A0B5B9H5 | *Geobacter pickeringii* | Uncharacterized protein |
| 280 | 471 | 0.02 | Q39U98 | *Geobacter metallireducens* | Phosphoglycerate kinase (EC 2.7.2.3) |
| 281 | 470 | 0.02 | Q39RM6 | *Geobacter metallireducens* | Glycosyltransferase |
| 282 | 469 | 0.02 | A0A0A8WLB0 | *Geobacter sp.* | Protein TraH |
| 283 | 469 | 0.02 | B9M1Q4 | *Geobacter daltonii* | Biotin-dependent acyl-CoA carboxylase, biotin carboxylase subunit |

| | | | | | |
|---|---|---|---|---|---|
| 284 | 469 | 0.02 | Q39ZP3 | *Geobacter metallireducens* | VWFA superfamily protein |
| 285 | 469 | 0.02 | Q39VY3 | *Geobacter metallireducens* | Aspartate--tRNA(Asp/Asn) ligase (EC 6.1.1.23) (Aspartyl-tRNA synthetase) (AspRS) (Non-discriminating aspartyl-tRNA synthetase) (ND-AspRS) |
| 286 | 468 | 0.02 | Q39ZH5 | *Geobacter metallireducens* | Cytochrome c/b |
| 287 | 468 | 0.02 | Q74B80 | *Geobacter sulfurreducens* | Uncharacterized protein |
| 288 | 466 | 0.02 | A5GCN0 | *Geobacter uraniireducens* | Short chain amide porin |
| 289 | 465 | 0.02 | Q39X71 | *Geobacter metallireducens* | Uncharacterized protein |
| 290 | 463 | 0.02 | A0A1G0LE89 | *Geobacteraceae bacterium* | Aconitate hydratase (Aconitase) (EC 4.2.1.3) |
| 291 | 462 | 0.02 | Q39X99 | *Geobacter metallireducens* | UDP-glucuronate 4-dehydrogenase, decarboxylating |
| 292 | 461 | 0.02 | A0A0B5BDE7 | *Geobacter pickeringii* | Fis family transcriptional regulator |
| 293 | 461 | 0.02 | A0A0B5BKE7 | *Geobacter pickeringii* | NADH-quinone oxidoreductase subunit N (EC 1.6.5.11) (NADH dehydrogenase I subunit N) (NDH-1 subunit N) |
| 294 | 459 | 0.02 | A5G8I3 | *Geobacter uraniireducens* | Chaperone protein ClpB |
| 295 | 458 | 0.02 | Q39Q79 | *Geobacter metallireducens* | Winged-helix transcriptional response regulator |
| 296 | 458 | 0.02 | Q747G2 | *Geobacter sulfurreducens* | Outer membrane channel OmpJ |
| 297 | 457 | 0.02 | Q74BB8 | *Geobacter sulfurreducens* | Metal-dependent phosphohydrolase |
| 298 | 456 | 0.02 | A0A1G0E0Z3 | *Gammaproteobacteria bacterium* | Conjugal transfer protein TrbE |
| 299 | 456 | 0.02 | A0A1G0LEY2 | *Geobacteraceae bacterium* | Uncharacterized protein |
| 300 | 456 | 0.02 | Q74AT0 | *Geobacter sulfurreducens* | Lysine--tRNA ligase (EC 6.1.1.6) (Lysyl-tRNA synthetase) (LysRS) |
| 301 | 456 | 0.02 | Q39WF6 | *Geobacter metallireducens* | Ribonuclease Y (RNase Y) (EC 3.1.-.-) |
| 302 | 455 | 0.02 | A0A0D5N8K2 | *Geobacter sulfurreducens* | RND transporter |
| 303 | 455 | 0.02 | B3EAX6 | *Geobacter lovleyi* | Uncharacterized protein |
| 304 | 455 | 0.02 | Q39S10 | *Geobacter metallireducens* | Endopeptidase La (EC 3.4.21.53) |
| 305 | 454 | 0.02 | A0A0A8WRD3 | *Geobacter sp.* | Putative ABC transporter ATP-binding protein YkpA |
| 306 | 454 | 0.02 | A0A0B5BG42 | *Geobacter pickeringii* | Vitamin B12-dependent ribonucleotide reductase (EC 1.17.4.1) |
| 307 | 454 | 0.02 | Q748B3 | *Geobacter sulfurreducens* | UDP-N-acetylglucosamine 1-carboxyvinyltransferase (EC 2.5.1.7) (Enoylpyruvate transferase) (UDP-N-acetylglucosamine enolpyruvyl transferase) (EPT) |
| 308 | 454 | 0.02 | Q39X25 | *Geobacter metallireducens* | Sensor diguanylate cyclase/phosphodiesterase, GAF domain-containing |
| 309 | 454 | 0.02 | Q39X31 | *Geobacter metallireducens* | Protein translocase subunit SecA |
| 310 | 453 | 0.02 | Q74BT7 | *Geobacter sulfurreducens* | Sensor histidine kinase cyclic nucleotide phosphodiesterase, GAF, GAF and HD-GYP-related domain-containing |
| 311 | 450 | 0.02 | Q39VA6 | *Geobacter metallireducens* | Translation initiation factor IF-2 |
| 312 | 449 | 0.02 | P61185 | *Geobacter sulfurreducens* | Chaperone protein HtpG (Heat shock protein HtpG) (High temperature protein G) |
| 313 | 448 | 0.02 | A0A0B5BE21 | *Geobacter pickeringii* | Histidine kinase |
| 314 | 446 | 0.02 | A0A077XLW5 | *Sphinx1.76-related DNA* | Replication protein |

| | | | | | |
|---|---|---|---|---|---|
| 315 | 446 | 0.02 | A0A0D5NAV3 | *Geobacter sulfurreducens* | Peptidase S41 |
| 316 | 445 | 0.02 | A0A0B5BJV3 | *Geobacter pickeringii* | ABC transporter substrate-binding protein |
| 317 | 445 | 0.02 | | | |
| 318 | 444 | 0.02 | Q746T2 | *Geobacter sulfurreducens* | NADH-quinone oxidoreductase subunit H 2 (EC 1.6.5.11) (NADH dehydrogenase I subunit H 2) (NDH-1 subunit H 2) |
| 319 | 444 | 0.02 | Q39YH3 | *Geobacter metallireducens* | UDP-N-acetylglucosamine 4,6-dehydratase and UDP-2-acetamido-2,6-dideoxy-alpha-D-xylo-4-hexulose 5-epimerase |
| 320 | 443 | 0.02 | A0A1T4RGQ7 | *Geobacter thiogenes* | NAD+---dinitrogen-reductase ADP-D-ribosyltransferase |
| 321 | 442 | 0.02 | B3EAX2 | *Geobacter lovleyi* | Efflux transporter, RND family, MFP subunit |
| 322 | 442 | 0.02 | Q39WZ1 | *Geobacter metallireducens* | Nitrogen fixation protein NifU |
| 323 | 441 | 0.02 | P61523 | *Geobacter sulfurreducens* | Argininosuccinate synthase (EC 6.3.4.5) (Citrulline--aspartate ligase) |
| 324 | 441 | 0.02 | Q39SL1 | *Geobacter metallireducens* | Lon protease (EC 3.4.21.53) (ATP-dependent protease La) |
| 325 | 441 | 0.02 | | | |
| 326 | 440 | 0.02 | A0A0B5BBZ4 | *Geobacter pickeringii* | DNA polymerase I (EC 2.7.7.7) |
| 327 | 440 | 0.02 | A0A0D5N949 | *Geobacter sulfurreducens* | Response regulator |
| 328 | 439 | 0.02 | A0A0A8WQV9 | *Geobacter sp.* | Protein TraU |
| 329 | 439 | 0.02 | Q39Y18 | *Geobacter metallireducens* | Transcription termination/antitermination protein NusG |
| 330 | 437 | 0.02 | A0A0C1TS26 | *Geobacter soli* | L-aspartate oxidase (EC 1.4.3.16) |
| 331 | 437 | 0.02 | A0A0T5YH86 | *Acinetobacter baumannii* | Mobilization protein A |
| 332 | 437 | 0.02 | Q748X2 | *Geobacter sulfurreducens* | 3-isopropylmalate dehydrogenase (EC 1.1.1.85) (3-IPM-DH) (Beta-IPM dehydrogenase) (IMDH) |
| 333 | 437 | 0.02 | Q39Q96 | *Geobacter metallireducens* | Type II secretion system inner membrane protein GspF |
| 334 | 436 | 0.02 | Q39Z54 | *Geobacter metallireducens* | Membrane protein, putative |
| 335 | 435 | 0.02 | A0A0B5BCB2 | *Geobacter pickeringii* | Nitrogen fixation protein NifE |
| 336 | 433 | 0.01 | A0A0B5B7K6 | *Geobacter pickeringii* | Cytochrome C biogenesis protein ResC |
| 337 | 433 | 0.01 | Q39QM2 | *Geobacter metallireducens* | K(+)-insensitive pyrophosphate-energized proton pump (EC 3.6.1.1) (Membrane-bound proton-translocating pyrophosphatase) (Pyrophosphate-energized inorganic pyrophosphatase) (H(+)-PPase) |
| 338 | 431 | 0.01 | A0A0B5BC52 | *Geobacter pickeringii* | General secretion pathway protein GspD |
| 339 | 431 | 0.01 | A0A0D5N4Y8 | *Geobacter sulfurreducens* | Pilus assembly protein PilB |
| 340 | 431 | 0.01 | Q39RF8 | *Geobacter metallireducens* | Carbon starvation protein CstA |
| 341 | 430 | 0.01 | Q39QZ9 | *Geobacter metallireducens* | Flagellar motor switch protein FliG |
| 342 | 430 | 0.01 | Q39UW3 | *Geobacter metallireducens* | Succinyl:acetate coenzyme A transferase |
| 343 | 430 | 0.01 | Q39W43 | *Geobacter metallireducens* | Oligoendopeptidase F |
| 344 | 430 | 0.01 | Q747G4 | *Geobacter sulfurreducens* | (R)-methylmalonyl-CoA mutase, isobutyryl-CoA mutase-like catalytic subunit |
| 345 | 428 | 0.01 | Q748Y3 | *Geobacter sulfurreducens* | 50S ribosomal protein L1 |
| 346 | 427 | 0.01 | A0A0B5B6P6 | *Geobacter pickeringii* | Histidine kinase |

| | | | | | |
|---|---|---|---|---|---|
| 347 | 426 | 0.01 | Q39QU4 | *Geobacter metallireducens* | ATP-dependent DNA helicase DinG |
| 348 | 426 | 0.01 | Q39XW9 | *Geobacter metallireducens* | Nitrogenase protein alpha chain (EC 1.18.6.1) |
| 349 | 424 | 0.01 | A0A0B5BGV8 | *Geobacter pickeringii* | Zinc protease |
| 350 | 424 | 0.01 | A5GDX5 | *Geobacter uraniireducens* | DNA gyrase subunit A (EC 5.99.1.3) |
| 351 | 424 | 0.01 | Q39VR6 | *Geobacter metallireducens* | DNA mismatch repair protein MutS |
| 352 | 424 | 0.01 | Q74D56 | *Geobacter sulfurreducens* | Aspartate--tRNA(Asp/Asn) ligase (EC 6.1.1.23) (Aspartyl-tRNA synthetase) (AspRS) (Non-discriminating aspartyl-tRNA synthetase) (ND-AspRS) |
| 353 | 423 | 0.01 | E1QK40 | *Desulfarculus baarsii* | DNA-cytosine methyltransferase |
| 354 | 422 | 0.01 | A0A0D5N3H2 | *Geobacter sulfurreducens* | DEAD/DEAH box helicase |
| 355 | 422 | 0.01 | A0A1F9PXI1 | *Desulfuromonadales bacterium* | Cytosine-specific methyltransferase (EC 2.1.1.37) |
| 356 | 422 | 0.01 | Q74F32 | *Geobacter sulfurreducens* | Periplasmically oriented, membrane-bound formate dehydrogenase, major subunit, selenocysteine-containing |
| 357 | 421 | 0.01 | Q39PS5 | *Geobacter metallireducens* | Quinolinate synthase A (EC 2.5.1.72) |
| 358 | 421 | 0.01 | Q39QK5 | *Geobacter metallireducens* | Uncharacterized protein |
| 359 | 421 | 0.01 | Q8VQM8 | *Prevotella intermedia* | Mobilization protein |
| 360 | 420 | 0.01 | K9DJ35 | *Veillonella seminalis* | DNA topoisomerase III |
| 361 | 420 | 0.01 | Q39Q46 | *Geobacter metallireducens* | ABC transporter, ATP-binding/membrane protein |
| 362 | 420 | 0.01 | Q39ZE2 | *Geobacter metallireducens* | Polyphosphate kinase (EC 2.7.4.1) (ATP-polyphosphate phosphotransferase) (Polyphosphoric acid kinase) |
| 363 | 418 | 0.01 | B9M5J7 | *Geobacter daltonii* | CRISPR-associated protein Csd2 |
| 364 | 418 | 0.01 | Q39PY9 | *Geobacter metallireducens* | Histidine kinase (EC 2.7.13.3) |
| 365 | 418 | 0.01 | Q39US2 | *Geobacter metallireducens* | Carbamoyl-phosphate synthase small chain (EC 6.3.5.5) (Carbamoyl-phosphate synthetase glutamine chain) |
| 366 | 418 | 0.01 | Q39X54 | *Geobacter metallireducens* | Peptidoglycan transpeptidase |
| 367 | 418 | 0.01 | Q39YS7 | *Geobacter metallireducens* | Peptidoglycan transglycosylase and transpeptidase MrcA |
| 368 | 416 | 0.01 | A5GE75 | *Geobacter metallireducens* | Heavy metal translocating P-type ATPase |
| 369 | 416 | 0.01 | Q39RQ8 | *Geobacter metallireducens* | Ribose-phosphate pyrophosphokinase (RPPK) (EC 2.7.6.1) (5-phospho-D-ribosyl alpha-1-diphosphate) (Phosphoribosyl diphosphate synthase) (Phosphoribosyl pyrophosphate synthase) (P-Rib-PP synthase) (PRPP synthase) (PRPPase) |
| 370 | 415 | 0.01 | A0A1T4YKH5 | *Prosthecobacter debontii* | Antirestriction protein ArdC |
| 371 | 415 | 0.01 | Q39ZP1 | *Geobacter metallireducens* | ATPase, AAA family |
| 372 | 415 | 0.01 | Q74GH2 | *Geobacter sulfurreducens* | Cytochrome c, and cytochrome b |
| 373 | 414 | 0.01 | C6DZ84 | *Geobacter sp.* | p-type conjugative transfer protein TrbG |
| 374 | 414 | 0.01 | Q39X42 | *Geobacter metallireducens* | Alpha-1,4 glucan phosphorylase (EC 2.4.1.1) |
| 375 | 414 | 0.01 | Q39XX9 | *Geobacter metallireducens* | DNA-directed RNA polymerase subunit alpha (RNAP subunit alpha) (EC 2.7.7.6) (RNA polymerase subunit alpha) (Transcriptase subunit alpha) |

| | | | | | |
|---|---|---|---|---|---|
| 376 | 413 | 0.01 | A0A0D5N8X6 | *Geobacter sulfurreducens* | Chemotaxis protein |
| 377 | 413 | 0.01 | Q39RS7 | *Geobacter metallireducens* | DUF748 repeat protein |
| 378 | 411 | 0.01 | A0A0B5BH19 | *Geobacter pickeringii* | Uncharacterized protein |
| 379 | 411 | 0.01 | Q39QS0 | *Geobacter metallireducens* | Radical SAM domain iron-sulfur cluster-binding oxidoreductase, DUF2344-containing |
| 380 | 411 | 0.01 | Q39RC6 | *Geobacter metallireducens* | DNA polymerase I (EC 2.7.7.7) |
| 381 | 411 | 0.01 | Q39RL8 | *Geobacter metallireducens* | Undecaprenyl-diphospho-oligosaccharide flippase |
| 382 | 410 | 0.01 | A0A0D5N732 | *Geobacter sulfurreducens* | Transcription-repair-coupling factor (TRCF) (EC 3.6.4.-) |
| 383 | 410 | 0.01 | Q39ZQ3 | *Geobacter metallireducens* | ATPase, AAA_5 family |
| 384 | 409 | 0.01 | C6DZ91 | *Geobacter sp.* | p-type conjugative transfer ATPase TrbB |
| 385 | 408 | 0.01 | Q748A7 | *Geobacter sulfurreducens* | Transcription termination factor Rho (EC 3.6.4.-) (ATP-dependent helicase Rho) |
| 386 | 407 | 0.01 | A0A0B5BJ17 | *Geobacter pickeringii* | UvrABC system protein C (Protein UvrC) (Excinuclease ABC subunit C) |
| 387 | 407 | 0.01 | Q39UC7 | *Geobacter metallireducens* | Response receiver sensor histidine kinase response regulator, PAS domain-containing |
| 388 | 405 | 0.01 | A0A0B5B8W1 | *Geobacter pickeringii* | ATP-dependent zinc metalloprotease FtsH (EC 3.4.24.-) |
| 389 | 405 | 0.01 | A0A0B5B9Q9 | *Geobacter pickeringii* | Glycosyl transferase |
| 390 | 404 | 0.01 | A0A0D5N645 | *Geobacter sulfurreducens* | Penicillin-binding protein |
| 391 | 404 | 0.01 | A0A1Q8EYA1 | *Aeromonas veronii* | Methyltransferase (EC 2.1.1.-) |
| 392 | 404 | 0.01 | B9M4C2 | *Geobacter daltonii* | Squalene cyclase |
| 393 | 404 | 0.01 | Q39T27 | *Geobacter metallireducens* | Enolase (EC 4.2.1.11) (2-phospho-D-glycerate hydro-lyase) (2-phosphoglycerate dehydratase) |
| 394 | 404 | 0.01 | Q39Q47 | *Geobacter metallireducens* | Selenocysteine-specific translation elongation factor |
| 395 | 403 | 0.01 | A0A0B5BFA5 | *Geobacter pickeringii* | Formate dehydrogenase |
| 396 | 403 | 0.01 | A0A0B5BK21 | *Geobacter pickeringii* | Inner membrane cytochrome H |
| 397 | 403 | 0.01 | P61946 | *Geobacter sulfurreducens* | S-adenosylmethionine synthase (AdoMet synthase) (EC 2.5.1.6) (MAT) (Methionine adenosyltransferase) |
| 398 | 401 | 0.01 | A0A0B5BFC0 | *Geobacter pickeringii* | Ni/Fe hydrogenase |
| 399 | 401 | 0.01 | A0A0B5BGX4 | *Geobacter pickeringii* | Stage II sporulation protein E |
| 400 | 401 | 0.01 | Q747K3 | *Geobacter sulfurreducens* | UvrABC system protein B (Protein UvrB) (Excinuclease ABC subunit B) |
| 401 | 400 | 0.01 | Q74H59 | *Geobacter sulfurreducens* | Chaperone protein DnaK (HSP70) (Heat shock 70 kDa protein) (Heat shock protein 70) |
| 402 | 400 | 0.01 | Q39WA8 | *Geobacter metallireducens* | Sodium/solute symporter, GAF and PP2C family protein serine/threonine phosphatase domain protein |
| 403 | 400 | 0.01 | Q39YM5 | *Geobacter metallireducens* | Peptidoglycan transglycosylase and transpeptidase FtsI |
| 404 | 399 | 0.01 | A0A0B5BK07 | *Geobacter pickeringii* | Histidine kinase |
| 405 | 399 | 0.01 | B0L930 | *uncultured Geobacter sp.* | Tuf (Fragment) |
| 406 | 398 | 0.01 | Q39UL6 | *Geobacter metallireducens* | Sodium/phosphate symporter, putative |
| 407 | 397 | 0.01 | A0A0D5N435 | *Geobacter sulfurreducens* | Multidrug transporter |
| 408 | 397 | 0.01 | Q74EP9 | *Geobacter sulfurreducens* | DNA-binding ATPase Uup |
| 409 | 397 | 0.01 | | | |

| | | | | | |
|---|---|---|---|---|---|
| 410 | 396 | 0.01 | Q39YP6 | *Geobacter metallireducens* | Histidinol-phosphate aminotransferase (EC 2.6.1.9) (Imidazole acetol-phosphate transaminase) |
| 411 | 395 | 0.01 | A0A0B5B6N9 | *Geobacter pickeringii* | Alanine--tRNA ligase (EC 6.1.1.7) (Alanyl-tRNA synthetase) (AlaRS) |
| 412 | 395 | 0.01 | Q39TA6 | *Geobacter metallireducens* | Inosine-5'-monophosphate dehydrogenase (IMP dehydrogenase) (IMPD) (IMPDH) (EC 1.1.1.205) |
| 413 | 394 | 0.01 | A0A0F2N896 | *Desulfobulbaceae bacterium* | Restriction endonuclease subunit R |
| 414 | 394 | 0.01 | Q39U52 | *Geobacter metallireducens* | Sensor histidine kinase, GAF domain-containing |
| 415 | 394 | 0.01 | | | |
| 416 | 393 | 0.01 | A0A0B5BCR8 | *Geobacter pickeringii* | Alpha-1,4 glucan phosphorylase (EC 2.4.1.1) |
| 417 | 392 | 0.01 | A0A0B5BGU8 | *Geobacter pickeringii* | Chemotaxis protein |
| 418 | 392 | 0.01 | Q39ZF6 | *Geobacter metallireducens* | Carbamoyltransferase HypF (EC 2.1.3.-) |
| 419 | 392 | 0.01 | Q39Z77 | *Geobacter metallireducens* | Alanine--tRNA ligase (EC 6.1.1.7) (Alanyl-tRNA synthetase) (AlaRS) |
| 420 | 392 | 0.01 | Q39QN3 | *Geobacter metallireducens* | UvrABC system protein C (Protein UvrC) (Excinuclease ABC subunit C) |
| 421 | 391 | 0.01 | A0A0D5N5H4 | *Geobacter sulfurreducens* | Cell division protein FtsK |
| 422 | 391 | 0.01 | Q74H61 | *Geobacter sulfurreducens* | Heat-inducible transcription repressor HrcA |
| 423 | 391 | 0.01 | Q39W55 | *Geobacter metallireducens* | RNA polymerase sigma-54 factor |
| 424 | 389 | 0.01 | Q39QK1 | *Geobacter metallireducens* | Adenylosuccinate synthetase (AMPSase) (AdSS) (EC 6.3.4.4) (IMP--aspartate ligase) |
| 425 | 389 | 0.01 | Q39QD6 | *Geobacter metallireducens* | Amino acid aminotransferase, putative |
| 426 | 388 | 0.01 | A0A0B5B672 | *Geobacter pickeringii* | FeS-binding protein |
| 427 | 388 | 0.01 | A0A1T4RHG8 | *Geobacter thiogenes* | Iron complex outermembrane recepter protein |
| 428 | 388 | 0.01 | A5GC25 | *Geobacter uraniireducens* | Propionyl-CoA carboxylase (EC 6.4.1.3) |
| 429 | 388 | 0.01 | B3EAB9 | *Geobacter lovleyi* | 4Fe-4S ferredoxin, iron-sulfur binding domain protein |
| 430 | 387 | 0.01 | A0A086P5I1 | *Sphingobium herbicidovorans* | Conjugal transfer protein TrbE |
| 431 | 387 | 0.01 | P61343 | *Geobacter sulfurreducens* | Aspartyl/glutamyl-tRNA(Asn/Gln) amidotransferase subunit B (Asp/Glu-ADT subunit B) (EC 6.3.5.-) |
| 432 | 387 | 0.01 | Q74E27 | *Geobacter sulfurreducens* | Ribonuclease Y (RNase Y) (EC 3.1.-.-) |
| 433 | 385 | 0.01 | Q39VU9 | *Geobacter metallireducens* | Polyprenyl-phospho-glycoside--protein O-glycosyltransferase DUF2723 membrane protein, putative |
| 434 | 384 | 0.01 | A0A0B5BJ50 | *Geobacter pickeringii* | Chemotaxis protein CheA |
| 435 | 384 | 0.01 | B3EA98 | *Geobacter lovleyi* | Transcriptional regulator, XRE family |
| 436 | 384 | 0.01 | Q39QD7 | *Geobacter metallireducens* | Methyl-accepting chemotaxis sensory transducer |
| 437 | 384 | 0.01 | Q74B89 | *Geobacter sulfurreducens* | Uncharacterized protein |
| 438 | 383 | 0.01 | A0A0D5N4L7 | *Geobacter sulfurreducens* | Outer membrane protein assembly factor BamA |
| 439 | 383 | 0.01 | A1AM25 | *Pelobacter propionicus* | Squalene-hopene cyclase |
| 440 | 383 | 0.01 | Q39WA5 | *Geobacter metallireducens* | TPR domain protein |
| 441 | 382 | 0.01 | A0A0B5BIU4 | *Geobacter pickeringii* | Probable glycine dehydrogenase (decarboxylating) subunit 1 (EC 1.4.4.2) (Glycine cleavage system P-protein subunit 1) (Glycine decarboxylase subunit 1) (Glycine dehydrogenase (aminomethyl-transferring) subunit 1) |

| | | | | | |
|---|---|---|---|---|---|
| 442 | 382 | 0.01 | A0A0B5BIW8 | *Geobacter pickeringii* | Aminotransferase |
| 443 | 382 | 0.01 | A0A1T4LT91 | *Geobacter thiogenes* | Putative transposase |
| 444 | 382 | 0.01 | Q39S32 | *Geobacter metallireducens* | DNA helicase (EC 3.6.4.12) |
| 445 | 382 | 0.01 | Q39SS9 | *Geobacter metallireducens* | Cytochrome c |
| 446 | 381 | 0.01 | A0A0B5BIH3 | *Geobacter pickeringii* | Uncharacterized protein |
| 447 | 381 | 0.01 | A1ANK6 | *Pelobacter propionicus* | Peptidase C1A, papain |
| 448 | 380 | 0.01 | A0A1T4RTR1 | *Geobacter thiogenes* | Iron complex outermembrane recepter protein |
| 449 | 380 | 0.01 | Q39RM7 | *Geobacter metallireducens* | Glycosyltransferase |
| 450 | 380 | 0.01 | Q39T88 | *Geobacter metallireducens* | Sensor histidine kinase CheA associated with MCPs of class 40H |
| 451 | 379 | 0.01 | A0A0B5BDG8 | *Geobacter pickeringii* | ABC transporter ATP-binding protein |
| 452 | 378 | 0.01 | A0A0B5B7G6 | *Geobacter pickeringii* | Cytochrome C |
| 453 | 378 | 0.01 | A0A0F2NAV5 | *Desulfobulbaceae bacterium* | Helicase |
| 454 | 378 | 0.01 | B5E8R5 | *Geobacter bemidjiensis* | Glutamine synthetase (EC 6.3.1.2) |
| 455 | 377 | 0.01 | A0A0B5BGY7 | *Geobacter pickeringii* | Phosphoglucomutase |
| 456 | 377 | 0.01 | Q39UI6 | *Geobacter metallireducens* | Type II secretion system secretin lipoprotein PulQ |
| 457 | 377 | 0.01 | Q74GF2 | *Geobacter sulfurreducens* | Metal-dependent phosphohydrolase, HDOD domain-containing |
| 458 | 376 | 0.01 | A0A0B5B942 | *Geobacter pickeringii* | Bifunctional uridylyltransferase/uridylyl-removing enzyme (UTase/UR) (Bifunctional [protein-PII] modification enzyme) (Bifunctional nitrogen sensor protein) [Includes: [Protein-PII]-UMP uridylyl-removing enzyme (UR) (EC 3.1.4.-); [Protein-PII] uridylyltransferase (PII uridylyltransferase) (UTase) (EC 2.7.7.59)] |
| 459 | 376 | 0.01 | A0A0B5BBH7 | *Geobacter pickeringii* | Endopeptidase La (EC 3.4.21.53) |
| 460 | 376 | 0.01 | A0A0B5BGI9 | *Geobacter pickeringii* | Lon protease (EC 3.4.21.53) (ATP-dependent protease La) |
| 461 | 376 | 0.01 | Q74A52 | *Geobacter sulfurreducens* | Carboxynorspermidine/carboxyspermidine dehydrogenase |
| 462 | 376 | 0.01 | Q74AR7 | *Geobacter sulfurreducens* | Membrane-associated metal-dependent phosphohydrolase, HDc domain-containing |
| 463 | 376 | 0.01 | P60401 | *Geobacter sulfurreducens* | 50S ribosomal protein L2 |
| 464 | 375 | 0.01 | A0A0B5BJG9 | *Geobacter pickeringii* | GntR family transcriptional regulator |
| 465 | 375 | 0.01 | B3EA94 | *Geobacter lovleyi* | Uncharacterized protein |
| 466 | 375 | 0.01 | Q74BF0 | *Geobacter sulfurreducens* | Phosphoribosylaminoimidazole-succinocarboxamide synthase (EC 6.3.2.6) (SAICAR synthetase) |
| 467 | 375 | 0.01 | Q74D04 | *Geobacter sulfurreducens* | Threonine--tRNA ligase (EC 6.1.1.3) (Threonyl-tRNA synthetase) (ThrRS) |
| 468 | 374 | 0.01 | A0A0D5NDV9 | *Geobacter sulfurreducens* | HD family phosphohydrolase |
| 469 | 374 | 0.01 | A0A0M3GK70 | *Rhizobium phaseoli* | Uncharacterized protein |
| 470 | 374 | 0.01 | N8YFI4 | *Acinetobacter gerneri* | Uncharacterized protein |
| 471 | 373 | 0.01 | A0A0B5B8X9 | *Geobacter pickeringii* | Phosphoenolpyruvate-protein phosphotransferase (EC 2.7.3.9) (Phosphotransferase system, enzyme I) |
| 472 | 373 | 0.01 | A0A0B5BDW0 | *Geobacter pickeringii* | Leucine--tRNA ligase (EC 6.1.1.4) (Leucyl-tRNA synthetase) (LeuRS) |

| | | | | | |
|---|---|---|---|---|---|
| 473 | 373 | 0.01 | Q39QC0 | *Geobacter metallireducens* | NADH-quinone oxidoreductase subunit N (EC 1.6.5.11) (NADH dehydrogenase I subunit N) (NDH-1 subunit N) |
| 474 | 372 | 0.01 | Q39Z96 | *Geobacter metallireducens* | S1 RNA-binding domain-containing transcriptional accessory protein |
| 475 | 371 | 0.01 | A0A0D5N8I2 | *Geobacter sulfurreducens* | Histidine kinase |
| 476 | 371 | 0.01 | A0A1G0MT57 | *Geobacteraceae bacterium* | Uncharacterized protein |
| 477 | 370 | 0.01 | A0A0B5BFH3 | *Geobacter pickeringii* | Polyribonucleotide nucleotidyltransferase (EC 2.7.7.8) (Polynucleotide phosphorylase) (PNPase) |
| 478 | 370 | 0.01 | A0A0C1TL33 | *Geobacter soli* | Uncharacterized protein |
| 479 | 370 | 0.01 | Q39WW8 | *Geobacter metallireducens* | Periplasmic serine endoprotease DegP-like (EC 3.4.21.107) |
| 480 | 369 | 0.01 | Q39RK5 | *Geobacter metallireducens* | ResB-like family cytochrome c |
| 481 | 369 | 0.01 | Q74EQ2 | *Geobacter sulfurreducens* | Aldehyde:ferredoxin oxidoreductase, tungsten-containing |
| 482 | 368 | 0.01 | A0A0A8WQH6 | *Geobacter sp. OR-1* | Uncharacterized protein |
| 483 | 368 | 0.01 | A0A0B5BI14 | *Geobacter pickeringii* | GTPase Der (GTP-binding protein EngA) |
| 484 | 368 | 0.01 | Q39QI4 | *Geobacter metallireducens* | DNA repair protein RadA |
| 485 | 368 | 0.01 | Q39QZ8 | *Geobacter metallireducens* | Flagellar M-ring protein |
| 486 | 367 | 0.01 | A0A077XNB6 | *Sphinx1.76-related DNA* | Replication protein |
| 487 | 367 | 0.01 | A0A0B5B9N3 | *Geobacter pickeringii* | 3-oxoacyl-[acyl-carrier-protein] synthase 2 (EC 2.3.1.179) |
| 488 | 367 | 0.01 | Q39W82 | *Geobacter metallireducens* | 1-deoxy-D-xylulose 5-phosphate reductoisomerase (DXP reductoisomerase) (EC 1.1.1.267) (1-deoxyxylulose-5-phosphate reductoisomerase) (2-C-methyl-D-erythritol 4-phosphate synthase) |
| 489 | 367 | 0.01 | Q39RG8 | *Geobacter metallireducens* | Methyl-accepting chemotaxis sensory transducer, class 40+24H |
| 490 | 366 | 0.01 | E1QK33 | *Desulfarculus baarsii* | ATP-binding region ATPase domain protein |
| 491 | 365 | 0.01 | A0A0B5BBF5 | *Geobacter pickeringii* | Twin-arginine translocation pathway signal protein |
| 492 | 365 | 0.01 | Q30VS6 | *Desulfovibrio alaskensis* | Site-specific DNA-methyltransferase (Adenine-specific) (EC 2.1.1.72) |
| 493 | 364 | 0.01 | A0A1G1GH24 | *Nitrospirae bacterium* | Uncharacterized protein |
| 494 | 364 | 0.01 | Q39UL3 | *Geobacter metallireducens* | Peptidoglycan-binding protein, OmpA family |
| 495 | 364 | 0.01 | Q39V80 | *Geobacter metallireducens* | Acyl-(Acyl carrier protein) ligase, acyl carrier, [acyl-]glycerolphosphate acyltransferase fusion protein |
| 496 | 364 | 0.01 | Q39ZS2 | *Geobacter metallireducens* | DNA polymerase III subunit beta (EC 2.7.7.7) |
| 497 | 364 | 0.01 | Q747L0 | *Geobacter sulfurreducens* | Glycoside hydrolase, family 57 |
| 498 | 364 | 0.01 | Q74GB8 | *Geobacter sulfurreducens* | Type II secretion system ATPase GspE |
| 499 | 364 | 0.01 | Q74H55 | *Geobacter sulfurreducens* | Serine--tRNA ligase (EC 6.1.1.11) (Seryl-tRNA synthetase) (SerRS) (Seryl-tRNA(Ser/Sec) synthetase) |
| 500 | 363 | 0.01 | A0A0B5B7W4 | *Geobacter pickeringii* | Transporter |

**Table S6b.** Functional overview of M4C community (top 500 rank only).

| Rank | Count | Abundance | ID | Organism | Protein Names |
|---|---|---|---|---|---|
| 1 | 2713 | 0.13 | | | |
| 2 | 2691 | 0.12 | Q90ZJ2 | *Oryzias latipes* | TAP2 protein (Fragment) |
| 3 | 2512 | 0.12 | | | |
| 4 | 2059 | 0.10 | | | |
| 5 | 1704 | 0.08 | A0A031IBP7 | *Micrococcus luteus* | Uncharacterized protein |
| 6 | 1366 | 0.06 | Q37953 | *Phage M13mp18* | LacZ protein (Fragment) |
| 7 | 1281 | 0.06 | | | |
| 8 | 1129 | 0.05 | N8ZVC9 | *Acinetobacter gerneri* | Uncharacterized protein |
| 9 | 979 | 0.05 | | | |
| 10 | 914 | 0.04 | N8ZNX3 | *Acinetobacter gerneri* | Uncharacterized protein |
| 11 | 812 | 0.04 | | | |
| 12 | 755 | 0.04 | | | |
| 13 | 748 | 0.03 | A0A0R3QEE4 | *Brugia timori* | Uncharacterized protein |
| 14 | 696 | 0.03 | Q8VQM8 | *Prevotella intermedia* | Mobilization protein |
| 15 | 692 | 0.03 | A0A0D5N8Q4 | *Geobacter sulfurreducens* | Transposase |
| 16 | 676 | 0.03 | | | |
| 17 | 668 | 0.03 | | | |
| 18 | 654 | 0.03 | | | |
| 19 | 640 | 0.03 | | | |
| 20 | 621 | 0.03 | A0A0D5N7W1 | *Geobacter sulfurreducens* | ATPase |
| 21 | 612 | 0.03 | Q39RM9 | *Geobacter metallireducens* | Membrane protein, putative |
| 22 | 580 | 0.03 | Q39Y12 | *Geobacter metallireducens* | DNA-directed RNA polymerase subunit beta' (RNAP subunit beta') (EC 2.7.7.6) (RNA polymerase subunit beta') (Transcriptase subunit beta') |
| 23 | 572 | 0.03 | Q748Y6 | *Geobacter sulfurreducens* | DNA-directed RNA polymerase subunit beta (RNAP subunit beta) (EC 2.7.7.6) (RNA polymerase subunit beta) (Transcriptase subunit beta) |
| 24 | 572 | 0.03 | | | |
| 25 | 570 | 0.03 | A0A077XLW5 | *Sphinx1.76-related DNA* | Replication protein |
| 26 | 532 | 0.02 | A5GAB4 | *Geobacter uraniireducens* | Putative GAF sensor protein |
| 27 | 527 | 0.02 | Q74AK2 | *Geobacter sulfurreducens* | Sodium/solute symporter family protein |
| 28 | 525 | 0.02 | | | |
| 29 | 513 | 0.02 | A0A0A8WNU3 | *Geobacter sp. OR-1* | Protein TraC |
| 30 | 508 | 0.02 | A0A090I137 | *Methanobacterium formicicum* | Transposase |
| 31 | 506 | 0.02 | Q747C7 | *Geobacter sulfurreducens* | 60 kDa chaperonin (GroEL protein) (Protein Cpn60) |

| | | | | | |
|---|---|---|---|---|---|
| 32 | 505 | 0.02 | A0LQ17 | *Syntrophobacter fumaroxidans* | Helicase domain protein |
| 33 | 468 | 0.02 | | | |
| 34 | 459 | 0.02 | A0A077XNB6 | *Sphinx1.76-related DNA* | Replication protein |
| 35 | 457 | 0.02 | A0A0T5YH86 | *Acinetobacter baumannii* | Mobilization protein A |
| 36 | 454 | 0.02 | A0A1G0M9J2 | *Geobacteraceae bacterium* | Uncharacterized protein |
| 37 | 452 | 0.02 | E1QK39 | *Desulfarculus baarsii* | Helicase domain protein |
| 38 | 447 | 0.02 | D8L7S1 | *Paramecium caudatum* | NADH-ubiquinone oxidoreductase chain 5 (EC 1.6.5.3) |
| 39 | 435 | 0.02 | | | |
| 40 | 434 | 0.02 | Q748X8 | *Geobacter sulfurreducens* | Elongation factor Tu (EF-Tu) |
| 41 | 432 | 0.02 | A0A0D5N8M0 | *Geobacter sulfurreducens* | MgtE integral membrane protein |
| 42 | 432 | 0.02 | A1AKN8 | *Pelobacter propionicus* | Resolvase, N-terminal domain protein |
| 43 | 429 | 0.02 | A0A0B5BF68 | *Geobacter pickeringii* | CRISPR-associated protein Cas3 |
| 44 | 425 | 0.02 | A0A088WWT4 | *Burkholderia oklahomensis* | Putative membrane protein |
| 45 | 420 | 0.02 | | | |
| 46 | 416 | 0.02 | F0FB90 | *Prevotella multiformis* | Initiator RepB protein (Fragment) |
| 47 | 408 | 0.02 | Q749A7 | *Geobacter sulfurreducens* | Protein translocase subunit SecY |
| 48 | 405 | 0.02 | A0A178FWP5 | *Acinetobacter sp. SFC* | Transposase |
| 49 | 405 | 0.02 | F0FB88 | *Prevotella multiformis* | Putative plasmid recombination enzyme |
| 50 | 403 | 0.02 | D8L7Q9 | *Paramecium caudatum* | Ymf66 |
| 51 | 403 | 0.02 | D8L7R1 | *Paramecium caudatum* | NADH-ubiquinone oxidoreductase chain 4 (EC 1.6.5.3) |
| 52 | 388 | 0.02 | A1AKK8 | *Pelobacter propionicus* | Uncharacterized protein |
| 53 | 388 | 0.02 | Q747H5 | *Geobacter sulfurreducens* | K(+)-insensitive pyrophosphate-energized proton pump (EC 3.6.1.1) (Membrane-bound proton-translocating pyrophosphatase) (Pyrophosphate-energized inorganic pyrophosphatase) (H(+)-PPase) |
| 54 | 387 | 0.02 | Q748Y8 | *Geobacter sulfurreducens* | Elongation factor G 2 (EF-G 2) |
| 55 | 386 | 0.02 | A0A0B5BF21 | *Geobacter pickeringii* | Glycosyl transferase family 1 |
| 56 | 382 | 0.02 | Q39Q43 | *Geobacter metallireducens* | Pyruvate-flavodoxin oxidoreductase (EC 1.2.7.-) |
| 57 | 381 | 0.02 | | | |
| 58 | 378 | 0.02 | A0A1F9PUL2 | *Desulfuromonadales bacterium* | Restriction endonuclease |
| 59 | 378 | 0.02 | Q39QB9 | *Geobacter metallireducens* | NADH dehydrogenase I, M subunit |
| 60 | 375 | 0.02 | A0A1D3L4G2 | *Methanobacterium congolense* | Uncharacterized protein |
| 61 | 370 | 0.02 | A7I7H9 | *Methanoregula boonei* | TPR repeat-containing protein |
| 62 | 368 | 0.02 | Q39U60 | *Geobacter metallireducens* | Translation elongation factor G |

| | | | | | |
|---|---|---|---|---|---|
| 63 | 362 | 0.02 | A0A0B5BJ81 | *Geobacter pickeringii* | NADH-quinone oxidoreductase subunit D (EC 1.6.5.11) (NADH dehydrogenase I subunit D) (NDH-1 subunit D) |
| 64 | 361 | 0.02 | A0A0U1CQ28 | *Chlamydia trachomatis* | Replication protein |
| 65 | 359 | 0.02 | | | |
| 66 | 354 | 0.02 | Q74FF1 | *Geobacter sulfurreducens* | Chaperone protein ClpB |
| 67 | 348 | 0.02 | Q39WU2 | *Geobacter metallireducens* | Phosphate-selective outer membrane channel |
| 68 | 348 | 0.02 | Q747X9 | *Geobacter sulfurreducens* | Isoleucine--tRNA ligase (EC 6.1.1.5) (Isoleucyl-tRNA synthetase) (IleRS) |
| 69 | 345 | 0.02 | Q39YK5 | *Geobacter metallireducens* | Flagellar biogenesis protein FlhA |
| 70 | 343 | 0.02 | A0A0B5BCT5 | *Geobacter pickeringii* | NADH dehydrogenase |
| 71 | 341 | 0.02 | B3EA87 | *Geobacter lovleyi* | RNA polymerase sigma factor |
| 72 | 339 | 0.02 | Q39QB2 | *Geobacter metallireducens* | NADH dehydrogenase I, F subunit |
| 73 | 338 | 0.02 | A0A1T4RHG2 | *Geobacter thiogenes* | Nif-specific regulatory protein |
| 74 | 338 | 0.02 | P15689 | *Paramecium tetraurelia* | NADH-ubiquinone oxidoreductase 49 kDa subunit (EC 1.6.5.3) (EC 1.6.99.3) (NADH dehydrogenase subunit 7) |
| 75 | 338 | 0.02 | Q39ZC8 | *Geobacter metallireducens* | Ferredoxin-dependent glutamate synthase |
| 76 | 335 | 0.02 | N8YFI4 | *Acinetobacter gerneri* | Uncharacterized protein |
| 77 | 332 | 0.02 | A0A0L8ETQ7 | *Achromobacter sp. DMS1* | Uncharacterized protein |
| 78 | 332 | 0.02 | | | |
| 79 | 330 | 0.02 | A0A0B5B7H3 | *Geobacter pickeringii* | Cytochrome C |
| 80 | 328 | 0.02 | A0A1S0XXD8 | *Rhodanobacter sp.* | Histidine kinase |
| 81 | 326 | 0.02 | | | |
| 82 | 324 | 0.02 | A7I4I9 | *Methanoregula boonei* | DNA polymerase II large subunit (Pol II) (EC 2.7.7.7) |
| 83 | 324 | 0.02 | Q74FK9 | *Geobacter sulfurreducens* | Sigma-54-dependent transcriptional response regulator |
| 84 | 323 | 0.01 | Q74C76 | *Geobacter sulfurreducens* | (R)-citramalate synthase (EC 2.3.1.182) (Citramalate synthase) |
| 85 | 323 | 0.01 | | | |
| 86 | 321 | 0.01 | A0A0D5NDJ8 | *Geobacter sulfurreducens* | Stage II sporulation protein E |
| 87 | 319 | 0.01 | A0A0D5N8Q2 | *Geobacter sulfurreducens* | Uncharacterized protein |
| 88 | 319 | 0.01 | Q74FM6 | *Geobacter sulfurreducens* | Pyruvate, phosphate dikinase (EC 2.7.9.1) |
| 89 | 317 | 0.01 | E1QK33 | *Desulfarculus baarsii* | ATP-binding region ATPase domain protein |
| 90 | 316 | 0.01 | A0A1G0MAK9 | *Geobacteraceae bacterium* | Sodium:proton exchanger |
| 91 | 316 | 0.01 | Q39QD0 | *Geobacter metallireducens* | Periplasmically oriented, membrane-bound [NiFe]-hydrogenase, large subunit |
| 92 | 315 | 0.01 | A5GFJ7 | *Geobacter uraniireducens* | Erythronolide synthase (EC 2.3.1.94) |
| 93 | 315 | 0.01 | A7IAM2 | *Methanoregula boonei* | Isoleucine--tRNA ligase (EC 6.1.1.5) (Isoleucyl-tRNA synthetase) (IleRS) |
| 94 | 314 | 0.01 | B3EB58 | *Geobacter lovleyi* | Uncharacterized protein |
| 95 | 310 | 0.01 | | | |

| | | | | | |
|---|---|---|---|---|---|
| 96 | 306 | 0.01 | D4H381 | *Denitrovibrio acetiphilus* | DNA-directed RNA polymerase subunit beta (RNAP subunit beta) (EC 2.7.7.6) (RNA polymerase subunit beta) (Transcriptase subunit beta) |
| 97 | 305 | 0.01 | Q74FD5 | *Geobacter sulfurreducens* | Hydroxylamine reductase (EC 1.7.99.1) (Hybrid-cluster protein) (HCP) (Prismane protein) |
| 98 | 303 | 0.01 | A0A0U1CQ01 | *Chlamydia trachomatis* | Uncharacterised protein |
| 99 | 303 | 0.01 | Q39RM8 | *Geobacter metallireducens* | Glycosyltransferase |
| 100 | 303 | 0.01 | Q39ZE7 | *Geobacter metallireducens* | UvrABC system protein A (UvrA protein) (Excinuclease ABC subunit A) |
| 101 | 303 | 0.01 | | | |
| 102 | 302 | 0.01 | A0A0D5NE26 | *Geobacter sulfurreducens* | Multidrug resistance protein B |
| 103 | 300 | 0.01 | A0A0B5B8A8 | *Geobacter pickeringii* | Valine--tRNA ligase (EC 6.1.1.9) (Valyl-tRNA synthetase) (ValRS) |
| 104 | 299 | 0.01 | Q74H82 | *Geobacter sulfurreducens* | Flagellar biogenesis master response receiver sensor histidine kinase, PAS and GAF domain-containing |
| 105 | 298 | 0.01 | A0A0D5NDD6 | *Geobacter sulfurreducens* | Type II secretion system protein E |
| 106 | 296 | 0.01 | D4H382 | *Denitrovibrio acetiphilus* | DNA-directed RNA polymerase subunit beta' (RNAP subunit beta') (EC 2.7.7.6) (RNA polymerase subunit beta') (Transcriptase subunit beta') |
| 107 | 295 | 0.01 | A0A1G0M577 | *Geobacteraceae bacterium* | Uncharacterized protein |
| 108 | 294 | 0.01 | Q74FW2 | *Geobacter sulfurreducens* | Succinyl:acetate coenzyme A transferase |
| 109 | 293 | 0.01 | A0A0B5BJ46 | *Geobacter pickeringii* | Uncharacterized protein |
| 110 | 292 | 0.01 | A0A0B5BB65 | *Geobacter pickeringii* | Uncharacterized protein |
| 111 | 292 | 0.01 | E8WU24 | *Geobacter sp.* | Helicase domain protein |
| 112 | 292 | 0.01 | Q39Y76 | *Geobacter metallireducens* | Sigma-54-dependent transcriptional response regulator |
| 113 | 290 | 0.01 | Q74H58 | *Geobacter sulfurreducens* | Chaperone protein DnaJ |
| 114 | 287 | 0.01 | A0A0B5BI89 | *Geobacter pickeringii* | Type II secretion system protein |
| 115 | 287 | 0.01 | Q746X3 | *Geobacter sulfurreducens* | Bifunctional protein PutA |
| 116 | 287 | 0.01 | Q748Z4 | *Geobacter sulfurreducens* | 30S ribosomal protein S3 |
| 117 | 286 | 0.01 | A0A0D5N712 | *Geobacter sulfurreducens* | Phosphoglucomutase |
| 118 | 286 | 0.01 | A0A1G0LE89 | *Geobacteraceae bacterium* | Aconitate hydratase (Aconitase) (EC 4.2.1.3) |
| 119 | 286 | 0.01 | | | |
| 120 | 283 | 0.01 | A0A0B5BDJ5 | *Geobacter pickeringii* | NADH-quinone oxidoreductase subunit H (EC 1.6.5.11) (NADH dehydrogenase I subunit H) (NDH-1 subunit H) |
| 121 | 282 | 0.01 | A0A0B5BH93 | *Geobacter pickeringii* | C-and b-type cytochrome L |
| 122 | 282 | 0.01 | A0A0C1TL30 | *Geobacter soli* | Flagellar basal body stator protein MotB |
| 123 | 282 | 0.01 | F3ZQL8 | *Bacteroides coprosuis* | Transposase IS4 family protein |
| 124 | 281 | 0.01 | | | |
| 125 | 280 | 0.01 | Q39T74 | *Geobacter metallireducens* | GTP/GDP 3'-pyrophosphokinase and (P)ppGpp 3'-pyrophosphohydrolase |
| 126 | 279 | 0.01 | A0A0D5N8G6 | *Geobacter sulfurreducens* | Cation transporter |

| | | | | | |
|---|---|---|---|---|---|
| 127 | 278 | 0.01 | Q39QJ8 | *Geobacter metallireducens* | Flagellar biogenesis master sigma-54-dependent transcriptional response regulator |
| 128 | 277 | 0.01 | F4BUN6 | *Methanosaeta concilii* | Transposase, IS116/IS110/IS902 family |
| 129 | 277 | 0.01 | Q8VQM9 | *Prevotella intermedia* | Replication protein |
| 130 | 276 | 0.01 | A0A0B5BAC1 | *Geobacter pickeringii* | Dihydrolipoyl dehydrogenase (EC 1.8.1.4) |
| 131 | 276 | 0.01 | A0A0D5N8I5 | *Geobacter sulfurreducens* | Signal recognition particle protein (Fifty-four homolog) |
| 132 | 273 | 0.01 | D8L7S0 | *Paramecium caudatum* | Cytochrome c oxidase subunit 2 |
| 133 | 273 | 0.01 | Q747M6 | *Geobacter sulfurreducens* | Ribonuclease G |
| 134 | 273 | 0.01 | | | |
| 135 | 272 | 0.01 | A0A0B5BDG0 | *Geobacter pickeringii* | ATP-dependent Clp protease proteolytic subunit (EC 3.4.21.92) (Endopeptidase Clp) |
| 136 | 272 | 0.01 | A0A0D5N8S3 | *Geobacter sulfurreducens* | NADH dehydrogenase |
| 137 | 272 | 0.01 | B5EFV6 | *Geobacter bemidjiensis* | Bifunctional protein PutA |
| 138 | 271 | 0.01 | A0A1Q9F786 | *Symbiodinium microadriaticum* | Uncharacterized protein |
| 139 | 270 | 0.01 | Q74B77 | *Geobacter sulfurreducens* | Uncharacterized protein |
| 140 | 269 | 0.01 | | | |
| 141 | 267 | 0.01 | A0A0B5BIS9 | *Geobacter pickeringii* | Multidrug transporter AcrB |
| 142 | 267 | 0.01 | Q39ZH1 | *Geobacter metallireducens* | Glutamine--fructose-6-phosphate aminotransferase [isomerizing] (EC 2.6.1.16) (D-fructose-6-phosphate amidotransferase) (GFAT) (Glucosamine-6-phosphate synthase) (Hexosephosphate aminotransferase) (L-glutamine--D-fructose-6-phosphate amidotransferase) |
| 143 | 267 | 0.01 | Q39ZK4 | *Geobacter metallireducens* | Bifunctional glutamine synthetase adenylyltransferase/adenylyl-removing enzyme (ATP:glutamine synthetase adenylyltransferase) (ATase) [Includes: Glutamine synthetase adenylyl-L-tyrosine phosphorylase (EC 2.7.7.89) (Adenylyl removase) (AR) (AT-N); Glutamine synthetase adenylyl transferase (EC 2.7.7.42) (Adenylyl transferase) (AT) (AT-C)] |
| 144 | 266 | 0.01 | A0A0D5N8E2 | *Geobacter sulfurreducens* | ATP synthase (EC 3.6.3.14) |
| 145 | 265 | 0.01 | A0A0C1U608 | *Geobacter soli* | Chemotaxis protein |
| 146 | 265 | 0.01 | A0A0D5ND20 | *Geobacter sulfurreducens* | Amino acid dehydrogenase |
| 147 | 265 | 0.01 | A0A0D5ND99 | *Geobacter sulfurreducens* | ATPase AAA |
| 148 | 265 | 0.01 | B3E9R4 | *Geobacter lovleyi* | Isocitrate dehydrogenase, NADP-dependent (EC 1.1.1.42) |
| 149 | 265 | 0.01 | | | |
| 150 | 262 | 0.01 | A0A0D5NE46 | *Geobacter sulfurreducens* | Molecular chaperone DnaJ |
| 151 | 262 | 0.01 | N8QHE7 | *Acinetobacter bohemicus* | Uncharacterized protein |
| 152 | 262 | 0.01 | Q39QB8 | *Geobacter metallireducens* | NADH dehydrogenase I, L subunit |
| 153 | 260 | 0.01 | Q39Q36 | *Geobacter metallireducens* | Heterodisulfide oxidoreductase, FAD-binding and iron-sulfur cluster-binding subunit A |
| 154 | 259 | 0.01 | A0A0B5BK17 | *Geobacter pickeringii* | Diguanylate cyclase |

| | | | | | |
|---|---|---|---|---|---|
| 155 | 259 | 0.01 | Q39XL9 | *Geobacter metallireducens* | Sensor sigma-54-dependent transcriptional regulator, GAF and GAF domain-containing |
| 156 | 257 | 0.01 | Q39RL8 | *Geobacter metallireducens* | Undecaprenyl-diphospho-oligosaccharide flippase |
| 157 | 257 | 0.01 | Q747G2 | *Geobacter sulfurreducens* | Outer membrane channel OmpJ |
| 158 | 256 | 0.01 | A5GB28 | *Geobacter sulfurreducens* | Plasmid maintenance system antidote protein, XRE family |
| 159 | 256 | 0.01 | Q74AK3 | *Geobacter sulfurreducens* | Cation-translocating P-type ATPase |
| 160 | 254 | 0.01 | Q39PS2 | *Geobacter metallireducens* | Transcription-repair-coupling factor (TRCF) (EC 3.6.4.-) |
| 161 | 253 | 0.01 | A0A0B5BH05 | *Geobacter pickeringii* | Malic enzyme (EC 1.1.1.40) |
| 162 | 253 | 0.01 | Q749N5 | *Geobacter sulfurreducens* | Acetate kinase (EC 2.7.2.1) (Acetokinase) |
| 163 | 253 | 0.01 | Q39V52 | *Geobacter metallireducens* | GAF sensor methyl-accepting chemotaxis sensory transducer, class 40H |
| 164 | 250 | 0.01 | A7IAV3 | *Methanoregula boonei* | V-type ATP synthase subunit I |
| 165 | 250 | 0.01 | Q74BC2 | *Geobacter sulfurreducens* | Integrative genetic element Gsu56, integrase |
| 166 | 249 | 0.01 | A0A0C5IZ87 | *Rugosibacter aromaticivorans* | Uncharacterized protein |
| 167 | 248 | 0.01 | A0A0A8WPS5 | *Geobacter sp.* | Efflux pump membrane transporter |
| 168 | 248 | 0.01 | A0A0D5N849 | *Geobacter sulfurreducens* | DNA polymerase III subunit gamma/tau (EC 2.7.7.7) |
| 169 | 248 | 0.01 | A5GFJ6 | *Geobacter uraniireducens* | Beta-hydroxyacyl-(Acyl-carrier-protein) dehydratase, FabA/FabZ |
| 170 | 248 | 0.01 | A7IAD8 | *Methanoregula boonei* | Chromosome partition protein Smc |
| 171 | 248 | 0.01 | Q74GY2 | *Geobacter sulfurreducens* | ATP synthase subunit alpha (EC 3.6.3.14) (ATP synthase F1 sector subunit alpha) (F-ATPase subunit alpha) |
| 172 | 248 | 0.01 | Q39UR9 | *Geobacter metallireducens* | Carbamoyl-phosphate synthase large chain (EC 6.3.5.5) (Carbamoyl-phosphate synthetase ammonia chain) |
| 173 | 248 | 0.01 | | | |
| 174 | 246 | 0.01 | A0A0B5BDE2 | *Geobacter pickeringii* | ATP phosphoribosyltransferase regulatory subunit |
| 175 | 246 | 0.01 | Q74GG6 | *Geobacter sulfurreducens* | Chromosomal replication initiator protein DnaA |
| 176 | 245 | 0.01 | A0A0D5N5P7 | *Geobacter sulfurreducens* | Chemotaxis protein |
| 177 | 244 | 0.01 | A7I9N6 | *Methanoregula boonei* | DNA-directed RNA polymerase subunit (EC 2.7.7.6) |
| 178 | 244 | 0.01 | F5UKT4 | *Microcoleus vaginatus* | Polymorphic outer membrane protein |
| 179 | 244 | 0.01 | Q39QP7 | *Geobacter metallireducens* | Lon protease (EC 3.4.21.53) (ATP-dependent protease La) |
| 180 | 244 | 0.01 | Q39YH3 | *Geobacter metallireducens* | UDP-N-acetylglucosamine 4,6-dehydratase and UDP-2-acetamido-2,6-dideoxy-alpha-D-xylo-4-hexulose 5-epimerase |
| 181 | 243 | 0.01 | A0A0B5B9H5 | *Geobacter pickeringii* | Uncharacterized protein |
| 182 | 243 | 0.01 | A1AM25 | *Pelobacter propionicus* | Squalene-hopene cyclase |
| 183 | 243 | 0.01 | | | |
| 184 | 242 | 0.01 | A0A0B5BD43 | *Geobacter pickeringii* | ABC transporter permease |

| | | | | | |
|---|---|---|---|---|---|
| 185 | 242 | 0.01 | A7I486 | *Methanoregula boonei* | K(+)-insensitive pyrophosphate-energized proton pump (EC 3.6.1.1) (Membrane-bound proton-translocating pyrophosphatase) (Pyrophosphate-energized inorganic pyrophosphatase) (H(+)-PPase) |
| 186 | 241 | 0.01 | A5GDX4 | *Geobacter uraniireducens* | DNA gyrase subunit B (EC 5.99.1.3) |
| 187 | 240 | 0.01 | B9M4C2 | *Geobacter daltonii* | Squalene cyclase |
| 188 | 240 | 0.01 | E1QK40 | *Desulfarculus baarsii* | DNA-cytosine methyltransferase |
| 189 | 240 | 0.01 | Q746S4 | *Geobacter sulfurreducens* | NADH-quinone oxidoreductase subunit B/C/D (EC 1.6.5.11) (NADH dehydrogenase I subunit B/C/D) (NDH-1 subunit B/C/D) |
| 190 | 239 | 0.01 | A0A0D5N8X4 | *Geobacter sulfurreducens* | Flagellar M-ring protein |
| 191 | 239 | 0.01 | E6VQ93 | *Rhodopseudomonas palustris* | Putative methylase/helicase |
| 192 | 238 | 0.01 | A0A1F8XVG3 | *Deltaproteobacteria bacterium* | Helicase |
| 193 | 238 | 0.01 | A5GF11 | *Geobacter uraniireducens* | Acetyl-CoA carboxylase carboxyltransferase subunit alpha (EC 6.3.4.14) |
| 194 | 238 | 0.01 | N8VN75 | *Acinetobacter sp.* | Uncharacterized protein |
| 195 | 237 | 0.01 | A0A0D5NAK5 | *Geobacter sulfurreducens* | Bifunctional uridylyltransferase/uridylyl-removing enzyme (UTase/UR) (Bifunctional [protein-PII] modification enzyme) (Bifunctional nitrogen sensor protein) [Includes: [Protein-PII]-UMP uridylyl-removing enzyme (UR) (EC 3.1.4.-); [Protein-PII] uridylyltransferase (PII uridylyltransferase) (UTase) (EC 2.7.7.59)] |
| 196 | 237 | 0.01 | A0R7X1 | *Pelobacter propionicus* | Type III restriction enzyme, res subunit |
| 197 | 237 | 0.01 | Q39R25 | *Geobacter metallireducens* | Sensor diguanylate cyclase/phosphodiesterase, PAS, PAS and PAS domain-containing |
| 198 | 236 | 0.01 | A0A0B5BGI9 | *Geobacter pickeringii* | Lon protease (EC 3.4.21.53) (ATP-dependent protease La) |
| 199 | 236 | 0.01 | A0A0B5BK56 | *Geobacter pickeringii* | Lon protease (EC 3.4.21.53) (ATP-dependent protease La) |
| 200 | 235 | 0.01 | D2VLN5 | *Naegleria gruberi (Amoeba)* | Predicted protein (Fragment) |
| 201 | 235 | 0.01 | | | |
| 202 | 234 | 0.01 | P61679 | *Geobacter sulfurreducens* | UDP-N-acetylmuramate--L-alanine ligase (EC 6.3.2.8) (UDP-N-acetylmuramoyl-L-alanine synthetase) |
| 203 | 234 | 0.01 | P61667 | *Geobacter sulfurreducens* | DNA mismatch repair protein MutS |
| 204 | 233 | 0.01 | A0A0D5NAW4 | *Geobacter sulfurreducens* | Cation transporter |
| 205 | 232 | 0.01 | A0A0B5BC86 | *Geobacter pickeringii* | Multicopper oxidase |
| 206 | 232 | 0.01 | A0A0B5BK89 | *Geobacter pickeringii* | FAD-binding protein |
| 207 | 232 | 0.01 | Q74F57 | *Geobacter sulfurreducens* | Membrane protein, major facilitator superfamily |
| 208 | 230 | 0.01 | Q74GY0 | *Geobacter sulfurreducens* | ATP synthase subunit beta (EC 3.6.3.14) (ATP synthase F1 sector subunit beta) (F-ATPase subunit beta) |
| 209 | 230 | 0.01 | Q39QJ5 | *Geobacter metallireducens* | Sensor histidine kinase CheA associated with MCPs of classes 40H and 40+24H |
| 210 | 229 | 0.01 | A0A0B5BDW0 | *Geobacter pickeringii* | Leucine--tRNA ligase (EC 6.1.1.4) (Leucyl-tRNA synthetase) (LeuRS) |

| | | | | | |
|---|---|---|---|---|---|
| 211 | 229 | 0.01 | A7IAB8 | *Methanoregula boonei* | Phosphoenolpyruvate synthase (PEP synthase) (EC 2.7.9.2) (Pyruvate, water dikinase) |
| 212 | 229 | 0.01 | B9M5J7 | *Geobacter daltonii* | CRISPR-associated protein Csd2 |
| 213 | 229 | 0.01 | Q39QY3 | *Geobacter metallireducens* | Efflux pump, RND family, inner membrane protein |
| 214 | 228 | 0.01 | A0A0B5B668 | *Geobacter pickeringii* | Peptidase U32 |
| 215 | 228 | 0.01 | Q39QU2 | *Geobacter metallireducens* | Phosphoenolpyruvate carboxykinase (ATP) (PCK) (PEP carboxykinase) (PEPCK) (EC 4.1.1.49) |
| 216 | 228 | 0.01 | Q39X91 | *Geobacter metallireducens* | DNA topoisomerase 1 (EC 5.99.1.2) (DNA topoisomerase I) |
| 217 | 227 | 0.01 | A0A0D5N6Q1 | *Geobacter sulfurreducens* | FeS-binding protein |
| 218 | 227 | 0.01 | A0A0L8EU90 | *Achromobacter sp. DMS1* | Uncharacterized protein |
| 219 | 227 | 0.01 | A7I927 | *Methanoregula boonei* | Valine--tRNA ligase (EC 6.1.1.9) (Valyl-tRNA synthetase) (ValRS) |
| 220 | 227 | 0.01 | B9M1D8 | *Geobacter daltonii* | Cyclic diguanylate phosphodiesterase |
| 221 | 226 | 0.01 | A0A0B5BBC9 | *Geobacter pickeringii* | Type I citrate synthase (EC 2.3.3.1) |
| 222 | 226 | 0.01 | A0A0D5N2S0 | *Geobacter sulfurreducens* | Uncharacterized protein |
| 223 | 226 | 0.01 | A0A1F9PXI1 | *Desulfuromonadales bacterium* | Cytosine-specific methyltransferase (EC 2.1.1.37) |
| 224 | 226 | 0.01 | A5G9A7 | *Geobacter uraniireducens* | NADH dehydrogenase subunit M (EC 1.6.5.3) |
| 225 | 226 | 0.01 | B5EGM4 | *Geobacter bemidjiensis* | Glycerol/1,2-propanediol dehydratase |
| 226 | 225 | 0.01 | A0A0B4XMH5 | *Alcanivorax pacificus* | Adenine-specific DNA methylase containing a Zn-ribbon |
| 227 | 225 | 0.01 | Q74C83 | *Geobacter sulfurreducens* | ATP-dependent Clp protease ATP-binding subunit ClpX |
| 228 | 225 | 0.01 | E1QK35 | *Desulfarculus baarsii* | Uncharacterized protein |
| 229 | 225 | 0.01 | Q74BB9 | *Geobacter sulfurreducens* | TraG family protein |
| 230 | 224 | 0.01 | A0A0B5BIQ7 | *Geobacter pickeringii* | Flagellar basal-body rod protein FlgF (Flagellar hook protein FlgE) |
| 231 | 224 | 0.01 | A5G7V6 | *Geobacter uraniireducens* | Dihydroxy-acid dehydratase (DAD) (EC 4.2.1.9) |
| 232 | 223 | 0.01 | A0A0D5NCW1 | *Geobacter sulfurreducens* | Histidine kinase |
| 233 | 223 | 0.01 | Q39Q79 | *Geobacter metallireducens* | Winged-helix transcriptional response regulator |
| 234 | 221 | 0.01 | A0A0B5BCL9 | *Geobacter pickeringii* | Uncharacterized protein |
| 235 | 221 | 0.01 | A7I9M8 | *Methanoregula boonei* | Leucine--tRNA ligase (EC 6.1.1.4) (Leucyl-tRNA synthetase) (LeuRS) |
| 236 | 221 | 0.01 | A7I4X4 | *Methanoregula boonei* | Elongation factor 2 (EF-2) |
| 237 | 221 | 0.01 | N8YF58 | *Acinetobacter gerneri* | Uncharacterized protein |
| 238 | 221 | 0.01 | Q39WG2 | *Geobacter metallireducens* | Chromosome partition protein Smc |
| 239 | 221 | 0.01 | | | |
| 240 | 220 | 0.01 | A0A0B5BCB2 | *Geobacter pickeringii* | Nitrogen fixation protein NifE |
| 241 | 220 | 0.01 | A0A0B5BCZ8 | *Geobacter pickeringii* | Succinate dehydrogenase (EC 1.3.5.1) |
| 242 | 220 | 0.01 | A0A0B5BJ26 | *Geobacter pickeringii* | Uncharacterized protein |

| | | | | | |
|---|---|---|---|---|---|
| 243 | 220 | 0.01 | A0A0D5N7I1 | *Geobacter sulfurreducens* | NADH-quinone oxidoreductase subunit N (EC 1.6.5.11) (NADH dehydrogenase I subunit N) (NDH-1 subunit N) |
| 244 | 220 | 0.01 | Q39U54 | *Geobacter metallireducens* | Sensor histidine kinase cyclic nucleotide phosphodiesterase, GAF, GAF and HD-GYP-related domain-containing |
| 245 | 220 | 0.01 | Q74AJ5 | *Geobacter sulfurreducens* | Glycoside hydrolase, family 57, DUF3536 domain-containing |
| 246 | 220 | 0.01 | A5GAY2 | *Geobacter uraniireducens* | DNA-directed RNA polymerase subunit beta' (RNAP subunit beta') (EC 2.7.7.6) (RNA polymerase subunit beta') (Transcriptase subunit beta') |
| 247 | 219 | 0.01 | A0A0B5BIF0 | *Geobacter pickeringii* | NADH-ubiquinone oxidoreductase subunit 3 |
| 248 | 219 | 0.01 | A0A1G0LEY2 | *Geobacteraceae bacterium* | Uncharacterized protein |
| 249 | 219 | 0.01 | A0A1T4LST4 | *Geobacter thiogenes* | Multidrug efflux pump subunit AcrB |
| 250 | 219 | 0.01 | A7I6J1 | *Methanoregula boonei* | Carbamoyl-phosphate synthase large chain (EC 6.3.5.5) (Carbamoyl-phosphate synthetase ammonia chain) |
| 251 | 219 | 0.01 | S0G1I5 | *Desulfotignum phosphitoxidans* | Type I restriction enzyme HindVIIP/EcoKI protein HsdR (EC 3.1.21.3) |
| 252 | 218 | 0.01 | A0A0B5BKE7 | *Geobacter pickeringii* | NADH-quinone oxidoreductase subunit N (EC 1.6.5.11) (NADH dehydrogenase I subunit N) (NDH-1 subunit N) |
| 253 | 218 | 0.01 | A0A0D5N7N0 | *Geobacter sulfurreducens* | Glutamate synthase |
| 254 | 218 | 0.01 | A0A154DNE7 | *Acinetobacter baumannii* | mRNA endoribonuclease LS (EC 3.1.-.-) |
| 255 | 218 | 0.01 | A5GCN0 | *Geobacter uraniireducens* | Short chain amide porin |
| 256 | 217 | 0.01 | A0A0B5BG42 | *Geobacter pickeringii* | Vitamin B12-dependent ribonucleotide reductase (EC 1.17.4.1) |
| 257 | 217 | 0.01 | A0A0B5BHV9 | *Geobacter pickeringii* | ATPase AAA |
| 258 | 217 | 0.01 | A0A0D5N400 | *Geobacter sulfurreducens* | Uncharacterized protein |
| 259 | 217 | 0.01 | A0A1T4RGQ7 | *Geobacter thiogenes* | NAD+---dinitrogen-reductase ADP-D-ribosyltransferase |
| 260 | 217 | 0.01 | Q39X64 | *Geobacter metallireducens* | Cell shape-determining protein MreB |
| 261 | 217 | 0.01 | A5GAY1 | *Geobacter uraniireducens* | DNA-directed RNA polymerase subunit beta (RNAP subunit beta) (EC 2.7.7.6) (RNA polymerase subunit beta) (Transcriptase subunit beta) |
| 262 | 216 | 0.01 | A0A0C1TL33 | *Geobacter soli* | Uncharacterized protein |
| 263 | 216 | 0.01 | A7I485 | *Methanoregula boonei* | Vitamin B12-dependent ribonucleotide reductase (EC 1.17.4.1) |
| 264 | 216 | 0.01 | A7I791 | *Methanoregula boonei* | UvrABC system protein A (UvrA protein) (Excinuclease ABC subunit A) |
| 265 | 216 | 0.01 | Q74CR5 | *Geobacter sulfurreducens* | Serine hydroxymethyltransferase (SHMT) (Serine methylase) (EC 2.1.2.1) |
| 266 | 216 | 0.01 | Q39SJ2 | *Geobacter metallireducens* | Nitrogen fixation master sigma-54-dependent transcriptional response regulator |
| 267 | 215 | 0.01 | A0A0D5N7R7 | *Geobacter sulfurreducens* | GntR family transcriptional regulator |
| 268 | 215 | 0.01 | Q39RM6 | *Geobacter metallireducens* | Glycosyltransferase |
| 269 | 215 | 0.01 | Q39SI5 | *Geobacter metallireducens* | Fumarate hydratase class I (EC 4.2.1.2) |
| 270 | 215 | 0.01 | Q39X99 | *Geobacter metallireducens* | UDP-glucuronate 4-dehydrogenase, decarboxylating |
| 271 | 214 | 0.01 | A0A0B5BDE7 | *Geobacter pickeringii* | Fis family transcriptional regulator |

| | | | | | |
|---|---|---|---|---|---|
| 272 | 214 | 0.01 | B9M1Q4 | *Geobacter daltonii* | Biotin-dependent acyl-CoA carboxylase, biotin carboxylase subunit |
| 273 | 214 | 0.01 | Q74C70 | *Geobacter sulfurreducens* | Phosphoglucosamine mutase (EC 5.4.2.10) |
| 274 | 214 | 0.01 | Q39Y82 | *Geobacter metallireducens* | 5-methyltetrahydrofolate--homocysteine S-methyltransferase, cobalamin-dependent |
| 275 | 214 | 0.01 | Q74F32 | *Geobacter sulfurreducens* | Periplasmically oriented, membrane-bound formate dehydrogenase, major subunit, selenocysteine-containing |
| 276 | 214 | 0.01 | A0A4W3 | *Geobacter sulfurreducens* | Biosynthetic arginine decarboxylase (ADC) (EC 4.1.1.19) |
| 277 | 214 | 0.01 | | | |
| 278 | 213 | 0.01 | A0A0B5BIA1 | *Geobacter pickeringii* | DNA-directed DNA polymerase (EC 2.7.7.7) |
| 279 | 213 | 0.01 | B9LZR8 | *Geobacter daltonii* | Succinyl:acetate coenzyme A transferase |
| 280 | 212 | 0.01 | A0A0B5BE38 | *Geobacter pickeringii* | Transcription-repair-coupling factor (TRCF) (EC 3.6.4.-) |
| 281 | 212 | 0.01 | A0A0D5N419 | *Geobacter sulfurreducens* | Uncharacterized protein |
| 282 | 212 | 0.01 | A1ANK6 | *Pelobacter propionicus* | Peptidase C1A, papain |
| 283 | 212 | 0.01 | Q39YR4 | *Geobacter metallireducens* | FtsK/SpoIIIE domain protein |
| 284 | 210 | 0.01 | A0A0C1QTJ3 | *Geobacter soli* | Hemolysin D |
| 285 | 210 | 0.01 | Q74EQ2 | *Geobacter sulfurreducens* | Aldehyde:ferredoxin oxidoreductase, tungsten-containing |
| 286 | 209 | 0.01 | Q748I6 | *Geobacter sulfurreducens* | Metalloprotease domain protein, M6 family |
| 287 | 208 | 0.01 | A0A0A8WNZ4 | *Geobacter sp. OR-1* | DNA gyrase subunit A (EC 5.99.1.3) |
| 288 | 208 | 0.01 | A0A0B5B8W1 | *Geobacter pickeringii* | ATP-dependent zinc metalloprotease FtsH (EC 3.4.24.-) |
| 289 | 208 | 0.01 | B3EB49 | *Geobacter lovleyi* | Uncharacterized protein |
| 290 | 208 | 0.01 | I9R3U4 | *Bacteroides cellulosilyticus* | Uncharacterized protein |
| 291 | 208 | 0.01 | Q39QK7 | *Geobacter metallireducens* | Outer membrane channel OmpJ |
| 292 | 208 | 0.01 | Q39ZQ2 | *Geobacter metallireducens* | Uncharacterized protein |
| 293 | 207 | 0.01 | A0A0A8WQV9 | *Geobacter sp.* | Protein TraU |
| 294 | 206 | 0.01 | A0A0B5BC52 | *Geobacter pickeringii* | General secretion pathway protein GspD |
| 295 | 206 | 0.01 | A0A0D5N732 | *Geobacter sulfurreducens* | Transcription-repair-coupling factor (TRCF) (EC 3.6.4.-) |
| 296 | 206 | 0.01 | A7I8Y5 | *Methanoregula boonei* | DEAD/DEAH box helicase domain protein |
| 297 | 206 | 0.01 | Q39QT7 | *Geobacter metallireducens* | Cytochrome c |
| 298 | 205 | 0.01 | A0A0B5BBF5 | *Geobacter pickeringii* | Twin-arginine translocation pathway signal protein |
| 299 | 205 | 0.01 | A0A0B5BCI8 | *Geobacter pickeringii* | Galactose-1-phosphate uridylyltransferase |
| 300 | 205 | 0.01 | A0A0E9G9S8 | *Chlamydia trachomatis* | Uncharacterised protein |
| 301 | 205 | 0.01 | P60789 | *Geobacter sulfurreducens* | Elongation factor 4 (EF-4) (EC 3.6.5.n1) (Ribosomal back-translocase LepA) |
| 302 | 205 | 0.01 | P05513 | *Paramecium tetraurelia* | NADH-ubiquinone oxidoreductase chain 1 (EC 1.6.5.3) (NADH dehydrogenase subunit 1) |
| 303 | 205 | 0.01 | Q39XG6 | *Geobacter metallireducens* | Pyruvate carboxylase (EC 6.4.1.1) |
| 304 | 205 | 0.01 | Q747L0 | *Geobacter sulfurreducens* | Glycoside hydrolase, family 57 |

| | | | | | |
|---|---|---|---|---|---|
| 305 | 204 | 0.01 | Q39RM7 | *Geobacter metallireducens* | Glycosyltransferase |
| 306 | 203 | 0.01 | Q39RQ5 | *Geobacter metallireducens* | Chaperone protein ClpB |
| 307 | 203 | 0.01 | | | |
| 308 | 202 | 0.01 | A0A0B5B822 | *Geobacter pickeringii* | Formate dehydrogenase |
| 309 | 202 | 0.01 | Q39RI3 | *Geobacter metallireducens* | Sensor histidine kinase, HAMP domain-containing, 2 heme-binding sites |
| 310 | 201 | 0.01 | A0A0D5N7M4 | *Geobacter sulfurreducens* | Chemotaxis protein |
| 311 | 201 | 0.01 | A0A0D5N8K2 | *Geobacter sulfurreducens* | RND transporter |
| 312 | 201 | 0.01 | A0A1G0MT57 | *Geobacteraceae bacterium* | Uncharacterized protein |
| 313 | 201 | 0.01 | A7I842 | *Methanoregula boonei* | Beta-lactamase domain protein |
| 314 | 201 | 0.01 | Q74AT0 | *Geobacter sulfurreducens* | Lysine--tRNA ligase (EC 6.1.1.6) (Lysyl-tRNA synthetase) (LysRS) |
| 315 | 201 | 0.01 | Q74B80 | *Geobacter sulfurreducens* | Uncharacterized protein |
| 316 | 201 | 0.01 | | | |
| 317 | 200 | 0.01 | A0A0A8WN28 | *Geobacter sp. OR-1* | Protein TraN |
| 318 | 200 | 0.01 | A0A0D5NB12 | *Geobacter sulfurreducens* | Aldehyde dehydrogenase |
| 319 | 200 | 0.01 | Q74BB8 | *Geobacter sulfurreducens* | Metal-dependent phosphohydrolase |
| 320 | 199 | 0.01 | A0A0B5BCH1 | *Geobacter pickeringii* | Hydrogenase 2 large subunit (EC 1.12.99.6) |
| 321 | 199 | 0.01 | A7I7B5 | *Methanoregula boonei* | DNA polymerase (EC 2.7.7.7) |
| 322 | 199 | 0.01 | Q74GT3 | *Geobacter sulfurreducens* | LL-diaminopimelate aminotransferase (DAP-AT) (DAP-aminotransferase) (LL-DAP-aminotransferase) (EC 2.6.1.83) |
| 323 | 199 | 0.01 | Q39UA2 | *Geobacter metallireducens* | Adenylosuccinate lyase (ASL) (EC 4.3.2.2) (Adenylosuccinase) |
| 324 | 198 | 0.01 | A1AN79 | *Pelobacter propionicus* | Adenosylmethionine-8-amino-7-oxononanoate aminotransferase (EC 2.6.1.62) (7,8-diamino-pelargonic acid aminotransferase) (DAPA AT) (DAPA aminotransferase) (7,8-diaminononanoate synthase) (DANS) (Diaminopelargonic acid synthase) |
| 325 | 198 | 0.01 | A5GC25 | *Geobacter uraniireducens* | Propionyl-CoA carboxylase (EC 6.4.1.3) |
| 326 | 198 | 0.01 | A7I766 | *Methanoregula boonei* | Efflux transporter, putative, hydrophobe/amphiphile efflux-3 (HAE3) family |
| 327 | 198 | 0.01 | Q39Z79 | *Geobacter metallireducens* | Twitching motility pilus retraction ATPase |
| 328 | 198 | 0.01 | Q74EP9 | *Geobacter sulfurreducens* | DNA-binding ATPase Uup |
| 329 | 198 | 0.01 | | | |
| 330 | 196 | 0.01 | A0A0D5N2P4 | *Geobacter sulfurreducens* | Uncharacterized protein |
| 331 | 196 | 0.01 | A0A1T4LT91 | *Geobacter thiogenes* | Putative transposase |
| 332 | 196 | 0.01 | A5G8I3 | *Geobacter uraniireducens* | Chaperone protein ClpB |
| 333 | 196 | 0.01 | A7I5U5 | *Methanoregula boonei* | 4Fe-4S ferredoxin, iron-sulfur binding domain protein |
| 334 | 196 | 0.01 | Q3A5L6 | *Pelobacter carbinolicus* | Uncharacterized protein |
| 335 | 195 | 0.01 | A0A0B5B7G6 | *Geobacter pickeringii* | Cytochrome C |
| 336 | 195 | 0.01 | A0A1F8XUV7 | *Deltaproteobacteria bacterium* | Uncharacterized protein |

| | | | | | |
|---|---|---|---|---|---|
| 337 | 195 | 0.01 | Q39ZI6 | *Geobacter metallireducens* | DNA helicase (EC 3.6.4.12) |
| 338 | 194 | 0.01 | A0A0B5BFT0 | *Geobacter pickeringii* | Phosphoribosylformylglycinamidine synthase subunit PurL (FGAM synthase) (EC 6.3.5.3) (Formylglycinamide ribonucleotide amidotransferase subunit II) (FGAR amidotransferase II) (FGAR-AT II) (Glutamine amidotransferase PurL) (Phosphoribosylformylglycinamidine synthase subunit II) |
| 339 | 194 | 0.01 | A0A1T4RTR1 | *Geobacter thiogenes* | Iron complex outermembrane recepter protein |
| 340 | 194 | 0.01 | A5G3L3 | *Geobacter uraniireducens* | Isocitrate dehydrogenase, NADP-dependent (EC 1.1.1.42) |
| 341 | 194 | 0.01 | A5GE75 | *Geobacter uraniireducens* | Heavy metal translocating P-type ATPase |
| 342 | 194 | 0.01 | B5EHY9 | *Geobacter bemidjiensis* | Uncharacterized protein |
| 343 | 194 | 0.01 | Q39UB0 | *Geobacter metallireducens* | Trans-isoprenyl diphosphate synthase |
| 344 | 193 | 0.01 | A0A0A8WRD3 | *Geobacter sp.* | Putative ABC transporter ATP-binding protein YkpA |
| 345 | 193 | 0.01 | A0A0B5BB89 | *Geobacter pickeringii* | Chromosome partition protein Smc |
| 346 | 193 | 0.01 | A0A0B5BFH3 | *Geobacter pickeringii* | Polyribonucleotide nucleotidyltransferase (EC 2.7.7.8) (Polynucleotide phosphorylase) (PNPase) |
| 347 | 193 | 0.01 | Q39WR2 | *Geobacter metallireducens* | CRISPR-associated endonuclease Cas1 (EC 3.1.-.-) |
| 348 | 193 | 0.01 | Q74AR7 | *Geobacter sulfurreducens* | Membrane-associated metal-dependent phosphohydrolase, HDc domain-containing |
| 349 | 192 | 0.01 | D4H4P2 | *Denitrovibrio acetiphilus* | Transposase IS3/IS911 family protein |
| 350 | 192 | 0.01 | D8L7U2 | *Paramecium caudatum* | Ymf65 |
| 351 | 192 | 0.01 | F4BZD0 | *Methanosaeta concilii* | Transposase, putative |
| 352 | 192 | 0.01 | Q39VY8 | *Geobacter metallireducens* | Glutamine synthetase (EC 6.3.1.2) |
| 353 | 192 | 0.01 | Q74AF7 | *Geobacter sulfurreducens* | Methyl viologen-reducing hydrogenase, large subunit |
| 354 | 191 | 0.01 | A0A0D5N5Y8 | *Geobacter sulfurreducens* | Cell division protein FtsA |
| 355 | 191 | 0.01 | P61946 | *Geobacter sulfurreducens* | S-adenosylmethionine synthase (AdoMet synthase) (EC 2.5.1.6) (MAT) (Methionine adenosyltransferase) |
| 356 | 190 | 0.01 | A0A0B5B5Z5 | *Geobacter pickeringii* | DNA polymerase III subunit beta (EC 2.7.7.7) |
| 357 | 190 | 0.01 | A7I9M7 | *Methanoregula boonei* | PilT protein domain protein |
| 358 | 190 | 0.01 | Q39VY0 | *Geobacter metallireducens* | Malate dehydrogenase (EC 1.1.1.37) |
| 359 | 190 | 0.01 | Q748B3 | *Geobacter sulfurreducens* | UDP-N-acetylglucosamine 1-carboxyvinyltransferase (EC 2.5.1.7) (Enoylpyruvate transferase) (UDP-N-acetylglucosamine enolpyruvyl transferase) (EPT) |
| 360 | 190 | 0.01 | Q39QB3 | *Geobacter metallireducens* | NADH dehydrogenase I, G subunit |
| 361 | 189 | 0.01 | A0A0B5BIX5 | *Geobacter pickeringii* | DNA polymerase II |
| 362 | 189 | 0.01 | A7I4Y9 | *Methanoregula boonei* | DEAD/DEAH box helicase domain protein |
| 363 | 189 | 0.01 | Q39W73 | *Geobacter metallireducens* | 2-isopropylmalate synthase (EC 2.3.3.13) (Alpha-IPM synthase) (Alpha-isopropylmalate synthase) |
| 364 | 189 | 0.01 | Q39ZL2 | *Geobacter metallireducens* | TrkA domain protein |

| | | | | | |
|---|---|---|---|---|---|
| 365 | 189 | 0.01 | Q74D18 | *Geobacter sulfurreducens* | ABC transporter, ATP-binding protein |
| 366 | 189 | 0.01 | Q74E27 | *Geobacter sulfurreducens* | Ribonuclease Y (RNase Y) (EC 3.1.-.-) |
| 367 | 189 | 0.01 | Q74FM7 | *Geobacter sulfurreducens* | Glycine--tRNA ligase beta subunit (EC 6.1.1.14) (Glycyl-tRNA synthetase beta subunit) (GlyRS) |
| 368 | 189 | 0.01 | Q74H55 | *Geobacter sulfurreducens* | Serine--tRNA ligase (EC 6.1.1.11) (Seryl-tRNA synthetase) (SerRS) (Seryl-tRNA(Ser/Sec) synthetase) |
| 369 | 189 | 0.01 | | | |
| 370 | 188 | 0.01 | A0A0B5BHJ9 | *Geobacter pickeringii* | Chaperone protein HtpG (Heat shock protein HtpG) (High temperature protein G) |
| 371 | 188 | 0.01 | A0A0D5N949 | *Geobacter sulfurreducens* | Response regulator |
| 372 | 188 | 0.01 | A0A1G0LYB2 | *Geobacteraceae bacterium* | Uncharacterized protein |
| 373 | 188 | 0.01 | B3EAB4 | *Geobacter lovleyi* | Cytochrome c-552 (EC 1.7.2.2) |
| 374 | 188 | 0.01 | Q39VR4 | *Geobacter metallireducens* | Bifunctional uridylyltransferase/uridylyl-removing enzyme (UTase/UR) (Bifunctional [protein-PII] modification enzyme) (Bifunctional nitrogen sensor protein) [Includes: [Protein-PII]-UMP uridylyl-removing enzyme (UR) (EC 3.1.4.-); [Protein-PII] uridylyltransferase (PII uridylyltransferase) (UTase) (EC 2.7.7.59)] |
| 375 | 188 | 0.01 | Q747G4 | *Geobacter sulfurreducens* | (R)-methylmalonyl-CoA mutase, isobutyryl-CoA mutase-like catalytic subunit |
| 376 | 188 | 0.01 | Q74D04 | *Geobacter sulfurreducens* | Threonine--tRNA ligase (EC 6.1.1.3) (Threonyl-tRNA synthetase) (ThrRS) |
| 377 | 187 | 0.01 | A0A0B5BCP9 | *Geobacter pickeringii* | Uncharacterized protein |
| 378 | 187 | 0.01 | A0A0B5BE21 | *Geobacter pickeringii* | Histidine kinase |
| 379 | 187 | 0.01 | A0A0C1TQJ7 | *Geobacter soli* | Uncharacterized protein |
| 380 | 187 | 0.01 | D4H844 | *Denitrovibrio acetiphilus* | Carbamoyl-phosphate synthase large chain (EC 6.3.5.5) (Carbamoyl-phosphate synthetase ammonia chain) |
| 381 | 186 | 0.01 | A0A0A8WNV3 | *Geobacter sp.* | Uncharacterized protein |
| 382 | 186 | 0.01 | A0A0B5BBZ4 | *Geobacter pickeringii* | DNA polymerase I (EC 2.7.7.7) |
| 383 | 186 | 0.01 | A0A0B5BEL7 | *Geobacter pickeringii* | Cytochrome C |
| 384 | 186 | 0.01 | A0A1T4RHG8 | *Geobacter thiogenes* | Iron complex outermembrane recepter protein |
| 385 | 186 | 0.01 | A5GDX5 | *Geobacter uraniireducens* | DNA gyrase subunit A (EC 5.99.1.3) |
| 386 | 186 | 0.01 | | | |
| 387 | 185 | 0.01 | A0A0B5BAI5 | *Geobacter pickeringii* | Ketose-bisphosphate aldolase |
| 388 | 185 | 0.01 | A7I6L6 | *Methanoregula boonei* | DNA topoisomerase 1 (EC 5.99.1.2) (DNA topoisomerase I) |
| 389 | 185 | 0.01 | Q74FM8 | *Geobacter sulfurreducens* | Glycine--tRNA ligase alpha subunit (EC 6.1.1.14) (Glycyl-tRNA synthetase alpha subunit) (GlyRS) |
| 390 | 185 | 0.01 | | | |
| 391 | 184 | 0.01 | A0A0D5N3S3 | *Geobacter sulfurreducens* | Chemotaxis protein CheY |
| 392 | 184 | 0.01 | A0A0W8FRW6 | *hydrocarbon metagenome* | Uncharacterized protein |
| 393 | 184 | 0.01 | H8MU17 | *Corallococcus coralloides* | Long-chain-fatty-acid--CoA ligase |

| | | | | | |
|---|---|---|---|---|---|
| 394 | 183 | 0.01 | A0A0B5BH19 | *Geobacter pickeringii* | Uncharacterized protein |
| 395 | 182 | 0.01 | A0A0B5BB11 | *Geobacter pickeringii* | Pyruvate carboxylase (EC 6.4.1.1) |
| 396 | 182 | 0.01 | A0A1D3L5C3 | *Methanobacterium congolense* | Uncharacterized protein |
| 397 | 182 | 0.01 | A7IB61 | *Methanoregula boonei* | ATP-dependent DNA helicase Hel308 (EC 3.6.4.12) |
| 398 | 182 | 0.01 | Q74GF2 | *Geobacter sulfurreducens* | Metal-dependent phosphohydrolase, HDOD domain-containing |
| 399 | 181 | 0.01 | A0A0B5BIH3 | *Geobacter pickeringii* | Uncharacterized protein |
| 400 | 181 | 0.01 | A0A0D5NAV3 | *Geobacter sulfurreducens* | Peptidase S41 |
| 401 | 181 | 0.01 | A7I6S1 | *Methanoregula boonei* | CoA-binding domain protein |
| 402 | 181 | 0.01 | B9M149 | *Geobacter daltonii* | Translation-regulating membrane GTPase TypA |
| 403 | 181 | 0.01 | Q39UH4 | *Geobacter metallireducens* | Lon protease (EC 3.4.21.53) (ATP-dependent protease La) |
| 404 | 181 | 0.01 | A7I6H7 | *Methanoregula boonei* | Methionine--tRNA ligase (EC 6.1.1.10) (Methionyl-tRNA synthetase) (MetRS) |
| 405 | 180 | 0.01 | A0A0B5BD12 | *Geobacter pickeringii* | Inorganic pyrophosphatase |
| 406 | 180 | 0.01 | A0A0B5BDG1 | *Geobacter pickeringii* | tRNA uridine 5-carboxymethylaminomethyl modification enzyme MnmG (Glucose-inhibited division protein A) |
| 407 | 180 | 0.01 | A0A0B5BDP4 | *Geobacter pickeringii* | Polya polymerase |
| 408 | 180 | 0.01 | A0A0B5BGV8 | *Geobacter pickeringii* | Zinc protease |
| 409 | 180 | 0.01 | A0A1F9P4Q4 | *Desulfuromonadales bacterium* | Uncharacterized protein |
| 410 | 180 | 0.01 | A7I4K7 | *Methanoregula boonei* | Lysine decarboxylase (EC 4.1.1.18) |
| 411 | 180 | 0.01 | A7I656 | *Methanoregula boonei* | Elongation factor 1-alpha (EF-1-alpha) (Elongation factor Tu) (EF-Tu) |
| 412 | 180 | 0.01 | Q39VA8 | *Geobacter metallireducens* | Transcription termination/antitermination protein NusA |
| 413 | 180 | 0.01 | Q39XL2 | *Geobacter metallireducens* | Phosphoenolpyruvate synthase (PEP synthase) (EC 2.7.9.2) (Pyruvate, water dikinase) |
| 414 | 179 | 0.01 | A0A0A8WV23 | *Geobacter sp.* | Protein translocase subunit SecD |
| 415 | 179 | 0.01 | A0A0B5BGX4 | *Geobacter pickeringii* | Stage II sporulation protein E |
| 416 | 179 | 0.01 | A7I5I6 | *Methanoregula boonei* | Coenzyme F420 hydrogenase (EC 1.12.98.1) |
| 417 | 179 | 0.01 | A7IAF8 | *Methanoregula boonei* | ABC-1 domain protein |
| 418 | 179 | 0.01 | B9M380 | *Geobacter daltonii* | 4-hydroxy-tetrahydrodipicolinate synthase (HTPA synthase) (EC 4.3.3.7) |
| 419 | 179 | 0.01 | Q74H61 | *Geobacter sulfurreducens* | Heat-inducible transcription repressor HrcA |
| 420 | 179 | 0.01 | Q39Y18 | *Geobacter metallireducens* | Transcription termination/antitermination protein NusG |
| 421 | 178 | 0.01 | A0A090I3J2 | *Methanobacterium formicicum* | Uncharacterized protein |
| 422 | 178 | 0.01 | A0A0B5B8J3 | *Geobacter pickeringii* | Rod shape-determining protein RodA |
| 423 | 178 | 0.01 | A0A0B5BK21 | *Geobacter pickeringii* | Inner membrane cytochrome H |
| 424 | 178 | 0.01 | Q39QA3 | *Geobacter metallireducens* | ATP synthase subunit a (ATP synthase F0 sector subunit a) (F-ATPase subunit 6) |

| | | | | | |
|---|---|---|---|---|---|
| 425 | 178 | 0.01 | N8P3X5 | *Acinetobacter bohemicus* | Uncharacterized protein |
| 426 | 178 | 0.01 | Q748A7 | *Geobacter sulfurreducens* | Transcription termination factor Rho (EC 3.6.4.-) (ATP-dependent helicase Rho) |
| 427 | 177 | 0.01 | A0A090JUS9 | *Methanobacterium formicicum* | Putative secreted protein |
| 428 | 177 | 0.01 | | | |
| 429 | 176 | 0.01 | A0A0A8WQH6 | *Geobacter sp.* | Uncharacterized protein |
| 430 | 176 | 0.01 | A0A0B5B6N9 | *Geobacter pickeringii* | Alanine--tRNA ligase (EC 6.1.1.7) (Alanyl-tRNA synthetase) (AlaRS) |
| 431 | 176 | 0.01 | A0A0D5N435 | *Geobacter sulfurreducens* | Multidrug transporter |
| 432 | 176 | 0.01 | A0A1G0MUH0 | *Geobacteraceae bacterium* | Integrase |
| 433 | 176 | 0.01 | Q39W76 | *Geobacter metallireducens* | Ketol-acid reductoisomerase (NADP(+)) (KARI) (EC 1.1.1.86) (Acetohydroxy-acid isomeroreductase) (AHIR) (Alpha-keto-beta-hydroxylacyl reductoisomerase) (Ketol-acid reductoisomerase type 1) (Ketol-acid reductoisomerase type I) |
| 434 | 176 | 0.01 | Q39TA6 | *Geobacter metallireducens* | Inosine-5'-monophosphate dehydrogenase (IMP dehydrogenase) (IMPD) (IMPDH) (EC 1.1.1.205) |
| 435 | 175 | 0.01 | A0A0A8WLB0 | *Geobacter sp.* | Protein TraH |
| 436 | 175 | 0.01 | A0A0B5BIW8 | *Geobacter pickeringii* | Aminotransferase |
| 437 | 175 | 0.01 | A7IB30 | *Methanoregula boonei* | AAA family ATPase, CDC48 subfamily (EC 3.6.4.6) |
| 438 | 175 | 0.01 | D4H7W9 | *Denitrovibrio acetiphilus* | Translation initiation factor IF-2 |
| 439 | 175 | 0.01 | Q39U98 | *Geobacter metallireducens* | Phosphoglycerate kinase (EC 2.7.2.3) |
| 440 | 175 | 0.01 | Q39PY9 | *Geobacter metallireducens* | Histidine kinase (EC 2.7.13.3) |
| 441 | 175 | 0.01 | Q39X31 | *Geobacter metallireducens* | Protein translocase subunit SecA |
| 442 | 175 | 0.01 | A7I7S4 | *Methanoregula boonei* | Alanine--tRNA ligase (EC 6.1.1.7) (Alanyl-tRNA synthetase) (AlaRS) |
| 443 | 175 | 0.01 | | | |
| 444 | 174 | 0.01 | A0A0B5BJ17 | *Geobacter pickeringii* | UvrABC system protein C (Protein UvrC) (Excinuclease ABC subunit C) |
| 445 | 174 | 0.01 | A0A0B5BK07 | *Geobacter pickeringii* | Histidine kinase |
| 446 | 174 | 0.01 | A0A1D3L0E4 | *Methanobacterium congolense* | WD40 domain-containing protein |
| 447 | 174 | 0.01 | D0SHF8 | *Acinetobacter johnsonii* | Uncharacterized protein |
| 448 | 174 | 0.01 | Q74D63 | *Geobacter sulfurreducens* | Uncharacterized protein |
| 449 | 174 | 0.01 | Q747K3 | *Geobacter sulfurreducens* | UvrABC system protein B (Protein UvrB) (Excinuclease ABC subunit B) |
| 450 | 173 | 0.01 | A0A0B5BDG8 | *Geobacter pickeringii* | ABC transporter ATP-binding protein |
| 451 | 173 | 0.01 | A0A0B5BJV3 | *Geobacter pickeringii* | ABC transporter substrate-binding protein |
| 452 | 173 | 0.01 | A0A0D5N6Q2 | *Geobacter sulfurreducens* | Lactamase |
| 453 | 173 | 0.01 | P05489 | *Paramecium tetraurelia* | Cytochrome c oxidase subunit 1 (EC 1.9.3.1) (Cytochrome c oxidase polypeptide I) |
| 454 | 173 | 0.01 | L0HAI4 | *Methanoregula formicica* | Formate dehydrogenase, alpha subunit, archaeal-type |
| 455 | 173 | 0.01 | Q74B66 | *Geobacter sulfurreducens* | TrfA family protein |

| | | | | | |
|---|---|---|---|---|---|
| 456 | 173 | 0.01 | | | |
| 457 | 172 | 0.01 | A0A0B5BGU8 | *Geobacter pickeringii* | Chemotaxis protein |
| 458 | 172 | 0.01 | A0A0B5BGY7 | *Geobacter pickeringii* | Phosphoglucomutase |
| 459 | 172 | 0.01 | A0A0D5N9G3 | *Geobacter sulfurreducens* | Multidrug ABC transporter substrate-binding protein |
| 460 | 172 | 0.01 | A0A0Q7S8L4 | *Pelomonas sp.* | Transposase |
| 461 | 172 | 0.01 | A7I4R1 | *Methanoregula boonei* | Thermosome |
| 462 | 172 | 0.01 | A5GAC6 | *Geobacter uraniireducens* | FMN-dependent NADH-azoreductase (EC 1.7.-.-) (Azo-dye reductase) (FMN-dependent NADH-azo compound oxidoreductase) |
| 463 | 172 | 0.01 | F4BY05 | *Methanosaeta concilii* | Transposase, IS116/IS110/IS902 family |
| 464 | 172 | 0.01 | Q748T0 | *Geobacter sulfurreducens* | 5-methyltetrahydrofolate--homocysteine S-methyltransferase, cobalamin-dependent |
| 465 | 172 | 0.01 | Q74D56 | *Geobacter sulfurreducens* | Aspartate--tRNA(Asp/Asn) ligase (EC 6.1.1.23) (Aspartyl-tRNA synthetase) (AspRS) (Non-discriminating aspartyl-tRNA synthetase) (ND-AspRS) |
| 466 | 172 | 0.01 | | | |
| 467 | 172 | 0.01 | | | |
| 468 | 171 | 0.01 | A0A0B5BHF0 | *Geobacter pickeringii* | 2-oxoglutarate dehydrogenase (EC 1.2.4.2) |
| 469 | 171 | 0.01 | B3EAZ5 | *Geobacter lovleyi* | TraG family protein |
| 470 | 171 | 0.01 | F4GG80 | *Alicycliphilus denitrificans* | Transposase IS4 family protein |
| 471 | 171 | 0.01 | Q74A32 | *Geobacter sulfurreducens* | PppGpp 5'-phosphohydrolase and exopolyphosphatase, HD domain-containing |
| 472 | 170 | 0.01 | A0A0A8WQR9 | *Geobacter sp.* | Helix-turn-helix domain protein |
| 473 | 170 | 0.01 | A0A0D5N645 | *Geobacter sulfurreducens* | Penicillin-binding protein |
| 474 | 170 | 0.01 | A0A1G1FB69 | *Nitrospirae bacterium* | Uncharacterized protein |
| 475 | 170 | 0.01 | B3EA94 | *Geobacter lovleyi* | Uncharacterized protein |
| 476 | 170 | 0.01 | Q746T2 | *Geobacter sulfurreducens* | NADH-quinone oxidoreductase subunit H 2 (EC 1.6.5.11) (NADH dehydrogenase I subunit H 2) (NDH-1 subunit H 2) |
| 477 | 170 | 0.01 | Q39VU9 | *Geobacter metallireducens* | Polyprenyl-phospho-glycoside--protein O-glycosyltransferase DUF2723 membrane protein, putative |
| 478 | 170 | 0.01 | Q74BT7 | *Geobacter sulfurreducens* | Sensor histidine kinase cyclic nucleotide phosphodiesterase, GAF, GAF and HD-GYP-related domain-containing |
| 479 | 170 | 0.01 | Q748Y3 | *Geobacter sulfurreducens* | 50S ribosomal protein L1 |
| 480 | 169 | 0.01 | A0A1B1ICB8 | *Prevotella scopos* | Uncharacterized protein |
| 481 | 169 | 0.01 | B9M5J8 | *Geobacter daltonii* | CRISPR-associated protein Csd1 |
| 482 | 169 | 0.01 | Q39QK5 | *Geobacter metallireducens* | Uncharacterized protein |
| 483 | 169 | 0.01 | Q39RM2 | *Geobacter metallireducens* | Glycosyltransferase |
| 484 | 169 | 0.01 | Q749D7 | *Geobacter sulfurreducens* | Nitrogenase molybdenum-iron cofactor biosynthesis protein NifEN |
| 485 | 169 | 0.01 | A7I6P0 | *Methanoregula boonei* | Threonine--tRNA ligase (EC 6.1.1.3) (Threonyl-tRNA synthetase) (ThrRS) |

| | | | | | |
|---|---|---|---|---|---|
| 486 | 168 | 0.01 | A0A0B5B792 | *Geobacter pickeringii* | Flagellar biosynthesis protein FlhF |
| 487 | 168 | 0.01 | A0A0B5BBX6 | *Geobacter pickeringii* | Ribonuclease E (RNase E) (EC 3.1.26.12) |
| 488 | 168 | 0.01 | A0A0B5BHF1 | *Geobacter pickeringii* | Antibiotic ABC transporter ATP-binding protein |
| 489 | 168 | 0.01 | A0A0D5N6L1 | *Geobacter sulfurreducens* | Histidine kinase |
| 490 | 168 | 0.01 | Q74D76 | *Geobacter sulfurreducens* | Histidine kinase (EC 2.7.13.3) |
| 491 | 167 | 0.01 | A0A0D5N5H4 | *Geobacter sulfurreducens* | Cell division protein FtsK |
| 492 | 167 | 0.01 | A0A1F9PKR5 | *Desulfuromonadales bacterium* | Uncharacterized protein |
| 493 | 167 | 0.01 | B9M7P3 | *Geobacter daltonii* | UvrABC system protein A (UvrA protein) (Excinuclease ABC subunit A) |
| 494 | 167 | 0.01 | Q39ZF6 | *Geobacter metallireducens* | Carbamoyltransferase HypF (EC 2.1.3.-) |
| 495 | 167 | 0.01 | Q746S9 | *Geobacter sulfurreducens* | NADH-quinone oxidoreductase (EC 1.6.5.11) |
| 496 | 167 | 0.01 | U6EA67 | *Methanobacterium sp.* | Uncharacterized protein |
| 497 | 166 | 0.01 | A0A0B5B7W4 | *Geobacter pickeringii* | Transporter |
| 498 | 166 | 0.01 | A0A0B5BBV1 | *Geobacter pickeringii* | Cytochrome C |
| 499 | 166 | 0.01 | A0A0D5NDV9 | *Geobacter sulfurreducens* | HD family phosphohydrolase |
| 500 | 166 | 0.01 | C9MTU8 | *Prevotella veroralis* | Initiator RepB protein |

**Table S6c.** Functional overview of M4P community (top 500 rank only).

| Rank | Count | Abundance | ID | Organism | Protein Names |
|---|---|---|---|---|---|
| 1 | 4631 | 0.17 | Q8VQM8 | *Prevotella intermedia* | Mobilization protein |
| 2 | 3649 | 0.14 | N8ZVC9 | *Acinetobacter gerneri* | Uncharacterized protein |
| 3 | 2964 | 0.11 | F0FB90 | *Prevotella multiformis* | Initiator RepB protein (Fragment) |
| 4 | 2821 | 0.11 | A0A0A2LWZ9 | *Flavobacterium suncheonense* | Transposase |
| 5 | 2600 | 0.10 | N8ZNX3 | *Acinetobacter gerneri* | Uncharacterized protein |
| 6 | 2509 | 0.09 | A0A0U1CQ28 | *Chlamydia trachomatis* | Replication protein |
| 7 | 2497 | 0.09 | F0FB88 | *Prevotella multiformis* | Putative plasmid recombination enzyme |
| 8 | 2288 | 0.09 | A0A0R3QEE4 | *Brugia timori* | Uncharacterized protein |
| 9 | 2060 | 0.08 | A0A0U1CQ01 | *Chlamydia trachomatis* | Uncharacterised protein |
| 10 | 1878 | 0.07 | | | |
| 11 | 1862 | 0.07 | Q8VQM9 | *Prevotella intermedia* | Replication protein |
| 12 | 1646 | 0.06 | A0A077XLW5 | *Sphinx1.76-related DNA* | Replication protein |
| 13 | 1478 | 0.06 | A0A0E9G9S8 | *Chlamydia trachomatis* | Uncharacterised protein |
| 14 | 1422 | 0.05 | A0A077XNB6 | *Sphinx1.76-related DNA* | Replication protein |
| 15 | 1400 | 0.05 | A0A0Q7S8L4 | *Pelomonas sp.* | Transposase |
| 16 | 1364 | 0.05 | | | |
| 17 | 1359 | 0.05 | A0A1B1ICB8 | *Prevotella scopos* | Uncharacterized protein |
| 18 | 1322 | 0.05 | C9MTU8 | *Prevotella veroralis* | Initiator RepB protein |
| 19 | 1322 | 0.05 | F4GG80 | *Alicycliphilus denitrificans* | Transposase IS4 family protein |
| 20 | 1154 | 0.04 | A0A1Q3RZJ5 | *Flavobacterium sp.* | Uncharacterized protein |
| 21 | 1123 | 0.04 | A0A023UP33 | *Magnusiomyces ingens* | Uncharacterized protein (Fragment) |
| 22 | 1109 | 0.04 | A0A1Q3SCF5 | *Flavobacterium sp.* | Uncharacterized protein |
| 23 | 1068 | 0.04 | N8YFI4 | *Acinetobacter gerneri* | Uncharacterized protein |
| 24 | 1004 | 0.04 | A0A1Q3S858 | *Flavobacterium sp.* | Uncharacterized protein |
| 25 | 917 | 0.03 | A0A1E4A455 | *Chryseobacterium sp.* | Cell surface protein SprA |
| 26 | 903 | 0.03 | A0A1Q3RWC6 | *Flavobacterium sp.* | Uncharacterized protein |
| 27 | 864 | 0.03 | A0A1Q3M6H6 | *Bacteroidales bacterium* | CRISPR-associated endonuclease Cas9 (EC 3.1.-.-) |
| 28 | 847 | 0.03 | A0A023UP27 | *Magnusiomyces ingens* | Uncharacterized protein |
| 29 | 847 | 0.03 | A0A0Q7S7D9 | *Pelomonas sp.* | Uncharacterized protein |
| 30 | 847 | 0.03 | A0A1L9GVN0 | *Cand. Micrarchaeum acidiphilum* | Uncharacterized protein |
| 31 | 809 | 0.03 | G9FB24 | *uncultured bacterium* | IS21 family transposition protein |
| 32 | 792 | 0.03 | A0A1E3ZQ34 | *Chryseobacterium sp.* | Cell envelope biogenesis protein OmpA |
| 33 | 764 | 0.03 | A0A0D2WNJ4 | *Capsaspora owczarzaki* | Paramyosin |

| | | | | | |
|---|---|---|---|---|---|
| 34 | 760 | 0.03 | A0A0T5YH86 | *Acinetobacter baumannii* | Mobilization protein A |
| 35 | 748 | 0.03 | E4T3F9 | *Paludibacter propionicigenes* | Glutamate synthase (NADH) large subunit (EC 1.4.1.14) (EC 1.4.7.1) |
| 36 | 737 | 0.03 | A0A0D8Y6E2 | *Dictyocaulus viviparus* | Uncharacterized protein |
| 37 | 722 | 0.03 | A0A1Q3S9F2 | *Flavobacterium sp.* | Uncharacterized protein |
| 38 | 698 | 0.03 | A0A1Q3S0Y5 | *Flavobacterium sp.* | Uncharacterized protein |
| 39 | 695 | 0.03 | A0A0Q7RZB1 | *Pelomonas sp. Root1444* | Uncharacterized protein |
| 40 | 637 | 0.02 | A0A0M8MCD5 | *Flavobacterium akiainvivens* | Integrase |
| 41 | 622 | 0.02 | A0A1J5AN19 | *Bacteroidetes bacterium* | IS110 family transposase |
| 42 | 615 | 0.02 | A0A023UMS7 | *Magnusiomyces ingens* | NADH-ubiquinone oxidoreductase chain 5 (EC 1.6.5.3) |
| 43 | 599 | 0.02 | A0A016UCK1 | *Ancylostoma ceylanicum* | Uncharacterized protein |
| 44 | 582 | 0.02 | A0A1E4A9F0 | *Chryseobacterium sp.* | Uncharacterized protein |
| 45 | 572 | 0.02 | A0A023UMT3 | *Magnusiomyces ingens* | Uncharacterized protein (Fragment) |
| 46 | 569 | 0.02 | A0A0R3R918 | *Brugia timori* | Uncharacterized protein |
| 47 | 568 | 0.02 | A0A1K2IH81 | *Chryseobacterium limigenitum* | Uncharacterized protein |
| 48 | 530 | 0.02 | A0A1E4ARQ1 | *Chryseobacterium sp.* | Uncharacterized protein |
| 49 | 523 | 0.02 | A0A1T5HUH3 | *Alkalitalea saponilacus* | Uncharacterized protein |
| 50 | 519 | 0.02 | G0IXG6 | *Cyclobacterium marinum* | Transposase IS116/IS110/IS902 family protein |
| 51 | 515 | 0.02 | A0A0Q6WNI0 | *Pelomonas sp.* | DNA-directed RNA polymerase subunit beta' (RNAP subunit beta') (EC 2.7.7.6) (RNA polymerase subunit beta') (Transcriptase subunit beta') |
| 52 | 511 | 0.02 | A0A1E4AD77 | *Chryseobacterium sp.* | Uncharacterized protein |
| 53 | 511 | 0.02 | A0A1Q3S812 | *Flavobacterium sp.* | Uncharacterized protein |
| 54 | 503 | 0.02 | E4T8J6 | *Paludibacter propionicigenes* | Cell surface receptor IPT/TIG domain protein |
| 55 | 501 | 0.02 | A0A077XLZ7 | *Sphinx1.76-related DNA* | Replication protein |
| 56 | 494 | 0.02 | A0A127MR59 | *Pseudomonas citronellolis* | Uncharacterized protein |
| 57 | 494 | 0.02 | V6RWN6 | *Flavobacterium cauense* | Uncharacterized protein |
| 58 | 491 | 0.02 | A0A1M3DNL6 | *Bacteroidales bacterium* | IS4 family transposase |
| 59 | 481 | 0.02 | A0A1Q3RUL1 | *Flavobacterium sp.* | Uncharacterized protein |
| 60 | 481 | 0.02 | F3ZQL8 | *Bacteroides coprosuis* | Transposase IS4 family protein |
| 61 | 472 | 0.02 | A0A1Q3S0Z6 | *Flavobacterium sp.* | Uncharacterized protein |
| 62 | 472 | 0.02 | E4T5D1 | *Paludibacter propionicigenes* | Methionine--tRNA ligase (EC 6.1.1.10) (Methionyl-tRNA synthetase) (MetRS) |
| 63 | 466 | 0.02 | | | |
| 64 | 459 | 0.02 | A0A174XH04 | *Bacteroides caccae* | Addiction module toxin, Txe/YoeB family |
| 65 | 451 | 0.02 | | | |
| 66 | 450 | 0.02 | A0A180EWA0 | *Bacteroidales bacterium* | Replication protein |
| 67 | 445 | 0.02 | A0A167ZLW1 | *Flavobacterium fryxellicola* | Uncharacterized protein |
| 68 | 442 | 0.02 | A0A1Q3S042 | *Flavobacterium sp.* | Peptidase C25 |

| | | | | | |
|---|---|---|---|---|---|
| 69 | 439 | 0.02 | A0A0K3A869 | *Xanthomonas translucens pv. poae* | Riboflavin biosynthesis protein (EC 2.7.7.2) |
| 70 | 438 | 0.02 | A0A1Q3S6M2 | *Flavobacterium sp.* | Uncharacterized protein |
| 71 | 436 | 0.02 | G9FB23 | *uncultured bacterium* | IS21 family transposition protein |
| 72 | 434 | 0.02 | | | |
| 73 | 432 | 0.02 | A0A1E4A330 | *Chryseobacterium sp.* | Acriflavin resistance protein |
| 74 | 431 | 0.02 | A0A1Q3RUG1 | *Flavobacterium sp.* | Uncharacterized protein |
| 75 | 430 | 0.02 | A0A1E3ZXL4 | *Chryseobacterium sp.* | Uncharacterized protein |
| 76 | 430 | 0.02 | A0A1Q3RVF6 | *Flavobacterium sp.* | Multidrug transporter AcrB |
| 77 | 429 | 0.02 | F0FB86 | *Prevotella multiformis DSM 16608* | Uncharacterized protein |
| 78 | 428 | 0.02 | A0A060RAF8 | *Mucinivorans hirudinis* | Mobile element protein |
| 79 | 425 | 0.02 | A0A1E4ADX1 | *Chryseobacterium sp.* | Uncharacterized protein |
| 80 | 424 | 0.02 | A0A177LQD3 | *Chryseobacterium sp. J200* | Uncharacterized protein |
| 81 | 423 | 0.02 | A0A1E4AFR9 | *Chryseobacterium sp.* | Uncharacterized protein |
| 82 | 420 | 0.02 | A0A1Q3RUG7 | *Flavobacterium sp.* | Uncharacterized protein |
| 83 | 416 | 0.02 | A0A1Q3RVI5 | *Flavobacterium sp.* | Uncharacterized protein |
| 84 | 415 | 0.02 | A0A1E4ABZ6 | *Chryseobacterium sp.* | Multifunctional fusion protein [Includes: Protein translocase subunit SecD; Protein-export membrane protein SecF] |
| 85 | 414 | 0.02 | A0A1E4AA02 | *Chryseobacterium sp.* | TonB-dependent receptor |
| 86 | 413 | 0.02 | A0A0K0WVI4 | *Litoditis aff. marina PmIII* | NADH dehydrogenase subunit 5 |
| 87 | 412 | 0.02 | | | |
| 88 | 411 | 0.02 | A0A1Q3S3Z3 | *Flavobacterium sp.* | DNA methylase |
| 89 | 407 | 0.02 | A0A1Q3S3S6 | *Flavobacterium sp.* | Uncharacterized protein |
| 90 | 406 | 0.02 | A0A0Q7SV37 | *Pelomonas sp. Root1444* | Glutamate synthase |
| 91 | 402 | 0.02 | A0A0Q8GZB4 | *Pelomonas sp. Root662* | UvrABC system protein A |
| 92 | 402 | 0.02 | A0A1Q3S6H8 | *Flavobacterium sp.* | Uncharacterized protein |
| 93 | 399 | 0.01 | | | |
| 94 | 397 | 0.01 | | | |
| 95 | 394 | 0.01 | A0A1Q3S3X4 | *Flavobacterium sp.* | Uncharacterized protein |
| 96 | 394 | 0.01 | N8YF58 | *Acinetobacter gerneri* | Uncharacterized protein |
| 97 | 393 | 0.01 | A0A1Q3M3N2 | *Bacteroidales bacterium* | Uncharacterized protein (Fragment) |
| 98 | 392 | 0.01 | A0A1E4AD73 | *Chryseobacterium sp.* | Outer membrane protein assembly factor BamA |
| 99 | 390 | 0.01 | A0A023UP41 | *Magnusiomyces ingens* | Cytochrome c oxidase subunit 1 (EC 1.9.3.1) |
| 100 | 387 | 0.01 | A0A1Q3RUH2 | *Flavobacterium sp.* | Uncharacterized protein |
| 101 | 385 | 0.01 | A0A127MVT9 | *Pseudomonas citronellolis* | Uncharacterized protein |

| | | | | | |
|---|---|---|---|---|---|
| 102 | 385 | 0.01 | A0A177LRY5 | *Chryseobacterium sp.* | Uncharacterized protein |
| 103 | 379 | 0.01 | K9L3G6 | *Magnusiomyces magnusii* | Cytochrome b |
| 104 | 377 | 0.01 | I9R3U4 | *Bacteroides cellulosilyticus* | Uncharacterized protein |
| 105 | 375 | 0.01 | A0A1B2YV26 | *uncultured bacterium* | Conserved repeat domain-containing protein |
| 106 | 373 | 0.01 | A0A023UPJ4 | *Magnusiomyces ingens* | Uncharacterized protein (Fragment) |
| 107 | 373 | 0.01 | A0A1Q3S919 | *Flavobacterium sp.* | SusC/RagA family protein |
| 108 | 373 | 0.01 | | | |
| 109 | 372 | 0.01 | A0A023UM92 | *Magnusiomyces ingens* | Uncharacterized protein |
| 110 | 370 | 0.01 | A0A140JXE2 | *Escherichia coli* | Uncharacterized protein |
| 111 | 370 | 0.01 | A0A1J5HM76 | *Porphyromonadaceae bacterium* | Multidrug transporter AcrB |
| 112 | 369 | 0.01 | A0A023UM78 | *Magnusiomyces ingens* | NADH dehydrogenase subunit 2 (EC 1.6.5.3) |
| 113 | 369 | 0.01 | A0A1Q3RYV8 | *Flavobacterium sp.* | 2-oxoglutarate dehydrogenase E1 component |
| 114 | 368 | 0.01 | A0A1E3ZQQ1 | *Chryseobacterium sp.* | Uncharacterized protein |
| 115 | 366 | 0.01 | Q45FH5 | *Adineta vaga* | Transposase |
| 116 | 366 | 0.01 | W2UCH0 | *Gammaproteobacteria bacterium* | Uncharacterized protein |
| 117 | 362 | 0.01 | A0A1Q3S7F1 | *Flavobacterium sp.* | Beta-N-acetylglucosaminidase |
| 118 | 362 | 0.01 | A0A1Q3S9B3 | *Flavobacterium sp.* | Uncharacterized protein |
| 119 | 360 | 0.01 | A0A154DNE7 | *Acinetobacter baumannii* | mRNA endoribonuclease LS (EC 3.1.-.-) |
| 120 | 360 | 0.01 | A0A1E4A6N2 | *Chryseobacterium sp.* | Alkaline phosphatase |
| 121 | 359 | 0.01 | A0A0A2M931 | *Flavobacterium suncheonsens* | Uncharacterized protein (Fragment) |
| 122 | 358 | 0.01 | A0A1Q3S993 | *Flavobacterium sp.* | Uncharacterized protein |
| 123 | 356 | 0.01 | A0A1E4AC15 | *Chryseobacterium sp.* | Uncharacterized protein |
| 124 | 353 | 0.01 | A0A1E4AHR4 | *Chryseobacterium sp.* | Uncharacterized protein |
| 125 | 352 | 0.01 | A0A1E4A2B9 | *Chryseobacterium sp.* | Quinol:cytochrome C oxidoreductase |
| 126 | 351 | 0.01 | A1W6Q0 | *Acidovorax sp.* | Integrase, catalytic region |
| 127 | 350 | 0.01 | A0A1E3ZYA8 | *Chryseobacterium sp.* | Uncharacterized protein |
| 128 | 348 | 0.01 | A0A1E4A8D6 | *Chryseobacterium sp.* | Metallophosphoesterase |
| 129 | 346 | 0.01 | A0A1E4A2A9 | *Chryseobacterium sp.* | Organic solvent tolerance protein OstA |
| 130 | 343 | 0.01 | A0A1Q3S1D7 | *Flavobacterium sp.* | TonB-dependent receptor |
| 131 | 343 | 0.01 | R7Y1V2 | *Nocardioides sp. CF8* | Uncharacterized protein |
| 132 | 342 | 0.01 | A0A1E4AC98 | *Chryseobacterium sp.* | Probable potassium transport system protein kup |
| 133 | 341 | 0.01 | A0A1Q3S8X4 | *Flavobacterium sp.* | Cation transporter |
| 134 | 340 | 0.01 | A0A0Q6WK82 | *Pelomonas sp.* | RND transporter |
| 135 | 339 | 0.01 | A0A1E4AAS0 | *Chryseobacterium sp.* | Uncharacterized protein |

| | | | | | |
|---|---|---|---|---|---|
| 136 | 338 | 0.01 | A0A1Q3RWE1 | *Flavobacterium sp.* | Penicillin-binding protein |
| 137 | 337 | 0.01 | A0A0Q7SRI3 | *Pelomonas sp.* | Efflux pump membrane transporter |
| 138 | 337 | 0.01 | A0A1Q3S143 | *Flavobacterium sp.* | Glycine dehydrogenase (decarboxylating) (EC 1.4.4.2) (Glycine cleavage system P-protein) (Glycine decarboxylase) (Glycine dehydrogenase (aminomethyl-transferring)) |
| 139 | 336 | 0.01 | A0A1Q3RVH4 | *Flavobacterium sp.* | Metallophosphoesterase |
| 140 | 335 | 0.01 | A0A015R892 | *Bacteroides fragilis* | Plasmid encoded toxin Txe family protein |
| 141 | 335 | 0.01 | A0A0Q7SJ43 | *Pelomonas sp.* | Transposase |
| 142 | 333 | 0.01 | A0A1Q3S2G4 | *Flavobacterium sp.* | Protein translocase subunit SecA |
| 143 | 333 | 0.01 | D0SHF8 | *Acinetobacter johnsonii* | Uncharacterized protein |
| 144 | 329 | 0.01 | A0A0Q7SS31 | *Pelomonas sp.* | Transposase |
| 145 | 328 | 0.01 | A0A1Q3RXG1 | *Flavobacterium sp.* | Uncharacterized protein |
| 146 | 327 | 0.01 | A0A1I1TKT3 | *Pseudomonas citronellolis* | Uncharacterized protein |
| 147 | 327 | 0.01 | E9I741 | *Daphnia pulex* | Putative uncharacterized protein (Fragment) |
| 148 | 325 | 0.01 | A0A077XLZ0 | *Sphinx1.76-related DNA* | Replication protein |
| 149 | 325 | 0.01 | A0A0R0LGW3 | *Ferrovum sp.* | ATP-dependent RNA helicase RhlE (EC 3.6.4.13) |
| 150 | 325 | 0.01 | A0A1E4ACQ6 | *Chryseobacterium sp.* | LmbE family protein |
| 151 | 324 | 0.01 | A0A1M3DC14 | *Bacteroidales bacterium* | Formate C-acetyltransferase |
| 152 | 324 | 0.01 | A0A1Q3S2W6 | *Flavobacterium sp.* | Cytochrome C biogenesis protein |
| 153 | 320 | 0.01 | E4T072 | *Paludibacter propionicigenes* | Pyruvate, phosphate dikinase (EC 2.7.9.1) |
| 154 | 319 | 0.01 | A0A1E4A9M1 | *Chryseobacterium sp.* | Cation transporter |
| 155 | 319 | 0.01 | A0A1Q3RYP3 | *Flavobacterium sp.* | DNA-directed DNA polymerase (EC 2.7.7.7) |
| 156 | 318 | 0.01 | A0A1Q3S3T4 | *Flavobacterium sp.* | TonB-dependent receptor |
| 157 | 317 | 0.01 | A0A177LPI8 | *Chryseobacterium sp.* | Uncharacterized protein |
| 158 | 315 | 0.01 | A0A1Q3SAL9 | *Flavobacterium sp.* | Peptidase M3 |
| 159 | 314 | 0.01 | A0A1E4AHQ8 | *Chryseobacterium sp.* | TonB-dependent receptor |
| 160 | 314 | 0.01 | A0A1Q3SAB4 | *Flavobacterium sp.* | TonB-dependent receptor |
| 161 | 314 | 0.01 | | | |
| 162 | 313 | 0.01 | A0A0Q7SDJ3 | *Pelomonas sp.* | Iron permease |
| 163 | 313 | 0.01 | A0A1E4AFY9 | *Chryseobacterium sp.* | Uncharacterized protein |
| 164 | 312 | 0.01 | A0A1E4A7M8 | *Chryseobacterium sp.* | Uncharacterized protein |
| 165 | 312 | 0.01 | A0A1Q3SAM2 | *Flavobacterium sp.* | Uncharacterized protein |
| 166 | 311 | 0.01 | F5UKT4 | *Microcoleus vaginatus* | Polymorphic outer membrane protein |
| 167 | 310 | 0.01 | E4T5D5 | *Paludibacter propionicigenes* | DNA-directed DNA polymerase (EC 2.7.7.7) |
| 168 | 309 | 0.01 | A0A0E9GA77 | *Chlamydia trachomatis* | Toxin RelK (EC 3.1.-.-) |
| 169 | 309 | 0.01 | A0A1Q3RXA0 | *Flavobacterium sp.* | Uncharacterized protein |

| | | | | | |
|---|---|---|---|---|---|
| 170 | 309 | 0.01 | A0A1Q3S7G4 | *Flavobacterium sp.* | Acriflavin resistance protein |
| 171 | 308 | 0.01 | A0A0M9VJW9 | *Flavobacterium akiainvivens* | Transposase |
| 172 | 308 | 0.01 | A0A0Q7SMD8 | *Pelomonas sp.* | Acriflavine resistance protein B |
| 173 | 308 | 0.01 | A0A1Q3S2Z4 | *Flavobacterium sp.* | Aminopeptidase |
| 174 | 307 | 0.01 | A0A1E3ZPZ7 | *Chryseobacterium sp.* | Peptidase S46 |
| 175 | 307 | 0.01 | A0A1Q3S5R1 | *Flavobacterium sp.* | Uncharacterized protein |
| 176 | 307 | 0.01 | A0A1Q3S7K6 | *Flavobacterium sp.* | Uncharacterized protein |
| 177 | 306 | 0.01 | A0A1Q3RXE2 | *Flavobacterium sp.* | Uncharacterized protein |
| 178 | 305 | 0.01 | A0A1E4ADZ5 | *Chryseobacterium sp.* | ABC transporter substrate-binding protein |
| 179 | 305 | 0.01 | A0A1J5HWN1 | *Porphyromonadaceae bacterium* | Uncharacterized protein |
| 180 | 305 | 0.01 | A0A1Q3LCT2 | *Bacteroidales bacterium* | Cell filamentation protein Fic |
| 181 | 305 | 0.01 | A0A1Q3RT16 | *Flavobacterium sp.* | S9 family peptidase |
| 182 | 304 | 0.01 | A0A0Q7S7G4 | *Pelomonas sp.* | Efflux pump membrane transporter |
| 183 | 304 | 0.01 | A0A1Q3S7E9 | *Flavobacterium sp.* | Formate acetyltransferase |
| 184 | 304 | 0.01 | A6GYU0 | *Flavobacterium psychrophilum* | DNA-directed RNA polymerase subunit beta (RNAP subunit beta) (EC 2.7.7.6) (RNA polymerase subunit beta) (Transcriptase subunit beta) |
| 185 | 302 | 0.01 | A0A0Q7RYH2 | *Pelomonas sp.* | 4-deoxy-4-formamido-L-arabinose-phospho-UDP deformylase |
| 186 | 302 | 0.01 | A0A1Q3M633 | *Bacteroidales bacterium* | ATP-dependent helicase |
| 187 | 301 | 0.01 | A0A0Q7SBL7 | *Pelomonas sp.* | Efflux pump membrane transporter |
| 188 | 299 | 0.01 | A0A1Q3S8S4 | *Flavobacterium sp.* | Sodium:proline symporter |
| 189 | 296 | 0.01 | A0A1E4AD61 | *Chryseobacterium sp.* | Uncharacterized protein |
| 190 | 296 | 0.01 | E4T5Y5 | *Paludibacter propionicigenes* | Multifunctional fusion protein [Includes: Protein translocase subunit SecD; Protein-export membrane protein SecF] |
| 191 | 293 | 0.01 | A0A1Q3S9A2 | *Flavobacterium sp.* | Uncharacterized protein |
| 192 | 293 | 0.01 | E4T450 | *Paludibacter propionicigenes* | UvrABC system protein A |
| 193 | 291 | 0.01 | A0A0Q7S584 | *Pelomonas sp.* | DNA-directed DNA polymerase (EC 2.7.7.7) |
| 194 | 291 | 0.01 | A0A0Q7SCT5 | *Pelomonas sp.* | Chromosome partition protein Smc |
| 195 | 291 | 0.01 | A0A1E4ACZ5 | *Chryseobacterium sp.* | TonB-dependent receptor |
| 196 | 291 | 0.01 | A0A1Q3RTV8 | *Flavobacterium sp.* | Sodium:proton antiporter |
| 197 | 289 | 0.01 | A0A0N8CUF4 | *Daphnia magna* | Uncharacterized protein (Fragment) |
| 198 | 288 | 0.01 | A0A1Q3RZ42 | *Flavobacterium sp.* | Cell envelope biogenesis protein OmpA |
| 199 | 288 | 0.01 | A0A1T5GFF7 | *Sphingobacterium nematocida* | Gliding motility-associated C-terminal domain-containing protein |
| 200 | 287 | 0.01 | A0A0L8VCE1 | *Sunxiuqinia dokdonensis* | Multidrug transporter AcrB |
| 201 | 286 | 0.01 | A0A1Q3M3K5 | *Bacteroidales bacterium* | Alpha-mannosidase |
| 202 | 285 | 0.01 | A0A0P6BFP2 | *Daphnia magna* | Uncharacterized protein |

| | | | | | |
|---|---|---|---|---|---|
| 203 | 285 | 0.01 | A0A1E4A183 | *Chryseobacterium sp.* | Tyrosine protein kinase |
| 204 | 285 | 0.01 | Q8HEC4 | *Caenorhabditis briggsae* | NADH-ubiquinone oxidoreductase chain 1 (EC 1.6.5.3) (NADH dehydrogenase subunit 1) |
| 205 | 284 | 0.01 | A0A1E3ZTP9 | *Chryseobacterium sp.* | Hybrid sensor histidine kinase/response regulator |
| 206 | 284 | 0.01 | A0A1Q3S9F5 | *Flavobacterium sp.* | DNA mismatch repair protein MutS |
| 207 | 283 | 0.01 | A0A1Q3S549 | *Flavobacterium sp.* | Phenylalanine--tRNA ligase beta subunit (EC 6.1.1.20) (Phenylalanyl-tRNA synthetase beta subunit) (PheRS) |
| 208 | 282 | 0.01 | A0A0Q7S6W6 | *Pelomonas sp.* | Indolepyruvate ferredoxin oxidoreductase (EC 1.2.7.8) |
| 209 | 282 | 0.01 | A0A1Q3SAJ2 | *Flavobacterium sp.* | Tail-specific protease |
| 210 | 280 | 0.01 | A0A1E4A600 | *Chryseobacterium sp.* | Recombinase |
| 211 | 280 | 0.01 | A0A1Q3M5E8 | *Bacteroidales bacterium* | Uncharacterized protein (Fragment) |
| 212 | 280 | 0.01 | A0A1Q3RZD7 | *Flavobacterium sp.* | ATP-dependent helicase |
| 213 | 280 | 0.01 | A0A1Q3S8N5 | *Flavobacterium sp.* | Uncharacterized protein |
| 214 | 279 | 0.01 | A0A1E3ZRN6 | *Chryseobacterium sp.* | Uncharacterized protein |
| 215 | 279 | 0.01 | N8QHE7 | *Acinetobacter bohemicus* | Uncharacterized protein |
| 216 | 278 | 0.01 | A0A0Q7SJM3 | *Pelomonas sp.* | tRNA modification GTPase MnmE (EC 3.6.-.-) |
| 217 | 278 | 0.01 | A0A1E4AD98 | *Chryseobacterium sp.* | Ribonuclease R (RNase R) (EC 3.1.13.1) |
| 218 | 278 | 0.01 | | | |
| 219 | 277 | 0.01 | A0A0Q7RZ30 | *Pelomonas sp.* | Acriflavin resistance protein |
| 220 | 277 | 0.01 | A0A0Q7RZF8 | *Pelomonas sp.* | Phosphoribosylformylglycinamidine synthase (FGAM synthase) (FGAMS) (EC 6.3.5.3) (Formylglycinamide ribonucleotide amidotransferase) (FGAR amidotransferase) (FGAR-AT) |
| 221 | 277 | 0.01 | A0A1G4VJQ3 | *Flavobacterium saliperosum* | Gliding motility-associated C-terminal domain-containing protein |
| 222 | 277 | 0.01 | A0A1M3MAE9 | *Paludibacter sp.* | DNA-directed RNA polymerase subunit beta' (RNAP subunit beta') (EC 2.7.7.6) (RNA polymerase subunit beta') (Transcriptase subunit beta') |
| 223 | 275 | 0.01 | A0A1Q3S511 | *Flavobacterium sp.* | Heat-shock protein Hsp70 |
| 224 | 275 | 0.01 | Q5CWD9 | *Cryptosporidium parvum* | Predicted secreted protein, signal peptide with several threonines, possible mucin |
| 225 | 274 | 0.01 | S2WV93 | *Delftia acidovorans* | Uncharacterized protein |
| 226 | 273 | 0.01 | A0A0Q6T477 | *Pelomonas sp.* | Uncharacterized protein |
| 227 | 273 | 0.01 | A0A0Q8H5F4 | *Pelomonas sp.* | TonB-dependent receptor |
| 228 | 272 | 0.01 | A0A0Q6UIJ2 | *Pelomonas sp.* | Nitrate reductase |
| 229 | 271 | 0.01 | A0A0Q6U523 | *Pelomonas sp.* | Nitric oxide reductase large subunit |
| 230 | 271 | 0.01 | A0A0Q7SA64 | *Pelomonas sp.* | Transcription-repair-coupling factor (TRCF) (EC 3.6.4.-) |
| 231 | 271 | 0.01 | A0A1E4A395 | *Chryseobacterium sp.* | Peptidase S46 |
| 232 | 271 | 0.01 | A0A1J5HML6 | *Porphyromonadaceae bacterium* | Penicillin-binding protein |

| | | | | | |
|---|---|---|---|---|---|
| 233 | 271 | 0.01 | A0A1Q3RWQ7 | *Flavobacterium sp.* | Cation/H(+) antiporter |
| 234 | 270 | 0.01 | A0A0Q8G4H5 | *Pelomonas sp.* | Elongation factor Tu (EF-Tu) |
| 235 | 270 | 0.01 | A0A1Q3RZB2 | *Flavobacterium sp.* | Uncharacterized protein |
| 236 | 270 | 0.01 | E4T5C1 | *Paludibacter propionicigenes* | DNA topoisomerase IV subunit A (EC 5.99.1.3) |
| 237 | 268 | 0.01 | | | |
| 238 | 267 | 0.01 | A0A1Q3RWZ9 | *Flavobacterium sp.* | Uncharacterized protein |
| 239 | 266 | 0.01 | E4T570 | *Paludibacter propionicigenes* | DNA-directed RNA polymerase subunit beta (RNAP subunit beta) (EC 2.7.7.6) (RNA polymerase subunit beta) (Transcriptase subunit beta) |
| 240 | 265 | 0.01 | A0A1E3ZRQ0 | *Chryseobacterium sp.* | Uncharacterized protein |
| 241 | 265 | 0.01 | A0A1E3ZY95 | *Chryseobacterium sp.* | Uncharacterized protein |
| 242 | 265 | 0.01 | A0A1E4AGS9 | *Chryseobacterium sp.* | TonB-dependent receptor |
| 243 | 265 | 0.01 | A0A1Q3S6Q0 | *Flavobacterium sp.* | DNA polymerase I (EC 2.7.7.7) |
| 244 | 264 | 0.01 | A0A1E4A8B6 | *Chryseobacterium sp.* | Type II and III secretion system protein |
| 245 | 264 | 0.01 | A0A1M3MBI7 | *Paludibacter sp.* | Cell surface protein SprA |
| 246 | 264 | 0.01 | A0A1Q3M609 | *Bacteroidales bacterium* | Uncharacterized protein |
| 247 | 264 | 0.01 | H8KUZ8 | *Solitalea canadensis* | Uncharacterized protein |
| 248 | 263 | 0.01 | A0A077XLW7 | *Sphinx1.76-related DNA* | Replication protein |
| 249 | 263 | 0.01 | A0A1E4AEN9 | *Chryseobacterium sp.* | Peptidase S9 |
| 250 | 262 | 0.01 | A0A0Q6UG42 | *Pelomonas sp.* | Pyruvate ferredoxin oxidoreductase |
| 251 | 261 | 0.01 | A0A1E4AAQ4 | *Chryseobacterium sp.* | Copper oxidase |
| 252 | 261 | 0.01 | A1W5C2 | *Acidovorax sp.* | IstB domain protein ATP-binding protein |
| 253 | 261 | 0.01 | U6RAE1 | *Bacteroides sp.* | Uncharacterized protein |
| 254 | 260 | 0.01 | A0A0Q7T0F9 | *Pelomonas sp.* | Lon protease (EC 3.4.21.53) (ATP-dependent protease La) |
| 255 | 260 | 0.01 | A0A1Q3S8T7 | *Flavobacterium sp.* | Nucleotidyltransferase |
| 256 | 260 | 0.01 | E4T1A7 | *Paludibacter propionicigenes* | Pyruvate phosphate dikinase PEP/pyruvate-binding protein |
| 257 | 259 | 0.01 | A0A1J5HGG6 | *Porphyromonadaceae bacterium* | DNA-directed RNA polymerase subunit beta (RNAP subunit beta) (EC 2.7.7.6) (RNA polymerase subunit beta) (Transcriptase subunit beta) |
| 258 | 258 | 0.01 | A0A1E4A7R6 | *Chryseobacterium sp.* | Uncharacterized protein |
| 259 | 258 | 0.01 | A0A1Q3S6R5 | *Flavobacterium sp.* | Uncharacterized protein |
| 260 | 257 | 0.01 | A0A1Q3S1W4 | *Flavobacterium sp.* | Sulfatase |
| 261 | 257 | 0.01 | K0XEC8 | *Lachnoanaerobaculum sp.* | Uncharacterized protein |
| 262 | 256 | 0.01 | A0A0Q7SFB9 | *Pelomonas sp.* | Multidrug transporter |
| 263 | 256 | 0.01 | A0A0Q8GR06 | *Pelomonas sp.* | Alpha-2-macroglobulin |
| 264 | 256 | 0.01 | A0A1E4AC51 | *Chryseobacterium sp.* | Helicase |
| 265 | 256 | 0.01 | E4T2N1 | *Paludibacter propionicigenes* | DNA gyrase subunit A (EC 5.99.1.3) |
| 266 | 255 | 0.01 | A0A0P6IS02 | *Daphnia magna* | Uncharacterized protein |

| | | | | | |
|---|---|---|---|---|---|
| 267 | 254 | 0.01 | A0A023UNG0 | *Magnusiomyces ingens* | Uncharacterized protein (Fragment) |
| 268 | 254 | 0.01 | A0A0D6TL26 | *Flavobacterium sp.* | Metalloprotease |
| 269 | 254 | 0.01 | A0A1E3ZX97 | *Chryseobacterium sp.* | Gliding motility protein |
| 270 | 253 | 0.01 | A0A1Q3S6W5 | *Flavobacterium sp.* | Peptidase M16 |
| 271 | 253 | 0.01 | | | |
| 272 | 252 | 0.01 | A0A0P0UB73 | *Flavobacterium psychrophilum* | Multidrug transporter AcrB |
| 273 | 252 | 0.01 | A0A0Q7SE50 | *Pelomonas sp.* | Bifunctional glutamine synthetase adenylyltransferase/adenylyl-removing enzyme (ATP:glutamine synthetase adenylyltransferase) (ATase) [Includes: Glutamine synthetase adenylyl-L-tyrosine phosphorylase (EC 2.7.7.89) (Adenylyl removase) (AR) (AT-N); Glutamine synthetase adenylyl transferase (EC 2.7.7.42) (Adenylyl transferase) (AT) (AT-C)] |
| 274 | 252 | 0.01 | A0A0Q7SEF7 | *Pelomonas sp.* | Translation initiation factor IF-2 |
| 275 | 252 | 0.01 | A0A1E3ZYM1 | *Chryseobacterium sp.* | Peptidylprolyl isomerase (EC 5.2.1.8) |
| 276 | 252 | 0.01 | A0A1E4A473 | *Chryseobacterium sp.* | Penicillin-binding protein 2 |
| 277 | 252 | 0.01 | A0A1Q3RY20 | *Flavobacterium sp.* | Collagen-binding protein |
| 278 | 252 | 0.01 | E4T2W8 | *Paludibacter propionicigenes* | Pyruvate carboxylase (EC 6.4.1.1) |
| 279 | 251 | 0.01 | A0A0Q6SZK0 | *Pelomonas sp.* | Protein translocase subunit SecA |
| 280 | 251 | 0.01 | A0A0Q7S238 | *Pelomonas sp.* | Histidine kinase |
| 281 | 251 | 0.01 | A0A1Q3S878 | *Flavobacterium sp.* | Patatin |
| 282 | 250 | 0.01 | A0A0D5CAE6 | *Oscheius chongmingensis* | NADH-ubiquinone oxidoreductase chain 4 (EC 1.6.5.3) |
| 283 | 250 | 0.01 | A0A0Q7SD92 | *Pelomonas sp.* | Uncharacterized protein |
| 284 | 250 | 0.01 | A0A0Q7SH01 | *Pelomonas sp.* | Lysine decarboxylase |
| 285 | 250 | 0.01 | A0A1E4ACD4 | *Chryseobacterium sp.* | Gliding motility lipoprotein GldJ |
| 286 | 250 | 0.01 | E4T571 | *Paludibacter propionicigenes* | DNA-directed RNA polymerase subunit beta' (RNAP subunit beta') (EC 2.7.7.6) (RNA polymerase subunit beta') (Transcriptase subunit beta') |
| 287 | 249 | 0.01 | A0A0Q7SEL2 | *Pelomonas sp.* | Uncharacterized protein |
| 288 | 249 | 0.01 | A0A1E4A2H0 | *Chryseobacterium sp.* | Peptidylprolyl isomerase |
| 289 | 249 | 0.01 | A0A1Q3S481 | *Flavobacterium sp.* | Uncharacterized protein |
| 290 | 248 | 0.01 | A0A1E4ABW2 | *Chryseobacterium sp.* | Peptidase M1 |
| 291 | 247 | 0.01 | A0A1E4A9N2 | *Chryseobacterium sp.* | Uncharacterized protein |
| 292 | 247 | 0.01 | A0A1L4BMB6 | *Didymosphenia geminata* | NADH dehydrogenase subunit 7 |
| 293 | 246 | 0.01 | A0A0Q7S7S5 | *Pelomonas sp.* | Valine--tRNA ligase (EC 6.1.1.9) (Valyl-tRNA synthetase) (ValRS) |
| 294 | 246 | 0.01 | E4T2R0 | *Paludibacter propionicigenes* | ATPase AAA-2 domain protein |
| 295 | 245 | 0.01 | A0A0Q7SRF6 | *Pelomonas sp.* | DNA topoisomerase III |
| 296 | 245 | 0.01 | A0A0Q8H4A2 | *Pelomonas sp.* | Uncharacterized protein |
| 297 | 245 | 0.01 | A0A1E4A0Q8 | *Chryseobacterium sp.* | Polysaccharide biosynthesis protein |

| | | | | | |
|---|---|---|---|---|---|
| 298 | 245 | 0.01 | A0A1Q3RZ46 | *Flavobacterium sp.* | Uncharacterized protein |
| 299 | 245 | 0.01 | A0A1Q3S3S5 | *Flavobacterium sp.* | Uncharacterized protein |
| 300 | 244 | 0.01 | K9L3F4 | *Magnusiomyces magnusii* | NADH-ubiquinone oxidoreductase chain 1 (EC 1.6.5.3) |
| 301 | 243 | 0.01 | E4T7X4 | *Paludibacter propionicigenes* | Alanine--tRNA ligase (EC 6.1.1.7) (Alanyl-tRNA synthetase) (AlaRS) |
| 302 | 242 | 0.01 | A0A0Q7T172 | *Pelomonas sp.* | Ribonucleoside-diphosphate reductase (EC 1.17.4.1) |
| 303 | 242 | 0.01 | A0A0Q8H242 | *Pelomonas sp.* | PAS domain-containing sensor histidine kinase |
| 304 | 242 | 0.01 | A0A1M3DNS5 | *Bacteroidales bacterium* | Putative K(+)-stimulated pyrophosphate-energized sodium pump (EC 3.6.1.1) (Membrane-bound sodium-translocating pyrophosphatase) (Pyrophosphate-energized inorganic pyrophosphatase) (Na(+)-PPase) |
| 305 | 242 | 0.01 | A0A1M3MCY6 | *Paludibacter sp.* | Glycosyl hydrolase |
| 306 | 242 | 0.01 | F3PIP2 | *Bacteroides clarus* | TonB-dependent receptor plug domain protein |
| 307 | 241 | 0.01 | A0A0Q7SKX9 | *Pelomonas sp.* | ATP-dependent helicase |
| 308 | 241 | 0.01 | A0A0Q7T1P6 | *Pelomonas sp.* | 3-hydroxyacyl-CoA dehydrogenase |
| 309 | 241 | 0.01 | A0A0T5VPN0 | *Pedobacter ginsenosidimutans* | SusC/RagA family TonB-linked outer membrane protein |
| 310 | 241 | 0.01 | A0A142L3A0 | *Bacteroidetes bacterium* | DEAD/DEAH box helicase |
| 311 | 241 | 0.01 | A0A1Q3RTV6 | *Flavobacterium sp.* | Uncharacterized protein |
| 312 | 241 | 0.01 | A0A1Q3RX80 | *Flavobacterium sp.* | Uncharacterized protein |
| 313 | 241 | 0.01 | A0A1Q3S955 | *Flavobacterium sp.* | Polyphosphate kinase (EC 2.7.4.1) (ATP-polyphosphate phosphotransferase) (Polyphosphoric acid kinase) |
| 314 | 239 | 0.01 | A0A1Q3S8C6 | *Flavobacterium sp.* | Uncharacterized protein |
| 315 | 239 | 0.01 | A0A1Q3S9B2 | *Flavobacterium sp.* | Uncharacterized protein |
| 316 | 239 | 0.01 | B1N6J8 | *uncultured beta proteobacterium* | Putative uncharacterized protein |
| 317 | 239 | 0.01 | E4T676 | *Paludibacter propionicigenes* | Collagenase |
| 318 | 238 | 0.01 | A0A0D5C9J6 | *Oscheius chongmingensis* | NADH dehydrogenase subunit 5 |
| 319 | 238 | 0.01 | E4T653 | *Paludibacter propionicigenes* | Peptidoglycan glycosyltransferase (EC 2.4.1.129) |
| 320 | 238 | 0.01 | E4T8D5 | *Paludibacter propionicigenes* | Orn/DAP/Arg decarboxylase 2 |
| 321 | 237 | 0.01 | A0A0Q6WEF8 | *Pelomonas sp.* | Uncharacterized protein |
| 322 | 237 | 0.01 | A0A0Q7T6T8 | *Pelomonas sp.* | Histidine kinase |
| 323 | 237 | 0.01 | A0A1Q3S478 | *Flavobacterium sp.* | Conjugal transfer protein TraG |
| 324 | 236 | 0.01 | A0A023UP21 | *Magnusiomyces ingens* | Ribosomal protein S3 |
| 325 | 236 | 0.01 | A0A1Q3RXD8 | *Flavobacterium sp.* | UvrABC system protein A |
| 326 | 236 | 0.01 | E4T0X8 | *Paludibacter propionicigenes* | DNA helicase (EC 3.6.4.12) |
| 327 | 235 | 0.01 | A0A0Q6TP10 | *Pelomonas sp.* | Uncharacterized protein |
| 328 | 234 | 0.01 | A0A1J5I0W6 | *Porphyromonadaceae bacterium* | DNA mismatch repair protein MutS |

| | | | | | |
|---|---|---|---|---|---|
| 329 | 234 | 0.01 | A0A1Q3S0W7 | *Flavobacterium sp.* | Uncharacterized protein |
| 330 | 234 | 0.01 | A0A1Q3SAK4 | *Flavobacterium sp.* | TonB-dependent receptor |
| 331 | 233 | 0.01 | A0A0Q7S4W2 | *Pelomonas sp.* | FAD-linked oxidase |
| 332 | 233 | 0.01 | A0A0Q7T373 | *Pelomonas sp.* | Ribonuclease R (RNase R) (EC 3.1.13.1) |
| 333 | 233 | 0.01 | A0A1J5HBV2 | *Porphyromonadaceae bacterium* | Uncharacterized protein |
| 334 | 233 | 0.01 | A0A1Q3SB52 | *Flavobacterium sp.* | Acriflavin resistance protein |
| 335 | 233 | 0.01 | E4T2H4 | *Paludibacter propionicigenes* | Phosphoribosylformylglycinamidine synthase (FGAM synthase) (FGAMS) (EC 6.3.5.3) (Formylglycinamide ribonucleotide amidotransferase) (FGAR amidotransferase) (FGAR-AT) |
| 336 | 233 | 0.01 | E4T3X9 | *Paludibacter propionicigenes* | DNA topoisomerase III (EC 5.99.1.2) |
| 337 | 231 | 0.01 | A0A0Q6T5H8 | *Pelomonas sp.* | Nodulation protein NolG |
| 338 | 231 | 0.01 | A0A0Q6XBI6 | *Pelomonas sp.* | Penicillin-binding protein 1C |
| 339 | 231 | 0.01 | A0A0Q7SN01 | *Pelomonas sp.* | ATPase |
| 340 | 231 | 0.01 | A0A0Q7SRK7 | *Pelomonas sp.* | Leucine--tRNA ligase (EC 6.1.1.4) (Leucyl-tRNA synthetase) (LeuRS) |
| 341 | 231 | 0.01 | I4ZNR1 | *Acinetobacter sp.* | Uncharacterized protein (Fragment) |
| 342 | 230 | 0.01 | E4T3F8 | *Paludibacter propionicigenes* | L-glutamine synthetase (EC 6.3.1.2) |
| 343 | 229 | 0.01 | D7W2K3 | *Chryseobacterium gleum* | Uncharacterized protein |
| 344 | 228 | 0.01 | A0A0Q7SLN7 | *Pelomonas sp.* | UvrABC system protein A (UvrA protein) (Excinuclease ABC subunit A) |
| 345 | 228 | 0.01 | A0A1E3ZYZ1 | *Chryseobacterium sp.* | Uncharacterized protein |
| 346 | 228 | 0.01 | E4T1U6 | *Paludibacter propionicigenes* | Carbamoyl-phosphate synthase L chain ATP-binding protein (EC 6.4.1.1) |
| 347 | 228 | 0.01 | I2JQL4 | *Brettanomyces bruxellensis* | Uncharacterized protein |
| 348 | 227 | 0.01 | A0A023UP13 | *Magnusiomyces ingens* | NADH-ubiquinone oxidoreductase chain 4 (EC 1.6.5.3) |
| 349 | 227 | 0.01 | A0A023UPL4 | *Magnusiomyces ingens* | Uncharacterized protein (Fragment) |
| 350 | 227 | 0.01 | A0A0M8MHN4 | *Flavobacterium akiainvivens* | Elongation factor G (EF-G) |
| 351 | 227 | 0.01 | A0A0Q7SAX3 | *Pelomonas sp.* | Glycine dehydrogenase (decarboxylating) (EC 1.4.4.2) (Glycine cleavage system P-protein) (Glycine decarboxylase) (Glycine dehydrogenase (aminomethyl-transferring)) |
| 352 | 227 | 0.01 | A0A1E4ABE8 | *Chryseobacterium sp.* | Methionine synthase |
| 353 | 227 | 0.01 | A0A1Q3RY34 | *Flavobacterium sp.* | TonB-dependent receptor |
| 354 | 227 | 0.01 | D8D4W5 | *Comamonas testosteroni* | Uncharacterized protein |
| 355 | 226 | 0.01 | A0A1B4SJP8 | *Burkholderia sp.* | Uncharacterized protein |
| 356 | 226 | 0.01 | A0A1E4A3F0 | *Chryseobacterium sp.* | DNA topoisomerase IV |
| 357 | 226 | 0.01 | A0A1M3DJX9 | *Bacteroidales bacterium* | Beta-galactosidase (EC 3.2.1.23) (Lactase) |
| 358 | 226 | 0.01 | A0A1Q3SAW0 | *Flavobacterium sp.* | Sugar hydrolase |
| 359 | 226 | 0.01 | | | |

| | | | | | |
|---|---|---|---|---|---|
| 360 | 225 | 0.01 | A0A0Q6WQC8 | *Pelomonas sp.* | Uncharacterized protein |
| 361 | 225 | 0.01 | A0A0Q7SCD0 | *Pelomonas sp.* | NADPH-dependent 2,4-dienoyl-CoA reductase (EC 1.3.1.34) |
| 362 | 225 | 0.01 | A0A0T5VPL4 | *Pedobacter ginsenosidimutans* | Uncharacterized protein |
| 363 | 225 | 0.01 | A0A1Q3S1U2 | *Flavobacterium sp.* | Phosphoribosylformylglycinamidine synthase (FGAM synthase) (FGAMS) (EC 6.3.5.3) (Formylglycinamide ribonucleotide amidotransferase) (FGAR amidotransferase) (FGAR-AT) |
| 364 | 225 | 0.01 | T1JK00 | *Strigamia maritima* | Uncharacterized protein |
| 365 | 224 | 0.01 | A0A1E4AGP3 | *Chryseobacterium sp.* | TonB-dependent receptor |
| 366 | 224 | 0.01 | A0A1E4AQU3 | *Chryseobacterium sp.* | Phage tail protein |
| 367 | 224 | 0.01 | A0A1J5HIK6 | *Porphyromonadaceae bacterium* | UDP-N-acetylglucosamine 2-epimerase |
| 368 | 224 | 0.01 | A0A1Q3RW26 | *Flavobacterium sp.* | Isocitrate dehydrogenase (NADP(+)) |
| 369 | 224 | 0.01 | A0A1Q3RX31 | *Flavobacterium sp.* | Copper-translocating P-type ATPase |
| 370 | 224 | 0.01 | A0A1Q3S1B3 | *Flavobacterium sp.* | S9 family peptidase |
| 371 | 224 | 0.01 | A0A1Q3S421 | *Flavobacterium sp.* | Uncharacterized protein (Fragment) |
| 372 | 224 | 0.01 | A0A1Q3SA66 | *Flavobacterium sp.* | Polyribonucleotide nucleotidyltransferase (EC 2.7.7.8) (Polynucleotide phosphorylase) (PNPase) |
| 373 | 224 | 0.01 | D8L7S1 | *Paramecium caudatum* | NADH-ubiquinone oxidoreductase chain 5 (EC 1.6.5.3) |
| 374 | 224 | 0.01 | R6DEG5 | *Bacteroides sp.* | RagB/SusD domain-containing protein |
| 375 | 223 | 0.01 | A0A0Q7S0G4 | *Pelomonas sp.* | AAA family ATPase |
| 376 | 223 | 0.01 | A0A0Q7SUY8 | *Pelomonas sp.* | Enoyl-CoA hydratase |
| 377 | 223 | 0.01 | A0A1E4AH32 | *Chryseobacterium sp.* | Endothelin-converting protein |
| 378 | 223 | 0.01 | A0A1Q3SAU5 | *Flavobacterium sp.* | S9 family peptidase |
| 379 | 222 | 0.01 | A0A0Q7SY69 | *Pelomonas sp.* | DNA helicase (EC 3.6.4.12) |
| 380 | 222 | 0.01 | A0A1Q3RU83 | *Flavobacterium sp.* | Primosomal protein N' (EC 3.6.4.-) (ATP-dependent helicase PriA) |
| 381 | 222 | 0.01 | A0A1Q3S9I3 | *Flavobacterium sp.* | Type IV secretion protein Rhs |
| 382 | 221 | 0.01 | A0A073CD69 | *Planktothrix agardhii* | Uncharacterized protein |
| 383 | 221 | 0.01 | A0A0Q8GMD4 | *Pelomonas sp.* | Uncharacterized protein |
| 384 | 221 | 0.01 | A0A1Q3S4W4 | *Flavobacterium sp.* | Oligopeptidase B |
| 385 | 220 | 0.01 | A0A0Q7T0V8 | *Pelomonas sp.* | Sodium:solute symporter |
| 386 | 220 | 0.01 | A0A178FWP5 | *Acinetobacter sp.* | Transposase |
| 387 | 220 | 0.01 | A0A1M3M9Z9 | *Paludibacter sp.* | Uncharacterized protein |
| 388 | 220 | 0.01 | A0A1Q3S6U9 | *Flavobacterium sp.* | Uncharacterized protein |
| 389 | 220 | 0.01 | K9L3A6 | *Magnusiomyces magnusii* | NADH-ubiquinone oxidoreductase chain 5 (EC 1.6.5.3) |
| 390 | 219 | 0.01 | A0A0Q5LD55 | *Acidovorax sp.* | Transposase (Fragment) |
| 391 | 219 | 0.01 | A0A0Q6TGN2 | *Pelomonas sp.* | DNA helicase (EC 3.6.4.12) |

| | | | | | |
|---|---|---|---|---|---|
| 392 | 219 | 0.01 | A0A0Q6WDL9 | *Pelomonas sp.* | Serine/threonine protein kinase |
| 393 | 219 | 0.01 | A0A0Q6WDT0 | *Pelomonas sp.* | Copper-transporting ATPase |
| 394 | 219 | 0.01 | A0A0Q7ST55 | *Pelomonas sp.* | Guanosine-3',5'-bis(Diphosphate) 3'-pyrophosphohydrolase |
| 395 | 219 | 0.01 | A0A0R0LL75 | *Ferrovum sp.* | N-6 DNA methylase |
| 396 | 219 | 0.01 | E4T3R5 | *Paludibacter propionicigenes* | Phenylalanine--tRNA ligase beta subunit (EC 6.1.1.20) (Phenylalanyl-tRNA synthetase beta subunit) (PheRS) |
| 397 | 218 | 0.01 | A0A0Q6TCI2 | *Pelomonas sp.* | Uncharacterized protein |
| 398 | 218 | 0.01 | A0A0Q7S0S4 | *Pelomonas sp.* | GTP pyrophosphokinase |
| 399 | 218 | 0.01 | A0A0Q7SL61 | *Pelomonas sp.* | 2-oxoglutarate dehydrogenase subunit E1 (EC 1.2.4.2) |
| 400 | 218 | 0.01 | A0A133PWA7 | *Prevotella corporis* | Transposase |
| 401 | 218 | 0.01 | A0A1E3ZWT8 | *Chryseobacterium sp.* | Translation initiation factor IF-2 |
| 402 | 218 | 0.01 | A0A1Q3RUB8 | *Flavobacterium sp.* | TonB-dependent receptor |
| 403 | 218 | 0.01 | A0A1Q3S932 | *Flavobacterium sp.* | Uncharacterized protein |
| 404 | 216 | 0.01 | A0A0Q7SL81 | *Pelomonas sp.* | Uncharacterized protein |
| 405 | 216 | 0.01 | A0A0Q7T0L4 | *Pelomonas sp.* | Primosomal protein N' (EC 3.6.4.-) (ATP-dependent helicase PriA) |
| 406 | 216 | 0.01 | A0A1E4ABB0 | *Chryseobacterium sp.* | Homoserine dehydrogenase |
| 407 | 216 | 0.01 | A0A1G0M577 | *Geobacteraceae bacterium* | Uncharacterized protein |
| 408 | 216 | 0.01 | E4T386 | *Paludibacter propionicigenes* | Heterodimeric methylmalonyl-CoA mutase large subunit (EC 5.4.99.2) |
| 409 | 216 | 0.01 | S3C5M7 | *Ophiostoma piceae* | Uncharacterized protein |
| 410 | 215 | 0.01 | A0A0Q7SDY4 | *Pelomonas sp.* | GCN5 family acetyltransferase |
| 411 | 215 | 0.01 | A0A1E4ADX4 | *Chryseobacterium sp.* | BatD protein |
| 412 | 215 | 0.01 | A0A1Q3S9E5 | *Flavobacterium sp.* | Signal peptide peptidase SppA |
| 413 | 214 | 0.01 | A0A0H5Q1D3 | *uncultured prokaryote* | Uncharacterized protein |
| 414 | 214 | 0.01 | A0A0Q7SNJ9 | *Pelomonas sp.* | Cyanophycin synthetase |
| 415 | 214 | 0.01 | A0A1Q3LBC0 | *Bacteroidales bacterium 45-6* | Uncharacterized protein |
| 416 | 214 | 0.01 | E4T1S7 | *Paludibacter propionicigenes* | Carbamoyl-phosphate synthase small chain (EC 6.3.5.5) (Carbamoyl-phosphate synthetase glutamine chain) |
| 417 | 213 | 0.01 | A0A0Q7SL07 | *Pelomonas sp.* | DNA polymerase I (EC 2.7.7.7) |
| 418 | 213 | 0.01 | A0A1J5HP94 | *Porphyromonadaceae bacterium* | UPF0313 protein AUK44_04705 |
| 419 | 213 | 0.01 | A0A1Q3RVM0 | *Flavobacterium sp.* | Flagellar motor protein MotB |
| 420 | 213 | 0.01 | A0A1Q3RW11 | *Flavobacterium sp.* | Magnesium-translocating P-type ATPase |
| 421 | 212 | 0.01 | A0A1E4AQU9 | *Chryseobacterium sp.* | Uncharacterized protein |
| 422 | 212 | 0.01 | A0A1Q3M5Y6 | *Bacteroidales bacterium* | Uncharacterized protein |
| 423 | 212 | 0.01 | I2F7Y4 | *Mesotoga prima* | Retron-type reverse transcriptase |
| 424 | 212 | 0.01 | | | |

| | | | | | |
|---|---|---|---|---|---|
| 425 | 211 | 0.01 | A0A0Q6TFR1 | *Pelomonas sp.* | Uncharacterized protein |
| 426 | 211 | 0.01 | A0A0Q7S182 | *Pelomonas sp.* | Phenylalanine--tRNA ligase beta subunit (EC 6.1.1.20) (Phenylalanyl-tRNA synthetase beta subunit) (PheRS) |
| 427 | 211 | 0.01 | A0A1E4AFH6 | *Chryseobacterium sp.* | Leucine--tRNA ligase (EC 6.1.1.4) (Leucyl-tRNA synthetase) (LeuRS) |
| 428 | 211 | 0.01 | A0A1M6UHC8 | *Chryseobacterium molle* | Uncharacterized protein |
| 429 | 211 | 0.01 | E4T682 | *Paludibacter propionicigenes* | UvrABC system protein C (Protein UvrC) (Excinuclease ABC subunit C) |
| 430 | 211 | 0.01 | G7ZJ76 | *Azospirillum lipoferum* | Uncharacterized protein |
| 431 | 211 | 0.01 | | | |
| 432 | 210 | 0.01 | A0A1E4A3S3 | *Chryseobacterium sp.* | Glycosyl transferase |
| 433 | 210 | 0.01 | A0A1J5HSL1 | *Porphyromonadaceae bacterium* | Pyruvate-flavodoxin oxidoreductase (EC 1.2.7.-) |
| 434 | 210 | 0.01 | A0A1Q3RZR6 | *Flavobacterium sp.* | Flagellar motor protein MotB |
| 435 | 210 | 0.01 | W1I7F0 | *uncultured bacterium* | Uncultured bacterium extrachromosomal DNA RGI01790 |
| 436 | 209 | 0.01 | A0A1E3ZZS5 | *Chryseobacterium sp.* | Amino acid transporter |
| 437 | 209 | 0.01 | A0A1F3LNZ9 | *Bacteroidetes bacterium* | Glucarate dehydratase |
| 438 | 209 | 0.01 | A0A1J5HRY1 | *Porphyromonadaceae bacterium* | Cell surface protein SprA |
| 439 | 209 | 0.01 | A0A1Q3S4I8 | *Flavobacterium sp.* | Uncharacterized protein |
| 440 | 209 | 0.01 | | | |
| 441 | 209 | 0.01 | | | |
| 442 | 208 | 0.01 | A0A023UMR4 | *Magnusiomyces ingens* | NADH-ubiquinone oxidoreductase chain 3 (EC 1.6.5.3) |
| 443 | 208 | 0.01 | A0A0Q6T0C3 | *Pelomonas sp.* | DNA repair protein RecN (Recombination protein N) |
| 444 | 208 | 0.01 | A0A0Q7SDQ5 | *Pelomonas sp.* | Efflux pump membrane transporter |
| 445 | 208 | 0.01 | A0A1E4ELA1 | *Paludibacter sp.* | Dihydrodipicolinate synthase family protein |
| 446 | 208 | 0.01 | A0A1Q3SA76 | *Flavobacterium sp.* | Gliding motility protein GldM |
| 447 | 208 | 0.01 | E4T409 | *Paludibacter propionicigenes* | Mg chelatase, subunit ChlI |
| 448 | 208 | 0.01 | E4T5B8 | *Paludibacter propionicigenes* | Translation elongation factor 2 (EF-2/EF-G) |
| 449 | 207 | 0.01 | A0A0Q5UMI6 | *Flavobacterium sp.* | Uncharacterized protein |
| 450 | 207 | 0.01 | A0A0Q7SNK2 | *Pelomonas sp.* | Short-chain dehydrogenase |
| 451 | 207 | 0.01 | A0A0Q8H0M9 | *Pelomonas sp.* | Uncharacterized protein |
| 452 | 207 | 0.01 | A0A1E4A4U4 | *Chryseobacterium sp.* | Uncharacterized protein |
| 453 | 207 | 0.01 | A0A1E4ACP7 | *Chryseobacterium sp.* | Sodium:solute symporter |
| 454 | 207 | 0.01 | A0A1E4AGM4 | *Chryseobacterium sp.* | Radical SAM protein |
| 455 | 207 | 0.01 | A0A1Q3S796 | *Flavobacterium sp.* | Dehydrogenase |
| 456 | 207 | 0.01 | E4T4N5 | *Paludibacter propionicigenes* | 2-succinyl-5-enolpyruvyl-6-hydroxy-3-cyclohexene-1-carboxylate synthase (SEPHCHC synthase) (EC 2.2.1.9) (Menaquinone biosynthesis protein MenD) |

| | | | | | |
|---|---|---|---|---|---|
| 457 | 207 | 0.01 | N8VN75 | *Acinetobacter sp.* | Uncharacterized protein |
| 458 | 206 | 0.01 | A0A0K0WVZ1 | *Litoditis aff. marina PmIV* | NADH dehydrogenase subunit 2 |
| 459 | 206 | 0.01 | A0A0Q6W7U3 | *Pelomonas sp.* | Uncharacterized protein |
| 460 | 206 | 0.01 | A0A0Q7S0T9 | *Pelomonas sp.* | MFS transporter |
| 461 | 206 | 0.01 | A0A1E4AQS9 | *Chryseobacterium sp.* | Uncharacterized protein |
| 462 | 206 | 0.01 | A0A1Q3RU69 | *Flavobacterium sp.* | Glycosyl transferase family 2 |
| 463 | 206 | 0.01 | E4T5N8 | *Paludibacter propionicigenes* | UvrABC system protein A |
| 464 | 205 | 0.01 | A0A0Q6TH52 | *Pelomonas sp.* | Uncharacterized protein |
| 465 | 205 | 0.01 | A0A1E4A4H6 | *Chryseobacterium sp.* | Uncharacterized protein |
| 466 | 205 | 0.01 | A0A1E4A7Y4 | *Chryseobacterium sp.* | Uncharacterized protein |
| 467 | 205 | 0.01 | A0A1Q3S6T7 | *Flavobacterium sp.* | Cytochrome C |
| 468 | 205 | 0.01 | A0A1Q3SA51 | *Flavobacterium sp.* | Gliding motility lipoprotein GldK |
| 469 | 205 | 0.01 | A0A1Q3SAE9 | *Flavobacterium sp.* | Penicillin-binding protein |
| 470 | 205 | 0.01 | A0A1Q3SAG2 | *Flavobacterium sp.* | Aminopeptidase |
| 471 | 205 | 0.01 | E4T592 | *Paludibacter propionicigenes* | Methionine synthase (B12-dependent) (EC 2.1.1.13) |
| 472 | 205 | 0.01 | P15689 | *Paramecium tetraurelia* | NADH-ubiquinone oxidoreductase 49 kDa subunit (EC 1.6.5.3) (EC 1.6.99.3) (NADH dehydrogenase subunit 7) |
| 473 | 205 | 0.01 | | | |
| 474 | 204 | 0.01 | A0A0Q7SJP7 | *Pelomonas sp.* | Phosphoenolpyruvate carboxylase (PEPC) (PEPCase) (EC 4.1.1.31) |
| 475 | 204 | 0.01 | T1KH75 | *Tetranychus urticae* | Mitochondrial NADH-quinone oxidoreductase 49 kDa subunit (EC 1.6.5.11) |
| 476 | 203 | 0.01 | A0A0Q6WW26 | *Pelomonas sp.* | Uncharacterized protein |
| 477 | 203 | 0.01 | A0A0Q7SA36 | *Pelomonas sp.* | Protein translocase subunit SecY |
| 478 | 203 | 0.01 | A0A1E4ADB7 | *Chryseobacterium sp.* | ATP-dependent RNA helicase |
| 479 | 203 | 0.01 | A0A1J5HA84 | *Porphyromonadaceae bacterium* | Potassium transporter |
| 480 | 203 | 0.01 | A0A1M3M7V7 | *Paludibacter sp.* | Carbamoyl-phosphate synthase (glutamine-hydrolyzing) (EC 6.3.5.5) |
| 481 | 203 | 0.01 | A0A1Q3RTW4 | *Flavobacterium sp.* | Two-component sensor histidine kinase |
| 482 | 202 | 0.01 | A0A085EMB6 | *Flavobacterium gilvum* | Uncharacterized protein |
| 483 | 202 | 0.01 | A0A0Q6U3Y0 | *Pelomonas sp.* | Uncharacterized protein |
| 484 | 202 | 0.01 | A0A0Q8GKD4 | *Pelomonas sp.* | Uncharacterized protein |
| 485 | 202 | 0.01 | A0A1E3ZYN2 | *Chryseobacterium sp.* | Uncharacterized protein |
| 486 | 202 | 0.01 | A0A1E4ACN4 | *Chryseobacterium sp.* | Peptidase S8 |
| 487 | 202 | 0.01 | A0A1Q3S2M1 | *Flavobacterium sp.* | Uncharacterized protein |
| 488 | 202 | 0.01 | A0A1Q3S6U3 | *Flavobacterium sp.* | Gliding motility protein RemB |
| 489 | 201 | 0.01 | A0A0Q6UIQ9 | *Pelomonas sp.* | Histidine kinase (EC 2.7.13.3) |
| 490 | 201 | 0.01 | A0A0Q8GXY5 | *Pelomonas sp.* | Uncharacterized protein |

| | | | | | |
|---|---|---|---|---|---|
| 491 | 201 | 0.01 | A0A1E4A1F8 | *Chryseobacterium sp.* | Cell division protein FtsZ |
| 492 | 201 | 0.01 | A0A1E4AHA9 | *Chryseobacterium sp.* | Glycerol acyltransferase |
| 493 | 201 | 0.01 | A0A1J5HT60 | *Porphyromonadaceae bacterium* | DNA polymerase I (EC 2.7.7.7) |
| 494 | 201 | 0.01 | A0A1J5HUJ2 | *Porphyromonadaceae bacterium* | Cytochrome C oxidase assembly protein |
| 495 | 200 | 0.01 | A0A0Q7RYM2 | *Pelomonas sp.* | Copper oxidase |
| 496 | 200 | 0.01 | A0A0Q7SCD2 | *Pelomonas sp.* | Helicase |
| 497 | 200 | 0.01 | A0A0Q7SED5 | *Pelomonas sp.* | Alanine--tRNA ligase (EC 6.1.1.7) (Alanyl-tRNA synthetase) (AlaRS) |
| 498 | 200 | 0.01 | A0A1E4A2L4 | *Chryseobacterium sp.* | Flagellar motor protein MotB |
| 499 | 200 | 0.01 | A0A1E4A659 | *Chryseobacterium sp.* | Peptidase M61 |
| 500 | 200 | 0.01 | A0A1Q3S7A3 | *Flavobacterium sp.* | Endonuclease MutS2 (EC 3.1.-.-) |

**Table S7.** Functional overview of M5A community (top 500 rank only).

| Rank | Count | Abundance | ID | Organism | Protein Names |
|---|---|---|---|---|---|
| 1 | 4376 | 0.14 | F4BUN6 | *Methanosaeta concilii* | Transposase, IS116/IS110/IS902 family |
| 2 | 3604 | 0.12 | F4BZD0 | *Methanosaeta concilii* | Transposase, putative |
| 3 | 2697 | 0.09 | F4BY05 | *Methanosaeta concilii* | Transposase, IS116/IS110/IS902 family |
| 4 | 1997 | 0.07 | F4BY15 | *Methanosaeta concilii* | Transposase, IS116/IS110/IS902 family |
| 5 | 1995 | 0.07 | I9R3U4 | *Bacteroides cellulosilyticus* | Uncharacterized protein |
| 6 | 1977 | 0.07 | F4BSV9 | *Methanosaeta concilii* | Transposase, IS4 family |
| 7 | 1641 | 0.05 | F4BZ63 | *Methanosaeta concilii* | Transposase, IS4 family |
| 8 | 1506 | 0.05 | F4BSV1 | *Methanosaeta concilii* | Transposase, IS4 family, putative |
| 9 | 1289 | 0.04 | F4BV05 | *Methanosaeta concilii* | Transposase, Rhodopirellula family protein |
| 10 | 1247 | 0.04 | F4BZW8 | *Methanosaeta concilii* | Transposase, IS605 OrfB family |
| 11 | 1232 | 0.04 | F4BX44 | *Methanosaeta concilii* | Transposase, IS605 OrfB family |
| 12 | 1164 | 0.04 | N8ZVC9 | *Acinetobacter gerneri* | Uncharacterized protein |
| 13 | 1132 | 0.04 | F3ZQL8 | *Bacteroides coprosuis* | Transposase IS4 family protein |
| 14 | 1125 | 0.04 | F4BYN0 | *Methanosaeta concilii* | Transposase, IS605 OrfB family |
| 15 | 1124 | 0.04 | N8ZNX3 | *Acinetobacter gerneri* | Uncharacterized protein |
| 16 | 1122 | 0.04 | F4BXC1 | *Methanosaeta concilii* | Transposase, IS4 |
| 17 | 981 | 0.03 | F4BV83 | *Methanosaeta concilii* | Uncharacterized protein |
| 18 | 916 | 0.03 | F4BUD0 | *Methanosaeta concilii* | Conserved domain transposase IS605 family |
| 19 | 872 | 0.03 | F4BVK0 | *Methanosaeta concilii* | Transposase, IS116/IS110/IS902 family |
| 20 | 803 | 0.03 | | | |
| 21 | 796 | 0.03 | F4C073 | *Methanosaeta concilii* | Transposase, IS605 OrfB family |
| 22 | 780 | 0.03 | A0A0R3QEE4 | *Brugia timori* | Uncharacterized protein |
| 23 | 738 | 0.02 | F4BZ72 | *Methanosaeta concilii* | Uncharacterized protein |
| 24 | 729 | 0.02 | F4BTW8 | *Methanosaeta concilii* | Transposase IS3 family protein, putative integrase |
| 25 | 716 | 0.02 | F4BUB6 | *Methanosaeta concilii* | Transposase, IS605 OrfB family |
| 26 | 661 | 0.02 | F4BUL3 | *Methanosaeta concilii* | Transposase |
| 27 | 624 | 0.02 | H8MU17 | *Corallococcus coralloides* | Long-chain-fatty-acid--CoA ligase |
| 28 | 608 | 0.02 | | | |
| 29 | 607 | 0.02 | A0A077XLW5 | *Sphinx1.76-related DNA* | Replication protein |
| 30 | 592 | 0.02 | F4C0V0 | *Methanosaeta concilii* | Uncharacterized protein |
| 31 | 586 | 0.02 | F4BTV1 | *Methanosaeta concilii* | Glycosyltransferase family 36 protein |
| 32 | 578 | 0.02 | A0A1Q2CUJ4 | *Tessaracoccus flavescens* | ISL3 family transposase |
| 33 | 570 | 0.02 | F4BZ82 | *Methanosaeta concilii* | Cna B domain protein |

| | | | | | |
|---|---|---|---|---|---|
| 34 | 555 | 0.02 | F4BV12 | *Methanosaeta concilii* | Transposase, Mutator family |
| 35 | 549 | 0.02 | F4BTX0 | *Methanosaeta concilii* | Conserved domain protein |
| 36 | 536 | 0.02 | A0A0T5YH86 | *Acinetobacter baumannii* | Mobilization protein A |
| 37 | 534 | 0.02 | A0A1Q3QH52 | *Bacteroidia bacterium* | Uncharacterized protein |
| 38 | 521 | 0.02 | F4BW50 | *Methanosaeta concilii* | Transposase, IS605 OrfB family domain protein |
| 39 | 520 | 0.02 | F4BT92 | *Methanosaeta concilii* | Transposase, Mutator family |
| 40 | 513 | 0.02 | A0A077XNB6 | *Sphinx1.76-related DNA* | Replication protein |
| 41 | 505 | 0.02 | A0A1Q3QCL0 | *Bacteroidia bacterium* | Uncharacterized protein |
| 42 | 504 | 0.02 | G9FB24 | *uncultured bacterium* | IS21 family transposition protein |
| 43 | 501 | 0.02 | H8MYX7 | *Corallococcus coralloides* | Non-ribosomal peptide synthetase/polyketide synthase |
| 44 | 487 | 0.02 | | | |
| 45 | 483 | 0.02 | A0A1Q3QJR2 | *Bacteroidia bacterium* | Cell surface protein SprA |
| 46 | 482 | 0.02 | | | |
| 47 | 481 | 0.02 | | | |
| 48 | 478 | 0.02 | F4BW44 | *Methanosaeta concilii* | Transposase, IS605 OrfB family |
| 49 | 469 | 0.02 | A0A178FWP5 | *Acinetobacter sp. SFC* | Transposase |
| 50 | 466 | 0.02 | F4BVI0 | *Methanosaeta concilii* | Methyltransferase/glycosyl transferase fusion protein |
| 51 | 437 | 0.01 | F4BXD7 | *Methanosaeta concilii* | Transposase, IS4 (Transposase, IS4 family) |
| 52 | 426 | 0.01 | A0A1M7BMK5 | *Fibrobacter sp. UWEL* | Transposase InsO and inactivated derivatives |
| 53 | 423 | 0.01 | | | |
| 54 | 418 | 0.01 | | | |
| 55 | 417 | 0.01 | N8QHE7 | *Acinetobacter bohemicus* | Uncharacterized protein |
| 56 | 414 | 0.01 | A0A1Q3QC75 | *Bacteroidia bacterium* | Uncharacterized protein |
| 57 | 409 | 0.01 | | | |
| 58 | 408 | 0.01 | | | |
| 59 | 405 | 0.01 | U6RUP9 | *Bacteroides sp. HPS0048* | Uncharacterized protein |
| 60 | 405 | 0.01 | | | |
| 61 | 402 | 0.01 | | | |
| 62 | 401 | 0.01 | | | |
| 63 | 399 | 0.01 | F4BUA5 | *Methanosaeta concilii* | Transposase, IS605 OrfB family |
| 64 | 393 | 0.01 | | | |
| 65 | 384 | 0.01 | A0A1Q3QBX8 | *Bacteroidia bacterium* | Uncharacterized protein |
| 66 | 383 | 0.01 | F4BUD1 | *Methanosaeta concilii* | Transposase, putative |
| 67 | 380 | 0.01 | | | |
| 68 | 379 | 0.01 | A0A1Q3QGY6 | *Bacteroidia bacterium* | Hybrid sensor histidine kinase/response regulator |
| 69 | 374 | 0.01 | I2F7Y4 | *Mesotoga prima MesG1.Ag.4.2* | Retron-type reverse transcriptase |

| | | | | | |
|---|---|---|---|---|---|
| 70 | 370 | 0.01 | T5L8A5 | *Microbacterium maritypicum* | Uncharacterized protein |
| 71 | 361 | 0.01 | H8MN58 | *Corallococcus coralloides* | Long-chain-fatty-acid--CoA ligase |
| 72 | 356 | 0.01 | A0A1Q3QI02 | *Bacteroidia bacterium* | Uncharacterized protein |
| 73 | 355 | 0.01 | F4BXC2 | *Methanosaeta concilii* | Transposase IS200-like protein (Transposase, IS200-like, putative MITE) |
| 74 | 355 | 0.01 | F4BY39 | *Methanosaeta concilii* | Transposase, IS605 OrfB family |
| 75 | 351 | 0.01 | A0A0C5RZ24 | *Pseudomonas putida* | Adhesin |
| 76 | 351 | 0.01 | | | |
| 77 | 348 | 0.01 | F3ZQK5 | *Bacteroides coprosuis* | Transposase IS4 family protein |
| 78 | 348 | 0.01 | F4BYV3 | *Methanosaeta concilii* | Riboflavin biosynthesis protein RibD C-domain protein |
| 79 | 346 | 0.01 | | | |
| 80 | 345 | 0.01 | G9FB23 | *uncultured bacterium* | IS21 family transposition protein |
| 81 | 345 | 0.01 | | | |
| 82 | 342 | 0.01 | F4BZ39 | *Methanosaeta concilii* | Transposase, IS116/IS110/IS902 family |
| 83 | 340 | 0.01 | A0A1M6CDY9 | *Tessaracoccus bendigoensis* | Cd2+/Zn2+-exporting ATPase |
| 84 | 336 | 0.01 | F4BXE4 | *Methanosaeta concilii* | Conserved domain, transposase IS3 family protein, putative integrase |
| 85 | 336 | 0.01 | | | |
| 86 | 332 | 0.01 | | | |
| 87 | 331 | 0.01 | N8P3X5 | *Acinetobacter bohemicus* | Uncharacterized protein |
| 88 | 330 | 0.01 | | | |
| 89 | 329 | 0.01 | F4BXF6 | *Methanosaeta concilii* | Uncharacterized protein |
| 90 | 327 | 0.01 | A0A1Q3QHS9 | *Bacteroidia bacterium* | Uncharacterized protein |
| 91 | 322 | 0.01 | H8MSC6 | *Corallococcus coralloides* | Non-ribosomal peptide synthetase |
| 92 | 321 | 0.01 | | | |
| 93 | 320 | 0.01 | | | |
| 94 | 314 | 0.01 | F4BVL8 | *Methanosaeta concilii* | Uncharacterized protein |
| 95 | 313 | 0.01 | | | |
| 96 | 311 | 0.01 | F4BZ62 | *Methanosaeta concilii* | Conserved domain protein |
| 97 | 307 | 0.01 | A0A1Q3QCN8 | *Bacteroidia bacterium* | Uncharacterized protein |
| 98 | 307 | 0.01 | A0A1Q3QIV6 | *Bacteroidia bacterium* | Uncharacterized protein |
| 99 | 307 | 0.01 | N8YFI4 | *Acinetobacter gerneri* | Uncharacterized protein |
| 100 | 305 | 0.01 | | | |
| 101 | 304 | 0.01 | A0A1Q3QBE7 | *Bacteroidia bacterium* | Uncharacterized protein |
| 102 | 302 | 0.01 | F4BVN9 | *Methanosaeta concilii* | Conserved repeat domain protein |
| 103 | 301 | 0.01 | N9GC16 | *Acinetobacter lwoffii* | Uncharacterized protein |
| 104 | 300 | 0.01 | F4BTW7 | *Methanosaeta concilii* | Conserved domain protein |
| 105 | 297 | 0.01 | A0A1Q3QDS6 | *Bacteroidia bacterium* | Uncharacterized protein |

| | | | | | |
|---|---|---|---|---|---|
| 106 | 297 | 0.01 | F4BUE8 | *Methanosaeta concilii* | Transposase, predicted |
| 107 | 291 | 0.01 | A0A1Q3QHX6 | *Bacteroidia bacterium* | Multidrug transporter AcrB |
| 108 | 290 | 0.01 | | | |
| 109 | 289 | 0.01 | | | |
| 110 | 288 | 0.01 | A0A1Q3QI54 | *Bacteroidia bacterium* | SusC/RagA family TonB-linked outer membrane protein |
| 111 | 287 | 0.01 | F4BXK5 | *Methanosaeta concilii* | Transposase |
| 112 | 284 | 0.01 | F4BX50 | *Methanosaeta concilii* | Type III restriction enzyme, res subunit |
| 113 | 284 | 0.01 | | | |
| 114 | 283 | 0.01 | D2E402 | *Kernia pachypleura* | Ribosomal protein 3 |
| 115 | 283 | 0.01 | | | |
| 116 | 282 | 0.01 | | | |
| 117 | 281 | 0.01 | A0A0W8FAI0 | *hydrocarbon metagenome* | Uncharacterized protein |
| 118 | 281 | 0.01 | A0A1Q3QIY4 | *Bacteroidia bacterium* | SusC/RagA family protein |
| 119 | 279 | 0.01 | A0A1Q3QGJ2 | *Bacteroidia bacterium* | Uncharacterized protein |
| 120 | 278 | 0.01 | | | |
| 121 | 277 | 0.01 | | | |
| 122 | 276 | 0.01 | N8YF58 | *Acinetobacter gerneri* | Uncharacterized protein |
| 123 | 276 | 0.01 | | | |
| 124 | 276 | 0.01 | | | |
| 125 | 275 | 0.01 | A0A1Q3QB47 | *Bacteroidia bacterium* | Deoxyribonuclease HsdR |
| 126 | 274 | 0.01 | F4BU67 | *Methanosaeta concilii* | Multi-sensor hybrid histidine kinase |
| 127 | 272 | 0.01 | F4BT93 | *Methanosaeta concilii* | Uncharacterized protein |
| 128 | 272 | 0.01 | | | |
| 129 | 271 | 0.01 | | | |
| 130 | 271 | 0.01 | | | |
| 131 | 269 | 0.01 | | | |
| 132 | 268 | 0.01 | | | |
| 133 | 267 | 0.01 | | | |
| 134 | 267 | 0.01 | | | |
| 135 | 266 | 0.01 | F4BXN8 | *Methanosaeta concilii* | CobN/magnesium chelatase domain protein |
| 136 | 266 | 0.01 | | | |
| 137 | 265 | 0.01 | | | |
| 138 | 263 | 0.01 | A0A1Q3QF20 | *Bacteroidia bacterium* | SusC/RagA family protein |
| 139 | 263 | 0.01 | A0A1Q3QI16 | *Bacteroidia bacterium* | SusC/RagA family TonB-linked outer membrane protein |
| 140 | 262 | 0.01 | A0A1Q3QH94 | *Bacteroidia bacterium* | DNA-directed DNA polymerase (EC 2.7.7.7) |
| 141 | 262 | 0.01 | | | |

| | | | | | |
|---|---|---|---|---|---|
| 142 | 262 | 0.01 | | | |
| 143 | 262 | 0.01 | | | |
| 144 | 261 | 0.01 | | | |
| 145 | 260 | 0.01 | F4BXH3 | *Methanosaeta concilii* | Cobaltochelatase, CobN subunit |
| 146 | 259 | 0.01 | A0A1Q3QIK1 | *Bacteroidia bacterium* | Phosphoribosylformylglycinamidine synthase (FGAM synthase) (FGAMS) (EC 6.3.5.3) (Formylglycinamide ribonucleotide amidotransferase) (FGAR amidotransferase) (FGAR-AT) |
| 147 | 257 | 0.01 | A0A1Q3Q9Q2 | *Bacteroidia bacterium* | SusC/RagA family protein |
| 148 | 257 | 0.01 | F4BT87 | *Methanosaeta concilii* | TPR-repeat protein |
| 149 | 257 | 0.01 | F4BXG6 | *Methanosaeta concilii* | Cobaltochelatase, CobN subunit (EC 6.6.1.2) |
| 150 | 256 | 0.01 | | | |
| 151 | 253 | 0.01 | A0A1Q3QAN2 | *Bacteroidia bacterium* | SusC/RagA family TonB-linked outer membrane protein |
| 152 | 251 | 0.01 | A0A1Q3Q9V3 | *Bacteroidia bacterium* | Peptidase S9 |
| 153 | 249 | 0.01 | A0A1M6CCR6 | *Tessaracoccus bendigoensis* | Uncharacterized protein |
| 154 | 249 | 0.01 | A0A1Q3QA74 | *Bacteroidia bacterium* | TonB-dependent receptor |
| 155 | 248 | 0.01 | H8MS99 | *Corallococcus coralloides* | Adventurous gliding motility protein AgmK |
| 156 | 247 | 0.01 | A0A1Q3QBC6 | *Bacteroidia bacterium* | Uncharacterized protein |
| 157 | 247 | 0.01 | A0A1Q3QE63 | *Bacteroidia bacterium* | Uncharacterized protein |
| 158 | 245 | 0.01 | A0A077XLZ7 | *Sphinx1.76-related DNA* | Replication protein |
| 159 | 245 | 0.01 | A0A1Q3QF86 | *Bacteroidia bacterium* | Multidrug transporter AcrB |
| 160 | 244 | 0.01 | F4C043 | *Methanosaeta concilii* | Membrane protein, putative |
| 161 | 243 | 0.01 | A0A1G4G9S9 | *Petrimonas mucosa* | TonB-dependent receptor SusC |
| 162 | 243 | 0.01 | A0A1Q3QGM8 | *Bacteroidia bacterium* | SusC/RagA family TonB-linked outer membrane protein |
| 163 | 243 | 0.01 | F4BV15 | *Methanosaeta concilii* | Transposase, IS605 OrfB family |
| 164 | 243 | 0.01 | | | |
| 165 | 242 | 0.01 | A0A1Q3QAH7 | *Bacteroidia bacterium* | DNA polymerase I (EC 2.7.7.7) |
| 166 | 242 | 0.01 | A0A1Q3QDC9 | *Bacteroidia bacterium* | SusC/RagA family TonB-linked outer membrane protein |
| 167 | 242 | 0.01 | | | |
| 168 | 241 | 0.01 | F4BWZ9 | *Methanosaeta concilii* | Two-component hybrid sensor and regulator |
| 169 | 240 | 0.01 | A0A1Q3QBL0 | *Bacteroidia bacterium* | Acriflavine resistance protein B |
| 170 | 240 | 0.01 | A0A1Q3QD12 | *Bacteroidia bacterium* | SusC/RagA family TonB-linked outer membrane protein |
| 171 | 240 | 0.01 | | | |
| 172 | 239 | 0.01 | A0A1Q3QFV8 | *Bacteroidia bacterium* | DNA helicase (EC 3.6.4.12) |
| 173 | 239 | 0.01 | F4BUA4 | *Methanosaeta concilii* | Transposase, IS605 OrfB family |
| 174 | 238 | 0.01 | F4BTN1 | *Methanosaeta concilii* | Two-component hybrid sensor and regulator |
| 175 | 238 | 0.01 | | | |
| 176 | 238 | 0.01 | | | |

| | | | | | |
|---|---|---|---|---|---|
| 177 | 237 | 0.01 | A0A1R3TE89 | *Proteiniphilum saccharofermentans* | Uncharacterized protein |
| 178 | 237 | 0.01 | F4C0L1 | *Methanosaeta concilii* | TPR-repeat protein |
| 179 | 236 | 0.01 | | | |
| 180 | 235 | 0.01 | A0A0N1L384 | *Novosphingobium sp.* | Glutamate dehydrogenase |
| 181 | 235 | 0.01 | A0A1Q3QG05 | *Bacteroidia bacterium* | Multifunctional fusion protein [Includes: Protein translocase subunit SecD; Protein-export membrane protein SecF] |
| 182 | 234 | 0.01 | A0A1M6CDN2 | *Tessaracoccus bendigoensis* | Multicopper oxidase with three cupredoxin domains (Includes cell division protein FtsP and spore coat protein CotA) |
| 183 | 234 | 0.01 | A0A1Q3QCP4 | *Bacteroidia bacterium* | SusC/RagA family TonB-linked outer membrane protein |
| 184 | 234 | 0.01 | | | |
| 185 | 233 | 0.01 | F4BY67 | *Methanosaeta concilii* | Uncharacterized protein |
| 186 | 232 | 0.01 | | | |
| 187 | 231 | 0.01 | F4BXB3 | *Methanosaeta concilii* | Transposase, IS605 family protein, putative MITE |
| 188 | 231 | 0.01 | | | |
| 189 | 231 | 0.01 | | | |
| 190 | 230 | 0.01 | F4BWV9 | *Methanosaeta concilii* | S-layer-related duplication domain protein |
| 191 | 230 | 0.01 | | | |
| 192 | 228 | 0.01 | A0A1Q3QGP0 | *Bacteroidia bacterium* | Glycosyl hydrolase family 35 |
| 193 | 228 | 0.01 | | | |
| 194 | 228 | 0.01 | | | |
| 195 | 226 | 0.01 | F4BVL9 | *Methanosaeta concilii* | Uncharacterized protein |
| 196 | 226 | 0.01 | | | |
| 197 | 225 | 0.01 | | | |
| 198 | 225 | 0.01 | | | |
| 199 | 224 | 0.01 | | | |
| 200 | 224 | 0.01 | | | |
| 201 | 223 | 0.01 | A0A1Q3QI74 | *Bacteroidia bacterium* | Peptidase M16 |
| 202 | 223 | 0.01 | A0A1Q3QIQ0 | *Bacteroidia bacterium* | Uncharacterized protein |
| 203 | 223 | 0.01 | | | |
| 204 | 222 | 0.01 | A0A1Q3QBF6 | *Bacteroidia bacterium* | SusC/RagA family protein |
| 205 | 222 | 0.01 | A0A1Q3QDK9 | *Bacteroidia bacterium* | Uncharacterized protein |
| 206 | 222 | 0.01 | A0A1Q3QJE6 | *Bacteroidia bacterium* | Cation transporter |
| 207 | 222 | 0.01 | | | |
| 208 | 222 | 0.01 | | | |
| 209 | 221 | 0.01 | A0A1Q3QD11 | *Bacteroidia bacterium* | SusC/RagA family protein |

| | | | | | |
|---|---|---|---|---|---|
| 210 | 220 | 0.01 | A0A1Q3QGR5 | *Bacteroidia bacterium* | SusC/RagA family TonB-linked outer membrane protein |
| 211 | 220 | 0.01 | F1Z681 | *Novosphingobium nitrogenifigens* | Type I site-specific deoxyribonuclease, HsdR family |
| 212 | 220 | 0.01 | F4BSZ2 | *Methanosaeta concilii* | Oligosaccharyl transferase STT3 subunit |
| 213 | 220 | 0.01 | F4BVN7 | *Methanosaeta concilii* | Pentapeptide repeat protein |
| 214 | 220 | 0.01 | F4C050 | *Methanosaeta concilii* | Asparagine synthase (Glutamine-hydrolyzing) (EC 6.3.5.4) |
| 215 | 220 | 0.01 | | | |
| 216 | 219 | 0.01 | A0A1Q3Q9V2 | *Bacteroidia bacterium* | Phosphate transporter |
| 217 | 219 | 0.01 | A0A1Q3QFQ4 | *Bacteroidia bacterium* | Uncharacterized protein |
| 218 | 219 | 0.01 | F4BVI9 | *Methanosaeta concilii* | IS3/IS66 family element, Orf1 protein |
| 219 | 219 | 0.01 | | | |
| 220 | 218 | 0.01 | A0A0W8FC74 | *hydrocarbon metagenome* | Sensor histidine kinase |
| 221 | 218 | 0.01 | F1ZDD0 | *Novosphingobium nitrogenifigens* | Penicillin amidase family protein |
| 222 | 218 | 0.01 | F4BZ35 | *Methanosaeta concilii* | Uncharacterized protein |
| 223 | 218 | 0.01 | | | |
| 224 | 218 | 0.01 | | | |
| 225 | 218 | 0.01 | | | |
| 226 | 218 | 0.01 | | | |
| 227 | 217 | 0.01 | A0A1Q3QAJ6 | *Bacteroidia bacterium* | Uncharacterized protein |
| 228 | 217 | 0.01 | F4BSW7 | *Methanosaeta concilii* | Multi-sensor hybrid histidine kinase |
| 229 | 217 | 0.01 | F4BW20 | *Methanosaeta concilii* | Glycosyl transferase group 2/group 1 fusion protein |
| 230 | 217 | 0.01 | | | |
| 231 | 216 | 0.01 | A0A1Q3QBY0 | *Bacteroidia bacterium* | Phosphoenolpyruvate synthase |
| 232 | 216 | 0.01 | A0A1Q3QCR5 | *Bacteroidia bacterium* | Isoleucine--tRNA ligase (EC 6.1.1.5) (Isoleucyl-tRNA synthetase) (IleRS) |
| 233 | 216 | 0.01 | F4BXZ6 | *Methanosaeta concilii* | S-layer-related duplication domain protein |
| 234 | 216 | 0.01 | | | |
| 235 | 216 | 0.01 | | | |
| 236 | 216 | 0.01 | | | |
| 237 | 215 | 0.01 | A0A0F0LYK2 | *Microbacterium ginsengisoli* | Putative cadmium-transporting ATPase (EC 3.6.3.3) |
| 238 | 215 | 0.01 | A0A1Q3QJ62 | *Bacteroidia bacterium* | Uncharacterized protein (Fragment) |
| 239 | 215 | 0.01 | | | |
| 240 | 214 | 0.01 | A0A1Q3QAN0 | *Bacteroidia bacterium* | Uncharacterized protein |
| 241 | 214 | 0.01 | | | |
| 242 | 214 | 0.01 | | | |
| 243 | 214 | 0.01 | | | |
| 244 | 213 | 0.01 | A0A1Q3Q9W7 | *Bacteroidia bacterium* | Uncharacterized protein |
| 245 | 213 | 0.01 | A0A1Q3QED9 | *Bacteroidia bacterium* | Uncharacterized protein |

| | | | | | |
|---|---|---|---|---|---|
| 246 | 213 | 0.01 | A0A1Q3QGE6 | *Bacteroidia bacterium* | Uncharacterized protein |
| 247 | 213 | 0.01 | A0A1Q3QGG2 | *Bacteroidia bacterium* | SusC/RagA family protein |
| 248 | 213 | 0.01 | | | |
| 249 | 212 | 0.01 | A0A1Q3QDG3 | *Bacteroidia bacterium* | SusC/RagA family TonB-linked outer membrane protein |
| 250 | 212 | 0.01 | F0M3U0 | *Pseudarthrobacter phenanthrenivorans (strain DSM 18606 / JCM 16027 / LMG 23796 / Sphe3) (Arthrobacter phenanthrenivorans)* | Phospholipase/Carboxylesterase |
| 251 | 212 | 0.01 | | | |
| 252 | 211 | 0.01 | A0A1Q3QC05 | *Bacteroidia bacterium* | Uncharacterized protein |
| 253 | 211 | 0.01 | A0A1Q3QES6 | *Bacteroidia bacterium* | Uncharacterized protein |
| 254 | 211 | 0.01 | F4BXL6 | *Methanosaeta concilii* | Conserved domain protein |
| 255 | 211 | 0.01 | | | |
| 256 | 211 | 0.01 | | | |
| 257 | 211 | 0.01 | | | |
| 258 | 210 | 0.01 | A0A0Q7S8L4 | *Pelomonas sp.* | Transposase |
| 259 | 210 | 0.01 | A0A1Q3QCH6 | *Bacteroidia bacterium* | Nitrate reductase subunit alpha |
| 260 | 210 | 0.01 | A0A1Q3QFU6 | *Bacteroidia bacterium* | TonB-dependent receptor |
| 261 | 210 | 0.01 | | | |
| 262 | 210 | 0.01 | | | |
| 263 | 209 | 0.01 | | | |
| 264 | 209 | 0.01 | | | |
| 265 | 208 | 0.01 | A0A0N1ANW5 | *Novosphingobium sp.* | Cellulose synthase |
| 266 | 208 | 0.01 | A0A1Q3QAN4 | *Bacteroidia bacterium* | Uncharacterized protein |
| 267 | 208 | 0.01 | F4BXF7 | *Methanosaeta concilii* | Transposase, putative integrase |
| 268 | 208 | 0.01 | | | |
| 269 | 207 | 0.01 | A0A1Q3QIQ9 | *Bacteroidia bacterium* | Uncharacterized protein |
| 270 | 207 | 0.01 | F4BYM4 | *Methanosaeta concilii* | Chromosome partition protein Smc |
| 271 | 207 | 0.01 | | | |
| 272 | 207 | 0.01 | | | |
| 273 | 206 | 0.01 | A0A1Q3QIR5 | *Bacteroidia bacterium* | Alpha-glucan phosphorylase |
| 274 | 206 | 0.01 | A0A1Q3QJ29 | *Bacteroidia bacterium* | DNA mismatch repair protein MutS |
| 275 | 205 | 0.01 | H8MFD7 | *Corallococcus coralloides* | Long-chain-fatty-acid--CoA ligase |
| 276 | 205 | 0.01 | L7LGQ0 | *Gordonia sihwensis* | Putative non-ribosomal peptide synthetase |
| 277 | 205 | 0.01 | | | |

| | | | | | |
|---|---|---|---|---|---|
| 278 | 205 | 0.01 | | | |
| 279 | 204 | 0.01 | | | |
| 280 | 204 | 0.01 | | | |
| 281 | 203 | 0.01 | A0A1G4GAX8 | *Petrimonas mucosa* | Uncharacterized protein |
| 282 | 203 | 0.01 | A0A1Q3QDL1 | *Bacteroidia bacterium* | Methionine synthase |
| 283 | 203 | 0.01 | P11408 | *Moraxella sp.* | Modification methylase MspI (M.MspI) (EC 2.1.1.37) (Cytosine-specific methyltransferase MspI) |
| 284 | 203 | 0.01 | | | |
| 285 | 202 | 0.01 | | | |
| 286 | 202 | 0.01 | | | |
| 287 | 201 | 0.01 | F4BY50 | *Methanosaeta concilii* | ATP-dependent RNA helicase, putative |
| 288 | 201 | 0.01 | | | |
| 289 | 201 | 0.01 | | | |
| 290 | 201 | 0.01 | | | |
| 291 | 200 | 0.01 | F4BWJ6 | *Methanosaeta concilii* | Peptidase families S8 and S53 protein |
| 292 | 200 | 0.01 | F4BWT2 | *Methanosaeta concilii* | DNA polymerase (EC 2.7.7.7) |
| 293 | 200 | 0.01 | F5XPC2 | *Microlunatus phosphovorus* | Uncharacterized protein |
| 294 | 200 | 0.01 | | | |
| 295 | 199 | 0.01 | A0A1Q3QB44 | *Bacteroidia bacterium* | Uncharacterized protein |
| 296 | 199 | 0.01 | A0A1Q3QBJ9 | *Bacteroidia bacterium* | Uncharacterized protein |
| 297 | 199 | 0.01 | A0A1R3T512 | *Proteiniphilum saccharofermentans* | TonB-linked outer membrane protein, SusC/RagA family |
| 298 | 199 | 0.01 | D8L7S1 | *Paramecium caudatum* | NADH-ubiquinone oxidoreductase chain 5 (EC 1.6.5.3) |
| 299 | 198 | 0.01 | A0A0R3R918 | *Brugia timori* | Uncharacterized protein |
| 300 | 198 | 0.01 | F4BZ75 | *Methanosaeta concilii* | Carbamoyl-phosphate synthase large chain (EC 6.3.5.5) (Carbamoyl-phosphate synthetase ammonia chain) |
| 301 | 198 | 0.01 | S2WV93 | *Delftia acidovorans* | Uncharacterized protein |
| 302 | 198 | 0.01 | | | |
| 303 | 197 | 0.01 | | | |
| 304 | 197 | 0.01 | | | |
| 305 | 196 | 0.01 | F4BU15 | *Methanosaeta concilii* | DNA mismatch repair protein MutS |
| 306 | 196 | 0.01 | F5XPA1 | *Microlunatus phosphovorus* | Uncharacterized protein |
| 307 | 196 | 0.01 | | | |
| 308 | 195 | 0.01 | A0A1Q3Q9V6 | *Bacteroidia bacterium* | Multidrug transporter AcrB |
| 309 | 195 | 0.01 | A0A1Q3QF64 | *Bacteroidia bacterium* | Alpha-rhamnosidase |
| 310 | 195 | 0.01 | U6RCL0 | *Bacteroides sp.* | Uncharacterized protein |
| 311 | 195 | 0.01 | | | |

| | | | | | |
|---|---|---|---|---|---|
| 312 | 194 | 0.01 | A0A1Q3QED5 | *Bacteroidia bacterium* | Multidrug transporter AcrB |
| 313 | 194 | 0.01 | A0A1Q3QF91 | *Bacteroidia bacterium* | Alpha-rhamnosidase |
| 314 | 194 | 0.01 | A0A1Q3QJP8 | *Bacteroidia bacterium* | Cytochrome C biogenesis protein |
| 315 | 194 | 0.01 | | | |
| 316 | 193 | 0.01 | A0A1Q3QGG1 | *Bacteroidia bacterium* | Uncharacterized protein |
| 317 | 193 | 0.01 | A0A1Q3QIS8 | *Bacteroidia bacterium* | NADH dehydrogenase |
| 318 | 193 | 0.01 | F4BVD8 | *Methanosaeta concilii* | Cation transporting P-type ATPase |
| 319 | 193 | 0.01 | F4BVJ1 | *Methanosaeta concilii* | Beta-propeller repeat protein |
| 320 | 193 | 0.01 | | | |
| 321 | 192 | 0.01 | A0A1Q3QFG4 | *Bacteroidia bacterium* | Alpha-galactosidase |
| 322 | 192 | 0.01 | A0A1Q3QFT2 | *Bacteroidia bacterium* | Uncharacterized protein |
| 323 | 192 | 0.01 | A0A1Q3QI00 | *Bacteroidia bacterium* | Tricorn protease homolog (EC 3.4.21.-) |
| 324 | 192 | 0.01 | | | |
| 325 | 192 | 0.01 | | | |
| 326 | 192 | 0.01 | | | |
| 327 | 191 | 0.01 | F4BVJ0 | *Methanosaeta concilii* | Uncharacterized protein |
| 328 | 191 | 0.01 | H8MSC8 | *Corallococcus coralloides* | Tyrocidine synthase |
| 329 | 191 | 0.01 | | | |
| 330 | 191 | 0.01 | | | |
| 331 | 190 | 0.01 | A0A154DNE7 | *Acinetobacter baumannii* | mRNA endoribonuclease LS (EC 3.1.-.-) |
| 332 | 190 | 0.01 | A0A1Q3QFK3 | *Bacteroidia bacterium* | Alpha-rhamnosidase |
| 333 | 190 | 0.01 | A0A1Q3QHQ6 | *Bacteroidia bacterium* | SusC/RagA family protein |
| 334 | 190 | 0.01 | A0A1Q3QIM7 | *Bacteroidia bacterium* | Uncharacterized protein |
| 335 | 190 | 0.01 | | | |
| 336 | 189 | 0.01 | A0A0E3T6K6 | *Gordonia phage Gmala1* | Tape measure protein |
| 337 | 189 | 0.01 | A0A1Q3Q9V8 | *Bacteroidia bacterium* | TonB-dependent receptor |
| 338 | 189 | 0.01 | | | |
| 339 | 189 | 0.01 | | | |
| 340 | 189 | 0.01 | | | |
| 341 | 189 | 0.01 | | | |
| 342 | 189 | 0.01 | | | |
| 343 | 188 | 0.01 | A0A1Q3QA64 | *Bacteroidia bacterium* | Uncharacterized protein |
| 344 | 188 | 0.01 | A0A1Q3QD21 | *Bacteroidia bacterium* | Beta-galactosidase (EC 3.2.1.23) (Lactase) |
| 345 | 188 | 0.01 | A0A1Q3QHZ1 | *Bacteroidia bacterium* | Peptidase M10A/M12B |
| 346 | 188 | 0.01 | A0A1Q3QI69 | *Bacteroidia bacterium* | Uncharacterized protein |
| 347 | 188 | 0.01 | A0A1Q3QIW6 | *Bacteroidia bacterium* | Uncharacterized protein |

| | | | | | |
|---|---|---|---|---|---|
| 348 | 188 | 0.01 | F4BZ38 | *Methanosaeta concilii* | Transposase, IS116/IS110/IS902 family |
| 349 | 188 | 0.01 | | | |
| 350 | 187 | 0.01 | A0A1Q3QD59 | *Bacteroidia bacterium* | Translation initiation factor IF-2 |
| 351 | 187 | 0.01 | F4BXQ5 | *Methanosaeta concilii* | Two-component hybrid sensor and regulator |
| 352 | 187 | 0.01 | F4BY09 | *Methanosaeta concilii* | DNA polymerase II large subunit (Pol II) (EC 2.7.7.7) |
| 353 | 187 | 0.01 | | | |
| 354 | 187 | 0.01 | | | |
| 355 | 187 | 0.01 | | | |
| 356 | 187 | 0.01 | | | |
| 357 | 187 | 0.01 | | | |
| 358 | 186 | 0.01 | A0A151BT14 | *Branchiibius sp.* | Mercuric reductase |
| 359 | 186 | 0.01 | A0A1Q3QCA7 | *Bacteroidia bacterium* | Glycosyl hydrolase family 31 |
| 360 | 186 | 0.01 | A0A1Q3QGC9 | *Bacteroidia bacterium* | Peptidase M16 |
| 361 | 186 | 0.01 | F4BU80 | *Methanosaeta concilii* | Transposase, IS605 OrfB family |
| 362 | 186 | 0.01 | | | |
| 363 | 185 | 0.01 | A0A1M3FLT2 | *Bacteroidia bacterium* | Uncharacterized protein |
| 364 | 185 | 0.01 | A0A1M6WBE7 | *Fibrobacter sp. UWOS* | Transposase (Or an inactivated derivative) |
| 365 | 185 | 0.01 | A0A1Q3QA25 | *Bacteroidia bacterium* | UvrABC system protein C (Protein UvrC) (Excinuclease ABC subunit C) |
| 366 | 185 | 0.01 | A0A1Q3QD60 | *Bacteroidia bacterium* | Alanine racemase (EC 5.1.1.1) |
| 367 | 185 | 0.01 | A0A1Q3QFA7 | *Bacteroidia bacterium* | Glycoside hydrolase family 2 |
| 368 | 185 | 0.01 | A0A1Q3QG11 | *Bacteroidia bacterium* | Capsular biosynthesis protein |
| 369 | 185 | 0.01 | A1W5C2 | *Acidovorax sp.* | IstB domain protein ATP-binding protein |
| 370 | 184 | 0.01 | A0A1G4G461 | *Petrimonas mucosa* | Outer membrane protein assembly factor BamA |
| 371 | 184 | 0.01 | A0A1Q3QAU4 | *Bacteroidia bacterium* | Uncharacterized protein |
| 372 | 184 | 0.01 | A0A1Q3QB09 | *Bacteroidia bacterium* | Glycoside hydrolase |
| 373 | 184 | 0.01 | A0A1Q3QGV1 | *Bacteroidia bacterium* | Uncharacterized protein |
| 374 | 184 | 0.01 | A1W6Q0 | *Acidovorax sp.* | Integrase, catalytic region |
| 375 | 184 | 0.01 | | | |
| 376 | 184 | 0.01 | | | |
| 377 | 184 | 0.01 | | | |
| 378 | 184 | 0.01 | | | |
| 379 | 183 | 0.01 | A0A0N0JFZ2 | *Novosphingobium sp.* | Glutamate synthase |
| 380 | 183 | 0.01 | A0A1Q3QI34 | *Bacteroidia bacterium* | Uncharacterized protein |
| 381 | 183 | 0.01 | | | |
| 382 | 183 | 0.01 | | | |
| 383 | 182 | 0.01 | A0A1Q3QBL5 | *Bacteroidia bacterium* | Uncharacterized protein |

| | | | | | |
|---|---|---|---|---|---|
| 384 | 182 | 0.01 | A0A1Q3QF81 | *Bacteroidia bacterium* | Uncharacterized protein |
| 385 | 182 | 0.01 | A0A1Q3QIV4 | *Bacteroidia bacterium* | Uncharacterized protein |
| 386 | 182 | 0.01 | F0M493 | *Pseudarthrobacter phenanthrenivorans* | Uncharacterized protein |
| 387 | 182 | 0.01 | F4BWG4 | *Methanosaeta concilii* | Uncharacterized protein |
| 388 | 182 | 0.01 | | | |
| 389 | 182 | 0.01 | | | |
| 390 | 181 | 0.01 | A0A1Q3QBI7 | *Bacteroidia bacterium* | Uncharacterized protein |
| 391 | 181 | 0.01 | A0A1Q3QIS1 | *Bacteroidia bacterium* | Uncharacterized protein |
| 392 | 181 | 0.01 | F4BW92 | *Methanosaeta concilii* | Cation-transporting P-type ATPase |
| 393 | 181 | 0.01 | | | |
| 394 | 180 | 0.01 | C5CEK1 | *Kosmotoga olearia* | IstB domain protein ATP-binding protein |
| 395 | 180 | 0.01 | F4BWV3 | *Methanosaeta concilii* | Alanine--tRNA ligase (EC 6.1.1.7) (Alanyl-tRNA synthetase) (AlaRS) |
| 396 | 180 | 0.01 | F4BXD3 | *Methanosaeta concilii* | S-layer-related duplication domain protein |
| 397 | 180 | 0.01 | | | |
| 398 | 179 | 0.01 | A0A1Q3QBV2 | *Bacteroidia bacterium* | Serine protease |
| 399 | 179 | 0.01 | A0A1Q3QF68 | *Bacteroidia bacterium* | Glycoside hydrolase |
| 400 | 179 | 0.01 | | | |
| 401 | 179 | 0.01 | | | |
| 402 | 178 | 0.01 | A0A1Q3QAC3 | *Bacteroidia bacterium* | Leucine--tRNA ligase (Fragment) |
| 403 | 178 | 0.01 | A0A1Q3QF16 | *Bacteroidia bacterium* | Beta-galactosidase (EC 3.2.1.23) (Lactase) |
| 404 | 178 | 0.01 | A0A1Q3QFY4 | *Bacteroidia bacterium* | Uncharacterized protein |
| 405 | 178 | 0.01 | A0A1Q3QIH3 | *Bacteroidia bacterium* | Uncharacterized protein |
| 406 | 178 | 0.01 | H8MH05 | *Corallococcus coralloides* | Long-chain-fatty-acid--CoA ligase |
| 407 | 178 | 0.01 | | | |
| 408 | 178 | 0.01 | | | |
| 409 | 177 | 0.01 | A0A1Q3QAJ9 | *Bacteroidia bacterium* | Uncharacterized protein |
| 410 | 177 | 0.01 | A0A1Q3QGA2 | *Bacteroidia bacterium* | Ferrous iron transport protein B |
| 411 | 177 | 0.01 | A0A1Q3QIL1 | *Bacteroidia bacterium* | Uncharacterized protein |
| 412 | 177 | 0.01 | A0A1Q3QIP6 | *Bacteroidia bacterium* | Transcription-repair-coupling factor (TRCF) (EC 3.6.4.-) |
| 413 | 177 | 0.01 | A7AI47 | *Parabacteroides merdae* | Uncharacterized protein |
| 414 | 177 | 0.01 | F4BYK4 | *Methanosaeta concilii* | Leucine--tRNA ligase (EC 6.1.1.4) (Leucyl-tRNA synthetase) (LeuRS) |
| 415 | 177 | 0.01 | P15689 | *Paramecium tetraurelia* | NADH-ubiquinone oxidoreductase 49 kDa subunit (EC 1.6.5.3) (EC 1.6.99.3) (NADH dehydrogenase subunit 7) |
| 416 | 177 | 0.01 | P11405 | *Moraxella sp.* | Type-2 restriction enzyme MspI (R.MspI) (EC 3.1.21.4) (Endonuclease MspI) (Type II restriction enzyme MspI) |

| | | | | | |
|---|---|---|---|---|---|
| 417 | 177 | 0.01 | | | |
| 418 | 177 | 0.01 | | | |
| 419 | 177 | 0.01 | | | |
| 420 | 177 | 0.01 | | | |
| 421 | 177 | 0.01 | | | |
| 422 | 176 | 0.01 | A0A1G4GAZ3 | *Petrimonas mucosa* | Uncharacterized protein |
| 423 | 176 | 0.01 | A0A1M6CEL6 | *Tessaracoccus bendigoensis* | Uncharacterized protein |
| 424 | 176 | 0.01 | A0A1Q3QAR6 | *Bacteroidia bacterium* | Mannose-6-phosphate isomerase |
| 425 | 176 | 0.01 | | | |
| 426 | 176 | 0.01 | | | |
| 427 | 175 | 0.01 | A0A1Q2CK56 | *Tessaracoccus sp.* | Uncharacterized protein |
| 428 | 175 | 0.01 | A0A1Q3QF30 | *Bacteroidia bacterium* | Hydrolase |
| 429 | 175 | 0.01 | A0A1Q3QGE3 | *Bacteroidia bacterium* | Uncharacterized protein |
| 430 | 175 | 0.01 | | | |
| 431 | 175 | 0.01 | | | |
| 432 | 175 | 0.01 | | | |
| 433 | 174 | 0.01 | F4BVH9 | *Methanosaeta concilii* | Glycosyl transferase, group 2 family protein |
| 434 | 174 | 0.01 | N1JKD7 | *Mesotoga infera* | RNA-directed DNA polymerase (Reverse transcriptase) |
| 435 | 174 | 0.01 | | | |
| 436 | 173 | 0.01 | A0A1Q3QBX1 | *Bacteroidia bacterium* | Polysaccharide biosynthesis protein |
| 437 | 173 | 0.01 | A0A1Q3QGI1 | *Bacteroidia bacterium* | Uncharacterized protein |
| 438 | 173 | 0.01 | A0A1Q3QHE9 | *Bacteroidia bacterium* | Glycosyl transferase |
| 439 | 173 | 0.01 | F4BTH0 | *Methanosaeta concilii* | Glycosyl transferase, group 2 family protein |
| 440 | 173 | 0.01 | F4BTY4 | *Methanosaeta concilii* | SNF2 family helicase (EC 3.6.1.-) |
| 441 | 173 | 0.01 | | | |
| 442 | 173 | 0.01 | | | |
| 443 | 172 | 0.01 | A0A180EWA0 | *Bacteroidales bacterium* | Replication protein |
| 444 | 172 | 0.01 | A0A1Q3QH96 | *Bacteroidia bacterium* | PAS domain-containing sensor histidine kinase |
| 445 | 172 | 0.01 | F0M490 | *Pseudarthrobacter phenanthrenivorans* | DNA primase/polymerase-like protein |
| 446 | 172 | 0.01 | F4BSY5 | *Methanosaeta concilii* | Uncharacterized protein |
| 447 | 172 | 0.01 | | | |
| 448 | 172 | 0.01 | | | |
| 449 | 171 | 0.01 | A0A1Q3Q9R3 | *Bacteroidia bacterium* | DUF5107 domain-containing protein |
| 450 | 171 | 0.01 | A0A1Q3QGP8 | *Bacteroidia bacterium* | Glycoside hydrolase |
| 451 | 171 | 0.01 | F4BUE7 | *Methanosaeta concilii* | Uncharacterized protein |

| | | | | | |
|---|---|---|---|---|---|
| 452 | 171 | 0.01 | | | |
| 453 | 171 | 0.01 | | | |
| 454 | 171 | 0.01 | | | |
| 455 | 170 | 0.01 | A0A1Q3QEW4 | *Bacteroidia bacterium* | Alanine--tRNA ligase (EC 6.1.1.7) (Alanyl-tRNA synthetase) (AlaRS) |
| 456 | 170 | 0.01 | A0A1Q3QGC5 | *Bacteroidia bacterium* | Uncharacterized protein |
| 457 | 170 | 0.01 | A0A1Q3QGL6 | *Bacteroidia bacterium* | Copper-translocating P-type ATPase |
| 458 | 170 | 0.01 | A0A1Q3QID2 | *Bacteroidia bacterium* | Uncharacterized protein |
| 459 | 170 | 0.01 | D8L7R1 | *Paramecium caudatum* | NADH-ubiquinone oxidoreductase chain 4 (EC 1.6.5.3) |
| 460 | 170 | 0.01 | F4BY71 | *Methanosaeta concilii* | WD40-like Beta Propeller Repeat protein |
| 461 | 170 | 0.01 | I4ZNR1 | *Acinetobacter sp.* | Uncharacterized protein (Fragment) |
| 462 | 170 | 0.01 | I8W058 | *Bacteroides cellulosilyticus* | Uncharacterized protein |
| 463 | 169 | 0.01 | A0A098BYW1 | *Fermentimonas caenicola* | Transposase |
| 464 | 169 | 0.01 | A0A1G4G8N7 | *Petrimonas mucosa* | ATP-dependent DNA helicase RecQ (EC 3.6.4.12) |
| 465 | 169 | 0.01 | A0A1Q3QBS3 | *Bacteroidia bacterium* | Uncharacterized protein |
| 466 | 169 | 0.01 | A0A1Q3QFW3 | *Bacteroidia bacterium* | Alpha-mannosidase |
| 467 | 169 | 0.01 | A0A1Q3QIT0 | *Bacteroidia bacterium* | Polyphosphate kinase (EC 2.7.4.1) (ATP-polyphosphate phosphotransferase) (Polyphosphoric acid kinase) |
| 468 | 169 | 0.01 | | | |
| 469 | 169 | 0.01 | V2USD3 | *Acinetobacter brisouii* | Uncharacterized protein |
| 470 | 169 | 0.01 | W9GKA2 | *Intrasporangium chromatireducens* | DNA primase (EC 2.7.7.-) |
| 471 | 168 | 0.01 | A0A023UP33 | *Magnusiomyces ingens* | Uncharacterized protein (Fragment) |
| 472 | 168 | 0.01 | A0A1Q3Q9X7 | *Bacteroidia bacterium* | Uncharacterized protein |
| 473 | 168 | 0.01 | A0A1Q3QFZ4 | *Bacteroidia bacterium* | Uncharacterized protein |
| 474 | 168 | 0.01 | F1Z7R9 | *Novosphingobium nitrogenifigens* | Type-IV secretion system protein TraC |
| 475 | 168 | 0.01 | F4BTS2 | *Methanosaeta concilii* | M6 family metalloprotease domain protein |
| 476 | 168 | 0.01 | F4BVQ0 | *Methanosaeta concilii* | Uncharacterized protein |
| 477 | 168 | 0.01 | | | |
| 478 | 168 | 0.01 | | | |
| 479 | 168 | 0.01 | | | |
| 480 | 167 | 0.01 | A0A0M3AFY3 | *Acinetobacter sp. AG1* | Cytochrome C peroxidase |
| 481 | 167 | 0.01 | A0A1Q3QAM8 | *Bacteroidia bacterium* | Uncharacterized protein |
| 482 | 167 | 0.01 | A0A1Q3QC30 | *Bacteroidia bacterium* | 4-alpha-glucanotransferase |
| 483 | 167 | 0.01 | A0A1Q3QC51 | *Bacteroidia bacterium* | Valine--tRNA ligase (EC 6.1.1.9) (Valyl-tRNA synthetase) (ValRS) |
| 484 | 167 | 0.01 | A0A1Q3QCU6 | *Bacteroidia bacterium* | Uncharacterized protein |
| 485 | 167 | 0.01 | F4BXE8 | *Methanosaeta concilii* | Conserved domain protein |
| 486 | 167 | 0.01 | F4BZQ7 | *Methanosaeta concilii* | S-layer-related duplication domain protein |

| | | | | | |
|---|---|---|---|---|---|
| 487 | 167 | 0.01 | A6LE80 | *Parabacteroides distasonis* | DNA-directed RNA polymerase subunit beta' (RNAP subunit beta') (EC 2.7.7.6) (RNA polymerase subunit beta') (Transcriptase subunit beta') |
| 488 | 167 | 0.01 | | | |
| 489 | 167 | 0.01 | | | |
| 490 | 166 | 0.01 | A0A1Q3QC45 | *Bacteroidia bacterium* | Uncharacterized protein |
| 491 | 166 | 0.01 | A0A1Q3QFI3 | *Bacteroidia bacterium* | DNA topoisomerase IV |
| 492 | 166 | 0.01 | F4BU69 | *Methanosaeta concilii* | Hydrophobe/amphiphile efflux-3 (HAE3) family transporter |
| 493 | 166 | 0.01 | F4BXP6 | *Methanosaeta concilii* | IS66 family element, transposase |
| 494 | 166 | 0.01 | F4C0R1 | *Methanosaeta concilii* | Uncharacterized protein |
| 495 | 166 | 0.01 | H8MK59 | *Corallococcus coralloides* | Linear gramicidin synthase subunit C |
| 496 | 166 | 0.01 | | | |
| 497 | 165 | 0.01 | A0A1Q3QDR2 | *Bacteroidia bacterium* | Uncharacterized protein |
| 498 | 165 | 0.01 | A0A1Q3QEA7 | *Bacteroidia bacterium* | Acriflavine resistance protein B |
| 499 | 165 | 0.01 | A0A1Q3QGG3 | *Bacteroidia bacterium* | Hydrolase |
| 500 | 165 | 0.01 | F4BWB4 | *Methanosaeta concilii* | TPR-repeat protein |
